# Reduced parenteral glucose supply in preterm neonatal infection ameliorates the pulmonary damage

**DOI:** 10.64898/2026.06.05.730350

**Authors:** Nguyen Phuoc Long, Ole Bæk, Karoline Aasmul-Olsen, Richard Doughty, Bjorn Klabunde, Nguyen Quang Thu, Le Hoang Bach Dat, Bui Thanh Liem, Klaus Bønnelykke, Duc Ninh Nguyen

## Abstract

Preterm infants are acutely susceptible to neonatal sepsis, a syndrome characterized by systemic pro-inflammatory activity and life-threatening multi-organ dysfunction. However, the specific pulmonary pathological response to sepsis and the potential for metabolic interventions to mitigate lung injury remain poorly characterized. Herein, we evaluated the impact of varying parenteral glucose regimens on pulmonary outcomes during severe infection using a preterm piglet model. Genome-wide gene expression analysis was used to characterize lung transcriptome profiles. The relationships between gene expression and circulating biochemical and immune profiles were also investigated. Our findings demonstrate that significant pulmonary tissue damage is a hallmark of neonatal sepsis. A reduced-glucose regimen markedly attenuated pulmonary tissue damage while simultaneously alleviating systemic metabolic acidosis and hyperlactatemia. Mechanistically, lung transcriptome profiling revealed a profound activation of pathways associated with inflammatory signaling, programmed cell death, and the dysregulation of glucose, amino acid, and lipid metabolism. The low-glucose intervention effectively mitigated these widespread molecular and metabolic disturbances, suggesting a restorative effect on the pulmonary transcriptome landscape. To facilitate further mechanistic exploration and the identification of novel therapeutic targets, we developed the NeoSepPulmoExplorer (https://pharmaco-omicslab.shinyapps.io/NeoSepPulmoExplorer/), an interactive web-based toolkit for better mechanistic understanding and the identification of potential treatment targets. These results collectively underscore the importance of metabolic modulation in preserving organ function, though further translational studies are requisite to improve clinical outcomes in septic neonates.

## 1. INTRODUCTION

Every year, 15 million infants are born prematurely (<37 weeks of gestation, 10% of all live births). These preterm infants, especially those below 1500 grams or born < 32 weeks of gestation, possess underdeveloped internal organs and immune systems, leading to a high risk of infection, sepsis, necrotizing enterocolitis, respiratory distress syndrome, and intraventricular hemorrhage. In particular, neonatal sepsis remains a substantial challenge, characterized by life-threatening organ dysfunction resulting from a dysregulated host response to infection (Fleiss et al., 2021). If not identified and treated promptly, neonatal sepsis can rapidly progress to severe complications, notably multi-organ impairment, including failure of vital organs. Moreover, neonatal sepsis significantly heightens the risk of long-term neurodevelopmental impairments, including cerebral palsy, cognitive and psychomotor delays, as well as vision and hearing deficits (Taneri et al., 2025).

Among organ injuries caused by sepsis, lung injury is the most common type. On the one hand, it may resemble indirect acute lung injury (ALI) seen in older populations, as it is driven by a complex, multi-factorial immune cell imbalance rather than primarily the bacterial load itself. This imbalance leads to impaired barrier function and subsequent neutrophil infiltration into remote organ systems (Fallon et al., 2021). On the other hand, respiratory complications in preterm neonates, including acute respiratory distress and frequent ventilation, may further contribute to these sepsis-induced morbidities. Further, neonatal sepsis is recognized as a major risk factor for bronchopulmonary dysplasia (BPD), a severe pulmonary disorder in premature infants characterized by disrupted alveolar and pulmonary vascular function. The key aspect of sepsis in this context is its role in triggering impaired lung maturation via persistent inflammation, marked by increased proinflammatory cytokines, e.g., interleukin (IL)-6, IL-1β, interferon (IFN)-γ, and tumor necrosis factor (TNF)-α, causing abnormal lung development by interfering with essential morphogenic processes such as angiogenesis, extracellular matrix formation, and alveologenesis (Ng et al., 2003). Mechanistically, sepsis-induced ALI in neonates is uniquely complicated by developmental arrest in the premature lung, which ultimately makes the infant more vulnerable to BPD (Salimi, Dummula, Tucker, Dela Cruz, & Sampath, 2022). Recognizing sepsis-induced ALI is crucial for research aimed at understanding its severity and identifying therapeutic windows to improve outcomes in preterm infants (Tucker et al., 2023). Targeting the necroptosis pathway through inhibition of receptor-interacting protein kinase 1 (RIPK1) represents a novel and warranted therapeutic approach to mitigating sepsis-mediated ALI and improving survival outcomes in this vulnerable patient population (Bolognese et al., 2018).

Despite this well-known background, the molecular alterations associated with lung injury in preterm infection and sepsis remain unexplored. There has been a lack of investigation into the lung transcriptome and immunometabolic alterations associated with lung tissue damage during infection in preterm neonates. Our group has previously investigated systemic immunometabolism and gene expression alterations of the liver and kidneys using a neonatal sepsis model in *Staphylococcus epidermidis*-infected preterm pigs. Of note, high glucose intake during infection can impair immune responses by triggering excessive glycolysis, which promotes inflammation, impairs cell repair mechanisms, and increases the risk of hyperglycemia. We also found that reducing parenteral glucose supply during neonatal severe infections and sepsis improved the outcomes in preterm piglets (Bæk et al., 2025; Z. Wu et al., 2024; Zhong et al., 2025). In this study, we performed a comprehensive analysis of the effects of systemic infection on lung tissue damage in preterm neonatal piglets using genome-wide gene expression analysis.

## 2. METHODS

### 2.1. Animal study approval

All experiments and studies involving blood and tissue samples from animals, along with all animal procedures, have been approved by the Danish National Committee on Animal Experimentation (License No. 2020-15-0201-00520), in accordance with Directive 2010/63/EU of the European Parliament.

### 2.2. Staphylococcus epidermidis culture preparation

*Staphylococcus epidermidis* stored in a frozen stock was thawed and cultured using tryptic soy broth overnight. The bacterial density was measured to check if it met a predefined criterion. Then, bacteria were collected via centrifugation and suspended in normal saline at a concentration of 3 X 10^8^ CFU/mL (OD_600_ = 1). The full protocol was described by Brunse *et al*. (Brunse, Worsøe, Pors, Skovgaard, & Sangild, 2019)

### 2.3. Animal cohort and experimental procedures

After delivery, Crossbred preterm piglets (Landrace X Large White X Duroc), those with 90% gestation and born by cesarean section, were immediately relocated to our intensive care unit at the University of Copenhagen. A cohort of 38 preterm piglets was used for this investigation. They were kept in individual heated (37 °C) and oxygenated (1-2 L/min) incubators. The tested subjects were resuscitated and fitted with umbilical arterial catheters. This provided the route for bacterial inoculation, parenteral glucose administration, and blood sampling. Within 2 h of birth, the piglets were randomly assigned to two primary groups: Sepsis (SE) and controls. In the SE group, piglets were inoculated with an *S. epidermidis* solution equivalent to 10^9^ CFU/kg via arterial infusion over 3 min. The piglets in the Control group were infused with an equivalent volume of sterile saline using the same protocol. Piglets were nourished using either high (21%, equivalent to 30 g/kg/d, High) or low (5%, equivalent to 7.2 g/kg/d, Low) glucose. Taken collectively, four groups of preterm piglets were established for the subsequent studies: High-glucose infection (n = 12); Low-glucose infection (n = 11); High-glucose control: Control subjects with high-dose glucose treatment (n = 8); and Low-glucose control: Control subjects with reduced-dose glucose treatment (n = 7).

Each animal was monitored closely for up to 22 h post-infection (experimental endpoint) or until sepsis diagnosis requiring humane euthanasia, which was defined by blood pH equal to or less than 7.1, together with clinical symptoms of deep lethargy, discoloration, and tachypnea. At the endpoint, Zoletil (0.1 mL/kg) was used to sedate the piglets, and euthanasia was performed using sodium pentobarbital (60 mg/kg) via intracardial injection. Lung tissue was collected after euthanasia, frozen in liquid nitrogen, and stored at –80 °C until use.

### 2.4. Serum biochemistry and inflammatory biomarkers

Blood samples were collected at 3, 6, 12, and 22 h post-infection to acquire serum and plasma. Serum samples from four time points were used for general hematology with the ADVIA 2120i Hematology System (Siemens Healthcare Diagnostics, NY, USA) and for blood gas analysis with the GEM Premier 3000 (Instrumentation Laboratory, MA, USA). Serum samples collected at the time of euthanasia (either experimental or humane endpoint) were used for biochemical measurements using the ADVIA 1800 Chemistry System (Siemens Healthcare Diagnostics, NY, USA).

### 2.5. Pulmonary lung pathological analysis

Fixed left lungs were paraffin-embedded, sectioned at 4 μm, and stained with hematoxylin and eosin (H&E). Five sections per animal, sampled systematically from the cranial lobe (cranial and caudal divisions) and caudal lobe (cranial, middle, and caudal divisions), were coded and scored by a single veterinary pathologist (RD) blinded to group allocation. Thirteen histopathological features were graded on a four-point ordinal scale (0 = absent to 3 = severe) across five high-power fields (×200) per section, focusing on peribronchiolar and peribronchial alveoli. The features comprised structural changes of impaired alveolar development (alveolar simplification, septal thickening, atelectasis, interstitial fibrosis, and chronic mononuclear inflammation) and acute injury patterns (alveolar hemorrhage, congestion, epithelial necrosis, hyaline membrane formation, neutrophilic infiltration, bronchiolar epithelial injury/denudation, pleural lymphatic dilatation, and microvascular thrombosis).

Quantitative morphometry was performed on H&E-stained sections from the middle caudal lobe and the cranial and caudal divisions of the cranial lobe using ImageJ (v1.54, NIH, USA) by a single blinded observer. Ten non-overlapping fields per section (five for alveolar density), selected by systematic uniform random sampling and excluding large airways, vessels, and atelectatic areas, were analyzed for four indices: mean linear intercept, measured at ×100 using a 2000 μm test-line grid (50 μm spacing) and calculated as total line length divided by alveolar-wall intercepts; radial alveolar count, the number of septa intersected by a perpendicular from the center of a respiratory bronchiole to the nearest septum or pleura (≥5 bronchioles per section); alveolar septal thickness, measured at ×400 by the orthogonal intercept method (20 random points per field, excluding areas of congestion or infiltration); and alveolar density, counted as complete alveolar profiles within a 0.25 mm² frame.

### 2.6. Statistical analysis

The statistical evaluation of both pathological and biochemical data was conducted to assess how parenteral glucose supply modulates the severe infection-induced lung injury and its associated biochemical alterations. We employed linear mixed-effects models (LMMs) from the lme4 package (1.1.38), which are well-suited to temporally correlated data, to analyze longitudinal changes in blood gases and related parameters (Laird & Ware, 1982). The experimental group, time, and their interaction were set as fixed effects to quantify how these levels changed over time between groups. Other fixed-effect variables (i.e., sex and birthweight) were also included to adjust for biological differences. Besides, to account for repeated measurements from the same animal and within-litter correlation, we assign individual piglets and litter as random effects. After model fitting, the significance of group-by-time interaction was assessed using the Wald chi-square test. Then, LMMs were modified to measure parameters at a single time point. The models included only the experimental group, sex, and birth weight as fixed predictors, while litter was retained as the sole random effect accounting for shared genetic backgrounds. Furthermore, the Wald chi-square tests were used to evaluate the overall significance of the group effect. For post hoc pairwise comparisons, estimated marginal means (EMMs) for each group were computed using the emmeans package (2.0.1), and the Kenward-Roger approximation was applied to adjust degrees of freedom introduced by random effects in mixed models, ensuring more accurate P-values given our small sample sizes (Kenward & Roger, 1997). Subsequently, pairwise contrasts between EMMs were performed to identify group differences.

The analysis of pathology scores followed a similar approach to the single-time-point analyses, using the exact same fixed and random effects, because the lung pathology assessment was performed once at the completion of the experiment. However, the cumulative link mixed models in the ordinal package (2025.12.29) were used to account for the ordinal nature of the data (Agresti & Natarajan, 2001; McCullagh, 1980). To assess the significance of the experimental groups, we employed likelihood ratio tests to obtain reliable P-values given the asymmetric distribution of our pathological data.

### 2.7. Pulmonary transcriptomics data acquisition and pre-processing

The experimental pipeline for pulmonary transcriptomics data acquisition and pre-processing was similar to that in our previous publication (Z. Wu et al., 2024). In brief, total RNA from lung tissue was extracted using the RNeasy Mini Kit (QIAGEN) and processed for library preparation with the VAHTS mRNA-seq V3 kit (Vazyme). Samples were sequenced (NovaSeq 6000, 150 bp paired-end reads) and analyzed using an established bioinformatic pipeline, including TrimGalore, Tophat2 (Sscrofa 11.1, Ensembl v99), and htseq-count, conducted by NOVOGENE (Cambridge, UK).

### 2.8. Transcriptomics data processing, statistical analysis, and visualization

We used protein-coding genes for all subsequent data processing and analysis. All subsequent analyses utilized protein-coding genes. Low-count genes (<10 in at least 7 samples) were pre-filtered. Data normalization and scaling were performed prior to the principal component analysis (PCA). DESeq2 was employed for differential expression analysis (DEA) across five comparisons to evaluate the impacts of infection and glucose supply. Treatment- and sepsis-associated pathways were identified. Briefly, the DEA results were sorted by fold change (on a log_2_ scale) regardless of the P-value. The false discovery rate (FDR) of 0.05 was chosen as the statistical threshold for the DEA and gene set enrichment analysis (GSEA). Then, GSEA was conducted using the Kyoto Encyclopedia of Genes and Genomes (KEGG, *Sus crofa*) and Biological Processes in Gene Ontology (GO:BP, *Sus crofa*). A positive normalized enrichment score (NES) indicated an “activated” or “up-regulated” pathway and *vice versa*.

The data normalization and DEA were implemented in ExpressAnalyst (Ewald et al., 2024). The GSEA was conducted using the ClusterProfiler package version 4.16.0 in R version 4.5.2 (R Core Team, 2023; T. Wu et al., 2021). PCA scores plots, volcano plots, and GSEA dot plots were visualized using the EasyPubPlot (Tien, Thu, Kim, Park, & Long, 2025), while the heatmaps of gene expression were made using the NeoSepPulmoExplorer. Each individual panel was assembled by EasyFigAssembler (Long & Thu, 2026).

### 2.9. Integrated web application elucidates the transcriptome mechanisms

We built an integrated web application, NeoSepPulmoExplorer, using R (version 4.5.1) to support comprehensive transcriptomics-based data mining. The graphical user interface was designed using the Shiny framework (version 1.12.1) (R Core Team, 2026a). The shinythemes package (version 1.2.0) defined the visual style of the interface (R Core Team, 2021). Dynamic layouts and interactive events on the frontend were managed using the shinyjs library (version 2.1.1) in combination with custom Cascading Style Sheets (R Core Team, 2026b). The shinyWidgets package (version 0.9.0) supplied advanced input controls, including multi-select dropdowns and an auto-completion feature (R Core Team, 2026c). To enhance the visualization experience, the server dynamically calculates the dimensions of the heatmap based on the size of the input data, namely, the number of queried genes and samples. The resulting heatmap is then displayed within a dedicated container with horizontal and vertical scrollbars, ensuring it is presented at its actual size across various screen sizes. Altogether, these components create a responsive, user-friendly interface that supports seamless exploration of the transcriptomics results.

The analytical pipeline extracts pathway-enriched genes using predefined KEGG (default) or GO:BP terms, mapped from Entrez IDs to gene symbols via a fixed *Ensembl* conversion file. Users may optionally filter features to retain only those with FDR < 0.05. Metadata-based mapping facilitates targeted cross-condition comparisons, with input data standardized via row-wise Z-scores. Final expression matrices are visualized using the ComplexHeatmap package (version 2.24.1), incorporating feature-wise hierarchical clustering (Gu, 2022). Annotation columns displaying FDR values from additional contrasts are integrated to provide a broader experimental context. Beyond the provided datasets (directly linked to the data generated by this study), the platform supports custom uploads of metadata, gene expression matrices, and pathway enrichment files for expanded data mining. It is worth noting that the application has no preprocessing function. Users must ensure their data is fully processed and normalized in advance using the same or a comparable pipeline.

Our web application also supports exporting graphical deliverables in TIFF, PDF, PNG, or JPEG formats. It offers two resolution options: 300 dots per inch (DPI) for standard use and 600 DPI for publication-quality figures. As a result, interested users can produce versatile deliverables tailored to their objectives, ranging from personal use to data presentation and formal publications.

## 3. RESULTS

### 3.1. Reduced-glucose supply in post-infection is associated with reduced severity of sepsis

The study design and workflow are illustrated in **Figure 1A**. We first examined lung pathology across the four experimental groups at the final time point (22h). The analysis underscored a profound increase in lung pathology in response to infection, irrespective of the glucose intervention. As shown in **Figure 2A**, the overall lung damage scores among the four experimental groups differed significantly. Both the Low-glucose and High-glucose infection groups exhibited significantly higher lung pathology scores than their respective control groups. Direct comparison of the control groups revealed no statistically significant difference in terminal lung pathology regardless of the glucose interventions. In contrast, the High-glucose infection group showed significantly greater lung pathology than the Low-glucose infection group. The main histopathological manifestations included lung congestion, inflammation, and edema (**Figures 2B-D**). Representative micrographs showing distinct morphological changes of the lungs across the infection and control groups (**Figures 2E-H)**. Significant pathological alterations were observed following infection. In the High-glucose infection group, apparent cellular infiltration around the bronchioles and in the interstitium/alveolar walls, thickened alveolar septa, and dilated and congested small blood vessels in the alveolar septa were observed (**Figure 2G**). The infection group treated with reduced glucose showed focal, sparse morphological changes and less distortion of the overall lung tissue architecture (**Figure 2H**). Taken collectively, our histopathological assessment suggests that the low-glucose treatment regimen significantly attenuates the severity of lung tissue damage in the context of severe infection.

**Figure 1.**
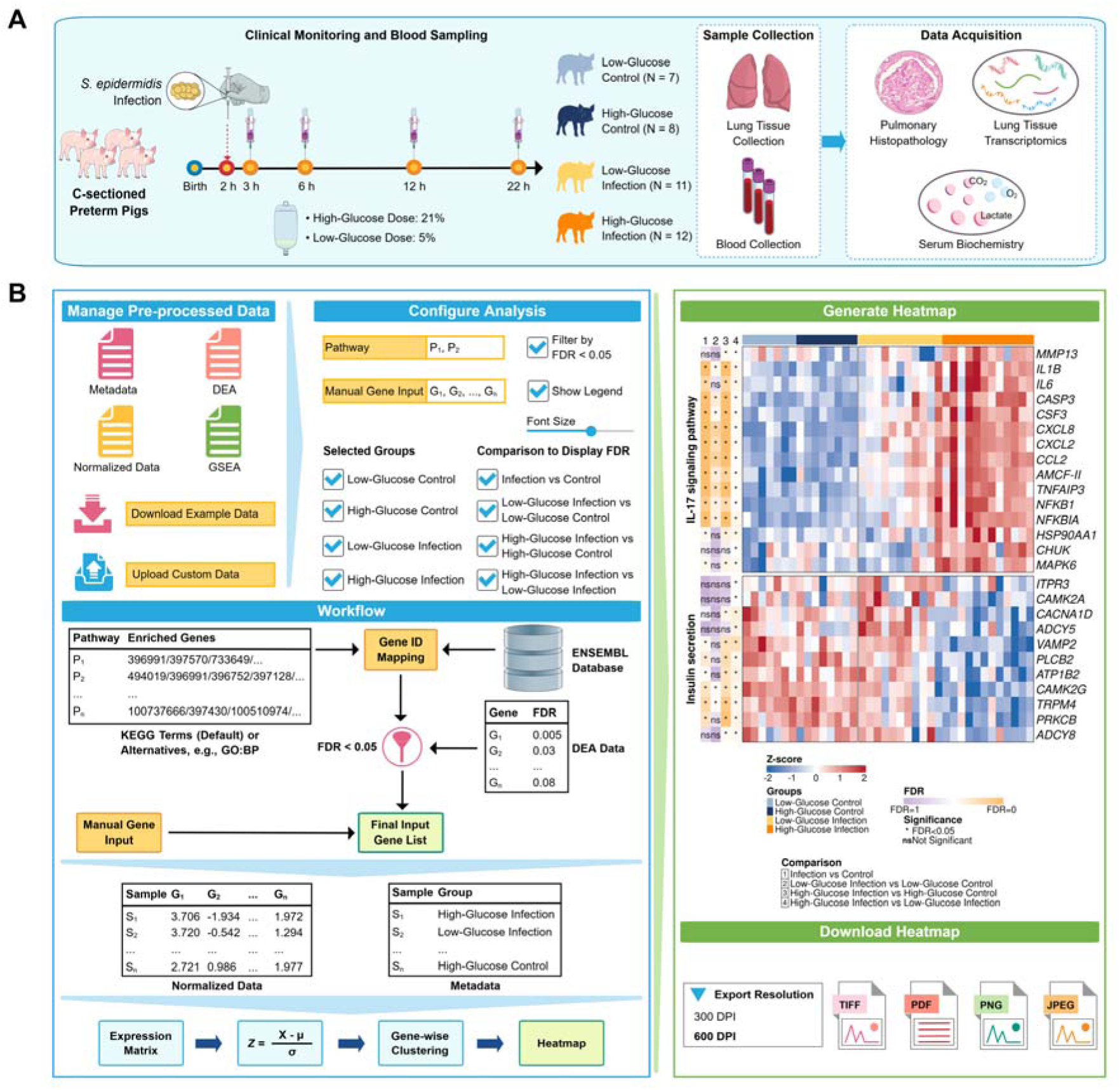
Study overview and the development of NeoSepPulmoExplorer. (A) Study overview and acquired data. (B) The architecture of NeoSepPulmoExplorer an interactive web-based toolkit for better mechanistic understanding of lung damage caused by infections. Abbreviations: DEA: Differential expression analysis; FDR: False discovery rate; GO:BP: Biological Processes in Gene Ontology; GSEA: Gene set enrichment analysis; IL-17: Interleukin-17; KEGG: Kyoto Encyclopedia of Genes and Genomes.

**Figure 2.**
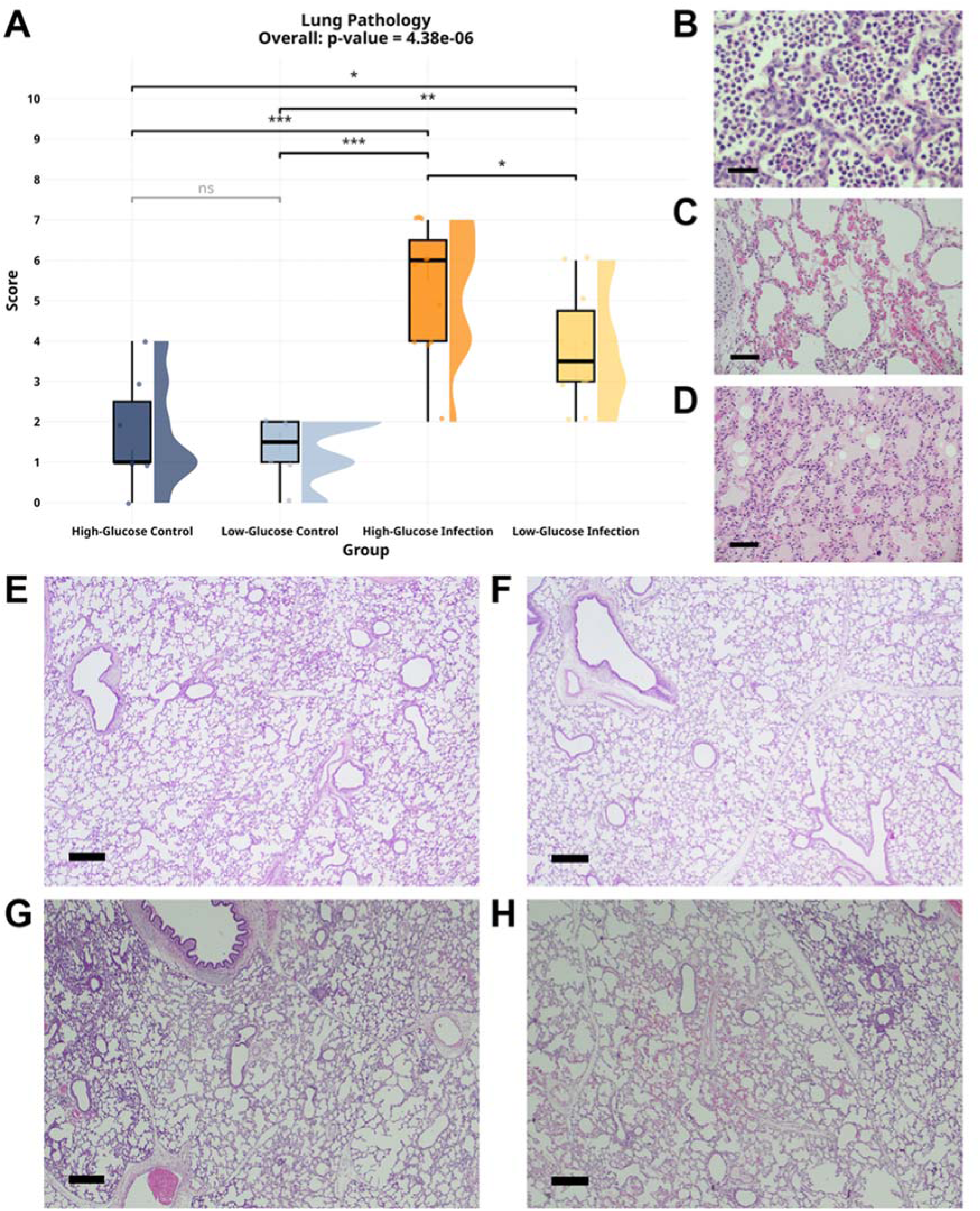
Histopathological assessment of the lung. (A) The pathological score of the lung upon infection and different glucose supply. (B-D) Representative H&E-stained micrographs show histopathological manifestations of the lung: inflammation (×600, scale bar: 50 μm), congestion (×200, scale bar: 150 μm), and edema (×200, scale bar: 150 μm). Representative H&E-stained micrographs of the lung tissue from the 4 groups: (E) High-glucose control (×40, scale bar: 500 μm), (F) Low-glucose control (×40, scale bar: 500 μm), (G) High-glucose infection (×40, scale bar: 500 μm), (H) Low-glucose infection (×40, scale bar: 500 μm). Abbreviation: H&E: Hematoxylin and eosin.

Furthermore, as previously documented (Bæk et al., 2025), infection led to both respiratory and lactic acidosis and an imbalance in the circulating acid-base status (**Supplementary Figure S1**). The low-glucose treatment did appear to partially mitigate the severity of acidosis, hyperlactatemia, and the decline in SBC or cBase (Ecf) compared with the High-Glucose Infection group at 12 h and 22 h. Based on clear evidence of how infection and glucose supply influence the respiratory acidotic state and histological alterations, we further conducted a lung transcriptome comparison among groups.

### 3.2. Infection induces profound lung transcriptome alterations

Comparison of the Infection and Control groups revealed substantial alterations in the lung transcriptome. PCA scores demonstrated distinct separation between groups **(Figure 3A)**, with a total explained variance of 41.7% (PC1: 28.5%). While the High-glucose infection and control groups exhibited distinct profiles, the Low-glucose infection group clustered centrally, suggesting an attenuated transcriptome shift **(Figure 3B)**. The analysis identified 6,398 differentially expressed genes (DEGs): 3,183 upregulated, 3,215 downregulated (**Supplementary Figure S2A**). Approximately 10% exhibited a fold-change of 2 or higher. Collectively, these data demonstrate a high magnitude of lung transcriptome flux by 22 h post-infection.

**Figure 3.**
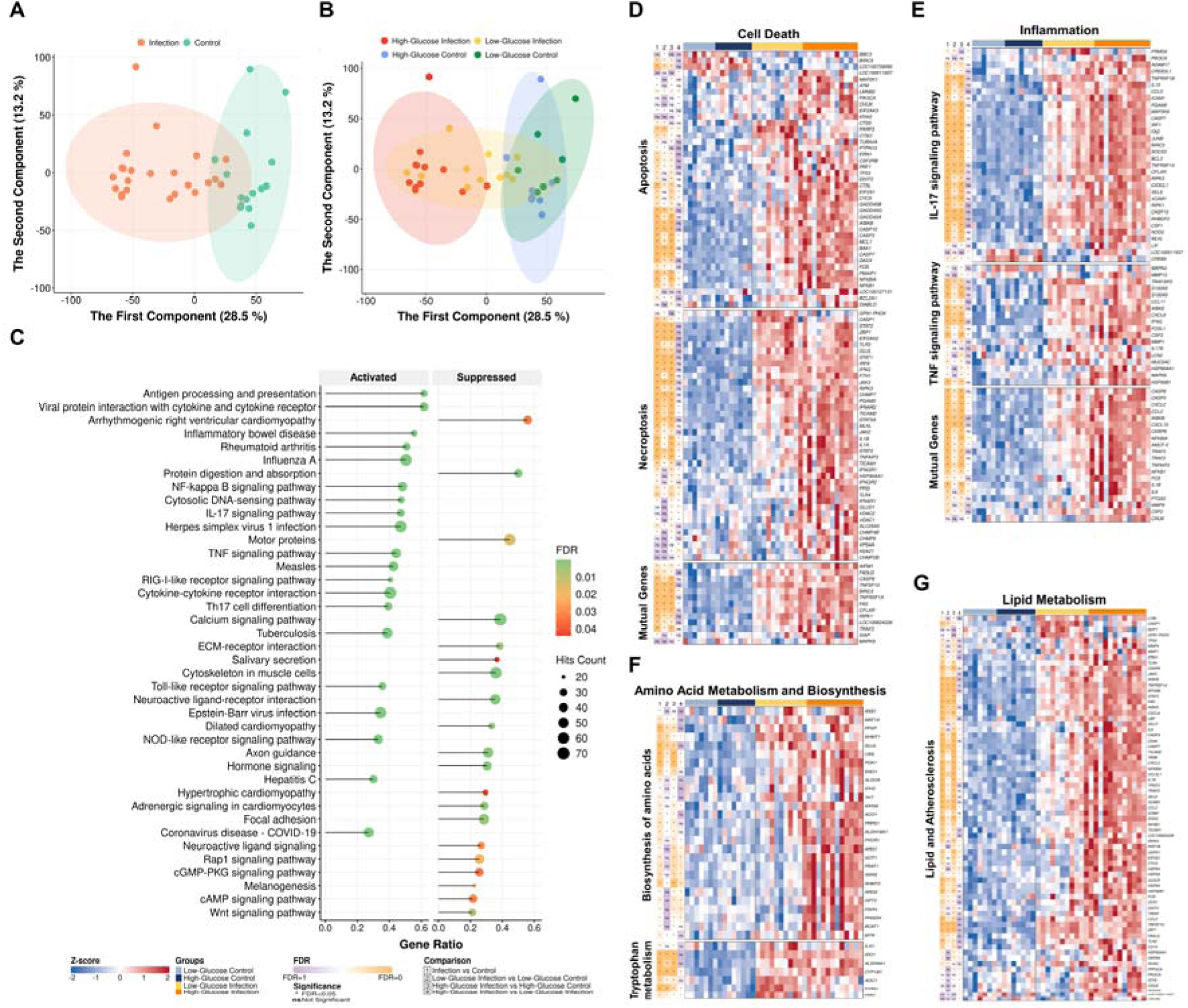
*Staphylococcus epidermidis* infection affected the pulmonary transcriptome of neonatal piglets at euthanasia. (A) PCA scores plot of transcriptome profiles between the Infection and Control groups. (B) PCA scores plot of transcriptome profiles across four subgroups. (C) GSEA dot plot of top activated and suppressed pathways related to the infection, KEGG database. (D-G) Heatmap of altered pathways related to (D) cell death; (E) inflammation; (F) amino acid metabolism and biosynthesis; and (G) lipid metabolism. Abbreviations: cAMP: Cyclic adenosine monophosphate; cGMP-PKG: Cyclic guanosine monophosphate-protein kinase G; ECM: Extracellular matrix; FDR: False discovery rate; GSEA: Gene set enrichment analysis; IL-17: Interleukin-17; KEGG: Kyoto Encyclopedia of Genes and Genomes; NF-kappa B: Nuclear factor-kappa B; NOD: Nucleotide-binding domain; PCA: Principal component analysis; Th17: T helper 17; TNF: Tumor necrosis factor.

We conducted GSEA to identify biological processes associated with the observed high level of lung transcriptome alterations. The GSEA results using the KEGG knowledgebase (GSEA-KEGG) showed 80 “activated” and 21 “suppressed” pathways **(Figure 3C)**. Neonatal sepsis triggered profound pulmonary transcriptome reprogramming characterized by the significant activation of pro-inflammatory processes (e.g., TNF, IL-17, and Toll-like receptor (TLR) signaling) and pathways related to programmed cell death. This immunological cascade, coupled with the upregulation of viral-defense and cytokine-mediated signaling, underscores a state of acute hyperinflammation. Conversely, the suppression of pathways that govern structural integrity and basal metabolism, such as extracellular matrix (ECM)-receptor interactions and calcium signaling, highlights a profound disruption of pulmonary homeostasis **(Figure 3D-G)**. GSEA-GO:BP revealed the upregulation of immune-related pathways in the sepsis piglets **(Supplementary Figures S2B-E)**. Complete lists of significant pathways in KEGG and GO:BP terms are provided in **Supplementary Tables S1 and S2**. Collectively, these molecular signatures delineate a landscape in which coordinated inflammatory activation and metabolic dysregulation drive ALI progression.

### 3.3. Conventional and reduced glucose supply affect the lung transcriptome differently in preterm neonatal infections

Here, we further conducted two subgroup analyses: High-glucose infection vs. High-glucose control and Low-glucose infection vs. Low-glucose control. In the first analysis comparing High-glucose infection and High-glucose control, High-glucose intervention during infection induced extensive shifts in the lung transcriptome relative to High-glucose control (**Figure 4A**). The High-glucose infection group exhibited 5,371 DEGs, with 2,880 up-regulated and 2,851 down-regulated features (**Supplementary Figure S3A**). Pathway enrichment analysis identified activation of inflammatory cascades, specifically the TNF and IL-17 signaling pathways, alongside dysregulation of amino acid and lipid metabolism (**Figures 4B-F**; **Supplementary Table S3**). The functional analysis using the GO:BP knowledgebase further confirmed the upregulation of immune-related signaling, including responses to cytokines and peptides (**Supplementary Figures S3B-E**, **Supplementary Table S4**).

**Figure 4.**
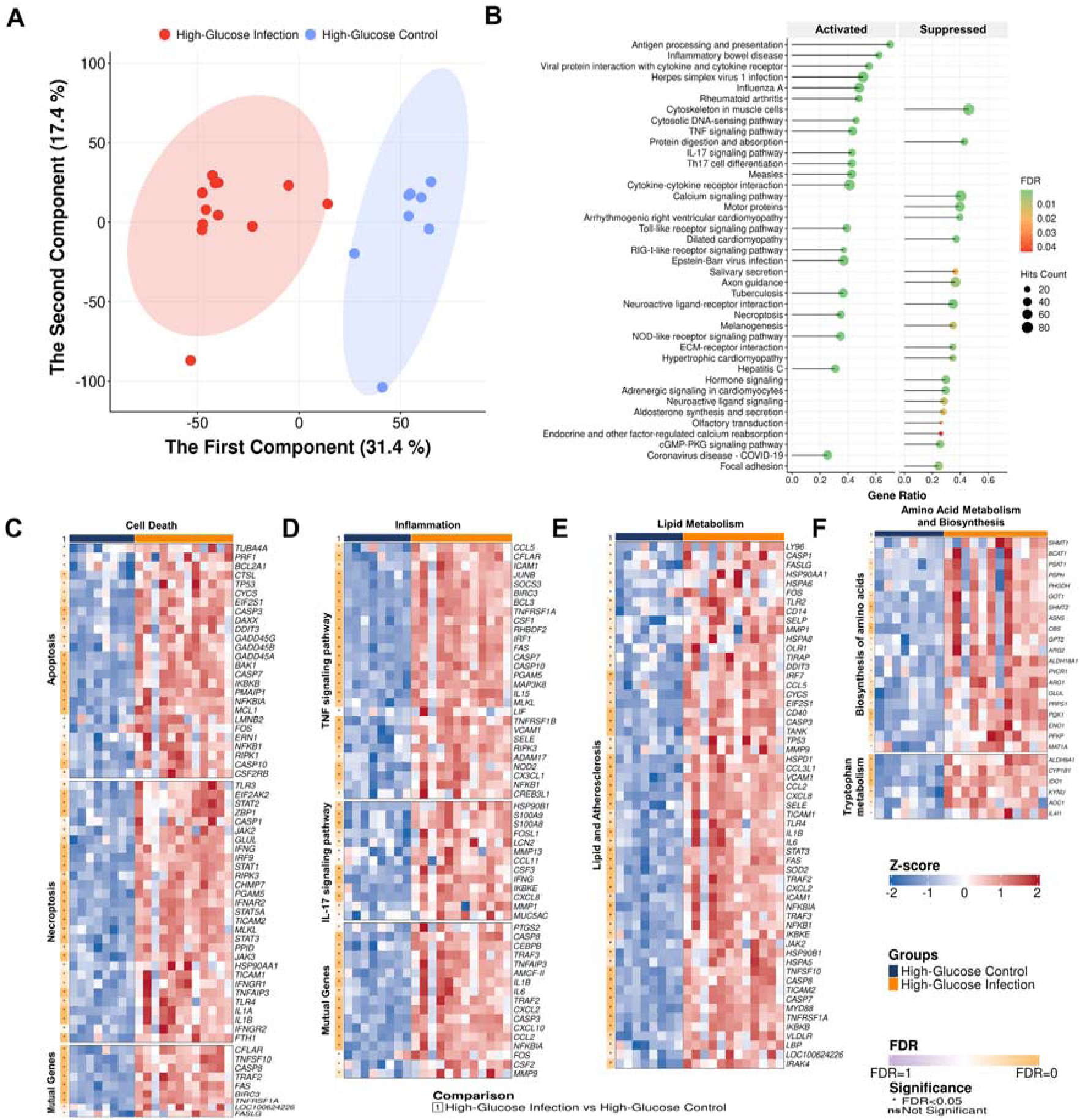
High-glucose supply results in profound alterations in the lung transcriptome profiles between infection and control animals. (A) PCA scores plot of transcriptome profiles between High-glucose infection and High-glucose control. (B) GSEA dot plot of top activated and suppressed pathways related to the infection, KEGG database. (C-F) Heatmap of altered pathways related to (C) cell death; (D) inflammation; (E) amino acid metabolism and biosynthesis; and (F) lipid metabolism. Abbreviations: cGMP-PKG: Cyclic guanosine monophosphate-protein kinase G; ECM: Extracellular matrix; FDR: False discovery rate; GSEA: Gene set enrichment analysis; IL-17: Interleukin-17; KEGG: Kyoto Encyclopedia of Genes and Genomes; NF-kappa B: Nuclear factor-kappa B; NOD: Nucleotide-binding domain; PCA: Principal component analysis; RIG-I: Retinoic acid-inducible gene I; Th17: T helper 17 cells; TNF: Tumor necrosis factor.

A reduced parenteral glucose supply attenuated the magnitude of these transcriptome alterations. Comparison of the Low-glucose infection and the corresponding control group revealed an observable separation (**Figure 5A**), with a substantially lower total of 2,955 identified DEGs (1,654 up-regulated and 1,391 down-regulated; **Supplementary Figure S4A**). While differences in core inflammatory and metabolic signatures remained detectable (**Figures 5B-F**, **Supplementary Figure S4B-E**), the extent of transcriptome change was reduced compared with that observed in the high-glucose intervention. Activated signaling pathways, including TLRs, were observed, whereas structural and basal pathways, such as protein digestion and ribosome function, were found suppressed (**Supplementary Tables S5-S6**). Altogether, the results suggested that two glucose-supplementation doses in infected preterm neonates had different effects, with the low-dose supplement resulting in fewer alterations in transcriptome profiles than the high-dose supplement.

**Figure 5.**
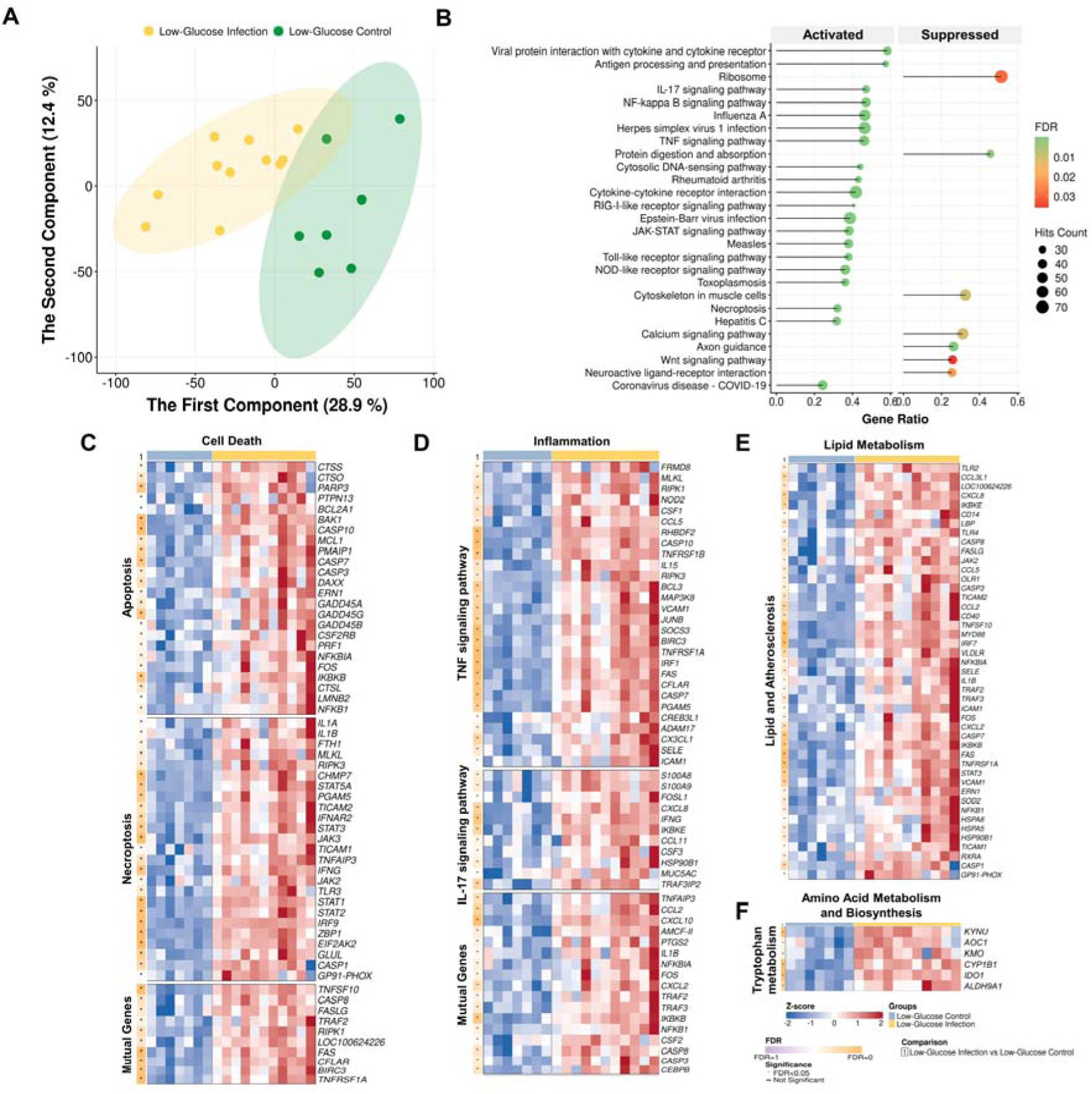
Reducing the glucose supply led to mild alterations in the lung transcriptome profiles between infection and control animals. (A) PCA scores plot of transcriptome profiles between the Low-glucose infection and the Low-glucose control. (B) GSEA dot plot of top activated and suppressed pathways related to the infection, KEGG database. (C-F) Heatmap of altered pathways related to (C) cell death; (D) inflammation; (E) amino acid metabolism and biosynthesis; and (F) lipid metabolism. Abbreviations: cGMP-PKG: Cyclic guanosine monophosphate-protein kinase G; ECM: Extracellular matrix; FDR: False discovery rate; GSEA: Gene set enrichment analysis; IL-17: Interleukin-17; JAK-STAT: Janus kinases-signal transducer and activator of transcription proteins; KEGG: Kyoto Encyclopedia of Genes and Genomes; NF-kappa B: Nuclear factor-kappa B; NOD: Nucleotide-binding domain; PCA: Principal component analysis; RIG-I: Retinoic acid-inducible gene I; TNF: Tumor necrosis factor.

### 3.4. Reduced glucose supply strategy attenuates the pulmonary inflammation of preterm neonatal infections

We also investigated the effects of reducing glucose supply on pulmonary inflammation in infected preterm piglets. The reduced parenteral glucose intervention significantly modulated the lung transcriptome response to infection. The PCA scores plot showed a slight difference between the two subgroups of infected piglets **(Figure 6A)**. The DEA revealed 2,689 differentially expressed genes between the two treatment doses, including 1,391 up-regulated and 1,298 down-regulated genes **(Supplementary Figure S5A)**. High-glucose treatment was characterized by robust activation of complex cascades associated with pro-inflammatory responses, specifically the IL-17 and TNF signaling pathways, along with significant upregulation of mediators implicated in apoptosis, necroptosis, and dysregulated amino acid and lipid metabolism. Furthermore, high-dose intervention led to greater activation of pathways governing ribosome biogenesis, protein processing in the endoplasmic reticulum, and oxidative phosphorylation, while suppressing homeostatic functions, including insulin secretion and ABC transporter activity (**Figures 6B-I**, **Supplementary Table S7**). Concordantly, GSEA-GO:BP enrichment confirmed the activation of cell cycle-related processes in the high-glucose piglets with infection (**Supplementary Figure S5B-F**; **Supplementary Table S8**). These results suggested that reduced-glucose treatment was associated with a lesser degree of pulmonary inflammation. The finding is consistent with the lower degree of lung damage in infected piglets receiving reduced-glucose treatment, as observed in the histological examination.

**Figure 6.**
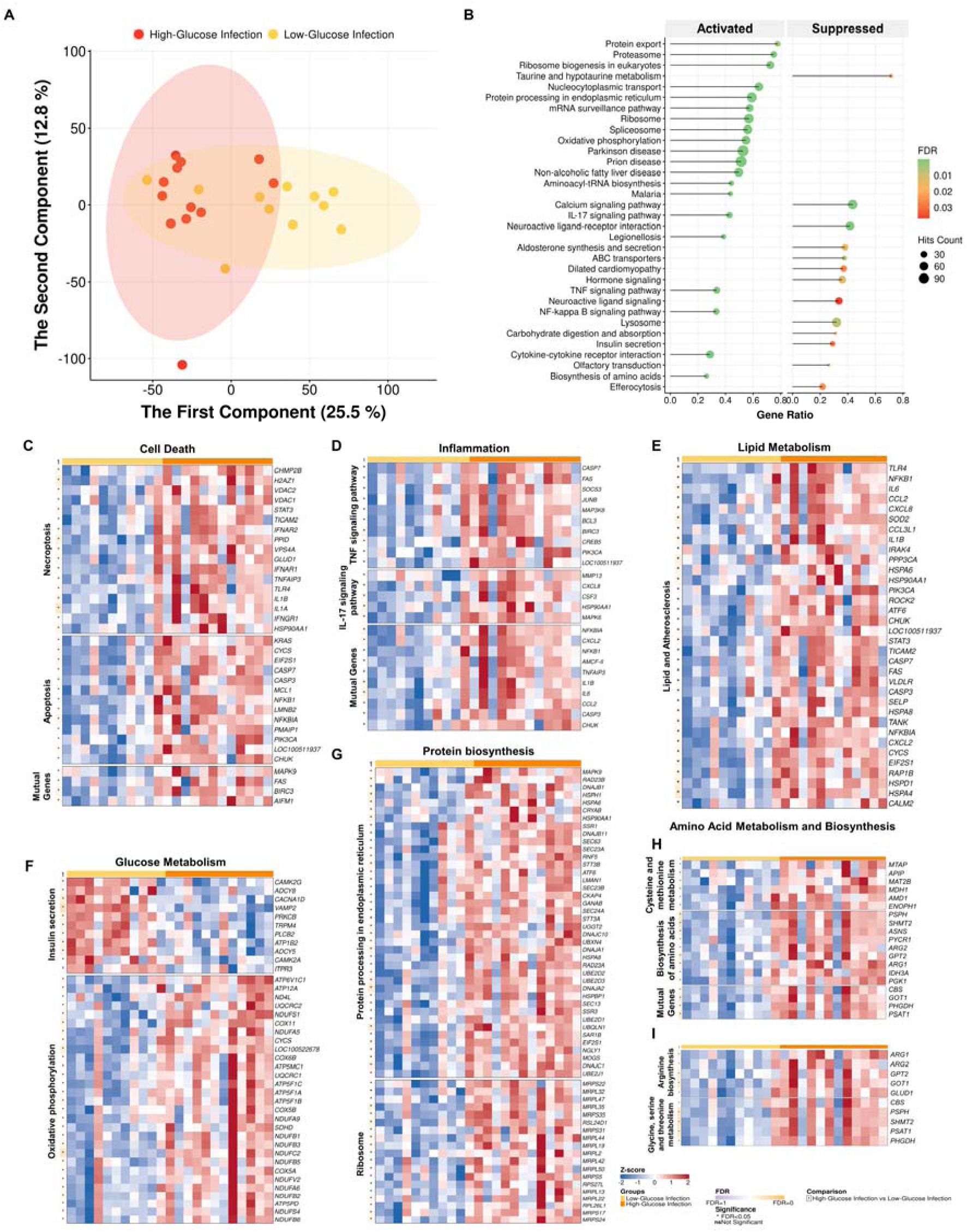
Low-glucose supply resulted in the reduction of inflammation and pulmonary OXPHOS in the infected piglets. (A) PCA scores plot of transcriptome profiles between High-glucose infection and Low-glucose infection. (B) GSEA dot plot of top activated and suppressed pathways associated with the treatment doses, KEGG database. (C-I) Heatmap of altered pathways related to (C) cell death; (D) inflammation; (E) lipid metabolism; (F) glucose metabolism; (G) Protein biosynthesis; (H-I) Amino acid metabolism and biosynthesis. Abbreviations: ABC: ATP-binding cassette; FDR: False discovery rate; GSEA: Gene set enrichment analysis; IL-17: Interleukin-17; KEGG: Kyoto Encyclopedia of Genes and Genomes; NF-kappa B: Nuclear factor-kappa B; PCA: Principal component analysis; TNF: Tumor necrosis factor.

Lastly, we examined whether different glucose-supply treatments altered the lung transcriptome in non-infected piglets. The PCA scores plot shows the overlap between the two clusters, High-glucose control and Low-glucose control, while no gene had an FDR below 0.05 from the DEA. The two analyses collectively indicated that different glucose doses did not alter the lung transcriptome profiles in the control animals **(Supplementary Figure S6)**.

### 3.5. Integrated web application elucidates the transcriptome mechanisms of infection-induced lung damage and provides novel therapeutic targets

Manual interpretation of transcriptomics data on lung injury mechanisms is often labor-intensive and prone to bias. We developed a user-friendly web-based platform that functions as a robust data-mining framework. It enables researchers to better understand the mechanisms of sepsis-induced lung damage and more efficiently bridge the gap between transcriptomics findings and the discovery of therapeutic interventions.

Our platform facilitates dynamic data exploration via an intuitive graphical interface that supports both in-house and custom datasets (**Figure 1B**). Gene selection is performed via pathway-based database queries or manual entry, with optional statistical filtering to refine biologically relevant signals. Visualizations are highly customizable, enabling selective inclusion of groups and the direct integration of statistical annotations for comparative analysis. Example heatmaps for IL-17 signaling and insulin secretion illustrate the mapping of gene-level significance across experimental conditions. Notably, the tool enables hybrid analysis by integrating pathway-driven and manually curated gene lists (**Figure 7A**). This utility was showcased by combining 20 literature-derived lipid-related genes with the 56-gene “Lipid and atherosclerosis” pathway, yielding a comprehensive comparison of transcriptome profiles (**Figures 7B-C**).

**Figure 7.**
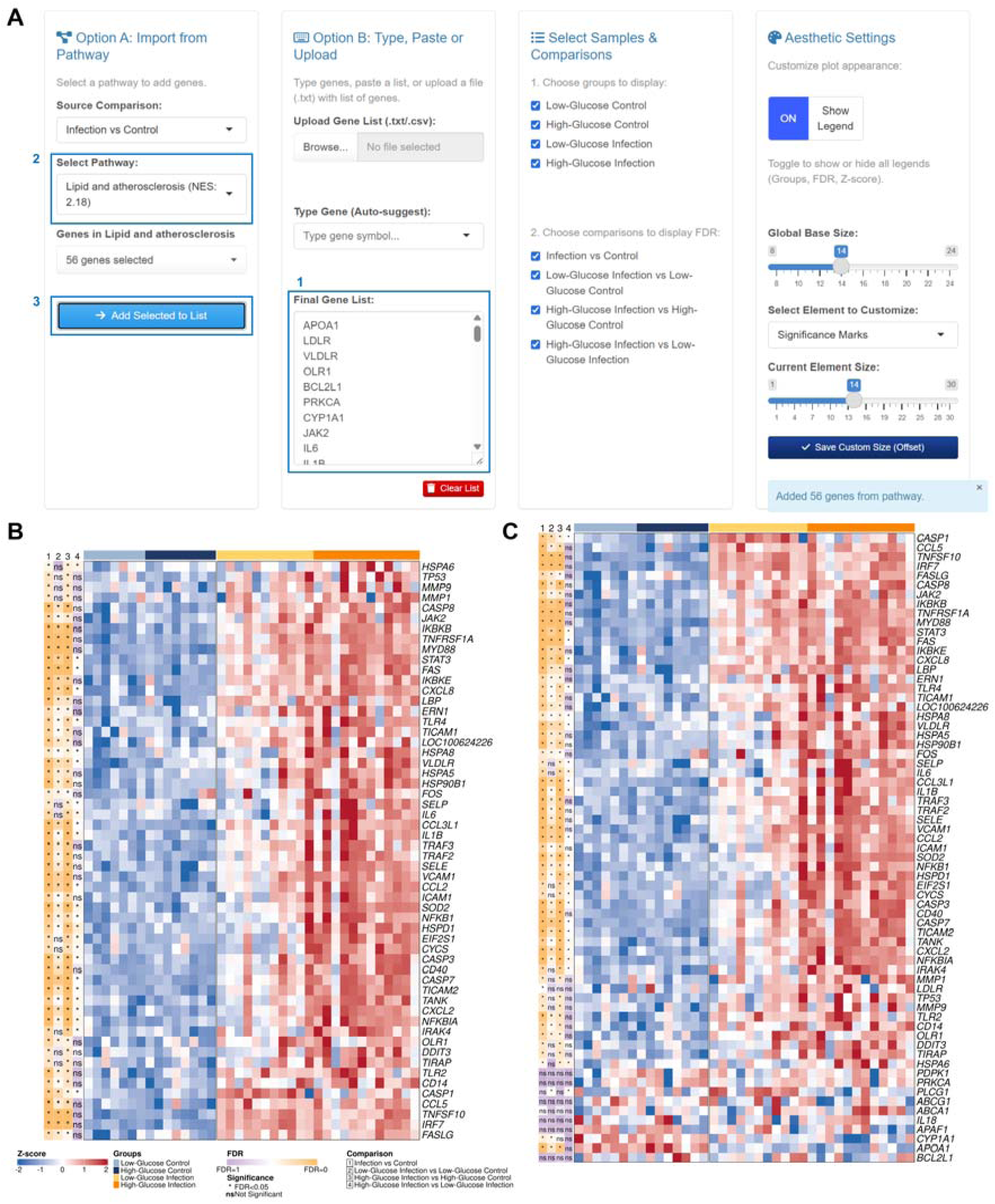
NeoSepPulmoExplorer showcase for visualization of selected genes from a pathway and manually entering a gene list. (A) The list of lipid-related genes was defined for heatmap visualization. 1) The list of 20 lipid-related genes was searched from the literature and manually inputted to the app; 2) The pathway “Lipid and atherosclerosis” was selected in the “Select Pathway”; and 3) Clicked “Add Selected to List” to combine both the pathway-driven gene list and the manually input gene list. (B-C) The heatmaps before (B) and after (C) the combination of the two gene lists. Abbreviations: FDR: False discovery rate; NES: Normalized enrichment score.

## 4. DISCUSSION

Using the preterm piglet model, our group has previously characterized the molecular mechanisms underlying damage to vital organs in severe preterm infections and sepsis, namely the liver, brain, and kidney. Briefly, the liver undergoes a profound metabolic shift during infection, from oxidative phosphorylation to aerobic glycolysis, thereby fueling intense pro-inflammatory cascades and collateral tissue damage. This immune-metabolic dysregulation impairs vital hepatic functions (Z. Wu et al., 2024). The immature blood-brain barrier in preterm neonates leaves the brain highly susceptible to systemic infections, triggering severe neuroinflammation driven by hyperactivation and M1-like polarization of microglial cells. This robust local immune response causes structural damage, including perivascular edema and cortical hyperemia, and actively impairs neuronal plasticity and overall neurodevelopment (Zhong et al., 2025). Structurally immature kidneys with highly fenestrated vasculature are directly exposed to systemic blood-borne pathogens, precipitating acute kidney injury fueled by upregulated local immune-cell glycolysis and severe intrarenal inflammation. This profound inflammatory state drives marked functional decline and significant tissue destruction, manifesting histopathologically as tubular dilatation, proximal tubule vacuolization, and extensive interstitial edema (Zhong et al., 2025). We also found that reducing parenteral glucose supply during severe preterm neonatal infections, while it might increase the risk for hypoglycemia and inadequate energy provision, significantly reduces sepsis severity and improves overall survival. This intervention works by shifting the host’s metabolic state. As a result, it restricts the excessive aerobic glycolysis that fuels hyper-inflammation and instead enhances hepatic oxidative phosphorylation (OXPHOS), the tricarboxylic acid (TCA) cycle, and gluconeogenesis. However, the extent of lung damage, the underlying mechanisms, and whether reduced parenteral glucose supply could reduce the disease severity remained unknown.

Pulmonary inflammation is implicated in the pathway leading to lung injury in preterm infants (Chakraborty, McGreal, & Kotecha, 2010). Proinflammatory cytokines such as TNF-α, IL-1, IL-6, IL-11, vascular endothelial growth factor (VEGF), transforming growth factor (TGF)-α, and TGF-β are inflammatory mediators that may cause severe damage to the capillary endothelium and alveolar epithelium, leading to hyaline membrane formation and edema fluid leakage. Our study contributed to the current body of literature on severe infections and sepsis in preterm infants by revealing profound activation of pathways associated with inflammatory signaling, programmed cell death, and dysregulation of glucose, amino acid, and lipid metabolism. We found that severe infection in preterm neonates causes profound structural lung damage driven by massive immunometabolic shifts at the tissue transcriptome level. In particular, the lung alters the expression of over 6,000 genes, fundamentally reprogramming its cellular environment. This is in line with our recent observations on the liver, which indicate that neonatal sepsis does not just trigger an inflammatory response. It induces a massive, systems-level reprogramming of the cellular environment (Bæk et al., 2025; Z. Wu et al., 2024). The primary driver of lung tissue injury may be due to a complex hyperinflammatory immune response. Infection significantly upregulates immune pathways, specifically TLR, TNF, and IL-17 signaling pathways, overwhelming the lungs with inflammatory cytokines and peptides. This intense immune response directly triggers programmed cell death pathways, including apoptosis and necroptosis, leading to physical lung damage. Markers of necroptosis, such as RIPK1 and IL-18, were previously reported to be significantly elevated during sepsis, whereas Caspase-8 levels were reduced, indicating a shift from silent apoptosis toward inflammatory necroptotic death (Briassoulis et al., 2025). A study investigating the effects of acute ozone exposure on the lungs of newborn mice also found a global suppression of functional genes involved in the cell cycle, cellular assembly, and tissue organization (Gabehart et al., 2014). Meanwhile, normal structural maintenance processes, such as ECM-receptor interactions, are severely suppressed. Our data also indicate that the infection causes severe systemic metabolic acidosis and hyperlactatemia, reflecting a massive disruption in how the body processes energy. In the lungs, we found significant alterations in lipid and amino acid metabolism. These immunometabolic mechanisms appeared to be highly sensitive to the amount of glucose supplied to the neonate. A high-glucose environment may worsen lung pathology by inducing hyperglycemia, elevated glycolysis, and inflammation, leading to metabolic acidosis and sepsis (Muk, Brunse, Henriksen, Aasmul-Olsen, & Nguyen, 2022). Conversely, safely restricting the glucose supply attenuates the severity of lung tissue damage, partially mitigates systemic lactic acidosis, and reduces the overall magnitude of inflammatory gene activation.

Neonatal sepsis is strongly associated with increased odds of chronic lung disease (Lahra, Beeby, & Jeffery, 2009). Sepsis in premature neonates induces alveolar simplification that manifests as BPD, the most common chronic lung disease in infants. BPD is driven by an inflammatory response in the immature lung that distorts alveolar, mesenchymal, and vascular structures (Holzfurtner et al., 2022), with inflammation serving as a common denominator (Balany & Bhandari, 2015). This process is mainly driven by redox-dependent inflammation and tissue injury in the developing pulmonary system. Mechanistically, lipopolysaccharide (LPS), a common bacterial component in sepsis, activates TLRs, thereby triggering complex signaling cascades that lead to an acute rise in reactive oxygen species production, primarily mediated by nicotinamide adenine dinucleotide phosphate oxidase 2 (NOX2). This activation stimulates typical TLR signaling, involving NOX2-regulated phosphorylation of key kinases, including inhibitor of NF-κB kinase-β (IKK-β) and mitogen-activated protein kinases (MAPKs) such as p38 and c-Jun N-terminal kinase (JNK), thereby promoting the nuclear translocation of proinflammatory transcription factors NF-κB and activator protein-1. Deficiency or inhibition of NOX2 reduces activation of this proinflammatory TLR pathway and lessens sepsis-induced alveolar remodeling by preventing the LPS-mediated increase in matrix metalloproteinase 9 and the decrease in crucial lung growth markers, such as elastin and fibroblast growth factor 7. However, the detailed mechanisms related to preterm lung damage in severe infection and sepsis, and their progression to BPD, warrant further investigation.

A major advantage of our study is the development of a user-friendly, web-based platform that streamlines the interpretation of complex lung transcriptome data, effectively bridging the gap between high-dimensional molecular findings and the discovery of novel therapeutic interventions. By automating the analysis of sepsis-induced lung damage, the application mitigates the labor-intensive nature and potential researcher bias associated with manual data mining. The tool’s intuitive interface enables dynamic exploration of biological pathways, such as IL-17 signaling, amino acid, and apoptosis, while also allowing researchers to integrate custom datasets and literature-derived gene lists for targeted mechanistic discovery. Ultimately, this integrated framework accelerates the identification of key molecular drivers behind acute pulmonary deterioration, providing a robust, reproducible resource for the neonatal research community.

Despite offering critical insights into pulmonary molecular changes, this study has several limitations. Firstly, assessing the lung transcriptome exclusively at the experimental endpoint provides a single-point snapshot that precludes establishing temporal associations between infection onset and lung injury. Consequently, these alterations may represent late-stage compensatory mechanisms rather than early molecular triggers. Reduced glucose supply was linked to both systemic metabolic improvement and attenuated lung injury in our study. However, whether this pulmonary protection stems from a direct lung-specific mechanism or is secondary to reduced systemic sepsis severity remains unknown. Future investigations should prioritize longitudinal multi-omic sampling to delineate the kinetic trajectory of host responses. Secondly, our analysis cannot completely rule out the influence of confounding factors inherent to the physiology of prematurity. In the preterm piglet model, which shares high anatomical and physiological fidelity with human neonates, conditions such as primary surfactant deficiency and acute cardiopulmonary instability are common. These prematurity-related issues can induce baseline pulmonary inflammation or mechanical stress that may synergize with, or mask, the specific molecular signatures of pathogen-induced lung injury. Thirdly, while our model using *S. epidermidis* accurately mimics common nosocomial coagulase-negative staphylococcal infections, the host’s lung defense strategies and transcriptome responses may differ significantly when challenged with different bacterial species or viral pathogens. Finally, while our study design allowed us to capture acute pulmonary deterioration at the molecular level, focusing on the immediate interplay between systemic metabolic disturbances and lung inflammatory events, it precludes assessment of the long-term pulmonary morbidity in surviving preterm neonates. Neonatal sepsis is a known risk factor for the development of chronic lung disease, such as bronchopulmonary dysplasia, yet the transition from acute molecular injury to chronic pathology warrants further investigation.

## 5. CONCLUSION

Our study provides a genome-wide characterization of the lung transcriptome in a well-validated preterm piglet model. The investigation revealed that severe infection and sepsis trigger profound molecular alterations, specifically the upregulation of pro-inflammatory signaling and significant metabolic alterations. Crucially, our findings highlight that a restricted-glucose-supply strategy significantly attenuates the magnitude of these transcriptome alterations and reduces the severity of lung tissue damage compared with conventional high-glucose interventions. This indicates that the pulmonary manifestations of neonatal sepsis are highly sensitive to glycemic control, with high glucose levels potentially exacerbating inflammatory and cell death pathways. Furthermore, we have provided a robust framework for researchers by integrating these complex datasets into a novel, user-friendly web application to bridge the gap between transcriptome signatures and the identification of therapeutic targets in immunometabolism. Ultimately, these results underscore the potential of metabolic modulation as a precision-medicine approach to mitigate sepsis-induced lung injury in the vulnerable preterm population, though further research is warranted to establish standardized pulmonary injury scores and clinical protocols for immunometabolic therapy.

## AUTHOR CONTRIBUTIONS

**Nguyen Phuoc Long:** Writing – review & editing, Writing – original draft, Software, Visualization, Software, Methodology, Investigation, Formal analysis, Data curation, Conceptualization. **Ole Bæk:** Writing – review & editing, Methodology, Data curation, Conceptualization. **Karoline Aasmul-Olsen:** Writing – review & editing, Data curation, Investigation. **Richard Doughty:** Writing – review & editing, Investigation, Formal analysis. **Bjorn Klabunde:** Writing – review & editing, Investigation. **Nguyen Quang Thu:** Writing – review & editing, Formal analysis, Data curation, Visualization. **Le Hoang Bach Dat:** Writing – review & editing, Formal analysis, Software, Visualization. **Bui Thanh Liem:** Writing – review & editing, Investigation. **Klaus Bønnelykke:** Writing – review & editing, Supervision, Resources. **Duc Ninh Nguyen:** Writing – review & editing, Validation, Supervision, Methodology, Resources, Conceptualization.

## ACKNOWLEDGMENTS

We acknowledge all the support of personnel from the Comparative Pediatrics group (University of Copenhagen) during the animal study.

## DECLARATION OF GENERATIVE AI IN SCIENTIFIC WRITING

Gemini and Grammarly were used to check grammar and improve the readability of the manuscript. The authors revised and approved the content, accepting full responsibility for the publication.

## DATA AVAILABILITY STATEMENT

The data supporting this study’s findings are available upon reasonable request.

## REFERENCES

Agresti, A., & Natarajan, R. (2001). Modeling Clustered Ordered Categorical Data: A Survey. International Statistical Review, 69(3), 345–371. 10.1111/j.1751-5823.2001.tb00463.x

Bæk, O., Muk, T., Wu, Z., Ye, Y., Khakimov, B., Casano, A. M.,…Nguyen, D. N. (2025). Altered hepatic metabolism mediates sepsis preventive effects of reduced glucose supply in infected preterm newborns. eLife, 13, RP97830. 10.7554/eLife.97830

Balany, J., & Bhandari, V. (2015). Understanding the Impact of Infection, Inflammation, and Their Persistence in the Pathogenesis of Bronchopulmonary Dysplasia. Frontiers in Medicine, Volume 2 - 2015 10.3389/fmed.2015.00090

Bolognese, A. C., Yang, W.-L., Hansen, L. W., Denning, N.-L., Nicastro, J. M., Coppa, G. F., & Wang, P. (2018). Inhibition of necroptosis attenuates lung injury and improves survival in neonatal sepsis. Surgery, 164(1), 110–116. 10.1016/j.surg.2018.02.017

Briassoulis, G., Tzermia, K., Bastaki, K., Miliaraki, M., Briassoulis, P., Damianaki, A.,…Ilia, S. (2025). Necroptotic and Apoptotic Pathways in Sepsis: A Comparative Analysis of Pediatric and Adult ICU Patients. Biomedicines, 13(7), 1747. 10.3390/biomedicines13071747

Brunse, A., Worsøe, P., Pors, S. E., Skovgaard, K., & Sangild, P. T. (2019). Oral Supplementation With Bovine Colostrum Prevents Septic Shock and Brain Barrier Disruption During Bloodstream Infection in Preterm Newborn Pigs. Shock, 51(3)10.1097/SHK.0000000000001131

Chakraborty, M., McGreal, E. P., & Kotecha, S. (2010). Acute Lung Injury in Preterm Newborn Infants: Mechanisms and Management. Paediatric Respiratory Reviews, 11(3), 162–170. 10.1016/j.prrv.2010.03.002

Ewald, J. D., Zhou, G., Lu, Y., Kolic, J., Ellis, C., Johnson, J. D.,…Xia, J. (2024). Web-based multi-omics integration using the Analyst software suite. Nature Protocols, 19(5), 1467–1497. 10.1038/s41596-023-00950-4

Fallon, E. A., Chung, C.-S., Heffernan, D. S., Chen, Y., De Paepe, M. E., & Ayala, A. (2021). Survival and Pulmonary Injury After Neonatal Sepsis: PD1/PDL1’s Contributions to Mouse and Human Immunopathology. Frontiers in Immunology, 12 10.3389/fimmu.2021.634529

Fleiss, N., Coggins, S. A., Lewis, A. N., Zeigler, A., Cooksey, K. E., Walker, L. A.,…Wynn, J. L. (2021). Evaluation of the Neonatal Sequential Organ Failure Assessment and Mortality Risk in Preterm Infants With Late-Onset Infection. JAMA Network Open, 4(2), e2036518–e2036518. 10.1001/jamanetworkopen.2020.36518

Gabehart, K., Correll, K. A., Yang, J., Collins, M. L., Loader, J. E., Leach, S.,…Dakhama, A. (2014). Transcriptome Profiling of the Newborn Mouse Lung Response to Acute Ozone Exposure. Toxicological Sciences, 138(1), 175–190. 10.1093/toxsci/kft276

Gu, Z. (2022). Complex heatmap visualization. iMeta, 1(3), e43. 10.1002/imt2.43

Holzfurtner, L., Shahzad, T., Dong, Y., Rekers, L., Selting, A., Staude, B.,…Ehrhardt, H. (2022). When inflammation meets lung development—an update on the pathogenesis of bronchopulmonary dysplasia. Molecular and Cellular Pediatrics, 9(1), 7. 10.1186/s40348-022-00137-z

Kenward, M. G., & Roger, J. H. (1997). Small Sample Inference for Fixed Effects from Restricted Maximum Likelihood. Biometrics, 53(3), 983–997. 10.2307/2533558

Lahra, M. M., Beeby, P. J., & Jeffery, H. E. (2009). Intrauterine Inflammation, Neonatal Sepsis, and Chronic Lung Disease: A 13-Year Hospital Cohort Study. Pediatrics, 123(5), 1314–1319. 10.1542/peds.2008-0656

Laird, N. M., & Ware, J. H. (1982). Random-Effects Models for Longitudinal Data. Biometrics, 38(4), 963–974. 10.2307/2529876

Long, N. P., & Thu, N. Q. (2026). EasyFigAssembler: Enhance Omics Data Storytelling through Effective Figure Assembly. Journal of Proteome Research, 25(4), 2193–2199. 10.1021/acs.jproteome.5c01022

McCullagh, P. (1980). Regression Models for Ordinal Data. Journal of the Royal Statistical Society: Series B (Methodological), 42(2), 109–127. 10.1111/j.2517-6161.1980.tb01109.x

Muk, T., Brunse, A., Henriksen, N. L., Aasmul-Olsen, K., & Nguyen, D. N. (2022). Glucose supply and glycolysis inhibition shape the clinical fate of Staphylococcus epidermidis-infected preterm newborns. JCI Insight, 7(11)10.1172/jci.insight.157234

Ng, P. C., Li, K., Wong, R. P. O., Chui, K., Wong, E., Li, G., & Fok, T. F. (2003). Proinflammatory and anti-inflammatory cytokine responses in preterm infants with systemic infections. Archives of Disease in Childhood - Fetal and Neonatal Edition, 88(3), F209–F213. 10.1136/fn.88.3.F209

R Core Team. (2021). shinythemes: Themes for Shiny. Vienna, Austria.

R Core Team. (2023). R: A Language and Environment for Statistical Computing. R Foundation for Statistical Computing. Vienna, Austria.

R Core Team. (2026a). shiny: Web Application Framework for R. Vienna, Austria.

R Core Team. (2026b). shinyjs: Easily Improve the User Experience of Your Shiny Apps in Seconds. Vienna, Austria.

R Core Team. (2026c). shinyWidgets: Custom Inputs Widgets for Shiny. Vienna, Austria.

Salimi, U., Dummula, K., Tucker, M. H., Dela Cruz, C. S., & Sampath, V. (2022). Postnatal Sepsis and Bronchopulmonary Dysplasia in Premature Infants: Mechanistic Insights into “New BPD”. American journal of respiratory cell and molecular biology, 66(2), 137–145. 10.1165/rcmb.2021-0353PS

Taneri, P. E., Biesty, L., Kirkham, J. J., Molloy, E. J., Polin, R. A., Branagan, A.,…Devane, D. (2025). Proposed Core Outcomes After Neonatal Sepsis: A Consensus Statement. JAMA Network Open, 8(2), e2461554–e2461554. 10.1001/jamanetworkopen.2024.61554

Tien, N. T. N., Thu, N. Q., Kim, D. H., Park, S., & Long, N. P. (2025). EasyPubPlot: A Shiny Web Application for Rapid Omics Data Exploration and Visualization. Journal of Proteome Research, 24(4), 2188–2195. 10.1021/acs.jproteome.4c01068

Tucker, M. H., Yeh, H.-W., Oh, D., Shaw, N., Kumar, N., & Sampath, V. (2023). Preterm sepsis is associated with acute lung injury as measured by pulmonary severity score. Pediatric Research, 93(4), 1050–1056. 10.1038/s41390-022-02218-1

Wu, T., Hu, E., Xu, S., Chen, M., Guo, P., Dai, Z.,…Yu, G. (2021). clusterProfiler 4.0: A universal enrichment tool for interpreting omics data. The Innovation, 2(3), 100141. 10.1016/j.xinn.2021.100141

Wu, Z., Tien, N. T. N., Bæk, O., Zhong, J., Klabunde, B., Nguyen, T. T.,…Nguyen, D. N. (2024). Regulation of host metabolism and defense strategies to survive neonatal infection. Biochimica et Biophysica Acta (BBA) - Molecular Basis of Disease, 1870(8), 167482. 10.1016/j.bbadis.2024.167482

Zhong, J., Bæk, O., Doughty, R., Jørgensen, B. M., Jensen, H. E., Thymann, T.,…Nguyen, D. N. (2025). Reduced parenteral glucose supply during neonatal infection attenuates neurological and renal pathology associated with modulation of innate and Th1 immunity. Biochimica et Biophysica Acta (BBA) - Molecular Basis of Disease, 1871(4), 167723. 10.1016/j.bbadis.2025.167723

