## Supplementary File for "Reduced parenteral glucose supply in preterm neonatal infection ameliorates the pulmonary damage"

### SUPPLEMENTARY FIGURES

**Supplementary Figure S1. Biochemical test results of the piglets after infection and parenteral nutrition for 22 h.** (A) Blood pH. (B) Blood standard base excess (cBase (Ecf)) (mmol/L). (C) Blood standard bicarbonate (SBC) (mmol/L). (D) Blood lactate (mmol/L). (E) Blood partial pressure of carbon dioxide ($pCO_2$) (kPa). (F) Blood partial pressure of oxygen ($pO_2$) (kPa). Abbreviations: ns: not significant; *: P-value < 0.05; **: P-value < 0.01; ***: P-value < 0.001.

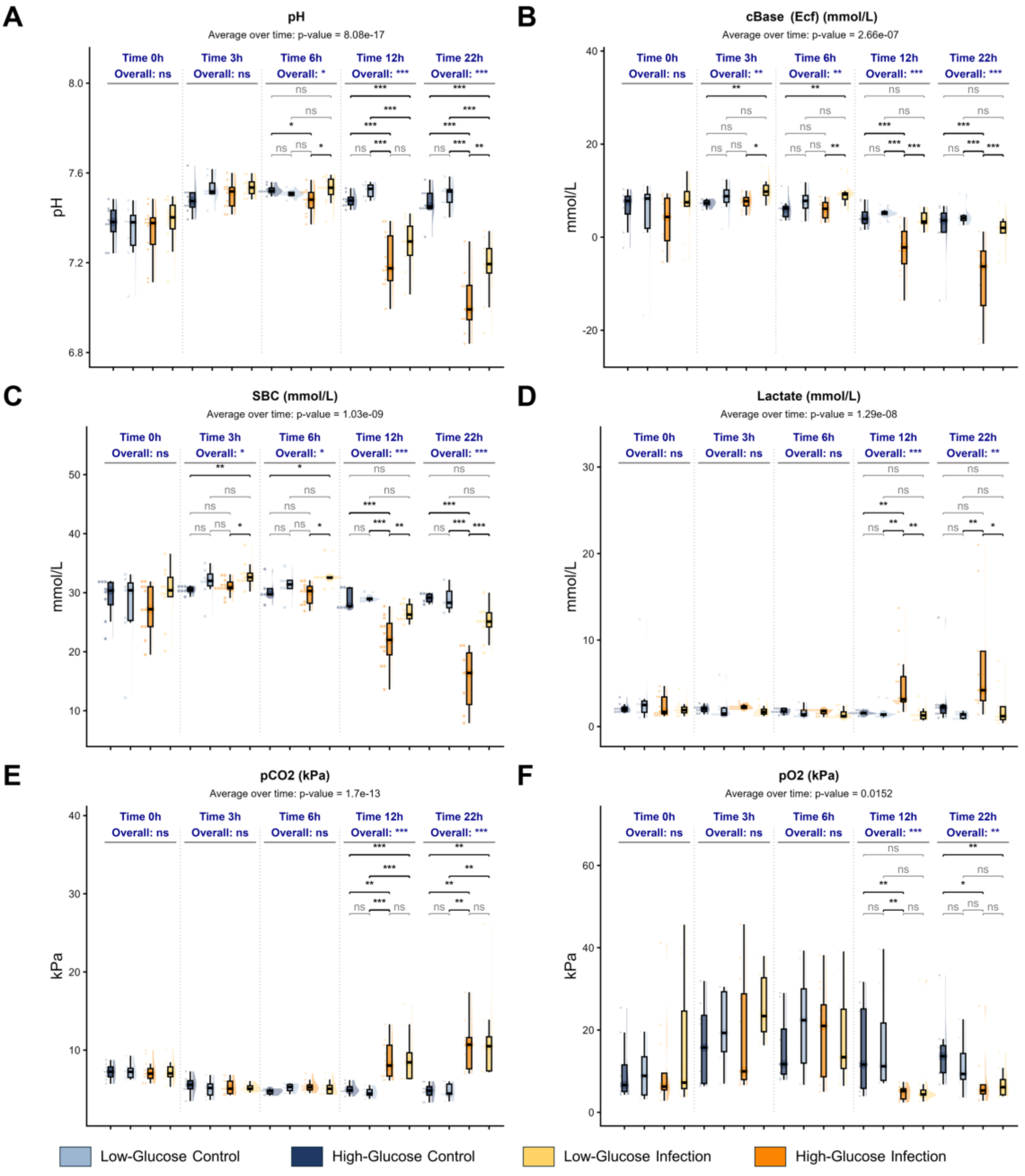

**Supplementary Figure S2. The alterations of pulmonary transcriptome profiles in neonatal piglets under sepsis.** (A) Volcano plot of DEGs between Infection and Control. (B) GSEA dot plot of top activated and suppressed pathways related to the infection, GO:BP database. (C-E) Heatmap of altered pathways related to (C) response to bacterium; (D) inflammation; (E) immune response. Abbreviations: DEGs: Differentially expressed genes; FC: Fold change; FDR: False discovery rate; GO:BP: Biological Processes in Gene Ontology; GSEA: Gene set enrichment analysis.

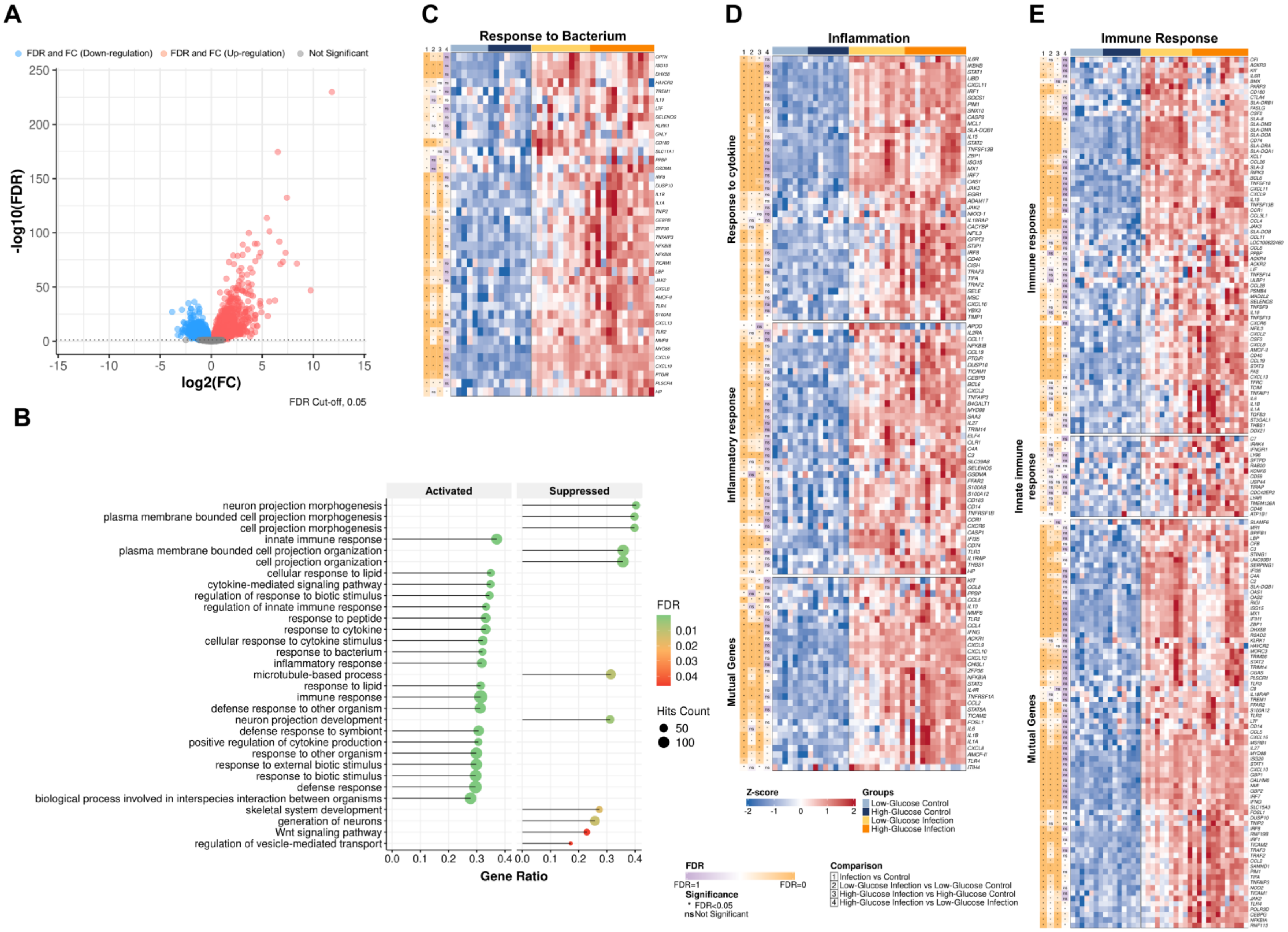
A
FDR and FC (Down-regulation)
FDR and FC (Up-regulation)
Not Significant
-log10(FDR)
log2(FC)
FDR Cut-off, 0.05
B
Activated
Suppressed
neuron projection morphogenesis
plasma membrane bounded cell projection morphogenesis
cell projection morphogenesis
innate immune response
plasma membrane bounded cell projection organization
cell projection organization
cellular response to lipid
cytokine-mediated signaling pathway
regulation of response to biotic stimulus
regulation of innate immune response
response to peptide
response to cytokine
cellular response to cytokine stimulus
response to bacterium
inflammatory response
microtubule-based process
response to lipid
immune response
defense response to other organism
neuron projection development
defense response to symbiont
positive regulation of cytokine production
response to other organism
response to external biotic stimulus
response to biotic stimulus
defense response
biological process involved in interspecies interaction between organisms
skeletal system development
generation of neurons
Wnt signaling pathway
regulation of vesicle-mediated transport
Gene Ratio
FDR
Hits Count
C
Response to Bacterium
D
Inflammation
Response to cytokine
Inflammatory response
Mutual Genes
E
Immune Response
Immune response
Innate immune response
Mutual Genes
Z-score
Groups
Low-Glucose Control
High-Glucose Control
Low-Glucose Infection
High-Glucose Infection
FDR
FDR=1
FDR=0
Significance
* FDR<0.05
ns Not Significant
Comparison
1 Infection vs Control
2 Low-Glucose Infection vs Low-Glucose Control
3 High-Glucose Infection vs High-Glucose Control
4 High-Glucose Infection vs Low-Glucose Infection

**Supplementary Figure S3. Conventional glucose dose led to profound alteration of pulmonary transcriptome profiles between sepsis and control piglets.** (A) Volcano plot of DEGs between High-glucose infection vs High-glucose control. (B) GSEA dot plot of top activated and suppressed pathways related to the infection and high-dose glucose treatment, GO:BP database. (C-E) Heatmap of altered pathways related to (C) response to bacterium; (D) inflammation; (E) immune response. Abbreviations: DEGs: Differentially expressed genes; FC: Fold change; FDR: False discovery rate; GO:BP: Biological Processes in Gene Ontology; GSEA: Gene set enrichment analysis.

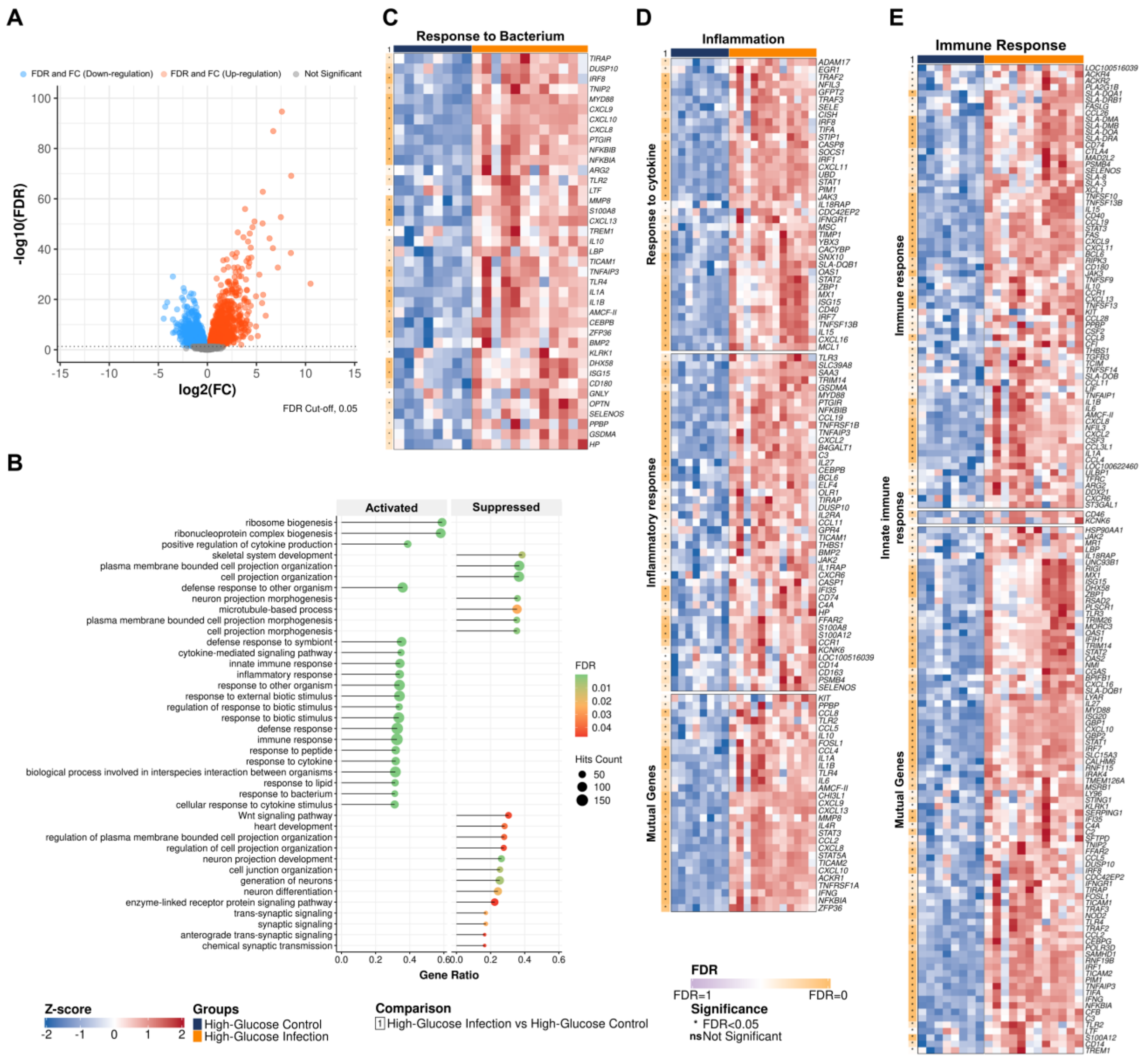

**Supplementary Figure S4. Reduced glucose dose resulted in mild alteration of lung transcriptome profiles between infected and healthy neonates.** (A) Volcano plot of DEGs between Low-glucose infection vs Low-glucose control. (B) GSEA dot plot of top activated and suppressed pathways related to the infection and low-dose glucose treatment, GO:BP database. (C-E) Heatmap of altered pathways related to (C) response to bacterium; (D) inflammation; (E) immune response. Abbreviations: DEGs: Differentially expressed genes; FC: Fold change; FDR: False discovery rate; GO:BP: Biological Processes in Gene Ontology; GSEA: Gene set enrichment analysis.

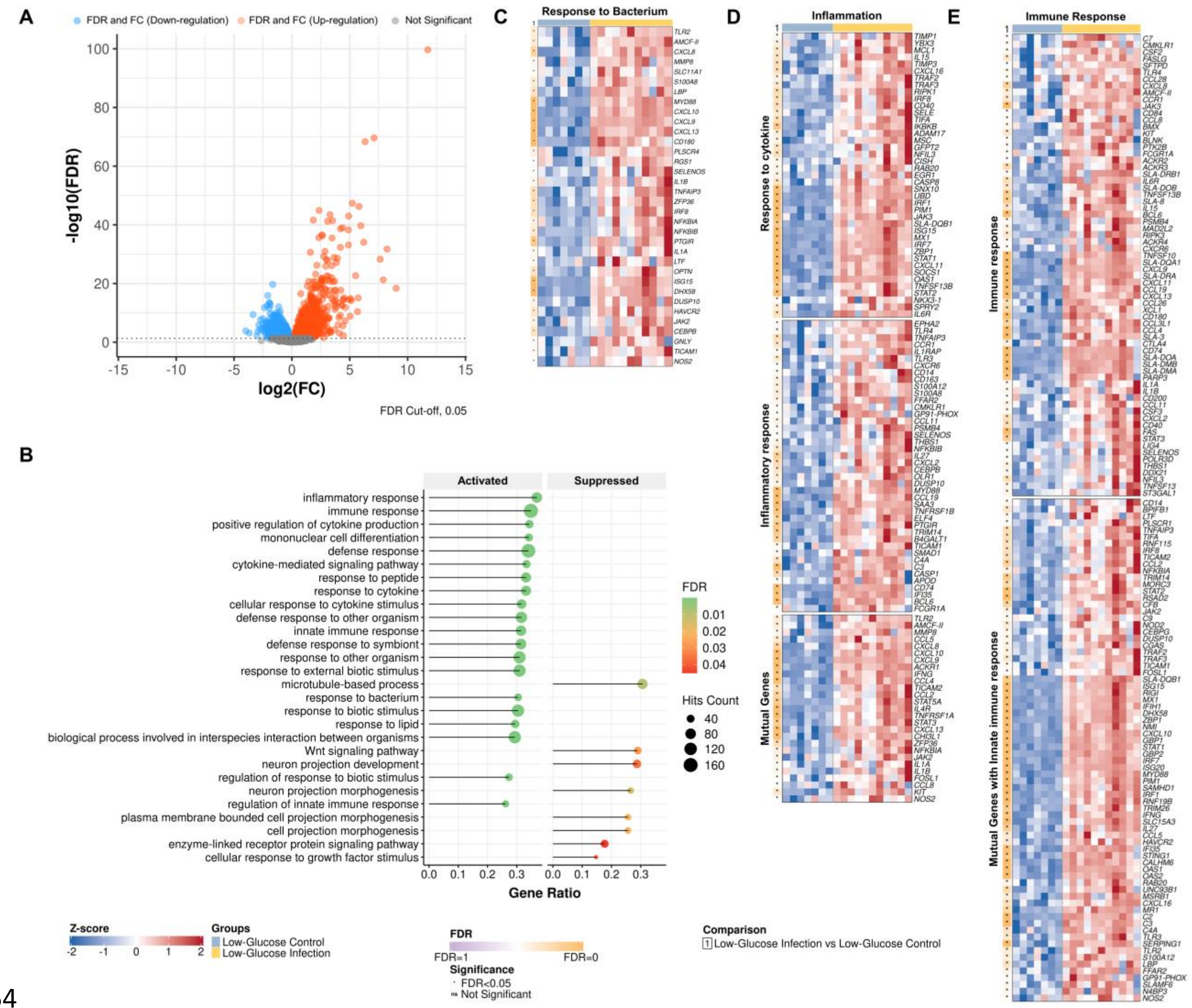

**Supplementary Figure S5. Low-glucose supply resulted in the suppression of cell-cycle-related pathways in the infected piglets.** (A) Volcano plot of DEGs between High-glucose infection vs Low-glucose infection. (B) GSEA dot plot of top activated and suppressed pathways related to the glucose treatment doses, GO:BP database. (c-e) Heatmap of altered pathways related to (C) response to bacterium; (D) inflammation; (E-F) cell cycle. Abbreviations: DEGs: Differentially expressed genes; FC: Fold change; FDR: False discovery rate; GO:BP: Biological Processes in Gene Ontology; GSEA: Gene set enrichment analysis.

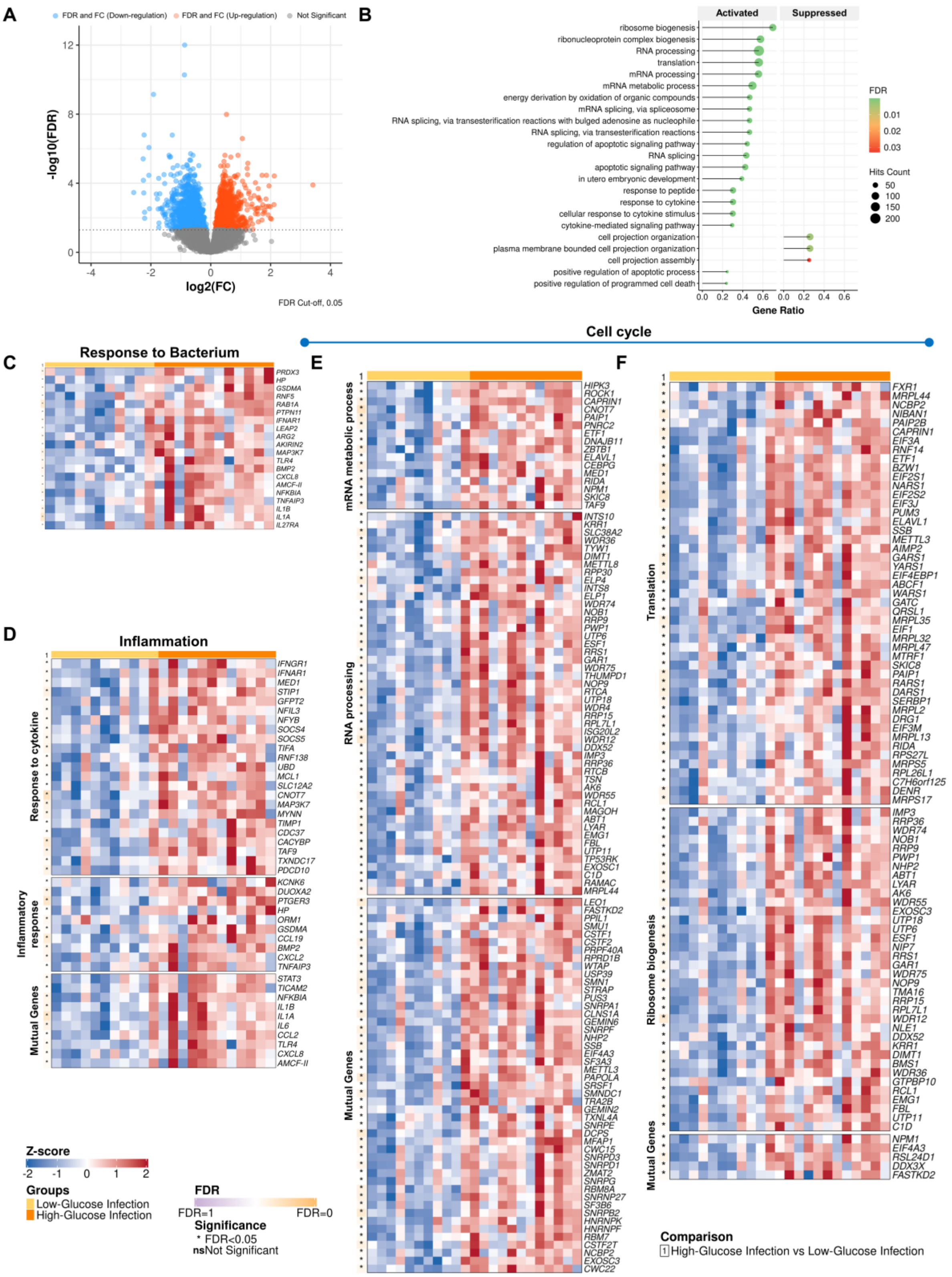
A
FDR and FC (Down-regulation)
FDR and FC (Up-regulation)
Not Significant
-log10(FDR)
log2(FC)
FDR Cut-off, 0.05
B
Activated
Suppressed
ribosome biogenesis
ribonucleoprotein complex biogenesis
RNA processing
translation
mRNA processing
mRNA metabolic process
energy derivation by oxidation of organic compounds
mRNA splicing, via spliceosome
RNA splicing, via transesterification reactions with bulged adenosine as nucleophile
RNA splicing, via transesterification reactions
regulation of apoptotic signaling pathway
RNA splicing
apoptotic signaling pathway
in utero embryonic development
response to peptide
response to cytokine
cellular response to cytokine stimulus
cytokine-mediated signaling pathway
cell projection organization
plasma membrane bounded cell projection organization
cell projection assembly
positive regulation of apoptotic process
positive regulation of programmed cell death
Gene Ratio
FDR
Hits Count
Cell cycle
C
Response to Bacterium
D
Inflammation
Response to cytokine
Inflammatory response
Mutual Genes
E
mRNA metabolic process
RNA processing
Mutual Genes
F
Translation
Ribosome biogenesis
Mutual Genes
Z-score
Groups
Low-Glucose Infection
High-Glucose Infection
FDR
FDR=1
FDR=0
Significance
* FDR<0.05
ns Not Significant
Comparison
1 High-Glucose Infection vs Low-Glucose Infection

62

**Supplementary Figure S6. Difference in glucose supplement doses caused no alterations in the lung transcriptome of control piglets.** (A) PCA scores plot. (B) Volcano plot. Abbreviations: FC: Fold change; FDR: False discovery rate; PCA: Principal component analysis.

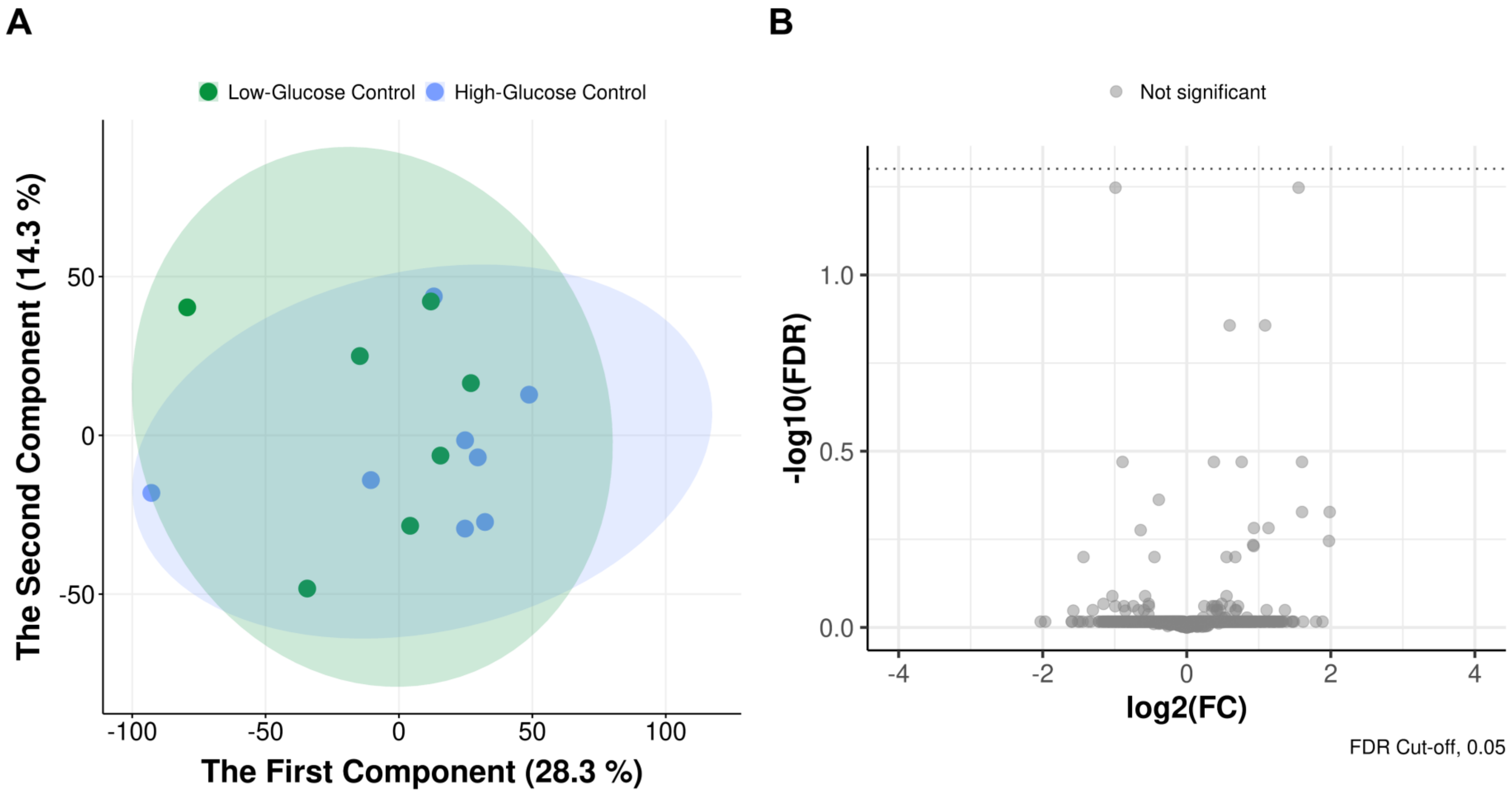

### SUPPLEMENTARY TABLES

**Supplementary Table S1. Significantly enriched pathways from the GSEA result of the KEGG database, Infection versus Control.**

Abbreviations: GSEA: Gene set enrichment analysis; KEGG: Kyoto Encyclopedia of Genes and Genomes.

| KEGG ID | Pathways | Set size | NES | P-value | FDR | Core enrichment gene |
|---|---|---|---|---|---|---|
| ssc05168 | Herpes simplex virus 1 infection | 140 | 2.78 | 2.17E-24 | 7.48E-22 | *IFNG, OAS1, TAP1, CCL2, OAS2, IRF7, RIGI, IL1B, STAT1, BST2, TAP2, SLA-DOA, BIRC3, IL6, FAS, SOCS3, IFIH1, SLA-DRA, EIF2AK2, SLA-3, SLA-DQA1, IRF9, SLA-DOB, CD74, SLA-DQB1, B2M, RNASEL, TAPBP, FASLG, SLA-DMB, STAT2, SLA-7, CCL5, PML, SLA-DMA, SLA-DRB1, C3, TAPBPL, IFNAR2, LOC100517129, NFKBIA, BAK1, TNFSF14, IKBKE, CASP3, PDIA3, CYCS, STING1, TRAF3, TRAF2, CGAS, MYD88, EIF4EBP1, TNFRSF1A, CASP8, ZC3HAV1, CALR, DAXX, TICAM1, ITGA5, SLA-8, TLR2, TLR3, NFKB1, IKBKB, JAK2* |
| ssc05164 | Influenza A | 127 | 2.74 | 1.16E-21 | 2.00E-19 | *CXCL10, IFNG, RSAD2, MX1, OAS1, CCL2, CXCL8, OAS2, IRF7, RIGI, IL1B, STAT1, SLA-DOA, IL6, FAS, SOCS3, IL1A, IFIH1, TNFSF10, SLA-DRA, EIF2AK2, SLA-DQA1, IRF9, SLA-DOB, SLA-DQB1, RNASEL, FASLG, SLA-DMB, STAT2, CCL5, ADAR, PML, SLA-DMA, SLA-DRB1, CASP1, NFKBIB, IFNAR2, LOC100624226, NFKBIA, TRIM25, BAK1, IKBKE, CASP3, CYCS, TRAF3, MYD88, TNFRSF1A, ICAM1, CASP8, TICAM1, TLR4, KPNA2, TLR3, NFKB1, IKBKB, JAK2, DNAJC3, CALCOCO2, EIF2S1, SLC25A5, IFNGR1, IRAK4, IFNGR2, TMPRSS4* |
| ssc04668 | TNF signaling pathway | 104 | 2.69 | 1.96E-18 | 2.25E-16 | *CXCL10, SELE, AMCF-II, IRF1, CXCL2, CCL2, IL1B, BIRC3, IL6, FAS, SOCS3, CX3CL1, RHBDF2, BCL3, MAP3K8, TNFAIP3, JUNB, VCAM1, CASP10, FOS, CASP7, IL15, CCL5, NOD2, CSF1, MMP9, CSF2, LIF, PTGS2, NFKBIA, MLKL, CASP3, PGAM5, TRAF3, TRAF2, CEBPB, CFLAR, ADAM17, TNFRSF1A, ICAM1, CASP8, RIPK3, CREB3L1, TNFRSF1B, NFKB1, IKBKB* |
| ssc04620 | Toll-like receptor signaling pathway | 87 | 2.54 | 7.96E-14 | 3.92E-12 | *CXCL9, CXCL11, CXCL10, CCL4, CXCL8, IRF7, IL1B, CCL3L1, STAT1, LBP, IL6, IRF9, MAP3K8, STAT2, FOS, CD40, CCL5, TICAM2, IFNAR2, NFKBIA, IKBKE, TRAF3, CD14, MYD88, CASP8, TICAM1, TLR2, TLR4, TLR3, NFKB1, IKBKB* |

| | | | | | | |
|---|---|---|---|---|---|---|
| ssc04657 | IL-17 signaling pathway | 70 | 2.54 | 1.70E-13 | 7.31E-12 | *CXCL10*, *IFNG*, *AMCF-II*, *CXCL2*, *CCL2*, *CXCL8*, *IL1B*, *S100A8*, *S100A9*, *CCL11*, *IL6*, *CSF3*, *FOSL1*, *TNFAIP3*, *MMP1*, *FOS*, *MUC5AC*, *MMP9*, *CSF2*, *PTGS2*, *NFKBIA*, *HSP90B1*, *IKBKE*, *CASP3*, *LCN2*, *TRAF3*, *TRAF2*, *CEBPB*, *CASP8*, *TRAF3IP2*, *MMP13*, *NFKB1*, *IKBKB* |
| ssc05169 | Epstein-Barr virus infection | 168 | 2.54 | 3.40E-17 | 2.35E-15 | *CXCL10*, *ISG15*, *OAS1*, *TAP1*, *OAS2*, *IRF7*, *RIGI*, *STAT1*, *TAP2*, *SLA-DOA*, *IL6*, *FAS*, *SLA-DRA*, *EIF2AK2*, *SLA-3*, *SLA-DQA1*, *IRF9*, *RELB*, *TNFAIP3*, *SLA-DOB*, *SLA-DQB1*, *CDKN1A*, *B2M*, *TAPBP*, *SLA-DMB*, *STAT2*, *CD40*, *SLA-7*, *GADD45G*, *SLA-DMA*, *SLA-DRB1*, *GADD45A*, *NFKBIB*, *NFKB2*, *TAPBPL*, *NFKBIE*, *IFNAR2*, *NFKBIA*, *STAT3*, *BAK1*, *IKBKE*, *CASP3*, *PDIA3*, *CYCS*, *TRAF3*, *TRAF2*, *JAK3*, *MYD88*, *ICAM1*, *CASP8*, *CALR*, *MYC*, *SLA-8*, *TLR2*, *PSMD14*, *GADD45B*, *NFKB1*, *IKBKB* |
| ssc04612 | Antigen processing and presentation | 47 | 2.53 | 6.96E-13 | 2.40E-11 | *IFNG*, *TAP1*, *TAP2*, *SLA-DOA*, *PSME2*, *SLA-DRA*, *SLA-3*, *CTSL*, *SLA-DQA1*, *SLA-DOB*, *CD74*, *SLA-DQB1*, *B2M*, *PSME1*, *TAPBP*, *SLA-DMB*, *SLA-7*, *SLA-DMA*, *SLA-DRB1*, *HSPA5*, *TAPBPL*, *IFI30*, *RFX5*, *HSPA6*, *PDIA3*, *LOC100523789*, *HSPA8*, *CALR*, *SLA-8* |
| ssc05171 | Coronavirus disease - COVID-19 | 181 | 2.52 | 4.29E-18 | 3.70E-16 | *CXCL10*, *ISG15*, *MX1*, *OAS1*, *CCL2*, *CXCL8*, *OAS2*, *RIGI*, *IL1B*, *STAT1*, *IL6*, *IFIH1*, *EIF2AK2*, *CSF3*, *IRF9*, *MMP1*, *C2*, *CFB*, *STAT2*, *F13A1*, *FOS*, *C4A*, *ADAR*, *C3*, *CASP1*, *SELP*, *NFKBIB*, *IFNAR2*, *CSF2*, *NFKBIA*, *STAT3*, *IKBKE*, *FGB*, *STING1*, *TRAF3*, *RPL13A*, *LOC100515049*, *CGAS*, *MYD88*, *ADAM17*, *TNFRSF1A*, *TLR2*, *TLR4*, *C9*, *TLR3*, *IL6R*, *NFKB1*, *IL6ST*, *IKBKB* |
| ssc04060 | Cytokine-cytokine receptor interaction | 170 | 2.50 | 2.65E-16 | 1.52E-14 | *CXCL9*, *CXCL11*, *CXCL10*, *IFNG*, *CCL19*, *CXCL13*, *AMCF-II*, *CXCL2*, *CCL2*, *CCL4*, *CCL26*, *CXCL8*, *IL1B*, *CCL3L1*, *IL27*, *TNFSF13B*, *CCL11*, *IL6*, *FAS*, *IL1A*, *TNFSF10*, *CX3CL1*, *CSF3*, *CCL8*, *LOC110260995*, *IL4R*, *IL1R2*, *CSF3R*, *FASLG*, *XCL1*, *ACKR4*, *CD40*, *IL15*, *CCL5*, *IL2RB*, *GDF15*, *IL1RAP*, *CSF1*, *IL1R1*, *CXCR6*, *IFNAR2*, *CCR1*, *OSMR*, *LOC100624226*, *CCL28*, *CSF2*, *LIF*, *CNTFR*, *TNFSF14*, *CXCL16*, *IL10*, *TNFSF13*, *ACVR1B*, *PPBP*, *CSF2RB*, *IL2RA*, *IL18RAP*, *TNFRSF1A*, *IL10RB*, *ACKR3*, *TNFSF9*, *TNFRSF12A*, *RELL1*, *TGFB3*, *TNFRSF1B*, *RELT*, *IL21R*, *IL6R*, *IL6ST* |
| ssc05323 | Rheumatoid arthritis | 65 | 2.46 | 1.05E-11 | 3.03E-10 | *IFNG*, *AMCF-II*, *CXCL2*, *CCL2*, *CXCL8*, *IL1B*, *CCL3L1*, *TNFSF13B*, *SLA-DOA*, *IL6*, *IL1A*, *SLA-DRA*, *CTSL*, *SLA-DQA1*, *MMP1*, *SLA-DOB*, *CTLA4*, *SLA-DQB1*, *SLA-DMB*, *FOS*, *IL15*, *CCL5*, *SLA-DMA*, *SLA-DRB1*, *CSF1*, *TCIRG1*, *CSF2*, *TNFSF13*, *ICAM1*, *ATP6V1C2*, *TLR2*, *TLR4*, *TGFB3* |

| | | | | | | |
|---|---|---|---|---|---|---|
| ssc05162 | Measles | 108 | 2.45 | 3.82E-13 | 1.46E-11 | *MX1*, *OAS1*, *OAS2*, *IRF7*, *RIGI*, *IL1B*, *STAT1*, *IL6*, *FAS*, *IL1A*, *IFIH1*, *SLAMF1*, *EIF2AK2*, *IRF9*, *TNFAIP3*, *FASLG*, *STAT2*, *FOS*, *IL2RB*, *ADAR*, *STAT5A*, *FCGR2B*, *NFKBIB*, *IFNAR2*, *NFKBIA*, *STAT3*, *BAK1*, *IKBKE*, *CASP3*, *HSPA6*, *CYCS*, *TRAF3*, *IL2RA*, *JAK3*, *MYD88*, *HSPA8*, *CASP8*, *TLR2*, *TLR4*, *NFKB1*, *IKBKB*, *IL2RG*, *EIF2S1*, *TP53*, *CD46*, *IRAK4* |
| ssc04621 | NOD-like receptor signaling pathway | 127 | 2.44 | 9.81E-13 | 3.08E-11 | *GBP1*, *GBP2*, *OAS1*, *CXCL2*, *CCL2*, *CXCL8*, *OAS2*, *IRF7*, *IL1B*, *STAT1*, *BIRC3*, *IL6*, *LOC100522887*, *IRF9*, *TNFAIP3*, *RNASEL*, *STAT2*, *CCL5*, *NOD2*, *CASP1*, *NFKBIB*, *NAMPT*, *IFNAR2*, *NFKBIA*, *IKBKE*, *GSDMD*, *TRPM2*, *STING1*, *TRAF3*, *TRAF2*, *TXNIP*, *MYD88*, *CASP8*, *TICAM1*, *RIPK2*, *RIPK3*, *TLR4*, *NFKB1*, *IKBKB*, *NLRP12*, *TANK*, *RIPK1* |
| ssc04061 | Viral protein interaction with cytokine and cytokine receptor | 60 | 2.41 | 2.31E-10 | 5.70E-09 | *CXCL9*, *CXCL11*, *CXCL10*, *CCL19*, *CXCL13*, *AMCF-II*, *CXCL2*, *CCL2*, *CCL4*, *CCL26*, *CXCL8*, *CCL3L1*, *CCL11*, *IL6*, *TNFSF10*, *CX3CL1*, *CCL8*, *XCL1*, *ACKR4*, *CCL5*, *IL2RB*, *CSF1*, *CCR1*, *LOC100624226*, *CCL28*, *TNFSF14*, *IL10*, *PPBP*, *IL2RA*, *IL18RAP*, *TNFRSF1A*, *IL10RB*, *ACKR3*, *TNFRSF1B*, *IL6R*, *IL6ST*, *IL2RG* |
| ssc04622 | RIG-I-like receptor signaling pathway | 54 | 2.39 | 2.30E-09 | 3.53E-08 | *CXCL10*, *ISG15*, *DHX58*, *CXCL8*, *IRF7*, *RIGI*, *IFIH1*, *ZNFX1*, *CASP10*, *ADAR*, *NFKBIB*, *NFKBIA*, *TRIM25*, *IKBKE*, *STING1*, *TRAF3*, *TRAF2*, *CASP8*, *NFKB1*, *IKBKB*, *TANK*, *RIPK1* |
| ssc04623 | Cytosolic DNA-sensing pathway | 59 | 2.38 | 1.49E-09 | 2.57E-08 | *CXCL10*, *CCL4*, *IRF7*, *ZBP1*, *RIGI*, *IL1B*, *SAMHD1*, *IL6*, *CASP7*, *CCL5*, *ADAR*, *CASP1*, *NFKBIB*, *NFKBIA*, *MLKL*, *IKBKE*, *TREX1*, *CASP3*, *GSDMD*, *STING1*, *CGAS*, *CASP8*, *POLR3D*, *RIPK3*, *NFKB1*, *IKBKB*, *POLR3E*, *RIPK1* |
| ssc05160 | Hepatitis C | 120 | 2.36 | 5.58E-11 | 1.48E-09 | *CXCL10*, *IFNG*, *RSAD2*, *MX1*, *IFIT1*, *OAS1*, *OAS2*, *IRF7*, *RIGI*, *STAT1*, *FAS*, *SOCS3*, *EIF2AK2*, *IRF9*, *CDKN1A*, *RNASEL*, *FASLG*, *STAT2*, *IFNAR2*, *NFKBIA*, *STAT3*, *BAK1*, *IKBKE*, *CASP3*, *CYCS*, *TRAF3*, *TRAF2*, *CFLAR*, *TNFRSF1A*, *CASP8*, *TICAM1*, *MYC*, *TLR3*, *CLDN5*, *NFKB1*, *IKBKB* |
| ssc04064 | NF-kappa B signaling pathway | 87 | 2.35 | 1.03E-09 | 1.86E-08 | *CCL19*, *CXCL2*, *CCL4*, *CXCL8*, *RIGI*, *IL1B*, *TNFSF13B*, *LBP*, *BIRC3*, *RELB*, *TNFAIP3*, *VCAM1*, *CD40*, *GADD45G*, *TICAM2*, *GADD45A*, *NFKB2*, *IL1R1*, *PTGS2*, *NFKBIA*, *TRIM25*, *TNFSF14*, *TRAF3*, *TRAF2*, *CD14*, *CFLAR*, *MYD88*, *BCL2A1*, *TNFRSF1A*, *ICAM1*, *TICAM1*, *TLR4*, *GADD45B*, *NFKB1*, *IKBKB*, *RIPK1*, *TIRAP*, *LTBR*, *IRAK4*, *ERC1*, *LY96*, *TAB3* |

| | | | | | | |
|---|---|---|---|---|---|---|
| ssc05321 | Inflammatory bowel disease | 45 | 2.32 | 4.58E-08 | 6.17E-07 | *IFNG*, *IL1B*, *STAT1*, *SLA-DOA*, *IL6*, *IL1A*, *SLA-DRA*, *SLA-DQA1*, *SLA-DOB*, *SLA-DQB1*, *IL4R*, *SLA-DMB*, *NOD2*, *SLA-DMA*, *SLA-DRB1*, *STAT3*, *IL10*, *IL18RAP*, *TBX21*, *TLR2*, *TLR4*, *TGFB3*, *IL21R*, *NFKB1*, *IL2RG* |
| ssc05152 | Tuberculosis | 137 | 2.28 | 3.66E-10 | 7.71E-09 | *IFNG*, *CLEC4E*, *LOC110258711*, *IL1B*, *STAT1*, *LBP*, *SLA-DOA*, *IL6*, *IL1A*, *SLA-DRA*, *SLA-DQA1*, *SLA-DOB*, *CD74*, *SLA-DQB1*, *SLA-DMB*, *CASP10*, *CYP27B1*, *NOD2*, *SLA-DMA*, *SLA-DRB1*, *C3*, *FCGR2B*, *TCIRG1*, *RFX5*, *IL10*, *CASP3*, *PLK3*, *CEBPG*, *CYCS*, *CEBPB*, *CD14*, *MYD88*, *HSPA9*, *TNFRSF1A*, *IL10RB*, *CASP8*, *RIPK2*, *CLEC7A*, *HSPD1*, *TLR2*, *TLR4*, *TGFB3*, *MRC1*, *NFKB1*, *JAK2*, *FCGR1A*, *TIRAP*, *IFNGR1*, *IRAK4*, *IFNGR2*, *VDR*, *CTSS*, *IRAK2* |
| ssc04659 | Th17 cell differentiation | 89 | 2.28 | 2.35E-09 | 3.53E-08 | *IFNG*, *IL1B*, *IL27*, *STAT1*, *SLA-DOA*, *IL6*, *FOXS1*, *SLA-DRA*, *SLA-DQA1*, *SLA-DOB*, *SLA-DQB1*, *IL4R*, *SLA-DMB*, *FOS*, *IL2RB*, *SLA-DMA*, *SLA-DRB1*, *STAT5A*, *IL1RAP*, *NFKBIB*, *NFKBIE*, *IL1R1*, *NFKBIA*, *STAT3*, *IL2RA*, *JAK3*, *TBX21*, *IL21R*, *IL6R*, *NFKB1*, *IL6ST*, *IKBKB*, *JAK2*, *IL2RG*, *HIF1A* |
| ssc04217 | Necroptosis | 109 | 2.27 | 1.69E-09 | 2.78E-08 | *IFNG*, *ZBP1*, *IL1B*, *STAT1*, *BIRC3*, *FAS*, *IL1A*, *TNFSF10*, *EIF2AK2*, *IRF9*, *TNFAIP3*, *FASLG*, *STAT2*, *STAT5A*, *TICAM2*, *CASP1*, *IFNAR2*, *LOC100624226*, *STAT3*, *GLUL*, *MLKL*, *PGAM5*, *FTH1*, *TRAF2*, *JAK3*, *CFLAR*, *CHMP7*, *TNFRSF1A*, *CASP8*, *TICAM1*, *RIPK3*, *TLR4*, *TLR3*, *JAK2*, *RIPK1*, *PPID*, *SLC25A5*, *IFNGR1*, *IFNGR2* |
| ssc05140 | Leishmaniasis | 59 | 2.26 | 1.10E-07 | 1.27E-06 | *IFNG*, *IL1B*, *STAT1*, *SLA-DOA*, *IL1A*, *SLA-DRA*, *SLA-DQA1*, *SLA-DOB*, *SLA-DQB1*, *SLA-DMB*, *FOS*, *SLA-DMA*, *SLA-DRB1*, *C3*, *NFKBIB*, *PTGS2*, *NFKBIA*, *IL10*, *MYD88*, *TLR2*, *TLR4*, *TGFB3*, *NFKB1*, *JAK2*, *FCGR1A*, *IFNGR1*, *IRAK4*, *IFNGR2* |
| ssc04630 | JAK-STAT signaling pathway | 110 | 2.25 | 4.03E-09 | 5.80E-08 | *IFNG*, *IL27*, *STAT1*, *IL6*, *SOCS1*, *SOCS3*, *CSF3*, *IRF9*, *CDKN1A*, *PIM1*, *IL4R*, *CSF3R*, *STAT2*, *IL15*, *IL2RB*, *STAT5A*, *IFNAR2*, *OSMR*, *CSF2*, *LIF*, *CISH*, *CNTFR*, *STAT3*, *IL10*, *CSF2RB*, *IL2RA*, *PTPN2*, *JAK3*, *IL10RB*, *MCL1*, *MYC*, *IL21R*, *IL6R*, *IL6ST*, *JAK2*, *IL2RG*, *PDGFRB*, *IFNGR1*, *SOCS7*, *IFNGR2* |
| ssc04141 | Protein processing in endoplasmic reticulum | 149 | 2.23 | 3.80E-10 | 7.71E-09 | *HYOU1*, *HSPA5*, *DNAJB11*, *HSP90B1*, *BAK1*, *PDIA4*, *HSPA6*, *WFS1*, *PDIA3*, *HSPH1*, *SEC24D*, *TRAF2*, *SEC23B*, *HERPUD1*, *DDIT3*, *CKAP4*, *HSPA8*, *DERL3*, *NGLY1*, *CALR*, *SELENOS*, *PREB*, *LMAN1*, *SEC61A1*, *DNAJC1*, *EDEM1*, *MOGS*, *SEC24A*, *DNAJA1*, *YOD1*, *DNAJC3*, *PDIA6*, *MAN1B1*, *UBE2D1*, *OSTC*, *RPN1*, *ERN1*, *RPN2*, *SSR1*, *P4HB*, *SSR3*, *UGGT1*, *EIF2S1*, *CANX*, |

| | | | | | | |
|---|---|---|---|---|---|---|
| | | | | | | *SEC13*, *SEC61B*, *TRAM1*, *STT3A*, *HSP90AA1*, *UBQLN1*, *UBE2D3*, *RAD23A*, *DNAJA2*, *VCP*, *HSPBP1*, *XBP1*, *ERO1B*, *STT3B*, *SAR1B*, *DNAJB1*, *EIF2AK3*, *RRBP1*, *PLAA*, *DERL2*, *SEC63*, *DAD1*, *SEL1L*, *ATF6*, *SEC24C*, *GANAB*, *MAGT1*, *EDEM2*, *CUL1*, *NPLOC4*, *BAG1*, *SEC31B*, *SYVN1* |
| ssc05145 | Toxoplasmosis | 94 | 2.22 | 4.65E-08 | 6.17E-07 | *IFNG*, *STAT1*, *SLA-DOA*, *BIRC3*, *SOCS1*, *SLA-DRA*, *SLA-DQA1*, *SLA-DOB*, *PPIF*, *SLA-DQB1*, *SLA-DMB*, *CD40*, *SLA-DMA*, *SLA-DRB1*, *NFKBIB*, *NFKBIA*, *STAT3*, *IL10*, *CASP3*, *HSPA6*, *CYCS*, *MYD88*, *HSPA8*, *TNFRSF1A*, *IL10RB*, *CASP8*, *TLR2*, *TLR4*, *TGFB3*, *NFKB1*, *IKBKB*, *ITGA6*, *JAK2*, *LAMB4*, *IFNGR1*, *IRAK4*, *IFNGR2*, *LY96* |
| ssc05332 | Graft-versus-host disease | 23 | 2.21 | 4.76E-07 | 5.28E-06 | *IFNG*, *IL1B*, *SLA-DOA*, *IL6*, *FAS*, *IL1A*, *SLA-DRA*, *SLA-3*, *SLA-DQA1*, *SLA-DOB*, *SLA-DQB1*, *FASLG*, *SLA-DMB*, *SLA-7*, *SLA-DMA*, *SLA-DRB1*, *PRF1*, *LOC100523789*, *SLA-8* |
| ssc05133 | Pertussis | 63 | 2.20 | 4.90E-07 | 5.28E-06 | *AMCF-II*, *IRF1*, *CXCL8*, *IL1B*, *IL6*, *IL1A*, *C2*, *IRF8*, *FOS*, *C4A*, *CASP7*, *TICAM2*, *SFTPA1*, *C3*, *CASP1*, *SERPING1*, *IL10*, *CASP3*, *C4BPB*, *CD14*, *MYD88*, *TICAM1*, *ITGA5*, *TLR4*, *NFKB1* |
| ssc04062 | Chemokine signaling pathway | 158 | 2.20 | 2.83E-10 | 6.52E-09 | *CXCL9*, *CXCL11*, *CXCL10*, *CCL19*, *CXCL13*, *AMCF-II*, *CXCL2*, *CCL2*, *CCL4*, *CCL26*, *CXCL8*, *CCL3L1*, *STAT1*, *CCL11*, *CX3CL1*, *CCL8*, *STAT2*, *XCL1*, *CCL5*, *NFKBIB*, *CXCR6*, *CCR1*, *CCL28*, *NFKBIA*, *STAT3*, *CXCL16*, *PPBP*, *JAK3*, *GNG10*, *HCK*, *FGR*, *LOC100621431*, *NFKB1*, *IKBKB*, *JAK2* |
| ssc03050 | Proteasome | 40 | 2.19 | 1.46E-06 | 1.44E-05 | *IFNG*, *PSMB9*, *PSMB8*, *PSME2*, *PSMB10*, *PSME1*, *PSMA5*, *PSMA2*, *PSMA3*, *PSMA1*, *PSMB3*, *PSMD14*, *PSMB2*, *PSMB4*, *PSMD3*, *PSMD12*, *PSMA4*, *PSMD11*, *PSMD7*, *PSMD8*, *PSMC4*, *PSMF1*, *PSMD1*, *PSMC6*, *PSME4*, *PSME3*, *PSMD2*, *PSMC1*, *PSMB1* |
| ssc05417 | Lipid and atherosclerosis | 171 | 2.18 | 8.05E-10 | 1.54E-08 | *SELE*, *CXCL2*, *CCL2*, *CXCL8*, *IRF7*, *IL1B*, *CCL3L1*, *LBP*, *IL6*, *FAS*, *TNFSF10*, *MMP1*, *SOD2*, *VCAM1*, *FASLG*, *FOS*, *CD40*, *CASP7*, *CCL5*, *TICAM2*, *CASP1*, *SELP*, *HSPA5*, *MMP9*, *LOC100624226*, *NFKBIA*, *HSP90B1*, *STAT3*, *OLR1*, *IKBKE*, *CASP3*, *HSPA6*, *CYCS*, *TRAF3*, *TRAF2*, *CD14*, *MYD88*, *DDIT3*, *HSPA8*, *VLDLR*, *TNFRSF1A*, *ICAM1*, *CASP8*, *TICAM1*, *HSPD1*, *TLR2*, *TLR4*, *NFKB1*, *IKBKB*, *JAK2*, *TANK*, *ERN1*, *TIRAP*, *EIF2S1*, *TP53*, *IRAK4* |
| ssc04940 | Type I diabetes mellitus | 25 | 2.17 | 7.58E-07 | 7.92E-06 | *IFNG*, *IL1B*, *SLA-DOA*, *FAS*, *IL1A*, *SLA-DRA*, *SLA-3*, *SLA-DQA1*, *SLA-DOB*, *SLA-DQB1*, *FASLG*, *SLA-DMB*, *SLA-7*, *SLA-DMA*, *SLA-DRB1*, *PRF1*, *SLA-8*, *HSPD1* |

| | | | | | | |
|---|---|---|---|---|---|---|
| ssc05134 | Legionellosis | 44 | 2.17 | 4.21E-06 | 3.71E-05 | *CXCL2*, *CXCL8*, *IL1B*, *IL6*, *CASP7*, *C3*, *CASP1*, *NFKB2*, *NFKBIA*, *CASP3*, *HSPA6*, *CYCS*, *CD14*, *MYD88*, *HSPA8*, *CASP8*, *HSPD1*, *TLR2*, *BNIP3*, *TLR4*, *NFKB1* |
| ssc04672 | Intestinal immune network for IgA production | 34 | 2.17 | 2.69E-06 | 2.44E-05 | *MADCAM1*, *PIGR*, *TNFSF13B*, *SLA-DOA*, *IL6*, *SLA-DRA*, *SLA-DQA1*, *SLA-DOB*, *SLA-DQB1*, *SLA-DMB*, *CD40*, *IL15*, *SLA-DMA*, *SLA-DRB1*, *CCL28*, *IL10*, *TNFSF13* |
| ssc05330 | Allograft rejection | 21 | 2.14 | 4.73E-06 | 3.98E-05 | *IFNG*, *SLA-DOA*, *FAS*, *SLA-DRA*, *SLA-3*, *SLA-DQA1*, *SLA-DOB*, *SLA-DQB1*, *FASLG*, *SLA-DMB*, *CD40*, *SLA-7*, *SLA-DMA*, *SLA-DRB1*, *IL10*, *PRF1* |
| ssc04658 | Th1 and Th2 cell differentiation | 73 | 2.09 | 5.59E-06 | 4.49E-05 | *IFNG*, *STAT1*, *SLA-DOA*, *SLA-DRA*, *SLA-DQA1*, *SLA-DOB*, *SLA-DQB1*, *IL4R*, *SLA-DMB*, *FOS*, *IL2RB*, *SLA-DMA*, *SLA-DRB1*, *STAT5A*, *NFKBIB*, *NFKBIE*, *NFKBIA*, *DLL4*, *IL2RA*, *JAK3*, *TBX21*, *NFKB1*, *IKBKB*, *JAK2*, *IL2RG* |
| ssc05320 | Autoimmune thyroid disease | 26 | 2.09 | 2.78E-05 | 1.88E-04 | *SLA-DOA*, *FAS*, *SLA-DRA*, *SLA-3*, *TG*, *SLA-DQA1*, *SLA-DOB*, *CTLA4*, *SLA-DQB1*, *FASLG*, *SLA-DMB*, *CD40*, *SLA-7*, *SLA-DMA*, *SLA-DRB1*, *IL10*, *PRF1* |
| ssc05144 | Malaria | 39 | 2.08 | 5.47E-05 | 3.56E-04 | *IFNG*, *ACKR1*, *SELE*, *CCL2*, *CXCL8*, *IL1B*, *IL6*, *CSF3*, *THBS1*, *VCAM1*, *CD40*, *SELP*, *IL10*, *MYD88*, *ICAM1*, *KLRK1*, *TLR2*, *TLR4*, *TGFB3* |
| ssc05167 | Kaposi sarcoma-associated herpesvirus infection | 156 | 2.07 | 9.33E-08 | 1.11E-06 | *CXCL2*, *CXCL8*, *IRF7*, *STAT1*, *IL6*, *FAS*, *EIF2AK2*, *SLA-3*, *CLEC2B*, *IRF9*, *CDKN1A*, *STAT2*, *FOS*, *SLA-7*, *ZFP36*, *C3*, *IFNAR2*, *CCR1*, *CSF2*, *PTGS2*, *NFKBIA*, *STAT3*, *BAK1*, *IKBKE*, *CASP3*, *CYCS*, *TRAF3*, *TRAF2*, *TNFRSF1A*, *ICAM1*, *CASP8*, *TICAM1*, *GNG10*, *HCK*, *MYC*, *SLA-8*, *LOC100621431*, *TLR3*, *NFKB1*, *IL6ST*, *IKBKB*, *JAK2*, *HIF1A* |
| ssc04210 | Apoptosis | 114 | 2.07 | 1.52E-06 | 1.45E-05 | *BIRC3*, *FAS*, *TNFSF10*, *PMAIP1*, *CTSL*, *FASLG*, *CASP10*, *FOS*, *CASP7*, *GADD45G*, *LOC100127131*, *GADD45A*, *LOC100624226*, *NFKBIA*, *BAK1*, *CASP3*, *PRF1*, *CYCS*, *CSF2RB*, *TRAF2*, *CFLAR*, *PARP3*, *DDIT3*, *BCL2A1*, *TNFRSF1A*, *CASP8*, *DAXX*, *MCL1*, *GADD45B*, *NFKB1*, *IKBKB*, *CTSO*, *TUBA4A*, *RIPK1*, *LMNB2*, *ERN1*, *EIF2S1*, *TP53* |
| ssc05142 | Chagas disease | 92 | 2.07 | 5.95E-06 | 4.66E-05 | *IFNG*, *CCL2*, *CXCL8*, *IL1B*, *CCL3L1*, *SERPINE1*, *IL6*, *FAS*, *FASLG*, *FOS*, *CCL5*, *C3*, *NFKBIA*, *IL10*, *LOC100515049*, *CFLAR*, *MYD88*, *TNFRSF1A*, *CASP8*, *CALR*, *TICAM1*, *TLR2*, *TLR4*, *TGFB3*, *NFKB1*, *IKBKB* |

| | | | | | | |
|---|---|---|---|---|---|---|
| ssc05161 | Hepatitis B | 141 | 2.07 | 4.91E-08 | 6.27E-07 | *CXCL8*, *IRF7*, *RIGI*, *STAT1*, *IL6*, *FAS*, *IFIH1*, *EGR2*, *CDKN1A*, *FASLG*, *STAT2*, *CASP10*, *FOS*, *STAT5A*, *TICAM2*, *MMP9*, *NFKBIA*, *STAT3*, *IKBKE*, *CASP3*, *CYCS*, *TRAF3*, *JAK3*, *MYD88*, *CASP8*, *TICAM1*, *MYC*, *TLR2*, *TLR4*, *CREB3L1*, *TGFB3*, *TLR3*, *NFKB1*, *IKBKB*, *JAK2*, *TIRAP*, *TP53*, *IRAK4*, *LOC100624590*, *E2F3*, *MAP2K3*, *PTK2B*, *IRAK1* |
| ssc03008 | Ribosome biogenesis in eukaryotes | 69 | 2.06 | 8.42E-06 | 6.46E-05 | *GNL3*, *POP1*, *EMG1*, *FBL*, *WDR43*, *NHP2*, *NOB1*, *GAR1*, *DKC1*, *GTPBP4*, *UTP4*, *WDR75*, *RBM28*, *XRN1*, *NOL6*, *BMS1*, *TBL3*, *IMP3*, *MPHOSPH10*, *WDR3*, *SPATA5*, *RCL1*, *NOP58*, *XPO1*, *GNL2*, *NAT10*, *UTP18*, *NOP56*, *HEATR1*, *PWP2*, *UTP15*, *CSNK2A1*, *MDN1*, *RRP7A*, *RPP40*, *UTP6*, *AK6*, *RAN*, *WDR36*, *NXT1*, *EIF6*, *NMD3*, *IMP4*, *LSG1* |
| ssc05310 | Asthma | 14 | 2.06 | 9.06E-06 | 6.79E-05 | *CCL11*, *SLA-DOA*, *SLA-DRA*, *FCER1A*, *SLA-DQA1*, *SLA-DOB*, *SLA-DQB1*, *SLA-DMB*, *CD40*, *SLA-DMA*, *SLA-DRB1*, *IL10* |
| ssc04936 | Alcoholic liver disease | 109 | 2.06 | 1.02E-06 | 1.04E-05 | *CXCL2*, *CXCL8*, *IL1B*, *LBP*, *IL6*, *FAS*, *CPT1C*, *C2*, *FASLG*, *C4A*, *C3*, *NFKBIA*, *IKBKE*, *CASP3*, *TRAF3*, *CD14*, *MYD88*, *ALDH9A1*, *TNFRSF1A*, *CASP8*, *TICAM1*, *TLR4*, *NFKB1*, *IKBKB*, *PPARGC1A*, *RIPK1*, *TIRAP*, *IRAK4*, *LY96*, *IL17RA*, *TAB3*, *ADH1C*, *MAP2K3*, *IRAK1*, *TRA2B* |
| ssc05322 | Systemic lupus erythematosus | 48 | 2.03 | 5.57E-05 | 3.56E-04 | *IFNG*, *SLA-DOA*, *SLA-DRA*, *SLA-DQA1*, *SLA-DOB*, *C2*, *SLA-DQB1*, *TRIM21*, *SLA-DMB*, *CD40*, *C4A*, *SLA-DMA*, *SLA-DRB1*, *C3*, *IL10* |
| ssc04625 | C-type lectin receptor signaling pathway | 90 | 2.01 | 5.17E-06 | 4.25E-05 | *CLEC4E*, *IRF1*, *LOC110258711*, *IL1B*, *STAT1*, *IL6*, *BCL3*, *IRF9*, *RELB*, *EGR2*, *STAT2*, *CASP1*, *NFKB2*, *PTGS2*, *NFKBIA*, *IKBKE*, *IL10*, *PLK3*, *CASP8*, *CLEC7A*, *NFKB1*, *IKBKB* |
| ssc05163 | Human cytomegalovirus infection | 190 | 2.00 | 6.86E-08 | 8.46E-07 | *TAP1*, *CCL2*, *CCL4*, *CXCL8*, *IL1B*, *CCL3L1*, *TAP2*, *IL6*, *FAS*, *CX3CL1*, *SLA-3*, *CDKN1A*, *B2M*, *TAPBP*, *FASLG*, *SLA-7*, *CCL5*, *TAPBPL*, *IL1R1*, *CCR1*, *PTGS2*, *NFKBIA*, *STAT3*, *BAK1*, *CASP3*, *PDIA3*, *CYCS*, *STING1*, *TRAF2*, *CGAS*, *EIF4EBP1*, *TNFRSF1A*, *IL10RB*, *CASP8*, *CALR*, *GNG10*, *MYC*, *SLA-8*, *CREB3L1*, *LOC100621431*, *IL6R*, *NFKB1*, *IKBKB* |
| ssc05150 | Staphylococcus aureus infection | 32 | 1.94 | 4.61E-04 | 2.57E-03 | *SLA-DOA*, *SLA-DRA*, *SLA-DQA1*, *SLA-DOB*, *C2*, *SLA-DQB1*, *CFB*, *SLA-DMB*, *C4A*, *SLA-DMA*, *SLA-DRB1*, *C3*, *FCGR2B*, *SELP*, *IL10*, *ICAM1*, *CFI*, *FCGR1A* |
| ssc04145 | Phagosome | 115 | 1.92 | 2.34E-05 | 1.62E-04 | *TAP1*, *TAP2*, *SLA-DOA*, *SLA-DRA*, *SLA-3*, *CTSL*, *SLA-DQA1*, *THBS1*, *SLA-DOB*, *SLA-DQB1*, *SLA-DMB*, *SLA-7*, *LOC100127131*, *SLA-DMA*, *SLA-DRB1*, *SFTPA1*, *C3*, *FCGR2B*, *TCIRG1*, *TFRC*, |

| | | | | | | |
|---|---|---|---|---|---|---|
| | | | | | | *OLR1*, *CD14*, *CALR*, *ATP6V1C2*, *ITGA5*, *CLEC7A*, *SLA-8*, *TLR2*, *TLR4*, *SEC61A1*, *MRC1*, *THBS2*, *FCGR1A*, *SFTPD*, *TUBA4A*, *SEC22B* |
| ssc05166 | Human T-cell leukemia virus 1 infection | 194 | 1.91 | 2.22E-06 | 2.07E-05 | *SLA-DOA*, *IL6*, *SLA-DRA*, *SLA-3*, *SLA-DQA1*, *RELB*, *FOSL1*, *SLA-DOB*, *EGR2*, *SLA-DQB1*, *CDKN1A*, *B2M*, *IL1R2*, *SLA-DMB*, *FOS*, *CD40*, *IL15*, *SLA-7*, *IL2RB*, *ZFP36*, *SLA-DMA*, *SLA-DRB1*, *STAT5A*, *NFKB2*, *IL1R1*, *CSF2*, *NFKBIA*, *EGR1*, *MSX1*, *IL2RA*, *JAK3*, *TNFRSF1A*, *ICAM1*, *CALR*, *MYC*, *SLA-8*, *CREB3L1*, *TGFB3*, *NFKB1*, *IKBKB*, *MSX2*, *IL2RG*, *TSPO*, *RANBP1*, *KAT2A*, *ETS2*, *SLC25A5*, *CANX*, *TP53*, *LTBR* |
| ssc04380 | Osteoclast differentiation | 115 | 1.90 | 3.68E-05 | 2.44E-04 | *IFNG*, *IL1B*, *STAT1*, *SOCS1*, *SOCS3*, *IL1A*, *IRF9*, *RELB*, *FOSL1*, *LOC100622460*, *JUNB*, *STAT2*, *FOS*, *FCGR2B*, *CSF1*, *OSCAR*, *NFKB2*, *IL1R1*, *IFNAR2*, *NFKBIA*, *TRAF2*, *LOC100522225*, *TNFRSF1A*, *FOSL2*, *NFKB1*, *IKBKB*, *FCGR1A* |
| ssc05235 | PD-L1 expression and PD-1 checkpoint pathway in cancer | 81 | 1.89 | 1.73E-04 | 1.04E-03 | *IFNG*, *BATF2*, *CD274*, *STAT1*, *BATF3*, *FOS*, *TICAM2*, *NFKBIB*, *NFKBIE*, *NFKBIA*, *STAT3*, *MYD88*, *TICAM1*, *TLR2*, *TLR4*, *NFKB1*, *IKBKB*, *JAK2*, *HIF1A*, *TIRAP*, *IFNGR1*, *IFNGR2*, *MAP2K3* |
| ssc04932 | Non-alcoholic fatty liver disease | 135 | 1.89 | 1.37E-05 | 1.01E-04 | *CXCL8*, *IL1B*, *IL6*, *FAS*, *SOCS3*, *IL1A*, *FASLG*, *FOS*, *CASP7*, *CASP3*, *CYCS*, *TRAF2*, *DDIT3*, *TNFRSF1A*, *CASP8*, *MLXIPL*, *IL6R*, *NFKB1*, *IKBKB*, *ERN1*, *EIF2S1*, *NDUFAB1*, *COX7A2*, *RXRA*, *XBP1*, *EIF2AK3*, *COX5A*, *BID*, *NDUFB2*, *UQCRC1*, *NR1H3*, *NDUFB3*, *RELA*, *LOC100522678*, *NDUFV2*, *MLXIP*, *COX6B*, *COX5B*, *SREBF1*, *PIK3CA*, *MAPK11*, *NDUFA9*, *IRS1* |
| ssc05416 | Viral myocarditis | 51 | 1.89 | 4.87E-04 | 2.65E-03 | *SLA-DOA*, *SLA-DRA*, *SLA-3*, *SLA-DQA1*, *SLA-DOB*, *SLA-DQB1*, *SLA-DMB*, *CD40*, *SLA-7*, *SLA-DMA*, *SLA-DRB1*, *CASP3*, *PRF1*, *CYCS*, *ICAM1*, *CASP8*, *SLA-8* |
| ssc04917 | Prolactin signaling pathway | 59 | 1.86 | 7.20E-04 | 3.60E-03 | *IRF1*, *STAT1*, *SOCS1*, *SOCS3*, *FOS*, *STAT5A*, *CISH*, *STAT3*, *GCK*, *NFKB1*, *JAK2* |
| ssc05143 | African trypanosomiasis | 28 | 1.85 | 9.56E-04 | 4.65E-03 | *IDO1*, *IFNG*, *SELE*, *IL1B*, *IL6*, *FAS*, *VCAM1*, *FASLG*, *IL10*, *MYD88*, *ICAM1* |
| ssc03060 | Protein export | 27 | 1.79 | 5.64E-03 | 2.40E-02 | *HSPA5*, *SRPRA*, *SEC61A1*, *SRPRB*, *SPCS3*, *SEC11C*, *SEC61B*, *SPCS1*, *SPCS2*, *SRP72*, *SRP14*, *SEC63*, *SRP54*, *SGTA*, *SRP68*, *SRP9* |

| | | | | | | |
|---|---|---|---|---|---|---|
| ssc04933 | AGE-RAGE signaling pathway in diabetic complications | 93 | 1.79 | 5.47E-04 | 2.86E-03 | *SELE*, *CCL2*, *CXCL8*, *IL1B*, *STAT1*, *SERPINE1*, *IL6*, *IL1A*, *VCAM1*, *PIM1*, *STAT5A*, *EGR1*, *STAT3*, *CASP3*, *ICAM1*, *TGFB3*, *NFKB1*, *JAK2* |
| ssc00970 | Aminoacyl-tRNA biosynthesis | 43 | 1.78 | 3.49E-03 | 1.54E-02 | *WARS1*, *CARS1*, *MARS1*, *GARS1*, *AARS1*, *TARS1*, *YARS1*, *FARSA*, *SARS1*, *NARS1*, *VARS1*, *MARS2*, *IARS1*, *EPRS1*, *YARS2*, *EARS2* |
| ssc00520 | Amino sugar and nucleotide sugar metabolism | 35 | 1.77 | 3.26E-03 | 1.50E-02 | *GFPT2*, *HK3*, *CHIA*, *GCK*, *GALE*, *UGP2*, *HK2*, *PGM2*, *GFPT1*, *PMM2*, *GNPNAT1*, *UAP1*, *UGDH* |
| ssc00591 | Linoleic acid metabolism | 12 | 1.76 | 9.90E-03 | 3.67E-02 | *PLA2G2D*, *CYP3A22* |
| ssc01230 | Biosynthesis of amino acids | 61 | 1.75 | 1.79E-03 | 8.46E-03 | *PSAT1*, *CBS*, *ASNS*, *SHMT1*, *SHMT2*, *MAT1A*, *GLUL*, *PYCR1*, *PHGDH*, *PSPH*, *PFKP*, *GPT2*, *ARG1*, *PGK1*, *PRPS1*, *ENO1*, *GOT1*, *ALDH18A1*, *ASS1*, *ARG2*, *IDH3A*, *BCAT1*, *TKT*, *CTH*, *ACO1*, *SDS*, *ALDOA*, *MTR*, *PAH*, *IDH2*, *PC*, *PGAM1*, *GAPDH*, *ALDOC*, *TALDO1* |
| ssc05203 | Viral carcinogenesis | 149 | 1.73 | 1.85E-04 | 1.08E-03 | *IRF7*, *PMAIP1*, *EIF2AK2*, *SLA-3*, *IRF9*, *EGR2*, *CDKN1A*, *SLA-7*, *STAT5A*, *C3*, *NFKB2*, *LOC100517129*, *NFKBIA*, *STAT3*, *BAK1*, *CASP3*, *TRAF3*, *TRAF2*, *JAK3*, *CASP8*, *SLA-8*, *CREB3L1*, *NFKB1*, *IL6ST*, *HDAC4*, *RANBP1*, *KAT2A*, *TP53*, *LTBR*, *H2BC21*, *LOC100624590*, *SND1*, *MAPKAPK2*, *H2BC15*, *TBPL1*, *CCND3*, *DNAJA3*, *JAK1* |
| ssc00380 | Tryptophan metabolism | 32 | 1.71 | 6.19E-03 | 2.51E-02 | *IDO1*, *AOC1*, *CYP1B1*, *KYNU*, *ALDH9A1*, *IL4I1*, *KMO* |
| ssc05418 | Fluid shear stress and atherosclerosis | 119 | 1.70 | 7.48E-04 | 3.69E-03 | *IFNG*, *SELE*, *CCL2*, *IL1B*, *IL1A*, *CTSL*, *VCAM1*, *IL1R2*, *FOS*, *MMP9*, *IL1R1*, *LOC100510917*, *HSP90B1*, *TNFRSF1A*, *ICAM1*, *SDC4*, *NFKB1*, *NQO1*, *IKBKB*, *DUSP1* |
| ssc01250 | Biosynthesis of nucleotide sugars | 34 | 1.70 | 7.80E-03 | 3.06E-02 | *GFPT2*, *HK3*, *GCK*, *CMAH*, *GALE*, *UGP2*, *HK2*, *PGM2*, *GFPT1*, *PMM2*, *GNPNAT1*, *UAP1*, *UGDH* |

| | | | | | | |
|---|---|---|---|---|---|---|
| ssc04640 | Hematopoietic cell lineage | 68 | 1.68 | 3.39E-03 | 1.54E-02 | *IL1B*, *SLA-DOA*, *IL6*, *IL1A*, *SLA-DRA*, *CSF3*, *SLA-DQA1*, *SLA-DOB*, *SLA-DQB1*, *IL4R*, *IL1R2*, *CSF3R*, *SLA-DMB*, *CD38*, *SLA-DMA*, *SLA-DRB1*, *CSF1*, *IL1R1*, *CSF2*, *TFRC*, *KIT*, *IL2RA*, *CD14*, *ITGA5*, *IL6R*, *ITGA6*, *FCGR1A* |
| ssc05132 | Salmonella infection | 215 | 1.68 | 1.59E-04 | 9.82E-04 | *CXCL8*, *IL1B*, *BIRC3*, *IL6*, *TNFSF10*, *LOC100522887*, *FOS*, *CASP7*, *LOC100127131*, *CASP1*, *LOC100624226*, *NFKBIA*, *HSP90B1*, *BAK1*, *MLKL*, *CASP3*, *MYL7*, *GSDMD*, *CYCS*, *RHOB*, *TRAF2*, *RHOJ*, *CD14*, *MYD88*, *TNFRSF1A*, *CASP8*, *RIPK2*, *MYC*, *TLR2*, *RIPK3*, *TLR4*, *NFKB1*, *IKBKB*, *TUBA4A*, *RIPK1*, *TIRAP*, *RHOH*, *ACTR3B*, *IRAK4*, *LY96*, *TAB3*, *MAP2K3*, *HSP90AA1*, *KPNA4*, *RHOG*, *IRAK1*, *TXN*, *ACBD3*, *EXOC5*, *ARPC5L*, *ACTR3*, *RRAS* |
| ssc05170 | Human immunodeficiency virus 1 infection | 175 | 1.66 | 4.98E-04 | 2.65E-03 | *TAP1*, *SAMHD1*, *BST2*, *TAP2*, *FAS*, *SLA-3*, *B2M*, *TAPBP*, *FASLG*, *FOS*, *SLA-7*, *TAPBPL*, *NFKBIA*, *BAK1*, *CASP3*, *PDIA3*, *CYCS*, *STING1*, *TRAF2*, *CGAS*, *MYD88*, *TNFRSF1A*, *CASP8*, *CALR*, *GNG10*, *SLA-8*, *TLR2*, *TLR4*, *LOC100621431*, *TNFRSF1B*, *NFKB1*, *IKBKB* |
| ssc04514 | Cell adhesion molecule (CAM) interaction | 114 | 1.65 | 2.65E-03 | 1.24E-02 | *MADCAM1*, *SELE*, *CD274*, *SLA-DOA*, *SLA-DRA*, *SLA-3*, *PDCD1LG2*, *SLA-DQA1*, *SLA-DOB*, *CTLA4*, *SLA-DQB1*, *VCAM1*, *SLA-DMB*, *CD40*, *SLA-7*, *PVR*, *SLA-DMA*, *SLA-DRB1*, *SELP*, *LRRC4* |
| ssc04920 | Adipocytokine signaling pathway | 59 | 1.65 | 1.06E-02 | 3.91E-02 | *SOCS3*, *CPT1C*, *PCK2*, *NFKBIB*, *ACSL1*, *NFKBIE*, *NFKBIA*, *STAT3*, *SLC2A4*, *ACSL5*, *TRAF2*, *TNFRSF1A*, *TNFRSF1B*, *NFKB1*, *IKBKB*, *JAK2*, *ACSL4*, *PPARGC1A* |
| ssc04115 | p53 signaling pathway | 65 | 1.63 | 7.64E-03 | 3.03E-02 | *SERPINE1*, *FAS*, *PMAIP1*, *THBS1*, *CDKN1A*, *SHISA5*, *GADD45G*, *GADD45A*, *LOC100624226*, *SESN2*, *CASP3*, *CYCS*, *CASP8*, *GADD45B*, *ZMAT3*, *TP53* |
| ssc04216 | Ferroptosis | 37 | 1.62 | 1.11E-02 | 3.96E-02 | *SLC39A14*, *SLC7A11*, *CP*, *ACSL1*, *TFRC*, *SLC39A8*, *FTH1*, *SLC3A2*, *ACSL5*, *GCLM*, *SLC11A2*, *ACSL4*, *SAT1*, *TP53*, *LPCAT3*, *STEAP3*, *GP91-PHOX*, *HMOX1* |
| ssc01524 | Platinum drug resistance | 58 | 1.59 | 1.25E-02 | 4.32E-02 | *BIRC3*, *FAS*, *PMAIP1*, *CDKN1A*, *FASLG*, *LOC100510917*, *BAK1*, *CASP3*, *CYCS*, *CASP8*, *SLC31A1* |
| ssc04650 | Natural killer cell mediated cytotoxicity | 79 | 1.58 | 6.13E-03 | 2.51E-02 | *IFNG*, *FAS*, *TNFSF10*, *FASLG*, *IFNAR2*, *LOC100624226*, *CSF2*, *SH3BP2*, *CASP3*, *PRF1*, *LOC100523789*, *CD48*, *ICAM1*, *ULBP1*, *KLRK1* |

| | | | | | | |
|---|---|---|---|---|---|---|
| ssc04068 | FoxO signaling pathway | 116 | 1.58 | 4.53E-03 | 1.98E-02 | *IL6*, *TNFSF10*, *SOD2*, *CDKN1A*, *PCK2*, *FASLG*, *GADD45G*, *GADD45A*, *S1PR1*, *STAT3*, *SLC2A4*, *DEPP1*, *IL10*, *PLK3*, *BCL6*, *BNIP3*, *TGFB3*, *GADD45B*, *IKBKB* |
| ssc04066 | HIF-1 signaling pathway | 96 | 1.58 | 6.13E-03 | 2.51E-02 | *IFNG*, *SERPINE1*, *IL6*, *HK3*, *CDKN1A*, *TIMP1*, *STAT3*, *TFRC*, *PFKP*, *EGLN3*, *EIF4EBP1*, *HK2*, *TLR4*, *PGK1*, *IL6R*, *NFKB1*, *HIF1A*, *ENO1*, *IFNGR1*, *LTBR*, *IFNGR2*, *EIF4E* |
| ssc05202 | Transcriptional misregulation in cancer | 145 | 1.50 | 8.21E-03 | 3.17E-02 | *ETV7*, *CXCL8*, *BIRC3*, *IL6*, *NUPR1*, *CDKN1A*, *NFKBIZ*, *IL1R2*, *CD40*, *IL2RB*, *GADD45G*, *PML*, *GADD45A*, *MMP9*, *CSF2*, *NR4A3*, *BAK1*, *MAX*, *BCL6*, *CEBPB*, *CD14*, *DDIT3*, *BCL2A1*, *MYC*, *ZBTB16*, *GADD45B*, *NFKB1*, *WNT16*, *ETV6*, *SMAD1*, *FCGR1A* |
| ssc05012 | Parkinson disease | 220 | 1.40 | 1.10E-02 | 3.94E-02 | *UBA7*, *UBE2L6*, *SLC39A14*, *PPIF*, *LOC100127131*, *HSPA5*, *PSMA5*, *SLC39A8*, *PSMA2*, *CASP3*, *PSMA3*, *GPR37*, *CYCS*, *DDIT3*, *DAXX*, *PSMA1*, *PSMB3*, *SLC11A2*, *PSMD14*, *PSMB2*, *PSMB4*, *DUSP1*, *SLC39A7*, *TUBA4A*, *PSMD3*, *ERN1*, *EIF2S1*, *SLC39A6*, *PSMD12*, *SLC25A5*, *PSMA4*, *TP53*, *SLC39A11*, *NDUFAB1*, *TXN*, *PSMD11*, *SLC39A1*, *PSMD7*, *PSMD8*, *PSMC4*, *COX7A2*, *ATP5MC1*, *XBP1*, *PSMD1*, *PSMC6*, *EIF2AK3*, *COX5A*, *NDUFB2*, *UQCRC1*, *ATF6*, *PSMD2*, *PSMC1*, *PSMB1*, *NDUFB3*, *LOC100522678*, *PSMC2*, *NDUFV2*, *MAOA*, *SNCAIP*, *COX6B*, *COX5B*, *ATP5F1B*, *PSMD13*, *PSMD4*, *ATP5F1C*, *ITPR1*, *NDUFA9*, *PSMD6*, *SLC18A1*, *KIF5B*, *UBE2J2*, *SDHD*, *SLC39A10*, *ATP5F1A*, *UBE2L3*, *UQCRQ*, *SLC18A2*, *MAPK10*, *NFE2L2* |
| ssc05200 | Pathways in cancer | 441 | 1.34 | 1.20E-02 | 4.21E-02 | *IFNG*, *CXCL8*, *STAT1*, *BIRC3*, *IL6*, *FAS*, *PMAIP1*, *MMP1*, *CDKN1A*, *PIM1*, *IL4R*, *CSF3R*, *FASLG*, *STAT2*, *FOS*, *CASP7*, *IL15*, *IL2RB*, *GADD45G*, *PML*, *STAT5A*, *GADD45A*, *FZD9*, *MMP9*, *NFKB2*, *IFNAR2*, *LOC100510917*, *PTGS2*, *NFKBIA*, *HSP90B1*, *STAT3*, *BAK1*, *F2R*, *KIT*, *CASP3*, *RALGDS*, *CYCS*, *MAX*, *CSF2RB*, *DLL4*, *TRAF3*, *TRAF2*, *RASSF1*, *IL2RA*, *JAK3*, *EGLN3*, *CASP8*, *GNG10*, *MYC*, *NKX3-1*, *LOC100621431*, *TGFB3*, *ZBTB16*, *LPAR3*, *IL6R*, *TXNRD1*, *GADD45B*, *NFKB1*, *NQO1*, *IL6ST*, *IKBKB*, *ITGA6*, *JAK2*, *WNT16*, *IL2RG*, *HIF1A*, *WNT8B*, *PGF*, *PDGFRB*, *GNA13*, *GLI2*, *LAMB4* |
| ssc04518 | Integrin signaling | 140 | -1.40 | 1.08E-02 | 3.92E-02 | *LAMC2*, *PDGFD*, *FREM2*, *CHAD*, *COL4A6*, *LIMS2*, *ITGA8*, *LIMS1*, *PARVG*, *VCL*, *EGF*, *ITGB4*, *FERMT1*, *ITGA3*, *FREM1*, *ITGAM*, *COL1A1*, *LAMA3*, *FLT4*, *COL4A4*, *COL1A2*, *LAMB2*, *RAC3*, *FN1*, *ITGB7*, *NPNT*, *COL4A3*, *RELN*, *TLN2*, *ITGA4*, *THBS3*, *LAMA2*, *IGF1*, *COMP*, *LAMA1*, *TNXB*, *VTN*, *COL6A6*, *COL9A3*, *VEGFD* |

| | | | | | | |
|---|---|---|---|---|---|---|
| ssc04024 | cAMP signaling pathway | 166 | -1.42 | 8.28E-03 | 3.17E-02 | *HTR1D*, *NPY1R*, *MYL9*, *GRIN3A*, *GABBR2*, *ATP2A3*, *HTR4*, *RAPGEF4*, *ATP2B2*, *SOX9*, *GRIN2B*, *CAMK2G*, *ADCY1*, *CNGA2*, *NFATC1*, *PLD1*, *TSHR*, *ATP1B2*, *LHB*, *ATP2B4*, *CREB5*, *RAC3*, *GRIA1*, *CACNA1S*, *BDNF*, *FXYD2*, *GRIN3B*, *ADRB1*, *CHRM2*, *GRIA4*, *HHIP*, *CNGA1*, *VIPR2*, *TNNI3*, *CALML4*, *EDN3* |
| ssc04022 | cGMP-PKG signaling pathway | 141 | -1.43 | 9.37E-03 | 3.51E-02 | *ADCY7*, *ADRA1B*, *CALM3*, *PDE3B*, *MEF2B*, *NPR1*, *ADCY8*, *MYLK4*, *SLC8A2*, *ADRB3*, *PDE5A*, *PRKG2*, *MYL9*, *GUCY1A2*, *PIK3R6*, *ATP2A3*, *ADRA1D*, *ATP2B2*, *LOC100736765*, *ADCY1*, *NFATC1*, *ATP1B2*, *ATP2B4*, *KCNMB4*, *CREB5*, *PIK3CG*, *NFATC2*, *PDE2A*, *PLCB2*, *CACNA1S*, *FXYD2*, *ADRB1*, *CNGA1*, *ADRA1A*, *CALML4*, *ADRA2C* |
| ssc04015 | Rap1 signaling pathway | 179 | -1.46 | 4.68E-03 | 2.02E-02 | *VEGFA*, *FGF21*, *ADCY7*, *CALM3*, *TIAM1*, *MAP2K6*, *EVL*, *PARD6A*, *ADCY8*, *KDR*, *RASSF5*, *MAGI3*, *EFNA5*, *PARD6G*, *PDGFD*, *RAPGEF4*, *TEK*, *PRKCZ*, *GRIN2B*, *LPAR2*, *MAGI2*, *EGF*, *SKAP1*, *ADCY1*, *RASGRP2*, *DOCK4*, *FGF18*, *EFNA2*, *ITGAM*, *PRKCG*, *FLT4*, *RAC3*, *FARP2*, *CSF1R*, *RGS14*, *KITLG*, *PRKCB*, *TLN2*, *CNR1*, *PLCB2*, *IGF1*, *ANGPT1*, *FGF1*, *NGFR*, *VEGFD*, *CALML4* |
| ssc04082 | Neuroactive ligand signaling | 116 | -1.47 | 7.31E-03 | 2.93E-02 | *GRM8*, *GABRG1*, *GABBR2*, *HTR4*, *ADRA1D*, *GRIN2B*, *P2RX5*, *SLC6A13*, *ADCY1*, *GRIK4*, *SLC1A1*, *PRKCG*, *GNG2*, *GABRR3*, *P2RX6*, *GABRA1*, *GNG4*, *PRKCB*, *PLCB2*, *GRIA1*, *GRIK2*, *GRIN3B*, *ADRB1*, *OPRL1*, *CHRM2*, *GRIA4*, *P2RY12*, *ADRA1A*, *P2RX3*, *ADRA2C*, *SLC6A4* |
| ssc04814 | Motor proteins | 141 | -1.51 | 3.48E-03 | 1.54E-02 | *DNAH3*, *ACTB*, *DNAH6*, *DNAH7*, *MYH7B*, *DCTN1*, *DYNLT1*, *DYNC2H1*, *KIFAP3*, *TUBB4B*, *DYNLT3*, *DNAI3*, *MYO9A*, *KIF23*, *KIF20B*, *MYH11*, *MYO1F*, *BICDL2*, *DNAH11*, *DYNC2LI1*, *KIF7*, *TUBAL3*, *MYO1C*, *DNAH1*, *KIF15*, *KIF2C*, *DNAH9*, *TPM1*, *DYNLRB2*, *DNAH10*, *KIF1C*, *DYNLT5*, *DYNLT4*, *ACTC1*, *TUBB2B*, *KIF18B*, *KIF13A*, *DNAI1*, *MYO7A*, *KIF11*, *MYO5C*, *MYL9*, *KIF24*, *MYL6*, *DNAI2*, *CENPE*, *KLC4*, *KIF3C*, *KIFC1*, *MYH14*, *KIF20A*, *TUBB4A*, *LOC100736765*, *MYO5B*, *BICDL1*, *KIF14*, *KIF22*, *KIFC2*, *MYO15B*, *TNNI1*, *KLC3*, *TNNI3*, *MYH2* |
| ssc05410 | Hypertrophic cardiomyopathy | 78 | -1.53 | 1.22E-02 | 4.23E-02 | *ACTC1*, *DMD*, *SLC8A2*, *SGCE*, *CACNA2D2*, *ITGA8*, *CACNB3*, *ATP2A3*, *ITGB4*, *LOC100736765*, *ITGA3*, *ITGB7*, *CACNB4*, *SGCA*, *ITGA4*, *CACNG6*, *LAMA2*, *CACNA1S*, *IGF1*, *ACE*, *LAMA1*, *TGFB2*, *TNNI3* |

| | | | | | | |
|---|---|---|---|---|---|---|
| ssc04310 | Wnt signaling pathway | 151 | -1.54 | 1.45E-03 | 6.94E-03 | *FZD2*, *WNT3*, *AXIN2*, *WNT7A*, *CAMK2G*, *WNT11*, *WNT9A*, *LOC100624648*, *NFATC1*, *TLE4*, *PRKCG*, *WNT10B*, *WNT3A*, *NOTUM*, *RAC3*, *TCF7*, *WNT2*, *APCDD1*, *CTNNBIP1*, *WNT5B*, *FZD8*, *PRKCB*, *NFATC2*, *SFRP5*, *CTNND2*, *PRICKLE1*, *PLCB2*, *WIF1*, *WNT4*, *SFRP4*, *DKK2*, *SFRP2* |
| ssc04916 | Melanogenesis | 89 | -1.55 | 5.96E-03 | 2.51E-02 | *FZD2*, *WNT3*, *WNT7A*, *CAMK2G*, *WNT11*, *WNT9A*, *ADCY1*, *PRKCG*, *WNT10B*, *WNT3A*, *TCF7*, *DCT*, *WNT2*, *WNT5B*, *KITLG*, *FZD8*, *PRKCB*, *PLCB2*, *WNT4*, *CALML4* |
| ssc04970 | Salivary secretion | 60 | -1.57 | 1.29E-02 | 4.40E-02 | *ADCY7*, *ADRA1B*, *CALM3*, *ADCY8*, *ADRB3*, *PRKG2*, *GUCY1A2*, *ADRA1D*, *ATP2B2*, *ADCY1*, *RYR3*, *PRKCG*, *ATP1B2*, *ATP2B4*, *PRKCB*, *CST3*, *PLCB2*, *FXYD2*, *ADRB1*, *BST1*, *ADRA1A*, *CALML4* |
| ssc04510 | Focal adhesion | 179 | -1.58 | 6.14E-04 | 3.16E-03 | *LAMC1*, *PDGFA*, *PXN*, *VEGFA*, *DIAPH1*, *COL6A5*, *FLNA*, *SHC3*, *KDR*, *MYLK4*, *LAMC2*, *PDGFD*, *MYL9*, *CHAD*, *COL4A6*, *ITGA8*, *PARVG*, *VCL*, *EGF*, *ITGB4*, *PPP1R12B*, *ITGA3*, *RASGRF1*, *PRKCG*, *COL1A1*, *LAMA3*, *FLT4*, *COL4A4*, *COL1A2*, *LAMB2*, *RAC3*, *FN1*, *CAV2*, *ITGB7*, *COL4A3*, *PRKCB*, *RELN*, *TLN2*, *ITGA4*, *THBS3*, *PAK6*, *LAMA2*, *IGF1*, *COMP*, *LAMA1*, *CAV1*, *TNXB*, *VTN*, *COL6A6*, *COL9A3*, *VEGFD* |
| ssc05412 | Arrhythmogenic right ventricular cardiomyopathy | 68 | -1.63 | 8.80E-03 | 3.34E-02 | *SLC8A3*, *CACNA1C*, *ITGAV*, *DAG1*, *ACTB*, *CTNNB1*, *CACNB2*, *CTNNA1*, *PKP2*, *ITGB5*, *ITGA11*, *CACNA1D*, *SNTB2*, *JUP*, *DSP*, *TCF7L1*, *ACTN2*, *DMD*, *SLC8A2*, *SGCE*, *DSC2*, *CACNA2D2*, *ITGA8*, *CACNB3*, *ATP2A3*, *ITGB4*, *DSG2*, *ITGA3*, *TCF7*, *ITGB7*, *CTNNA3*, *CACNB4*, *SGCA*, *ITGA4*, *CACNG6*, *LAMA2*, *CACNA1S*, *LAMA1* |
| ssc04261 | Adrenergic signaling in cardiomyocytes | 122 | -1.65 | 4.93E-04 | 2.65E-03 | *TPM1*, *ADCY7*, *ADRA1B*, *CALM3*, *ACTC1*, *KCNQ1*, *ADCY8*, *SLC8A2*, *RPS6KA5*, *CACNA2D2*, *CACNB3*, *PIK3R6*, *ATP2A3*, *RAPGEF4*, *ADRA1D*, *ATP2B2*, *KCNE1*, *CAMK2G*, *LOC100736765*, *ADCY1*, *ATP1B2*, *ATP2B4*, *CREB5*, *PIK3CG*, *CACNB4*, *PPP2R2C*, *CACNG6*, *PLCB2*, *CACNA1S*, *FXYD2*, *ADRB1*, *AGTR2*, *ADRA1A*, *TNNI3*, *CALML4* |
| ssc04360 | Axon guidance | 167 | -1.66 | 2.25E-04 | 1.30E-03 | *SEMA5B*, *PLXNB1*, *EPHB3*, *CDK5*, *SLIT2*, *SEMA6D*, *PARD6A*, *NGEF*, *ROBO2*, *CXCR4*, *RHOD*, *NTNG2*, *EFNA5*, *PARD6G*, *EPHA4*, *MYL9*, *EPHB2*, *BMPR2*, *BOC*, *SRGAP1*, *ROBO1*, *PRKCZ*, *CAMK2G*, *SSH1*, *NEO1*, *BMP7*, *SEMA4G*, *SEMA3C*, *SEMA6A*, *EFNA2*, *PLXNB3*, *UNC5C*, *SEMA3E*, *SEMA6C*, *RAC3*, *EPHA7*, *PLXNA3*, *SEMA3B*, *WNT5B*, *EPHB1*, *SHH*, *NFATC2*, *PAK6*, *EFNB3*, *SLIT3*, *SRGAP3*, *SEMA3D*, *NTNG1*, *WNT4*, *EPHA3*, *LOC100620540*, *NTN3* |

| | | | | | | |
|---|---|---|---|---|---|---|
| ssc04080 | Neuroactive ligand-receptor interaction | 149 | -1.67 | 1.56E-04 | 9.80E-04 | *ADRB3*, *HTR1D*, *NPY1R*, *PTH1R*, *GRIN3A*, *PTGER1*, *GRM8*, *GABRG1*, *GABBR2*, *HTR4*, *S1PR5*, *ADRA1D*, *GRIN2B*, *LPAR2*, *P2RX5*, *PAQR6*, *P2RY10*, *GCGR*, *TRHR*, *GRIK4*, *F2RL1*, *TSHR*, *LEPR*, *GALR1*, *LHB*, *NPFF*, *CALCB*, *GABRR3*, *P2RX6*, *GABRA1*, *P2RX7*, *GRID2*, *CNR1*, *P2RY8*, *GRIA1*, *PYY*, *GRIK2*, *GRIN3B*, *LYPD6*, *APLN*, *ADRB1*, *OPRL1*, *CHRM2*, *GRIA4*, *AGTR2*, *GPR156*, *VIPR2*, *ADRA1A*, *QRFPR*, *EDN3*, *P2RX3*, *APLNR*, *ADRA2C* |
| ssc04081 | Hormone signaling | 128 | -1.67 | 1.75E-04 | 1.04E-03 | *ADCY7*, *ADRA1B*, *GNG8*, *HEPH*, *SMAD9*, *GNA14*, *SLC16A10*, *NPR1*, *ADCY8*, *VIPR1*, *ADRB3*, *PTH1R*, *PRKG2*, *RAPGEF4*, *ADRA1D*, *IGF2*, *ESR2*, *ADCY1*, *SLC40A1*, *GCGR*, *TRHR*, *EPOR*, *TSHR*, *LEPR*, *LHB*, *GNG2*, *CALCB*, *SLC16A2*, *ESR1*, *GNG4*, *PLCB2*, *IGF1*, *RAMP3*, *ADRB1*, *OPRL1*, *AGTR2*, *VIPR2*, *ADRA1A*, *ADRA2C* |
| ssc04820 | Cytoskeleton in muscle cells | 178 | -1.74 | 1.68E-05 | 1.21E-04 | *FBN1*, *DIAPH1*, *TPM1*, *FLNC*, *AGRN*, *COL6A5*, *ACTC1*, *SYNE3*, *DMD*, *PDLIM2*, *COL11A1*, *FBLN1*, *SGCE*, *DSC2*, *PDLIM4*, *MYL9*, *SDC2*, *COL4A6*, *SYNPO2*, *ITGA8*, *ENO3*, *VCL*, *SPTBN1*, *FHL2*, *BGN*, *MYH14*, *SDC1*, *ITGB4*, *LOC100736765*, *CSRP1*, *DSG2*, *ITGA3*, *COL1A1*, *COL4A4*, *VIM*, *COL3A1*, *ATP1B2*, *COL1A2*, *MYOM3*, *FN1*, *DCN*, *FBN3*, *ITGB7*, *COL4A3*, *TNNI1*, *SGCA*, *TLN2*, *TMOD2*, *ITGA4*, *SPTBN4*, *THBS3*, *LAMA2*, *SDC3*, *COMP*, *LAMA1*, *FHL1*, *FBLN2*, *PDLIM3*, *OBSCN*, *COL6A6*, *COL9A3*, *TNNI3*, *MYH2*, *TRIM55* |
| ssc04512 | ECM-receptor interaction | 78 | -1.74 | 6.98E-04 | 3.54E-03 | *LAMC2*, *FREM2*, *CHAD*, *COL4A6*, *ITGA8*, *SDC1*, *ITGB4*, *ITGA3*, *FREM1*, *COL1A1*, *LAMA3*, *COL4A4*, *COL1A2*, *LAMB2*, *FN1*, *ITGB7*, *NPNT*, *COL4A3*, *GP9*, *RELN*, *ITGA4*, *THBS3*, *LAMA2*, *COMP*, *LAMA1*, *TNXB*, *VTN*, *COL6A6*, *COL9A3*, *LOC100511343* |
| ssc04020 | Calcium signaling pathway | 186 | -1.78 | 4.30E-06 | 3.71E-05 | *ERBB3*, *CACNA1D*, *MCOLN1*, *PLCE1*, *HRC*, *ERBB2*, *CALM1*, *GRIN2D*, *PTGFR*, *TACR3*, *CAMK4*, *CYSLTR1*, *PDGFA*, *VEGFA*, *FGF21*, *ADCY7*, *ADRA1B*, *CALM3*, *ERBB4*, *GNA14*, *PHKA2*, *ADCY8*, *CXCR4*, *KDR*, *MYLK4*, *SLC8A2*, *CACNA1G*, *ADRB3*, *PHKG1*, *PLCD3*, *PDGFD*, *MST1*, *ITPKB*, *GRIN3A*, *PTGER1*, *ATP2A3*, *HTR4*, *ADRA1D*, *ATP2B2*, *GRIN2B*, *GDNF*, *PLCD1*, *P2RX5*, *EGF*, *CAMK2G*, *ADCY1*, *FGF18*, *TRHR*, *NFATC1*, *RYR3*, *PRKCG*, *FLT4*, *ATP2B4*, *MCOLN2*, *P2RX6*, *P2RX7*, *PRKCB*, *NFATC2*, *NTRK2*, *PLCB2*, *CACNA1S*, *CASQ1*, *CACNA1E*, *FGF1*, *GRIN3B*, *ADRB1*, *CHRM2*, *ADRA1A*, *VEGFD*, *CALML4*, *P2RX3*, *CACNA1B* |

| | | | | | | |
|---|---|---|---|---|---|---|
| ssc05414 | Dilated cardiomyopathy | 81 | -1.79 | 4.15E-04 | 2.34E-03 | *TPM1*, *ADCY7*, *ACTC1*, *DMD*, *ADCY8*, *SLC8A2*, *SGCE*, *CACNA2D2*, *ITGA8*, *CACNB3*, *ATP2A3*, *ITGB4*, *LOC100736765*, *ADCY1*, *ITGA3*, *ITGB7*, *CACNB4*, *SGCA*, *ITGA4*, *CACNG6*, *LAMA2*, *CACNA1S*, *IGF1*, *LAMA1*, *TGFB2*, *ADRB1*, *TNNI3* |
| ssc04974 | Protein digestion and absorption | 70 | -2.01 | 1.84E-05 | 1.29E-04 | *COL16A1*, *COL8A1*, *COL28A1*, *PRCP*, *COL6A5*, *KCNQ1*, *SLC16A10*, *COL11A1*, *SLC8A2*, *COL4A6*, *MME*, *COL23A1*, *DPP4*, *CTRB2*, *SLC1A1*, *COL14A1*, *COL1A1*, *COL4A4*, *COL3A1*, *ATP1B2*, *COL1A2*, *COL26A1*, *COL4A3*, *SLC7A9*, *COL25A1*, *CPA1*, *FXYD2*, *COL19A1*, *COL21A1*, *COL8A2*, *COL6A6*, *COL9A3*, *SLC7A8*, *COL15A1*, *KCNJ13* |

Abbreviations: AGE-RAGE: Advanced glycation end products-receptor for AGEs; cAMP: Cyclic adenosine monophosphate; cGMP-PKG: Cyclic guanosine monophosphate-protein kinase G; ECM: Extracellular matrix; FDR: False discovery rate; FoxO: Forkhead box O; HIF-1: Hypoxia-inducible factor 1; IgA: Immunoglobulin A; IL-17: Interleukin-17; JAK-STAT: Janus kinases-signal transducer and activator of transcription proteins; KEGG: Kyoto Encyclopedia of Genes and Genomes; NES: Normalized enrichment score; NF-kappa B: Nuclear factor-kappa B; NOD: Nucleotide-binding domain; PD-1: Programmed cell death protein 1; PD-L1: Programmed cell death ligand 1; Rap1: Ras-associated protein-1; Th1/Th2/Th17: T helper 1/2/17 cells; TNF: Tumor necrosis factor.

**Supplementary Table S2. Significantly enriched pathways from the GSEA result of the GO:BP database, Infection versus Control.**

Abbreviations: GO:BP: Biological Processes in Gene Ontology; GSEA: Gene set enrichment analysis.

| GO:BP ID | Pathways | Set size | NES | P-value | FDR | Core enrichment gene |
|---|---|---|---|---|---|---|
| GO:0019221 | Cytokine-mediated signaling pathway | 129 | 2.70 | 1.00E-10 | 2.09E-09 | *CXCL9*, *CXCL11*, *CXCL10*, *IFNG*, *ACKR1*, *ISG15*, *CXCL13*, *AMCF-II*, *IRF1*, *OAS1*, *CCL2*, *CCL4*, *CXCL8*, *IRF7*, *ZBP1*, *IL1B*, *STAT1*, *MMP8*, *TNFSF13B*, *IL6*, *SOCS1*, *IL1A*, *FOSL1*, *TIFA*, *CCL8*, *IL4R*, *STAT2*, *CD40*, *IL15*, *CCL5*, *STAT5A*, *TICAM2*, *CISH*, *NFKBIA*, *EGR1*, *STAT3*, *IL10*, *KIT*, *PPBP*, *TRAF3*, *TRAF2*, *IL18RAP*, *JAK3*, *ADAM17*, *TNFRSF1A* |
| GO:0034097 | Response to cytokine | 214 | 2.69 | 1.00E-10 | 2.09E-09 | *CXCL9*, *CXCL11*, *CXCL10*, *UBD*, *IFNG*, *ACKR1*, *SELE*, *ISG15*, *MX1*, *CXCL13*, *AMCF-II*, *IRF1*, *OAS1*, *CHI3L1*, *CCL2*, *CCL4*, *CXCL8*, *IRF7*, *ZBP1*, *IL1B*, *STAT1*, *MMP8*, *TNFSF13B*, *IL6*, *SOCS1*, *IL1A*, *GFPT2*, *SNX10*, *FOSL1*, *TIFA*, *CCL8*, *SLA-DQB1*, *PIM1*, *IL4R*, *IRF8*, *MSC*, *STAT2*, *CD40*, *IL15*, *CCL5*, *ZFP36*, *TIMP1*, *NFIL3*, *STAT5A*, *TICAM2*, *CISH*, *NFKBIA*, *EGR1*, *STAT3*, *CXCL16*, *IL10*, *KIT*, *PPBP*, *TRAF3*, *TRAF2*, *IL18RAP*, *JAK3*, *ADAM17*, *TNFRSF1A*, *CASP8*, *YBX3*, *MCL1*, *ITIH4*, *TLR2*, *TLR4*, *NKX3-1*, *CACYBP*, *IL6R*, *IKBKB*, *STIP1*, *JAK2* |
| GO:1901652 | Response to peptide | 214 | 2.69 | 1.00E-10 | 2.09E-09 | *CXCL9*, *CXCL11*, *CXCL10*, *UBD*, *IFNG*, *ACKR1*, *SELE*, *ISG15*, *MX1*, *CXCL13*, *AMCF-II*, *IRF1*, *OAS1*, *CHI3L1*, *CCL2*, *CCL4*, *CXCL8*, *IRF7*, *ZBP1*, *IL1B*, *STAT1*, *MMP8*, *TNFSF13B*, *IL6*, *SOCS1*, *IL1A*, *GFPT2*, *SNX10*, *FOSL1*, *TIFA*, *CCL8*, *SLA-DQB1*, *PIM1*, *IL4R*, *IRF8*, *MSC*, *STAT2*, *CD40*, *IL15*, *CCL5*, *ZFP36*, *TIMP1*, *NFIL3*, *STAT5A*, *TICAM2*, *CISH*, *NFKBIA*, *EGR1*, *STAT3*, *CXCL16*, *IL10*, *KIT*, *PPBP*, *TRAF3*, *TRAF2*, *IL18RAP*, *JAK3*, *ADAM17*, *TNFRSF1A*, *CASP8*, *YBX3*, *MCL1*, *ITIH4*, *TLR2*, *TLR4*, *NKX3-1*, *CACYBP*, *IL6R*, *IKBKB*, *STIP1*, *JAK2* |
| GO:0043207 | Response to external biotic stimulus | 365 | 2.64 | 1.00E-10 | 2.09E-09 | *CXCL9*, *CXCL10*, *ISG20*, *GBP1*, *IFNG*, *BATF2*, *RSAD2*, *ISG15*, *MX1*, *GBP2*, *CXCL13*, *MX2*, *AMCF-II*, *IRF1*, *IFIT1*, *OAS1*, *CCL2*, *CCL4*, *DHX58*, *CXCL8*, *OAS2*, *IRF7*, *ZBP1*, *RIGI*, *IL1B*, *S100A12*, *S100A8*, *IL27*, *STAT1*, *MMP8*, *SAMHD1*, *BST2*, *LBP*, *CALHM6*, *IL1A*, *IFIH1*, *IFIT2*, *PTGIR*, *BPIFB1*, *RNF19B*, *FOSL1*, *TNFAIP3*, *TIFA*, *CCL8*, *NMI*, *C2*, *SLC15A3*, *SLA-DQB1*, *PIM1*, *IRF8*, *CD180*, *IFI35*, *CFB*, *STAT2*, *CD40*, *C4A*, *LTF*, *CCL5*, *TRIM14*, *FFAR2*, *ZFP36*, |

| | | | | | | |
|---|---|---|---|---|---|---|
| | | | | | | *NOD2*, *IFIT5*, *TICAM2*, *C3*, *GSDMA*, *NFKBIB*, *UNC93B1*, *HP*, *DUSP10*, *NFKBIA*, *SERPING1*, *BPIFA1*, *TREM1*, *TRIM26*, *MLKL*, *CXCL16*, *IL10*, *GNLY*, *MR1*, *PPBP*, *CEBPG*, *OPTN*, *STING1*, *TRAF3*, *TRAF2*, *HAVCR2*, *DDX21*, *NDUFAF4*, *CEBPB*, *IL18RAP*, *MSRB1*, *MORC3*, *CD14*, *CGAS*, *MYD88*, *SELENOS*, *TICAM1*, *KLRK1*, *PLSCR1*, *POLR3D*, *SLAMF6*, *CLEC7A*, *TLR2*, *NKG7*, *RIPK3*, *TLR4*, *C9*, *RNF115* |
| GO:0051707 | Response to other organism | 365 | 2.64 | 1.00E-10 | 2.09E-09 | *CXCL9*, *CXCL10*, *ISG20*, *GBP1*, *IFNG*, *BATF2*, *RSAD2*, *ISG15*, *MX1*, *GBP2*, *CXCL13*, *MX2*, *AMCF-II*, *IRF1*, *IFIT1*, *OAS1*, *CCL2*, *CCL4*, *DHX58*, *CXCL8*, *OAS2*, *IRF7*, *ZBP1*, *RIGI*, *IL1B*, *S100A12*, *S100A8*, *IL27*, *STAT1*, *MMP8*, *SAMHD1*, *BST2*, *LBP*, *CALHM6*, *IL1A*, *IFIH1*, *IFIT2*, *PTGIR*, *BPIFB1*, *RNF19B*, *FOSL1*, *TNFAIP3*, *TIFA*, *CCL8*, *NMI*, *C2*, *SLC15A3*, *SLA-DQB1*, *PIM1*, *IRF8*, *CD180*, *IFI35*, *CFB*, *STAT2*, *CD40*, *C4A*, *LTF*, *CCL5*, *TRIM14*, *FFAR2*, *ZFP36*, *NOD2*, *IFIT5*, *TICAM2*, *C3*, *GSDMA*, *NFKBIB*, *UNC93B1*, *HP*, *DUSP10*, *NFKBIA*, *SERPING1*, *BPIFA1*, *TREM1*, *TRIM26*, *MLKL*, *CXCL16*, *IL10*, *GNLY*, *MR1*, *PPBP*, *CEBPG*, *OPTN*, *STING1*, *TRAF3*, *TRAF2*, *HAVCR2*, *DDX21*, *NDUFAF4*, *CEBPB*, *IL18RAP*, *MSRB1*, *MORC3*, *CD14*, *CGAS*, *MYD88*, *SELENOS*, *TICAM1*, *KLRK1*, *PLSCR1*, *POLR3D*, *SLAMF6*, *CLEC7A*, *TLR2*, *NKG7*, *RIPK3*, *TLR4*, *C9*, *RNF115* |
| GO:0071345 | Cellular response to cytokine stimulus | 199 | 2.64 | 1.00E-10 | 2.09E-09 | *CXCL9*, *CXCL11*, *CXCL10*, *IFNG*, *ACKR1*, *ISG15*, *MX1*, *CXCL13*, *AMCF-II*, *IRF1*, *OAS1*, *CHI3L1*, *CCL2*, *CCL4*, *CXCL8*, *IRF7*, *ZBP1*, *IL1B*, *STAT1*, *MMP8*, *TNFSF13B*, *IL6*, *SOCS1*, *IL1A*, *GFPT2*, *SNX10*, *FOSL1*, *TIFA*, *CCL8*, *SLA-DQB1*, *PIM1*, *IL4R*, *IRF8*, *MSC*, *STAT2*, *CD40*, *IL15*, *CCL5*, *ZFP36*, *NFIL3*, *STAT5A*, *TICAM2*, *CISH*, *NFKBIA*, *EGR1*, *STAT3*, *IL10*, *KIT*, *PPBP*, *TRAF3*, *TRAF2*, *IL18RAP*, *JAK3*, *ADAM17*, *TNFRSF1A*, *YBX3*, *TLR2*, *TLR4*, *NKX3-1*, *CACYBP*, *IL6R*, *IKBKB*, *STIP1*, *JAK2* |
| GO:0098542 | Defense response to other organism | 304 | 2.63 | 1.00E-10 | 2.09E-09 | *CXCL9*, *CXCL10*, *ISG20*, *GBP1*, *IFNG*, *BATF2*, *RSAD2*, *ISG15*, *MX1*, *GBP2*, *CXCL13*, *MX2*, *AMCF-II*, *IRF1*, *OAS1*, *CCL2*, *CCL4*, *DHX58*, *CXCL8*, *OAS2*, *IRF7*, *ZBP1*, *RIGI*, *S100A12*, *IL27*, *STAT1*, *SAMHD1*, *BST2*, *LBP*, *CALHM6*, *IFIH1*, *BPIFB1*, *RNF19B*, *FOSL1*, *TNFAIP3*, *TIFA*, *CCL8*, *NMI*, *C2*, *SLC15A3*, *SLA-DQB1*, *PIM1*, *IRF8*, *IFI35*, *CFB*, *STAT2*, *CD40*, *C4A*, *LTF*, *CCL5*, *TRIM14*, *FFAR2*, *NOD2*, *TICAM2*, *C3*, *GSDMA*, *UNC93B1*, *HP*, *DUSP10*, |

| | | | | | | |
|---|---|---|---|---|---|---|
| | | | | | | *NFKBIA*, *SERPING1*, *BPIFA1*, *TREM1*, *TRIM26*, *MLKL*, *CXCL16*, *IL10*, *GNLY*, *MR1*, *PPBP*, *CEBPG*, *OPTN*, *STING1*, *TRAF3*, *TRAF2*, *HAVCR2*, *NDUFAF4*, *CEBPB*, *IL18RAP*, *MSRB1*, *MORC3*, *CD14*, *CGAS*, *MYD88*, *TICAM1*, *KLRK1*, *PLSCR1*, *POLR3D*, *SLAMF6*, *TLR2*, *NKG7*, *RIPK3*, *TLR4*, *C9*, *RNF115* |
| GO:0009607 | Response to biotic stimulus | 378 | 2.62 | 1.00E-10 | 2.09E-09 | *CXCL9*, *CXCL10*, *ISG20*, *GBP1*, *IFNG*, *BATF2*, *RSAD2*, *ISG15*, *MX1*, *GBP2*, *CXCL13*, *MX2*, *AMCF-II*, *IRF1*, *IFIT1*, *OAS1*, *CCL2*, *CCL4*, *DHX58*, *CXCL8*, *OAS2*, *IRF7*, *ZBP1*, *RIGI*, *IL1B*, *S100A12*, *S100A8*, *IL27*, *STAT1*, *MMP8*, *SAMHD1*, *BST2*, *LBP*, *CALHM6*, *IL1A*, *IFIH1*, *IFIT2*, *PTGIR*, *BPIFB1*, *RNF19B*, *FOSL1*, *TNFAIP3*, *TIFA*, *CCL8*, *NMI*, *C2*, *SLC15A3*, *SLA-DQB1*, *PIM1*, *IRF8*, *CD180*, *IFI35*, *CFB*, *STAT2*, *CD40*, *C4A*, *LTF*, *CCL5*, *TRIM14*, *FFAR2*, *ZFP36*, *NOD2*, *IFIT5*, *TICAM2*, *C3*, *GSDMA*, *NFKBIB*, *UNC93B1*, *HP*, *DUSP10*, *NFKBIA*, *SERPING1*, *BPIFA1*, *TREM1*, *TRIM26*, *MLKL*, *CXCL16*, *IL10*, *GNLY*, *MR1*, *PPBP*, *CEBPG*, *OPTN*, *STING1*, *TRAF3*, *TIGAR*, *TRAF2*, *TXNIP*, *HAVCR2*, *DDX21*, *NDUFAF4*, *CEBPB*, *IL18RAP*, *MSRB1*, *MORC3*, *CD14*, *CGAS*, *MYD88*, *DDIT3*, *SELENOS*, *TICAM1*, *KLRK1*, *PLSCR1*, *POLR3D*, *SLAMF6*, *CLEC7A*, *TLR2*, *NKG7*, *RIPK3*, *TLR4*, *C9*, *RNF115* |
| GO:0140546 | Defense response to symbiont | 264 | 2.59 | 1.00E-10 | 2.09E-09 | *CXCL9*, *CXCL10*, *ISG20*, *GBP1*, *IFNG*, *RSAD2*, *ISG15*, *MX1*, *GBP2*, *CXCL13*, *AMCF-II*, *IRF1*, *OAS1*, *CCL2*, *CCL4*, *DHX58*, *CXCL8*, *OAS2*, *IRF7*, *ZBP1*, *RIGI*, *S100A12*, *IL27*, *STAT1*, *SAMHD1*, *LBP*, *CALHM6*, *IFIH1*, *BPIFB1*, *RNF19B*, *FOSL1*, *TNFAIP3*, *TIFA*, *CCL8*, *NMI*, *C2*, *SLC15A3*, *SLA-DQB1*, *PIM1*, *IRF8*, *IFI35*, *CFB*, *STAT2*, *C4A*, *LTF*, *CCL5*, *TRIM14*, *FFAR2*, *NOD2*, *TICAM2*, *C3*, *UNC93B1*, *DUSP10*, *NFKBIA*, *SERPING1*, *BPIFA1*, *TREM1*, *TRIM26*, *CXCL16*, *MR1*, *PPBP*, *CEBPG*, *STING1*, *TRAF3*, *TRAF2*, *HAVCR2*, *IL18RAP*, *MSRB1*, *MORC3*, *CD14*, *CGAS*, *MYD88*, *TICAM1*, *KLRK1*, *PLSCR1*, *POLR3D*, *SLAMF6*, *TLR2*, *TLR4*, *C9*, *RNF115* |
| GO:0006955 | Immune response | 468 | 2.59 | 1.00E-10 | 2.09E-09 | *CXCL9*, *CXCL11*, *CXCL10*, *ISG20*, *GBP1*, *IFNG*, *CCL19*, *RSAD2*, *ISG15*, *MX1*, *GBP2*, *CXCL13*, *AMCF-II*, *IRF1*, *OAS1*, *CXCL2*, *CCL2*, *CCL4*, *CCL26*, *DHX58*, *CXCL8*, *OAS2*, *IRF7*, *ZBP1*, *RIGI*, *IL1B*, *CCL3L1*, *S100A12*, *IL27*, *STAT1*, *SAMHD1*, *TNFSF13B*, *LBP*, *CALHM6*, *CCL11*, *SLA-DOA*, *IL6*, *FAS*, *IL1A*, *IFIH1*, *TNFSF10*, *SLA-DRA*, *SLA-3*, *BPIFB1*, *CSF3*, |

| | | | | | | |
|---|---|---|---|---|---|---|
| | | | | | | *RNF19B*, *SLA-DQA1*, *THBS1*, *FOSL1*, *TNFAIP3*, *LOC100622460*, *TIFA*, *CCL8*, *SLA-DOB*, *NMI*, *C2*, *CTLA4*, *CD74*, *SLC15A3*, *SLA-DQB1*, *PIM1*, *IRF8*, *CD180*, *IFI35*, *CFB*, *FASLG*, *SLA-DMB*, *STAT2*, *XCL1*, *ACKR4*, *CD40*, *C4A*, *IL15*, *LTF*, *CCL5*, *TRIM14*, *FFAR2*, *NOD2*, *NFIL3*, *SLA-DMA*, *SLA-DRB1*, *TICAM2*, *C3*, *UNC93B1*, *CXCR6*, *CCR1*, *CCL28*, *CSF2*, *LIF*, *DUSP10*, *NFKBIA*, *SERPING1*, *BPIFA1*, *TREM1*, *STAT3*, *TFRC*, *TNFSF14*, *TRIM26*, *CXCL16*, *IL10*, *KIT*, *TNFSF13*, *MR1*, *PPBP*, *CEBPG*, *ACKR2*, *BCL6*, *STING1*, *MAD2L2*, *TRAF3*, *TRAF2*, *HAVCR2*, *DDX21*, *IL18RAP*, *MSRB1*, *MORC3*, *JAK3*, *CD14*, *CGAS*, *PARP3*, *MYD88*, *TCIM*, *SELENOS*, *ACKR3*, *TICAM1*, *ULBP1*, *KLRK1*, *PLSCR1*, *POLR3D*, *SLAMF6*, *TNFSF9*, *TNFAIP1*, *SLA-8*, *BMX*, *TLR2*, *RIPK3*, *TLR4*, *ST3GAL1*, *C9*, *RNF115*, *TGFB3*, *TLR3*, *IL6R*, *TNIP2*, *CFI*, *PSMB4*, *JAK2* |
| GO:0044419 | Biological process involved in interspecies interaction between organisms | 399 | 2.58 | 1.00E-10 | 2.09E-09 | *CXCL9*, *CXCL10*, *ISG20*, *GBP1*, *IFNG*, *BATF2*, *RSAD2*, *ISG15*, *MX1*, *GBP2*, *CXCL13*, *MX2*, *AMCF-II*, *IRF1*, *IFIT1*, *OAS1*, *CCL2*, *CCL4*, *DHX58*, *CXCL8*, *OAS2*, *IRF7*, *ZBP1*, *RIGI*, *IL1B*, *S100A12*, *S100A8*, *IL27*, *STAT1*, *MMP8*, *SAMHD1*, *BST2*, *LBP*, *CALHM6*, *IL1A*, *IFIH1*, *IFIT2*, *PTGIR*, *BPIFB1*, *RNF19B*, *FOSL1*, *TNFAIP3*, *TIFA*, *CCL8*, *NMI*, *C2*, *SLC15A3*, *SLA-DQB1*, *PIM1*, *IRF8*, *CD180*, *IFI35*, *CFB*, *STAT2*, *CD40*, *C4A*, *LTF*, *CCL5*, *TRIM14*, *FFAR2*, *ZFP36*, *NOD2*, *IFIT5*, *TICAM2*, *C3*, *GSDMA*, *NFKBIB*, *UNC93B1*, *HP*, *DUSP10*, *NFKBIA*, *SERPING1*, *BPIFA1*, *TREM1*, *TRIM26*, *MLKL*, *CXCL16*, *ST6GALNAC1*, *IL10*, *GNLY*, *MR1*, *PRF1*, *PPBP*, *CEBPG*, *OPTN*, *STING1*, *TRAF3*, *TRAF2*, *HAVCR2*, *DDX21*, *NDUFAF4*, *CEBPB*, *IL18RAP*, *MSRB1*, *MORC3*, *CD14*, *CGAS*, *MYD88*, *SELENOS*, *TICAM1*, *KLRK1*, *PLSCR1*, *POLR3D*, *SLAMF6*, *CLEC7A*, *TLR2*, *NKG7*, *RIPK3*, *TLR4*, *C9*, *RNF115* |
| GO:0006952 | Defense response | 450 | 2.57 | 1.00E-10 | 2.09E-09 | *CXCL9*, *CXCL11*, *CXCL10*, *SAA3*, *ISG20*, *GBP1*, *IFNG*, *BATF2*, *CCL19*, *ACKR1*, *RSAD2*, *ISG15*, *MX1*, *GBP2*, *CXCL13*, *MX2*, *AMCF-II*, *IRF1*, *OAS1*, *CXCL2*, *CHI3L1*, *CCL2*, *CCL4*, *DHX58*, *CXCL8*, *OAS2*, *IRF7*, *ZBP1*, *RIGI*, *IL1B*, *S100A12*, *S100A8*, *IL27*, *STAT1*, *MMP8*, *SAMHD1*, *BST2*, *LBP*, *CALHM6*, *CCL11*, *IL6*, *IL1A*, *IFIH1*, *PTGIR*, *BPIFB1*, *RNF19B*, *THBS1*, *FOSL1*, *TNFAIP3*, *TIFA*, *CCL8*, *NMI*, *C2*, *CD74*, *SLC15A3*, *SLA-DQB1*, *PIM1*, *IL4R*, *IRF8*, *IFI35*, *CFB*, *STAT2*, *CD40*, *C4A*, *LTF*, *CCL5*, *TRIM14*, *FFAR2*, *ZFP36*, *NOD2*, *STAT5A*, |

| | | | | | | |
|---|---|---|---|---|---|---|
| | | | | | | *TICAM2*, *C3*, *IL1RAP*, *CASP1*, *GSDMA*, *NFKBIB*, *UNC93B1*, *HP*, *CXCR6*, *CCR1*, *DUSP10*, *NFKBIA*, *SERPING1*, *BPIFA1*, *TREM1*, *STAT3*, *OLR1*, *SLC39A8*, *TRIM26*, *MLKL*, *CXCL16*, *IL10*, *KIT*, *GNLY*, *MR1*, *CD163*, *PPBP*, *APOD*, *CEBPG*, *OPTN*, *BCL6*, *STING1*, *TRAF3*, *TRAF2*, *HAVCR2*, *NDUFAF4*, *CEBPB*, *IL2RA*, *IL18RAP*, *MSRB1*, *MORC3*, *CD14*, *CGAS*, *B4GALT1*, *MYD88*, *TNFRSF1A*, *SELENOS*, *TICAM1*, *KLRK1*, *PLSCR1*, *POLR3D*, *SLAMF6*, *ITIH4*, *TLR2*, *NKG7*, *RIPK3*, *TLR4*, *ELF4*, *C9*, *RNF115*, *TNFRSF1B*, *TLR3* |
| GO:0045087 | Innate immune response | 248 | 2.48 | 1.00E-10 | 2.09E-09 | *CXCL10*, *ISG20*, *GBP1*, *IFNG*, *RSAD2*, *ISG15*, *MX1*, *GBP2*, *IRF1*, *OAS1*, *CCL2*, *DHX58*, *OAS2*, *IRF7*, *ZBP1*, *RIGI*, *S100A12*, *IL27*, *STAT1*, *SAMHD1*, *LBP*, *CALHM6*, *IFIH1*, *BPIFB1*, *RNF19B*, *FOSL1*, *TNFAIP3*, *TIFA*, *NMI*, *C2*, *SLC15A3*, *SLA-DQB1*, *PIM1*, *IRF8*, *IFI35*, *CFB*, *STAT2*, *C4A*, *LTF*, *CCL5*, *TRIM14*, *FFAR2*, *NOD2*, *TICAM2*, *C3*, *UNC93B1*, *DUSP10*, *NFKBIA*, *SERPING1*, *BPIFA1*, *TREM1*, *TRIM26*, *CXCL16*, *MR1*, *CEBPG*, *STING1*, *TRAF3*, *TRAF2*, *HAVCR2*, *IL18RAP*, *MSRB1*, *MORC3*, *CD14*, *CGAS*, *MYD88*, *TICAM1*, *KLRK1*, *PLSCR1*, *POLR3D*, *SLAMF6*, *TLR2*, *TLR4*, *C9*, *RNF115*, *TLR3*, *TNIP2*, *JAK2*, *LYAR*, *SFTPD*, *RAB20*, *TIRAP*, *CDC42EP2*, *CD59*, *ATP1B1*, *TMEM126A*, *IFNGR1*, *CD46*, *IRAK4*, *LY96*, *KCNK6*, *C7*, *USP44* |
| GO:0006954 | Inflammatory response | 217 | 2.36 | 1.00E-10 | 2.09E-09 | *CXCL9*, *CXCL10*, *SAA3*, *IFNG*, *CCL19*, *ACKR1*, *CXCL13*, *AMCF-II*, *CXCL2*, *CHI3L1*, *CCL2*, *CCL4*, *CXCL8*, *IL1B*, *S100A12*, *S100A8*, *IL27*, *MMP8*, *CCL11*, *IL6*, *IL1A*, *PTGIR*, *THBS1*, *FOSL1*, *TNFAIP3*, *CCL8*, *CD74*, *IL4R*, *IFI35*, *C4A*, *CCL5*, *TRIM14*, *FFAR2*, *ZFP36*, *STAT5A*, *TICAM2*, *C3*, *IL1RAP*, *CASP1*, *GSDMA*, *NFKBIB*, *HP*, *CXCR6*, *CCR1*, *DUSP10*, *NFKBIA*, *STAT3*, *OLR1*, *SLC39A8*, *IL10*, *KIT*, *CD163*, *PPBP*, *APOD*, *BCL6*, *CEBPB*, *IL2RA*, *CD14*, *B4GALT1*, *MYD88*, *TNFRSF1A*, *SELENOS*, *TICAM1*, *ITIH4*, *TLR2*, *TLR4*, *ELF4*, *TNFRSF1B*, *TLR3* |
| GO:0033993 | Response to lipid | 156 | 2.29 | 1.00E-10 | 2.09E-09 | *CXCL9*, *CXCL10*, *CXCL13*, *AMCF-II*, *CXCL8*, *IL1B*, *S100A8*, *MMP8*, *IL1A*, *PTGIR*, *TNFAIP3*, *AQP3*, *JUNB*, *PIM1*, *IRF8*, *CD180*, *LTF*, *FFAR2*, *ZFP36*, *AARD*, *NFKBIB*, *DUSP10*, *NFKBIA*, *IL10*, *PPBP*, *HAVCR2*, *GPR155*, *CEBPB*, *CALR*, *SELENOS*, *TICAM1*, *KLRK1*, *GDAP1*, *TLR4*, |

| | | | | | | |
|---|---|---|---|---|---|---|
| | | | | | | *NKX3-1*, *TGFB3*, *KLF9*, *TNIP2*, *INSIG1*, *SLC11A1*, *JAK2*, *PLSCR4*, *DDIT4*, *PLSCR3*, *GOT1*, *TIRAP*, *PTGES3*, *SPP1*, *CYP26B1* |
| GO:0002831 | Regulation of response to biotic stimulus | 139 | 2.26 | 9.09E-10 | 1.47E-08 | *RSAD2*, *ISG15*, *IRF1*, *DHX58*, *IRF7*, *ZBP1*, *SAMHD1*, *CALHM6*, *BPIFB1*, *FOSL1*, *TNFAIP3*, *TIFA*, *SLC15A3*, *PIM1*, *CD180*, *IFI35*, *STAT2*, *LTF*, *FFAR2*, *TICAM2*, *UNC93B1*, *DUSP10*, *NFKBIA*, *SERPING1*, *OPTN*, *STING1*, *TRAF3*, *TIGAR*, *HAVCR2*, *IL18RAP*, *CGAS*, *MYD88*, *TICAM1*, *KLRK1*, *PLSCR1*, *POLR3D*, *SLAMF6*, *TLR2*, *TLR4*, *RNF115*, *TNIP2*, *LYAR*, *TIRAP*, *TMEM126A*, *CD46*, *ARG2*, *LY96*, *KCNK6* |
| GO:0045088 | Regulation of innate immune response | 126 | 2.22 | 3.70E-09 | 5.27E-08 | *RSAD2*, *ISG15*, *IRF1*, *DHX58*, *IRF7*, *ZBP1*, *SAMHD1*, *CALHM6*, *BPIFB1*, *FOSL1*, *TNFAIP3*, *TIFA*, *SLC15A3*, *PIM1*, *IFI35*, *STAT2*, *FFAR2*, *TICAM2*, *UNC93B1*, *DUSP10*, *NFKBIA*, *SERPING1*, *STING1*, *HAVCR2*, *IL18RAP*, *CGAS*, *MYD88*, *TICAM1*, *KLRK1*, *PLSCR1*, *POLR3D*, *SLAMF6*, *TLR2*, *TLR4*, *RNF115*, *TNIP2*, *LYAR*, *TIRAP*, *TMEM126A*, *CD46*, *LY96*, *KCNK6* |
| GO:0001819 | Positive regulation of cytokine production | 134 | 2.20 | 2.69E-09 | 3.99E-08 | *IFNG*, *RSAD2*, *ISG15*, *IRF1*, *CHI3L1*, *DHX58*, *IRF7*, *IL1B*, *STAT1*, *MMP8*, *IL1A*, *THBS1*, *CD74*, *IRF8*, *CD40*, *IL15*, *FFAR2*, *STAT5A*, *TICAM2*, *CASP1*, *ADRA2A*, *EGR1*, *IL10*, *KIT*, *CEBPG*, *PANX2*, *STING1*, *TRAF2*, *HAVCR2*, *DDX21*, *CEBPB*, *CGAS*, *MYD88*, *DDIT3*, *CASP8*, *TICAM1*, *KLRK1*, *POLR3D*, *SLAMF6*, *TLR2*, *TLR4* |
| GO:0071396 | Cellular response to lipid | 120 | 2.20 | 2.69E-08 | 2.74E-07 | *CXCL9*, *CXCL10*, *CXCL13*, *AMCF-II*, *CXCL8*, *IL1B*, *MMP8*, *IL1A*, *TNFAIP3*, *PIM1*, *IRF8*, *CD180*, *LTF*, *FFAR2*, *ZFP36*, *NFKBIB*, *NFKBIA*, *IL10*, *PPBP*, *HAVCR2*, *GPR155*, *CALR*, *SELENOS*, *TICAM1*, *KLRK1*, *NKX3-1*, *KLF9*, *TNIP2*, *INSIG1*, *JAK2*, *PLSCR4*, *DDIT4*, *PLSCR3*, *PTGES3*, *SPP1*, *CYP26B1*, *LY96*, *CRY1*, *VDR*, *AQP1*, *PTK2B*, *MGST1* |
| GO:0009617 | Response to bacterium | 128 | 2.19 | 1.82E-08 | 2.03E-07 | *CXCL9*, *CXCL10*, *ISG15*, *CXCL13*, *AMCF-II*, *DHX58*, *CXCL8*, *IL1B*, *S100A8*, *MMP8*, *LBP*, *IL1A*, *PTGIR*, *TNFAIP3*, *IRF8*, *CD180*, *LTF*, *ZFP36*, *GSDMA*, *NFKBIB*, *HP*, *DUSP10*, *NFKBIA*, *BPIFA1*, *TREM1*, *IL10*, *GNLY*, *PPBP*, *OPTN*, *HAVCR2*, *CEBPB*, *MYD88*, *SELENOS*, *TICAM1*, *KLRK1*, *TLR2*, *TLR4*, *TNIP2*, *SLC11A1*, *JAK2*, *PLSCR4* |

| | | | | | | |
|---|---|---|---|---|---|---|
| GO:0002682 | Regulation of immune system process | 390 | 2.18 | 1.00E-10 | 2.09E-09 | *CXCL10*, *MADCAM1*, *IFNG*, *SELE*, *RSAD2*, *ISG15*, *CD274*, *IRF1*, *CCL2*, *DHX58*, *IRF7*, *ZBP1*, *IL1B*, *IL27*, *STAT1*, *MMP8*, *SAMHD1*, *TNFSF13B*, *CALHM6*, *SOCS1*, *IL1A*, *BPIFB1*, *CSF3*, *THBS1*, *FOSL1*, *TNFAIP3*, *AQP3*, *LOC100622460*, *TIFA*, *C2*, *CTLA4*, *SLC15A3*, *SLA-DQB1*, *PIM1*, *IFI35*, *CFB*, *SLA-DMB*, *STAT2*, *CD40*, *C4A*, *IL15*, *LTF*, *CCL5*, *FFAR2*, *ZFP36*, *STAT5A*, *TICAM2*, *C3*, *GDF15*, *UNC93B1*, *LRRC32*, *CCR1*, *LIF*, *DUSP10*, *NFKBIA*, *SERPING1*, *STAT3*, *TFRC*, *TNFSF14*, *IL10*, *KIT*, *CEACAM1*, *APOD*, *BCL6*, *STING1*, *MAD2L2*, *TRAF2*, *TMEM176B*, *HAVCR2*, *SLC7A1*, *DDX21*, *FBXO7*, *CEBPB*, *IL18RAP*, *JAK3*, *CGAS*, *PRDM1*, *PARP3*, *MYD88*, *HSPA9*, *TCIM*, *CASP8*, *CALR*, *SELENOS*, *TICAM1*, *ULBP1*, *KLRK1*, *SOX11*, *PLSCR1*, *POLR3D*, *SLAMF6*, *MYC*, *TNFSF9*, *BMX*, *TLR2*, *RIPK3*, *TLR4*, *C9*, *RNF115*, *TGFB3*, *TNIP2*, *CFI*, *IGFBP2*, *PSMB4*, *JAK2* |
| GO:0001775 | Cell activation | 261 | 2.16 | 1.00E-10 | 2.09E-09 | *CXCL10*, *IFNG*, *BATF2*, *RSAD2*, *CD274*, *AMCF-II*, *IRF1*, *CXCL8*, *IL1B*, *IL27*, *MMP8*, *TNFSF13B*, *IL6*, *SOCS1*, *IL1A*, *THBS1*, *TNFAIP3*, *CTLA4*, *SLA-DQB1*, *IRF8*, *CD180*, *IFI35*, *SLA-DMB*, *CD40*, *IL15*, *CCL5*, *NFIL3*, *STAT5A*, *ADRA2A*, *LRRC32*, *HSH2D*, *CTPS1*, *DUSP10*, *EGR1*, *STAT3*, *TFRC*, *TNFSF14*, *IL10*, *KIT*, *GJA1*, *CEBPG*, *FGB*, *BCL6*, *DLL4*, *MAD2L2*, *TRAF2*, *HAVCR2*, *SLC7A1*, *FBXO7*, *CEBPB*, *IL18RAP*, *JAK3*, *PRDM1*, *PARP3*, *TCIM*, *TICAM1*, *KLRK1*, *SOX11*, *PLSCR1*, *SLAMF6*, *TNFSF9*, *RIPK3*, *TLR4*, *ELF4*, *ST3GAL1*, *IL6R*, *TMEM106A*, *IGFBP2*, *JAK2*, *LIG4*, *CD200*, *GNA13*, *LRRC8A*, *CYP26B1*, *KAT2A*, *SLC25A5* |
| GO:0045321 | Leukocyte activation | 226 | 2.16 | 1.95E-10 | 3.86E-09 | *IFNG*, *BATF2*, *RSAD2*, *CD274*, *AMCF-II*, *IRF1*, *CXCL8*, *IL1B*, *IL27*, *MMP8*, *TNFSF13B*, *IL6*, *SOCS1*, *IL1A*, *THBS1*, *TNFAIP3*, *CTLA4*, *SLA-DQB1*, *IRF8*, *CD180*, *IFI35*, *SLA-DMB*, *CD40*, *IL15*, *CCL5*, *NFIL3*, *STAT5A*, *LRRC32*, *HSH2D*, *CTPS1*, *DUSP10*, *EGR1*, *STAT3*, *TFRC*, *TNFSF14*, *IL10*, *KIT*, *GJA1*, *CEBPG*, *BCL6*, *DLL4*, *MAD2L2*, *TRAF2*, *HAVCR2*, *SLC7A1*, *FBXO7*, *CEBPB*, *IL18RAP*, *JAK3*, *PRDM1*, *PARP3*, *TICAM1*, *KLRK1*, *SOX11*, *PLSCR1*, *SLAMF6*, *TNFSF9*, *RIPK3*, *TLR4*, *ELF4*, *ST3GAL1*, *IL6R*, *TMEM106A*, *IGFBP2*, *JAK2* |
| GO:0042254 | Ribosome biogenesis | 129 | 2.15 | 6.37E-08 | 5.53E-07 | *ISG20*, *NPM3*, *RRP9*, *EMG1*, *FBL*, *MRTO4*, *EXOSC5*, *UTP11*, *NHP2*, *NOP2*, *RRS1*, *NOB1*, *LYAR*, *PPAN*, *GAR1*, *NOC4L*, *GTPBP4*, *WDR74*, *WDR75*, *GRWD1*, *NOL6*, *BMS1*, *REXO4*, *IMP3*, *RRP36*, *BUD23*, *RCL1*, *WDR12*, *NLE1*, *NOP9*, *MTERF3*, *NAT10*, *RSL24D1*, *UTP18*, |

| | | | | | | |
|---|---|---|---|---|---|---|
| | | | | | | *NIP7*, *NOP56*, *RRP15*, *RPL7L1*, *WDR46*, *UTP23*, *UTP15*, *DDX3X*, *KRI1*, *ESF1*, *SDAD1*, *DDX49*, *PNO1*, *NPM1*, *ABT1*, *EBNA1BP2*, *DDX31*, *RPP40*, *NOL9*, *EXOSC3*, *UTP6*, *AK6*, *TSR2*, *WDR36*, *EIF6*, *TMA16*, *DDX52*, *DDX28*, *LOC100621115*, *PWP1*, *EIF4A3*, *MRPS7*, *MTREX*, *EXOSC4*, *FCF1*, *NSUN4*, *LSM6* |
| GO:1903131 | Mononuclear cell differentiation | 123 | 2.15 | 5.77E-08 | 5.27E-07 | *IFNG*, *BATF2*, *RSAD2*, *IRF1*, *IL1B*, *IL27*, *TNFSF13B*, *IL6*, *SOCS1*, *IL1A*, *SLA-DQB1*, *IRF8*, *NFIL3*, *STAT5A*, *LIF*, *DUSP10*, *EGR1*, *STAT3*, *IL10*, *KIT*, *CEBPG*, *BCL6*, *DLL4*, *TMEM176B*, *FBXO7*, *CEBPB*, *JAK3*, *PRDM1*, *CASP8*, *SLAMF6*, *MYC*, *TNFSF9*, *RIPK3*, *TLR4*, *ST3GAL1*, *IL6R*, *LIG4*, *LRRC8A*, *CYP26B1*, *KAT2A*, *SLC25A5* |
| GO:0032103 | Positive regulation of response to external stimulus | 166 | 2.12 | 1.27E-08 | 1.51E-07 | *CXCL10*, *IFNG*, *RSAD2*, *IRF1*, *DHX58*, *CXCL8*, *IRF7*, *ZBP1*, *S100A8*, *MMP8*, *BPIFB1*, *THBS1*, *FOSL1*, *TNFAIP3*, *TIFA*, *CD74*, *SLC15A3*, *CD180*, *IFI35*, *CCL5*, *FFAR2*, *STAT5A*, *TICAM2*, *UNC93B1*, *CCR1*, *NFKBIA*, *TNFSF14*, *OPTN*, *STING1*, *HAVCR2*, *CEBPB*, *IL18RAP*, *CGAS*, *MYD88*, *TNFRSF1A*, *CALR*, *TICAM1*, *KLRK1*, *PLSCR1*, *POLR3D*, *SLAMF6*, *TLR2*, *TLR4*, *RNF115* |
| GO:0022613 | Ribonucleoprotein complex biogenesis | 172 | 2.11 | 1.18E-08 | 1.45E-07 | *ISG20*, *NPM3*, *RRP9*, *EMG1*, *FBL*, *MRTO4*, *EXOSC5*, *UTP11*, *NHP2*, *NOP2*, *RRS1*, *NOB1*, *LYAR*, *PPAN*, *GAR1*, *NOC4L*, *GTPBP4*, *WDR74*, *WDR75*, *PTGES3*, *GRWD1*, *SNRPD1*, *NOL6*, *BMS1*, *REXO4*, *IMP3*, *SNRPF*, *SHQ1*, *RRP36*, *BUD23*, *NOPCHAP1*, *RCL1*, *WDR12*, *EIF3I*, *NLE1*, *NOP9*, *MTERF3*, *NAT10*, *RSL24D1*, *UTP18*, *NIP7*, *NOP56*, *RRP15*, *RPL7L1*, *WDR46*, *UTP23*, *UTP15*, *DDX3X*, *KRI1*, *ESF1*, *EIF3J*, *SDAD1*, *DDX49*, *PNO1*, *NPM1*, *ABT1*, *EBNA1BP2*, *SNRPG*, *DDX31*, *RPP40*, *NOL9*, *EXOSC3*, *UTP6*, *AK6*, *STRAP*, *SNRPD3*, *TSR2*, *EIF3A*, *GEMIN6*, *SRSF1*, *WDR36*, *EIF6*, *TMA16*, *DDX52*, *DDX28*, *USP39*, *TAF9*, *NUFIP1*, *LOC100621115*, *PWP1*, *EIF4A3*, *MRPS7*, *MTREX*, *EXOSC4*, *CLNS1A*, *FCF1*, *NSUN4*, *LSM6*, *SNRPC* |
| GO:0002684 | Positive regulation of immune system process | 265 | 2.11 | 2.14E-10 | 4.01E-09 | *CXCL10*, *MADCAM1*, *IFNG*, *RSAD2*, *IRF1*, *DHX58*, *IRF7*, *ZBP1*, *IL1B*, *MMP8*, *TNFSF13B*, *SOCS1*, *IL1A*, *BPIFB1*, *THBS1*, *FOSL1*, *TNFAIP3*, *AQP3*, *TIFA*, *C2*, *SLC15A3*, *SLA-DQB1*, *IFI35*, *CFB*, *SLA-DMB*, *CD40*, *C4A*, *IL15*, *CCL5*, *FFAR2*, *STAT5A*, *TICAM2*, *C3*, *UNC93B1*, *CCR1*, *LIF*, *DUSP10*, *NFKBIA*, *SERPING1*, *TFRC*, *TNFSF14*, *IL10*, *KIT*, *BCL6*, *STING1*, |

| | | | | | | |
|---|---|---|---|---|---|---|
| | | | | | | *MAD2L2*, *TRAF2*, *HAVCR2*, *SLC7A1*, *DDX21*, *IL18RAP*, *CGAS*, *MYD88*, *CASP8*, *CALR*, *TICAM1*, *ULBP1*, *KLRK1*, *PLSCR1*, *POLR3D*, *SLAMF6*, *TNFSF9*, *BMX*, *TLR2*, *TLR4*, *C9*, *RNF115*, *TNIP2*, *CFI*, *IGFBP2*, *JAK2* |
| GO:0050776 | Regulation of immune response | 232 | 2.11 | 8.07E-10 | 1.37E-08 | *IFNG*, *RSAD2*, *ISG15*, *IRF1*, *DHX58*, *IRF7*, *ZBP1*, *IL1B*, *IL27*, *SAMHD1*, *TNFSF13B*, *CALHM6*, *BPIFB1*, *FOSL1*, *TNFAIP3*, *LOC100622460*, *TIFA*, *C2*, *SLC15A3*, *SLA-DQB1*, *PIM1*, *IFI35*, *CFB*, *SLA-DMB*, *STAT2*, *CD40*, *C4A*, *IL15*, *FFAR2*, *TICAM2*, *C3*, *UNC93B1*, *DUSP10*, *NFKBIA*, *SERPING1*, *TFRC*, *IL10*, *KIT*, *BCL6*, *STING1*, *MAD2L2*, *TRAF2*, *HAVCR2*, *IL18RAP*, *JAK3*, *CGAS*, *PARP3*, *MYD88*, *SELENOS*, *TICAM1*, *ULBP1*, *KLRK1*, *PLSCR1*, *POLR3D*, *SLAMF6*, *BMX*, *TLR2*, *RIPK3*, *TLR4*, *C9*, *RNF115*, *TNIP2*, *CFI*, *PSMB4* |
| GO:0031347 | Regulation of defense response | 201 | 2.10 | 4.24E-09 | 5.59E-08 | *IFNG*, *RSAD2*, *ISG15*, *IRF1*, *DHX58*, *IRF7*, *ZBP1*, *S100A8*, *MMP8*, *SAMHD1*, *CALHM6*, *BPIFB1*, *FOSL1*, *TNFAIP3*, *TIFA*, *CD74*, *SLC15A3*, *PIM1*, *IFI35*, *STAT2*, *FFAR2*, *ZFP36*, *STAT5A*, *TICAM2*, *CASP1*, *UNC93B1*, *DUSP10*, *NFKBIA*, *SERPING1*, *SLC39A8*, *IL10*, *OPTN*, *BCL6*, *STING1*, *TRAF3*, *HAVCR2*, *CEBPB*, *IL18RAP*, *CGAS*, *MYD88*, *TNFRSF1A*, *SELENOS*, *TICAM1*, *KLRK1*, *PLSCR1*, *POLR3D*, *SLAMF6*, *TLR2*, *TLR4*, *ELF4*, *RNF115*, *TNIP2*, *PSMB4*, *JAK2*, *LYAR*, *ADORA1*, *RIPK1*, *TIRAP*, *TMEM126A*, *GPR4*, *CD46*, *ARG2*, *LY96*, *KCNK6* |
| GO:0001816 | Cytokine production | 207 | 2.08 | 4.78E-09 | 6.08E-08 | *IFNG*, *RSAD2*, *ISG15*, *IRF1*, *CHI3L1*, *DHX58*, *IRF7*, *IL1B*, *STAT1*, *MMP8*, *CALHM6*, *IL6*, *SOCS1*, *IL1A*, *THBS1*, *TNFAIP3*, *CD74*, *IRF8*, *CD40*, *IL15*, *LTF*, *FFAR2*, *ZFP36*, *STAT5A*, *TICAM2*, *CASP1*, *ADRA2A*, *LRRC32*, *CSF2*, *EGR1*, *TSKU*, *IL10*, *KIT*, *APOD*, *CEBPG*, *PANX2*, *BCL6*, *STING1*, *TRAF3*, *TRAF2*, *HAVCR2*, *DDX21*, *CEBPB*, *MORC3*, *CGAS*, *MYD88*, *DDIT3*, *CASP8*, *TICAM1*, *KLRK1*, *POLR3D*, *SLAMF6*, *TLR2*, *RIPK3*, *TLR4*, *ELF4*, *TGFB3*, *IL6R*, *TMEM106A* |
| GO:0046649 | Lymphocyte activation | 192 | 2.07 | 2.14E-08 | 2.31E-07 | *IFNG*, *RSAD2*, *CD274*, *IRF1*, *IL1B*, *IL27*, *TNFSF13B*, *IL6*, *SOCS1*, *IL1A*, *TNFAIP3*, *CTLA4*, *SLA-DQB1*, *IRF8*, *CD180*, *SLA-DMB*, *CD40*, *IL15*, *CCL5*, *NFIL3*, *STAT5A*, *LRRC32*, *HSH2D*, *CTPS1*, *DUSP10*, *EGR1*, *STAT3*, *TFRC*, *TNFSF14*, *IL10*, *KIT*, *GJA1*, *CEBPG*, *BCL6*, *DLL4*, *MAD2L2*, *TRAF2*, *HAVCR2*, *SLC7A1*, *FBXO7*, *CEBPB*, *JAK3*, *PRDM1*, *PARP3*, *TICAM1*, |

| | | | | | | |
|---|---|---|---|---|---|---|
| | | | | | | *KLRK1*, *SOX11*, *SLAMF6*, *TNFSF9*, *RIPK3*, *TLR4*, *ELF4*, *ST3GAL1*, *IL6R*, *IGFBP2*, *JAK2*, *LIG4*, *LRRC8A*, *CYP26B1*, *KAT2A*, *SLC25A5* |
| GO:0032101 | Regulation of response to external stimulus | 258 | 2.07 | 4.15E-10 | 7.39E-09 | *CXCL10*, *IFNG*, *RSAD2*, *ISG15*, *IRF1*, *CCL2*, *DHX58*, *CXCL8*, *IRF7*, *ZBP1*, *S100A8*, *MMP8*, *SERPINE1*, *SAMHD1*, *CALHM6*, *BPIFB1*, *THBS1*, *FOSL1*, *TNFAIP3*, *TIFA*, *CD74*, *SLC15A3*, *PIM1*, *CD180*, *IFI35*, *STAT2*, *LTF*, *CCL5*, *FFAR2*, *ZFP36*, *STAT5A*, *TICAM2*, *CASP1*, *UNC93B1*, *CCR1*, *DUSP10*, *NFKBIA*, *SERPING1*, *SLC39A8*, *TNFSF14*, *IL10*, *OPTN*, *BCL6*, *STING1*, *TRAF3*, *HAVCR2*, *CEBPB*, *IL18RAP*, *CGAS*, *MYD88*, *TNFRSF1A*, *CALR*, *SELENOS*, *TICAM1*, *KLRK1*, *PLSCR1*, *POLR3D*, *SLAMF6*, *TLR2*, *TLR4*, *ELF4*, *RNF115* |
| GO:0002521 | Leukocyte differentiation | 147 | 2.06 | 1.20E-07 | 1.02E-06 | *IFNG*, *BATF2*, *RSAD2*, *IRF1*, *IL1B*, *IL27*, *TNFSF13B*, *IL6*, *SOCS1*, *IL1A*, *SNX10*, *JUNB*, *SLA-DQB1*, *IRF8*, *LTF*, *NFIL3*, *STAT5A*, *CCR1*, *LIF*, *DUSP10*, *EGR1*, *STAT3*, *IL10*, *KIT*, *CEBPG*, *BCL6*, *DLL4*, *TMEM176B*, *FBXO7*, *CEBPB*, *JAK3*, *PRDM1*, *CASP8*, *SLAMF6*, *MYC*, *TNFSF9*, *RIPK3*, *TLR4*, *ST3GAL1* |
| GO:0006935 | Chemotaxis | 121 | 2.06 | 1.75E-06 | 1.06E-05 | *CXCL9*, *CXCL11*, *CXCL10*, *CXCL13*, *AMCF-II*, *CCL2*, *CCL4*, *CXCL8*, *S100A8*, *THBS1*, *CCL8*, *ACKR4*, *CCL5*, *FFAR2*, *CXCR6*, *CCR1*, *TREM1*, *TNFSF14*, *CXCL16*, *KIT*, *PPBP*, *ACKR2*, *CALR*, *ACKR3*, *KLRK1* |
| GO:0042330 | Taxis | 121 | 2.06 | 1.75E-06 | 1.06E-05 | *CXCL9*, *CXCL11*, *CXCL10*, *CXCL13*, *AMCF-II*, *CCL2*, *CCL4*, *CXCL8*, *S100A8*, *THBS1*, *CCL8*, *ACKR4*, *CCL5*, *FFAR2*, *CXCR6*, *CCR1*, *TREM1*, *TNFSF14*, *CXCL16*, *KIT*, *PPBP*, *ACKR2*, *CALR*, *ACKR3*, *KLRK1* |
| GO:0045089 | Positive regulation of innate immune response | 105 | 2.04 | 5.19E-06 | 2.68E-05 | *RSAD2*, *IRF1*, *DHX58*, *IRF7*, *ZBP1*, *BPIFB1*, *FOSL1*, *TNFAIP3*, *TIFA*, *SLC15A3*, *IFI35*, *FFAR2*, *TICAM2*, *UNC93B1*, *NFKBIA*, *STING1*, *HAVCR2*, *IL18RAP*, *CGAS*, *MYD88*, *TICAM1*, *KLRK1*, *PLSCR1*, *POLR3D*, *SLAMF6*, *TLR2*, *TLR4*, *RNF115*, *TNIP2* |
| GO:0031349 | Positive regulation of defense response | 138 | 2.04 | 5.97E-07 | 3.89E-06 | *IFNG*, *RSAD2*, *IRF1*, *DHX58*, *IRF7*, *ZBP1*, *S100A8*, *MMP8*, *BPIFB1*, *FOSL1*, *TNFAIP3*, *TIFA*, *CD74*, *SLC15A3*, *IFI35*, *FFAR2*, *STAT5A*, *TICAM2*, *UNC93B1*, *NFKBIA*, *OPTN*, *STING1*, *HAVCR2*, *CEBPB*, *IL18RAP*, *CGAS*, *MYD88*, *TNFRSF1A*, *TICAM1*, *KLRK1*, *PLSCR1*, *POLR3D*, *SLAMF6*, *TLR2*, *TLR4*, *RNF115* |

| | | | | | | |
|---|---|---|---|---|---|---|
| GO:0001817 | Regulation of cytokine production | 204 | 2.04 | 3.11E-08 | 3.07E-07 | *IFNG*, *RSAD2*, *ISG15*, *IRF1*, *CHI3L1*, *DHX58*, *IRF7*, *IL1B*, *STAT1*, *MMP8*, *CALHM6*, *SOCS1*, *IL1A*, *THBS1*, *TNFAIP3*, *CD74*, *IRF8*, *CD40*, *IL15*, *LTF*, *FFAR2*, *ZFP36*, *STAT5A*, *TICAM2*, *CASP1*, *ADRA2A*, *LRRC32*, *EGR1*, *TSKU*, *IL10*, *KIT*, *APOD*, *CEBPG*, *PANX2*, *BCL6*, *STING1*, *TRAF3*, *TRAF2*, *HAVCR2*, *DDX21*, *CEBPB*, *MORC3*, *CGAS*, *MYD88*, *DDIT3*, *CASP8*, *TICAM1*, *KLRK1*, *POLR3D*, *SLAMF6*, *TLR2*, *RIPK3*, *TLR4*, *ELF4*, *TGFB3*, *TMEM106A* |
| GO:0002694 | Regulation of leukocyte activation | 134 | 2.03 | 4.40E-07 | 3.07E-06 | *IFNG*, *CD274*, *IRF1*, *IL1B*, *IL27*, *MMP8*, *TNFSF13B*, *SOCS1*, *IL1A*, *THBS1*, *TNFAIP3*, *CTLA4*, *SLA-DQB1*, *SLA-DMB*, *CD40*, *IL15*, *CCL5*, *STAT5A*, *LRRC32*, *DUSP10*, *TFRC*, *TNFSF14*, *IL10*, *BCL6*, *MAD2L2*, *HAVCR2*, *SLC7A1*, *FBXO7*, *CEBPB*, *JAK3*, *PRDM1*, *PARP3*, *TICAM1*, *SOX11*, *PLSCR1*, *TNFSF9*, *RIPK3*, *TLR4* |
| GO:0042110 | T cell activation | 133 | 2.03 | 4.29E-07 | 3.06E-06 | *IFNG*, *RSAD2*, *CD274*, *IRF1*, *IL1B*, *IL27*, *TNFSF13B*, *IL6*, *SOCS1*, *IL1A*, *CTLA4*, *SLA-DQB1*, *SLA-DMB*, *IL15*, *CCL5*, *STAT5A*, *LRRC32*, *HSH2D*, *CTPS1*, *DUSP10*, *EGR1*, *STAT3*, *TFRC*, *TNFSF14*, *KIT*, *GJA1*, *BCL6*, *DLL4*, *TRAF2*, *HAVCR2*, *SLC7A1*, *CEBPB*, *JAK3*, *PRDM1*, *SLAMF6*, *TNFSF9*, *RIPK3*, *ELF4*, *IL6R*, *IGFBP2*, *JAK2* |
| GO:0002833 | Positive regulation of response to biotic stimulus | 110 | 2.02 | 3.85E-06 | 2.11E-05 | *RSAD2*, *IRF1*, *DHX58*, *IRF7*, *ZBP1*, *BPIFB1*, *FOSL1*, *TNFAIP3*, *TIFA*, *SLC15A3*, *CD180*, *IFI35*, *FFAR2*, *TICAM2*, *UNC93B1*, *NFKBIA*, *STING1*, *HAVCR2*, *IL18RAP*, *CGAS*, *MYD88*, *TICAM1*, *KLRK1*, *PLSCR1*, *POLR3D*, *SLAMF6*, *TLR2*, *TLR4*, *RNF115*, *TNIP2* |
| GO:0002250 | Adaptive immune response | 118 | 2.01 | 3.82E-06 | 2.11E-05 | *IFNG*, *RSAD2*, *IRF7*, *IL1B*, *IL27*, *TNFSF13B*, *SLA-DOA*, *IL6*, *SLA-DRA*, *SLA-DQA1*, *TNFAIP3*, *SLA-DOB*, *C2*, *CTLA4*, *SLA-DQB1*, *CFB*, *SLA-DMB*, *CD40*, *C4A*, *SLA-DMA*, *SLA-DRB1*, *C3*, *DUSP10*, *TREM1*, *STAT3*, *TFRC*, *BCL6*, *MAD2L2*, *TRAF2*, *HAVCR2*, *IL18RAP*, *JAK3*, *PARP3*, *ULBP1*, *KLRK1*, *SLAMF6*, *BMX*, *RIPK3*, *C9*, *IL6R*, *JAK2*, *FCGR1A*, *LIG4* |
| GO:0002252 | Immune effector process | 159 | 1.99 | 8.03E-07 | 5.10E-06 | *IFNG*, *RSAD2*, *DHX58*, *IRF7*, *IL1B*, *IL27*, *CALHM6*, *IL6*, *RNF19B*, *C2*, *SLA-DQB1*, *IRF8*, *CD180*, *IFI35*, *CFB*, *CD40*, *C4A*, *FFAR2*, *C3*, *DUSP10*, *SERPING1*, *TREM1*, *STAT3*, *TFRC*, *IL10*, *KIT*, *PRF1*, *CEBPG*, *BCL6*, *MAD2L2*, *TRAF2*, *HAVCR2*, *DDX21*, *IL18RAP*, *JAK3*, *PARP3*, *TCIM*, *TICAM1*, *ULBP1*, *KLRK1*, *SLAMF6*, *RIPK3*, *TLR4*, *ST3GAL1*, *C9*, *TGFB3*, *IL6R*, *CFI* |

| | | | | | | |
|---|---|---|---|---|---|---|
| GO:0051249 | Regulation of lymphocyte activation | 120 | 1.99 | 6.42E-06 | 3.19E-05 | *IFNG*, *CD274*, *IRF1*, *IL1B*, *IL27*, *TNFSF13B*, *SOCS1*, *IL1A*, *TNFAIP3*, *CTLA4*, *SLA-DQB1*, *SLA-DMB*, *CD40*, *IL15*, *CCL5*, *STAT5A*, *LRRC32*, *DUSP10*, *TFRC*, *TNFSF14*, *IL10*, *BCL6*, *MAD2L2*, *HAVCR2*, *SLC7A1*, *FBXO7*, *CEBPB*, *JAK3*, *PRDM1*, *PARP3*, *TICAM1*, *SOX11*, *TNFSF9*, *RIPK3*, *TLR4*, *IGFBP2*, *JAK2* |
| GO:0050865 | Regulation of cell activation | 146 | 1.99 | 2.07E-06 | 1.21E-05 | *IFNG*, *CD274*, *IRF1*, *IL1B*, *IL27*, *MMP8*, *TNFSF13B*, *SOCS1*, *IL1A*, *THBS1*, *TNFAIP3*, *CTLA4*, *SLA-DQB1*, *SLA-DMB*, *CD40*, *IL15*, *CCL5*, *STAT5A*, *LRRC32*, *DUSP10*, *TFRC*, *TNFSF14*, *IL10*, *BCL6*, *MAD2L2*, *HAVCR2*, *SLC7A1*, *FBXO7*, *CEBPB*, *JAK3*, *PRDM1*, *PARP3*, *TICAM1*, *SOX11*, *PLSCR1*, *TNFSF9*, *RIPK3*, *TLR4* |
| GO:0080134 | Regulation of response to stress | 333 | 1.98 | 2.67E-09 | 3.99E-08 | *IFNG*, *RSAD2*, *ISG15*, *IRF1*, *DHX58*, *IRF7*, *ZBP1*, *S100A8*, *MMP8*, *SERPINE1*, *SAMHD1*, *CALHM6*, *FAS*, *BPIFB1*, *THBS1*, *FOSL1*, *TNFAIP3*, *TIFA*, *CD74*, *SLC15A3*, *PIM1*, *IFI35*, *STAT2*, *FFAR2*, *ZFP36*, *STAT5A*, *TICAM2*, *CASP1*, *UNC93B1*, *ADRA2A*, *DUSP10*, *NFKBIA*, *SERPING1*, *SESN2*, *SLC39A8*, *IL10*, *CEBPG*, *OPTN*, *BCL6*, *STING1*, *MAD2L2*, *TRAF3*, *TIGAR*, *HAVCR2*, *CEBPB*, *IL18RAP*, *CGAS*, *PARP3*, *MYD88*, *TNFRSF1A*, *YBX3*, *SELENOS*, *ACKR3*, *TICAM1*, *KLRK1*, *PLSCR1*, *POLR3D*, *SLAMF6*, *MYC*, *TLR2*, *RNASEH2B*, *TLR4*, *ELF4*, *PSMD10*, *FICD*, *RNF115*, *TNIP2*, *INSIG1*, *PSMB4*, *JAK2*, *LYAR*, *FBXO4*, *ADORA1*, *RIPK1*, *TIRAP*, *TMEM126A*, *GPR4*, *CD46*, *CLDN4*, *ARG2*, *LY96*, *CRY1*, *KCNK6* |
| GO:0050778 | Positive regulation of immune response | 190 | 1.98 | 2.94E-07 | 2.23E-06 | *IFNG*, *RSAD2*, *IRF1*, *DHX58*, *IRF7*, *ZBP1*, *IL1B*, *TNFSF13B*, *BPIFB1*, *FOSL1*, *TNFAIP3*, *TIFA*, *C2*, *SLC15A3*, *SLA-DQB1*, *IFI35*, *CFB*, *SLA-DMB*, *CD40*, *C4A*, *IL15*, *FFAR2*, *TICAM2*, *C3*, *UNC93B1*, *NFKBIA*, *SERPING1*, *TFRC*, *IL10*, *STING1*, *MAD2L2*, *TRAF2*, *HAVCR2*, *IL18RAP*, *CGAS*, *MYD88*, *TICAM1*, *ULBP1*, *KLRK1*, *PLSCR1*, *POLR3D*, *SLAMF6*, *BMX*, *TLR2*, *TLR4*, *C9*, *RNF115*, *TNIP2*, *CFI* |
| GO:0030097 | Hemopoiesis | 236 | 1.95 | 1.51E-07 | 1.22E-06 | *IFNG*, *BATF2*, *RSAD2*, *ISG15*, *IRF1*, *IL1B*, *IL27*, *STAT1*, *TNFSF13B*, *IL6*, *SOCS1*, *IL1A*, *CSF3*, *SNX10*, *JUNB*, *SLA-DQB1*, *PIM1*, *IRF8*, *LTF*, *ZFP36*, *NFIL3*, *STAT5A*, *CCR1*, *CSF2*, *CEBPD*, *LIF*, *DUSP10*, *NFKBIA*, *EGR1*, *STAT3*, *IL10*, *SLC1A5*, *KIT*, *CASP3*, *CEBPG*, *BCL6*, *DLL4*, *TMEM176B*, *FBXO7*, *CEBPB*, *JAK3*, *PRDM1*, *HSPA9*, *TCIM*, *CASP8*, *SLAMF6*, *MYC*, *TNFSF9*, |

| | | | | | | |
|---|---|---|---|---|---|---|
| | | | | | | *GPATCH4*, *RIPK3*, *TLR4*, *ST3GAL1*, *IL6R*, *JAK2*, *LYAR*, *ETV6*, *LIG4*, *LRRC8A*, *CYP26B1*, *KAT2A*, *SLC25A5* |
| GO:0002764 | Immune response-regulating signaling pathway | 121 | 1.95 | 1.70E-05 | 7.84E-05 | *IFNG*, *RSAD2*, *IRF1*, *DHX58*, *IRF7*, *BPIFB1*, *FOSL1*, *TNFAIP3*, *LOC100622460*, *TIFA*, *SLC15A3*, *IFI35*, *CD40*, *FFAR2*, *TICAM2*, *UNC93B1*, *NFKBIA*, *KIT*, *STING1*, *HAVCR2*, *CGAS*, *MYD88*, *TICAM1*, *KLRK1*, *PLSCR1*, *BMX*, *TLR2*, *TLR4*, *RNF115*, *TNIP2* |
| GO:0022407 | Regulation of cell-cell adhesion | 103 | 1.95 | 2.39E-05 | 1.08E-04 | *IFNG*, *SELE*, *CD274*, *IRF1*, *IL1B*, *TNFSF13B*, *SOCS1*, *IL1A*, *SLA-DQB1*, *SLA-DMB*, *IL15*, *CCL5*, *STAT5A*, *LRRC32*, *DUSP10*, *NR4A3*, *TFRC*, *TNFSF14*, *IL10*, *BCL6*, *MAD2L2*, *HAVCR2*, *SLC7A1*, *CEBPB*, *JAK3* |
| GO:0002683 | Negative regulation of immune system process | 116 | 1.94 | 8.50E-06 | 4.04E-05 | *ISG15*, *CD274*, *IRF1*, *CCL2*, *DHX58*, *SOCS1*, *BPIFB1*, *THBS1*, *TNFAIP3*, *PIM1*, *STAT2*, *LTF*, *GDF15*, *LRRC32*, *DUSP10*, *SERPING1*, *IL10*, *APOD*, *BCL6*, *TMEM176B*, *HAVCR2*, *FBXO7*, *CEBPB*, *JAK3*, *CGAS*, *PARP3*, *HSPA9*, *SELENOS*, *KLRK1*, *SOX11*, *MYC*, *RNF115*, *TGFB3*, *PSMB4*, *LYAR*, *ADORA1*, *CD200*, *CD59*, *CD46*, *ARG2* |
| GO:1901700 | Response to oxygen-containing compound | 311 | 1.93 | 2.28E-08 | 2.39E-07 | *CXCL9*, *CXCL10*, *CXCL13*, *AMCF-II*, *CXCL8*, *IL1B*, *S100A8*, *STAT1*, *MMP8*, *SOCS1*, *IL1A*, *LPL*, *PTGIR*, *THBS1*, *TNFAIP3*, *AQP3*, *PIM1*, *IRF8*, *CD180*, *STAT2*, *LTF*, *FFAR2*, *ZFP36*, *AARD*, *STAT5A*, *GDF15*, *NFKBIB*, *ADRA2A*, *DUSP10*, *NFKBIA*, *EGR1*, *STAT3*, *SESN2*, *NR4A3*, *SLC2A4*, *IL10*, *GPR37*, *PLK3*, *PPBP*, *APOD*, *GJA1*, *RHOB*, *TRAF2*, *HAVCR2*, *GPR155*, *CEBPB*, *JAK3*, *CHRNA3*, *SELENOS*, *TICAM1*, *KLRK1*, *GDAP1*, *TLR2*, *RIPK3*, *TLR4*, *NKX3-1*, *TGFB3*, *KLF9*, *SLC25A33*, *TNIP2*, *INSIG1*, *SLC11A1*, *IGFBP2*, *JAK2*, *PLSCR4*, *DDIT4*, *PLSCR3*, *GOT1*, *TIRAP*, *SPP1*, *CYP26B1* |
| GO:0042127 | Regulation of cell population proliferation | 374 | 1.92 | 3.95E-09 | 5.41E-08 | *CXCL9*, *CXCL11*, *CXCL10*, *IFNG*, *CD274*, *IRF1*, *CCL2*, *IL1B*, *IL27*, *STAT1*, *TNFSF13B*, *IL6*, *IL1A*, *PTGIR*, *CSF3*, *THBS1*, *TNFAIP3*, *JUNB*, *CTLA4*, *PIM1*, *IFI35*, *STAT2*, *CD40*, *IL15*, *LTF*, *CCL5*, *ZFP36*, *TIMP1*, *STAT5A*, *TRNP1*, *LRRC32*, *IFI30*, *CSF2*, *LIF*, *DUSP10*, *NFKBIA*, *STAT3*, *ATF5*, *ESM1*, *NR4A3*, *TFRC*, *IL10*, *APOD*, *GJA1*, *PHLDA2*, *NME1*, *BCL6*, *DLL4*, *HAVCR2*, *SLC7A1*, *CEBPB*, *MORC3*, *EGLN3*, *PRDM1*, *B4GALT1*, *ADM*, *CALR*, *ACKR3*, *TICAM1*, *SOX11*, *MYC*, *TNFSF9*, *RIPK3*, *RNASEH2B*, *TLR4*, *NKX3-1*, *TGFB3*, *FAM98A*, *KLF9*, |

| | | | | | | |
|---|---|---|---|---|---|---|
| | | | | | | *IL6R*, *CLDN5*, *SLC25A33*, *NOP2*, *IGFBP2*, *JAK2*, *FUT2*, *FBXO4*, *TIPIN*, *SMAD1*, *PPARGC1A*, *GTPBP4* |
| GO:0016477 | Cell migration | 339 | 1.90 | 5.54E-08 | 5.19E-07 | *CXCL9*, *CXCL11*, *CXCL10*, *MADCAM1*, *IFNG*, *SELE*, *CXCL13*, *AMCF-II*, *CCL2*, *CCL4*, *RASGEF1A*, *CXCL8*, *S100A12*, *S100A8*, *THBS1*, *TNFAIP3*, *CCL8*, *ACKR4*, *CD40*, *CCL5*, *FFAR2*, *CDH12*, *GDF15*, *CXCR6*, *ADRA2A*, *CCR1*, *DUSP10*, *TREM1*, *GLUL*, *NR4A3*, *TNFSF14*, *CXCL16*, *KIT*, *PPBP*, *APOD*, *GJA1*, *PHLDA2*, *ACKR2*, *RHOB*, *DLL4*, *B4GALT1*, *DDIT3*, *CASP8*, *CALR*, *ACKR3*, *KLRK1*, *ITGA5*, *TNFAIP1*, *SDC4*, *NARS1*, *RIPK3*, *TLR4* |
| GO:0010628 | Positive regulation of gene expression | 304 | 1.90 | 1.23E-07 | 1.02E-06 | *IFNG*, *RSAD2*, *ISG15*, *IRF1*, *CHI3L1*, *DHX58*, *CXCL8*, *IRF7*, *IL1B*, *STAT1*, *MMP8*, *IL1A*, *ATF3*, *THBS1*, *CD74*, *IRF8*, *CD40*, *IL15*, *FFAR2*, *ZFP36*, *NFIL3*, *STAT5A*, *TICAM2*, *CASP1*, *ADRA2A*, *LRRC32*, *LIF*, *EGR1*, *TFRC*, *IL10*, *KIT*, *ARID5A*, *GJA1*, *CEBPG*, *PANX2*, *DLL4*, *STING1*, *MAD2L2*, *TRAF2*, *HAVCR2*, *DDX21*, *CEBPB*, *CGAS*, *PRDM1*, *MYD88*, *DDIT3*, *CASP8*, *CALR*, *TICAM1*, *KLRK1*, *SOX11*, *IGF2BP1*, *POLR3D*, *SLAMF6*, *TLR2*, *TLR4*, *NKX3-1*, *FAM98A*, *CLDN5*, *TMEM106A*, *JAK2*, *WNT16*, *EIF5A*, *SMAD1*, *PPARGC1A*, *TMED10*, *TIRAP*, *SPRY2*, *CYP26B1*, *KAT2A*, *ZPR1*, *LY96*, *INHBA*, *VDR*, *RUNX1*, *HSP90AA1*, *UAP1*, *SLC38A2*, *RAB1A*, *NAT10*, *GATA2*, *TRA2B*, *FMR1*, *BLNK*, *BMP2* |
| GO:1901701 | Cellular response to oxygen-containing compound | 243 | 1.89 | 5.73E-07 | 3.85E-06 | *CXCL9*, *CXCL10*, *CXCL13*, *AMCF-II*, *CXCL8*, *IL1B*, *STAT1*, *MMP8*, *SOCS1*, *IL1A*, *TNFAIP3*, *PIM1*, *IRF8*, *CD180*, *LTF*, *FFAR2*, *ZFP36*, *STAT5A*, *GDF15*, *NFKBIB*, *ADRA2A*, *NFKBIA*, *STAT3*, *SESN2*, *NR4A3*, *SLC2A4*, *IL10*, *GPR37*, *PPBP*, *GJA1*, *RHOB*, *TRAF2*, *HAVCR2*, *GPR155*, *JAK3*, *CHRNA3*, *SELENOS*, *TICAM1*, *KLRK1*, *TLR2*, *RIPK3*, *TLR4*, *KLF9*, *SLC25A33*, *TNIP2*, *INSIG1*, *JAK2*, *PLSCR4*, *DDIT4*, *PLSCR3*, *GOT1*, *SPP1*, *CYP26B1* |
| GO:0040011 | Locomotion | 285 | 1.89 | 3.05E-07 | 2.26E-06 | *CXCL9*, *CXCL11*, *CXCL10*, *MADCAM1*, *IFNG*, *SELE*, *CXCL13*, *AMCF-II*, *CCL2*, *CCL4*, *CXCL8*, *S100A8*, *THBS1*, *CCL8*, *ACKR4*, *CD40*, *CCL5*, *FFAR2*, *BVES*, *GDF15*, *CXCR6*, *ADRA2A*, *CCR1*, *DUSP10*, *TREM1*, *GLUL*, *NR4A3*, *TNFSF14*, *CXCL16*, *KIT*, *GPR37*, *PPBP*, *APOD*, *GJA1*, *PHLDA2*, *ACKR2*, *RHOB*, *BCL6*, *DLL4*, *FBXO7*, *ARRDC3*, *CASP8*, *CALR*, *ACKR3*, *KLRK1*, *RIPK3*, *TLR4* |

| | | | | | | |
|---|---|---|---|---|---|---|
| GO:0008283 | Cell population proliferation | 444 | 1.87 | 1.75E-08 | 2.01E-07 | *CXCL9*, *CXCL11*, *CXCL10*, *IFNG*, *CD274*, *IRF1*, *CCL2*, *IL1B*, *IL27*, *STAT1*, *TNFSF13B*, *GNAT1*, *IL6*, *IL1A*, *PTGIR*, *CSF3*, *THBS1*, *TNFAIP3*, *JUNB*, *CTLA4*, *PIM1*, *CD180*, *IFI35*, *STAT2*, *CD40*, *IL15*, *LTF*, *CCL5*, *ZFP36*, *TIMP1*, *STAT5A*, *TRNP1*, *LRRC32*, *IFI30*, *CSF2*, *LIF*, *CTPS1*, *DUSP10*, *NFKBIA*, *STAT3*, *ATF5*, *ESM1*, *NR4A3*, *TFRC*, *IL10*, *APOD*, *DRAM2*, *GJA1*, *PHLDA2*, *NME1*, *BCL6*, *DLL4*, *HAVCR2*, *SLC7A1*, *SDCBP2*, *CEBPB*, *IL18RAP*, *MORC3*, *EGLN3*, *PRDM1*, *B4GALT1*, *DDIT3*, *ADM*, *CALR*, *ACKR3*, *TICAM1*, *SOX11*, *IGF2BP1*, *MYC*, *TNFSF9*, *EIF2S2*, *RIPK3*, *RNASEH2B*, *TLR4*, *ELF4*, *NKX3-1*, *TGFB3*, *FAM98A*, *KLF9*, *IL6R*, *CLDN5*, *SLC25A33*, *NOP2*, *IGFBP2*, *JAK2*, *FUT2*, *WNT16*, *FBXO4*, *TIPIN*, *ETV6*, *SMAD1*, *PPARGC1A*, *GTPBP4*, *ENPEP*, *LIG4*, *GLI2*, *SPRY2*, *PTGES3*, *KAT2A*, *SLC25A5*, *ZPR1* |
| GO:0002757 | Immune response-activating signaling pathway | 116 | 1.87 | 3.61E-05 | 1.57E-04 | *IFNG*, *RSAD2*, *IRF1*, *DHX58*, *IRF7*, *BPIFB1*, *FOSL1*, *TNFAIP3*, *TIFA*, *SLC15A3*, *IFI35*, *FFAR2*, *TICAM2*, *UNC93B1*, *NFKBIA*, *STING1*, *HAVCR2*, *CGAS*, *MYD88*, *TICAM1*, *KLRK1*, *PLSCR1*, *BMX*, *TLR2*, *TLR4*, *RNF115*, *TNIP2* |
| GO:0030099 | Myeloid cell differentiation | 128 | 1.86 | 6.37E-05 | 2.64E-04 | *IFNG*, *BATF2*, *ISG15*, *STAT1*, *SOCS1*, *CSF3*, *SNX10*, *JUNB*, *IRF8*, *LTF*, *ZFP36*, *STAT5A*, *CCR1*, *CSF2*, *LIF*, *NFKBIA*, *STAT3*, *SLC1A5*, *KIT*, *CASP3*, *CEBPG*, *BCL6*, *CEBPB*, *HSPA9*, *CASP8*, *MYC*, *TLR4*, *JAK2*, *LYAR*, *SLC25A5*, *INHBA*, *TFE3*, *RUNX1*, *PTK2B*, *KLF10*, *GATA2*, *RASSF2*, *BMP2*, *CHMP5*, *SH2B3*, *HSF1* |
| GO:0002253 | Activation of immune response | 145 | 1.86 | 5.34E-05 | 2.27E-04 | *IFNG*, *RSAD2*, *IRF1*, *DHX58*, *IRF7*, *BPIFB1*, *FOSL1*, *TNFAIP3*, *TIFA*, *C2*, *SLC15A3*, *IFI35*, *CFB*, *C4A*, *FFAR2*, *TICAM2*, *C3*, *UNC93B1*, *NFKBIA*, *SERPING1*, *STING1*, *HAVCR2*, *CGAS*, *MYD88*, *TICAM1*, *KLRK1*, *PLSCR1*, *BMX*, *TLR2*, *TLR4*, *C9*, *RNF115*, *TNIP2*, *CFI*, *TIRAP*, *CD59*, *TMEM126A*, *CD46*, *LY96*, *KCNK6*, *C7* |
| GO:0072594 | Establishment of protein localization to organelle | 116 | 1.85 | 5.58E-05 | 2.34E-04 | *NEURL3*, *IFNG*, *ISG15*, *TNFAIP3*, *NFKBIA*, *STAT3*, *ZFAND2A*, *APOD*, *TIMM8A*, *TXNIP*, *FBXO7*, *DDIT3*, *TOMM40*, *IPO4*, *GDAP1*, *SRPRA*, *SEC61A1*, *KPNA2*, *LOC100157391*, *SSR3*, *ZPR1*, *TMEM126A*, *CHCHD4*, *SPCS1*, *TRAM1* |
| GO:0051050 | Positive regulation of transport | 149 | 1.85 | 2.59E-05 | 1.15E-04 | *CXCL9*, *CXCL11*, *CXCL10*, *IFNG*, *CCL2*, *IL1B*, *S100A8*, *IL6*, *IL1A*, *LIPG*, *IL15*, *CCL5*, *FFAR2*, *C3*, *ADRA2A*, *CCR1*, *NFKBIA*, *NR4A3*, *PLK3*, *P2RY2*, *PDZK1*, *CEBPB*, *F2RL2*, *CALR*, *SOX11*, |

| | | | | | | |
|---|---|---|---|---|---|---|
| | | | | | | *TLR2*, *TLR4*, *NKX3-1*, *CADPS2*, *TGFB3*, *LYAR*, *SFTPD*, *TMED10*, *LRRC8A*, *SPP1*, *ATP1B1*, *ZPR1* |
| GO:0042981 | Regulation of apoptotic process | 344 | 1.83 | 6.01E-07 | 3.89E-06 | *CXCL10*, *IFNG*, *CD274*, *CCL2*, *S100A8*, *STAT1*, *FAS*, *LOC100522887*, *ATF3*, *THBS1*, *TNFAIP3*, *PIM1*, *FASLG*, *CASP10*, *CD40*, *BCL2L14*, *LTF*, *CCL5*, *ZFP36*, *NOD2*, *GADD45G*, *STAT5A*, *CASP1*, *HSH2D*, *ANGPTL4*, *EGR1*, *ATF5*, *BAK1*, *IL10*, *CASP3*, *PLK3*, *RNF122*, *PHLDA2*, *RHOB*, *BCL6*, *TRAF3*, *TIGAR*, *TRAF2*, *FBXO7*, *CEBPB*, *JAK3*, *CFLAR*, *B4GALT1*, *DDIT3*, *HSPA9*, *TCIM*, *BCL2A1*, *CASP8*, *YBX3*, *SELENOS*, *HIGD1A*, *ACKR3*, *MCL1*, *RIPK2*, *ITGA5*, *MYC*, *RIPK3*, *TLR4*, *ST3GAL1*, *PSMD10*, *NKX3-1*, *TNFRSF12A*, *GHITM*, *GADD45B*, *TNIP2*, *UTP11*, *JAK2*, *WNT16*, *PPARGC1A*, *LIG4*, *PLSCR3*, *GLI2*, *SPRY2*, *SLC25A5*, *ZPR1*, *G0S2*, *TP53*, *ARG2*, *INHBA*, *VDR*, *SLC35F6*, *F3*, *HSP90AA1*, *AQP1* |
| GO:0043065 | Positive regulation of apoptotic process | 130 | 1.82 | 3.55E-05 | 1.56E-04 | *IFNG*, *CD274*, *CCL2*, *S100A8*, *STAT1*, *ATF3*, *THBS1*, *FASLG*, *CD40*, *CCL5*, *GADD45G*, *IL10*, *CASP3*, *RNF122*, *PHLDA2*, *RHOB*, *BCL6*, *TIGAR*, *TRAF2*, *B4GALT1*, *DDIT3*, *HSPA9*, *BCL2A1*, *CASP8*, *MCL1*, *RIPK3*, *TLR4*, *NKX3-1*, *TNFRSF12A*, *GADD45B*, *UTP11*, *JAK2* |
| GO:0030155 | Regulation of cell adhesion | 171 | 1.82 | 6.63E-05 | 2.71E-04 | *IFNG*, *SELE*, *CD274*, *IRF1*, *RND1*, *CXCL8*, *IL1B*, *TNFSF13B*, *SOCS1*, *IL1A*, *THBS1*, *SLA-DQB1*, *SLA-DMB*, *IL15*, *CCL5*, *STAT5A*, *LRRC32*, *LIF*, *DUSP10*, *TNC*, *NR4A3*, *TFRC*, *TNFSF14*, *IL10*, *APOD*, *BCL6*, *MAD2L2*, *HAVCR2*, *SLC7A1*, *CEBPB*, *JAK3*, *CALR*, *TNFSF9* |
| GO:0070887 | Cellular response to chemical stimulus | 463 | 1.80 | 3.44E-08 | 3.31E-07 | *CXCL9*, *CXCL11*, *CXCL10*, *CXCL13*, *AMCF-II*, *CCL2*, *CCL4*, *CXCL8*, *IL1B*, *S100A8*, *STAT1*, *MMP8*, *SOCS1*, *IL1A*, *THBS1*, *TNFAIP3*, *AQP3*, *CCL8*, *JUNB*, *PIM1*, *IRF8*, *CD180*, *ACKR4*, *LTF*, *CCL5*, *FFAR2*, *ZFP36*, *MT-2B*, *STAT5A*, *GDF15*, *NFKBIB*, *CXCR6*, *ADRA2A*, *CCR1*, *NFKBIA*, *TREM1*, *STAT3*, *SESN2*, *NR4A3*, *SLC39A8*, *SLC2A4*, *TNFSF14*, *CXCL16*, *IL10*, *KIT*, *ZFAND2A*, *ARID5A*, *GPR37*, *PPBP*, *GJA1*, *ACKR2*, *RHOB*, *TIGAR*, *TRAF2*, *HAVCR2*, *GPR155*, *CEBPB*, *JAK3*, *CGAS*, *DDIT3*, *YBX3*, *CALR*, *CHRNA3*, *SELENOS*, *ACKR3*, *TICAM1*, *KLRK1*, *TLR2*, *RIPK3*, *TLR4*, *NKX3-1*, *SLC41A1*, *KLF9*, *SLC25A33*, *TMEM38B*, *TNIP2*, *INSIG1*, *JAK2*, *WNT16*, *ITPKC*, *PLSCR4*, *DDIT4*, *PPARGC1A*, *FBP1*, *PLSCR3*, *GOT1*, *PTGES3*, *EIF2S1*, *SPP1*, *CYP26B1* |

| | | | | | | |
|---|---|---|---|---|---|---|
| GO:0055085 | Transmembrane transport | 259 | 1.79 | 4.49E-06 | 2.39E-05 | *CXCL9*, *CXCL11*, *CXCL10*, *SLC25A47*, *CALHM6*, *THBS1*, *CNGA4*, *AQP3*, *SLC15A3*, *AQP7*, *KCNJ2*, *SLC25A25*, *GLRB*, *C3*, *SLC25A22*, *KCNMA1*, *ADRA2A*, *SLC38A5*, *BPIFA1*, *SESN2*, *NR4A3*, *SLC39A8*, *SLC2A4*, *SLC25A28*, *SLC1A5*, *GJA1*, *PANX2*, *PDZK1*, *ATP1A2*, *STING1*, *GPR155*, *SLC7A1*, *SLC5A6*, *DDIT3*, *TOMM40*, *SLC1A3*, *SLC22A4*, *SEC61A1*, *SLC31A1*, *GHITM*, *KCNA2*, *KCNJ14*, *SLC25A33*, *TMEM38B*, *SLC11A1*, *LOC100157391*, *SLC7A2*, *NALF2*, *SLC30A7*, *LRRC8A*, *ATP1B1*, *SLC25A5*, *CHCHD4*, *KCNK6*, *KCNK1*, *TRAM1*, *VDR*, *KCNK7*, *SLC5A1*, *AQP1*, *PTK2B*, *SLC4A4*, *SLC33A1*, *AQP9*, *ABCB7*, *ABCG2*, *SLC35B3*, *SLC30A4*, *ATP13A5*, *LRRC8B* |
| GO:0031399 | Regulation of protein modification process | 104 | 1.79 | 5.04E-04 | 1.71E-03 | *IFNG*, *ISG15*, *CHI3L1*, *TNFAIP3*, *STAT2*, *IL15*, *C3*, *LIF*, *EGR1*, *GNL3*, *TICAM1*, *ITGA5*, *PSMD10*, *JAK2*, *FBXO4*, *UBE2D1*, *GTPBP4*, *SPRY2*, *KAT2A*, *CRY1*, *NDFIP2*, *NRP1*, *SEMA4D*, *NCOA7* |
| GO:0043067 | Regulation of programmed cell death | 353 | 1.79 | 1.86E-06 | 1.10E-05 | *CXCL10*, *IFNG*, *CD274*, *CCL2*, *S100A8*, *STAT1*, *FAS*, *LOC100522887*, *ATF3*, *THBS1*, *TNFAIP3*, *PIM1*, *FASLG*, *CASP10*, *CD40*, *BCL2L14*, *LTF*, *CCL5*, *ZFP36*, *NOD2*, *GADD45G*, *STAT5A*, *CASP1*, *HSH2D*, *ANGPTL4*, *EGR1*, *ATF5*, *BAK1*, *IL10*, *CASP3*, *PLK3*, *FTH1*, *RNF122*, *PHLDA2*, *RHOB*, *BCL6*, *TRAF3*, *TIGAR*, *TRAF2*, *FBXO7*, *CEBPB*, *JAK3*, *CFLAR*, *B4GALT1*, *DDIT3*, *HSPA9*, *TCIM*, *BCL2A1*, *CASP8*, *YBX3*, *SELENOS*, *HIGD1A*, *ACKR3*, *MCL1*, *RIPK2*, *ITGA5*, *MYC*, *RIPK3*, *TLR4*, *ST3GAL1*, *PSMD10*, *NKX3-1*, *TNFRSF12A*, *GHITM*, *GADD45B*, *TNIP2*, *UTP11*, *JAK2*, *WNT16*, *PPARGC1A*, *LIG4*, *PLSCR3*, *GLI2*, *SPRY2*, *SLC25A5*, *ZPR1*, *G0S2*, *TP53*, *ARG2*, *INHBA*, *VDR*, *SLC35F6*, *F3*, *HSP90AA1*, *AQP1* |
| GO:0009057 | Macromolecule catabolic process | 375 | 1.78 | 4.53E-07 | 3.10E-06 | *ISG20*, *IFNG*, *ISG15*, *CHI3L1*, *PSMB9*, *PSMB8*, *PSME2*, *PSMB10*, *RNF19B*, *TNFAIP3*, *PSME1*, *ZFP36*, *TIMP1*, *ADRA2A*, *PSMA5*, *HSP90B1*, *PSMA2*, *IL10*, *ZFAND2A*, *ARID5A*, *PSMA3*, *PLK3*, *RNF122*, *MAD2L2*, *TRAF2*, *DNAJB9*, *FBXO7*, *POP1*, *DDIT3*, *DERL3*, *CASP8*, *CALR*, *SELENOS*, *IGF2BP1*, *PSMA1*, *TNFAIP1*, *RNASEH2B*, *PSMD10*, *PSMB3*, *RNF115*, *AGO2*, *PNPT1*, *KLHDC3*, *YOD1*, *PSMB4*, *PGM2*, *FBXO4*, *UBE2D1*, *PSMD3*, *ETF1*, *TIRAP*, *PAQR3*, *PELO*, *ZPR1*, *TMEM126A*, *PSMA4*, *REXO4*, *NDFIP2*, *USP44*, *OVGP1*, *IDE*, *TIMP3*, *UBE2D3*, *PTK2B*, *PDE12*, *RBM8A*, *RAD23A*, *FMR1*, *VCP*, *DESI1*, *DXO*, *PSMD8*, *SKIC2*, *RNF114*, |

| | | | | | | |
|---|---|---|---|---|---|---|
| | | | | | | *CHMP5*, *TBRG4*, *SQSTM1*, *STT3B*, *PSMF1*, *RLIM*, *HYAL2*, *PSMD1*, *ADAM10*, *NOS2*, *PLAA*, *DDX49*, *NPM1*, *RIDA*, *CHMP4A*, *EXOSC3*, *UCHL5*, *UBE2B*, *TRAF7*, *SIAH2*, *ATM*, *PSME3*, *EDEM2*, *PSMD2*, *PSMC1*, *PSMB1* |
| GO:0048870 | Cell motility | 387 | 1.77 | 9.32E-07 | 5.82E-06 | *CXCL9*, *CXCL11*, *CXCL10*, *MADCAM1*, *IFNG*, *SELE*, *CXCL13*, *AMCF-II*, *CCL2*, *CCL4*, *RASGEF1A*, *CXCL8*, *S100A12*, *S100A8*, *THBS1*, *TNFAIP3*, *CCL8*, *ACKR4*, *CD40*, *CCL5*, *FFAR2*, *CDH12*, *GDF15*, *CXCR6*, *ADRA2A*, *CCR1*, *DUSP10*, *TREM1*, *GLUL*, *NR4A3*, *TNFSF14*, *CXCL16*, *KIT*, *PPBP*, *APOD*, *GJA1*, *PHLDA2*, *ACKR2*, *RHOB*, *BCL6*, *DLL4*, *B4GALT1*, *DDIT3*, *CASP8*, *CALR*, *ACKR3*, *KLRK1*, *ITGA5*, *TNFAIP1*, *SDC4*, *NARS1*, *RIPK3*, *TLR4* |
| GO:0071705 | Nitrogen compound transport | 495 | 1.77 | 6.17E-08 | 5.49E-07 | *NEURL3*, *IFNG*, *RSAD2*, *TAP1*, *S100A8*, *CALHM6*, *GNAT1*, *IL6*, *IL1A*, *RHBDF2*, *SNX10*, *AQP3*, *STX11*, *CD74*, *SLC15A3*, *EGR2*, *CCL5*, *FFAR2*, *ZFP36*, *SLC25A22*, *ADRA2A*, *SLC38A5*, *NFKBIA*, *STAT3*, *TFRC*, *ARL5B*, *SLC1A5*, *RAB3D*, *PLK3*, *APOD*, *GJA1*, *TIMM8A*, *OPTN*, *PDZK1*, *RHOB*, *SEC23B*, *TXNIP*, *SLC7A1*, *ARRDC3*, *F2RL2*, *HSPA9*, *TOMM40*, *GPM6B*, *DERL3*, *STXBP1*, *CALR*, *SELENOS*, *IPO4*, *IGF2BP1*, *PREB*, *RAB39B*, *SLC1A3*, *SCFD2*, *TLR2*, *SRPRA*, *TLR4*, *SLC22A4*, *SEC61A1*, *CADPS2*, *KPNA2*, *TGFB3*, *SEC24A*, *ARF4*, *KCNA2*, *SLC25A33*, *INSIG1*, *LOC100157391*, *SLC7A2*, *USO1*, *AP3S2*, *XPO6*, *SELENBP1*, *TMED5*, *TMED10*, *SEC22B*, *RAB20*, *LRRC8A*, *NOL6*, *SLC25A5*, *ZPR1*, *TMED2*, *TP53*, *SYS1*, *SEC61B*, *CHCHD4*, *NDFIP2*, *KCNK1*, *ARCN1*, *TRAM1*, *IGF2BP2*, *EIF4E*, *AQP1*, *SLC38A2*, *C2CD2L*, *SLC18B1*, *MGST1*, *RAB1A*, *COPE*, *TVP23B*, *SLC33A1*, *TMED9*, *RBM8A*, *KPNB1*, *SNX19*, *FMR1*, *VCP*, *DESI1*, *ABCG2*, *XPO5*, *GOSR2*, *ATP13A5*, *COPB1*, *CHGA*, *SQSTM1*, *STX17*, *GDI2*, *HYAL2*, *NUP188*, *SAR1B*, *ARF6*, *NOS2*, *COPB2*, *SDAD1*, *MACIR*, *YIF1A*, *COPZ1*, *SEC63*, *C7H6orf125*, *MAGOHB* |
| GO:0051246 | Regulation of protein metabolic process | 316 | 1.76 | 4.49E-06 | 2.39E-05 | *IFNG*, *ISG15*, *CHI3L1*, *SERPINE1*, *PSME2*, *THBS1*, *TNFAIP3*, *PSME1*, *LIPG*, *STAT2*, *IL15*, *ZFP36*, *TIMP1*, *C3*, *ADRA2A*, *LIF*, *NFKBIA*, *EGR1*, *SESN2*, *IL10*, *CASP3*, *ZFAND2A*, *PLK3*, *APOD*, *SOAT1*, *RGS2*, *GNL3*, *MAD2L2*, *EIF4EBP1*, *CASP8*, *CALR*, *TICAM1*, *IGF2BP1*, *ITGA5*, *PSMD10*, *AGO2*, *JAK2*, *EIF5A*, *FBXO4*, *UBE2D1*, *GTPBP4*, *PSMD3*, *ETF1*, *TMED10*, *SPRY2*, |

| | | | | | | |
|---|---|---|---|---|---|---|
| | | | | | | *PAQR3*, *EIF2S1*, *KAT2A*, *PUM3*, *CRY1*, *NDFIP2*, *INHBA*, *IGF2BP2*, *TIMP3*, *PFDN6*, *PTK2B*, *NRP1*, *NAT10*, *SEMA4D*, *NCOA7*, *FMR1*, *DESI1*, *OGFOD1*, *SQSTM1*, *PPP1R15B*, *DDX3X*, *PSMD1*, *NOS2*, *PLAA*, *EIF1*, *ACO1*, *NPM1*, *RIDA*, *C7H6orf125*, *PIWIL4*, *BZW1*, *WBP1L*, *CD84*, *UBE2B*, *TRAF7*, *STK35*, *PSME3*, *PTCD3*, *ABCF1*, *PLGRKT* |
| GO:0006915 | Apoptotic process | 450 | 1.76 | 3.72E-07 | 2.70E-06 | *CXCL10*, *IFNG*, *CD274*, *IRF1*, *CHI3L1*, *CCL2*, *S100A8*, *STAT1*, *FAS*, *IL1A*, *LOC100522887*, *ATF3*, *THBS1*, *TNFAIP3*, *PIM1*, *FASLG*, *CASP10*, *CD40*, *BCL2L14*, *LTF*, *CCL5*, *ZFP36*, *NOD2*, *SHISA5*, *GADD45G*, *STAT5A*, *CASP1*, *HSH2D*, *LOC100624226*, *ANGPTL4*, *EGR1*, *AEN*, *ATF5*, *BAK1*, *IL10*, *CSRNP1*, *CASP3*, *PLK3*, *CYCS*, *RNF122*, *PHLDA2*, *RHOB*, *BCL6*, *TRAF3*, *TIGAR*, *TRAF2*, *FBXO7*, *CEBPB*, *JAK3*, *NEK6*, *EGLN3*, *CFLAR*, *B4GALT1*, *DDIT3*, *HSPA9*, *TCIM*, *BCL2A1*, *TNFRSF1A*, *CASP8*, *YBX3*, *SELENOS*, *HIGD1A*, *ACKR3*, *TICAM1*, *PLSCR1*, *MCL1*, *RIPK2*, *ITGA5*, *MYC*, *RIPK3*, *TLR4*, *ST3GAL1*, *PSMD10*, *NKX3-1*, *TNFRSF12A*, *GHITM*, *GADD45B*, *TNIP2*, *UTP11*, *JAK2*, *WNT16*, *ETV6*, *DDIT4*, *PPARGC1A*, *LIG4*, *PLSCR3*, *GLI2*, *SPRY2*, *SLC25A5*, *ZPR1*, *G0S2*, *TP53*, *ARG2*, *INHBA*, *VDR*, *SLC35F6*, *F3*, *XKR8*, *HSP90AA1*, *AQP1* |
| GO:0007186 | G protein-coupled receptor signaling pathway | 166 | 1.76 | 1.26E-04 | 4.91E-04 | *CXCL9*, *CXCL11*, *CXCL10*, *ACKR1*, *CXCL13*, *AMCF-II*, *CCL2*, *CCL4*, *CXCL8*, *GNAT1*, *PTGIR*, *CCL8*, *RGS16*, *CCL5*, *FFAR2*, *GLRB*, *C3*, *ADRA2A*, *PROKR2*, *GPR37*, *PPBP*, *P2RY2*, *RGS2*, *ARRDC3*, *ADM*, *HRH2*, *MRAP2* |
| GO:0098609 | Cell-cell adhesion | 182 | 1.75 | 8.74E-05 | 3.50E-04 | *MADCAM1*, *IFNG*, *SELE*, *CD274*, *IRF1*, *IL1B*, *S100A8*, *TNFSF13B*, *SOCS1*, *IL1A*, *CLSTN3*, *SLA-DQB1*, *VCAM1*, *SLA-DMB*, *IL15*, *CCL5*, *CDH12*, *STAT5A*, *BVES*, *LRRC32*, *DUSP10*, *NR4A3*, *TFRC*, *OLR1*, *SLC39A8*, *TNFSF14*, *IL10*, *KIT*, *BCL6*, *MAD2L2*, *HAVCR2*, *SLC7A1*, *CEBPB*, *JAK3*, *ICAM1*, *ITGA5*, *TNFSF9* |
| GO:0008219 | Cell death | 463 | 1.74 | 2.31E-07 | 1.79E-06 | *CXCL10*, *IFNG*, *CD274*, *IRF1*, *CHI3L1*, *CCL2*, *S100A8*, *STAT1*, *FAS*, *IL1A*, *LOC100522887*, *ATF3*, *THBS1*, *TNFAIP3*, *PIM1*, *FASLG*, *CASP10*, *CD40*, *BCL2L14*, *LTF*, *CCL5*, *ZFP36*, *NOD2*, *SHISA5*, *GADD45G*, *STAT5A*, *CASP1*, *GSDMA*, *HSH2D*, *LOC100624226*, *ANGPTL4*, *EGR1*, *AEN*, *ATF5*, *BAK1*, *MLKL*, *IL10*, *CSRNP1*, *CASP3*, *PLK3*, *FTH1*, *CYCS*, *RNF122*, *PHLDA2*, *RHOB*, *BCL6*, *TRAF3*, *TIGAR*, *TRAF2*, *FBXO7*, *CEBPB*, *JAK3*, *NEK6*, *EGLN3*, *CFLAR*, |

| | | | | | | |
|---|---|---|---|---|---|---|
| | | | | | | *B4GALT1*, *DDIT3*, *HSPA9*, *TCIM*, *BCL2A1*, *TNFRSF1A*, *CASP8*, *YBX3*, *SELENOS*, *HIGD1A*, *ACKR3*, *TICAM1*, *PLSCR1*, *MCL1*, *RIPK2*, *ITGA5*, *MYC*, *RIPK3*, *TLR4*, *ST3GAL1*, *PSMD10*, *NKX3-1*, *TNFRSF12A*, *GHITM*, *GADD45B*, *TNIP2*, *UTP11*, *JAK2*, *WNT16*, *ETV6*, *DDIT4*, *PPARGC1A*, *LIG4*, *PLSCR3*, *GLI2*, *SPRY2*, *CYP26B1*, *SLC25A5*, *ZPR1*, *G0S2*, *TP53*, *ARG2*, *INHBA*, *VDR*, *SLC35F6*, *F3*, *XKR8*, *HSP90AA1*, *AQP1* |
| GO:0012501 | Programmed cell death | 463 | 1.74 | 2.31E-07 | 1.79E-06 | *CXCL10*, *IFNG*, *CD274*, *IRF1*, *CHI3L1*, *CCL2*, *S100A8*, *STAT1*, *FAS*, *IL1A*, *LOC100522887*, *ATF3*, *THBS1*, *TNFAIP3*, *PIM1*, *FASLG*, *CASP10*, *CD40*, *BCL2L14*, *LTF*, *CCL5*, *ZFP36*, *NOD2*, *SHISA5*, *GADD45G*, *STAT5A*, *CASP1*, *GSDMA*, *HSH2D*, *LOC100624226*, *ANGPTL4*, *EGR1*, *AEN*, *ATF5*, *BAK1*, *MLKL*, *IL10*, *CSRNP1*, *CASP3*, *PLK3*, *FTH1*, *CYCS*, *RNF122*, *PHLDA2*, *RHOB*, *BCL6*, *TRAF3*, *TIGAR*, *TRAF2*, *FBXO7*, *CEBPB*, *JAK3*, *NEK6*, *EGLN3*, *CFLAR*, *B4GALT1*, *DDIT3*, *HSPA9*, *TCIM*, *BCL2A1*, *TNFRSF1A*, *CASP8*, *YBX3*, *SELENOS*, *HIGD1A*, *ACKR3*, *TICAM1*, *PLSCR1*, *MCL1*, *RIPK2*, *ITGA5*, *MYC*, *RIPK3*, *TLR4*, *ST3GAL1*, *PSMD10*, *NKX3-1*, *TNFRSF12A*, *GHITM*, *GADD45B*, *TNIP2*, *UTP11*, *JAK2*, *WNT16*, *ETV6*, *DDIT4*, *PPARGC1A*, *LIG4*, *PLSCR3*, *GLI2*, *SPRY2*, *CYP26B1*, *SLC25A5*, *ZPR1*, *G0S2*, *TP53*, *ARG2*, *INHBA*, *VDR*, *SLC35F6*, *F3*, *XKR8*, *HSP90AA1*, *AQP1* |
| GO:0043068 | Positive regulation of programmed cell death | 134 | 1.74 | 3.08E-04 | 1.13E-03 | *IFNG*, *CD274*, *CCL2*, *S100A8*, *STAT1*, *ATF3*, *THBS1*, *FASLG*, *CD40*, *CCL5*, *GADD45G*, *IL10*, *CASP3*, *RNF122*, *PHLDA2*, *RHOB*, *BCL6*, *TIGAR*, *TRAF2*, *B4GALT1*, *DDIT3*, *HSPA9*, *BCL2A1*, *CASP8*, *MCL1*, *RIPK3*, *TLR4*, *NKX3-1*, *TNFRSF12A*, *GADD45B*, *UTP11*, *JAK2* |
| GO:1902533 | Positive regulation of intracellular signal transduction | 271 | 1.74 | 3.71E-05 | 1.59E-04 | *SAA3*, *RSAD2*, *CHI3L1*, *DHX58*, *RASGEF1A*, *S100A12*, *S100A8*, *THBS1*, *TIFA*, *CD74*, *SLC15A3*, *PIM1*, *IFI35*, *FASLG*, *CD40*, *CCL5*, *GADD45G*, *TICAM2*, *GDF15*, *ADRA2A*, *CCR1*, *CSF2*, *LIF*, *SESN2*, *TNFSF14*, *GPR37*, *TRAF2*, *HAVCR2*, *GPR155*, *DDX21*, *ARRDC3*, *CFLAR*, *MYD88*, *DDIT3*, *TCIM*, *ADAM17*, *CASP8*, *CALR*, *ACKR3*, *TICAM1*, *SOX11*, *MCL1*, *TLR2*, *RIPK3*, *TLR4*, *NKX3-1*, *LPAR3*, *TMEM106A*, *GADD45B*, *TNIP2*, *IKBKB*, *JAK2*, *WNT16*, *RIPK1*, *TIRAP*, *SPRY2*, *GPR4*, *TP53*, *LTBR*, *INHBA*, *KCNK6*, *VDR*, *F3*, *TERF2IP*, *PTK2B*, *NRP1* |

| | | | | | | |
|---|---|---|---|---|---|---|
| GO:0009967 | Positive regulation of signal transduction | 357 | 1.73 | 5.04E-06 | 2.64E-05 | *SAA3*, *RSAD2*, *CHI3L1*, *DHX58*, *RASGEF1A*, *IRF7*, *ZBP1*, *S100A12*, *S100A8*, *MMP8*, *IL6*, *ATF3*, *THBS1*, *TNFAIP3*, *TIFA*, *CD74*, *SLC15A3*, *PIM1*, *CD180*, *IFI35*, *FASLG*, *CD40*, *CCL5*, *GADD45G*, *TICAM2*, *C3*, *GDF15*, *ADRA2A*, *CCR1*, *CSF2*, *LIF*, *STAT3*, *ESM1*, *SESN2*, *TNFSF14*, *IL10*, *KIT*, *GPR37*, *DLL4*, *TRAF2*, *HAVCR2*, *GPR155*, *DDX21*, *ARRDC3*, *CFLAR*, *MYD88*, *DDIT3*, *TCIM*, *ADAM17*, *CASP8*, *CALR*, *ACKR3*, *TICAM1*, *SOX11*, *MRAP2*, *MCL1*, *TLR2*, *RIPK3*, *TLR4*, *NKX3-1*, *LPAR3*, *TMEM106A*, *GADD45B*, *TNIP2*, *IKBKB*, *JAK2*, *WNT16*, *RIPK1*, *TIRAP*, *SPRY2*, *G0S2*, *GPR4*, *TP53*, *LTBR*, *LY96*, *INHBA*, *KCNK6*, *VDR*, *RALB*, *F3*, *NLE1*, *TERF2IP*, *PTK2B*, *NRP1* |
| GO:0023056 | Positive regulation of signaling | 394 | 1.73 | 3.14E-06 | 1.78E-05 | *SAA3*, *RSAD2*, *CHI3L1*, *CCL2*, *DHX58*, *RASGEF1A*, *IRF7*, *ZBP1*, *IL1B*, *S100A12*, *S100A8*, *MMP8*, *IL6*, *ATF3*, *THBS1*, *TNFAIP3*, *TIFA*, *CD74*, *SLC15A3*, *PIM1*, *CD180*, *IFI35*, *FASLG*, *CD40*, *CCL5*, *FFAR2*, *GADD45G*, *LRRTM2*, *TICAM2*, *C3*, *BVES*, *GDF15*, *ADRA2A*, *CCR1*, *CSF2*, *LIF*, *STAT3*, *ESM1*, *SESN2*, *TNFSF14*, *IL10*, *KIT*, *GPR37*, *DLL4*, *TRAF2*, *HAVCR2*, *GPR155*, *DDX21*, *ARRDC3*, *CFLAR*, *F2RL2*, *MYD88*, *DDIT3*, *TCIM*, *ADAM17*, *CASP8*, *CALR*, *ACKR3*, *TICAM1*, *SOX11*, *MRAP2*, *MCL1*, *TLR2*, *RIPK3*, *TLR4*, *NKX3-1*, *LPAR3*, *TMEM106A*, *GADD45B*, *TNIP2*, *IKBKB*, *JAK2*, *WNT16*, *ADORA1*, *RIPK1*, *GRIA3*, *TIRAP*, *SPRY2*, *LRRC8A*, *SPP1*, *G0S2*, *GPR4*, *TP53*, *LTBR*, *LY96*, *INHBA*, *KCNK6*, *VDR*, *RALB*, *F3*, *NLE1*, *C2CD2L*, *TERF2IP*, *PTK2B*, *NRP1* |
| GO:0030163 | Protein catabolic process | 266 | 1.72 | 2.05E-05 | 9.35E-05 | *IFNG*, *ISG15*, *PSMB9*, *PSMB8*, *PSME2*, *PSMB10*, *RNF19B*, *TNFAIP3*, *PSME1*, *TIMP1*, *ADRA2A*, *PSMA5*, *HSP90B1*, *PSMA2*, *IL10*, *ZFAND2A*, *PSMA3*, *PLK3*, *RNF122*, *MAD2L2*, *TRAF2*, *DNAJB9*, *FBXO7*, *DDIT3*, *DERL3*, *CASP8*, *CALR*, *SELENOS*, *PSMA1*, *TNFAIP1*, *PSMD10*, *PSMB3*, *RNF115*, *KLHDC3*, *YOD1*, *PSMB4*, *FBXO4*, *UBE2D1*, *PSMD3*, *PAQR3*, *TMEM126A*, *PSMA4*, *NDFIP2*, *USP44*, *IDE*, *TIMP3*, *UBE2D3*, *PTK2B*, *RAD23A*, *VCP*, *DESI1*, *PSMD8*, *RNF114*, *CHMP5*, *SQSTM1*, *STT3B*, *PSMF1*, *RLIM*, *PSMD1*, *ADAM10*, *NOS2*, *PLAA*, *CHMP4A*, *UCHL5*, *UBE2B*, *TRAF7*, *SIAH2*, *PSME3*, *EDEM2*, *PSMD2*, *PSMC1*, *PSMB1* |

| | | | | | | |
|---|---|---|---|---|---|---|
| GO:0010647 | Positive regulation of cell communication | 392 | 1.72 | 3.12E-06 | 1.78E-05 | *SAA3*, *RSAD2*, *CHI3L1*, *CCL2*, *DHX58*, *RASGEF1A*, *IRF7*, *ZBP1*, *IL1B*, *S100A12*, *S100A8*, *MMP8*, *IL6*, *ATF3*, *THBS1*, *TNFAIP3*, *TIFA*, *CD74*, *SLC15A3*, *PIM1*, *CD180*, *IFI35*, *FASLG*, *CD40*, *CCL5*, *FFAR2*, *GADD45G*, *LRRTM2*, *TICAM2*, *C3*, *GDF15*, *ADRA2A*, *CCR1*, *CSF2*, *LIF*, *STAT3*, *ESM1*, *SESN2*, *TNFSF14*, *IL10*, *KIT*, *GPR37*, *DLL4*, *TRAF2*, *HAVCR2*, *GPR155*, *DDX21*, *ARRDC3*, *CFLAR*, *F2RL2*, *MYD88*, *DDIT3*, *TCIM*, *ADAM17*, *CASP8*, *CALR*, *ACKR3*, *TICAM1*, *SOX11*, *MRAP2*, *MCL1*, *TLR2*, *RIPK3*, *TLR4*, *NKX3-1*, *LPAR3*, *TMEM106A*, *GADD45B*, *TNIP2*, *IKBKB*, *JAK2*, *WNT16*, *ADORA1*, *RIPK1*, *GRIA3*, *TIRAP*, *SPRY2*, *LRRC8A*, *SPP1*, *G0S2*, *GPR4*, *TP53*, *LTBR*, *LY96*, *INHBA*, *KCNK6*, *VDR*, *RALB*, *F3*, *NLE1*, *C2CD2L*, *TERF2IP*, *PTK2B*, *NRP1* |
| GO:0051240 | Positive regulation of multicellular organismal process | 368 | 1.72 | 7.12E-06 | 3.47E-05 | *IFNG*, *RSAD2*, *ISG15*, *IRF1*, *CHI3L1*, *DHX58*, *CXCL8*, *IRF7*, *IL1B*, *STAT1*, *MMP8*, *TNFSF13B*, *IL6*, *SOCS1*, *IL1A*, *THBS1*, *CD74*, *EGR2*, *SLA-DQB1*, *PIM1*, *IRF8*, *LIPG*, *SLA-DMB*, *CD40*, *IL15*, *LTF*, *CCL5*, *FFAR2*, *LRRTM2*, *STAT5A*, *TICAM2*, *CASP1*, *ADRA2A*, *CCR1*, *LIF*, *DUSP10*, *EGR1*, *TFRC*, *TNFSF14*, *TSKU*, *IL10*, *KIT*, *P2RY2*, *GJA1*, *RGS2*, *CEBPG*, *PANX2*, *RHOB*, *BCL6*, *STING1*, *MAD2L2*, *TRAF2*, *HAVCR2*, *SLC7A1*, *DDX21*, *CEBPB*, *CGAS*, *MYD88*, *DDIT3*, *ADM*, *CASP8*, *YBX3*, *TICAM1*, *HRH2*, *KLRK1*, *SOX11*, *POLR3D*, *ITGA5*, *SLAMF6*, *B4GALT5*, *TNFSF9*, *TLR2*, *TLR4*, *NKX3-1*, *TGFB3*, *LPAR3*, *TMEM106A*, *IGFBP2*, *JAK2* |
| GO:0006396 | RNA processing | 357 | 1.72 | 7.81E-06 | 3.76E-05 | *ISG20*, *NPM3*, *RNASEL*, *RRP9*, *POP1*, *QTRT1*, *WDR4*, *EMG1*, *QTRT2*, *FBL*, *AGO2*, *PAPOLA*, *EXOSC5*, *PNPT1*, *UTP11*, *NHP2*, *NOP2*, *RRS1*, *NOB1*, *LYAR*, *PPAN*, *GAR1*, *DKC1*, *GTPBP4*, *PUS3*, *WDR74*, *WDR75*, *CCNL1*, *SNRPD1*, *RTCA*, *NOL6*, *TRMT6*, *ZPR1*, *SNRPA1*, *REXO4*, *IMP3*, *SNRPF*, *RRP36*, *SMNDC1*, *RTCB*, *BUD23*, *RCL1*, *WDR12*, *SLC38A2*, *NOP9*, *TYW1*, *NAT10*, *UTP18*, *TRA2B*, *RBM8A*, *RRP15*, *RPL7L1*, *WDR46*, *FMR1*, *NSUN2*, *UTP23*, *RPRD1B*, *TBRG4*, *UTP15*, *PUS1*, *KRI1*, *TRMT11*, *ESF1*, *RBM14*, *DDX49*, *ABT1*, *SNRPG*, *MAGOHB*, *RPP40*, *PRPF40A*, *NOL9*, *EXOSC3*, *UTP6*, *AK6*, *ISG20L2*, *STRAP*, *SMN1*, *SNRPD3*, *TSR2*, *SNRPA*, *GEMIN6*, *SRSF1*, *WTAP*, *WDR36*, *TYW5*, *CDK12*, *RBM25*, *THUMPD1*, *HSD17B10*, *DDX52*, *SNRNP27*, *USP39*, *HNRNPF*, *CSTF2*, *SRSF7*, *RBM7*, *MTO1*, *AHCYL1*, *METTL3*, *TENT2*, *RAMAC*, *RNPS1*, *SMU1*, *SNRNP40*, *PWP1*, *EIF4A3*, *PRPF3*, *SF3B6*, *TRMT2A*, |

| | | | | | | |
|---|---|---|---|---|---|---|
| | | | | | | *MTREX*, *EXOSC4*, *ZMAT2*, *LARP6*, *CLNS1A*, *FCF1*, *RBBP6*, *MRPL44*, *NSUN4*, *LSM6*, *CSTF3*, *SNRPC*, *INTS8*, *RBM47*, *TRMT5*, *PCF11*, *SRSF2*, *RBM39*, *RPP21*, *ZBTB7A*, *SCAF11*, *SSB*, *ZRANB2*, *PRPF19*, *PTBP1*, *FTSJ1*, *SNRPB2*, *MRM1*, *CWC22*, *ELP5*, *TGS1*, *SYMPK*, *METTL8*, *WDR83*, *PAN2*, *RBM15*, *U2AF1L4*, *SPOUT1*, *GEMIN8*, *U2SURP*, *CDC40*, *NSRP1*, *KRR1*, *WDR55*, *SBDS*, *ELP2*, *TSEN15*, *SFSWAP*, *RPP30*, *CSTF2T*, *DDX39B*, *DHX8*, *PABPC1* |
| GO:0051603 | Proteolysis involved in protein catabolic process | 215 | 1.71 | 1.24E-04 | 4.91E-04 | *ISG15*, *PSMB9*, *PSMB8*, *PSME2*, *PSMB10*, *RNF19B*, *TNFAIP3*, *PSME1*, *PSMA5*, *HSP90B1*, *PSMA2*, *ZFAND2A*, *PSMA3*, *RNF122*, *DNAJB9*, *FBXO7*, *DDIT3*, *DERL3*, *CASP8*, *CALR*, *SELENOS*, *PSMA1*, *TNFAIP1*, *PSMD10*, *PSMB3*, *RNF115*, *KLHDC3*, *YOD1*, *PSMB4*, *FBXO4*, *UBE2D1*, *PSMD3*, *PAQR3*, *TMEM126A*, *PSMA4*, *NDFIP2*, *USP44*, *IDE*, *UBE2D3*, *PTK2B*, *RAD23A*, *VCP*, *DESI1*, *PSMD8*, *RNF114*, *CHMP5*, *STT3B*, *PSMF1*, *RLIM*, *PLAA*, *CHMP4A*, *UCHL5*, *UBE2B*, *TRAF7*, *SIAH2*, *PSME3*, *EDEM2*, *PSMC1*, *PSMB1* |
| GO:0034220 | Monoatomic ion transmembrane transport | 131 | 1.68 | 7.59E-04 | 2.35E-03 | *CXCL9*, *CXCL11*, *CXCL10*, *CALHM6*, *CNGA4*, *KCNJ2*, *GLRB*, *KCNMA1*, *ADRA2A*, *BPIFA1*, *SLC39A8*, *SLC25A28*, *PANX2*, *PDZK1*, *ATP1A2*, *STING1*, *DDIT3*, *SLC31A1*, *GHITM*, *KCNA2*, *KCNJ14*, *TMEM38B*, *SLC11A1*, *NALF2*, *SLC30A7*, *LRRC8A*, *ATP1B1*, *KCNK6*, *KCNK1*, *KCNK7*, *SLC5A1* |
| GO:0098771 | Inorganic ion homeostasis | 133 | 1.68 | 4.62E-04 | 1.60E-03 | *CXCL9*, *CXCL11*, *CXCL10*, *IL1A*, *SNX10*, *CCL8*, *CD40*, *CP*, *CCL5*, *MT-2B*, *STC2*, *CCR1*, *TFRC*, *SLC39A8*, *FTH1*, *ATP1A2*, *DDIT3*, *MYC*, *SLC41A1*, *SLC31A1*, *GHITM*, *TMEM38B* |
| GO:0048878 | Chemical homeostasis | 242 | 1.68 | 1.50E-04 | 5.80E-04 | *CXCL9*, *CXCL11*, *CXCL10*, *IL6*, *IL1A*, *LPL*, *SNX10*, *HK3*, *AQP3*, *CCL8*, *AQP7*, *LIPG*, *CD40*, *CP*, *CCL5*, *FFAR2*, *MT-2B*, *STC2*, *ADRA2A*, *CCR1*, *ANGPTL4*, *BPIFA1*, *STAT3*, *SESN2*, *TFRC*, *SLC39A8*, *TSKU*, *GCK*, *SOAT1*, *FTH1*, *ATP1A2*, *DDIT3*, *HK2*, *MYC*, *ORMDL1*, *SLC41A1*, *SLC31A1*, *GHITM*, *TMEM38B*, *INSIG1*, *ABHD5*, *SFTPD*, *PLSCR3*, *GOT1*, *SLC30A7*, *RAB20*, *LRRC8A*, *ATP1B1*, *CRY1*, *FITM2*, *NUS1*, *VDR* |
| GO:0030003 | Intracellular monoatomic cation homeostasis | 133 | 1.67 | 4.83E-04 | 1.65E-03 | *CXCL9*, *CXCL11*, *CXCL10*, *IL1A*, *CCL8*, *CD40*, *CP*, *CCL5*, *MT-2B*, *STC2*, *CCR1*, *TFRC*, *SLC39A8*, *FTH1*, *ATP1A2*, *DDIT3*, *MYC*, *SLC41A1*, *SLC31A1*, *GHITM*, *TMEM38B* |

| | | | | | | |
|---|---|---|---|---|---|---|
| GO:0042592 | Homeostatic process | 401 | 1.67 | 1.62E-05 | 7.58E-05 | *CXCL9*, *CXCL11*, *CXCL10*, *ISG15*, *IL1B*, *STAT1*, *TNFSF13B*, *IL6*, *FAS*, *IL1A*, *LPL*, *SNX10*, *TNFAIP3*, *HK3*, *AQP3*, *CCL8*, *AQP7*, *LIPG*, *CD40*, *CP*, *CCL5*, *FFAR2*, *ZFP36*, *MT-2B*, *STC2*, *STAT5A*, *GDF15*, *ADRA2A*, *CCR1*, *ANGPTL4*, *EGR1*, *BPIFA1*, *STAT3*, *SESN2*, *TFRC*, *SLC39A8*, *ST6GALNAC1*, *TSKU*, *SLC1A5*, *KIT*, *CASP3*, *GCK*, *DRAM2*, *SOAT1*, *GJA1*, *FTH1*, *CEBPG*, *BCL6*, *ATP1A2*, *CEBPB*, *ARRDC3*, *DDIT3*, *HSPA9*, *SELENOS*, *HK2*, *MYC*, *RIPK3*, *TLR4*, *ORMDL1*, *SLC41A1*, *SLC31A1*, *GHITM*, *CLDN5*, *TXNRD1*, *TMEM38B*, *INSIG1*, *ABHD5*, *JAK2*, *LYAR*, *FBXO4*, *SMAD1*, *PPARGC1A*, *SFTPD*, *PLSCR3*, *GOT1*, *SLC30A7*, *RAB20*, *LRRC8A*, *SPP1*, *ATP1B1*, *SLC25A5*, *G0S2*, *CRY1*, *FITM2*, *INHBA*, *NUS1*, *VDR*, *E2F4*, *IDE*, *AQP1* |
| GO:0006873 | Intracellular monoatomic ion homeostasis | 134 | 1.67 | 8.40E-04 | 2.54E-03 | *CXCL9*, *CXCL11*, *CXCL10*, *IL1A*, *CCL8*, *CD40*, *CP*, *CCL5*, *MT-2B*, *STC2*, *CCR1*, *TFRC*, *SLC39A8*, *FTH1*, *ATP1A2*, *DDIT3*, *MYC*, *SLC41A1*, *SLC31A1*, *GHITM*, *TMEM38B* |
| GO:1902531 | Regulation of intracellular signal transduction | 476 | 1.67 | 5.54E-06 | 2.82E-05 | *SAA3*, *RSAD2*, *CHI3L1*, *DHX58*, *RASGEF1A*, *S100A12*, *S100A8*, *STAT1*, *FAS*, *DUSP5*, *ATF3*, *THBS1*, *TNFAIP3*, *TIFA*, *CD74*, *SLC15A3*, *PIM1*, *IFI35*, *FASLG*, *CD40*, *CCL5*, *GADD45G*, *TICAM2*, *GDF15*, *NFKBIB*, *ADRA2A*, *CCR1*, *CSF2*, *LIF*, *DUSP10*, *NFKBIA*, *SESN2*, *SLC39A8*, *TNFSF14*, *IL10*, *GPR37*, *CHN2*, *OPTN*, *BCL6*, *TRAF3*, *TNIP1*, *TRAF2*, *HAVCR2*, *GPR155*, *DDX21*, *FBXO7*, *ARRDC3*, *CGAS*, *CFLAR*, *MYD88*, *DDIT3*, *TCIM*, *ADAM17*, *CASP8*, *YBX3*, *CALR*, *SELENOS*, *ACKR3*, *TICAM1*, *SOX11*, *MCL1*, *MYC*, *TNFAIP1*, *TLR2*, *RIPK3*, *RNASEH2B*, *TLR4*, *PSMD10*, *NKX3-1*, *FICD*, *LPAR3*, *TMEM106A*, *GADD45B*, *TNIP2*, *INSIG1*, *NOP2*, *IKBKB*, *JAK2*, *WNT16*, *RRS1*, *FBXO4*, *UBE2D1*, *DDIT4*, *CDC42SE1*, *FBP1*, *RIPK1*, *TIRAP*, *SPRY2*, *PAQR3*, *CYP26B1*, *GPR4*, *TP53*, *LTBR*, *CRY1*, *INHBA*, *KCNK6*, *USP44*, *VDR* |
| GO:0055082 | Intracellular chemical homeostasis | 167 | 1.64 | 8.38E-04 | 2.54E-03 | *CXCL9*, *CXCL11*, *CXCL10*, *IL1A*, *HK3*, *CCL8*, *CD40*, *CP*, *CCL5*, *MT-2B*, *STC2*, *ADRA2A*, *CCR1*, *TFRC*, *SLC39A8*, *GCK*, *FTH1*, *ATP1A2*, *DDIT3*, *HK2*, *MYC*, *ORMDL1*, *SLC41A1*, *SLC31A1*, *GHITM*, *TMEM38B*, *SLC30A7*, *RAB20*, *LRRC8A*, *ATP1B1*, *FITM2*, *VDR*, *AQP1*, *C2CD2L*, *PTK2B*, *SLC4A4*, *TRA2B*, *ATP6V0A2* |

| | | | | | | |
|---|---|---|---|---|---|---|
| GO:0009896 | Positive regulation of catabolic process | 128 | 1.64 | 2.03E-03 | 5.44E-03 | *SAA3*, *IFNG*, *TNFAIP3*, *ZFP36*, *ADRA2A*, *SESN2*, *ZFAND2A*, *PLK3*, *OPTN*, *STING1*, *FBXO7*, *TICAM1*, *IGF2BP1*, *PSMD10*, *AGO2*, *ABHD5*, *PAQR3*, *SLC25A5*, *RALB*, *PTK2B*, *PDE12* |
| GO:0033365 | Protein localization to organelle | 237 | 1.63 | 5.96E-04 | 1.96E-03 | *NEURL3*, *IFNG*, *ISG15*, *SNX10*, *TNFAIP3*, *NFKBIA*, *STAT3*, *SESN2*, *ARL5B*, *ZFAND2A*, *APOD*, *TIMM8A*, *OPTN*, *GNL3*, *TXNIP*, *FBXO7*, *MORC3*, *PARP3*, *DDIT3*, *TOMM40*, *CALR*, *IPO4*, *GDAP1*, *SRPRA*, *SEC61A1*, *KPNA2*, *ABHD17B*, *INSIG1*, *LOC100157391*, *RRS1*, *FBXO4*, *TMED10*, *PAQR3*, *SSR3*, *ZPR1*, *TMEM126A*, *CHCHD4*, *SPCS1*, *TRAM1*, *TERF2IP*, *NRP1*, *KPNB1*, *VCP*, *ARL1*, *SQSTM1*, *DDX3X*, *GDI2*, *HYAL2*, *ARF6*, *ADAM10*, *SRP14*, *SEC63*, *CHMP4A*, *ORMDL3*, *BRD2*, *EMC3*, *ARL2BP*, *SRP54*, *IPO13* |
| GO:0055080 | Monoatomic cation homeostasis | 147 | 1.63 | 1.46E-03 | 4.21E-03 | *CXCL9*, *CXCL11*, *CXCL10*, *IL1A*, *SNX10*, *CCL8*, *CD40*, *CP*, *CCL5*, *MT-2B*, *STC2*, *CCR1*, *TFRC*, *SLC39A8*, *FTH1*, *ATP1A2*, *DDIT3*, *MYC*, *SLC41A1*, *SLC31A1*, *GHITM*, *TMEM38B* |
| GO:0050801 | Monoatomic ion homeostasis | 148 | 1.63 | 1.37E-03 | 4.00E-03 | *CXCL9*, *CXCL11*, *CXCL10*, *IL1A*, *SNX10*, *CCL8*, *CD40*, *CP*, *CCL5*, *MT-2B*, *STC2*, *CCR1*, *TFRC*, *SLC39A8*, *FTH1*, *ATP1A2*, *DDIT3*, *MYC*, *SLC41A1*, *SLC31A1*, *GHITM*, *TMEM38B* |
| GO:0043412 | Macromolecule modification | 499 | 1.62 | 6.44E-06 | 3.19E-05 | *IFNG*, *ISG15*, *UBA7*, *CHI3L1*, *USP18*, *SOCS1*, *SOCS3*, *RNF19B*, *TNFAIP3*, *PARP12*, *GALNT15*, *EGR2*, *TRIM21*, *STAT2*, *IL15*, *C3*, *LIF*, *CISH*, *EGR1*, *DNAJB11*, *GLUL*, *PARP11*, *SLC39A8*, *ST6GALNAC1*, *RNF122*, *GNL3*, *TRAF2*, *FBXO7*, *NEK6*, *B4GALT1*, *PARP3*, *DERL3*, *QTRT1*, *WDR4*, *USP15*, *TICAM1*, *EMG1*, *QTRT2*, *ITGA5*, *B4GALT5*, *TNFAIP1*, *WSB2*, *ST3GAL1*, *PSMD10*, *FBL*, *RNF115*, *POMGNT2*, *MOGS*, *ALG3*, *NHP2*, *NOP2*, *JAK2*, *FUT2*, *FBXO4*, *GAR1*, *UBE2D1*, *OSTC*, *B3GNT5*, *DKC1*, *GTPBP4*, *PUS3*, *RPN1*, *RPN2*, *USP2*, *SPRY2*, *KAT2A*, *TRMT6*, *SOCS7*, *OTUB2*, *CRY1*, *NDFIP2*, *PADI4*, *NUS1*, *USP44*, *KLHL25*, *BUD23*, *MPDU1*, *UBE2D3*, *NRP1*, *EIPR1*, *NAT10*, *SEMA4D*, *NCOA7*, *GALNT1*, *RNF138*, *VCP*, *MGAT1*, *NSUN2*, *DESI1*, *RNF114*, *B3GALT6*, *B3GAT2*, *PPP2R2A*, *RXYLT1*, *SQSTM1*, *DDX3X*, *PUS1*, *NPC1*, *STT3B*, *RLIM*, *ST8SIA2*, *B4GALT3*, *MGAT2*, *A4GALT*, *NOS2*, *PUS7L*, *CTH*, *PLAA*, *MGAT4C*, *DPAGT1*, *DAD1*, *EXOSC3*, *WBP1L*, *UCHL5*, *DCAF11*, *UBE2B*, *TRAF7*, *SIAH2*, *ATM*, *FUT8*, *MAGT1*, *WTAP*, *EDEM2*, *PARP8*, *TYW5*, *PPP1CB*, *DCUN1D5*, *THUMPD1*, *HSD17B10*, *PHF23*, *USP39*, *HINT1*, *FOXO1*, *C1GALT1*, *TMEM258*, *PPP2CA*, *TPST1*, *MTO1*, *PLA2G1B*, *KLHL15*, *METTL3*, *DCUN1D3*, *TRIM27*, *SIAH1*, *ST8SIA4*, *RAMAC*, |

| | | | | | | |
|---|---|---|---|---|---|---|
| | | | | | | *OGT*, *FBXO33*, *OTUD6B*, *BCL10*, *EXOSC4*, *DR1*, *GAPDH*, *CUL5*, *MSL2*, *DDOST*, *RNF14*, *RBBP6*, *MINDY3*, *NSUN3*, *UBA3*, *TMEM129*, *NSUN4*, *CD80*, *UBE3A*, *UBA6*, *MDM2*, *TRMT5*, *MTOR*, *OTUB1*, *ABL1*, *PRKAR2A*, *RNF141*, *WWP1*, *B4GALT7*, *USP5*, *PRPF19*, *N4BP1*, *LOC100625207*, *NFE2L2*, *TOPORS*, *FTSJ1*, *MRM1*, *ASB5*, *KEAP1*, *DOLPP1*, *RNF5*, *UFC1*, *B3GALT1*, *ELP5*, *TGS1*, *DPM1*, *ST6GAL2*, *GGTA1* |
| GO:0097190 | Apoptotic signaling pathway | 153 | 1.62 | 2.07E-03 | 5.51E-03 | *IFNG*, *S100A8*, *IL1A*, *ATF3*, *THBS1*, *TNFAIP3*, *FASLG*, *AEN*, *IL10*, *TRAF2*, *FBXO7*, *CEBPB*, *CFLAR*, *DDIT3*, *BCL2A1*, *CASP8*, *YBX3*, *SELENOS*, *ACKR3*, *TICAM1*, *MCL1*, *MYC*, *RIPK3*, *TLR4*, *PSMD10*, *NKX3-1*, *GHITM*, *JAK2*, *WNT16*, *DDIT4*, *PLSCR3*, *SLC25A5*, *G0S2*, *TP53*, *INHBA*, *SLC35F6* |
| GO:0019725 | Cellular homeostasis | 199 | 1.61 | 1.37E-03 | 4.00E-03 | *CXCL9*, *CXCL11*, *CXCL10*, *IL1A*, *SNX10*, *HK3*, *CCL8*, *CD40*, *CP*, *CCL5*, *MT-2B*, *STC2*, *ADRA2A*, *CCR1*, *TFRC*, *SLC39A8*, *GCK*, *FTH1*, *ATP1A2*, *DDIT3*, *SELENOS*, *HK2*, *MYC*, *ORMDL1*, *SLC41A1*, *SLC31A1*, *GHITM*, *TXNRD1*, *TMEM38B*, *FBXO4*, *SLC30A7*, *RAB20*, *LRRC8A*, *ATP1B1*, *FITM2*, *VDR*, *E2F4*, *IDE*, *AQP1*, *C2CD2L*, *PTK2B*, *SLC4A4*, *TRA2B*, *ATP6V0A2* |
| GO:0034655 | Nucleobase-containing compound catabolic process | 148 | 1.61 | 1.74E-03 | 4.85E-03 | *ISG20*, *SAMHD1*, *HK3*, *ZFP36*, *ARID5A*, *CNP*, *GCK*, *TIGAR*, *TRAF2*, *POP1*, *HK2*, *IGF2BP1*, *RNASEH2B*, *AGO2*, *PGK1*, *PNPT1*, *DPYS*, *DDIT4*, *FBP1*, *ETF1*, *TIRAP*, *PELO*, *ZPR1*, *PDE4B*, *REXO4*, *ENTPD1*, *FITM2*, *DCTPP1*, *PDE12*, *RBM8A*, *NT5C*, *PDE4A*, *FMR1*, *DXO*, *SKIC2*, *TBRG4*, *DDX49*, *LOC100515984*, *NPM1*, *RIDA*, *EXOSC3*, *ATM*, *ITPA*, *PDE4D*, *HINT1*, *ZFP36L1*, *PPP2CA*, *ZFP36L2*, *NUDT7*, *TENT2*, *OGT*, *EIF4A3*, *TENT5A*, *PHAX*, *MTREX*, *CNOT9*, *EXOSC4*, *PGAM1*, *GAPDH*, *ALDOC*, *ELAVL1* |
| GO:0051247 | Positive regulation of protein metabolic process | 172 | 1.60 | 1.02E-03 | 3.03E-03 | *IFNG*, *CHI3L1*, *THBS1*, *TNFAIP3*, *IL15*, *C3*, *ADRA2A*, *LIF*, *NFKBIA*, *EGR1*, *CASP3*, *ZFAND2A*, *PLK3*, *SOAT1*, *GNL3*, *CASP8*, *TICAM1*, *IGF2BP1*, *ITGA5*, *PSMD10*, *JAK2*, *EIF5A*, *FBXO4*, *UBE2D1*, *PAQR3*, *CRY1*, *NDFIP2*, *INHBA*, *PTK2B*, *NRP1*, *NAT10*, *SEMA4D*, *FMR1*, *DDX3X*, *NPM1*, *C7H6orf125*, *WBP1L*, *TRAF7*, *ABCF1*, *PLGRKT*, *DCUN1D5*, *PHF23*, *FOXO1*, *PLA2G1B*, *METTL3*, *DCUN1D3*, *OGT*, *FBXO33*, *EIF4A3*, *BCL10*, *FGFR4*, *ELAVL1*, *CD80*, *UBE3A*, *GUF1*, *MDM2*, *MTOR*, *LPCAT1*, *ABL1*, *SSB*, *AGTR1*, *CCBE1* |

| | | | | | | |
|---|---|---|---|---|---|---|
| GO:0006886 | Intracellular protein transport | 189 | 1.60 | 1.58E-03 | 4.49E-03 | *IFNG*, *SNX10*, *STX11*, *CD74*, *EGR2*, *NFKBIA*, *STAT3*, *ARL5B*, *RAB3D*, *PLK3*, *APOD*, *TIMM8A*, *RHOB*, *SEC23B*, *TXNIP*, *HSPA9*, *DERL3*, *CALR*, *SELENOS*, *IPO4*, *SRPRA*, *SEC61A1*, *KPNA2*, *SEC24A*, *INSIG1*, *USO1*, *AP3S2*, *XPO6*, *TMED10*, *RAB20*, *ZPR1*, *SEC61B*, *TRAM1*, *RAB1A*, *TMED9*, *KPNB1*, *VCP*, *DESI1*, *XPO5*, *COPB1*, *SQSTM1*, *STX17*, *HYAL2*, *SAR1B*, *ARF6*, *COPB2*, *COPZ1*, *SEC63* |
| GO:0098655 | Monoatomic cation transmembrane transport | 121 | 1.59 | 4.74E-03 | 1.12E-02 | *CXCL9*, *CXCL11*, *CXCL10*, *CNGA4*, *KCNJ2*, *KCNMA1*, *ADRA2A*, *BPIFA1*, *SLC39A8*, *SLC25A28*, *PDZK1*, *ATP1A2*, *DDIT3*, *SLC31A1*, *GHITM*, *KCNA2*, *KCNJ14*, *TMEM38B*, *SLC11A1*, *NALF2*, *SLC30A7*, *ATP1B1*, *KCNK6*, *KCNK1*, *KCNK7*, *SLC5A1* |
| GO:0051241 | Negative regulation of multicellular organismal process | 230 | 1.59 | 6.37E-04 | 2.06E-03 | *CXCL10*, *IFNG*, *CD274*, *IRF1*, *DHX58*, *STAT1*, *SERPINE1*, *SOCS1*, *THBS1*, *TNFAIP3*, *LTF*, *ZFP36*, *GDF15*, *KCNMA1*, *LRRC32*, *CCR1*, *DUSP10*, *DNAJB11*, *TSKU*, *IL10*, *APOD*, *GJA1*, *RGS2*, *BCL6*, *MAD2L2*, *TMEM176B*, *HAVCR2*, *FBXO7*, *CEBPB*, *ARRDC3*, *MORC3*, *JAK3*, *PARP3*, *DDIT3*, *HSPA9*, *ADM*, *YBX3*, *CALR*, *SOX11*, *MYC*, *TLR4*, *ELF4*, *CREB3L1*, *NKX3-1*, *TGFB3*, *CLDN5* |
| GO:0006811 | Monoatomic ion transport | 220 | 1.59 | 7.02E-04 | 2.21E-03 | *CXCL9*, *CXCL11*, *CXCL10*, *CCL2*, *CALHM6*, *STEAP4*, *SLC39A14*, *CNGA4*, *CCL8*, *KCNJ2*, *LTF*, *CCL5*, *GLRB*, *KCNMA1*, *ADRA2A*, *CCR1*, *BPIFA1*, *TFRC*, *SLC39A8*, *SLC25A28*, *ATP1B3*, *FTH1*, *PANX2*, *PDZK1*, *ATP1A2*, *STING1*, *SLC5A6*, *DDIT3*, *SLC22A4*, *SLC11A2*, *SLC31A1*, *GHITM*, *KCNA2*, *KCNJ14*, *SLCO4C1*, *TMEM38B*, *SLC11A1*, *SLC39A7*, *NALF2*, *SLC30A7*, *LRRC8A*, *ATP1B1*, *SLC17A7*, *NDFIP2*, *KCNK6*, *KCNK1*, *VDR*, *KCNK7*, *SLC5A1*, *SLC10A7*, *SLC41A2*, *SLC38A2* |
| GO:0045184 | Establishment of protein localization | 452 | 1.59 | 8.63E-05 | 3.49E-04 | *NEURL3*, *IFNG*, *RSAD2*, *ISG15*, *IL6*, *IL1A*, *RHBDF2*, *SNX10*, *TNFAIP3*, *STX11*, *CD74*, *EGR2*, *CCL5*, *FFAR2*, *ADRA2A*, *LRRC32*, *NFKBIA*, *STAT3*, *TFRC*, *ARL5B*, *ZFAND2A*, *RAB3D*, *PLK3*, *APOD*, *TIMM8A*, *OPTN*, *RHOB*, *SEC23B*, *TXNIP*, *FBXO7*, *ARRDC3*, *F2RL2*, *DDIT3*, *HSPA9*, *TOMM40*, *DERL3*, *STXBP1*, *CALR*, *SELENOS*, *IPO4*, *PREB*, *RAB39B*, *SCFD2*, *GDAP1*, *TLR2*, *SRPRA*, *TLR4*, *SEC61A1*, *CADPS2*, *KPNA2*, *TGFB3*, *SEC24A*, *ARF4*, *INSIG1*, *LOC100157391*, *USO1*, *AP3S2*, *XPO6*, *SELENBP1*, *TMED5*, *TMED10*, *SEC22B*, *RAB20*, *SSR3*, *LRRC8A*, *ZPR1*, *TMED2*, *TMEM126A*, *SYS1*, *SEC61B*, *CHCHD4*, *NDFIP2*, *SPCS1*, *ARCN1*, *TRAM1*, *C2CD2L*, |

| | | | | | | |
|---|---|---|---|---|---|---|
| | | | | | | *RAB1A*, *COPE*, *TVP23B*, *TMED9*, *KPNB1*, *SNX19*, *VCP*, *DESI1*, *XPO5*, *GOSR2*, *COPB1*, *CHGA*, *SQSTM1*, *STX17*, *GDI2*, *HYAL2*, *NUP188*, *SAR1B*, *YWHAG*, *ARF6*, *NOS2*, *COPB2*, *SDAD1*, *MACIR*, *YIF1A*, *COPZ1*, *SRP14*, *SEC63*, *C7H6orf125*, *CHMP4A* |
| GO:0046903 | Secretion | 189 | 1.59 | 1.97E-03 | 5.32E-03 | *IFNG*, *RSAD2*, *IL1B*, *S100A8*, *GNAT1*, *IL6*, *IL1A*, *RHBDF2*, *SNX10*, *CCL8*, *STX11*, *CCL5*, *FFAR2*, *STAT5A*, *ADRA2A*, *CCR1*, *LIF*, *KIT*, *P2RY2*, *GJA1*, *NME1*, *F2RL2*, *HRH2*, *SOX11*, *TLR2*, *TLR4*, *NKX3-1*, *CADPS2*, *TGFB3*, *KCNA2*, *TMEM38B*, *ADORA1*, *TMED10*, *LRRC8A*, *SPP1*, *TP53*, *CRY1*, *INHBA*, *KCNK1*, *VDR*, *AQP1*, *C2CD2L*, *RAB1A*, *TVP23B* |
| GO:0019941 | Modification-dependent protein catabolic process | 173 | 1.58 | 2.10E-03 | 5.53E-03 | *ISG15*, *PSMB9*, *RNF19B*, *TNFAIP3*, *PSMA5*, *PSMA2*, *ZFAND2A*, *PSMA3*, *RNF122*, *FBXO7*, *DDIT3*, *PSMA1*, *TNFAIP1*, *PSMD10*, *PSMB3*, *RNF115*, *KLHDC3*, *FBXO4*, *UBE2D1*, *PSMD3*, *PAQR3*, *PSMA4*, *NDFIP2*, *USP44*, *UBE2D3*, *PTK2B*, *RAD23A*, *VCP*, *DESI1*, *PSMD8*, *RNF114*, *CHMP5*, *PSMF1*, *RLIM*, *PLAA*, *CHMP4A*, *UCHL5*, *UBE2B*, *TRAF7*, *SIAH2*, *PSMC1*, *PSMC2*, *ARK2N*, *KLHL15*, *TAF9*, *SIAH1*, *OGT*, *PSMD4*, *CUL5*, *RNF14*, *RBBP6*, *TMEM129*, *UBE3A*, *MDM2*, *CACUL1*, *WWP1*, *TMEM168*, *N4BP1*, *NFE2L2*, *TOPORS*, *KEAP1*, *RNF5*, *AKIRIN2*, *NEMF*, *FBXO9*, *ELOC*, *RAD23B*, *LTN1*, *SOCS5*, *KLHL2* |
| GO:0040017 | Positive regulation of locomotion | 143 | 1.58 | 3.51E-03 | 8.68E-03 | *CXCL10*, *MADCAM1*, *IFNG*, *CCL2*, *CCL4*, *CXCL8*, *THBS1*, *CCL8*, *CD40*, *CCL5*, *BVES*, *ADRA2A*, *CCR1*, *NR4A3*, *TNFSF14*, *CXCL16*, *KIT*, *RHOB*, *BCL6*, *CASP8*, *CALR* |
| GO:0051049 | Regulation of transport | 313 | 1.58 | 2.29E-04 | 8.59E-04 | *CXCL9*, *CXCL11*, *CXCL10*, *IFNG*, *RSAD2*, *CCL2*, *IL1B*, *S100A8*, *IL6*, *IL1A*, *RHBDF2*, *THBS1*, *LIPG*, *IL15*, *KCNJ2*, *CCL5*, *FFAR2*, *LRRTM2*, *C3*, *ADRA2A*, *CCR1*, *LIF*, *NFKBIA*, *BPIFA1*, *NR4A3*, *PLK3*, *P2RY2*, *APOD*, *GJA1*, *PDZK1*, *CEBPB*, *F2RL2*, *GPM6B*, *DERL3*, *CALR*, *SOX11*, *TLR2*, *TLR4*, *NKX3-1*, *CADPS2*, *TGFB3*, *KCNA2*, *KCNJ14*, *TMEM38B*, *INSIG1*, *LYAR*, *SFTPD*, *ADORA1*, *SIGLEC1*, *TMED10*, *RUBCN*, *LRRC8A*, *SPP1*, *ATP1B1*, *SLC25A5*, *ZPR1*, *TP53*, *CRY1*, *NDFIP2*, *INHBA*, *NUS1* |
| GO:0045597 | Positive regulation of cell differentiation | 170 | 1.58 | 1.86E-03 | 5.13E-03 | *CXCL9*, *IFNG*, *ISG15*, *STAT1*, *SOCS1*, *CSF3*, *JUNB*, *EGR2*, *PIM1*, *LTF*, *ZFP36*, *STAT5A*, *CCR1*, *CEBPD*, *LIF*, *DUSP10*, *STAT3*, *TNFSF14*, *IL10*, *GJA1*, *BCL6*, *CEBPB*, *ADM*, *CASP8*, *SOX11*, *TNFSF9*, *TGFB3*, *LPAR3*, *CLDN5*, *SMAD1*, *LIG4*, *GLI2*, *LRRC8A*, *CYP26B1*, *INHBA*, *VDR*, *RUNX1*, *S1PR2*, *NRP1*, *KLF10*, *GATA2*, *SEMA4D*, *FMR1*, *BMP2*, *RRAS* |

| | | | | | | |
|---|---|---|---|---|---|---|
| GO:0051094 | Positive regulation of developmental process | 269 | 1.58 | 6.08E-04 | 1.98E-03 | *CXCL9*, *IFNG*, *ISG15*, *CHI3L1*, *CXCL8*, *STAT1*, *TNFSF13B*, *SOCS1*, *IL1A*, *CSF3*, *THBS1*, *JUNB*, *EGR2*, *PIM1*, *CD40*, *LTF*, *ZFP36*, *LRRTM2*, *STAT5A*, *GDF15*, *CCR1*, *CEBPD*, *LIF*, *DUSP10*, *STAT3*, *TFRC*, *TNFSF14*, *IL10*, *GJA1*, *RHOB*, *BCL6*, *MAD2L2*, *CEBPB*, *ADM*, *CASP8*, *YBX3*, *SOX11*, *ITGA5*, *B4GALT5*, *TNFSF9*, *C13H21orf91*, *TGFB3*, *LPAR3*, *CLDN5*, *MIEF1* |
| GO:0051248 | Negative regulation of protein metabolic process | 103 | 1.58 | 8.59E-03 | 1.89E-02 | *ISG15*, *SERPINE1*, *THBS1*, *TNFAIP3*, *ZFP36*, *TIMP1*, *SESN2*, *IL10*, *APOD*, *MAD2L2*, *EIF4EBP1*, *CALR*, *IGF2BP1*, *AGO2*, *GTPBP4*, *SPRY2*, *EIF2S1*, *CRY1*, *TIMP3*, *PFDN6*, *NCOA7*, *SQSTM1*, *DDX3X*, *NOS2*, *PLAA*, *RIDA*, *CD84*, *UBE2B*, *PAIP2B*, *DCUN1D3*, *SRP9*, *TAF9*, *OGT*, *SERBP1*, *EIF4A3* |
| GO:0043632 | Modification-dependent macromolecule catabolic process | 174 | 1.58 | 3.21E-03 | 8.09E-03 | *ISG15*, *PSMB9*, *RNF19B*, *TNFAIP3*, *PSMA5*, *PSMA2*, *ZFAND2A*, *PSMA3*, *RNF122*, *FBXO7*, *DDIT3*, *PSMA1*, *TNFAIP1*, *PSMD10*, *PSMB3*, *RNF115*, *KLHDC3*, *FBXO4*, *UBE2D1*, *PSMD3*, *PAQR3*, *PSMA4*, *NDFIP2*, *USP44*, *UBE2D3*, *PTK2B*, *RAD23A*, *VCP*, *DESI1*, *PSMD8*, *RNF114*, *CHMP5*, *PSMF1*, *RLIM*, *PLAA*, *CHMP4A*, *UCHL5*, *UBE2B*, *TRAF7*, *SIAH2*, *PSMC1*, *PSMC2*, *ARK2N*, *KLHL15*, *TAF9*, *SIAH1*, *OGT*, *PSMD4*, *CUL5*, *RNF14*, *RBBP6*, *TMEM129*, *UBE3A*, *MDM2*, *CACUL1*, *WWP1*, *TMEM168*, *N4BP1*, *NFE2L2*, *TOPORS*, *KEAP1*, *RNF5*, *AKIRIN2*, *NEMF*, *FBXO9*, *ELOC*, *RAD23B*, *LTN1*, *SOCS5*, *KLHL2* |
| GO:0006812 | Monoatomic cation transport | 196 | 1.57 | 2.85E-03 | 7.34E-03 | *CXCL9*, *CXCL11*, *CXCL10*, *CCL2*, *STEAP4*, *SLC39A14*, *CNGA4*, *CCL8*, *KCNJ2*, *LTF*, *CCL5*, *KCNMA1*, *ADRA2A*, *CCR1*, *BPIFA1*, *TFRC*, *SLC39A8*, *SLC25A28*, *ATP1B3*, *FTH1*, *PANX2*, *PDZK1*, *ATP1A2*, *SLC5A6*, *DDIT3*, *SLC22A4*, *SLC11A2*, *SLC31A1*, *GHITM*, *KCNA2*, *KCNJ14*, *TMEM38B*, *SLC11A1*, *SLC39A7*, *NALF2*, *SLC30A7*, *ATP1B1*, *SLC17A7*, *NDFIP2*, *KCNK6*, *KCNK1*, *VDR*, *KCNK7*, *SLC5A1*, *SLC10A7*, *SLC41A2*, *SLC38A2*, *PTK2B*, *SLC4A4* |
| GO:2000026 | Regulation of multicellular organismal development | 306 | 1.57 | 5.81E-04 | 1.93E-03 | *CXCL10*, *IFNG*, *ISG15*, *IRF1*, *CHI3L1*, *CXCL8*, *IL27*, *STAT1*, *SERPINE1*, *SOCS1*, *IL1A*, *CSF3*, *THBS1*, *EGR2*, *PIM1*, *CD40*, *LTF*, *ZFP36*, *LRRTM2*, *STAT5A*, *MAFF*, *CCR1*, *LIF*, *DUSP10*, *NFKBIA*, *DNAJB11*, *STAT3*, *GLUL*, *IL10*, *GJA1*, *VASH2*, *PHLDA2*, *RHOB*, *BCL6*, *DLL4*, *TMEM176B*, *FBXO7*, *CEBPB*, *JAK3*, *PRDM1*, *HSPA9*, *ADM*, *TCIM*, *CASP8*, *SOX11*, *ITGA5*, *B4GALT5*, *MYC*, *TNFSF9*, *CREB3L1*, *LPAR3*, *CLDN5*, *IKBKB*, *SMAD1*, *LIG4*, *GLI2*, *SPRY2*, |

| | | | | | | |
|---|---|---|---|---|---|---|
| | | | | | | *CYP26B1*, *KAT2A*, *ZPR1*, *GPR4*, *TP53*, *INHBA*, *VDR*, *TFE3*, *F3*, *RUNX1*, *AQP1*, *S1PR2*, *PTK2B*, *NRP1*, *KLF10*, *GATA2*, *SEMA4D*, *RASSF2*, *FMR1*, *HTATIP2*, *BMP2*, *RRAS* |
| GO:0008285 | Negative regulation of cell population proliferation | 156 | 1.57 | 2.53E-03 | 6.61E-03 | *IFNG*, *CD274*, *CCL2*, *STAT1*, *IL1A*, *PTGIR*, *THBS1*, *TNFAIP3*, *IFI35*, *LRRC32*, *IFI30*, *LIF*, *DUSP10*, *ATF5*, *IL10*, *APOD*, *BCL6*, *DLL4*, *HAVCR2*, *CEBPB*, *MORC3*, *B4GALT1*, *ACKR3*, *SOX11*, *NKX3-1*, *TGFB3*, *KLF9*, *JAK2*, *FBXO4*, *SMAD1*, *PPARGC1A*, *GTPBP4* |
| GO:0098660 | Inorganic ion transmembrane transport | 132 | 1.56 | 3.73E-03 | 9.04E-03 | *CXCL9*, *CXCL11*, *CXCL10*, *KCNJ2*, *GLRB*, *KCNMA1*, *ADRA2A*, *BPIFA1*, *SLC39A8*, *SLC25A28*, *ATP1A2*, *SLC5A6*, *DDIT3*, *SLC31A1*, *GHITM*, *KCNA2*, *KCNJ14*, *TMEM38B*, *SLC11A1*, *NALF2*, *SLC30A7*, *LRRC8A*, *ATP1B1*, *KCNK6*, *KCNK1*, *VDR*, *KCNK7*, *SLC5A1*, *AQP1* |
| GO:0061061 | Muscle structure development | 141 | 1.56 | 3.66E-03 | 8.94E-03 | *CXCL9*, *CXCL10*, *MYBPH*, *ATF3*, *EGR2*, *MSC*, *UNC45B*, *BVES*, *GDF15*, *MAFF*, *LIF*, *EGR1*, *TNFSF14*, *GJA1*, *DLL4*, *DDIT3*, *ADM*, *YBX3*, *SOX11* |
| GO:0030001 | Metal ion transport | 182 | 1.56 | 1.72E-03 | 4.85E-03 | *CXCL9*, *CXCL11*, *CXCL10*, *CCL2*, *STEAP4*, *SLC39A14*, *CCL8*, *KCNJ2*, *LTF*, *CCL5*, *KCNMA1*, *ADRA2A*, *CCR1*, *BPIFA1*, *TFRC*, *SLC39A8*, *SLC25A28*, *ATP1B3*, *FTH1*, *ATP1A2*, *SLC5A6*, *DDIT3*, *SLC22A4*, *SLC11A2*, *SLC31A1*, *GHITM*, *KCNA2*, *KCNJ14*, *TMEM38B*, *SLC11A1*, *SLC39A7*, *NALF2*, *SLC30A7*, *ATP1B1*, *SLC17A7*, *NDFIP2*, *KCNK6*, *KCNK1*, *VDR*, *KCNK7*, *SLC5A1*, *SLC10A7*, *SLC41A2*, *SLC38A2*, *PTK2B*, *SLC4A4* |
| GO:0051046 | Regulation of secretion | 116 | 1.56 | 4.49E-03 | 1.07E-02 | *IFNG*, *RSAD2*, *IL1B*, *S100A8*, *IL6*, *IL1A*, *RHBDF2*, *CCL5*, *FFAR2*, *ADRA2A*, *LIF*, *P2RY2*, *GJA1*, *F2RL2*, *SOX11*, *TLR2*, *TLR4*, *NKX3-1*, *CADPS2*, *TGFB3*, *KCNA2*, *ADORA1*, *TMED10*, *LRRC8A*, *SPP1*, *TP53*, *CRY1*, *INHBA*, *AQP1*, *C2CD2L* |
| GO:0036211 | Protein modification process | 430 | 1.56 | 1.64E-04 | 6.27E-04 | *IFNG*, *ISG15*, *UBA7*, *CHI3L1*, *USP18*, *SOCS1*, *SOCS3*, *RNF19B*, *TNFAIP3*, *PARP12*, *GALNT15*, *EGR2*, *TRIM21*, *STAT2*, *IL15*, *C3*, *LIF*, *CISH*, *EGR1*, *GLUL*, *PARP11*, *SLC39A8*, *ST6GALNAC1*, *RNF122*, *GNL3*, *TRAF2*, *FBXO7*, *NEK6*, *B4GALT1*, *PARP3*, *DERL3*, *USP15*, *TICAM1*, *ITGA5*, *B4GALT5*, *TNFAIP1*, *WSB2*, *ST3GAL1*, *PSMD10*, *RNF115*, *POMGNT2*, *MOGS*, *ALG3*, *JAK2*, *FUT2*, *FBXO4*, *UBE2D1*, *OSTC*, *B3GNT5*, *GTPBP4*, *RPN1*, *RPN2*, *USP2*, *SPRY2*, *KAT2A*, *SOCS7*, *OTUB2*, *CRY1*, *NDFIP2*, *PADI4*, *NUS1*, *USP44*, *KLHL25*, *MPDU1*, *UBE2D3*, *NRP1*, *EIPR1*, *SEMA4D*, *NCOA7*, *GALNT1*, *RNF138*, *VCP*, *MGAT1*, *DESI1*, *RNF114*, |

| | | | | | | |
|---|---|---|---|---|---|---|
| | | | | | | *B3GALT6*, *B3GAT2*, *PPP2R2A*, *RXYLT1*, *SQSTM1*, *DDX3X*, *NPC1*, *STT3B*, *RLIM*, *ST8SIA2*, *B4GALT3*, *MGAT2*, *A4GALT*, *NOS2*, *CTH*, *PLAA*, *MGAT4C*, *DPAGT1*, *DAD1*, *WBP1L*, *UCHL5*, *DCAF11*, *UBE2B*, *TRAF7*, *SIAH2*, *ATM*, *FUT8*, *MAGT1*, *EDEM2*, *PARP8*, *PPP1CB*, *DCUN1D5*, *PHF23*, *USP39*, *HINT1*, *FOXO1*, *C1GALT1*, *TMEM258*, *PPP2CA*, *TPST1*, *PLA2G1B*, *KLHL15*, *DCUN1D3*, *TRIM27*, *SIAH1*, *ST8SIA4*, *OGT*, *FBXO33*, *OTUD6B*, *BCL10*, *DR1*, *GAPDH*, *CUL5*, *MSL2*, *DDOST*, *RNF14*, *RBBP6*, *MINDY3*, *UBA3*, *TMEM129*, *CD80*, *UBE3A*, *UBA6*, *MDM2*, *MTOR*, *OTUB1*, *ABL1*, *PRKAR2A*, *RNF141*, *WWP1*, *B4GALT7*, *USP5*, *PRPF19*, *N4BP1*, *LOC100625207*, *NFE2L2*, *TOPORS*, *ASB5*, *KEAP1*, *DOLPP1*, *RNF5*, *UFC1*, *B3GALT1* |
| GO:0015031 | Protein transport | 395 | 1.55 | 3.82E-04 | 1.37E-03 | *NEURL3*, *IFNG*, *RSAD2*, *IL6*, *IL1A*, *RHBDF2*, *SNX10*, *STX11*, *CD74*, *EGR2*, *CCL5*, *FFAR2*, *ADRA2A*, *NFKBIA*, *STAT3*, *TFRC*, *ARL5B*, *RAB3D*, *PLK3*, *APOD*, *TIMM8A*, *OPTN*, *RHOB*, *SEC23B*, *TXNIP*, *ARRDC3*, *F2RL2*, *HSPA9*, *TOMM40*, *DERL3*, *STXBP1*, *CALR*, *SELENOS*, *IPO4*, *PREB*, *RAB39B*, *SCFD2*, *TLR2*, *SRPRA*, *TLR4*, *SEC61A1*, *CADPS2*, *KPNA2*, *TGFB3*, *SEC24A*, *ARF4*, *INSIG1*, *LOC100157391*, *USO1*, *AP3S2*, *XPO6*, *SELENBP1*, *TMED5*, *TMED10*, *SEC22B*, *RAB20*, *LRRC8A*, *ZPR1*, *TMED2*, *SYS1*, *SEC61B*, *CHCHD4*, *NDFIP2*, *ARCN1*, *TRAM1*, *C2CD2L*, *RAB1A*, *COPE*, *TVP23B*, *TMED9*, *KPNB1*, *SNX19*, *VCP*, *DESI1*, *XPO5*, *GOSR2*, *COPB1*, *CHGA*, *SQSTM1*, *STX17*, *GDI2*, *HYAL2*, *NUP188*, *SAR1B*, *ARF6*, *NOS2*, *COPB2*, *SDAD1*, *MACIR*, *YIF1A*, *COPZ1*, *SEC63*, *C7H6orf125* |
| GO:0010629 | Negative regulation of gene expression | 279 | 1.55 | 8.09E-04 | 2.48E-03 | *SAA3*, *IFNG*, *DHX58*, *CXCL8*, *SERPINE1*, *THBS1*, *TNFAIP3*, *LTF*, *ZFP36*, *MOV10*, *LRRC32*, *CCR1*, *SESN2*, *TSKU*, *IL10*, *ARID5A*, *APOD*, *GJA1*, *BCL6*, *DLL4*, *TRAF2*, *BAZ1A*, *HAVCR2*, *MORC3*, *PRDM1*, *PARP3*, *DDIT3*, *EIF4EBP1*, *CALR*, *SOX11*, *IGF2BP1*, *RNASEH2B*, *TLR4*, *ELF4*, *CREB3L1*, *NKX3-1*, *AGO2*, *TGFB3*, *PNPT1*, *CLDN5*, *ETF1*, *TIRAP*, *PELO*, *EIF2S1*, *ARG2*, *LOC100624590*, *NDFIP2*, *PDE12*, *GATA2*, *RBM8A*, *FMR1*, *BMP2*, *DXO*, *PDCD4*, *CMKLR1*, *TBRG4*, *SETDB2*, *DDX3X*, *RLIM*, *ADAM10*, *NOS2*, *EPHA2*, *MACIR*, *NPM1*, *RIDA*, *IL27RA*, *PIWIL4*, *EXOSC3*, *CD84* |

| | | | | | | |
|---|---|---|---|---|---|---|
| GO:0045595 | Regulation of cell differentiation | 312 | 1.55 | 7.23E-04 | 2.26E-03 | *CXCL9*, *CXCL10*, *IFNG*, *ISG15*, *IRF1*, *IL27*, *STAT1*, *SOCS1*, *CSF3*, *JUNB*, *EGR2*, *PIM1*, *LTF*, *ZFP36*, *STAT5A*, *MAFF*, *CCR1*, *CEBPD*, *LIF*, *DUSP10*, *NFKBIA*, *DNAJB11*, *STAT3*, *TNFSF14*, *TSKU*, *IL10*, *GJA1*, *BCL6*, *DLL4*, *MAD2L2*, *TMEM176B*, *FBXO7*, *CEBPB*, *JAK3*, *PRDM1*, *DDIT3*, *HSPA9*, *ADM*, *TCIM*, *CASP8*, *CALR*, *SOX11*, *MYC*, *TNFSF9*, *TGFB3*, *LPAR3*, *CLDN5*, *INSIG1*, *IKBKB*, *SMAD1*, *LIG4*, *GLI2*, *SPRY2*, *LRRC8A*, *CYP26B1*, *KAT2A*, *INHBA*, *VDR*, *TFE3*, *RUNX1*, *S1PR2*, *PTK2B*, *NRP1*, *TWIST2*, *KLF10*, *GATA2*, *SEMA4D*, *RASSF2*, *FMR1*, *BMP2*, *RBPMS2*, *RRAS*, *PDCD4* |
| GO:0030335 | Positive regulation of cell migration | 133 | 1.54 | 3.58E-03 | 8.79E-03 | *CXCL10*, *MADCAM1*, *IFNG*, *CCL2*, *CCL4*, *THBS1*, *CCL8*, *CD40*, *CCL5*, *ADRA2A*, *CCR1*, *NR4A3*, *TNFSF14*, *CXCL16*, *KIT*, *RHOB*, *CASP8*, *CALR* |
| GO:1902532 | Negative regulation of intracellular signal transduction | 170 | 1.54 | 3.15E-03 | 8.02E-03 | *DHX58*, *STAT1*, *DUSP5*, *ATF3*, *THBS1*, *TNFAIP3*, *IFI35*, *GDF15*, *LIF*, *DUSP10*, *NFKBIA*, *SESN2*, *SLC39A8*, *IL10*, *OPTN*, *BCL6*, *FBXO7*, *CGAS*, *DDIT3*, *CASP8*, *YBX3*, *CALR*, *SELENOS*, *ACKR3*, *MYC*, *TNFAIP1*, *TLR4*, *PSMD10*, *INSIG1*, *UBE2D1*, *DDIT4*, *FBP1*, *SPRY2*, *PAQR3*, *CYP26B1* |
| GO:0098662 | Inorganic cation transmembrane transport | 122 | 1.53 | 9.18E-03 | 2.00E-02 | *CXCL9*, *CXCL11*, *CXCL10*, *KCNJ2*, *KCNMA1*, *ADRA2A*, *BPIFA1*, *SLC39A8*, *SLC25A28*, *ATP1A2*, *DDIT3*, *SLC31A1*, *GHITM*, *KCNA2*, *KCNJ14*, *TMEM38B*, *SLC11A1*, *NALF2*, *SLC30A7*, *ATP1B1*, *KCNK6*, *KCNK1*, *KCNK7*, *SLC5A1*, *AQP1* |
| GO:0032879 | Regulation of localization | 413 | 1.53 | 2.73E-04 | 1.01E-03 | *CXCL9*, *CXCL11*, *CXCL10*, *IFNG*, *RSAD2*, *CCL2*, *IL1B*, *S100A8*, *IL6*, *IL1A*, *RHBDF2*, *THBS1*, *LIPG*, *IL15*, *KCNJ2*, *CCL5*, *FFAR2*, *ZFP36*, *LRRTM2*, *C3*, *SLC5A3*, *ADRA2A*, *CCR1*, *LIF*, *NFKBIA*, *BPIFA1*, *SESN2*, *NR4A3*, *PLIN2*, *PLK3*, *P2RY2*, *APOD*, *GJA1*, *GNL3*, *PDZK1*, *CEBPB*, *F2RL2*, *DDIT3*, *GPM6B*, *DERL3*, *CALR*, *SOX11*, *MRAP2*, *TLR2*, *TLR4*, *NKX3-1*, *CADPS2*, *TGFB3*, *ABHD17B*, *GHITM*, *KCNA2*, *KCNJ14*, *TMEM38B*, *INSIG1*, *ABHD5*, *LYAR*, *FBXO4*, *SFTPD*, *ADORA1*, *SIGLEC1*, *TMED10*, *RUBCN*, *LRRC8A*, *SPP1*, *ATP1B1*, *SLC25A5*, *ZPR1*, *TP53*, *CRY1*, *NDFIP2*, *FITM2*, *INHBA*, *NUS1* |
| GO:0006508 | Proteolysis | 432 | 1.53 | 5.80E-04 | 1.93E-03 | *IFNG*, *ISG15*, *PSMB9*, *USP18*, *MMP8*, *SERPINE1*, *MMP12*, *PSMB8*, *PSME2*, *PSMB10*, *CTSL*, *LOC100522887*, *RNF19B*, *THBS1*, *TNFAIP3*, *MMP1*, *C2*, *PSME1*, *CFB*, *CASP10*, *LTF*, *TIMP1*, *CASP1*, *MMP9*, *ADRA2A*, *MMP27*, *PSMA5*, *HSP90B1*, *DPP3*, *PSMA2*, *IL10*, *CASP3*, |

| | | | | | |
|---|---|---|---|---|---|
| | | | | | *ZFAND2A*, *CNDP2*, *PSMA3*, *RNF122*, *ASRGL1*, *DNAJB9*, *FBXO7*, *GZMA*, *CFLAR*, *DDIT3*, *DERL3*, *ADAM17*, *CASP8*, *CALR*, *SELENOS*, *USP15*, *PSMA1*, *TNFAIP1*, *PSMD10*, *PSMB3*, *RNF115*, *MMP13*, *KLHDC3*, *CFI*, *YOD1*, *PSMB4*, *FBXO4*, *SPCS3*, *UBE2D1*, *PISD*, *ENPEP*, *PSMD3*, *RHBDL2*, *USP2*, *PAQR3*, *ADAMTS9*, *TMEM126A*, *PSMA4*, *DNPEP*, *SEC11C*, *CD46*, *OTUB2*, *AEBP1*, *TMPRSS4*, *NDFIP2*, *SPCS1*, *USP44*, *F3*, *IDE*, *TIMP3*, *UBE2D3* |
| GO:0060284 | Regulation of cell development | 177 | 1.52 | 2.77E-03 | 7.19E-03 | *IFNG*, *ISG15*, *IRF1*, *IL27*, *STAT1*, *SOCS1*, *CSF3*, *EGR2*, *LTF*, *ZFP36*, *STAT5A*, *CCR1*, *LIF*, *DUSP10*, *NFKBIA*, *DNAJB11*, *STAT3*, *IL10*, *BCL6*, *DLL4*, *TMEM176B*, *FBXO7*, *CEBPB*, *JAK3*, *PRDM1*, *HSPA9*, *TCIM*, *CASP8*, *SOX11*, *MYC*, *TNFSF9*, *LPAR3*, *CLDN5*, *IKBKB*, *LIG4*, *CYP26B1*, *KAT2A*, *INHBA*, *TFE3*, *RUNX1*, *S1PR2*, *PTK2B*, *NRP1*, *KLF10*, *GATA2*, *SEMA4D*, *RASSF2*, *FMR1*, *BMP2* |
| GO:0010498 | Proteasomal protein catabolic process | 131 | 1.52 | 5.83E-03 | 1.34E-02 | *PSMB9*, *PSME2*, *PSME1*, *PSMA5*, *HSP90B1*, *ZFAND2A*, *RNF122*, *DNAJB9*, *FBXO7*, *DDIT3*, *DERL3*, *CALR*, *SELENOS*, *TNFAIP1*, *PSMD10*, *PSMB3*, *KLHDC3*, *YOD1*, *FBXO4*, *UBE2D1*, *PAQR3*, *USP44*, *RAD23A*, *VCP*, *DESI1*, *PSMD8*, *STT3B*, *PSMF1*, *UBE2B*, *SIAH2*, *PSME3*, *EDEM2*, *PSMC1*, *KLHL15*, *TAF9*, *SIAH1*, *OGT*, *PSMD4*, *ERLEC1*, *TMEM129*, *MDM2*, *WWP1*, *N4BP1*, *NFE2L2*, *TOPORS*, *KEAP1*, *RNF5*, *AKIRIN2*, *TMEM259*, *NEMF*, *FBXO9*, *RAD23B*, *LTN1*, *SOCS5*, *KLHL2*, *ARIH2*, *UBXN4*, *DDRGK1*, *PSMB7* |
| GO:0048585 | Negative regulation of response to stimulus | 401 | 1.52 | 6.42E-04 | 2.06E-03 | *ISG15*, *CCL2*, *DHX58*, *CXCL8*, *STAT1*, *SERPINE1*, *SOCS1*, *SOCS3*, *PTGIR*, *BPIFB1*, *DUSP5*, *ATF3*, *THBS1*, *TNFAIP3*, *RGS16*, *PIM1*, *IFI35*, *STAT2*, *LTF*, *DKK1*, *ZFP36*, *GDF15*, *ADRA2A*, *LIF*, *CISH*, *DUSP10*, *NFKBIA*, *EGR1*, *SERPING1*, *SESN2*, *SLC39A8*, *TSKU*, *IL10*, *APOD*, *RGS2*, *OPTN*, *BCL6*, *DLL4*, *MAD2L2*, *HAVCR2*, *GPR155*, *FBXO7*, *ARRDC3*, *JAK3*, *CGAS*, *CFLAR*, *PARP3*, *DDIT3*, *ADM*, *TCIM*, *PAK1IP1*, *CASP8*, *YBX3*, *CALR*, *SELENOS*, *ACKR3*, *KLRK1*, *MRAP2*, *MCL1*, *MYC*, *TNFAIP1*, *TLR4*, *ELF4*, *CREB3L1*, *PSMD10*, *NKX3-1*, *RNF115*, *GHITM*, *INSIG1*, *PSMB4*, *WNT16*, *LYAR*, *UBE2D1*, *DDIT4*, *ADORA1*, *FBP1*, *GLI2*, *SPRY2*, *PAQR3*, *CYP26B1*, *CD59*, *SLC25A5*, *SOCS7*, *CD46* |

| | | | | | | |
|---|---|---|---|---|---|---|
| GO:0040012 | Regulation of locomotion | 233 | 1.51 | 3.25E-03 | 8.15E-03 | *CXCL10*, *MADCAM1*, *IFNG*, *SELE*, *CCL2*, *CCL4*, *CXCL8*, *THBS1*, *CCL8*, *CD40*, *CCL5*, *BVES*, *GDF15*, *ADRA2A*, *CCR1*, *DUSP10*, *GLUL*, *NR4A3*, *TNFSF14*, *CXCL16*, *KIT*, *APOD*, *GJA1*, *PHLDA2*, *RHOB*, *BCL6*, *DLL4*, *FBXO7*, *ARRDC3*, *CASP8*, *CALR*, *KLRK1*, *RIPK3*, *TLR4* |
| GO:0010648 | Negative regulation of cell communication | 352 | 1.50 | 9.03E-04 | 2.70E-03 | *ISG15*, *DHX58*, *CXCL8*, *STAT1*, *SOCS1*, *SOCS3*, *PTGIR*, *BPIFB1*, *DUSP5*, *ATF3*, *THBS1*, *TNFAIP3*, *RGS16*, *IFI35*, *STAT2*, *LTF*, *DKK1*, *FFAR2*, *GDF15*, *ADRA2A*, *LIF*, *CISH*, *DUSP10*, *NFKBIA*, *EGR1*, *SESN2*, *SLC39A8*, *TSKU*, *IL10*, *APOD*, *GJA1*, *RGS2*, *OPTN*, *BCL6*, *DLL4*, *MAD2L2*, *GPR155*, *FBXO7*, *ARRDC3*, *CGAS*, *CFLAR*, *DDIT3*, *ADM*, *TCIM*, *PAK1IP1*, *CASP8*, *YBX3*, *CALR*, *SELENOS*, *ACKR3*, *MRAP2*, *MCL1*, *MYC*, *TNFAIP1*, *TLR4*, *CREB3L1*, *PSMD10*, *NKX3-1*, *RNF115*, *GHITM*, *INSIG1*, *WNT16*, *UBE2D1*, *DDIT4*, *ADORA1*, *FBP1*, *GLI2*, *SPRY2*, *PAQR3*, *CYP26B1*, *SLC25A5*, *SOCS7*, *TP53* |
| GO:2000147 | Positive regulation of cell motility | 139 | 1.50 | 7.09E-03 | 1.59E-02 | *CXCL10*, *MADCAM1*, *IFNG*, *CCL2*, *CCL4*, *THBS1*, *CCL8*, *CD40*, *CCL5*, *ADRA2A*, *CCR1*, *NR4A3*, *TNFSF14*, *CXCL16*, *KIT*, *RHOB*, *BCL6*, *CASP8*, *CALR* |
| GO:0006511 | Ubiquitin-dependent protein catabolic process | 170 | 1.50 | 5.36E-03 | 1.24E-02 | *PSMB9*, *RNF19B*, *TNFAIP3*, *PSMA5*, *PSMA2*, *ZFAND2A*, *PSMA3*, *RNF122*, *FBXO7*, *DDIT3*, *PSMA1*, *TNFAIP1*, *PSMD10*, *PSMB3*, *RNF115*, *KLHDC3*, *FBXO4*, *UBE2D1*, *PSMD3*, *PAQR3*, *PSMA4*, *NDFIP2*, *USP44*, *UBE2D3*, *PTK2B*, *RAD23A*, *VCP*, *DESI1*, *PSMD8*, *RNF114*, *CHMP5*, *PSMF1*, *RLIM*, *PLAA*, *CHMP4A*, *UCHL5*, *UBE2B*, *TRAF7*, *SIAH2*, *PSMC1*, *PSMC2*, *ARK2N*, *KLHL15*, *TAF9*, *SIAH1*, *OGT*, *PSMD4*, *CUL5*, *RNF14*, *RBBP6*, *TMEM129*, *UBE3A*, *MDM2*, *CACUL1*, *WWP1*, *TMEM168*, *N4BP1*, *NFE2L2*, *TOPORS*, *KEAP1*, *RNF5*, *AKIRIN2*, *NEMF*, *FBXO9*, *ELOC*, *RAD23B*, *LTN1*, *SOCS5*, *KLHL2* |
| GO:0009894 | Regulation of catabolic process | 237 | 1.49 | 5.98E-03 | 1.36E-02 | *SAA3*, *IFNG*, *PSME2*, *TNFAIP3*, *PSME1*, *ZFP36*, *TIMP1*, *ADRA2A*, *SESN2*, *DEPP1*, *IL10*, *ZFAND2A*, *ARID5A*, *PLK3*, *DRAM2*, *OPTN*, *STING1*, *DRAM1*, *MAD2L2*, *TIGAR*, *TRAF2*, *FBXO7*, *TICAM1*, *IGF2BP1*, *MCL1*, *RAB39B*, *PSMD10*, *AGO2*, *ABHD5*, *DDIT4*, *FBP1*, *PSMD3*, *RUBCN*, *TIRAP*, *PAQR3*, *SLC25A5*, *TMEM39A*, *RALB*, *TIMP3*, *PTK2B*, *PDE12* |
| GO:0023057 | Negative regulation of signaling | 354 | 1.49 | 1.96E-03 | 5.32E-03 | *ISG15*, *DHX58*, *CXCL8*, *STAT1*, *SOCS1*, *SOCS3*, *PTGIR*, *BPIFB1*, *DUSP5*, *ATF3*, *THBS1*, *TNFAIP3*, *RGS16*, *IFI35*, *STAT2*, *LTF*, *DKK1*, *FFAR2*, *GDF15*, *ADRA2A*, *LIF*, *CISH*, *DUSP10*, *NFKBIA*, *EGR1*, *SESN2*, *SLC39A8*, *TSKU*, *IL10*, *APOD*, *GJA1*, *RGS2*, *OPTN*, *BCL6*, *DLL4*, |

| | | | | | | |
|---|---|---|---|---|---|---|
| | | | | | | *MAD2L2*, *GPR155*, *FBXO7*, *ARRDC3*, *CGAS*, *CFLAR*, *DDIT3*, *ADM*, *TCIM*, *PAK1IP1*, *CASP8*, *YBX3*, *CALR*, *SELENOS*, *ACKR3*, *MRAP2*, *MCL1*, *MYC*, *TNFAIP1*, *TLR4*, *CREB3L1*, *PSMD10*, *NKX3-1*, *RNF115*, *GHITM*, *INSIG1*, *WNT16*, *UBE2D1*, *DDIT4*, *ADORA1*, *FBP1*, *GLI2*, *SPRY2*, *PAQR3*, *CYP26B1*, *SLC25A5*, *SOCS7*, *TP53* |
| GO:0006412 | Translation | 229 | 1.49 | 5.34E-03 | 1.24E-02 | *WARS1*, *THBS1*, *ZFP36*, *SESN2*, *GARS1*, *RGS2*, *RPL13A*, *YARS1*, *FARSA*, *EIF4EBP1*, *CALR*, *IGF2BP1*, *EIF2S2*, *NARS1*, *VARS1*, *AGO2*, *EIF4A1*, *EIF5A*, *ETF1*, *EIF2S1*, *PUM3*, *MRPL15*, *IGF2BP2*, *MRPL17*, *EIF3I*, *NAT10*, *RSL24D1*, *EARS2*, *FMR1*, *GFM2*, *OGFOD1*, *DDX3X*, *EIF3J*, *DRG1*, *MRPL35*, *EIF1*, *ACO1*, *NPM1*, *RIDA*, *C7H6orf125*, *PIWIL4*, *BZW1*, *RARS1*, *EIF3A*, *ATM*, *STK35*, *PTCD3*, *ABCF1*, *MRPS11*, *PAIP2B*, *AIMP2*, *METTL3*, *SRP9*, *HARS1*, *OGT*, *SERBP1*, *EIF4A3*, *RPL22L1*, *MRPS7*, *DARS1*, *MRPL20*, *GAPDH*, *RNF14*, *MRPS18B*, *ELAVL1*, *MRPL44*, *MRPL47*, *GUF1*, *MTOR*, *SSB*, *FTSJ1*, *QRSL1*, *MTRF1*, *MRPS5*, *MRPL2*, *DAP*, *MRPL1*, *MRPL32*, *NEMF*, *CAPRIN1*, *RPS6KB1*, *LTN1*, *MRPL18*, *PARS2*, *DDRGK1*, *HARS2*, *EIF3M*, *UPF3B* |
| GO:0048646 | Anatomical structure formation involved in morphogenesis | 275 | 1.49 | 1.56E-03 | 4.49E-03 | *CXCL9*, *CXCL10*, *CHI3L1*, *CXCL8*, *STAT1*, *MMP8*, *SERPINE1*, *TNFSF13B*, *IL1A*, *MYBPH*, *DUSP5*, *THBS1*, *TNFAIP3*, *JUNB*, *PIM1*, *CD40*, *GDF15*, *ANGPTL4*, *ESM1*, *GLUL*, *TNFSF14*, *IL10*, *CHN2*, *GJA1*, *VASH2*, *RHOB*, *BCL6*, *DLL4*, *B4GALT1*, *ADM*, *GARIN1A*, *ACKR3*, *SOX11*, *ITGA5*, *CREB3L1*, *NKX3-1*, *SLC31A1*, *PLEKHO1*, *CLDN5*, *SMAD1*, *WDR74*, *SAT1*, *GNA13*, *GLI2*, *SPRY2* |
| GO:1903530 | Regulation of secretion by cell | 107 | 1.49 | 1.01E-02 | 2.20E-02 | *IFNG*, *RSAD2*, *IL1B*, *IL6*, *IL1A*, *RHBDF2*, *CCL5*, *FFAR2*, *ADRA2A*, *LIF*, *GJA1*, *F2RL2*, *SOX11*, *TLR2*, *TLR4*, *NKX3-1*, *CADPS2*, *TGFB3*, *KCNA2*, *TMED10*, *LRRC8A*, *SPP1*, *TP53*, *CRY1*, *INHBA* |
| GO:0032940 | Secretion by cell | 162 | 1.48 | 1.49E-02 | 3.06E-02 | *IFNG*, *RSAD2*, *IL1B*, *GNAT1*, *IL6*, *IL1A*, *RHBDF2*, *CCL8*, *STX11*, *CCL5*, *FFAR2*, *ADRA2A*, *CCR1*, *LIF*, *KIT*, *GJA1*, *F2RL2*, *SOX11*, *TLR2*, *TLR4*, *NKX3-1*, *CADPS2*, *TGFB3*, *KCNA2*, *TMEM38B*, *TMED10*, *LRRC8A*, *SPP1*, *TP53*, *CRY1*, *INHBA*, *KCNK1*, *C2CD2L*, *RAB1A*, *TVP23B*, *SNX19*, *BMP2*, *SYN1* |

| | | | | | | |
|---|---|---|---|---|---|---|
| GO:2001233 | Regulation of apoptotic signaling pathway | 108 | 1.48 | 1.02E-02 | 2.20E-02 | *S100A8*, *ATF3*, *THBS1*, *TNFAIP3*, *FASLG*, *IL10*, *TRAF2*, *FBXO7*, *CFLAR*, *DDIT3*, *YBX3*, *SELENOS*, *ACKR3*, *MCL1*, *RIPK3*, *TLR4*, *PSMD10*, *NKX3-1*, *GHITM*, *JAK2*, *WNT16*, *SLC25A5*, *G0S2*, *TP53*, *INHBA*, *SLC35F6*, *NRP1*, *DDX3X*, *HYAL2*, *HMOX1*, *CTSC*, *BID*, *TRAF7*, *SIAH2*, *PSME3* |
| GO:0008284 | Positive regulation of cell population proliferation | 215 | 1.48 | 6.49E-03 | 1.46E-02 | *IL1B*, *STAT1*, *TNFSF13B*, *IL6*, *IL1A*, *CSF3*, *THBS1*, *TNFAIP3*, *PIM1*, *CD40*, *IL15*, *LTF*, *CCL5*, *TIMP1*, *STAT5A*, *CSF2*, *LIF*, *ESM1*, *NR4A3*, *TFRC*, *IL10*, *GJA1*, *NME1*, *BCL6*, *DLL4*, *HAVCR2*, *SLC7A1*, *B4GALT1*, *ADM*, *CALR*, *TICAM1*, *SOX11*, *MYC*, *TNFSF9*, *RNASEH2B*, *TLR4*, *NKX3-1*, *TGFB3*, *FAM98A*, *IL6R*, *CLDN5*, *SLC25A33*, *NOP2*, *IGFBP2*, *JAK2*, *TIPIN* |
| GO:0009968 | Negative regulation of signal transduction | 333 | 1.48 | 1.87E-03 | 5.13E-03 | *ISG15*, *DHX58*, *CXCL8*, *STAT1*, *SOCS1*, *SOCS3*, *PTGIR*, *BPIFB1*, *DUSP5*, *ATF3*, *THBS1*, *TNFAIP3*, *RGS16*, *IFI35*, *STAT2*, *LTF*, *DKK1*, *GDF15*, *LIF*, *CISH*, *DUSP10*, *NFKBIA*, *EGR1*, *SESN2*, *SLC39A8*, *TSKU*, *IL10*, *APOD*, *RGS2*, *OPTN*, *BCL6*, *DLL4*, *MAD2L2*, *GPR155*, *FBXO7*, *ARRDC3*, *CGAS*, *CFLAR*, *DDIT3*, *ADM*, *TCIM*, *PAK1IP1*, *CASP8*, *YBX3*, *CALR*, *SELENOS*, *ACKR3*, *MRAP2*, *MCL1*, *MYC*, *TNFAIP1*, *TLR4*, *CREB3L1*, *PSMD10*, *NKX3-1*, *RNF115*, *GHITM*, *INSIG1*, *WNT16*, *UBE2D1*, *DDIT4*, *FBP1*, *GLI2*, *SPRY2*, *PAQR3*, *CYP26B1*, *SLC25A5* |
| GO:0030334 | Regulation of cell migration | 209 | 1.47 | 6.03E-03 | 1.37E-02 | *CXCL10*, *MADCAM1*, *IFNG*, *SELE*, *CCL2*, *CCL4*, *THBS1*, *CCL8*, *CD40*, *CCL5*, *GDF15*, *ADRA2A*, *CCR1*, *DUSP10*, *GLUL*, *NR4A3*, *TNFSF14*, *CXCL16*, *KIT*, *APOD*, *GJA1*, *PHLDA2*, *RHOB*, *DLL4*, *CASP8*, *CALR*, *KLRK1*, *RIPK3*, *TLR4* |
| GO:0043687 | Post-translational protein modification | 268 | 1.47 | 3.43E-03 | 8.55E-03 | *ISG15*, *UBA7*, *USP18*, *SOCS1*, *SOCS3*, *RNF19B*, *TNFAIP3*, *PARP12*, *EGR2*, *TRIM21*, *CISH*, *EGR1*, *PARP11*, *RNF122*, *GNL3*, *TRAF2*, *FBXO7*, *PARP3*, *USP15*, *TICAM1*, *TNFAIP1*, *WSB2*, *PSMD10*, *RNF115*, *JAK2*, *FBXO4*, *UBE2D1*, *GTPBP4*, *USP2*, *SPRY2*, *SOCS7*, *OTUB2*, *CRY1*, *NDFIP2*, *PADI4*, *USP44*, *KLHL25*, *UBE2D3*, *EIPR1*, *RNF138*, *VCP*, *DESI1*, *RNF114*, *SQSTM1*, *DDX3X*, *STT3B*, *RLIM*, *PLAA*, *WBP1L*, *UCHL5*, *DCAF11*, *UBE2B*, *TRAF7*, *SIAH2*, *PARP8*, *DCUN1D5*, *PHF23*, *USP39*, *HINT1*, *TPST1*, *KLHL15*, *DCUN1D3*, *TRIM27*, *SIAH1*, *OGT*, *FBXO33*, *OTUD6B*, *BCL10*, *CUL5*, *MSL2*, *RNF14*, *RBBP6*, *MINDY3*, *UBA3*, *TMEM129*, *UBE3A*, *UBA6*, *MDM2*, *OTUB1*, *RNF141*, *WWP1*, *USP5*, *PRPF19*, *N4BP1*, *LOC100625207*, *NFE2L2*, *TOPORS*, *ASB5*, *KEAP1*, *RNF5*, *UFC1* |

| | | | | | | |
|---|---|---|---|---|---|---|
| GO:0141124 | Intracellular signaling cassette | 454 | 1.46 | 6.47E-04 | 2.06E-03 | *SAA3*, *RND1*, *CHI3L1*, *RASGEF1A*, *S100A12*, *STAT1*, *FAS*, *DUSP5*, *ATF3*, *THBS1*, *TNFAIP3*, *TIFA*, *CD74*, *IFI35*, *FASLG*, *ACKR4*, *CD40*, *CCL5*, *ZFP36*, *GADD45G*, *TICAM2*, *GDF15*, *NFKBIB*, *CXCR6*, *ADRA2A*, *CCR1*, *CSF2*, *LIF*, *DUSP10*, *NFKBIA*, *SESN2*, *SLC39A8*, *TNFSF14*, *GPR37*, *CHN2*, *OPTN*, *ACKR2*, *RHOB*, *BCL6*, *TRAF3*, *TNIP1*, *TRAF2*, *RASSF1*, *HAVCR2*, *DDX21*, *RHOJ*, *CGAS*, *CFLAR*, *MYD88*, *DDIT3*, *TCIM*, *ADAM17*, *CASP8*, *CALR*, *ACKR3*, *TICAM1*, *RAB39B*, *MYC*, *TNFAIP1*, *TLR2*, *CDC42EP3*, *RIPK3*, *TLR4*, *PSMD10*, *NKX3-1*, *LPAR3*, *TMEM106A*, *GADD45B*, *TMEM38B*, *TNIP2*, *SLC11A1*, *IKBKB*, *JAK2*, *WNT16*, *SMAD1*, *CDC42SE1*, *FBP1*, *RIPK1*, *GNA13*, *TIRAP*, *SPRY2*, *CDC42EP2*, *EIF2S1*, *ATP1B1*, *GPR4*, *LTBR* |
| GO:0007155 | Cell adhesion | 326 | 1.45 | 4.04E-03 | 9.64E-03 | *MADCAM1*, *IFNG*, *SELE*, *CD274*, *IRF1*, *RND1*, *CXCL8*, *IL1B*, *S100A8*, *TNFSF13B*, *SOCS1*, *IL1A*, *CLSTN3*, *THBS1*, *SLA-DQB1*, *VCAM1*, *SLA-DMB*, *IL15*, *CCL5*, *CDH12*, *STAT5A*, *BVES*, *SELP*, *LRRC32*, *AMBN*, *LIF*, *DUSP10*, *TNC*, *NR4A3*, *TFRC*, *OLR1*, *SLC39A8*, *TNFSF14*, *IL10*, *KIT*, *APOD*, *RHOB*, *BCL6*, *MAD2L2*, *HAVCR2*, *SLC7A1*, *CEBPB*, *JAK3*, *B4GALT1*, *ICAM1*, *CALR*, *VCAN*, *ITGA5*, *TNFSF9*, *ICAM4*, *CLDN5*, *IGFBP2*, *JAK2*, *FUT2*, *AMIGO2*, *LYVE1*, *THBS2*, *GTPBP4*, *SIGLEC1*, *SNED1*, *SPP1*, *ATP1B1*, *GPR4*, *CLDN4*, *ARG2*, *ENTPD1* |
| GO:0022603 | Regulation of anatomical structure morphogenesis | 188 | 1.45 | 1.22E-02 | 2.60E-02 | *CXCL10*, *CHI3L1*, *CCL2*, *CXCL8*, *STAT1*, *SERPINE1*, *TNFSF13B*, *IL1A*, *THBS1*, *TNFAIP3*, *STAT2*, *CD40*, *BVES*, *LIF*, *GLUL*, *IL10*, *KIT*, *SLC26A5*, *GJA1*, *VASH2*, *RHOB*, *ADM*, *ITGA5*, *CDC42EP3*, *CREB3L1*, *PLEKHO1*, *LPAR3*, *CLDN5*, *MIEF1*, *SMAD1*, *CDC42SE1*, *GNA13*, *SPRY2*, *CDC42EP2*, *GPR4*, *CLDN4*, *FITM2*, *F3*, *AQP1*, *PTK2B*, *NRP1*, *GATA2*, *SEMA4D*, *HTATIP2*, *BMP2*, *RRAS* |
| GO:0016071 | MRNA metabolic process | 244 | 1.43 | 1.23E-02 | 2.60E-02 | *RNASEL*, *ZFP36*, *DNAJB11*, *ARID5A*, *CEBPG*, *TRAF2*, *IGF2BP1*, *AGO2*, *PAPOLA*, *PNPT1*, *NHP2*, *PUS3*, *ETF1*, *TIRAP*, *PELO*, *SNRPD1*, *SNRPA1*, *SNRPF*, *SMNDC1*, *VDR*, *PDE12*, *TRA2B*, *RBM8A*, *FMR1*, *DXO*, *RPRD1B*, *TBRG4*, *PUS1*, *RBM14*, *NPM1*, *RIDA*, *SNRPG*, *PRPF40A*, *EXOSC3*, *STRAP*, *SMN1*, *SNRPD3*, *SNRPA*, *ATM*, *GEMIN6*, *SRSF1*, *WTAP*, *CDK12*, *RBM25*, *SNRNP27*, *USP39*, *HNRNPF*, *ZFP36L1*, *CSTF2*, *SRSF7*, *ZFP36L2*, *RBM7*, *AHCYL1*, *METTL3*, *TENT2*, *TAF9*, *RNPS1*, *SMU1*, *SNRNP40*, *EIF4A3*, *PRPF3*, *SF3B6*, *TENT5A*, *CNOT9*, |

| | | | | | | |
|---|---|---|---|---|---|---|
| | | | | | | *EXOSC4*, *ZMAT2*, *HIPK3*, *CLNS1A*, *RBBP6*, *ELAVL1*, *LSM6*, *CSTF3*, *SNRPC*, *RBM47*, *PCF11*, *SRSF2*, *RBM39*, *ZBTB7A*, *MED1*, *SCAF11*, *SSB*, *ZRANB2*, *PRPF19*, *CNOT7*, *PTBP1*, *SNRPB2*, *CWC22*, *SYMPK*, *WDR83*, *PAN2*, *RBM15*, *U2AF1L4*, *CAPRIN1*, *GEMIN8*, *CDC40*, *NSRP1*, *PCID2*, *SFSWAP*, *UPF3B*, *SMG9*, *CSTF2T*, *DCP2*, *DDX39B*, *DHX8*, *PABPC1*, *SNU13*, *ZC3H14*, *DCPS*, *ROCK1*, *LEO1*, *SENP1*, *DYRK1A*, *ZBTB1* |
| GO:0048871 | Multicellular organismal-level homeostasis | 185 | 1.43 | 1.09E-02 | 2.34E-02 | *ISG15*, *IL1B*, *STAT1*, *TNFSF13B*, *FAS*, *IL1A*, *SNX10*, *TNFAIP3*, *AQP3*, *AQP7*, *ZFP36*, *STAT5A*, *GDF15*, *EGR1*, *BPIFA1*, *STAT3*, *SLC39A8*, *ST6GALNAC1*, *TSKU*, *SLC1A5*, *KIT*, *CASP3*, *DRAM2*, *GJA1*, *CEBPG*, *BCL6*, *CEBPB*, *ARRDC3*, *DDIT3*, *HSPA9*, *RIPK3*, *TLR4*, *CLDN5*, *JAK2*, *LYAR*, *PPARGC1A*, *SFTPD* |
| GO:0000165 | MAPK cascade | 169 | 1.43 | 1.41E-02 | 2.94E-02 | *SAA3*, *CHI3L1*, *FAS*, *DUSP5*, *ATF3*, *CD40*, *ZFP36*, *GADD45G*, *GDF15*, *ADRA2A*, *CCR1*, *CSF2*, *LIF*, *DUSP10*, *GPR37*, *TRAF2*, *HAVCR2*, *ADAM17*, *ACKR3*, *MYC*, *TLR4*, *PSMD10*, *LPAR3*, *TMEM106A*, *GADD45B*, *SLC11A1*, *WNT16*, *SMAD1*, *TIRAP*, *SPRY2* |
| GO:0043408 | Regulation of MAPK cascade | 154 | 1.42 | 1.48E-02 | 3.06E-02 | *SAA3*, *CHI3L1*, *FAS*, *DUSP5*, *ATF3*, *CD40*, *GADD45G*, *GDF15*, *ADRA2A*, *CCR1*, *CSF2*, *LIF*, *DUSP10*, *GPR37*, *TRAF2*, *HAVCR2*, *ADAM17*, *ACKR3*, *MYC*, *TLR4*, *PSMD10*, *LPAR3*, *TMEM106A*, *GADD45B*, *WNT16* |
| GO:0006914 | Autophagy | 155 | 1.42 | 1.73E-02 | 3.50E-02 | *IFNG*, *S100A8*, *FUNDC1*, *SESN2*, *DEPP1*, *IL10*, *PLK3*, *DRAM2*, *OPTN*, *TMEM74*, *STING1*, *DRAM1*, *TIGAR*, *FBXO7*, *TICAM1*, *MCL1*, *RAB39B*, *CALCOCO2*, *RUBCN*, *EIF2S1*, *SLC25A5*, *TMEM39A*, *RALB*, *GABARAPL1*, *RAB1A*, *ATG2A*, *VCP*, *CHMP5*, *SQSTM1*, *STX17* |
| GO:0061919 | Process utilizing autophagic mechanism | 155 | 1.42 | 1.73E-02 | 3.50E-02 | *IFNG*, *S100A8*, *FUNDC1*, *SESN2*, *DEPP1*, *IL10*, *PLK3*, *DRAM2*, *OPTN*, *TMEM74*, *STING1*, *DRAM1*, *TIGAR*, *FBXO7*, *TICAM1*, *MCL1*, *RAB39B*, *CALCOCO2*, *RUBCN*, *EIF2S1*, *SLC25A5*, *TMEM39A*, *RALB*, *GABARAPL1*, *RAB1A*, *ATG2A*, *VCP*, *CHMP5*, *SQSTM1*, *STX17* |
| GO:2000145 | Regulation of cell motility | 225 | 1.41 | 1.37E-02 | 2.86E-02 | *CXCL10*, *MADCAM1*, *IFNG*, *SELE*, *CCL2*, *CCL4*, *THBS1*, *CCL8*, *CD40*, *CCL5*, *GDF15*, *ADRA2A*, *CCR1*, *DUSP10*, *GLUL*, *NR4A3*, *TNFSF14*, *CXCL16*, *KIT*, *APOD*, *GJA1*, *PHLDA2*, *RHOB*, *BCL6*, *DLL4*, *CASP8*, *CALR*, *KLRK1*, *RIPK3*, *TLR4* |
| GO:0070647 | Protein modification by small protein | 256 | 1.41 | 1.62E-02 | 3.32E-02 | *ISG15*, *UBA7*, *USP18*, *SOCS1*, *SOCS3*, *RNF19B*, *TNFAIP3*, *EGR2*, *TRIM21*, *CISH*, *EGR1*, *RNF122*, *GNL3*, *TRAF2*, *FBXO7*, *USP15*, *TICAM1*, *TNFAIP1*, *WSB2*, *PSMD10*, *RNF115*, |

| | | | | | | |
|---|---|---|---|---|---|---|
| | conjugation or removal | | | | | *FBXO4*, *UBE2D1*, *GTPBP4*, *USP2*, *SPRY2*, *SOCS7*, *OTUB2*, *CRY1*, *NDFIP2*, *USP44*, *KLHL25*, *UBE2D3*, *EIPR1*, *RNF138*, *VCP*, *DESI1*, *RNF114*, *SQSTM1*, *DDX3X*, *RLIM*, *PLAA*, *WBP1L*, *UCHL5*, *DCAF11*, *UBE2B*, *TRAF7*, *SIAH2*, *DCUN1D5*, *PHF23*, *USP39*, *HINT1*, *KLHL15*, *DCUN1D3*, *TRIM27*, *SIAH1*, *OGT*, *FBXO33*, *OTUD6B*, *BCL10*, *CUL5*, *MSL2*, *RNF14*, *RBBP6*, *MINDY3*, *UBA3*, *TMEM129*, *UBE3A*, *UBA6*, *MDM2*, *OTUB1*, *RNF141*, *WWP1*, *USP5*, *PRPF19*, *N4BP1*, *LOC100625207*, *NFE2L2*, *TOPORS*, *ASB5*, *KEAP1*, *RNF5*, *UFC1* |
| GO:0009725 | Response to hormone | 167 | 1.40 | 2.07E-02 | 4.13E-02 | *STAT1*, *SOCS1*, *JUNB*, *PIM1*, *STAT2*, *ZFP36*, *TIMP1*, *STAT5A*, *GDF15*, *ADRA2A*, *EGR1*, *STAT3*, *SESN2*, *NR4A3*, *SLC2A4*, *IL10*, *ARID5A*, *JAK3*, *CALR*, *NKX3-1*, *TGFB3*, *KLF9*, *SLC25A33*, *IGFBP2*, *JAK2*, *DDIT4*, *GOT1*, *PTGES3*, *CYP26B1*, *CRY1*, *VDR*, *AQP1*, *TIMP3* |
| GO:0009890 | Negative regulation of biosynthetic process | 481 | 1.39 | 2.97E-03 | 7.60E-03 | *SAA3*, *IFNG*, *IRF1*, *DHX58*, *CXCL8*, *SERPINE1*, *THBS1*, *TNFAIP3*, *FASLG*, *LTF*, *ZFP36*, *MOV10*, *LRRC32*, *CCR1*, *CEBPD*, *EGR1*, *RFX5*, *ATF5*, *SESN2*, *TSKU*, *IL10*, *ARID5A*, *MFSD12*, *PLK3*, *APOD*, *GJA1*, *BCL6*, *DLL4*, *MAD2L2*, *TRAF2*, *TXNIP*, *BAZ1A*, *HAVCR2*, *CEBPB*, *MORC3*, *PRDM1*, *PARP3*, *DDIT3*, *EIF4EBP1*, *YBX3*, *CALR*, *SOX11*, *IGF2BP1*, *RNASEH2B*, *TLR4*, *ELF4*, *CREB3L1*, *PSMD10*, *NKX3-1*, *ORMDL1*, *AGO2*, *TGFB3*, *PNPT1*, *CLDN5*, *INSIG1*, *LYAR*, *FBP1*, *ETF1*, *TIRAP*, *GLI2*, *USP2*, *NKRF*, *PELO*, *EIF2S1*, *KAT2A*, *TP53*, *ARG2*, *LOC100624590*, *CRY1*, *NDFIP2*, *VDR*, *PDE12*, *MTERF3*, *NAT10*, *KLF10*, *CIPC*, *GATA2*, *RBM8A*, *SEMA4D*, *FMR1*, *KDM8*, *BMP2*, *DXO*, *PDCD4*, *CMKLR1*, *TBRG4*, *SUV39H2*, *SETDB2*, *SQSTM1*, *DDX3X*, *RLIM*, *GATAD2A*, *ADAM10*, *DNAJB1*, *NOS2*, *BASP1*, *EPHA2*, *MACIR*, *NPM1*, *RIDA*, *IL27RA*, *PPARD*, *PIWIL4*, *ORMDL3*, *EXOSC3*, *STRAP*, *CD84*, *PID1* |
| GO:0010558 | Negative regulation of macromolecule biosynthetic process | 465 | 1.38 | 7.21E-03 | 1.59E-02 | *SAA3*, *IFNG*, *IRF1*, *DHX58*, *CXCL8*, *SERPINE1*, *THBS1*, *TNFAIP3*, *FASLG*, *LTF*, *ZFP36*, *MOV10*, *LRRC32*, *CCR1*, *CEBPD*, *EGR1*, *RFX5*, *ATF5*, *SESN2*, *TSKU*, *IL10*, *ARID5A*, *PLK3*, *APOD*, *GJA1*, *BCL6*, *DLL4*, *MAD2L2*, *TRAF2*, *TXNIP*, *BAZ1A*, *HAVCR2*, *CEBPB*, *MORC3*, *PRDM1*, *PARP3*, *DDIT3*, *EIF4EBP1*, *YBX3*, *CALR*, *SOX11*, *IGF2BP1*, *RNASEH2B*, *TLR4*, *ELF4*, *CREB3L1*, *PSMD10*, *NKX3-1*, *AGO2*, *TGFB3*, *PNPT1*, *CLDN5*, *LYAR*, *FBP1*, *ETF1*, *TIRAP*, *GLI2*, *USP2*, *NKRF*, *PELO*, *EIF2S1*, *KAT2A*, *TP53*, *ARG2*, *LOC100624590*, *CRY1*, *NDFIP2*, *VDR*, *PDE12*, *MTERF3*, *NAT10*, *KLF10*, *CIPC*, *GATA2*, *RBM8A*, *SEMA4D*, *FMR1*, |

| | | | | | | |
|---|---|---|---|---|---|---|
| | | | | | | *KDM8*, *BMP2*, *DXO*, *PDCD4*, *CMKLR1*, *TBRG4*, *SUV39H2*, *SETDB2*, *SQSTM1*, *DDX3X*, *RLIM*, *GATAD2A*, *ADAM10*, *DNAJB1*, *NOS2*, *BASP1*, *EPHA2*, *MACIR*, *NPM1*, *RIDA*, *IL27RA*, *PPARD*, *PIWIL4*, *EXOSC3*, *STRAP*, *CD84* |
| GO:0006796 | Phosphate-containing compound metabolic process | 389 | 1.34 | 1.35E-02 | 2.84E-02 | *IDO1*, *IFNG*, *CHI3L1*, *SAMHD1*, *GFPT2*, *PPA1*, *THBS1*, *HK3*, *STAT2*, *IL15*, *CCL5*, *C3*, *SLC25A22*, *NAMPT*, *LIF*, *CTPS1*, *STAT3*, *GK*, *CNP*, *GCK*, *NME1*, *TIGAR*, *UGP2*, *PLPPR2*, *NEK6*, *HK2*, *ITGA5*, *RNASEH2B*, *ORMDL1*, *PGK1*, *TMEM38B*, *PRPS1*, *ABHD5*, *JAK2*, *ITPKC*, *PDXK*, *DPYS*, *KMO*, *GFPT1*, *ACSL4*, *DDIT4*, *PISD*, *DCTD*, *FBP1*, *PLSCR3*, *GOT1*, *CHKA*, *ATP1B1*, *CDS1*, *PDE4B*, *RFK*, *MDH1*, *ENTPD1*, *LPCAT3*, *GNPNAT1*, *FITM2*, *NUS1*, *DCTPP1*, *UAP1*, *UGDH*, *NRP1*, *SEMA4D*, *NT5C*, *ME1*, *PDE4A*, *AGPAT4*, *SAMD8*, *AK3*, *ATP5MC1*, *ACLY*, *PPP2R2A*, *AASS*, *ITPK1*, *PDHX* |
| GO:0006793 | Phosphorus metabolic process | 390 | 1.33 | 2.05E-02 | 4.13E-02 | *IDO1*, *IFNG*, *CHI3L1*, *SAMHD1*, *GFPT2*, *PPA1*, *THBS1*, *HK3*, *STAT2*, *IL15*, *CCL5*, *C3*, *SLC25A22*, *NAMPT*, *LIF*, *CTPS1*, *STAT3*, *GK*, *CNP*, *GCK*, *NME1*, *TIGAR*, *UGP2*, *PLPPR2*, *NEK6*, *HK2*, *ITGA5*, *RNASEH2B*, *ORMDL1*, *PGK1*, *TMEM38B*, *PRPS1*, *ABHD5*, *JAK2*, *ITPKC*, *PDXK*, *DPYS*, *KMO*, *GFPT1*, *ACSL4*, *DDIT4*, *PISD*, *DCTD*, *FBP1*, *PLSCR3*, *GOT1*, *CHKA*, *ATP1B1*, *CDS1*, *PDE4B*, *RFK*, *MDH1*, *ENTPD1*, *LPCAT3*, *GNPNAT1*, *FITM2*, *NUS1*, *DCTPP1*, *UAP1*, *UGDH*, *NRP1*, *SEMA4D*, *NT5C*, *ME1*, *PDE4A*, *AGPAT4*, *SAMD8*, *AK3*, *ATP5MC1*, *ACLY*, *PPP2R2A*, *AASS*, *ITPK1*, *PDHX* |
| GO:0048699 | Generation of neurons | 268 | -1.33 | 5.35E-03 | 1.24E-02 | *SYT3*, *EPHB3*, *NFIB*, *ITM2C*, *FLOT1*, *NTF3*, *VEGFA*, *ATAT1*, *LOC100525798*, *HOXA2*, *SEMA6D*, *BBS10*, *B4GALT6*, *DMD*, *ACP4*, *RTN4RL1*, *KREMEN1*, *BMP4*, *BBS1*, *PROX1*, *TYRO3*, *NR2E1*, *ID4*, *BMPR2*, *CHODL*, *VCL*, *SOX9*, *WNT7A*, *ID2*, *ACSL6*, *NYAP1*, *WNT11*, *WNT9A*, *DCX*, *BMP7*, *RTN4R*, *UCHL1*, *GPRASP3*, *RASGRF1*, *SEMA6A*, *GBX1*, *SEMA3E*, *SEMA6C*, *SPOCK1*, *DCT*, *EPHA7*, *WNT2*, *SEMA3B*, *STMN1*, *FOLR3*, *BEND6*, *CNTN1*, *MAP1B*, *CNR1*, *EFNB3*, *HEY1*, *BDNF*, *WNT4*, *CNTN5*, *EPHA3*, *MDGA1*, *ATCAY*, *ZNF365*, *NDP*, *MYCN*, *HHIP*, *CRB2*, *VEGFD*, *SFRP2* |

| | | | | | | |
|---|---|---|---|---|---|---|
| GO:0016055 | Wnt signaling pathway | 131 | -1.35 | 2.33E-02 | 4.60E-02 | *LGR4*, *WLS*, *KREMEN1*, *NRARP*, *PKD2*, *LGR5*, *FZD2*, *CPE*, *SOX9*, *DKK3*, *WNT7A*, *CALCOCO1*, *SDC1*, *WNT11*, *WNT9A*, *TLE4*, *FAM53B*, *GATA3*, *WNT10B*, *WNT3A*, *WNT2*, *FOLR3*, *SFRP5*, *WIF1*, *ALPK2*, *WNT4*, *SFRP4*, *DKK2*, *NDP*, *SFRP2* |
| GO:0060627 | Regulation of vesicle-mediated transport | 105 | -1.36 | 2.30E-02 | 4.57E-02 | *HYAL3*, *BICD1*, *WNT7A*, *SYT5*, *SDC1*, *HIP1R*, *SYT17*, *PLD1*, *F2RL1*, *LGI3*, *TUB*, *EPHA3*, *PYCARD*, *COLEC11*, *APLN*, *SYT12*, *CCR2*, *APOA1* |
| GO:0007017 | Microtubule-based process | 245 | -1.38 | 3.97E-03 | 9.56E-03 | *MAD2L1*, *TSSK4*, *BICDL2*, *TACC3*, *WDR62*, *SPACA9*, *DYNC2LI1*, *FOXJ1*, *PIERCE1*, *CFAP58*, *SPEF2*, *CCNB2*, *ARMC3*, *SPC25*, *HOOK2*, *CFAP20*, *CFAP91*, *HSPB1*, *SGO1*, *TEKT2*, *CDK1*, *KIF2C*, *SPDL1*, *ODAD3*, *ATAT1*, *NUF2*, *CCNB1*, *MAP10*, *NCKAP5L*, *CCDC68*, *MID1*, *DYNLT5*, *DYNLT4*, *TUBB2B*, *KIF18B*, *IQUB*, *CEP250*, *CGN*, *PARD6G*, *KIF11*, *BBS1*, *KIF24*, *CCNF*, *CFAP141*, *SLC22A16*, *DNAI2*, *CATSPER4*, *GEN1*, *KLC4*, *IFT46*, *CENPH*, *KIFC1*, *BICD1*, *AURKB*, *KIF20A*, *FAM110A*, *BIRC5*, *TUBB4A*, *MEIG1*, *DCX*, *WRAP73*, *UCHL1*, *CENPA*, *KIF22*, *CFAP276*, *CFAP97D1*, *DYNC2I2*, *TUB*, *SKA1*, *STMN1*, *TPPP3*, *RGS14*, *MAP1B*, *TPPP*, *ODAD1*, *EPHA3*, *DEFB1* |
| GO:0120036 | Plasma membrane bounded cell projection organization | 296 | -1.48 | 4.47E-04 | 1.56E-03 | *BORCS7*, *FGD1*, *NEK1*, *LCA5*, *GFAP*, *SMO*, *SEMA4F*, *TPBG*, *CCDC113*, *PLPPR4*, *SPATA6*, *NEDD9*, *SPACA9*, *FUZ*, *DYNC2LI1*, *EFNA4*, *ENPP2*, *FOXJ1*, *EXT1*, *NTN4*, *PIERCE1*, *SFRP1*, *NTF4*, *INPP5J*, *CFAP58*, *SPEF2*, *ARHGEF25*, *CFAP20*, *TACSTD2*, *ARL6*, *SRCIN1*, *CFAP91*, *BTBD3*, *EHD2*, *TEKT2*, *SYT3*, *EPHB3*, *NFIB*, *AIF1*, *FLOT1*, *NTF3*, *ODAD3*, *HOXA2*, *SEMA6D*, *PALM*, *IQUB*, *BBS10*, *B4GALT6*, *ACP4*, *RTN4RL1*, *KREMEN1*, *CEP250*, *PODXL*, *RHOD*, *PKD2*, *TMEM107*, *BBS1*, *NR2E1*, *TTC39C*, *BMPR2*, *CHODL*, *EMP3*, *VCL*, *DNAI2*, *IFT46*, *WNT7A*, *NYAP1*, *MEIG1*, *DCX*, *WRAP73*, *BMP7*, *RTN4R*, *UCHL1*, *GPRASP3*, *PLD1*, *RASGRF1*, *SEMA6A*, *GBX1*, *F2RL1*, *SEMA3E*, *GMNC*, *CFAP97D1*, *SEMA6C*, *DYNC2I2*, *TUB*, *SPOCK1*, *INPP5K*, *CCNO*, *EPHA7*, *SEMA3B*, *STMN1*, *P2RX7*, *FOLR3*, *CNTN1*, *MAP1B*, *CNR1*, *EFNB3*, *ODAD1*, *BDNF*, *CNTN5*, *EPHA3*, *ATCAY*, *ZNF365*, *P2RY12*, *NDP*, *SFRP2* |
| GO:0030030 | Cell projection organization | 311 | -1.48 | 3.49E-04 | 1.27E-03 | *BORCS7*, *FGD1*, *NEK1*, *LCA5*, *GFAP*, *TCTN2*, *SMO*, *SEMA4F*, *TPBG*, *CCDC113*, *PLPPR4*, *SPATA6*, *NEDD9*, *SPACA9*, *FUZ*, *DYNC2LI1*, *EFNA4*, *ENPP2*, *FOXJ1*, *EXT1*, *NTN4*, |

| | | | | | | |
|---|---|---|---|---|---|---|
| | | | | | | *PIERCE1*, *SFRP1*, *NTF4*, *INPP5J*, *CFAP58*, *SPEF2*, *ARHGEF25*, *CFAP20*, *TACSTD2*, *ARL6*, *SRCIN1*, *CFAP91*, *BTBD3*, *EHD2*, *TEKT2*, *SYT3*, *EPHB3*, *NFIB*, *PDGFA*, *AIF1*, *FLOT1*, *NTF3*, *ODAD3*, *HOXA2*, *SEMA6D*, *PALM*, *IQUB*, *BBS10*, *B4GALT6*, *ACP4*, *RTN4RL1*, *KREMEN1*, *CEP250*, *PODXL*, *RHOD*, *TCTN1*, *PKD2*, *TMEM107*, *BBS1*, *NR2E1*, *PARVG*, *TTC39C*, *BMPR2*, *CHODL*, *EMP3*, *VCL*, *DNAI2*, *IFT46*, *MKS1*, *WNT7A*, *NYAP1*, *MEIG1*, *DCX*, *WRAP73*, *BMP7*, *RTN4R*, *UCHL1*, *GPRASP3*, *PLD1*, *RASGRF1*, *SEMA6A*, *GBX1*, *F2RL1*, *SEMA3E*, *GMNC*, *CFAP97D1*, *SEMA6C*, *DYNC2I2*, *TUB*, *SPOCK1*, *INPP5K*, *CCNO*, *EPHA7*, *SEMA3B*, *STMN1*, *P2RX7*, *FOLR3*, *CNTN1*, *MAP1B*, *CNR1*, *EFNB3*, *ODAD1*, *BDNF*, *CNTN5*, *EPHA3*, *ATCAY*, *ZNF365*, *P2RY12*, *NDP*, *SFRP2* |
| GO:0001501 | Skeletal system development | 117 | -1.49 | 7.15E-03 | 1.59E-02 | *ACVR2B*, *KDR*, *BMP4*, *TMEM107*, *BBS1*, *CHAD*, *PAPPA2*, *HOXC4*, *BMPR2*, *RFLNA*, *SOX9*, *SMAD7*, *HOXC6*, *BGN*, *WNT7A*, *WNT11*, *HOXB8*, *BMP7*, *OSR2*, *FGF18*, *OGN*, *LEPR*, *TRPV4*, *MMP16*, *P2RX7*, *HOXB7*, *NFATC2*, *MYCN*, *HHIP*, *CCN3*, *LRRC17*, *SFRP2* |
| GO:0031175 | Neuron projection development | 167 | -1.51 | 1.74E-03 | 4.85E-03 | *EXT1*, *NTN4*, *SFRP1*, *NTF4*, *INPP5J*, *ARHGEF25*, *SRCIN1*, *BTBD3*, *SYT3*, *EPHB3*, *NFIB*, *FLOT1*, *NTF3*, *HOXA2*, *SEMA6D*, *B4GALT6*, *ACP4*, *RTN4RL1*, *KREMEN1*, *BBS1*, *NR2E1*, *BMPR2*, *CHODL*, *VCL*, *WNT7A*, *NYAP1*, *DCX*, *BMP7*, *RTN4R*, *UCHL1*, *GPRASP3*, *RASGRF1*, *SEMA6A*, *GBX1*, *SEMA3E*, *SEMA6C*, *SPOCK1*, *EPHA7*, *SEMA3B*, *STMN1*, *FOLR3*, *CNTN1*, *MAP1B*, *CNR1*, *EFNB3*, *BDNF*, *CNTN5*, *EPHA3*, *ATCAY*, *ZNF365*, *NDP*, *SFRP2* |
| GO:0048858 | Cell projection morphogenesis | 113 | -1.72 | 3.90E-04 | 1.37E-03 | *SMO*, *SEMA4F*, *TPBG*, *PLPPR4*, *EFNA4*, *ENPP2*, *EXT1*, *NTN4*, *NTF4*, *ARHGEF25*, *SRCIN1*, *BTBD3*, *SYT3*, *EPHB3*, *NFIB*, *FLOT1*, *NTF3*, *HOXA2*, *SEMA6D*, *B4GALT6*, *NR2E1*, *BMPR2*, *CHODL*, *VCL*, *WNT7A*, *NYAP1*, *DCX*, *BMP7*, *RTN4R*, *UCHL1*, *GPRASP3*, *SEMA6A*, *GBX1*, *SEMA3E*, *SEMA6C*, *EPHA7*, *SEMA3B*, *CNTN1*, *MAP1B*, *EFNB3*, *BDNF*, *CNTN5*, *EPHA3*, *ZNF365*, *NDP* |
| GO:0120039 | Plasma membrane bounded cell projection morphogenesis | 113 | -1.72 | 3.90E-04 | 1.37E-03 | *SMO*, *SEMA4F*, *TPBG*, *PLPPR4*, *EFNA4*, *ENPP2*, *EXT1*, *NTN4*, *NTF4*, *ARHGEF25*, *SRCIN1*, *BTBD3*, *SYT3*, *EPHB3*, *NFIB*, *FLOT1*, *NTF3*, *HOXA2*, *SEMA6D*, *B4GALT6*, *NR2E1*, *BMPR2*, *CHODL*, *VCL*, *WNT7A*, *NYAP1*, *DCX*, *BMP7*, *RTN4R*, *UCHL1*, *GPRASP3*, *SEMA6A*, *GBX1*, |

| | | | | | | |
|---|---|---|---|---|---|---|
| | | | | | | *SEMA3E*, *SEMA6C*, *EPHA7*, *SEMA3B*, *CNTN1*, *MAP1B*, *EFNB3*, *BDNF*, *CNTN5*, *EPHA3*, *ZNF365*, *NDP* |
| GO:0048812 | Neuron projection morphogenesis | 109 | -1.75 | 1.79E-04 | 6.77E-04 | *SMO*, *SEMA4F*, *TPBG*, *PLPPR4*, *EFNA4*, *EXT1*, *NTN4*, *NTF4*, *ARHGEF25*, *SRCIN1*, *BTBD3*, *SYT3*, *EPHB3*, *NFIB*, *FLOT1*, *NTF3*, *HOXA2*, *SEMA6D*, *B4GALT6*, *NR2E1*, *BMPR2*, *CHODL*, *VCL*, *WNT7A*, *NYAP1*, *DCX*, *BMP7*, *RTN4R*, *UCHL1*, *GPRASP3*, *SEMA6A*, *GBX1*, *SEMA3E*, *SEMA6C*, *EPHA7*, *SEMA3B*, *CNTN1*, *MAP1B*, *EFNB3*, *BDNF*, *CNTN5*, *EPHA3*, *ZNF365*, *NDP* |

Abbreviations: GO:BP: Biological Processes in Gene Ontology; FDR: False discovery rate; MAPK: Mitogen-activated protein kinase; NES: Normalized enrichment score.

73

**Supplementary Table S3. Significantly enriched pathways from the GSEA result of the KEGG database, High-glucose infection versus High-glucose control.** Abbreviations: GSEA: Gene set enrichment analysis; KEGG: Kyoto Encyclopedia of Genes and Genomes.

| KEGG ID | Description | Set Size | NES | P-value | FDR | Core enrichment gene |
|---|---|---|---|---|---|---|
| ssc05164 | Influenza A | 127 | 2.888 | 1.14E-23 | 1.96E-21 | *CXCL10, IFNG, CCL2, IL1B, MX1, CXCL8, RSAD2, IL6, IL1A, IRF7, STAT1, OAS1, FAS, RIGI, OAS2, SOCS3, IFIH1, SLA-DOA, TNFSF10, EIF2AK2, SLA-DQA1, SLA-DRA, IRF9, SLA-DOB, SLA-DQB1, CCL5, RNASEL, STAT2, SLA-DRB1, NFKBIB, IFNAR2, SLA-DMB, NFKBIA, PML, FASLG, ADAR, CYCS, CASP3, BAK1, SLA-DMA, LOC100624226, IKBKE, TRAF3, CASP1, ICAM1, TRIM25, KPNA2, CASP8, TICAM1, MYD88, TLR4, TNFRSF1A, EIF2S1, NFKB1, IFNGR1, IRAK4, DNAJC3, TLR3, JAK2, IFNGR2, IKBKB* |
| ssc05168 | Herpes simplex virus 1 infection | 140 | 2.887 | 1.20E-24 | 4.14E-22 | *IFNG, CCL2, IL1B, TAP1, IL6, BIRC3, IRF7, STAT1, OAS1, FAS, TAP2, RIGI, OAS2, SOCS3, BST2, IFIH1, SLA-DOA, EIF2AK2, SLA-DQA1, SLA-DRA, SLA-3, IRF9, SLA-DOB, SLA-DQB1, CCL5, SLA-7, RNASEL, TAPBP, STAT2, SLA-DRB1, B2M, CD74, IFNAR2, SLA-DMB, C3, NFKBIA, TNFSF14, PML, FASLG, TAPBPL, CYCS, CASP3, BAK1, SLA-DMA, EIF4EBP1, TRAF2, IKBKE, TRAF3, PDIA3, LOC100517129, CASP8, TICAM1, MYD88, ZC3HAV1, CGAS, TNFRSF1A, CALR, DAXX, ITGA5, TLR2, EIF2S1, STING1, NFKB1, IFNGR1, IRAK4, SLA-8, TP53, TLR3, JAK2, IFNGR2, IKBKB* |
| ssc04668 | TNF signaling pathway | 104 | 2.738 | 5.40E-18 | 6.21E-16 | *CXCL10, SELE, CCL2, IRF1, AMCF-II, CXCL2, IL1B, IL6, BIRC3, FAS, SOCS3, CX3CL1, MAP3K8, BCL3, TNFAIP3, JUNB, RHBDF2, VCAM1, IL15, MMP9, CASP7, NOD2, CCL5, CASP10, CSF1, FOS, NFKBIA, LIF, CSF2, CASP3, MLKL, PTGS2, PGAM5, TRAF2, TRAF3, ICAM1, CEBPB, CASP8, ADAM17, CFLAR, TNFRSF1A, RIPK3, CREB3L1, NFKB1, TNFRSF1B* |
| ssc04657 | IL-17 signaling pathway | 70 | 2.667 | 5.68E-15 | 2.80E-13 | *CXCL10, IFNG, CCL2, AMCF-II, CXCL2, IL1B, CXCL8, S100A8, S100A9, IL6, CSF3, CCL11, MMP1, FOSL1, TNFAIP3, MMP9, MUC5AC, LCN2, MMP13, FOS, NFKBIA, HSP90B1, CSF2, CASP3, PTGS2, TRAF2, IKBKE, TRAF3, CEBPB, CASP8* |

| | | | | | | |
|---|---|---|---|---|---|---|
| ssc05169 | Epstein-Barr virus infection | 168 | 2.592 | 3.34E-17 | 2.88E-15 | *CXCL10, ISG15, TAP1, IL6, IRF7, STAT1, OAS1, FAS, TAP2, RIGI, OAS2, RELB, SLA-DOA, TNFAIP3, EIF2AK2, SLA-DQA1, SLA-DRA, CDKN1A, SLA-3, IRF9, SLA-DOB, CD40, SLA-DQB1, GADD45A, SLA-7, TAPBP, STAT2, SLA-DRB1, NFKBIE, B2M, NFKBIB, NFKB2, IFNAR2, SLA-DMB, NFKBIA, GADD45G, STAT3, TAPBPL, CYCS, CASP3, BAK1, SLA-DMA, TRAF2, IKBKE, TRAF3, PDIA3, ICAM1, CASP8, MYD88, JAK3, PSMD14, CALR, MYC, TLR2, GADD45B, NFKB1, IRAK4, SLA-8, PSMD12, TP53, PSMD3, IKBKB* |
| ssc04620 | Toll-like receptor signaling pathway | 87 | 2.588 | 2.02E-13 | 7.76E-12 | *CXCL9, CXCL11, CXCL10, CCL3L1, IL1B, CXCL8, CCL4, IL6, IRF7, STAT1, LBP, MAP3K8, IRF9, CD40, TICAM2, CCL5, STAT2, IFNAR2, FOS, NFKBIA, IKBKE, TRAF3, CD14, CASP8, TICAM1, MYD88, TLR4, TLR2, TIRAP, NFKB1, IRAK4, LY96, TLR3, IKBKB* |
| ssc04060 | Cytokine-cytokine receptor interaction | 170 | 2.574 | 1.04E-16 | 7.17E-15 | *CXCL9, CXCL11, CXCL10, IFNG, CCL19, CCL2, AMCF-II, CXCL2, CCL3L1, CXCL13, IL1B, CXCL8, CCL4, IL27, IL6, IL1A, TNFSF13B, FAS, CSF3, CCL26, CCL11, CCL8, CX3CL1, TNFSF10, IL1R2, IL15, LOC110260995, IL4R, CSF3R, CD40, PPBP, CCL5, ACKR4, GDF15, CSF1, XCL1, IL10, IFNAR2, IL1RAP, CNTFR, IL2RB, OSMR, TNFSF14, IL1R1, LIF, CCR1, TNFSF13, FASLG, CSF2, CCL28, CXCL16, IL2RA, CXCR6, ACVR1B, LOC100624226, IL10RB, CSF2RB, IL18RAP, TNFSF9, TGFB3, TNFRSF12A, TNFRSF1A, RELL1, IL6ST, IL21R, BMP2, IFNGR1, RELT, TNFRSF1B, LOC100516039* |
| ssc04612 | Antigen processing and presentation | 47 | 2.569 | 3.29E-12 | 9.46E-11 | *IFNG, TAP1, TAP2, PSME2, CTSL, SLA-DOA, SLA-DQA1, SLA-DRA, SLA-3, SLA-DOB, SLA-DQB1, PSME1, SLA-7, TAPBP, SLA-DRB1, B2M, HSPA6, CD74, SLA-DMB, HSPA5, LOC100523789, TAPBPL, RFX5, SLA-DMA, PDIA3, IFI30, HSPA8, CD8B, CALR, HSP90AA1, SLA-8, HSPA4, CANX* |
| ssc05323 | Rheumatoid arthritis | 65 | 2.552 | 2.83E-12 | 8.89E-11 | *IFNG, CCL2, AMCF-II, CXCL2, CCL3L1, IL1B, CXCL8, IL6, IL1A, TNFSF13B, MMP1, CTSL, SLA-DOA, SLA-DQA1, SLA-DRA, IL15, CTLA4, SLA-DOB, SLA-DQB1, CCL5, SLA-DRB1, CSF1, SLA-DMB, FOS, TNFSF13, CSF2, TCIRG1, SLA-DMA, ICAM1, TGFB3, TLR4* |
| ssc05162 | Measles | 108 | 2.538 | 3.75E-13 | 1.29E-11 | *IL1B, MX1, SLAMF1, IL6, IL1A, IRF7, STAT1, OAS1, FAS, RIGI, OAS2, IFIH1, TNFAIP3, EIF2AK2, IRF9, FCGR2B, STAT2, STAT5A, NFKBIB, HSPA6, IFNAR2, FOS, NFKBIA, IL2RB,* |

| | | | | | | |
|---|---|---|---|---|---|---|
| | | | | | | *FASLG, ADAR, STAT3, CYCS, CASP3, BAK1, IL2RA, IKBKE, TRAF3, CASP8, MYD88, JAK3, HSPA8, TLR4, TLR2, EIF2S1, NFKB1, IRAK4, TP53, IL2RG, CD46, IKBKB* |
| ssc04621 | NOD-like receptor signaling pathway | 127 | 2.525 | 1.13E-13 | 4.89E-12 | *GBP1, GBP2, CCL2, CXCL2, IL1B, CXCL8, IL6, BIRC3, IRF7, STAT1, OAS1, OAS2, LOC100522887, TNFAIP3, IRF9, NOD2, CCL5, RNASEL, STAT2, NFKBIB, IFNAR2, NFKBIA, NAMPT, GSDMD, TRAF2, IKBKE, TRAF3, CASP1, RIPK2, CASP8, TICAM1, MYD88, TXNIP, TLR4, TANK, RIPK3, TRPM2, NLRP12, STING1, NFKB1, IRAK4, HSP90AA1, IKBKB, TXN* |
| ssc05171 | Coronavirus disease - COVID-19 | 181 | 2.497 | 2.01E-15 | 1.15E-13 | *CXCL10, CCL2, IL1B, MX1, ISG15, CXCL8, IL6, STAT1, OAS1, RIGI, OAS2, CSF3, MMP1, IFIH1, EIF2AK2, IRF9, CFB, F13A1, SELP, STAT2, FGB, NFKBIB, C2, IFNAR2, FOS, C3, NFKBIA, C4A, ADAR, STAT3, CSF2, IKBKE, TRAF3, CASP1, MYD88, LOC100515049, TLR4, ADAM17, CGAS, TNFRSF1A, IL6ST, RPL13A, TLR2, STING1, NFKB1, IRAK4* |
| ssc04061 | Viral protein interaction with cytokine and cytokine receptor | 60 | 2.478 | 2.27E-10 | 4.36E-09 | *CXCL9, CXCL11, CXCL10, CCL19, CCL2, AMCF-II, CXCL2, CCL3L1, CXCL13, CXCL8, CCL4, IL6, CCL26, CCL11, CCL8, CX3CL1, TNFSF10, PPBP, CCL5, ACKR4, CSF1, XCL1, IL10, IL2RB, TNFSF14, CCR1, CCL28, IL2RA, LOC100624226, IL10RB, IL18RAP, TNFRSF1A, IL6ST* |
| ssc05160 | Hepatitis C | 120 | 2.445 | 1.96E-11 | 4.51E-10 | *CXCL10, IFNG, MX1, RSAD2, IFIT1, IRF7, STAT1, OAS1, FAS, RIGI, OAS2, SOCS3, EIF2AK2, CDKN1A, IRF9, RNASEL, STAT2, IFNAR2, NFKBIA, FASLG, STAT3, CYCS, CASP3, BAK1, TRAF2, IKBKE, TRAF3, CASP8, TICAM1, CFLAR, TNFRSF1A, MYC, EIF2S1, NFKB1, TP53, TLR3, IKBKB* |
| ssc05321 | Inflammatory bowel disease | 45 | 2.427 | 2.61E-09 | 3.92E-08 | *IFNG, IL1B, IL6, IL1A, STAT1, SLA-DOA, SLA-DQA1, SLA-DRA, SLA-DOB, IL4R, SLA-DQB1, NOD2, SLA-DRB1, IL10, SLA-DMB, STAT3, SLA-DMA, IL18RAP, TGFB3, TLR4, TBX21, IL21R, TLR2, NFKB1, IFNGR1, IL2RG, IFNGR2, SMAD3* |
| ssc04623 | Cytosolic DNA-sensing pathway | 59 | 2.410 | 3.05E-09 | 4.38E-08 | *CXCL10, IL1B, CCL4, IL6, IRF7, SAMHD1, ZBP1, RIGI, CASP7, CCL5, NFKBIB, NFKBIA, ADAR, CASP3, MLKL, GSDMD, TREX1, IKBKE, CASP1, POLR3D, CASP8, CGAS, RIPK3, STING1, NFKB1, POLR3E, IKBKB* |
| ssc05152 | Tuberculosis | 137 | 2.408 | 1.74E-11 | 4.30E-10 | *IFNG, CLEC4E, IL1B, LOC110258711, IL6, IL1A, STAT1, LBP, SLA-DOA, SLA-DQA1, SLA-DRA, SLA-DOB, FCGR2B, SLA-DQB1, NOD2, CASP10, SLA-DRB1, CD74, IL10, SLA-DMB, C3,* |

| | | | | | | |
|---|---|---|---|---|---|---|
| | | | | | | *CYP27B1, CYCS, CASP3, RFX5, PLK3, TCIRG1, CEBPG, SLA-DMA, HSPA9, IL10RB, CD14, CEBPB, RIPK2, HSPD1, CASP8, MYD88, TGFB3, CLEC7A, TLR4, TNFRSF1A, TLR2, MRC1, TIRAP, NFKB1, IFNGR1, IRAK4, VDR, JAK2, IFNGR2* |
| ssc04622 | RIG-I-like receptor signaling pathway | 54 | 2.406 | 2.43E-09 | 3.81E-08 | *CXCL10, ISG15, CXCL8, IRF7, DHX58, RIGI, IFIH1, ZNFX1, CASP10, NFKBIB, NFKBIA, ADAR, TRAF2, IKBKE, TRAF3, TRIM25, CASP8, TANK, STING1, NFKB1* |
| ssc04659 | Th17 cell differentiation | 89 | 2.381 | 1.81E-10 | 3.67E-09 | *IFNG, IL1B, IL27, IL6, STAT1, SLA-DOA, SLA-DQA1, SLA-DRA, FOXS1, SLA-DOB, IL4R, SLA-DQB1, SLA-DRB1, NFKBIE, STAT5A, NFKBIB, IL1RAP, SLA-DMB, FOS, NFKBIA, IL2RB, IL1R1, STAT3, SLA-DMA, IL2RA, JAK3, TBX21, IL6ST, IL21R, HIF1A, NFKB1, IFNGR1, HSP90AA1, JAK2, IL2RG, IFNGR2, IKBKB, SMAD3* |
| ssc04217 | Necroptosis | 109 | 2.376 | 6.52E-10 | 1.12E-08 | *IFNG, IL1B, IL1A, BIRC3, STAT1, ZBP1, FAS, TNFAIP3, TNFSF10, EIF2AK2, IRF9, TICAM2, STAT2, STAT5A, IFNAR2, FASLG, STAT3, FTH1, MLKL, PGAM5, LOC100624226, TRAF2, GLUL, CASP1, CASP8, TICAM1, JAK3, TLR4, CHMP7, CFLAR, TNFRSF1A, RIPK3, IFNGR1, HSP90AA1, TLR3, JAK2, PPID, IFNGR2* |
| ssc05140 | Leishmaniasis | 59 | 2.374 | 1.08E-08 | 1.43E-07 | *IFNG, IL1B, IL1A, STAT1, SLA-DOA, SLA-DQA1, SLA-DRA, SLA-DOB, SLA-DQB1, SLA-DRB1, NFKBIB, IL10, SLA-DMB, FOS, C3, NFKBIA, SLA-DMA, PTGS2, MYD88, TGFB3, TLR4, TLR2, NFKB1, IFNGR1, IRAK4, JAK2, IFNGR2* |
| ssc04064 | NF-kappa B signaling pathway | 87 | 2.373 | 1.52E-09 | 2.49E-08 | *CCL19, CXCL2, IL1B, CXCL8, CCL4, BIRC3, TNFSF13B, RIGI, LBP, RELB, TNFAIP3, VCAM1, CD40, TICAM2, GADD45A, NFKB2, NFKBIA, TNFSF14, IL1R1, GADD45G, PTGS2, TRAF2, TRAF3, ICAM1, TRIM25, CD14, TICAM1, MYD88, TLR4, BCL2A1, CFLAR, TNFRSF1A, TIRAP, GADD45B, NFKB1, IRAK4, LY96, IKBKB* |
| ssc05417 | Lipid and atherosclerosis | 171 | 2.346 | 4.77E-12 | 1.26E-10 | *SELE, CCL2, CXCL2, CCL3L1, IL1B, CXCL8, IL6, IRF7, FAS, LBP, MMP1, TNFSF10, SOD2, VCAM1, MMP9, CD40, CASP7, SELP, TICAM2, CCL5, HSPA6, FOS, NFKBIA, HSPA5, FASLG, HSP90B1, STAT3, CYCS, CASP3, LOC100624226, TRAF2, DDIT3, IKBKE, TRAF3, OLR1, CASP1, ICAM1, CD14, HSPD1, CASP8, TICAM1, MYD88, HSPA8, TLR4, TNFRSF1A, VLDLR, TANK, TLR2, TIRAP, EIF2S1, NFKB1, IRAK4, HSP90AA1, LY96, TP53, JAK2, IKBKB* |

| | | | | | | |
|---|---|---|---|---|---|---|
| ssc04141 | Protein processing in endoplasmic reticulum | 149 | 2.344 | 4.10E-11 | 8.85E-10 | *HSPA6, HYOU1, DNAJB11, HSPA5, HSP90B1, BAK1, PDIA4, HSPH1, TRAF2, DDIT3, WFS1, PDIA3, NGLY1, SEC23B, DERL3, HERPUD1, CKAP4, SEC24D, SELENOS, DNAJC1, HSPA8, PREB, CALR, SEC24A, MOGS, OSTC, DNAJA1, LMAN1, UBE2D1, SEC61A1, SEC61B, EIF2S1, MAN1B1, SSR3, YOD1, RPN1, PDIA6, HSP90AA1, DNAJC3, RPN2, SSR1, EDEM1, SEC13, DNAJA2, UBE2D3, UBQLN1, CANX, UGGT1, P4HB, HSPBP1, ERN1, STT3A, DAD1, TRAM1, DNAJB1, TMEM258, RAD23A, VCP, XBP1, DERL2, STT3B, SAR1B, RRBP1, SEC63, PLAA, BAG1, ATF6, SEL1L, UBE2J2, UBE2D2, RNF5, GANAB, UBXN4, ERO1B, MAGT1, EIF2AK3, EDEM2, SEC24C, UBXN8, NFE2L2, NPLOC4* |
| ssc03050 | Proteasome | 40 | 2.339 | 2.64E-07 | 2.76E-06 | *IFNG, PSMB9, PSME2, PSMB8, PSMB10, PSME1, PSMA5, PSMA2, PSMA3, PSMB3, PSMA1, PSMD14, PSMB2, PSMB4, PSMA4, PSMD12, PSMD3, PSMD8, PSMC4, PSMB1, PSMD7, PSMD11, PSMC6, PSMD1, PSMC1, PSME3, PSMC2, PSMB6, PSMD13, PSMF1, PSMD6, PSMD2, PSMD4* |
| ssc05133 | Pertussis | 63 | 2.330 | 1.27E-08 | 1.62E-07 | *IRF1, AMCF-II, IL1B, CXCL8, IL6, IL1A, IRF8, CASP7, TICAM2, C2, IL10, FOS, C3, C4A, SFTPA1, CASP3, SERPING1, CASP1, CD14, TICAM1, C4BPB, MYD88, TLR4, ITGA5, TIRAP, NFKB1, IRAK4, LY96* |
| ssc05332 | Graft-versus-host disease | 23 | 2.323 | 2.56E-08 | 3.15E-07 | *IFNG, IL1B, IL6, IL1A, FAS, SLA-DOA, SLA-DQA1, SLA-DRA, SLA-3, SLA-DOB, SLA-DQB1, SLA-7, SLA-DRB1, SLA-DMB, LOC100523789, FASLG, SLA-DMA, PRF1* |
| ssc04630 | JAK-STAT signaling pathway | 110 | 2.288 | 5.12E-09 | 7.06E-08 | *IFNG, IL27, IL6, STAT1, SOCS3, CSF3, SOCS1, IL15, CDKN1A, IRF9, PIM1, IL4R, CSF3R, STAT2, STAT5A, IL10, IFNAR2, CNTFR, IL2RB, OSMR, CISH, LIF, STAT3, CSF2, IL2RA, PTPN2, IL10RB, CSF2RB, JAK3, MCL1, IL6ST, IL21R, MYC, IFNGR1, JAK2, IL2RG, IFNGR2* |
| ssc03008 | Ribosome biogenesis in eukaryotes | 69 | 2.255 | 1.02E-07 | 1.14E-06 | *GNL3, EMG1, NHP2, POP1, GAR1, FBL, WDR43, NOB1, DKC1, WDR75, UTP4, SPATA5, GTPBP4, WDR3, RBM28, NOP58, NOL6, MPHOSPH10, IMP3, BMS1, TBL3, RCL1, NAT10, NOP56, XRN1, UTP18, GNL2, RAN, PWP2, RRP7A, CSNK2A1, EIF6, RPP40, UTP6, UTP15, AK6, NXT1, HEATR1, WDR36, LSG1, NMD3, FCF1, IMP4* |
| ssc05134 | Legionellosis | 44 | 2.250 | 8.73E-07 | 7.53E-06 | *CXCL2, IL1B, CXCL8, IL6, CASP7, NFKB2, HSPA6, C3, NFKBIA, CYCS, CASP3, CASP1, CD14, HSPD1, CASP8, MYD88, HSPA8, TLR4, TLR2, NFKB1, SEC22B, BNIP3, RAB1A* |

| | | | | | | |
|---|---|---|---|---|---|---|
| ssc04672 | Intestinal immune network for IgA production | 34 | 2.235 | 2.47E-06 | 2.03E-05 | *MADCAM1, IL6, TNFSF13B, PIGR, SLA-DOA, SLA-DQA1, SLA-DRA, IL15, SLA-DOB, CD40, SLA-DQB1, SLA-DRB1, IL10, SLA-DMB, TNFSF13, CCL28, SLA-DMA* |
| ssc04062 | Chemokine signaling pathway | 158 | 2.223 | 3.69E-10 | 6.70E-09 | *CXCL9, CXCL11, CXCL10, CCL19, CCL2, AMCF-II, CXCL2, CCL3L1, CXCL13, CXCL8, CCL4, STAT1, CCL26, CCL11, CCL8, CX3CL1, PPBP, CCL5, STAT2, NFKBIB, XCL1, NFKBIA, CCR1, STAT3, CCL28, CXCL16, CXCR6, GNG10, HCK, JAK3, FGR* |
| ssc05330 | Allograft rejection | 21 | 2.198 | 6.84E-07 | 6.38E-06 | *IFNG, FAS, SLA-DOA, SLA-DQA1, SLA-DRA, SLA-3, SLA-DOB, CD40, SLA-DQB1, SLA-7, SLA-DRB1, IL10, SLA-DMB, FASLG, SLA-DMA, PRF1* |
| ssc04940 | Type I diabetes mellitus | 25 | 2.187 | 2.74E-06 | 2.15E-05 | *IFNG, IL1B, IL1A, FAS, SLA-DOA, SLA-DQA1, SLA-DRA, SLA-3, SLA-DOB, SLA-DQB1, SLA-7, SLA-DRB1, SLA-DMB, FASLG, SLA-DMA, PRF1, HSPD1* |
| ssc05144 | Malaria | 39 | 2.171 | 7.04E-06 | 4.96E-05 | *IFNG, SELE, ACKR1, CCL2, IL1B, CXCL8, IL6, CSF3, THBS1, VCAM1, CD40, SELP, IL10, ICAM1, MYD88, TGFB3, TLR4, KLRK1, TLR2, THBS2* |
| ssc05320 | Autoimmune thyroid disease | 26 | 2.168 | 2.06E-05 | 1.37E-04 | *FAS, SLA-DOA, TG, SLA-DQA1, SLA-DRA, SLA-3, CTLA4, SLA-DOB, CD40, SLA-DQB1, SLA-7, SLA-DRB1, IL10, SLA-DMB, FASLG, SLA-DMA, PRF1* |
| ssc05145 | Toxoplasmosis | 94 | 2.160 | 4.25E-07 | 4.19E-06 | *IFNG, BIRC3, STAT1, SOCS1, SLA-DOA, SLA-DQA1, SLA-DRA, PPIF, SLA-DOB, CD40, SLA-DQB1, SLA-DRB1, NFKBIB, HSPA6, IL10, SLA-DMB, NFKBIA, STAT3, CYCS, CASP3, SLA-DMA, IL10RB, CASP8, MYD88, TGFB3, HSPA8, TLR4, TNFRSF1A, TLR2, NFKB1, IFNGR1, IRAK4, LY96, JAK2, IFNGR2, IKBKB* |
| ssc05142 | Chagas disease | 92 | 2.158 | 4.58E-07 | 4.39E-06 | *IFNG, CCL2, CCL3L1, IL1B, CXCL8, IL6, SERPINE1, FAS, CCL5, IL10, FOS, C3, NFKBIA, FASLG, CASP8, TICAM1, MYD88, TGFB3, LOC100515049, TLR4, CFLAR, TNFRSF1A, CALR, TLR2, NFKB1, IFNGR1, IRAK4, IFNGR2, IKBKB* |
| ssc04658 | Th1 and Th2 cell differentiation | 73 | 2.142 | 9.48E-07 | 7.98E-06 | *IFNG, STAT1, SLA-DOA, SLA-DQA1, SLA-DRA, SLA-DOB, IL4R, SLA-DQB1, SLA-DRB1, NFKBIE, STAT5A, NFKBIB, SLA-DMB, FOS, NFKBIA, IL2RB, SLA-DMA, IL2RA, JAK3, TBX21, NFKB1, IFNGR1, DLL4, JAK2, IL2RG, IFNGR2, IKBKB* |
| ssc05310 | Asthma | 14 | 2.127 | 2.58E-06 | 2.07E-05 | *FCER1A, CCL11, SLA-DOA, SLA-DQA1, SLA-DRA, SLA-DOB, CD40, SLA-DQB1, SLA-DRB1, IL10, SLA-DMB, SLA-DMA* |

| | | | | | | |
|---|---|---|---|---|---|---|
| ssc05161 | Hepatitis B | 141 | 2.101 | 1.27E-07 | 1.37E-06 | *CXCL8, IL6, IRF7, STAT1, FAS, RIGI, IFIH1, CDKN1A, EGR2, MMP9, TICAM2, CASP10, STAT2, STAT5A, FOS, NFKBIA, FASLG, STAT3, CYCS, CASP3, IKBKE, TRAF3, CASP8, TICAM1, MYD88, TGFB3, JAK3, TLR4, CREB3L1, MYC, TLR2, TIRAP, NFKB1, IRAK4, TP53, TLR3, JAK2, IKBKB, SMAD3* |
| ssc05167 | Kaposi sarcoma-associated herpesvirus infection | 156 | 2.091 | 3.85E-08 | 4.58E-07 | *CXCL2, CXCL8, IL6, IRF7, STAT1, FAS, CLEC2B, EIF2AK2, CDKN1A, SLA-3, IRF9, SLA-7, ZFP36, STAT2, IFNAR2, FOS, C3, NFKBIA, CCR1, STAT3, CSF2, CYCS, CASP3, BAK1, PTGS2, TRAF2, IKBKE, TRAF3, ICAM1, GNG10, CASP8, TICAM1, HCK, TNFRSF1A, IL6ST, HIF1A, MYC, NFKB1, IFNGR1, SLA-8, TP53, TLR3, JAK2, IKBKB, ANGPT2, LOC100621431* |
| ssc04936 | Alcoholic liver disease | 109 | 2.085 | 3.28E-06 | 2.51E-05 | *CXCL2, IL1B, CXCL8, IL6, FAS, LBP, CPT1C, C2, C3, NFKBIA, C4A, ADH1C, FASLG, CASP3, IKBKE, TRAF3, CD14, CASP8, TICAM1, MYD88, TLR4, ALDH9A1, TNFRSF1A, TIRAP, NFKB1, IRAK4, LY96, IKBKB, PPARGC1A, RIPK1, IRAK1, IL17RA, MAP2K3, TAB3, TRA2B* |
| ssc04625 | C-type lectin receptor signaling pathway | 90 | 2.063 | 7.58E-06 | 5.23E-05 | *CLEC4E, IRF1, IL1B, LOC110258711, IL6, STAT1, RELB, BCL3, IRF9, EGR2, STAT2, NFKB2, IL10, NFKBIA, PLK3, PTGS2, IKBKE, CASP1, CASP8, CLEC7A* |
| ssc04210 | Apoptosis | 114 | 2.044 | 8.04E-07 | 7.11E-06 | *BIRC3, FAS, PMAIP1, CTSL, TNFSF10, CASP7, GADD45A, CASP10, FOS, NFKBIA, FASLG, GADD45G, CYCS, CASP3, BAK1, LOC100624226, TRAF2, DDIT3, CSF2RB, PRF1, CASP8, LOC100127131, MCL1, BCL2A1, CFLAR, TNFRSF1A, DAXX, GADD45B, EIF2S1, NFKB1, TUBA4A, TP53, IKBKB, LMNB2, RIPK1, ERN1* |
| ssc05150 | Staphylococcus aureus infection | 32 | 2.038 | 8.81E-05 | 5.07E-04 | *SLA-DOA, SLA-DQA1, SLA-DRA, CFB, SLA-DOB, FCGR2B, SLA-DQB1, SELP, SLA-DRB1, C2, IL10, SLA-DMB, C3, C4A, SLA-DMA, ICAM1, KRT18, CFI* |
| ssc04932 | Non-alcoholic fatty liver disease | 135 | 2.004 | 4.28E-06 | 3.21E-05 | *IL1B, CXCL8, IL6, IL1A, FAS, SOCS3, CASP7, FOS, FASLG, CYCS, CASP3, TRAF2, DDIT3, CASP8, TNFRSF1A, EIF2S1, NFKB1, MLXIPL, IKBKB, NDUFB2, NDUFB3, COX7A2, ERN1, IL6R, COX5A, UQCRC1, NDUFAB1, COX6B, XBP1, COX5B, NDUFV2, UQCRQ, NDUFA6, LOC100522678, NDUFA9, COX7B, BID, SDHD, RELA, EIF2AK3, UQCR11, SREBF1, NDUFB1, NDUFB4* |

| | | | | | | |
|---|---|---|---|---|---|---|
| ssc05322 | Systemic lupus erythematosus | 48 | 1.987 | 1.57E-04 | 8.32E-04 | *IFNG, SLA-DOA, SLA-DQA1, SLA-DRA, SLA-DOB, CD40, SLA-DQB1, TRIM21, SLA-DRB1, C2, IL10, SLA-DMB, C3, C4A, SLA-DMA* |
| ssc05235 | PD-L1 expression and PD-1 checkpoint pathway in cancer | 81 | 1.975 | 3.82E-05 | 2.31E-04 | *IFNG, BATF2, CD274, STAT1, BATF3, TICAM2, NFKBIE, NFKBIB, FOS, NFKBIA, STAT3, TICAM1, MYD88, TLR4, HIF1A, TLR2, TIRAP, NFKB1, IFNGR1, JAK2, IFNGR2, IKBKB* |
| ssc05163 | Human cytomegalovirus infection | 190 | 1.967 | 3.09E-07 | 3.14E-06 | *CCL2, CCL3L1, IL1B, TAP1, CXCL8, CCL4, IL6, FAS, TAP2, CX3CL1, CDKN1A, SLA-3, CCL5, SLA-7, TAPBP, B2M, NFKBIA, IL1R1, CCR1, FASLG, STAT3, TAPBPL, CYCS, CASP3, BAK1, PTGS2, EIF4EBP1, TRAF2, PDIA3, IL10RB, GNG10, CASP8, CGAS, TNFRSF1A, CALR, CREB3L1, MYC, STING1, NFKB1, SLA-8, TP53, GNA13, IKBKB* |
| ssc04380 | Osteoclast differentiation | 115 | 1.918 | 2.69E-05 | 1.72E-04 | *IFNG, IL1B, IL1A, STAT1, SOCS3, LOC100622460, SOCS1, FOSL1, RELB, JUNB, IRF9, FCGR2B, STAT2, NFKB2, CSF1, IFNAR2, FOS, NFKBIA, IL1R1, TRAF2, OSCAR, TNFRSF1A, LOC100522225, NFKB1, IFNGR1, IFNGR2, IKBKB* |
| ssc01230 | Biosynthesis of amino acids | 61 | 1.913 | 2.18E-04 | 1.14E-03 | *PSAT1, CBS, ASNS, SHMT2, ARG1, PSPH, PYCR1, PHGDH, MAT1A, GPT2, GLUL, SHMT1, PGK1, PFKP, ARG2, GOT1, PRPS1, ENO1, BCAT1, ASS1, ALDH18A1* |
| ssc04145 | Phagosome | 115 | 1.902 | 3.76E-05 | 2.31E-04 | *TAP1, TAP2, CTSL, SLA-DOA, THBS1, SLA-DQA1, SLA-DRA, SLA-3, SLA-DOB, FCGR2B, SLA-DQB1, SLA-7, SLA-DRB1, SLA-DMB, C3, TFRC, SFTPA1, TCIRG1, SLA-DMA, OLR1, CD14, LOC100127131, CLEC7A, TLR4, CALR, ITGA5, TLR2, THBS2, MRC1, SEC61A1, SEC61B, SEC22B, TUBA4A, SLA-8, SFTPD* |
| ssc00970 | Aminoacyl-tRNA biosynthesis | 43 | 1.893 | 5.07E-04 | 2.43E-03 | *WARS1, CARS1, MARS1, GARS1, AARS1, SARS1, TARS1, YARS1, FARSA, NARS1, VARS1, IARS1, EPRS1, FARSB, YARS2, MARS2, RARS1, LARS1* |
| ssc05143 | African trypanosomiasis | 28 | 1.882 | 1.89E-03 | 7.94E-03 | *IFNG, IDO1, SELE, IL1B, IL6, FAS, VCAM1, IL10, FASLG, ICAM1, MYD88* |
| ssc05166 | Human T-cell leukemia virus 1 infection | 194 | 1.868 | 6.56E-06 | 4.72E-05 | *IL6, FOSL1, RELB, SLA-DOA, IL1R2, SLA-DQA1, SLA-DRA, IL15, CDKN1A, SLA-3, EGR2, SLA-DOB, CD40, SLA-DQB1, SLA-7, ZFP36, SLA-DRB1, STAT5A, B2M, NFKB2, SLA-DMB, FOS, NFKBIA, IL2RB, IL1R1, MSX1, CSF2, SLA-DMA, IL2RA, EGR1, ICAM1, TGFB3, JAK3,* |

| | | | | | | |
|---|---|---|---|---|---|---|
| | | | | | | TNFRSF1A, CALR, RANBP1, CREB3L1, MYC, NFKB1, SLA-8, TP53, IL2RG, IKBKB, SMAD3, KAT2A, SLC25A5, CANX, CDC26, LTBR, ETS2, E2F3, RAN, MSX2, NRP1, TBPL1, JAK1, CREB3L4, TSPO, POLB, CCND2 |
| ssc05418 | Fluid shear stress and atherosclerosis | 119 | 1.836 | 1.44E-04 | 7.88E-04 | IFNG, SELE, CCL2, IL1B, IL1A, CTSL, IL1R2, VCAM1, MMP9, FOS, IL1R1, HSP90B1, ICAM1, TNFRSF1A, LOC100510917, SDC4, NQO1, NFKB1, HSP90AA1, TP53, ASS1, IKBKB, TXN, MGST1, GSTM3 |
| ssc04933 | AGE-RAGE signaling pathway in diabetic complications | 93 | 1.836 | 3.10E-04 | 1.60E-03 | SELE, CCL2, IL1B, CXCL8, IL6, SERPINE1, IL1A, STAT1, VCAM1, PIM1, STAT5A, STAT3, CASP3, EGR1, ICAM1, TGFB3 |
| ssc04917 | Prolactin signaling pathway | 59 | 1.824 | 8.90E-04 | 4.04E-03 | IRF1, STAT1, SOCS3, SOCS1, STAT5A, FOS, CISH, STAT3, GCK |
| ssc05416 | Viral myocarditis | 51 | 1.802 | 1.12E-03 | 4.96E-03 | SLA-DOA, SLA-DQA1, SLA-DRA, SLA-3, SLA-DOB, CD40, SLA-DQB1, SLA-7, SLA-DRB1, SLA-DMB, CYCS, CASP3, SLA-DMA, ICAM1, PRF1, CASP8 |
| ssc03060 | Protein export | 27 | 1.784 | 4.81E-03 | 1.75E-02 | HSPA5, SRPRA, SRPRB, SEC61A1, SEC61B, SPCS3, SEC11C, SPCS1, SPCS2, SRP14, SRP72, SRP54, SEC63, SGTA, SRP9, SRP68, SEC11A, SRP19, TMEM208 |
| ssc00520 | Amino sugar and nucleotide sugar metabolism | 35 | 1.763 | 4.06E-03 | 1.53E-02 | GFPT2, HK3, CHIA, GALE, GCK, UGP2, PGM2, GFPT1, HK2, GNPNAT1, PMM2, UGDH, UAP1 |
| ssc01250 | Biosynthesis of nucleotide sugars | 34 | 1.754 | 3.51E-03 | 1.39E-02 | GFPT2, HK3, GALE, GCK, UGP2, CMAH, PGM2, GFPT1, HK2, GNPNAT1, PMM2, UGDH, UAP1, NANS, GMDS, GMPPA, GFUS |
| ssc03040 | Spliceosome | 123 | 1.743 | 4.76E-04 | 2.35E-03 | HSPA6, SNRPF, HSPA8, SNRPD1, RBM8A, SNRPA1, SMNDC1, SNRPG, THOC6, LOC102162880, SNRNP27, SNRPD3, HNRNPA3, NCBP1, TRA2B, HNRNPC, SNRPA, U2AF1, ZMAT2, MAGOHB, SRSF3, SF3B6, PRPF40A, ALYREF, SNRPC, DDX5, SNRPB2, USP39, PHF5A, DHX15, SRSF1, TCERG1, DDX23, EIF4A3, SF3B3, SRSF10, SNRPE, SNRNP40, SRSF7, LSM6, PRPF19, MAGOH, SNU13, RBMX2, CWC25, LSM7, RBM25, DHX38, THOC3, LSM5, PPIE, U2AF1L4, THOC7, |

| | | | | | | |
|---|---|---|---|---|---|---|
| | | | | | | *PRPF3, PPIL1, SRSF2, LSM8, HNRNPK, BUD31, NCBP2, U2AF2, SRSF9, SF3A3, LSM2, U2SURP, RBMX, SYF2, SNIP1, CDC40, DDX39B, BUD13, EFTUD2, SF3B4, CWC15, SNRNP200, WBP11, TRA2A* |
| ssc04216 | Ferroptosis | 37 | 1.722 | 3.78E-03 | 1.48E-02 | *SLC39A14, SLC7A11, CP, SLC39A8, ACSL1, TFRC, FTH1, SLC3A2, ACSL5, SLC11A2, GCLM, TP53, LPCAT3, ACSL4, STEAP3, SAT1* |
| ssc00591 | Linoleic acid metabolism | 12 | 1.716 | 9.13E-03 | 2.98E-02 | *PLA2G2D, CYP3A22, PLA2G1B* |
| ssc05203 | Viral carcinogenesis | 149 | 1.716 | 3.58E-04 | 1.79E-03 | *IRF7, PMAIP1, EIF2AK2, CDKN1A, SLA-3, IRF9, EGR2, SLA-7, STAT5A, NFKB2, C3, NFKBIA, STAT3, CASP3, BAK1, TRAF2, TRAF3, LOC100517129, CASP8, JAK3, IL6ST, RANBP1, CREB3L1, NFKB1, SLA-8, TP53, KAT2A, LTBR, SND1, MAPKAPK2, DDX3X, YWHAG, H2BC5, DNAJA3, H2BC15, HDAC4, TBPL1, JAK1, CREB3L4, POLB, CCND2, PSMC1* |
| ssc00380 | Tryptophan metabolism | 32 | 1.712 | 7.61E-03 | 2.57E-02 | *IDO1, AOC1, CYP1B1, KYNU, ALDH9A1, IL4I1* |
| ssc04640 | Hematopoietic cell lineage | 68 | 1.709 | 4.13E-03 | 1.53E-02 | *IL1B, IL6, IL1A, CSF3, SLA-DOA, IL1R2, SLA-DQA1, SLA-DRA, SLA-DOB, IL4R, CSF3R, SLA-DQB1, SLA-DRB1, CSF1, SLA-DMB, CD38, TFRC, IL1R1, CSF2, SLA-DMA, IL2RA, CD14, CD8B, KIT, ITGA5* |
| ssc05012 | Parkinson disease | 220 | 1.707 | 6.06E-05 | 3.54E-04 | *UBE2L6, UBA7, SLC39A14, PPIF, SLC39A8, HSPA5, PSMA5, CYCS, CASP3, PSMA2, PSMA3, DDIT3, PSMB3, PSMA1, PSMD14, LOC100127131, SLC11A2, GPR37, DAXX, PSMB2, PSMB4, EIF2S1, TUBA4A, SLC39A7, PSMA4, PSMD12, TP53, PSMD3, SLC39A6, NDUFB2, TXN, ATP5MC1, SLC25A5, NDUFB3, COX7A2, ERN1, SLC39A11, SLC39A1, PSMD8, PSMC4, COX5A, UQCRC1, NDUFAB1, COX6B, SLC18A1, XBP1, PSMB1, PSMD7, PSMD11, COX5B, NDUFV2, PSMC6, PSMD1, PSMC1, ATF6, PSMC2, UQCRQ, NDUFA6, ATP5F1C, SLC18A2, LOC100522678, NDUFA9, PSMB6, UBE2J2, DUSP1, PSMD13, ATP5F1B, PSMD6, COX7B, UBE2L3, SOD1, PSMD2, PSMD4, SDHD, ATP5F1D, KIF5B, ATP5PB, ATP5F1A, EIF2AK3, TUBB1, UQCR11, VDAC1, NDUFB1, NDUFB4, CYC1, NFE2L2, LRRK2, PSMB7, ATP5MC3,* |

| | | | | | | |
|---|---|---|---|---|---|---|
| | | | | | | *PSMC5, NDUFB9, MFN1, TUBB6, PSMC3, NDUFC2, VDAC2, ATP5PD, ITPR1, NDUFA4, UQCR10, MAOA* |
| ssc01524 | Platinum drug resistance | 58 | 1.700 | 2.46E-03 | 1.02E-02 | *BIRC3, FAS, PMAIP1, CDKN1A, FASLG, CYCS, CASP3, BAK1, CASP8, LOC100510917, SLC31A1, TP53, MGST1, GSTM3, GSTO1, GSTO2, MGST3* |
| ssc04215 | Apoptosis - multiple species | 25 | 1.695 | 8.47E-03 | 2.81E-02 | *BIRC3, PMAIP1, CASP7, CYCS, CASP3, BAK1, CASP8, TNFRSF1A* |
| ssc05132 | Salmonella infection | 215 | 1.677 | 1.19E-04 | 6.71E-04 | *IL1B, CXCL8, IL6, BIRC3, LOC100522887, TNFSF10, CASP7, FOS, NFKBIA, HSP90B1, CYCS, CASP3, MLKL, BAK1, LOC100624226, MYL7, GSDMD, TRAF2, RHOB, RHOJ, CASP1, CD14, RIPK2, CASP8, MYD88, LOC100127131, TLR4, TNFRSF1A, RIPK3, MYC, TLR2, TIRAP, NFKB1, IRAK4, ARPC5L, TUBA4A, HSP90AA1, LY96, IKBKB, TXN, KPNA4, RIPK1, IRAK1, ACTR3, RHOH, MAP2K3, ACBD3, TAB3, EXOC5, PFN1, ARF6* |
| ssc03010 | Ribosome | 144 | 1.660 | 1.05E-03 | 4.70E-03 | *RPL13A, MRPL38, MRPS27, MRPL17, MRPL34, RSL24D1, MRPS35, MRPL55, MRPL35, MRPL15, MRPL42, MRPS23, MRPS7, MRPL20, MRPL50, DAP3, MRPL48, MRPL2, MRPS11, RPS27L, MRPS15, MRPS30, MRPL19, MRPL51, MRPS6, MRPL47, MRPL28, MRPL40, MRPS18B, MRPL46, MRPS18C, MRPS10, MRPL32, MRPL44, RPL27, MRPL18, MRPL24, MRPL22, MRPS22, RPL26L1, MRPL1, MRPL13, RPL22L1, MRPL54, RPS2, MRPL11, MRPS24, RPS18, MRPS5, MRPS33, MRPL33, MRPS34, MRPL37, RPS23, MRPS26, MRPS18A, MRPS31, RPL23, MRPS17, RPL38, RPSA, RPLP0, RPL37A, MRPL36, RPL10A, MRPL27, MRPL45, MRPL4* |
| ssc04514 | Cell adhesion molecule (CAM) interaction | 114 | 1.650 | 1.44E-03 | 6.27E-03 | *MADCAM1, SELE, CD274, SLA-DOA, SLA-DQA1, SLA-DRA, VCAM1, SLA-3, CTLA4, PDCD1LG2, SLA-DOB, LRRC4, CD40, PVR, SLA-DQB1, SELP, SLA-7, SLA-DRB1, SLA-DMB, SLA-DMA, VCAN, ICAM1* |
| ssc04920 | Adipocytokine signaling pathway | 59 | 1.641 | 5.86E-03 | 2.08E-02 | *SOCS3, PCK2, CPT1C, NFKBIE, NFKBIB, ACSL1, NFKBIA, STAT3, TRAF2, ACSL5, TNFRSF1A, SLC2A4, NFKB1, TNFRSF1B, JAK2, IKBKB, ACSL4, PPARGC1A* |
| ssc04115 | p53 signaling pathway | 65 | 1.639 | 6.21E-03 | 2.19E-02 | *SERPINE1, FAS, PMAIP1, THBS1, CDKN1A, GADD45A, SHISA5, SESN2, GADD45G, CYCS, CASP3, LOC100624226, CASP8* |

| | | | | | | |
|---|---|---|---|---|---|---|
| ssc04066 | HIF-1 signaling pathway | 96 | 1.629 | 6.69E-03 | 2.28E-02 | *IFNG, IL6, SERPINE1, CDKN1A, HK3, TIMP1, TFRC, STAT3, EIF4EBP1, TLR4, PGK1, PFKP, HIF1A, NFKB1, IFNGR1, ENO1, EIF4E, HK2, IFNGR2, ANGPT2, LTBR, MKNK2, IL6R, EGLN3* |
| ssc00510 | N-Glycan biosynthesis | 52 | 1.627 | 9.47E-03 | 3.05E-02 | *B4GALT1, MOGS, OSTC, MAN1B1, ALG3, RPN1, RPN2, STT3A, DAD1, MGAT2, DHRSX, TMEM258, MGAT1, STT3B, DPAGT1, B4GALT3, ST6GAL2, DPM1, GANAB, FUT8, MAGT1, MGAT4A, DOLPP1, ALG5, ALG8, DDOST* |
| ssc04650 | Natural killer cell mediated cytotoxicity | 79 | 1.619 | 7.67E-03 | 2.57E-02 | *IFNG, FAS, TNFSF10, IFNAR2, LOC100523789, FASLG, CSF2, CASP3, SH3BP2, LOC100624226, ULBP1, ICAM1, PRF1, KLRK1, CD48, IFNGR1, IFNGR2* |
| ssc04068 | FoxO signaling pathway | 116 | 1.604 | 3.81E-03 | 1.48E-02 | *IL6, TNFSF10, SOD2, CDKN1A, PCK2, GADD45A, IL10, S1PR1, FASLG, GADD45G, STAT3, PLK3, BCL6, TGFB3, SLC2A4, GADD45B, PRMT1, BNIP3, IKBKB, SMAD3, DEPP1* |
| ssc05170 | Human immunodeficiency virus 1 infection | 175 | 1.596 | 1.63E-03 | 7.01E-03 | *TAP1, SAMHD1, FAS, TAP2, BST2, SLA-3, SLA-7, TAPBP, B2M, FOS, NFKBIA, FASLG, TAPBPL, CYCS, CASP3, BAK1, TRAF2, PDIA3, GNG10, CASP8, MYD88, TLR4, CGAS, TNFRSF1A, CALR, TLR2, STING1, NFKB1, IRAK4, TNFRSF1B, SLA-8, IKBKB* |
| ssc05202 | Transcriptional misregulation in cancer | 145 | 1.560 | 3.97E-03 | 1.52E-02 | *ETV7, CXCL8, IL6, BIRC3, NUPR1, IL1R2, CDKN1A, NFKBIZ, MMP9, CD40, NR4A3, GADD45A, IL2RB, PML, GADD45G, CSF2, BAK1, DDIT3, MAX, BCL6, CD14, CEBPB, BCL2A1, ETV6, MYC, GADD45B, NFKB1* |
| ssc03013 | Nucleocytoplasmic transport | 95 | 1.557 | 9.68E-03 | 3.09E-02 | *KPNA2, IPO4, RBM8A, TMEM33, KPNA4, SEC13, IPO7, THOC6, XPOT, NCBP1, RAN, XPO6, KPNB1, MAGOHB, POM121C, NXT1, UPF2, XPO5, PHAX, ALYREF, NMD3, NUP54, IPO11, EIF4A3, NUP50, NUP188, TNPO1, RNPS1, RAE1, MAGOH, KPNA6, NUP98, SAP18, XPO1, AHCTF1, TNPO3, IPO13, THOC3, SUMO1, SUMO2, KPNA1, THOC7, NUP42, NUP153, UPF3B, RANGAP1, CSE1L, SRRM1, TNPO2, XPO7, NUP35, NCBP2, PYM1, LOC100511376, IPO5, TPR* |
| ssc01240 | Biosynthesis of cofactors | 122 | 1.510 | 1.21E-02 | 3.81E-02 | *IDO1, CMPK2, PSAT1, SHMT2, AK2, CTPS1, NME1, MTHFD2, MAT1A, MTHFD1L, KYNU, UGP2, SHMT1, AK5, GCLM, NQO1, PDXK, EPRS1, BCAT1, UGT2B31, RFK, UMPS, PNPO, PMM2, LIPT2, UGDH, UGT1A6, AK3, NME2, DHODH, SDR16C5, AK6, KMO, GMPPA, COX10, CAD, EARS2, COQ2, NADK, CTPS2, GCH1, GPHN, BCO1, RDH13, FPGS, MAT2B, ADSS2, PPCDC, MPI, NMNAT1, CMPK1* |

| | | | | | | |
|---|---|---|---|---|---|---|
| ssc05135 | Yersinia infection | 124 | 1.509 | 1.20E-02 | 3.81E-02 | *CCL2, IL1B, CXCL8, IL6, IL10, FOS, NFKBIA, TRAF2, CASP1, TICAM1, MYD88, TLR4, FYB1, CD8B, ITGA5, NFKB1, IRAK4, ARPC5L, IKBKB* |
| ssc05014 | Amyotrophic lateral sclerosis | 296 | 1.475 | 1.86E-03 | 7.94E-03 | *HSPA5, PSMA5, GPX3, CYCS, CASP3, PSMA2, PSMA3, TRAF2, DDIT3, CASP1, PSMB3, OPTN, PSMA1, TOMM40, PSMD14, LOC100127131, SETX, TNFRSF1A, DAXX, PSMB2, TANK, PSMB4, RAB39B, EIF2S1, TUBA4A, TNFRSF1B, DNAH17, PSMA4, C9orf72, PSMD12, TP53, PSMD3, NDUFB2, RAB1A, ATP5MC1, SEC13, UBQLN1, NDUFB3, COX7A2, HNRNPA2B1, ERN1, MAP2K3, PSMD8, PSMC4, COX5A, UQCRC1, HNRNPA3, NDUFAB1, NCBP1, COX6B, VCP, XBP1, PSMB1, PFN1, PSMD7, PSMD11, COX5B, NDUFV2, PSMC6, POM121C, NXT1, PSMD1, SRSF3, PSMC1, ATF6, PSMC2, UQCRQ, PPP3CA, ALYREF, NDUFA6, ATP5F1C, LOC100522678, NDUFA9, PSMB6, NUP54, PSMD13, ATP5F1B, SIGMAR1, PSMD6, COX7B, ATG101, GPX7, SOD1, PSMD2, PSMD4, GPX8, SQSTM1, BID, SDHD, ATP5F1D, NRBF2, TARDBP, NUP50, KIF5B, SRSF7, ATP5PB, ATP5F1A, NUP188, PFN2, CHCHD10, EIF2AK3, TUBB1, UQCR11, VDAC1, CCS, NDUFB1, NDUFB4, RAE1, CYC1, RB1CC1, ATXN2L, PSMB7, ATP5MC3, DCTN6, DCTN5, ACTR1A, PSMC5, RAB8A, NUP98, NDUFB9, TBK1, TUBB6, PSMC3, NDUFC2, MAPK11, ATP5PD, RAB5A, NDUFA4, UQCR10* |
| ssc05010 | Alzheimer disease | 319 | 1.438 | 3.14E-03 | 1.26E-02 | *IL1B, IL6, IL1A, FAS, LPL, SLC39A14, EIF2AK2, PPIF, CASP7, SLC39A8, CSF1, PSMA5, CYCS, CASP3, PSMA2, PSMA3, PTGS2, FZD9, TRAF2, DDIT3, DKK1, PSMB3, CASP8, PSMA1, PSMD14, LOC100127131, ADAM17, TNFRSF1A, SLC11A2, PSMB2, PSMB4, EIF2S1, NFKB1, TUBA4A, SLC39A7, PSMA4, PSMD12, PPID, PSMD3, IKBKB, SLC39A6, NDUFB2, ATP5MC1, SLC25A5, WNT16, NDUFB3, COX7A2, ERN1, SLC39A11, SLC39A1, PSMD8, PSMC4, COX5A, UQCRC1, NDUFAB1, COX6B, XBP1, PSMB1, CSNK2A1, IDE, HSD17B10, PSMD7, PSMD11, COX5B, NDUFV2, PSMC6, PSMD1, PSMC1, BACE2, ATF6, PSMC2, UQCRQ, PPP3CA, NDUFA6, ATP5F1C, LOC100522678, NDUFA9, PSMB6, PSMD13, WNT7B, ATP5F1B, CHUK, PSMD6, COX7B, ATG101, PSMD2, PSMD4, BID, SDHD, ATP5F1D, NRBF2, CSNK1A1, KIF5B, RELA, ATP5PB, ATP5F1A, GAPDH, EIF2AK3, TUBB1, UQCR11, VDAC1, NDUFB1, NDUFB4* |

| | | | | | | |
|---|---|---|---|---|---|---|
| ssc04015 | Rap1 signaling pathway | 179 | -1.387 | 1.63E-02 | 4.98E-02 | *CTNND1, PLCB4, EFNA5, LPAR2, ADORA2A, PDGFD, TIAM1, ARAP3, FGF9, VEGFA, CALM3, PARD6A, ADCY7, EVL, EGF, FGF18, MAGI3, RASSF5, RAPGEF4, KDR, IGF1, FGF21, ANGPT4, PRKCZ, PRKCG, ADCY1, RAC3, EFNA2, MAP2K6, DOCK4, MAGI2, TEK, ITGAM, FLT4, RASGRP2, KITLG, FARP2, ADCY8, CSF1R, TLN2, RGS14, ANGPT1, CNR1, PRKCB, NGFR, FGF1, PLCB2, CALML4, VEGFD* |
| ssc04518 | Integrin signaling | 140 | -1.414 | 1.48E-02 | 4.57E-02 | *ITGB5, LAMC3, FRAS1, ITGA11, ACTN4, PDGFA, LAMA5, COL9A2, ICAM3, ERBB2, LAMC1, PDGFD, VEGFA, LIMS1, COL4A5, PXN, FBN1, COL2A1, LIMS2, LAMC2, EGF, FREM2, FLNA, FERMT1, KDR, VCL, IGF1, COL1A1, PARVG, RAC3, RELN, FN1, ITGA3, ITGA8, CHAD, COL4A6, LAMA3, ITGB4, COL1A2, LAMB2, COL4A4, ITGAM, FLT4, ITGA4, THBS3, NPNT, COL4A3, FREM1, TLN2, LAMA2, ITGB7, TNXB, COL6A6, LAMA1, VTN, COMP, COL9A3, VEGFD* |
| ssc04024 | cAMP signaling pathway | 166 | -1.450 | 6.48E-03 | 2.24E-02 | *GLI1, ADCY7, MYL9, HTR1D, LHB, CREB5, NPY1R, CACNA1D, TSHR, RAPGEF4, HTR4, SOX9, ATP2B2, ADCY1, RAC3, CAMK2G, PDE3B, FXYD2, CREB3L3, ATP2B4, NFATC1, CNGA2, NPR1, PLD1, GRIA1, ATP2A3, CACNA1S, ADCY8, GRIA4, BDNF, ATP1B2, CHRM2, ADRB1, GRIN3B, TNNI3, HHIP, CNGA1, CALML4, VIPR2, EDN3* |
| ssc04310 | Wnt signaling pathway | 151 | -1.474 | 4.32E-03 | 1.58E-02 | *AXIN2, APC2, SFRP5, WNT11, WNT2, PRKCG, RAC3, NFATC2, CAMK2G, WNT10B, TLE4, WNT7A, WNT9A, CTNND2, NFATC1, WNT3A, PRICKLE1, TCF7, WNT5B, NOTUM, CTNNBIP1, APCDD1, FZD8, PRKCB, WNT4, WIF1, PLCB2, SFRP4, DKK2, SFRP2* |
| ssc04713 | Circadian entrainment | 74 | -1.564 | 6.37E-03 | 2.22E-02 | *ADCY7, RPS6KA5, RYR3, PRKG2, CACNA1G, GUCY1A2, CACNA1D, GNG2, GNG8, PRKCG, ADCY1, CAMK2G, GNG4, GRIA1, PER3, ADCY8, GRIA4, PER2, PRKCB, PLCB2, CALML4* |
| ssc04360 | Axon guidance | 167 | -1.565 | 8.76E-04 | 4.03E-03 | *EFNB2, SEMA4F, PLXNC1, RHOD, ABLIM1, LRIG2, EPHA5, CDK5, SMO, EFNA5, ABLIM3, PTCH1, BMPR2, NTNG2, EPHA4, SLIT2, PLXNB1, PARD6A, BMPR1B, ROBO1, ROBO2, MYL9, NGEF, BMP7, SEMA3C, SEMA5B, CXCR4, EPHB3, SEMA6D, SEMA3E, SEMA5A, SRGAP1, BOC, EPHB1, PRKCZ, SEMA3B, RAC3, NEO1, NFATC2, CAMK2G, PLXNB3, EFNA2, SEMA6A, SEMA3D, UNC5C, EPHA7, SSH1, SEMA4G, PLXNA3, WNT5B, SHH, SEMA6C, PAK6, NTNG1, SLIT3, EFNB3, NTN3, EPHA3, SRGAP3, WNT4, LOC100620540* |

| | | | | | | |
|---|---|---|---|---|---|---|
| ssc04082 | Neuroactive ligand signaling | 116 | -1.578 | 2.58E-03 | 1.05E-02 | *SLC6A13, P2RX5, GABRA1, TACR3, GNG2, HTR4, GNG8, GRM8, ADRA1D, PRKCG, ADCY1, GABRG1, SLC1A1, GNG4, GABRR3, GRIA1, P2RX6, SLC6A2, GRIK2, ADCY8, GRIK4, GRIA4, PRKCB, PLCB2, CHRM2, ADRB1, OPRL1, GRIN3B, P2RY12, ADRA1A, P2RX3, ADRA2C, SLC6A4* |
| ssc04510 | Focal adhesion | 179 | -1.615 | 6.89E-04 | 3.26E-03 | *MYL9, LAMC2, EGF, SHC3, FLNA, KDR, VCL, IGF1, PRKCG, COL1A1, PARVG, RAC3, RELN, FN1, ITGA3, ITGA8, CHAD, CAV2, COL4A6, LAMA3, ITGB4, COL1A2, PPP1R12B, LAMB2, COL4A4, MYLK4, FLT4, ITGA4, THBS3, COL4A3, PAK6, TLN2, RASGRF1, CAV1, LAMA2, ITGB7, TNXB, COL6A6, PRKCB, LAMA1, VTN, COMP, COL9A3, VEGFD* |
| ssc04925 | Aldosterone synthesis and secretion | 79 | -1.633 | 4.10E-03 | 1.53E-02 | *CALM3, STAR, CYP11A1, ADCY7, CACNA1G, CREB5, CACNA1D, ATP2B2, PRKCG, ADCY1, CAMK2G, CREB3L3, ATP2B4, CAMK1G, NPR1, CACNA1S, ADCY8, PDE2A, PRKCB, ATP1B2, PLCB2, CALML4* |
| ssc04961 | Endocrine and other factor-regulated calcium reabsorption | 42 | -1.658 | 1.40E-02 | 4.37E-02 | *PTH1R, ATP2B2, PRKCG, FXYD2, DNM1, ATP2B4, SLC8A2, ESR1, PRKCB, ATP1B2, PLCB2* |
| ssc04916 | Melanogenesis | 89 | -1.661 | 2.48E-03 | 1.02E-02 | *WNT3, EDNRB, WNT2B, FZD7, PLCB4, TCF7L1, DVL2, CALM3, ADCY7, MITF, FZD2, WNT11, WNT2, PRKCG, ADCY1, CAMK2G, WNT10B, WNT7A, WNT9A, CREB3L3, WNT3A, DCT, KITLG, TCF7, WNT5B, ADCY8, FZD8, PRKCB, WNT4, PLCB2, CALML4* |
| ssc04022 | cGMP-PKG signaling pathway | 141 | -1.669 | 3.46E-04 | 1.75E-03 | *ADCY7, MYL9, PRKG2, GUCY1A2, CREB5, CACNA1D, PDE5A, MEF2B, ADRA1D, ADRB3, ATP2B2, ADCY1, NFATC2, PDE3B, FXYD2, CREB3L3, ATP2B4, PIK3R6, NFATC1, MYLK4, NPR1, ATP2A3, LOC100736765, KCNMB4, CACNA1S, ADCY8, SLC8A2, PIK3CG, PDE2A, ATP1B2, PLCB2, ADRB1, CNGA1, CALML4, ADRA1A, ADRA2C* |
| ssc04970 | Salivary secretion | 60 | -1.671 | 5.21E-03 | 1.87E-02 | *CALM3, VAMP2, ADCY7, RYR3, PRKG2, GUCY1A2, ADRA1D, ADRB3, ATP2B2, PRKCG, ADCY1, FXYD2, ATP2B4, CST3, ADCY8, PRKCB, ATP1B2, PLCB2, BST1, ADRB1, CALML4, ADRA1A* |

| | | | | | | |
|---|---|---|---|---|---|---|
| ssc04740 | Olfactory transduction | 38 | -1.672 | 9.16E-03 | 2.98E-02 | *CAMK2G, CNGA2, SLC8A2, LOC100736710, NCALD, LOC100738112, PDE2A, LOC110260676, CALML4, SLC24A4* |
| ssc04814 | Motor proteins | 141 | -1.717 | 1.36E-04 | 7.59E-04 | *KIF20B, MYO1C, TUBE1, MYH11, DYNLRB2, MYO1F, DYNC2H1, KIF7, MYO9A, DNAI3, KIF1C, TNNC2, DNAH12, TPM1, BICDL1, DNAH2, DNAH3, KIF15, DNAH5, KIF2C, DNAH6, KIF13A, MYL9, KLC4, DNAH11, DNAH7, MYO5C, KIF11, MYH14, DYNLT5, MYH7B, CENPE, DYNLT4, DNAI1, KIFC1, KIF22, KIF3C, KIF18B, KIF20A, MYO5B, MYO7A, DNAH9, TNNI1, TUBB4A, DNAH10, KIF24, DNAI2, ACTC1, DNAH1, KIF14, KLC3, KIFC2, LOC100736765, MYO15B, MYH2, TNNI3* |
| ssc04081 | Hormone signaling | 128 | -1.719 | 1.55E-04 | 8.32E-04 | *NPR2, THRA, BMPR1B, THPO, ADCY7, SLC16A10, PRKG2, LHB, HEPH, TSHR, IGF2, RAPGEF4, GNG2, PTH1R, GNG8, IGF1, TRHR, ADRA1D, ADRB3, EPOR, ADCY1, GNG4, NPR1, GCGR, SLC40A1, SLC16A2, LEPR, VIPR1, ADCY8, RAMP3, ESR1, AGTR2, PLCB2, ADRB1, OPRL1, ADRA1A, VIPR2, ADRA2C* |
| ssc04261 | Adrenergic signaling in cardiomyocytes | 122 | -1.789 | 4.41E-05 | 2.62E-04 | *TPM1, CACNB2, CACNB3, ADCY7, RPS6KA5, KCNQ1, KCNE1, CREB5, CACNA1D, CACNG5, RAPGEF4, CACNA2D2, ADRA1D, ATP2B2, ADCY1, CAMK2G, FXYD2, CREB3L3, ATP2B4, PIK3R6, ACTC1, ATP2A3, LOC100736765, CACNA1S, ADCY8, CACNB4, SLC8A2, CACNG6, PIK3CG, AGTR2, ATP1B2, PLCB2, ADRB1, TNNI3, CALML4, ADRA1A* |
| ssc05410 | Hypertrophic cardiomyopathy | 78 | -1.789 | 8.56E-04 | 3.99E-03 | *DMD, TPM1, SGCE, CACNB2, CACNB3, CACNA1D, CACNG5, CACNA2D2, IGF1, ITGA3, ITGA8, ITGB4, ACTC1, ITGA4, ATP2A3, LOC100736765, CACNA1S, CACNB4, SLC8A2, CACNG6, TGFB2, LAMA2, ITGB7, ACE, SGCA, LAMA1, TNNI3* |
| ssc04820 | Cytoskeleton in muscle cells | 178 | -1.809 | 1.42E-05 | 9.63E-05 | *ITGB5, PDLIM4, ITGA11, SPTAN1, SPTBN2, ENO3, MYH11, FBN2, SUN2, FMNL2, COL9A2, FBLN1, FLNC, ELN, SUN1, TNNC2, MYOZ1, DIAPH1, ANKRD23, DSP, PLEC, SDC1, COL4A5, JUP, SDC2, FBN1, DMD, TPM1, SGCE, MYL9, BGN, DSC2, SYNPO2, AGRN, SPTBN1, MYH14, MYOM3, COL11A1, MYH7B, FHL2, PDLIM2, VCL, COL1A1, FN1, DSG2, ITGA3, ITGA8, COL4A6, CSRP1, ITGB4, COL1A2, VIM, SYNE3, TNNI1, COL4A4, COL3A1, ACTC1, ITGA4, THBS3, DCN, SDC3, COL4A3, LOC100736765, PDLIM3, FBN3, TMOD2, TLN2, SPTBN4, FBLN2,* |

| | | | | | | |
|---|---|---|---|---|---|---|
| | | | | | | *FHL1, LAMA2, ITGB7, COL6A6, ATP1B2, SGCA, OBSCN, MYH2, LAMA1, TNNI3, COMP, COL9A3, TRIM55* |
| ssc04512 | ECM-receptor interaction | 78 | -1.822 | 4.97E-04 | 2.41E-03 | *COL1A1, RELN, FN1, ITGA3, ITGA8, CHAD, COL4A6, LAMA3, ITGB4, COL1A2, LAMB2, COL4A4, ITGA4, THBS3, NPNT, COL4A3, FREM1, GP9, LAMA2, ITGB7, TNXB, COL6A6, LAMA1, VTN, COMP, COL9A3, LOC100511343* |
| ssc04080 | Neuroactive ligand-receptor interaction | 149 | -1.849 | 4.80E-06 | 3.52E-05 | *LHB, NPY1R, P2RX5, TSHR, GABRA1, CYSLTR1, LTB4R2, TACR3, HTR4, PTH1R, GRM8, TRHR, ADRA1D, F2RL1, ADRB3, CNR2, PAQR6, GABRG1, NMUR1, GRID1, P2RY10, GABRR3, NPFF, GCGR, GRIA1, P2RX6, LEPR, VIPR1, GRIK2, GALR1, GRIK4, P2RY8, APLN, GRIA4, AGTR2, CNR1, GRID2, P2RX7, LYPD6, CHRM2, ADRB1, OPRL1, GRIN3B, GPR156, ADRA1A, VIPR2, P2RX3, F2, QRFPR, APLNR, EDN3, ADRA2C* |
| ssc05412 | Arrhythmogenic right ventricular cardiomyopathy | 68 | -1.921 | 3.70E-05 | 2.31E-04 | *DSP, TCF7L1, JUP, DMD, SGCE, CACNB2, CACNB3, DSC2, CACNA1D, CACNG5, CACNA2D2, DSG2, ITGA3, ITGA8, ITGB4, ITGA4, ATP2A3, TCF7, CACNA1S, CTNNA3, CACNB4, SLC8A2, CACNG6, LAMA2, ITGB7, SGCA, LAMA1* |
| ssc04020 | Calcium signaling pathway | 186 | -1.991 | 4.46E-08 | 5.13E-07 | *PDGFA, EDNRB, TBXA2R, ADRA1B, PLCB4, GRIN3A, ERBB2, ADORA2A, PDGFD, PTGFR, FGF9, PTGER1, TNNC2, VEGFA, CALM3, TPCN2, TPCN1, ADCY7, MST1, RYR3, EGF, CXCR4, CACNA1G, FGF18, CACNA1D, PLCD3, P2RX5, PHKA2, CYSLTR1, LTB4R2, TACR3, KDR, HTR4, ERBB4, HRC, TRHR, FGF21, ADRA1D, ADRB3, ATP2B2, PRKCG, ADCY1, ITPKB, NFATC2, PHKG1, CAMK2G, ATP2B4, NFATC1, MYLK4, FLT4, PLCD1, CAMK1G, MCOLN2, GDNF, NTRK3, P2RX6, ATP2A3, NTRK2, CACNA1S, ADCY8, SLC8A2, CASQ1, P2RX7, PRKCB, FGF1, PLCB2, CHRM2, CACNA1E, ADRB1, GRIN3B, CALML4, ADRA1A, P2RX3, VEGFD, CACNA1B* |
| ssc04974 | Protein digestion and absorption | 70 | -2.005 | 2.66E-05 | 1.72E-04 | *SLC16A10, COL28A1, KCNQ1, DPP4, MME, COL11A1, COL26A1, COL23A1, COL1A1, COL4A6, SLC1A1, COL14A1, FXYD2, COL1A2, COL4A4, COL3A1, CTRB2, COL4A3, SLC8A2, COL25A1, SLC7A9, COL6A6, ATP1B2, COL19A1, SLC7A8, COL21A1, COL15A1, COL8A2, COL9A3, KCNJ13* |

| | | | | | | |
|---|---|---|---|---|---|---|
| ssc05414 | Dilated cardiomyopathy | 81 | -2.094 | 7.88E-07 | 7.11E-06 | *DMD, TPM1, SGCE, CACNB2, CACNB3, ADCY7, CACNA1D, CACNG5, CACNA2D2, IGF1, ADCY1, ITGA3, ITGA8, ITGB4, ACTC1, ITGA4, ATP2A3, LOC100736765, CACNA1S, ADCY8, CACNB4, SLC8A2, CACNG6, TGFB2, LAMA2, ITGB7, SGCA, ADRB1, LAMA1, TNNI3* |

Abbreviations: AGE-RAGE: Advanced glycation end products-receptor for AGEs; cAMP: Cyclic adenosine monophosphate; cGMP-PKG: Cyclic guanosine monophosphate-protein kinase G; ECM: Extracellular matrix; FDR: False discovery rate; FoxO: Forkhead box O; HIF-1: Hypoxia-inducible factor 1; IL-17: Interleukin-17; JAK-STAT: Janus kinases-signal transducer and activator of transcription proteins; KEGG: Kyoto Encyclopedia of Genes and Genomes; NES: Normalized enrichment score; NF-kappa B: Nuclear factor-kappa B; NOD: Nucleotide-binding domain; PD-1: Programmed cell death protein 1; PD-L1: Programmed cell death ligand 1; Rap1: Ras-associated protein-1; RIG-I: Retinoic acid-inducible gene I; Th1/Th2/Th17: T helper 1/2/17 cells; TNF: Tumor necrosis factor.

**Supplementary Table S4. Significantly enriched pathways from the GSEA result of the GO:BP database, High-glucose infection versus High-glucose control.** Abbreviations: GO:BP: Biological Processes in Gene Ontology; GSEA: Gene set enrichment analysis.

| GO:BP ID | Description | Set Size | NES | P-value | FDR | Core enrichment gene |
|---|---|---|---|---|---|---|
| GO:0034097 | Response to cytokine | 214 | 2.779 | 1.00E-10 | 1.87E-09 | *CXCL9, CXCL11, CXCL10, UBD, IFNG, SELE, ACKR1, CCL2, IRF1, AMCF-II, CXCL13, IL1B, CHI3L1, MMP8, MX1, ISG15, CXCL8, CCL4, IL6, IL1A, TNFSF13B, IRF7, STAT1, ZBP1, OAS1, SOCS1, GFPT2, CCL8, FOSL1, TIFA, SNX10, IL15, IRF8, PIM1, IL4R, TIMP1, CD40, SLA-DQB1, MSC, TICAM2, PPBP, CCL5, NFIL3, ZFP36, STAT2, STAT5A, IL10, NFKBIA, CISH, STAT3, CXCL16, EGR1, TRAF2, TRAF3, IL18RAP, CASP8, YBX3, CACYBP, JAK3, MCL1, TLR4, ADAM17, TNFRSF1A, KIT, STIP1, TLR2, CDC42EP2, IFNGR1* |
| GO:1901652 | Response to peptide | 214 | 2.779 | 1.00E-10 | 1.87E-09 | *CXCL9, CXCL11, CXCL10, UBD, IFNG, SELE, ACKR1, CCL2, IRF1, AMCF-II, CXCL13, IL1B, CHI3L1, MMP8, MX1, ISG15, CXCL8, CCL4, IL6, IL1A, TNFSF13B, IRF7, STAT1, ZBP1, OAS1, SOCS1, GFPT2, CCL8, FOSL1, TIFA, SNX10, IL15, IRF8, PIM1, IL4R, TIMP1, CD40, SLA-DQB1, MSC, TICAM2, PPBP, CCL5, NFIL3, ZFP36, STAT2, STAT5A, IL10, NFKBIA, CISH, STAT3, CXCL16, EGR1, TRAF2, TRAF3, IL18RAP, CASP8, YBX3, CACYBP, JAK3, MCL1, TLR4, ADAM17, TNFRSF1A, KIT, STIP1, TLR2, CDC42EP2, IFNGR1* |
| GO:0019221 | Cytokine-mediated signaling pathway | 129 | 2.777 | 1.00E-10 | 1.87E-09 | *CXCL9, CXCL11, CXCL10, IFNG, ACKR1, CCL2, IRF1, AMCF-II, CXCL13, IL1B, MMP8, ISG15, CXCL8, CCL4, IL6, IL1A, TNFSF13B, IRF7, STAT1, ZBP1, OAS1, SOCS1, CCL8, FOSL1, TIFA, IL15, IL4R, CD40, TICAM2, PPBP, CCL5, STAT2, STAT5A, IL10, NFKBIA, CISH, STAT3, EGR1, TRAF2, TRAF3, IL18RAP, JAK3, ADAM17, TNFRSF1A, KIT* |
| GO:0043207 | Response to external biotic stimulus | 365 | 2.767 | 1.00E-10 | 1.87E-09 | *CXCL9, CXCL10, GBP1, ISG20, IFNG, BATF2, GBP2, CCL2, IRF1, AMCF-II, S100A12, CXCL13, IL1B, MMP8, MX1, ISG15, CXCL8, S100A8, CCL4, RSAD2, IL27, MX2, IFIT1, IL1A, IRF7, DHX58, SAMHD1, STAT1, ZBP1, OAS1, RIGI, OAS2, BST2, LBP, CALHM6, CCL8, PTGIR, GSDMA, FOSL1, RNF19B, IFIH1, BPIFB1, TNFAIP3, TIFA, NMI, SLC15A3, IRF8,* |

| | | | | | |
|---|---|---|---|---|---|
| | | | | | *CFB, IFIT2, PIM1, FFAR2, CD40, SLA-DQB1, NOD2, IFI35, TICAM2, PPBP, CCL5, ZFP36, HP, STAT2, NFKBIB, C2, IL10, TRIM14, C3, NFKBIA, IFIT5, LTF, C4A, CD180, DUSP10, MLKL, CXCL16, CEBPG, TREM1, NDUFAF4, BPIFA1, TRAF2, UNC93B1, SERPING1, TRAF3, DDX21, TRIM26, CD14, MSRB1, POLR3D, IL18RAP, OPTN, CEBPB, TICAM1, MYD88, SELENOS, GNLY, CLEC7A, PLSCR1, TLR4, TNIP2, CGAS, KLRK1, MORC3, ARG2, DDIT4, NKG7, LYAR, RIPK3, RNF115, TMEM126A, TLR2, TIRAP, BMP2, CDC42EP2, STING1, IFNGR1, IRAK4, MR1, LOC100516039, HSP90AA1, LY96, SFTPD, TLR3, JAK2, CD46, KCNK6* |
| GO:0051707 | Response to other organism | 365 | 2.767 | 1.00E-10 | 1.87E-09 | *CXCL9, CXCL10, GBP1, ISG20, IFNG, BATF2, GBP2, CCL2, IRF1, AMCF-II, S100A12, CXCL13, IL1B, MMP8, MX1, ISG15, CXCL8, S100A8, CCL4, RSAD2, IL27, MX2, IFIT1, IL1A, IRF7, DHX58, SAMHD1, STAT1, ZBP1, OAS1, RIGI, OAS2, BST2, LBP, CALHM6, CCL8, PTGIR, GSDMA, FOSL1, RNF19B, IFIH1, BPIFB1, TNFAIP3, TIFA, NMI, SLC15A3, IRF8, CFB, IFIT2, PIM1, FFAR2, CD40, SLA-DQB1, NOD2, IFI35, TICAM2, PPBP, CCL5, ZFP36, HP, STAT2, NFKBIB, C2, IL10, TRIM14, C3, NFKBIA, IFIT5, LTF, C4A, CD180, DUSP10, MLKL, CXCL16, CEBPG, TREM1, NDUFAF4, BPIFA1, TRAF2, UNC93B1, SERPING1, TRAF3, DDX21, TRIM26, CD14, MSRB1, POLR3D, IL18RAP, OPTN, CEBPB, TICAM1, MYD88, SELENOS, GNLY, CLEC7A, PLSCR1, TLR4, TNIP2, CGAS, KLRK1, MORC3, ARG2, DDIT4, NKG7, LYAR, RIPK3, RNF115, TMEM126A, TLR2, TIRAP, BMP2, CDC42EP2, STING1, IFNGR1, IRAK4, MR1, LOC100516039, HSP90AA1, LY96, SFTPD, TLR3, JAK2, CD46, KCNK6* |
| GO:0009607 | Response to biotic stimulus | 378 | 2.741 | 1.00E-10 | 1.87E-09 | *CXCL9, CXCL10, GBP1, ISG20, IFNG, BATF2, GBP2, CCL2, IRF1, AMCF-II, S100A12, CXCL13, IL1B, MMP8, MX1, ISG15, CXCL8, S100A8, CCL4, RSAD2, IL27, MX2, IFIT1, IL1A, IRF7, DHX58, SAMHD1, STAT1, ZBP1, OAS1, RIGI, OAS2, BST2, LBP, CALHM6, CCL8, PTGIR, GSDMA, FOSL1, RNF19B, IFIH1, BPIFB1, TNFAIP3, TIFA, NMI, SLC15A3, IRF8, CFB, IFIT2, PIM1, FFAR2, CD40, SLA-DQB1, NOD2, IFI35, TICAM2, PPBP, CCL5, ZFP36, HP, STAT2, NFKBIB, C2, IL10, TRIM14, C3, NFKBIA, IFIT5, LTF, C4A, CD180, DUSP10,* |

| | | | | | | |
|---|---|---|---|---|---|---|
| | | | | | | *MLKL, CXCL16, CEBPG, TREM1, NDUFAF4, BPIFA1, TRAF2, DDIT3, UNC93B1, SERPING1, TRAF3, DDX21, TRIM26, TIGAR, CD14, MSRB1, POLR3D, IL18RAP, OPTN, CEBPB, TICAM1, MYD88, TXNIP, SELENOS, GNLY, CLEC7A, PLSCR1, TLR4, TNIP2, CGAS, KLRK1, MORC3, ARG2, DDIT4, NKG7, LYAR, RIPK3, RNF115, TMEM126A, TLR2, TIRAP, BMP2, CDC42EP2, STING1, IFNGR1, IRAK4, MR1, LOC100516039, HSP90AA1, LY96, SFTPD, TLR3, JAK2, CD46, KCNK6* |
| GO:0098542 | Defense response to other organism | 304 | 2.739 | 1.00E-10 | 1.87E-09 | *CXCL9, CXCL10, GBP1, ISG20, IFNG, BATF2, GBP2, CCL2, IRF1, AMCF-II, S100A12, CXCL13, MX1, ISG15, CXCL8, CCL4, RSAD2, IL27, MX2, IRF7, DHX58, SAMHD1, STAT1, ZBP1, OAS1, RIGI, OAS2, BST2, LBP, CALHM6, CCL8, GSDMA, FOSL1, RNF19B, IFIH1, BPIFB1, TNFAIP3, TIFA, NMI, SLC15A3, IRF8, CFB, PIM1, FFAR2, CD40, SLA-DQB1, NOD2, IFI35, TICAM2, PPBP, CCL5, HP, STAT2, C2, IL10, TRIM14, C3, NFKBIA, LTF, C4A, DUSP10, MLKL, CXCL16, CEBPG, TREM1, NDUFAF4, BPIFA1, TRAF2, UNC93B1, SERPING1, TRAF3, TRIM26, CD14, MSRB1, POLR3D, IL18RAP, OPTN, CEBPB, TICAM1, MYD88, GNLY, PLSCR1, TLR4, TNIP2, CGAS, KLRK1, MORC3, ARG2, DDIT4, NKG7, LYAR, RIPK3, RNF115, TMEM126A, TLR2, TIRAP, CDC42EP2, STING1, IFNGR1, IRAK4, MR1, LOC100516039, HSP90AA1, LY96, SFTPD, TLR3, JAK2, CD46, KCNK6* |
| GO:0071345 | Cellular response to cytokine stimulus | 199 | 2.726 | 1.00E-10 | 1.87E-09 | *CXCL9, CXCL11, CXCL10, IFNG, ACKR1, CCL2, IRF1, AMCF-II, CXCL13, IL1B, CHI3L1, MMP8, MX1, ISG15, CXCL8, CCL4, IL6, IL1A, TNFSF13B, IRF7, STAT1, ZBP1, OAS1, SOCS1, GFPT2, CCL8, FOSL1, TIFA, SNX10, IL15, IRF8, PIM1, IL4R, CD40, SLA-DQB1, MSC, TICAM2, PPBP, CCL5, NFIL3, ZFP36, STAT2, STAT5A, IL10, NFKBIA, CISH, STAT3, EGR1, TRAF2, TRAF3, IL18RAP, YBX3, CACYBP, JAK3, TLR4, ADAM17, TNFRSF1A, KIT, STIP1, TLR2, CDC42EP2, IFNGR1* |
| GO:0044419 | Biological process involved in interspecies | 399 | 2.710 | 1.00E-10 | 1.87E-09 | *CXCL9, CXCL10, GBP1, ISG20, IFNG, BATF2, GBP2, CCL2, IRF1, AMCF-II, S100A12, CXCL13, IL1B, MMP8, MX1, ISG15, CXCL8, S100A8, CCL4, RSAD2, IL27, MX2, IFIT1, IL1A, IRF7, DHX58, SAMHD1, STAT1, ZBP1, OAS1, RIGI, OAS2, BST2, LBP, CALHM6, CCL8, PTGIR, GSDMA, FOSL1, RNF19B, IFIH1, BPIFB1, TNFAIP3, TIFA, NMI, SLC15A3, IRF8,* |

| | | | | | | |
|---|---|---|---|---|---|---|
| | interaction between organisms | | | | | *CFB, IFIT2, PIM1, FFAR2, CD40, SLA-DQB1, NOD2, IFI35, TICAM2, PPBP, CCL5, ZFP36, HP, STAT2, NFKBIB, C2, IL10, TRIM14, C3, NFKBIA, IFIT5, LTF, C4A, CD180, DUSP10, MLKL, CXCL16, CEBPG, TREM1, NDUFAF4, BPIFA1, TRAF2, UNC93B1, SERPING1, ST6GALNAC1, TRAF3, DDX21, TRIM26, CD14, MSRB1, POLR3D, IL18RAP, OPTN, CEBPB, PRF1, TICAM1, MYD88, SELENOS, GNLY, CLEC7A, PLSCR1, TLR4, TNIP2, CGAS, KLRK1, MORC3, ARG2, DDIT4, NKG7, LYAR, RIPK3, RNF115, TMEM126A, TLR2, TIRAP, BMP2, CDC42EP2, STING1, IFNGR1, IRAK4, MR1, LOC100516039, HSP90AA1, LY96, SFTPD, TLR3, JAK2, CD46, KCNK6* |
| GO:0140546 | Defense response to symbiont | 264 | 2.694 | 1.00E-10 | 1.87E-09 | *CXCL9, CXCL10, GBP1, ISG20, IFNG, GBP2, CCL2, IRF1, AMCF-II, S100A12, CXCL13, MX1, ISG15, CXCL8, CCL4, RSAD2, IL27, IRF7, DHX58, SAMHD1, STAT1, ZBP1, OAS1, RIGI, OAS2, LBP, CALHM6, CCL8, FOSL1, RNF19B, IFIH1, BPIFB1, TNFAIP3, TIFA, NMI, SLC15A3, IRF8, CFB, PIM1, FFAR2, SLA-DQB1, NOD2, IFI35, TICAM2, PPBP, CCL5, STAT2, C2, TRIM14, C3, NFKBIA, LTF, C4A, DUSP10, CXCL16, CEBPG, TREM1, BPIFA1, TRAF2, UNC93B1, SERPING1, TRAF3, TRIM26, CD14, MSRB1, POLR3D, IL18RAP, TICAM1, MYD88, PLSCR1, TLR4, TNIP2, CGAS, KLRK1, MORC3, LYAR, RNF115, TMEM126A, TLR2, TIRAP, CDC42EP2, STING1, IFNGR1, IRAK4, MR1, LOC100516039, HSP90AA1, LY96, SFTPD, TLR3, JAK2, CD46, KCNK6* |
| GO:0006955 | Immune response | 468 | 2.677 | 1.00E-10 | 1.87E-09 | *CXCL9, CXCL11, CXCL10, GBP1, ISG20, IFNG, CCL19, GBP2, CCL2, IRF1, AMCF-II, CXCL2, S100A12, CCL3L1, CXCL13, IL1B, MX1, ISG15, CXCL8, CCL4, RSAD2, IL27, IL6, IL1A, TNFSF13B, IRF7, DHX58, SAMHD1, STAT1, ZBP1, OAS1, FAS, RIGI, OAS2, LOC100622460, CSF3, LBP, CALHM6, CCL26, CCL11, CCL8, FOSL1, RNF19B, IFIH1, SLA-DOA, BPIFB1, TNFAIP3, TIFA, TNFSF10, THBS1, SLA-DQA1, SLA-DRA, NMI, IL15, SLA-3, SLC15A3, CTLA4, IRF8, CFB, SLA-DOB, PIM1, FFAR2, CD40, SLA-DQB1, NOD2, IFI35, TICAM2, PPBP, CCL5, ACKR4, NFIL3, STAT2, SLA-DRB1, C2, CD74, XCL1, IL10, TRIM14, SLA-DMB, C3, NFKBIA, LTF, TFRC, TNFSF14, C4A, LIF, CCR1, TNFSF13, CD180, DUSP10, FASLG, STAT3, CSF2, CCL28, CXCL16, CEBPG, SLA-DMA, TREM1, CXCR6, BPIFA1,* |

| | | | | | | |
|---|---|---|---|---|---|---|
| | | | | | | *TRAF2, ULBP1, UNC93B1, SERPING1, PLA2G1B, TCIM, TRAF3, DDX21, ACKR2, TRIM26, BCL6, MAD2L2, CD14, MSRB1, POLR3D, IL18RAP, TNFSF9, TICAM1, MYD88, TNFAIP1, TGFB3, SELENOS, JAK3, PLSCR1, TLR4, TNIP2, CGAS, KLRK1, MORC3, ARG2, KIT, LYAR, RIPK3, RNF115, TMEM126A, PSMB4, TLR2, ST3GAL1, TIRAP, CDC42EP2, CFI, STING1, IFNGR1, IRAK4, MR1, LOC100516039, HSP90AA1, LY96, SLA-8, SFTPD, TLR3, JAK2* |
| GO:0006952 | Defense response | 450 | 2.665 | 1.00E-10 | 1.87E-09 | *CXCL9, CXCL11, SAA3, CXCL10, GBP1, ISG20, IFNG, BATF2, CCL19, ACKR1, GBP2, CCL2, IRF1, AMCF-II, CXCL2, S100A12, CXCL13, IL1B, CHI3L1, MMP8, MX1, ISG15, CXCL8, S100A8, CCL4, RSAD2, IL27, MX2, IL6, IL1A, IRF7, DHX58, SAMHD1, STAT1, ZBP1, OAS1, RIGI, OAS2, BST2, LBP, CALHM6, CCL11, CCL8, PTGIR, GSDMA, FOSL1, RNF19B, IFIH1, BPIFB1, TNFAIP3, TIFA, THBS1, NMI, SLC15A3, IRF8, CFB, PIM1, IL4R, FFAR2, CD40, SLA-DQB1, NOD2, IFI35, TICAM2, PPBP, CCL5, ZFP36, HP, STAT2, SLC39A8, STAT5A, NFKBIB, C2, CD74, IL10, TRIM14, IL1RAP, C3, NFKBIA, LTF, C4A, CCR1, DUSP10, STAT3, MLKL, CXCL16, CEBPG, TREM1, NDUFAF4, IL2RA, CXCR6, BPIFA1, TRAF2, UNC93B1, SERPING1, TRAF3, TRIM26, OLR1, CD163, BCL6, CASP1, CD14, MSRB1, POLR3D, IL18RAP, OPTN, CEBPB, TICAM1, MYD88, SELENOS, B4GALT1, GNLY, PLSCR1, TLR4, TNIP2, CGAS, KLRK1, MORC3, TNFRSF1A, ARG2, DDIT4, NKG7, KIT, LYAR, RIPK3, RNF115, TMEM126A, PSMB4, TLR2, TIRAP, BMP2, CDC42EP2, STING1, IFNGR1, IRAK4, MR1, TNFRSF1B, LOC100516039, GPR4, HSP90AA1, LY96, SFTPD, TLR3, JAK2, ELF4, CD46, KCNK6* |
| GO:0045087 | Innate immune response | 248 | 2.572 | 1.00E-10 | 1.87E-09 | *CXCL10, GBP1, ISG20, IFNG, GBP2, CCL2, IRF1, S100A12, MX1, ISG15, RSAD2, IL27, IRF7, DHX58, SAMHD1, STAT1, ZBP1, OAS1, RIGI, OAS2, LBP, CALHM6, FOSL1, RNF19B, IFIH1, BPIFB1, TNFAIP3, TIFA, NMI, SLC15A3, IRF8, CFB, PIM1, FFAR2, SLA-DQB1, NOD2, IFI35, TICAM2, CCL5, STAT2, C2, TRIM14, C3, NFKBIA, LTF, C4A, DUSP10, CXCL16, CEBPG, TREM1, BPIFA1, TRAF2, UNC93B1, SERPING1, TRAF3, TRIM26, CD14, MSRB1, POLR3D, IL18RAP, TICAM1, MYD88, PLSCR1, TLR4, TNIP2, CGAS, KLRK1, MORC3, LYAR,* |

| | | | | | | |
|---|---|---|---|---|---|---|
| | | | | | | *RNF115, TMEM126A, TLR2, TIRAP, CDC42EP2, STING1, IFNGR1, IRAK4, MR1, HSP90AA1, LY96, SFTPD, TLR3, JAK2, CD46, KCNK6* |
| GO:0022613 | Ribonucleoprotein complex biogenesis | 172 | 2.440 | 1.00E-10 | 1.87E-09 | *ISG20, NPM3, EMG1, RRP9, NHP2, SNRPF, GAR1, FBL, MRTO4, SNRPD1, NOB1, PPAN, RRS1, NOP2, LYAR, EXOSC5, NLE1, GRWD1, WDR75, NOC4L, WDR74, UTP11, GTPBP4, PTGES3, NIP7, REXO4, BUD23, RSL24D1, EIF3I, NOL6, IMP3, NOP9, BMS1, NOPCHAP1, WDR12, RCL1, RRP36, EIF3J, NAT10, RPL7L1, NOP56, WDR46, DDX49, RRP15, SNRPG, EXOSC3, ABT1, UTP18, UTP23, EBNA1BP2, MTERF3, SHQ1, PNO1, SNRPD3, DDX3X, ESF1, GEMIN6, DDX52, NPM1, MRPS7, EIF6, RPP40, UTP6, STRAP, SDAD1, TMA16, UTP15, TSR2, AK6, NUFIP1, PWP1, WDR36, TAF9, CLNS1A, EIF3A, SNRPC, USP39, EXOSC4, NOL9, FCF1, WDR55, SRSF1, KRI1, EIF4A3, SNRPE, LSM6, DDX31, RPL27, TRMT112, LOC100621115, FASTKD2, EIF3M, LOC100738684, SNU13, MTREX, MRM1, GTPBP10, KRR1, DENR, DDX28* |
| GO:0042254 | Ribosome biogenesis | 129 | 2.425 | 1.00E-10 | 1.87E-09 | *ISG20, NPM3, EMG1, RRP9, NHP2, GAR1, FBL, MRTO4, NOB1, PPAN, RRS1, NOP2, LYAR, EXOSC5, NLE1, GRWD1, WDR75, NOC4L, WDR74, UTP11, GTPBP4, NIP7, REXO4, BUD23, RSL24D1, NOL6, IMP3, NOP9, BMS1, WDR12, RCL1, RRP36, NAT10, RPL7L1, NOP56, WDR46, DDX49, RRP15, EXOSC3, ABT1, UTP18, UTP23, EBNA1BP2, MTERF3, PNO1, DDX3X, ESF1, DDX52, NPM1, MRPS7, EIF6, RPP40, UTP6, SDAD1, TMA16, UTP15, TSR2, AK6, PWP1, WDR36, EXOSC4, NOL9, FCF1, WDR55, KRI1, EIF4A3, LSM6, DDX31, RPL27, TRMT112, LOC100621115, FASTKD2, SNU13, MTREX, MRM1, GTPBP10* |
| GO:0006954 | Inflammatory response | 217 | 2.419 | 1.00E-10 | 1.87E-09 | *CXCL9, SAA3, CXCL10, IFNG, CCL19, ACKR1, CCL2, AMCF-II, CXCL2, S100A12, CXCL13, IL1B, CHI3L1, MMP8, CXCL8, S100A8, CCL4, IL27, IL6, IL1A, CCL11, CCL8, PTGIR, GSDMA, FOSL1, TNFAIP3, THBS1, IL4R, FFAR2, IFI35, TICAM2, PPBP, CCL5, ZFP36, HP, SLC39A8, STAT5A, NFKBIB, CD74, IL10, TRIM14, IL1RAP, C3, NFKBIA, C4A, CCR1, DUSP10, STAT3, IL2RA, CXCR6, OLR1, CD163, BCL6, CASP1, CD14, CEBPB, TICAM1, MYD88, SELENOS, B4GALT1, TLR4, TNFRSF1A, KIT, PSMB4, TLR2, TIRAP, BMP2, TNFRSF1B, LOC100516039, GPR4, TLR3, JAK2, ELF4, KCNK6* |

| | | | | | | |
|---|---|---|---|---|---|---|
| GO:0033993 | Response to lipid | 156 | 2.331 | 1.00E-10 | 1.87E-09 | *CXCL9, CXCL10, AMCF-II, CXCL13, IL1B, MMP8, CXCL8, S100A8, IL1A, PTGIR, TNFAIP3, AQP3, JUNB, IRF8, PIM1, FFAR2, AARD, PPBP, ZFP36, NFKBIB, IL10, NFKBIA, LTF, CD180, DUSP10, CEBPB, TICAM1, TGFB3, SELENOS, TLR4, TNIP2, KLRK1, CALR, DDIT4, GOT1, INSIG1, GPR155, TIRAP, PTGES3, LY96, VDR, JAK2, KLF9, CRY1, GDAP1, MGST1, NKX3-1, PLSCR4, PLSCR3* |
| GO:0009617 | Response to bacterium | 128 | 2.287 | 2.43E-09 | 2.88E-08 | *CXCL9, CXCL10, AMCF-II, CXCL13, IL1B, MMP8, ISG15, CXCL8, S100A8, IL1A, DHX58, LBP, PTGIR, GSDMA, TNFAIP3, IRF8, PPBP, ZFP36, HP, NFKBIB, IL10, NFKBIA, LTF, CD180, DUSP10, TREM1, BPIFA1, OPTN, CEBPB, TICAM1, MYD88, SELENOS, GNLY, TLR4, TNIP2, KLRK1, ARG2, TLR2, TIRAP, BMP2* |
| GO:0002831 | Regulation of response to biotic stimulus | 139 | 2.287 | 4.23E-10 | 6.27E-09 | *IRF1, ISG15, RSAD2, IRF7, DHX58, SAMHD1, ZBP1, CALHM6, FOSL1, BPIFB1, TNFAIP3, TIFA, SLC15A3, PIM1, FFAR2, IFI35, TICAM2, STAT2, NFKBIA, LTF, CD180, DUSP10, UNC93B1, SERPING1, TRAF3, TIGAR, POLR3D, IL18RAP, OPTN, TICAM1, MYD88, PLSCR1, TLR4, TNIP2, CGAS, KLRK1, ARG2, LYAR, RNF115, TMEM126A, TLR2, TIRAP, STING1, HSP90AA1, LY96, CD46, KCNK6* |
| GO:0001819 | Positive regulation of cytokine production | 134 | 2.244 | 3.14E-09 | 3.61E-08 | *IFNG, IRF1, IL1B, CHI3L1, MMP8, ISG15, RSAD2, IL1A, IRF7, DHX58, STAT1, THBS1, IL15, IRF8, FFAR2, CD40, TICAM2, STAT5A, CD74, IL10, CEBPG, PANX2, EGR1, TRAF2, DDIT3, PLA2G1B, DDX21, CASP1, ADRA2A, POLR3D, CEBPB, CASP8, TICAM1, MYD88, TLR4, CGAS, KLRK1, KIT, TMEM106A, TLR2, TIRAP, STING1, HSP90AA1, LY96, JAK2, HILPDA, RAB1A, TMED10, KAT2A, IL16, HAVCR2, UAP1* |
| GO:0045088 | Regulation of innate immune response | 126 | 2.244 | 3.40E-09 | 3.78E-08 | *IRF1, ISG15, RSAD2, IRF7, DHX58, SAMHD1, ZBP1, CALHM6, FOSL1, BPIFB1, TNFAIP3, TIFA, SLC15A3, PIM1, FFAR2, IFI35, TICAM2, STAT2, NFKBIA, DUSP10, UNC93B1, SERPING1, POLR3D, IL18RAP, TICAM1, MYD88, PLSCR1, TLR4, TNIP2, CGAS, KLRK1, LYAR, RNF115, TMEM126A, TLR2, TIRAP, STING1, HSP90AA1, LY96, CD46, KCNK6* |
| GO:0071396 | Cellular response to lipid | 120 | 2.228 | 2.86E-08 | 2.61E-07 | *CXCL9, CXCL10, AMCF-II, CXCL13, IL1B, MMP8, CXCL8, IL1A, TNFAIP3, IRF8, PIM1, FFAR2, PPBP, ZFP36, NFKBIB, IL10, NFKBIA, LTF, CD180, TICAM1, SELENOS, TNIP2,* |

| | | | | | | |
|---|---|---|---|---|---|---|
| | | | | | | *KLRK1, CALR, DDIT4, INSIG1, GPR155, PTGES3, LY96, VDR, JAK2, KLF9, CRY1, MGST1, NKX3-1, PLSCR4, PLSCR3, HAVCR2* |
| GO:0002682 | Regulation of immune system process | 390 | 2.217 | 1.00E-10 | 1.87E-09 | *CXCL10, MADCAM1, IFNG, SELE, CD274, CCL2, IRF1, IL1B, MMP8, ISG15, RSAD2, IL27, IL1A, TNFSF13B, IRF7, DHX58, SAMHD1, STAT1, ZBP1, LOC100622460, CSF3, SOCS1, CALHM6, FOSL1, BPIFB1, TNFAIP3, TIFA, THBS1, AQP3, IL15, SLC15A3, CTLA4, CFB, PIM1, FFAR2, CD40, SLA-DQB1, IFI35, TICAM2, CCL5, GDF15, ZFP36, STAT2, STAT5A, C2, IL10, SLA-DMB, C3, NFKBIA, LTF, TFRC, TNFSF14, C4A, LIF, CCR1, DUSP10, STAT3, LRRC32, CEACAM1, TRAF2, ULBP1, UNC93B1, SERPING1, PLA2G1B, TCIM, HSPA9, DDX21, SOX11, BCL6, MAD2L2, SLC7A1, POLR3D, IL18RAP, CEBPB, TNFSF9, CASP8, TICAM1, MYD88, FBXO7, TGFB3, SELENOS, TMEM176B, JAK3, PLSCR1, TLR4, TNIP2, CGAS, KLRK1, CALR, ARG2, KIT, LYAR, RIPK3, RNF115, TMEM126A, MYC, PSMB4, TLR2, IGFBP2, TIRAP, CFI, STING1* |
| GO:0045321 | Leukocyte activation | 226 | 2.157 | 1.62E-10 | 2.75E-09 | *IFNG, BATF2, CD274, IRF1, AMCF-II, IL1B, MMP8, CXCL8, RSAD2, IL27, IL6, IL1A, TNFSF13B, SOCS1, TNFAIP3, THBS1, IL15, CTLA4, IRF8, CD40, SLA-DQB1, IFI35, CCL5, NFIL3, STAT5A, IL10, HSH2D, SLA-DMB, TFRC, TNFSF14, GJA1, CTPS1, CD180, DUSP10, STAT3, LRRC32, CEBPG, EGR1, TRAF2, SOX11, BCL6, MAD2L2, SLC7A1, IL18RAP, CEBPB, TNFSF9, TICAM1, FBXO7, JAK3, PLSCR1, TLR4, KLRK1, ARG2, KIT, TMEM106A, RIPK3, ST3GAL1, IGFBP2, PRDM1, DLL4, JAK2, ELF4, CD200, LIG4, KAT2A, SLC25A5* |
| GO:0001775 | Cell activation | 261 | 2.144 | 1.26E-10 | 2.25E-09 | *CXCL10, IFNG, BATF2, CD274, IRF1, AMCF-II, IL1B, MMP8, CXCL8, RSAD2, IL27, IL6, IL1A, TNFSF13B, SOCS1, TNFAIP3, THBS1, IL15, CTLA4, IRF8, CD40, SLA-DQB1, IFI35, CCL5, NFIL3, STAT5A, FGB, IL10, HSH2D, SLA-DMB, TFRC, TNFSF14, GJA1, CTPS1, CD180, DUSP10, STAT3, LRRC32, CEBPG, EGR1, TRAF2, TCIM, SOX11, BCL6, MAD2L2, ADRA2A, SLC7A1, IL18RAP, CEBPB, TNFSF9, TICAM1, FBXO7, JAK3, PLSCR1, TLR4, KLRK1, ARG2, KIT, TMEM106A, RIPK3, ST3GAL1, IGFBP2, PRDM1, DLL4, JAK2, ELF4, GNA13* |

| | | | | | | |
|---|---|---|---|---|---|---|
| GO:0050776 | Regulation of immune response | 232 | 2.144 | 2.72E-10 | 4.40E-09 | *IFNG, IRF1, IL1B, ISG15, RSAD2, IL27, TNFSF13B, IRF7, DHX58, SAMHD1, ZBP1, LOC100622460, CALHM6, FOSL1, BPIFB1, TNFAIP3, TIFA, IL15, SLC15A3, CFB, PIM1, FFAR2, CD40, SLA-DQB1, IFI35, TICAM2, STAT2, C2, IL10, SLA-DMB, C3, NFKBIA, TFRC, C4A, DUSP10, TRAF2, ULBP1, UNC93B1, SERPING1, PLA2G1B, BCL6, MAD2L2, POLR3D, IL18RAP, TICAM1, MYD88, SELENOS, JAK3, PLSCR1, TLR4, TNIP2, CGAS, KLRK1, ARG2, KIT, LYAR, RIPK3, RNF115, TMEM126A, PSMB4, TLR2, TIRAP, CFI, STING1, HSP90AA1, LY96, CD46, KCNK6* |
| GO:0031347 | Regulation of defense response | 201 | 2.137 | 1.49E-09 | 1.97E-08 | *IFNG, IRF1, MMP8, ISG15, S100A8, RSAD2, IRF7, DHX58, SAMHD1, ZBP1, CALHM6, FOSL1, BPIFB1, TNFAIP3, TIFA, SLC15A3, PIM1, FFAR2, IFI35, TICAM2, ZFP36, STAT2, SLC39A8, STAT5A, CD74, IL10, NFKBIA, DUSP10, UNC93B1, SERPING1, TRAF3, BCL6, CASP1, POLR3D, IL18RAP, OPTN, CEBPB, TICAM1, MYD88, SELENOS, PLSCR1, TLR4, TNIP2, CGAS, KLRK1, TNFRSF1A, ARG2, LYAR, RNF115, TMEM126A, PSMB4, TLR2, TIRAP, STING1, GPR4, HSP90AA1, LY96, JAK2, ELF4, CD46, KCNK6* |
| GO:0002684 | Positive regulation of immune system process | 265 | 2.134 | 4.01E-10 | 6.21E-09 | *CXCL10, MADCAM1, IFNG, IRF1, IL1B, MMP8, RSAD2, IL1A, TNFSF13B, IRF7, DHX58, ZBP1, SOCS1, FOSL1, BPIFB1, TNFAIP3, TIFA, THBS1, AQP3, IL15, SLC15A3, CFB, FFAR2, CD40, SLA-DQB1, IFI35, TICAM2, CCL5, STAT5A, C2, IL10, SLA-DMB, C3, NFKBIA, TFRC, TNFSF14, C4A, LIF, CCR1, DUSP10, TRAF2, ULBP1, UNC93B1, SERPING1, PLA2G1B, DDX21, BCL6, MAD2L2, SLC7A1, POLR3D, IL18RAP, TNFSF9, CASP8, TICAM1, MYD88, PLSCR1, TLR4, TNIP2, CGAS, KLRK1, CALR, KIT, RNF115, TMEM126A, TLR2, IGFBP2, TIRAP, CFI, STING1, HSP90AA1, LY96, JAK2, CD46, KCNK6* |
| GO:0032103 | Positive regulation of response to external stimulus | 166 | 2.124 | 6.90E-08 | 5.34E-07 | *CXCL10, IFNG, IRF1, MMP8, CXCL8, S100A8, RSAD2, IRF7, DHX58, ZBP1, FOSL1, BPIFB1, TNFAIP3, TIFA, THBS1, SLC15A3, FFAR2, IFI35, TICAM2, CCL5, STAT5A, CD74, NFKBIA, TNFSF14, CCR1, CD180, UNC93B1, POLR3D, IL18RAP, OPTN, CEBPB, TICAM1, MYD88, PLSCR1, TLR4, TNIP2, CGAS, KLRK1, TNFRSF1A, CALR, RNF115, TMEM126A, TLR2, TIRAP, STING1, GPR4, HSP90AA1, LY96* |

| | | | | | | |
|---|---|---|---|---|---|---|
| GO:1903131 | Mononuclear cell differentiation | 123 | 2.101 | 1.54E-06 | 8.43E-06 | *IFNG, BATF2, IRF1, IL1B, RSAD2, IL27, IL6, IL1A, TNFSF13B, SOCS1, IRF8, SLA-DQB1, NFIL3, STAT5A, IL10, LIF, DUSP10, STAT3, CEBPG, EGR1, BCL6, CEBPB, TNFSF9, CASP8, FBXO7, TMEM176B, JAK3, TLR4, KIT, RIPK3, MYC, ST3GAL1, PRDM1, DLL4, LIG4, KAT2A, SLC25A5* |
| GO:0032101 | Regulation of response to external stimulus | 258 | 2.088 | 1.93E-09 | 2.37E-08 | *CXCL10, IFNG, CCL2, IRF1, MMP8, ISG15, CXCL8, S100A8, RSAD2, SERPINE1, IRF7, DHX58, SAMHD1, ZBP1, CALHM6, FOSL1, BPIFB1, TNFAIP3, TIFA, THBS1, SLC15A3, PIM1, FFAR2, IFI35, TICAM2, CCL5, ZFP36, STAT2, SLC39A8, STAT5A, CD74, IL10, NFKBIA, LTF, TNFSF14, CCR1, CD180, DUSP10, UNC93B1, SERPING1, TRAF3, BCL6, CASP1, POLR3D, IL18RAP, OPTN, CEBPB, TICAM1, MYD88, SELENOS, PLSCR1, TLR4, TNIP2, CGAS, KLRK1, TNFRSF1A, CALR, ARG2, LYAR, RNF115, TMEM126A, PSMB4, TLR2, TIRAP, STING1, GPR4, HSP90AA1, LY96, JAK2, ELF4, CD46, KCNK6* |
| GO:0031349 | Positive regulation of defense response | 138 | 2.084 | 5.25E-07 | 3.34E-06 | *IFNG, IRF1, MMP8, S100A8, RSAD2, IRF7, DHX58, ZBP1, FOSL1, BPIFB1, TNFAIP3, TIFA, SLC15A3, FFAR2, IFI35, TICAM2, STAT5A, CD74, NFKBIA, UNC93B1, POLR3D, IL18RAP, OPTN, CEBPB, TICAM1, MYD88, PLSCR1, TLR4, TNIP2, CGAS, KLRK1, TNFRSF1A, RNF115, TMEM126A, TLR2, TIRAP, STING1, GPR4, HSP90AA1, LY96* |
| GO:0001816 | Cytokine production | 207 | 2.079 | 8.15E-09 | 8.54E-08 | *IFNG, IRF1, IL1B, CHI3L1, MMP8, ISG15, RSAD2, IL6, IL1A, IRF7, DHX58, STAT1, SOCS1, CALHM6, TNFAIP3, THBS1, IL15, IRF8, FFAR2, CD40, TICAM2, ZFP36, STAT5A, CD74, IL10, TSKU, LTF, LRRC32, CSF2, CEBPG, PANX2, EGR1, TRAF2, DDIT3, PLA2G1B, TRAF3, DDX21, BCL6, CASP1, ADRA2A, POLR3D, CEBPB, CASP8, TICAM1, MYD88, TGFB3, TLR4, CGAS, KLRK1, MORC3, ARG2, KIT, TMEM106A, RIPK3, TLR2, TIRAP, STING1, HSP90AA1, LY96, JAK2, HILPDA, ELF4, RAB1A, TMED10* |
| GO:0002694 | Regulation of leukocyte activation | 134 | 2.074 | 2.96E-07 | 2.07E-06 | *IFNG, CD274, IRF1, IL1B, MMP8, IL27, IL1A, TNFSF13B, SOCS1, TNFAIP3, THBS1, IL15, CTLA4, CD40, SLA-DQB1, CCL5, STAT5A, IL10, SLA-DMB, TFRC, TNFSF14, DUSP10, LRRC32, SOX11, BCL6, MAD2L2, SLC7A1, CEBPB, TNFSF9, TICAM1, FBXO7, JAK3, PLSCR1, TLR4, ARG2, RIPK3, IGFBP2* |

| | | | | | | |
|---|---|---|---|---|---|---|
| GO:0006396 | RNA processing | 357 | 2.070 | 1.00E-10 | 1.87E-09 | *ISG20, NPM3, RNASEL, QTRT1, EMG1, RRP9, WDR4, NHP2, POP1, SNRPF, GAR1, FBL, QTRT2, SNRPD1, PAPOLA, NOB1, PPAN, RRS1, NOP2, LYAR, PNPT1, EXOSC5, DKC1, PUS3, WDR75, RTCA, WDR74, UTP11, GTPBP4, ZPR1, RBM8A, SNRPA1, REXO4, TRMT6, BUD23, NOL6, SMNDC1, IMP3, NOP9, CCNL1, RTCB, AGO2, WDR12, TYW1, RCL1, RRP36, NAT10, RPL7L1, WDR46, DDX49, RRP15, SNRPG, EXOSC3, ABT1, UTP18, UTP23, SNRNP27, SNRPD3, SLC38A2, TBRG4, NSUN2, TRMT11, TRA2B, ESF1, GEMIN6, DDX52, PUS1, SMN1, SNRPA, RPRD1B, HSD17B10, RPP40, UTP6, ZMAT2, STRAP, MAGOHB, UTP15, THUMPD1, TSR2, RAMAC, AK6, WTAP, SF3B6, ISG20L2, PWP1, WDR36, PRPF40A, CLNS1A, METTL8, FMR1, SNRPC, SNRPB2, SMU1, USP39, EXOSC4, TRMT2A, TYW5, NOL9, FCF1, WDR55, ELP5, HNRNPF, CSTF2, SRSF1, FTSJ1, KRI1, EIF4A3, RPP21, RBM7, MTO1, SSB, SNRPE, SNRNP40, SRSF7, LSM6, RBM14, MRPL44, RPL27, DCPS, TRMT112, PRPF19, RNPS1, TSEN15, RBM47, CSTF3, MAGOH, FASTKD2, TP53RK, METTL3, NSRP1, SNU13, RBMX2, MTREX, RPP30, MRM1, RBM25, WDR83, KRR1, TRMT5, TPRKB, RBM42, TENT2, CDK12, INTS8, LSM5, SCAF11, SLX9, CSTF2T, LEO1, AHCYL1, GEMIN8, U2AF1L4, CWC22, SBDS, DTWD1, RBM39, TYW3, MBNL1, THOC7, PRPF3, EXOSC2, RBBP6, SYMPK, NAF1, UBL5, ELP4, ELP2, ZRANB2, PPIL1, SRSF2, LSM8, C1H9orf78, RPP25L, HNRNPK, PTBP1, BUD31, PAN2, TSN, GEMIN2, SDE2, PCF11, RPP38, PTBP3, INTS10, NSUN4, NOP10, MTFMT, DIMT1, RBM15, NCBP2, ZBTB8OS, PABPC1, C1D, U2AF2, SSU72, TGS1, RBM41, POP4, ALKBH8, MRM2, SF3A3, LSM2, U2SURP, RBMX, CLP1, EXOSC1, SON, BCDIN3D* |
| GO:0002521 | Leukocyte differentiation | 147 | 2.068 | 1.66E-07 | 1.20E-06 | *IFNG, BATF2, IRF1, IL1B, RSAD2, IL27, IL6, IL1A, TNFSF13B, SOCS1, SNX10, JUNB, IRF8, SLA-DQB1, NFIL3, STAT5A, IL10, LTF, LIF, CCR1, DUSP10, STAT3, CEBPG, EGR1, BCL6, CEBPB, TNFSF9, CASP8, FBXO7, TMEM176B, JAK3, TLR4, KIT, RIPK3, MYC, ST3GAL1, BMP2, PRDM1, DLL4, LIG4, KAT2A, SLC25A5, KLF10, RASSF2, TFE3, IL6R, CYP26B1, RUNX1, LRRC8A, USP44* |

| | | | | | | |
|---|---|---|---|---|---|---|
| GO:0006935 | Chemotaxis | 121 | 2.065 | 1.97E-06 | 1.03E-05 | *CXCL9, CXCL11, CXCL10, CCL2, AMCF-II, CXCL13, CXCL8, S100A8, CCL4, CCL8, THBS1, FFAR2, PPBP, CCL5, ACKR4, TNFSF14, CCR1, CXCL16, TREM1, CXCR6, PLA2G1B, ACKR2, KLRK1, CALR, KIT* |
| GO:0042330 | Taxis | 121 | 2.065 | 1.97E-06 | 1.03E-05 | *CXCL9, CXCL11, CXCL10, CCL2, AMCF-II, CXCL13, CXCL8, S100A8, CCL4, CCL8, THBS1, FFAR2, PPBP, CCL5, ACKR4, TNFSF14, CCR1, CXCL16, TREM1, CXCR6, PLA2G1B, ACKR2, KLRK1, CALR, KIT* |
| GO:0022407 | Regulation of cell-cell adhesion | 103 | 2.051 | 5.44E-06 | 2.65E-05 | *IFNG, SELE, CD274, IRF1, IL1B, IL1A, TNFSF13B, SOCS1, IL15, NR4A3, SLA-DQB1, CCL5, STAT5A, IL10, SLA-DMB, TFRC, TNFSF14, DUSP10, LRRC32, BCL6, MAD2L2, SLC7A1, CEBPB, TNFSF9, JAK3, ARG2, IGFBP2, BMP2, GTPBP4, JAK2* |
| GO:0050865 | Regulation of cell activation | 146 | 2.043 | 1.58E-06 | 8.55E-06 | *IFNG, CD274, IRF1, IL1B, MMP8, IL27, IL1A, TNFSF13B, SOCS1, TNFAIP3, THBS1, IL15, CTLA4, CD40, SLA-DQB1, CCL5, STAT5A, IL10, SLA-DMB, TFRC, TNFSF14, DUSP10, LRRC32, SOX11, BCL6, MAD2L2, SLC7A1, CEBPB, TNFSF9, TICAM1, FBXO7, JAK3, PLSCR1, TLR4, ARG2, RIPK3, IGFBP2* |
| GO:0046649 | Lymphocyte activation | 192 | 2.041 | 9.41E-08 | 7.13E-07 | *IFNG, CD274, IRF1, IL1B, RSAD2, IL27, IL6, IL1A, TNFSF13B, SOCS1, TNFAIP3, IL15, CTLA4, IRF8, CD40, SLA-DQB1, CCL5, NFIL3, STAT5A, IL10, HSH2D, SLA-DMB, TFRC, TNFSF14, GJA1, CTPS1, CD180, DUSP10, STAT3, LRRC32, CEBPG, EGR1, TRAF2, SOX11, BCL6, MAD2L2, SLC7A1, CEBPB, TNFSF9, TICAM1, FBXO7, JAK3, TLR4, KLRK1, ARG2, KIT, RIPK3, ST3GAL1, IGFBP2, PRDM1, DLL4, JAK2, ELF4, LIG4, KAT2A, SLC25A5* |
| GO:0001817 | Regulation of cytokine production | 204 | 2.038 | 3.08E-08 | 2.74E-07 | *IFNG, IRF1, IL1B, CHI3L1, MMP8, ISG15, RSAD2, IL1A, IRF7, DHX58, STAT1, SOCS1, CALHM6, TNFAIP3, THBS1, IL15, IRF8, FFAR2, CD40, TICAM2, ZFP36, STAT5A, CD74, IL10, TSKU, LTF, LRRC32, CEBPG, PANX2, EGR1, TRAF2, DDIT3, PLA2G1B, TRAF3, DDX21, BCL6, CASP1, ADRA2A, POLR3D, CEBPB, CASP8, TICAM1, MYD88, TGFB3, TLR4, CGAS, KLRK1, MORC3, ARG2, KIT, TMEM106A, RIPK3, TLR2, TIRAP, STING1, HSP90AA1, LY96, JAK2, HILPDA, ELF4, RAB1A, TMED10* |

| | | | | | | |
|---|---|---|---|---|---|---|
| GO:0045089 | Positive regulation of innate immune response | 105 | 2.033 | 8.44E-06 | 3.95E-05 | *IRF1, RSAD2, IRF7, DHX58, ZBP1, FOSL1, BPIFB1, TNFAIP3, TIFA, SLC15A3, FFAR2, IFI35, TICAM2, NFKBIA, UNC93B1, POLR3D, IL18RAP, TICAM1, MYD88, PLSCR1, TLR4, TNIP2, CGAS, KLRK1, RNF115, TMEM126A, TLR2, TIRAP, STING1, HSP90AA1, LY96, KCNK6* |
| GO:0042110 | T cell activation | 133 | 2.030 | 1.54E-06 | 8.43E-06 | *IFNG, CD274, IRF1, IL1B, RSAD2, IL27, IL6, IL1A, TNFSF13B, SOCS1, IL15, CTLA4, SLA-DQB1, CCL5, STAT5A, HSH2D, SLA-DMB, TFRC, TNFSF14, GJA1, CTPS1, DUSP10, STAT3, LRRC32, EGR1, TRAF2, BCL6, SLC7A1, CEBPB, TNFSF9, JAK3, ARG2, KIT, RIPK3, IGFBP2, PRDM1, DLL4, JAK2, ELF4, LIG4, KAT2A* |
| GO:0051249 | Regulation of lymphocyte activation | 120 | 2.025 | 5.00E-06 | 2.47E-05 | *IFNG, CD274, IRF1, IL1B, IL27, IL1A, TNFSF13B, SOCS1, TNFAIP3, IL15, CTLA4, CD40, SLA-DQB1, CCL5, STAT5A, IL10, SLA-DMB, TFRC, TNFSF14, DUSP10, LRRC32, SOX11, BCL6, MAD2L2, SLC7A1, CEBPB, TNFSF9, TICAM1, FBXO7, JAK3, TLR4, ARG2, RIPK3, IGFBP2, PRDM1* |
| GO:0002833 | Positive regulation of response to biotic stimulus | 110 | 1.995 | 1.76E-05 | 7.48E-05 | *IRF1, RSAD2, IRF7, DHX58, ZBP1, FOSL1, BPIFB1, TNFAIP3, TIFA, SLC15A3, FFAR2, IFI35, TICAM2, NFKBIA, CD180, UNC93B1, POLR3D, IL18RAP, TICAM1, MYD88, PLSCR1, TLR4, TNIP2, CGAS, KLRK1, RNF115, TMEM126A, TLR2, TIRAP, STING1, HSP90AA1, LY96, KCNK6* |
| GO:0050778 | Positive regulation of immune response | 190 | 1.985 | 3.12E-07 | 2.14E-06 | *IFNG, IRF1, IL1B, RSAD2, TNFSF13B, IRF7, DHX58, ZBP1, FOSL1, BPIFB1, TNFAIP3, TIFA, IL15, SLC15A3, CFB, FFAR2, CD40, SLA-DQB1, IFI35, TICAM2, C2, IL10, SLA-DMB, C3, NFKBIA, TFRC, C4A, TRAF2, ULBP1, UNC93B1, SERPING1, PLA2G1B, MAD2L2, POLR3D, IL18RAP, TICAM1, MYD88, PLSCR1, TLR4, TNIP2, CGAS, KLRK1, RNF115, TMEM126A, TLR2, TIRAP, CFI, STING1, HSP90AA1, LY96, CD46, KCNK6* |
| GO:0002764 | Immune response-regulating signaling pathway | 121 | 1.978 | 8.86E-06 | 4.05E-05 | *IFNG, IRF1, RSAD2, IRF7, DHX58, LOC100622460, FOSL1, BPIFB1, TNFAIP3, TIFA, SLC15A3, FFAR2, CD40, IFI35, TICAM2, NFKBIA, UNC93B1, TICAM1, MYD88, PLSCR1, TLR4, TNIP2, CGAS, KLRK1, KIT, RNF115, TMEM126A, TLR2, TIRAP, STING1, LY96, KCNK6* |
| GO:0043065 | Positive regulation of apoptotic process | 130 | 1.973 | 4.56E-06 | 2.29E-05 | *IFNG, CD274, CCL2, S100A8, STAT1, ATF3, THBS1, CD40, CCL5, IL10, FASLG, RNF122, GADD45G, CASP3, PHLDA2, TRAF2, DDIT3, PLA2G1B, RHOB, HSPA9, BCL6, TIGAR,* |

| | | | | | | |
|---|---|---|---|---|---|---|
| | | | | | | *CASP8, B4GALT1, MCL1, TLR4, TNFRSF12A, BCL2A1, RIPK3, BMP2, GADD45B, UTP11, VDR, TP53, JAK2, INHBA, NKX3-1, RASSF2, PDCD4, G0S2, SIAH1* |
| GO:0080134 | Regulation of response to stress | 333 | 1.972 | 3.76E-09 | 4.06E-08 | *IFNG, IRF1, MMP8, ISG15, S100A8, RSAD2, SERPINE1, IRF7, DHX58, SAMHD1, ZBP1, FAS, CALHM6, FOSL1, BPIFB1, TNFAIP3, TIFA, THBS1, SLC15A3, PIM1, FFAR2, IFI35, TICAM2, ZFP36, STAT2, SLC39A8, STAT5A, CD74, IL10, NFKBIA, DUSP10, SESN2, CEBPG, UNC93B1, SERPING1, TRAF3, BCL6, CASP1, MAD2L2, TIGAR, ADRA2A, POLR3D, IL18RAP, OPTN, CEBPB, TICAM1, MYD88, YBX3, SELENOS, PSMD10, PLSCR1, TLR4, TNIP2, CGAS, KLRK1, TNFRSF1A, ARG2, LYAR, FICD, FBXO4, RNF115, TMEM126A, MYC, PSMB4, TLR2, RNASEH2B, INSIG1, TIRAP, STING1, GPR4, HSP90AA1, LY96, JAK2, MORF4L2, ELF4, CD46, KCNK6, CRY1* |
| GO:0042127 | Regulation of cell population proliferation | 374 | 1.966 | 1.32E-09 | 1.81E-08 | *CXCL9, CXCL11, CXCL10, IFNG, CD274, CCL2, IRF1, IL1B, IL27, IL6, IL1A, TNFSF13B, STAT1, CSF3, PTGIR, TNFAIP3, THBS1, JUNB, IL15, CTLA4, PIM1, TIMP1, CD40, NR4A3, IFI35, CCL5, ZFP36, STAT2, STAT5A, IL10, ATF5, NFKBIA, LTF, TFRC, GJA1, LIF, DUSP10, NME1, STAT3, LRRC32, CSF2, TRNP1, PHLDA2, ESM1, PLA2G1B, SOX11, BCL6, SLC7A1, IFI30, CEBPB, TNFSF9, TICAM1, TGFB3, B4GALT1, TLR4, MORC3, CALR, ARG2, FAM98A, NOP2, SLC25A33, FBXO4, RIPK3, MYC, RNASEH2B, IGFBP2, TIPIN, BMP2, PRDM1, GTPBP4, VDR, DLL4, JAK2, HILPDA, KLF9, SMAD1, INHBA, LIG4, ADM, NKX3-1, SLC25A5, FUT2, PPARGC1A* |
| GO:0010628 | Positive regulation of gene expression | 304 | 1.964 | 9.76E-09 | 9.93E-08 | *IFNG, IRF1, IL1B, CHI3L1, MMP8, ISG15, CXCL8, RSAD2, IL1A, IRF7, DHX58, STAT1, ATF3, THBS1, IL15, IRF8, FFAR2, CD40, TICAM2, NFIL3, ZFP36, STAT5A, CD74, IL10, TFRC, GJA1, LIF, LRRC32, ARID5A, CEBPG, PANX2, EGR1, TRAF2, DDIT3, PLA2G1B, DDX21, SOX11, CASP1, MAD2L2, ADRA2A, POLR3D, CEBPB, CASP8, TICAM1, MYD88, TLR4, CGAS, KLRK1, CALR, KIT, FAM98A, TMEM106A, TLR2, TIRAP, BMP2, STING1, EIF5A, PRDM1, ZPR1, HSP90AA1, LY96, VDR, DLL4, JAK2, HILPDA, RAB1A, SMAD1, TMED10, INHBA, KAT2A, IGF2BP1, NKX3-1, PPARGC1A, WNT16, C7H6orf125, NAT10, IL16,* |

| | | | | | | |
|---|---|---|---|---|---|---|
| | | | | | | *HAVCR2, UAP1, EXOSC3, PLGRKT, SPRY2, CYP26B1, POU2F2, SLC38A2, CLDN5, RUNX1, DDX3X, TRA2B* |
| GO:0002252 | Immune effector process | 159 | 1.961 | 3.65E-06 | 1.86E-05 | *IFNG, IL1B, RSAD2, IL27, IL6, IRF7, DHX58, CALHM6, RNF19B, IRF8, CFB, FFAR2, CD40, SLA-DQB1, IFI35, C2, IL10, C3, TFRC, C4A, CD180, DUSP10, STAT3, CEBPG, TREM1, TRAF2, ULBP1, SERPING1, PLA2G1B, TCIM, DDX21, BCL6, MAD2L2, IL18RAP, PRF1, TICAM1, TGFB3, JAK3, TLR4, KLRK1, KIT, RIPK3, ST3GAL1, CFI* |
| GO:1901700 | Response to oxygen-containing compound | 311 | 1.950 | 1.22E-08 | 1.21E-07 | *CXCL9, CXCL10, AMCF-II, CXCL13, IL1B, MMP8, CXCL8, S100A8, IL1A, STAT1, SOCS1, PTGIR, TNFAIP3, LPL, THBS1, AQP3, IRF8, PIM1, FFAR2, NR4A3, AARD, PPBP, GDF15, ZFP36, STAT2, STAT5A, NFKBIB, IL10, NFKBIA, LTF, GJA1, CD180, DUSP10, SESN2, CHRNA3, STAT3, PLK3, EGR1, TRAF2, PLA2G1B, RHOB, ADRA2A, CEBPB, TICAM1, TGFB3, SELENOS, JAK3, TLR4, TNIP2, KLRK1, GPR37, DDIT4, SLC2A4, GOT1, SLC25A33, RIPK3, TLR2, INSIG1, IGFBP2, GPR155, TIRAP, LY96, VDR, JAK2, KLF9, CRY1, GDAP1, MGST1* |
| GO:0002683 | Negative regulation of immune system process | 116 | 1.946 | 1.23E-05 | 5.48E-05 | *CD274, CCL2, IRF1, ISG15, DHX58, SOCS1, BPIFB1, TNFAIP3, THBS1, PIM1, GDF15, STAT2, IL10, LTF, DUSP10, LRRC32, SERPING1, HSPA9, SOX11, BCL6, CEBPB, FBXO7, TGFB3, SELENOS, TMEM176B, JAK3, CGAS, KLRK1, ARG2, LYAR, RNF115, MYC, PSMB4* |
| GO:1901701 | Cellular response to oxygen-containing compound | 243 | 1.945 | 2.04E-07 | 1.45E-06 | *CXCL9, CXCL10, AMCF-II, CXCL13, IL1B, MMP8, CXCL8, IL1A, STAT1, SOCS1, TNFAIP3, IRF8, PIM1, FFAR2, NR4A3, PPBP, GDF15, ZFP36, STAT5A, NFKBIB, IL10, NFKBIA, LTF, GJA1, CD180, SESN2, CHRNA3, STAT3, TRAF2, PLA2G1B, RHOB, ADRA2A, TICAM1, SELENOS, JAK3, TLR4, TNIP2, KLRK1, GPR37, DDIT4, SLC2A4, GOT1, SLC25A33, RIPK3, TLR2, INSIG1, GPR155, LY96, VDR, JAK2, KLF9* |
| GO:0051050 | Positive regulation of transport | 149 | 1.935 | 6.18E-06 | 2.97E-05 | *CXCL9, CXCL11, CXCL10, IFNG, CCL2, IL1B, S100A8, IL6, IL1A, IL15, FFAR2, LIPG, NR4A3, CCL5, PDZK1, P2RY2, C3, NFKBIA, CCR1, PLK3, PLA2G1B, SOX11, ADRA2A, CEBPB, TGFB3, TLR4, CALR, LYAR, TLR2, BMP2, EIPR1, ZPR1, SFTPD, F2RL2* |
| GO:0030097 | Hemopoiesis | 236 | 1.923 | 3.54E-07 | 2.38E-06 | *IFNG, BATF2, IRF1, IL1B, ISG15, RSAD2, IL27, IL6, IL1A, TNFSF13B, STAT1, CSF3, SOCS1, SNX10, JUNB, IRF8, PIM1, SLA-DQB1, NFIL3, ZFP36, STAT5A, IL10, CEBPD, NFKBIA, LTF,* |

| | | | | | | |
|---|---|---|---|---|---|---|
| | | | | | | *LIF, CCR1, DUSP10, STAT3, CSF2, CASP3, CEBPG, EGR1, SLC1A5, TCIM, HSPA9, BCL6, CEBPB, TNFSF9, CASP8, FBXO7, TMEM176B, JAK3, GPATCH4, TLR4, KIT, LYAR, RIPK3, ETV6, MYC, ST3GAL1, BMP2* |
| GO:0008283 | Cell population proliferation | 444 | 1.921 | 9.74E-10 | 1.39E-08 | *CXCL9, CXCL11, CXCL10, IFNG, CD274, CCL2, IRF1, IL1B, IL27, IL6, IL1A, TNFSF13B, STAT1, CSF3, GNAT1, PTGIR, TNFAIP3, THBS1, JUNB, IL15, CTLA4, PIM1, TIMP1, CD40, NR4A3, IFI35, CCL5, ZFP36, STAT2, STAT5A, IL10, ATF5, NFKBIA, LTF, TFRC, GJA1, CTPS1, LIF, CD180, DUSP10, NME1, STAT3, LRRC32, CSF2, DRAM2, TRNP1, PHLDA2, ESM1, DDIT3, PLA2G1B, SOX11, BCL6, EIF2S2, SLC7A1, IFI30, IL18RAP, CEBPB, TNFSF9, TICAM1, TGFB3, B4GALT1, SDCBP2, TLR4, MORC3, CALR, ARG2, FAM98A, NOP2, SLC25A33, FBXO4, RIPK3, ETV6, MYC, RNASEH2B, IGFBP2, TIPIN, BMP2, PRDM1, GTPBP4, PTGES3, FAM83B, ZPR1, VDR, DLL4, ENPEP, JAK2, HILPDA, KLF9, ELF4, SMAD1, INHBA, LIG4, ADM, KAT2A, IGF2BP1, NKX3-1, SLC25A5, FUT2, PPARGC1A, WNT16* |
| GO:0002250 | Adaptive immune response | 118 | 1.919 | 2.60E-05 | 1.03E-04 | *IFNG, IL1B, RSAD2, IL27, IL6, TNFSF13B, IRF7, SLA-DOA, TNFAIP3, SLA-DQA1, SLA-DRA, CTLA4, CFB, SLA-DOB, CD40, SLA-DQB1, SLA-DRB1, C2, SLA-DMB, C3, TFRC, C4A, DUSP10, STAT3, SLA-DMA, TREM1, TRAF2, ULBP1, BCL6, MAD2L2, IL18RAP, JAK3, KLRK1* |
| GO:0016477 | Cell migration | 339 | 1.916 | 1.89E-08 | 1.81E-07 | *CXCL9, CXCL11, CXCL10, MADCAM1, IFNG, SELE, CCL2, AMCF-II, S100A12, CXCL13, CXCL8, S100A8, CCL4, RASGEF1A, CCL8, TNFAIP3, THBS1, FFAR2, CD40, NR4A3, PPBP, CCL5, ACKR4, GDF15, TNFSF14, GJA1, CCR1, DUSP10, CXCL16, TREM1, PHLDA2, CXCR6, DDIT3, GLUL, PLA2G1B, RHOB, ACKR2, ADRA2A, CDH12, NARS1, CASP8, TNFAIP1, B4GALT1, TLR4, KLRK1, CALR, KIT, SDC4, RIPK3, ITGA5, TWIST2, BMP2, GTPBP4, LOC100516039, DLL4, ENPEP, GNA13, LAMTOR2, RAB1A, ANGPT2, PLEKHO1* |
| GO:0006412 | Translation | 229 | 1.911 | 1.29E-06 | 7.31E-06 | *WARS1, THBS1, ZFP36, SESN2, GARS1, YARS1, EIF4EBP1, EIF2S2, FARSA, RGS2, NARS1, VARS1, CALR, RPL13A, EIF4A1, EIF2S1, EIF5A, ETF1, MRPL17, PUM3, RSL24D1, EIF3I, IGF2BP1, MRPL35, AGO2, MRPL15, C7H6orf125, EIF3J, NAT10, EIF1, RIDA, DDX3X,* |

| | | | | | | |
|---|---|---|---|---|---|---|
| | | | | | | *BZW1, RARS1, DRG1, NPM1, MRPS7, MRPL20, IGF2BP2, GFM2, MRPL2, MRPS11, ABCF1, RPS27L, FMR1, EIF3A, OGFOD1, EARS2, MRPL47, ATM, ACO1, FTSJ1, EIF4A3, DARS1, SRP9, MRPS18B, SERBP1, MRPS18C, SSB, MRPL32, PTCD3, QRSL1, GAPDH, MRPL44, RPL27, HARS1, DDRGK1, MRPL18, OGT, PAIP2B, RNF14, ELAVL1, SKIC8, FASTKD2, AIMP2, EIF3M, MTRF1, METTL3, MRPL24, LOC100738684, RPL26L1, MRPL58, DAP, DENR, MRPL1, MRPL13, CAPRIN1, RPL22L1, NEMF, MRPL11, STK35, RPS18, GUF1, MRPS5, MRPL33, PAIP1, UPF3B, GATC, RPS23, HARS2, MRPS18A, EIF2S3, MTOR, PAIP2, PARS2, IGFBP5, VARS2, RPL23, USP16, MRPS17, CDKAL1, LTN1, CHCHD1, FXR1, RPL38, MRRF, NCBP2, NCK1* |
| GO:0070887 | Cellular response to chemical stimulus | 463 | 1.901 | 1.73E-09 | 2.20E-08 | *CXCL9, CXCL11, CXCL10, CCL2, AMCF-II, CXCL13, IL1B, MMP8, CXCL8, S100A8, CCL4, IL1A, STAT1, SOCS1, CCL8, TNFAIP3, THBS1, AQP3, JUNB, IRF8, PIM1, FFAR2, NR4A3, MT-2B, PPBP, CCL5, ACKR4, GDF15, ZFP36, SLC39A8, STAT5A, NFKBIB, IL10, NFKBIA, LTF, TNFSF14, GJA1, CCR1, CD180, SESN2, ZFAND2A, CHRNA3, STAT3, ARID5A, CXCL16, TREM1, CXCR6, TRAF2, DDIT3, PLA2G1B, RHOB, ACKR2, TIGAR, ADRA2A, CEBPB, TICAM1, YBX3, SELENOS, JAK3, TLR4, TNIP2, CGAS, KLRK1, GPR37, CALR, DDIT4, KIT, SLC2A4, GOT1, SLC25A33, RIPK3, TLR2, TMEM38B, INSIG1, GPR155, EIF2S1, SLC41A1, PTGES3, LOC100516039, LY96, AQP5, VDR, JAK2, KLF9, LAMTOR2, CRY1, MGST1, ITPKC, NKX3-1, GSTM3, PPARGC1A, WNT16, PLSCR4, PLSCR3, IL16, HAVCR2, NCOA7, RALB, FBP1, PDCD4, GSTO1, CPNE8, SUV39H2, C2CD2L, PDE12, CYP26B1* |
| GO:0002757 | Immune response-activating signaling pathway | 116 | 1.895 | 3.12E-05 | 1.22E-04 | *IFNG, IRF1, RSAD2, IRF7, DHX58, FOSL1, BPIFB1, TNFAIP3, TIFA, SLC15A3, FFAR2, IFI35, TICAM2, NFKBIA, UNC93B1, TICAM1, MYD88, PLSCR1, TLR4, TNIP2, CGAS, KLRK1, RNF115, TMEM126A, TLR2, TIRAP, STING1, LY96, KCNK6* |
| GO:0043068 | Positive regulation of programmed cell death | 134 | 1.884 | 1.98E-05 | 8.09E-05 | *IFNG, CD274, CCL2, S100A8, STAT1, ATF3, THBS1, CD40, CCL5, IL10, FASLG, RNF122, GADD45G, CASP3, PHLDA2, TRAF2, DDIT3, PLA2G1B, RHOB, HSPA9, BCL6, TIGAR, CASP8, B4GALT1, MCL1, TLR4, TNFRSF12A, BCL2A1, RIPK3, BMP2, GADD45B, UTP11, VDR, TP53, JAK2, INHBA, NKX3-1, RASSF2, PDCD4, G0S2, SIAH1* |

| | | | | | | |
|---|---|---|---|---|---|---|
| GO:0040011 | Locomotion | 285 | 1.884 | 5.01E-07 | 3.24E-06 | *CXCL9, CXCL11, CXCL10, MADCAM1, IFNG, SELE, CCL2, AMCF-II, CXCL13, CXCL8, S100A8, CCL4, CCL8, THBS1, FFAR2, CD40, NR4A3, PPBP, CCL5, ACKR4, GDF15, BVES, TNFSF14, GJA1, CCR1, DUSP10, CXCL16, TREM1, PHLDA2, CXCR6, GLUL, PLA2G1B, RHOB, ACKR2, BCL6, ADRA2A, CASP8, FBXO7, TLR4, ARRDC3, KLRK1, GPR37, CALR, KIT, RIPK3, TWIST2, BMP2, GTPBP4, LOC100516039* |
| GO:0031399 | Regulation of protein modification process | 104 | 1.881 | 1.86E-04 | 6.26E-04 | *IFNG, CHI3L1, ISG15, TNFAIP3, IL15, STAT2, C3, LIF, EGR1, GNL3, PLA2G1B, TICAM1, PSMD10, FBXO4, ITGA5, UBE2D1, GTPBP4, JAK2, CRY1, NDFIP2, KAT2A, NCOA7, SPRY2, DDX3X, NRP1, UBE2B, DCUN1D5, PLAA, PHF23, FBXO33, SQSTM1, DR1, BCL10, PIK3R5, WBP1L, SEMA4D, OGT, DCUN1D3, PPIA* |
| GO:0042981 | Regulation of apoptotic process | 344 | 1.880 | 6.83E-08 | 5.34E-07 | *CXCL10, IFNG, CD274, CCL2, S100A8, STAT1, FAS, ATF3, LOC100522887, TNFAIP3, THBS1, PIM1, CD40, NOD2, CCL5, ZFP36, CASP10, STAT5A, ANGPTL4, IL10, ATF5, HSH2D, LTF, FASLG, RNF122, GADD45G, CASP3, PLK3, BCL2L14, BAK1, EGR1, PHLDA2, TRAF2, DDIT3, PLA2G1B, TCIM, RHOB, TRAF3, HSPA9, HIGD1A, BCL6, CASP1, TIGAR, CEBPB, RIPK2, CASP8, FBXO7, YBX3, SELENOS, B4GALT1, GHITM, JAK3, PSMD10, MCL1, TLR4, TNIP2, TNFRSF12A, BCL2A1, CFLAR, ARG2, RIPK3, MYC, ITGA5, ST3GAL1, BMP2, GADD45B, UTP11, ZPR1, HSP90AA1, VDR, TP53, JAK2, INHBA, LIG4, NKX3-1, SLC25A5, PPARGC1A, WNT16, PLSCR3* |
| GO:0009057 | Macromolecule catabolic process | 375 | 1.869 | 5.51E-08 | 4.46E-07 | *ISG20, IFNG, CHI3L1, ISG15, PSMB9, PSME2, PSMB8, RNF19B, PSMB10, TNFAIP3, TIMP1, PSME1, ZFP36, IL10, PSMA5, RNF122, ZFAND2A, HSP90B1, PSMA2, PLK3, ARID5A, PSMA3, TRAF2, DDIT3, DNAJB9, MAD2L2, ADRA2A, PSMB3, DERL3, CASP8, PSMA1, POP1, FBXO7, TNFAIP1, SELENOS, PSMD10, CALR, PGM2, FBXO4, RNF115, TMEM126A, PNPT1, PSMB4, RNASEH2B, UBE2D1, TIRAP, KLHDC3, PELO, PAQR3, YOD1, ZPR1, RBM8A, PSMA4, ETF1, PSMD3, REXO4, NDFIP2, UBE2D3, IGF2BP1, AGO2, DXO, DDX49, EXOSC3, PSMD8, PDE12, SIAH1, RIDA, TBRG4, RAD23A, RNF114, USP44, VCP, NPM1, PSMB1, CHMP5, IDE, DESI1, STT3B, UBE2B, PSMD1, HYAL2, OVGP1, PSMC1, UCHL5, PSME3, PLAA, TAF9, SIAH2, PSMC2, PHAX, FMR1, PSMB6, EXOSC4, SNF8, PTK2B,* |

| | | | | | | |
|---|---|---|---|---|---|---|
| | | | | | | *PHKG2, SKIC2, ATM, PSMF1, AKIRIN2, EIF4A3, PSMD2, PSMD4, SQSTM1, CNOT7, RLIM, RNF5, SSB, TRAF7, CHMP4A, ELOC, UBXN4, CNOT9, CUL5, DCPS, DDRGK1, OGT, EDEM2, ARK2N, RNF14, ELAVL1, NFE2L2, TMEM129, SKIC8, FASTKD2, PSMB7, FBXO9, PDCL3, N4BP1, MTREX, RAD23B, PSMC5, UBE3A, SOCS5, TOPORS, CACUL1, RIC1, UCHL3* |
| GO:0051603 | Proteolysis involved in protein catabolic process | 215 | 1.865 | 2.73E-06 | 1.41E-05 | *ISG15, PSMB9, PSME2, PSMB8, RNF19B, PSMB10, TNFAIP3, PSME1, PSMA5, RNF122, ZFAND2A, HSP90B1, PSMA2, PSMA3, DDIT3, DNAJB9, PSMB3, DERL3, CASP8, PSMA1, FBXO7, TNFAIP1, SELENOS, PSMD10, CALR, FBXO4, RNF115, TMEM126A, PSMB4, UBE2D1, KLHDC3, PAQR3, YOD1, PSMA4, PSMD3, NDFIP2, UBE2D3, PSMD8, SIAH1, RAD23A, RNF114, USP44, VCP, PSMB1, CHMP5, IDE, DESI1, STT3B, UBE2B, PSMC1, UCHL5, PSME3, PLAA, TAF9, SIAH2, PSMC2, PSMB6, SNF8, PTK2B, PSMF1, AKIRIN2, PSMD4, RLIM, RNF5, TRAF7, CHMP4A, ELOC, UBXN4, CUL5, DDRGK1, OGT, EDEM2, ARK2N, RNF14, NFE2L2, TMEM129, PSMB7, FBXO9, PDCL3, N4BP1, RAD23B, PSMC5, UBE3A, SOCS5, TOPORS, CACUL1, UCHL3, ARIH2* |
| GO:0030155 | Regulation of cell adhesion | 171 | 1.860 | 8.88E-06 | 4.05E-05 | *IFNG, SELE, RND1, CD274, IRF1, IL1B, CXCL8, IL1A, TNFSF13B, SOCS1, THBS1, IL15, NR4A3, SLA-DQB1, CCL5, STAT5A, IL10, SLA-DMB, TFRC, TNFSF14, LIF, TNC, DUSP10, LRRC32, BCL6, MAD2L2, SLC7A1, CEBPB, TNFSF9, JAK3, CALR, ARG2, IGFBP2, BMP2, GTPBP4, GPR4* |
| GO:0072594 | Establishment of protein localization to organelle | 116 | 1.856 | 6.17E-05 | 2.34E-04 | *IFNG, NEURL3, ISG15, TNFAIP3, NFKBIA, ZFAND2A, STAT3, TIMM8A, DDIT3, KPNA2, FBXO7, TXNIP, TOMM40, IPO4, LOC100157391, SRPRA, TMEM126A, SEC61A1, CHCHD4, SSR3, ZPR1, GDAP1, SPCS1* |
| GO:0051246 | Regulation of protein metabolic process | 316 | 1.852 | 5.70E-07 | 3.56E-06 | *IFNG, CHI3L1, ISG15, SERPINE1, PSME2, TNFAIP3, THBS1, IL15, TIMP1, LIPG, PSME1, ZFP36, STAT2, IL10, C3, NFKBIA, LIF, SESN2, ZFAND2A, CASP3, PLK3, EGR1, GNL3, EIF4EBP1, SOAT1, PLA2G1B, MAD2L2, ADRA2A, RGS2, CASP8, TICAM1, PSMD10, CALR, FBXO4, ITGA5, PFDN6, UBE2D1, EIF2S1, PAQR3, EIF5A, GTPBP4, ETF1, JAK2, PUM3, PSMD3, CRY1, NDFIP2, TMED10, INHBA, KAT2A, IGF2BP1, AGO2, C7H6orf125, NAT10,* |

| | | | | | | |
|---|---|---|---|---|---|---|
| | | | | | | *NCOA7, EIF1, PLGRKT, SPRY2, RIDA, DDX3X, BZW1, NPM1, PPP1R15B, NRP1, DESI1, UBE2B, PSMD1, DCUN1D5, IGF2BP2, PSME3, PLAA, TAF9, ABCF1, PHF23, FBXO33, FMR1, OGFOD1, SNF8, PTK2B, ACO1, EIF4A3, PSMD2, SQSTM1, SRP9, DR1, SERBP1, BCL10, SSB, TRAF7, PIK3R5, WBP1L, SEMA4D, PTCD3, GAPDH, DDRGK1, OGT, PAIP2B, DCUN1D3, HDAC2, APOD, PPIA* |
| GO:0043067 | Regulation of programmed cell death | 353 | 1.840 | 1.51E-07 | 1.12E-06 | *CXCL10, IFNG, CD274, CCL2, S100A8, STAT1, FAS, ATF3, LOC100522887, TNFAIP3, THBS1, PIM1, CD40, NOD2, CCL5, ZFP36, CASP10, STAT5A, ANGPTL4, IL10, ATF5, HSH2D, LTF, FASLG, RNF122, GADD45G, CASP3, FTH1, PLK3, BCL2L14, BAK1, EGR1, PHLDA2, TRAF2, DDIT3, PLA2G1B, TCIM, RHOB, TRAF3, HSPA9, HIGD1A, BCL6, CASP1, TIGAR, CEBPB, RIPK2, CASP8, FBXO7, YBX3, SELENOS, B4GALT1, GHITM, JAK3, PSMD10, MCL1, TLR4, TNIP2, TNFRSF12A, BCL2A1, CFLAR, ARG2, RIPK3, MYC, ITGA5, ST3GAL1, BMP2, GADD45B, UTP11, ZPR1, HSP90AA1, VDR, TP53, JAK2, INHBA, LIG4, NKX3-1, SLC25A5, PPARGC1A, WNT16, PLSCR3* |
| GO:0030099 | Myeloid cell differentiation | 128 | 1.831 | 1.60E-04 | 5.42E-04 | *IFNG, BATF2, ISG15, STAT1, CSF3, SOCS1, SNX10, JUNB, IRF8, ZFP36, STAT5A, NFKBIA, LTF, LIF, CCR1, STAT3, CSF2, CASP3, CEBPG, SLC1A5, HSPA9, BCL6, CEBPB, CASP8, TLR4, KIT, LYAR, MYC, BMP2* |
| GO:0002253 | Activation of immune response | 145 | 1.830 | 7.86E-05 | 2.94E-04 | *IFNG, IRF1, RSAD2, IRF7, DHX58, FOSL1, BPIFB1, TNFAIP3, TIFA, SLC15A3, CFB, FFAR2, IFI35, TICAM2, C2, C3, NFKBIA, C4A, UNC93B1, SERPING1, TICAM1, MYD88, PLSCR1, TLR4, TNIP2, CGAS, KLRK1, RNF115, TMEM126A, TLR2, TIRAP, CFI, STING1, HSP90AA1, LY96, CD46, KCNK6* |
| GO:0098609 | Cell-cell adhesion | 182 | 1.823 | 1.68E-05 | 7.19E-05 | *MADCAM1, IFNG, SELE, CD274, IRF1, IL1B, S100A8, IL1A, TNFSF13B, SOCS1, CLSTN3, VCAM1, IL15, NR4A3, SLA-DQB1, CCL5, SLC39A8, BVES, STAT5A, IL10, SLA-DMB, TFRC, TNFSF14, DUSP10, LRRC32, OLR1, BCL6, MAD2L2, ICAM1, CDH12, SLC7A1, CEBPB, TNFSF9, JAK3, ICAM4, ARG2, KIT, ITGA5, IGFBP2, BMP2* |
| GO:0006915 | Apoptotic process | 450 | 1.818 | 2.02E-08 | 1.89E-07 | *CXCL10, IFNG, CD274, CCL2, IRF1, CHI3L1, S100A8, IL1A, STAT1, FAS, ATF3, LOC100522887, TNFAIP3, THBS1, PIM1, CD40, NOD2, CCL5, ZFP36, CASP10, STAT5A,* |

| | | | | | | |
|---|---|---|---|---|---|---|
| | | | | | | *ANGPTL4, IL10, ATF5, HSH2D, LTF, SHISA5, AEN, FASLG, RNF122, GADD45G, CYCS, CASP3, CSRNP1, PLK3, BCL2L14, BAK1, EGR1, PHLDA2, LOC100624226, TRAF2, DDIT3, PLA2G1B, TCIM, RHOB, TRAF3, HSPA9, NEK6, HIGD1A, BCL6, CASP1, TIGAR, CEBPB, RIPK2, CASP8, TICAM1, FBXO7, YBX3, SELENOS, B4GALT1, GHITM, JAK3, PSMD10, MCL1, PLSCR1, TLR4, TNIP2, TNFRSF12A, BCL2A1, CFLAR, TNFRSF1A, ARG2, DDIT4, RIPK3, ETV6, MYC, ITGA5, ST3GAL1, BMP2, GADD45B, UTP11, ZPR1, HSP90AA1, VDR, TP53, JAK2* |
| GO:0008219 | Cell death | 463 | 1.816 | 4.57E-08 | 3.88E-07 | *CXCL10, IFNG, CD274, CCL2, IRF1, CHI3L1, S100A8, IL1A, STAT1, FAS, GSDMA, ATF3, LOC100522887, TNFAIP3, THBS1, PIM1, CD40, NOD2, CCL5, ZFP36, CASP10, STAT5A, ANGPTL4, IL10, ATF5, HSH2D, LTF, SHISA5, AEN, FASLG, RNF122, GADD45G, CYCS, CASP3, FTH1, CSRNP1, MLKL, PLK3, BCL2L14, BAK1, EGR1, PHLDA2, LOC100624226, TRAF2, DDIT3, PLA2G1B, TCIM, RHOB, TRAF3, HSPA9, NEK6, HIGD1A, BCL6, CASP1, TIGAR, CEBPB, RIPK2, CASP8, TICAM1, FBXO7, YBX3, SELENOS, B4GALT1, GHITM, JAK3, PSMD10, MCL1, PLSCR1, TLR4, TNIP2, TNFRSF12A, BCL2A1, CFLAR, TNFRSF1A, ARG2, DDIT4, RIPK3, ETV6, MYC, ITGA5, ST3GAL1, BMP2, GADD45B, UTP11, ZPR1, HSP90AA1, VDR, TP53, JAK2* |
| GO:0012501 | Programmed cell death | 463 | 1.816 | 4.57E-08 | 3.88E-07 | *CXCL10, IFNG, CD274, CCL2, IRF1, CHI3L1, S100A8, IL1A, STAT1, FAS, GSDMA, ATF3, LOC100522887, TNFAIP3, THBS1, PIM1, CD40, NOD2, CCL5, ZFP36, CASP10, STAT5A, ANGPTL4, IL10, ATF5, HSH2D, LTF, SHISA5, AEN, FASLG, RNF122, GADD45G, CYCS, CASP3, FTH1, CSRNP1, MLKL, PLK3, BCL2L14, BAK1, EGR1, PHLDA2, LOC100624226, TRAF2, DDIT3, PLA2G1B, TCIM, RHOB, TRAF3, HSPA9, NEK6, HIGD1A, BCL6, CASP1, TIGAR, CEBPB, RIPK2, CASP8, TICAM1, FBXO7, YBX3, SELENOS, B4GALT1, GHITM, JAK3, PSMD10, MCL1, PLSCR1, TLR4, TNIP2, TNFRSF12A, BCL2A1, CFLAR, TNFRSF1A, ARG2, DDIT4, RIPK3, ETV6, MYC, ITGA5, ST3GAL1, BMP2, GADD45B, UTP11, ZPR1, HSP90AA1, VDR, TP53, JAK2* |

| | | | | | | |
|---|---|---|---|---|---|---|
| GO:0071705 | Nitrogen compound transport | 495 | 1.795 | 4.80E-08 | 3.97E-07 | *IFNG, NEURL3, TAP1, S100A8, RSAD2, IL6, IL1A, CALHM6, GNAT1, SNX10, AQP3, RHBDF2, STX11, SLC15A3, EGR2, FFAR2, CCL5, ZFP36, PDZK1, SLC38A5, CD74, NFKBIA, TFRC, GJA1, SLC25A22, STAT3, TIMM8A, ARL5B, PLK3, SLC1A5, RAB3D, RHOB, HSPA9, ADRA2A, SLC1A3, GPM6B, SEC23B, SLC7A1, DERL3, KPNA2, OPTN, TXNIP, TOMM40, SCFD2, TGFB3, SELENOS, IPO4, TLR4, PREB, ARRDC3, LOC100157391, SRPRA, CALR, SEC24A, ARF4, SLC25A33, TLR2, RAB39B, INSIG1, SEC61A1, SEC61B, CHCHD4, TMED5, STXBP1, SYS1, USO1, SEC22B, EIF4E, AP3S2, ZPR1, RBM8A, TP53, F2RL2, KCNK1, RAB1A, NDFIP2, NOL6, ARCN1, MGST1, TMED2, TMED10, DNAJC15, IGF2BP1, SLC25A5, COPE, C7H6orf125, CADPS2, KCNA2, C2CD2L, TRAM1, SLC38A2, TMED9, TVP23B, GOSR2, XPO6, SLC33A1, IFT20, LRRC8A, COPB1, VCP, KPNB1, ARF6, COPB2, SLC7A2, MAGOHB, DESI1, SDAD1, YIF1A, SAR1B, GDI2, NXT1, HYAL2, SLC18B1, HTR1B, TIMM13, IGF2BP2, SRP54, COPZ1, SEC63, XPO5, PHAX, FMR1, SLC18A2, SNX19, MACIR, MTX1, PDCD10, SNF8, SLC22A4, SRI, TIMM44, RAB9A, RAB20, AKIRIN2, RAB13, EIF4A3, SQSTM1, RAB2A, SNX5, TIMM8B, NUTF2, TOMM20, FYTTD1, TMED1, SSB, STX17, NUP188, ENY2, ARF1, TNPO1, TIMM21, MRPL18, SFXN1, VTI1B, SEC24C, YIPF5, CLTC, NSF, APOD, ARF2, MTX2, AP3M1, ELAVL1, PHB2, RAE1, BBS5, MAGOH, TMEM129, MPC2, VPS26A, M6PR, NHERF1* |
| GO:0048870 | Cell motility | 387 | 1.789 | 4.72E-07 | 3.11E-06 | *CXCL9, CXCL11, CXCL10, MADCAM1, IFNG, SELE, CCL2, AMCF-II, S100A12, CXCL13, CXCL8, S100A8, CCL4, RASGEF1A, CCL8, TNFAIP3, THBS1, FFAR2, CD40, NR4A3, PPBP, CCL5, ACKR4, GDF15, TNFSF14, GJA1, CCR1, DUSP10, CXCL16, TREM1, PHLDA2, CXCR6, DDIT3, GLUL, PLA2G1B, RHOB, ACKR2, BCL6, ADRA2A, CDH12, NARS1, CASP8, TNFAIP1, B4GALT1, TLR4, KLRK1, CALR, KIT, SDC4, RIPK3, ITGA5, TWIST2, BMP2, GTPBP4, LOC100516039, DLL4, ENPEP, CCNYL1, GNA13, LAMTOR2, RAB1A, ANGPT2, PLEKHO1* |
| GO:0030163 | Protein catabolic process | 266 | 1.783 | 1.05E-05 | 4.73E-05 | *IFNG, ISG15, PSMB9, PSME2, PSMB8, RNF19B, PSMB10, TNFAIP3, TIMP1, PSME1, IL10, PSMA5, RNF122, ZFAND2A, HSP90B1, PSMA2, PLK3, PSMA3, TRAF2, DDIT3, DNAJB9, MAD2L2, ADRA2A, PSMB3, DERL3, CASP8, PSMA1, FBXO7, TNFAIP1, SELENOS, PSMD10,* |

| | | | | | | |
|---|---|---|---|---|---|---|
| | | | | | | *CALR, FBXO4, RNF115, TMEM126A, PSMB4, UBE2D1, KLHDC3, PAQR3, YOD1, PSMA4, PSMD3, NDFIP2, UBE2D3, PSMD8, SIAH1, RAD23A, RNF114, USP44, VCP, PSMB1, CHMP5, IDE, DESI1, STT3B, UBE2B, PSMD1, PSMC1, UCHL5, PSME3, PLAA, TAF9, SIAH2, PSMC2, PSMB6, SNF8, PTK2B, PSMF1, AKIRIN2, PSMD2, PSMD4, SQSTM1, RLIM, RNF5, TRAF7, CHMP4A, ELOC, UBXN4, CUL5, DDRGK1, OGT, EDEM2, ARK2N, RNF14, NFE2L2, TMEM129, PSMB7, FBXO9, PDCL3, N4BP1, RAD23B, PSMC5, UBE3A, SOCS5, TOPORS, CACUL1, RIC1, UCHL3, ARIH2, STX5, ADAM10* |
| GO:0009967 | Positive regulation of signal transduction | 357 | 1.780 | 7.44E-07 | 4.57E-06 | *SAA3, S100A12, CHI3L1, MMP8, S100A8, RSAD2, RASGEF1A, IL6, IRF7, DHX58, ZBP1, ATF3, TNFAIP3, TIFA, THBS1, SLC15A3, PIM1, CD40, IFI35, TICAM2, CCL5, GDF15, CD74, IL10, C3, TNFSF14, LIF, CCR1, CD180, SESN2, FASLG, GADD45G, STAT3, CSF2, ESM1, TRAF2, DDIT3, PLA2G1B, TCIM, DDX21, SOX11, ADRA2A, CASP8, TICAM1, MYD88, LPAR3, MCL1, TLR4, TNIP2, ADAM17, ARRDC3, CFLAR, GPR37, CALR, KIT, TMEM106A, RIPK3, MRAP2, TLR2, GPR155, NLE1, TIRAP, BMP2, GADD45B, FAM83B, GPR4, LY96, VDR, DLL4, TP53, JAK2, LAMTOR2, IKBKB, KCNK6, INHBA, NKX3-1, WNT16, RIPK1, HAVCR2, TERF2IP, RALB, LTBR, RASSF2, PDCD4, G0S2, SPRY2, SIAH1* |
| GO:0023056 | Positive regulation of signaling | 394 | 1.771 | 1.28E-06 | 7.31E-06 | *SAA3, CCL2, S100A12, IL1B, CHI3L1, MMP8, S100A8, RSAD2, RASGEF1A, IL6, IRF7, DHX58, ZBP1, ATF3, TNFAIP3, TIFA, THBS1, SLC15A3, PIM1, FFAR2, CD40, IFI35, TICAM2, CCL5, GDF15, BVES, CD74, IL10, C3, TNFSF14, LIF, CCR1, CD180, SESN2, FASLG, GADD45G, STAT3, CSF2, LRRTM2, ESM1, TRAF2, DDIT3, PLA2G1B, TCIM, DDX21, SOX11, ADRA2A, CASP8, TICAM1, MYD88, LPAR3, MCL1, TLR4, TNIP2, ADAM17, ARRDC3, CFLAR, GPR37, CALR, KIT, TMEM106A, RIPK3, MRAP2, TLR2, GPR155, NLE1, TIRAP, BMP2, GADD45B, FAM83B, GPR4, LY96, VDR, DLL4, TP53, JAK2, F2RL2, LAMTOR2, IKBKB, KCNK6, INHBA, ADORA1, NKX3-1, WNT16, RIPK1, HAVCR2, TERF2IP, RALB, LTBR, RASSF2, PDCD4, G0S2, SPRY2, C2CD2L, SIAH1* |
| GO:0055085 | Transmembrane transport | 259 | 1.766 | 2.59E-05 | 1.03E-04 | *CXCL9, CXCL11, CXCL10, SLC25A47, CALHM6, THBS1, AQP3, CNGA4, SLC15A3, NR4A3, SLC25A25, SLC39A8, KCNJ2, PDZK1, SLC38A5, C3, GJA1, SLC25A22, GLRB, SESN2,* |

| | | | | | | |
|---|---|---|---|---|---|---|
| | | | | | | *KCNMA1, PANX2, SLC1A5, BPIFA1, DDIT3, AQP7, SLC25A28, PLA2G1B, ADRA2A, SLC1A3, SLC7A1, TOMM40, GHITM, SLC5A6, LOC100157391, SLC2A4, SLC31A1, SLC25A33, KCNJ14, TMEM38B, GPR155, SEC61A1, CHCHD4, STING1* |
| GO:1902533 | Positive regulation of intracellular signal transduction | 271 | 1.765 | 1.83E-05 | 7.68E-05 | *SAA3, S100A12, CHI3L1, S100A8, RSAD2, RASGEF1A, DHX58, TIFA, THBS1, SLC15A3, PIM1, CD40, IFI35, TICAM2, CCL5, GDF15, CD74, TNFSF14, LIF, CCR1, SESN2, FASLG, GADD45G, CSF2, TRAF2, DDIT3, PLA2G1B, TCIM, DDX21, SOX11, ADRA2A, CASP8, TICAM1, MYD88, LPAR3, MCL1, TLR4, TNIP2, ADAM17, ARRDC3, CFLAR, GPR37, CALR, TMEM106A, RIPK3, TLR2, GPR155, TIRAP, BMP2, GADD45B, GPR4, VDR, TP53, JAK2, LAMTOR2, IKBKB, KCNK6, INHBA, NKX3-1, WNT16, RIPK1, HAVCR2, TERF2IP, LTBR, RASSF2, PDCD4, SPRY2, SIAH1, ACKR3, DDX3X* |
| GO:0051240 | Positive regulation of multicellular organismal process | 368 | 1.759 | 1.02E-06 | 6.16E-06 | *IFNG, IRF1, IL1B, CHI3L1, MMP8, ISG15, CXCL8, RSAD2, IL6, IL1A, TNFSF13B, IRF7, DHX58, STAT1, SOCS1, THBS1, IL15, IRF8, EGR2, PIM1, FFAR2, LIPG, CD40, SLA-DQB1, TICAM2, CCL5, STAT5A, P2RY2, CD74, IL10, SLA-DMB, TSKU, LTF, TFRC, TNFSF14, GJA1, LIF, CCR1, DUSP10, LRRTM2, CEBPG, PANX2, EGR1, TRAF2, DDIT3, PLA2G1B, RHOB, DDX21, SOX11, BCL6, CASP1, MAD2L2, ADRA2A, SLC7A1, RGS2, POLR3D, CEBPB, TNFSF9, CASP8, TICAM1, MYD88, LPAR3, YBX3, TGFB3, B4GALT5, TLR4, HRH2, CGAS, KLRK1, KIT, TMEM106A, ITGA5, TLR2, IGFBP2, TIRAP, BMP2, STING1, HSP90AA1, LY96, VDR, JAK2, HILPDA, RAB1A, SMAD1, ANGPT2, TMED10, INHBA, LIG4, ADM, KAT2A, NKX3-1, PPARGC1A* |
| GO:0010647 | Positive regulation of cell communication | 392 | 1.753 | 1.22E-06 | 7.17E-06 | *SAA3, CCL2, S100A12, IL1B, CHI3L1, MMP8, S100A8, RSAD2, RASGEF1A, IL6, IRF7, DHX58, ZBP1, ATF3, TNFAIP3, TIFA, THBS1, SLC15A3, PIM1, FFAR2, CD40, IFI35, TICAM2, CCL5, GDF15, CD74, IL10, C3, TNFSF14, LIF, CCR1, CD180, SESN2, FASLG, GADD45G, STAT3, CSF2, LRRTM2, ESM1, TRAF2, DDIT3, PLA2G1B, TCIM, DDX21, SOX11, ADRA2A, CASP8, TICAM1, MYD88, LPAR3, MCL1, TLR4, TNIP2, ADAM17, ARRDC3, CFLAR, GPR37, CALR, KIT, TMEM106A, RIPK3, MRAP2, TLR2, GPR155, NLE1, TIRAP, BMP2, GADD45B, FAM83B, GPR4, LY96, VDR, DLL4, TP53, JAK2, F2RL2, LAMTOR2,* |

| | | | | | | |
|---|---|---|---|---|---|---|
| | | | | | | *IKBKB, KCNK6, INHBA, ADORA1, NKX3-1, WNT16, RIPK1, HAVCR2, TERF2IP, RALB, LTBR, RASSF2, PDCD4, G0S2, SPRY2, C2CD2L, SIAH1* |
| GO:0019941 | Modification-dependent protein catabolic process | 173 | 1.742 | 1.53E-04 | 5.25E-04 | *ISG15, PSMB9, RNF19B, TNFAIP3, PSMA5, RNF122, ZFAND2A, PSMA2, PSMA3, DDIT3, PSMB3, PSMA1, FBXO7, TNFAIP1, PSMD10, FBXO4, RNF115, UBE2D1, KLHDC3, PAQR3, PSMA4, PSMD3, NDFIP2, UBE2D3, PSMD8, SIAH1, RAD23A, RNF114, USP44, VCP, CHMP5, DESI1, UBE2B, PSMC1, UCHL5, PLAA, TAF9, SIAH2, PSMC2, SNF8, PTK2B, PSMF1, AKIRIN2, PSMD4, RLIM, RNF5, TRAF7, CHMP4A, ELOC, CUL5, DDRGK1, OGT, ARK2N, RNF14, NFE2L2, TMEM129, PSMB7, FBXO9, PDCL3, N4BP1, RAD23B, PSMC5, UBE3A, SOCS5, TOPORS, CACUL1, UCHL3, ARIH2* |
| GO:0043632 | Modification-dependent macromolecule catabolic process | 174 | 1.739 | 1.21E-04 | 4.23E-04 | *ISG15, PSMB9, RNF19B, TNFAIP3, PSMA5, RNF122, ZFAND2A, PSMA2, PSMA3, DDIT3, PSMB3, PSMA1, FBXO7, TNFAIP1, PSMD10, FBXO4, RNF115, UBE2D1, KLHDC3, PAQR3, PSMA4, PSMD3, NDFIP2, UBE2D3, PSMD8, SIAH1, RAD23A, RNF114, USP44, VCP, CHMP5, DESI1, UBE2B, PSMC1, UCHL5, PLAA, TAF9, SIAH2, PSMC2, SNF8, PTK2B, PSMF1, AKIRIN2, PSMD4, RLIM, RNF5, TRAF7, CHMP4A, ELOC, CUL5, DDRGK1, OGT, ARK2N, RNF14, NFE2L2, TMEM129, PSMB7, FBXO9, PDCL3, N4BP1, RAD23B, PSMC5, UBE3A, SOCS5, TOPORS, CACUL1, UCHL3, ARIH2* |
| GO:0006886 | Intracellular protein transport | 189 | 1.736 | 9.58E-05 | 3.41E-04 | *IFNG, SNX10, STX11, EGR2, CD74, NFKBIA, STAT3, TIMM8A, ARL5B, PLK3, RAB3D, RHOB, HSPA9, SEC23B, DERL3, KPNA2, TXNIP, SELENOS, IPO4, SRPRA, CALR, SEC24A, INSIG1, SEC61A1, SEC61B, USO1, AP3S2, ZPR1, RAB1A, TMED10, TRAM1, TMED9, XPO6, COPB1, VCP, KPNB1, ARF6, COPB2, DESI1, SAR1B, NXT1, HYAL2, SRP54, COPZ1, SEC63, XPO5, PDCD10, SNF8, RAB20, AKIRIN2, SQSTM1, SNX5, TOMM20, TMED1, STX17, TNPO1, VTI1B, SEC24C, CLTC, APOD, AP3M1, ELAVL1, PHB2, TMEM129, VPS26A, M6PR, AKAP5, RIC1, STX12, HIKESHI, TNPO3, SEC23A, STX5, IPO13, MED1, RAB5A, COPA, AHCYL1, SNX9, RANGRF, MTCH1, KPNA1* |

| | | | | | | |
|---|---|---|---|---|---|---|
| GO:0007186 | G protein-coupled receptor signaling pathway | 166 | 1.722 | 3.12E-04 | 1.03E-03 | *CXCL9, CXCL11, CXCL10, ACKR1, CCL2, AMCF-II, CXCL13, CXCL8, CCL4, GNAT1, CCL8, PTGIR, RGS16, FFAR2, PPBP, CCL5, P2RY2, C3, PROKR2, GLRB* |
| GO:0016071 | MRNA metabolic process | 244 | 1.707 | 6.16E-05 | 2.34E-04 | *RNASEL, ZFP36, DNAJB11, ARID5A, CEBPG, TRAF2, NHP2, SNRPF, SNRPD1, PAPOLA, PNPT1, TIRAP, PUS3, PELO, RBM8A, SNRPA1, VDR, ETF1, SMNDC1, IGF2BP1, AGO2, DXO, SNRPG, EXOSC3, PDE12, SNRNP27, RIDA, SNRPD3, TBRG4, TRA2B, GEMIN6, NPM1, PUS1, SMN1, SNRPA, RPRD1B, ZMAT2, STRAP, WTAP, SF3B6, TAF9, PRPF40A, CLNS1A, FMR1, SNRPC, SNRPB2, SMU1, USP39, EXOSC4, ATM, HNRNPF, CSTF2, SRSF1, EIF4A3, CNOT7, RBM7, SSB, SNRPE, SNRNP40, SRSF7, HIPK3, LSM6, CNOT9, RBM14, DCPS, PRPF19, RNPS1, ELAVL1, RBM47, CSTF3, SKIC8, FASTKD2, METTL3, NSRP1, SNU13, RBMX2, RBM25, WDR83, MED1, ZFP36L2, CAPRIN1, RBM42, TENT2, CDK12, LSM5, SCAF11, CSTF2T, LEO1, AHCYL1, GEMIN8, U2AF1L4, ZBTB1, CWC22, RBM39, THOC7, PCID2, PRPF3, RBBP6, PAIP1, UPF3B, SYMPK, UBL5, ROCK1, ZRANB2, PPIL1, SRSF2, LSM8, C1H9orf78, HNRNPK, PTBP1, BUD31, PAN2, GEMIN2, SDE2, PCF11, SMG9, ZFP36L1, PTBP3, TAF7, RBM15, NCBP2, PABPC1, U2AF2, SSU72, RBM41, SF3A3, ZC3H14, LSM2, RBMX, TAF11, CLP1, SON, CSDE1* |
| GO:0043412 | Macromolecule modification | 499 | 1.699 | 1.23E-06 | 7.17E-06 | *IFNG, CHI3L1, ISG15, UBA7, USP18, SOCS3, SOCS1, RNF19B, TNFAIP3, IL15, PARP12, EGR2, TRIM21, STAT2, SLC39A8, GALNT15, C3, CISH, LIF, DNAJB11, RNF122, QTRT1, PARP11, EGR1, GNL3, EMG1, TRAF2, GLUL, ST6GALNAC1, PLA2G1B, NEK6, WDR4, NHP2, DERL3, TICAM1, FBXO7, GAR1, TNFAIP1, B4GALT1, FBL, PSMD10, B4GALT5, QTRT2, WSB2, USP15, NOP2, FBXO4, RNF115, MOGS, OSTC, ITGA5, UBE2D1, ST3GAL1, DKC1, PUS3, EIPR1, GTPBP4, B3GNT5, POMGNT2, ALG3, RPN1, OTUB2, JAK2, TRMT6, RPN2, CRY1, KLHL25, BUD23, NDFIP2, USP2, UBE2D3, RXYLT1, KAT2A, FUT2, MPDU1, NAT10, NCOA7, NUS1, EXOSC3, SOCS7, SPRY2, SIAH1, DAD1, MGAT2, PARP3, RNF138, NSUN2, DDX3X, TMEM258, B3GALT6, GALNT1, RNF114, USP44, VCP, PUS1, PUS7L, HSD17B10, NRP1, DESI1, PPP2R2A, MGAT1, A4GALT, THUMPD1, STT3B, RAMAC, UBE2B,* |

| | | | | | | |
|---|---|---|---|---|---|---|
| | | | | | | *NPC1, WTAP, ST8SIA2, DCUN1D5, B3GALNT1, DPAGT1, UCHL5, HINT1, PLAA, SIAH2, PHF23, FBXO33, USP39, EXOSC4, PPP2CA, TYW5, ATM, ELP5, C1GALT1, PARP8, FTSJ1, B4GALT3, UBA3, PPP1CB, SQSTM1, RLIM, DR1, RNF5, OTUB1, OTUD6B, ST6GAL2, MTO1, BCL10, TRAF7, PIK3R5, DPM1, WBP1L, TPST1, FUT8, SEMA4D, COPS6, GAPDH, CUL5, MAGT1, DDRGK1, OGT, PADI4, DCUN1D3, EDEM2, RNF14, TRMT112, PPIA* |
| GO:0051248 | Negative regulation of protein metabolic process | 103 | 1.691 | 2.32E-03 | 5.94E-03 | *ISG15, SERPINE1, TNFAIP3, THBS1, TIMP1, ZFP36, IL10, SESN2, EIF4EBP1, MAD2L2, CALR, PFDN6, EIF2S1, GTPBP4, CRY1, IGF2BP1, AGO2, NCOA7, SPRY2, RIDA, DDX3X, UBE2B, PLAA, TAF9, EIF4A3, SQSTM1, SRP9, SERBP1, GAPDH, DDRGK1, OGT, PAIP2B, DCUN1D3, APOD, PPIA* |
| GO:0097190 | Apoptotic signaling pathway | 153 | 1.677 | 5.68E-04 | 1.79E-03 | *IFNG, S100A8, IL1A, ATF3, TNFAIP3, THBS1, IL10, AEN, FASLG, TRAF2, DDIT3, CEBPB, CASP8, TICAM1, FBXO7, YBX3, SELENOS, GHITM, PSMD10, MCL1, TLR4, BCL2A1, CFLAR, DDIT4, RIPK3, MYC, TP53, JAK2, INHBA, NKX3-1, SLC25A5, WNT16, PLSCR3, G0S2, SIAH1, ACKR3, DDX3X, NRP1, SLC35F6, HYAL2, MLLT11, HINT1, SNAI2, PSME3, SIAH2, RPS27L* |
| GO:0034655 | Nucleobase-containing compound catabolic process | 148 | 1.676 | 7.07E-04 | 2.11E-03 | *ISG20, SAMHD1, HK3, ZFP36, ARID5A, GCK, TRAF2, TIGAR, POP1, CNP, PGK1, DDIT4, PNPT1, RNASEH2B, TIRAP, PELO, ZPR1, RBM8A, HK2, PDE4B, ETF1, REXO4, DCTPP1, IGF2BP1, LOC100515984, AGO2, DXO, DDX49, FBP1, EXOSC3, PDE12, NT5C, FITM2, RIDA, TBRG4, ENTPD1, NPM1, ITPA, PDE4D, HINT1, PHAX, FMR1, PDE4A, EXOSC4, PPP2CA, TPI1, SKIC2, ATM, NUDT11, DPYS, EIF4A3, CNOT7, SSB, CNOT9, GAPDH, DCPS, OGT, ELAVL1, SKIC8, FASTKD2* |
| GO:1902531 | Regulation of intracellular signal transduction | 476 | 1.675 | 6.54E-06 | 3.11E-05 | *SAA3, S100A12, CHI3L1, S100A8, RSAD2, RASGEF1A, DHX58, STAT1, FAS, ATF3, DUSP5, TNFAIP3, TIFA, THBS1, SLC15A3, PIM1, CD40, IFI35, TICAM2, CCL5, GDF15, SLC39A8, NFKBIB, CD74, IL10, NFKBIA, TNFSF14, LIF, CCR1, DUSP10, SESN2, FASLG, GADD45G, CSF2, CHN2, TRAF2, DDIT3, PLA2G1B, TCIM, TRAF3, DDX21, SOX11, BCL6, ADRA2A, TNIP1, OPTN, CASP8, TICAM1, MYD88, FBXO7, LPAR3, YBX3, TNFAIP1, SELENOS, PSMD10, MCL1, TLR4, TNIP2, ADAM17, ARRDC3, CGAS, CFLAR, GPR37, CALR, DDIT4,* |

| | | | | | | |
|---|---|---|---|---|---|---|
| | | | | | | *RRS1, NOP2, FICD, TMEM106A, FBXO4, RIPK3, MYC, TLR2, RNASEH2B, INSIG1, UBE2D1, GPR155, TIRAP, BMP2, GADD45B, PAQR3, GPR4, VDR, TP53, JAK2, LAMTOR2, IKBKB, CDC42SE1, KCNK6, CRY1* |
| GO:0006511 | Ubiquitin-dependent protein catabolic process | 170 | 1.673 | 9.74E-04 | 2.80E-03 | *PSMB9, RNF19B, TNFAIP3, PSMA5, RNF122, ZFAND2A, PSMA2, PSMA3, DDIT3, PSMB3, PSMA1, FBXO7, TNFAIP1, PSMD10, FBXO4, RNF115, UBE2D1, KLHDC3, PAQR3, PSMA4, PSMD3, NDFIP2, UBE2D3, PSMD8, SIAH1, RAD23A, RNF114, USP44, VCP, CHMP5, DESI1, UBE2B, PSMC1, UCHL5, PLAA, TAF9, SIAH2, PSMC2, SNF8, PTK2B, PSMF1, AKIRIN2, PSMD4, RLIM, RNF5, TRAF7, CHMP4A, ELOC, CUL5, DDRGK1, OGT, ARK2N, RNF14, NFE2L2, TMEM129, PSMB7, FBXO9, PDCL3, N4BP1, RAD23B, PSMC5, UBE3A, SOCS5, TOPORS, CACUL1, UCHL3, ARIH2* |
| GO:0048878 | Chemical homeostasis | 242 | 1.673 | 7.96E-05 | 2.95E-04 | *CXCL9, CXCL11, CXCL10, IL6, IL1A, CCL8, LPL, SNX10, AQP3, HK3, CP, FFAR2, LIPG, CD40, MT-2B, CCL5, SLC39A8, ANGPTL4, STC2, TSKU, TFRC, CCR1, SESN2, STAT3, FTH1, SOAT1, BPIFA1, GCK, DDIT3, AQP7, ADRA2A, GHITM, ORMDL1, SLC31A1, GOT1, ABHD5, MYC, TMEM38B, INSIG1, SLC41A1, SFTPD, VDR, HK2* |
| GO:0033365 | Protein localization to organelle | 237 | 1.665 | 1.52E-04 | 5.25E-04 | *IFNG, NEURL3, ISG15, TNFAIP3, SNX10, NFKBIA, SESN2, ZFAND2A, STAT3, TIMM8A, ARL5B, GNL3, DDIT3, KPNA2, OPTN, FBXO7, TXNIP, TOMM40, IPO4, LOC100157391, MORC3, SRPRA, CALR, RRS1, FBXO4, TMEM126A, INSIG1, SEC61A1, ABHD17B, CHCHD4, PAQR3, SSR3, ZPR1, GDAP1, SPCS1, TMED10, SRP14, TERF2IP, TRAM1, PARP3, DDX3X, IFT20, EMC3, VCP, KPNB1, ARL1, ARF6, NRP1, GDI2, HYAL2, SRP54, SEC63, TBRG1, BCS1L, SNF8, TIMM44, BRD2, AKIRIN2, ARL2BP, SQSTM1, SRP9, ORMDL3, SSB, CHMP4A, SRP68, CHCHD10, TNPO1, TIMM21, DDRGK1, OGT, APOD, ELAVL1, PHB2, M6PR, MFSD1, EMC4, MMGT1, SRP19, TOPORS, TIMM17B, HIKESHI, TNPO3, ADAM10, IPO13* |
| GO:0006873 | Intracellular monoatomic ion homeostasis | 134 | 1.663 | 7.99E-04 | 2.35E-03 | *CXCL9, CXCL11, CXCL10, IL1A, CCL8, CP, CD40, MT-2B, CCL5, SLC39A8, STC2, TFRC, CCR1, FTH1, DDIT3, GHITM, SLC31A1, MYC, TMEM38B, SLC41A1* |

| | | | | | | |
|---|---|---|---|---|---|---|
| GO:0051247 | Positive regulation of protein metabolic process | 172 | 1.660 | 7.25E-04 | 2.15E-03 | *IFNG, CHI3L1, TNFAIP3, THBS1, IL15, C3, NFKBIA, LIF, ZFAND2A, CASP3, PLK3, EGR1, GNL3, SOAT1, PLA2G1B, ADRA2A, CASP8, TICAM1, PSMD10, FBXO4, ITGA5, UBE2D1, PAQR3, EIF5A, JAK2, CRY1, NDFIP2, INHBA, IGF2BP1, C7H6orf125, NAT10, PLGRKT, DDX3X, NPM1, NRP1, DCUN1D5, ABCF1, PHF23, FBXO33, FMR1, SNF8, PTK2B, EIF4A3, BCL10, SSB, TRAF7, PIK3R5, WBP1L, SEMA4D, DDRGK1, OGT, DCUN1D3, HDAC2, PPIA, ELAVL1* |
| GO:0030003 | Intracellular monoatomic cation homeostasis | 133 | 1.660 | 1.04E-03 | 2.95E-03 | *CXCL9, CXCL11, CXCL10, IL1A, CCL8, CP, CD40, MT-2B, CCL5, SLC39A8, STC2, TFRC, CCR1, FTH1, DDIT3, GHITM, SLC31A1, MYC, TMEM38B, SLC41A1* |
| GO:0042592 | Homeostatic process | 401 | 1.658 | 1.37E-05 | 5.95E-05 | *CXCL9, CXCL11, CXCL10, IL1B, ISG15, IL6, IL1A, TNFSF13B, STAT1, FAS, CCL8, TNFAIP3, LPL, SNX10, AQP3, HK3, CP, FFAR2, LIPG, CD40, MT-2B, CCL5, GDF15, ZFP36, SLC39A8, STAT5A, ANGPTL4, STC2, TSKU, TFRC, GJA1, CCR1, SESN2, STAT3, CASP3, FTH1, CEBPG, DRAM2, EGR1, SLC1A5, SOAT1, BPIFA1, GCK, DDIT3, AQP7, ST6GALNAC1, HSPA9, BCL6, ADRA2A, CEBPB, SELENOS, GHITM, TLR4, ARRDC3, ORMDL1, KIT, SLC31A1, LYAR, GOT1, FBXO4, RIPK3, ABHD5, MYC, TMEM38B, INSIG1, NLE1, TXNRD1, SLC41A1, SFTPD, VDR, HK2, JAK2* |
| GO:0006397 | MRNA processing | 169 | 1.654 | 4.13E-04 | 1.32E-03 | *RNASEL, NHP2, SNRPF, SNRPD1, PAPOLA, RBM8A, SNRPA1, SMNDC1, SNRPG, SNRNP27, SNRPD3, TBRG4, TRA2B, GEMIN6, SMN1, SNRPA, RPRD1B, ZMAT2, STRAP, WTAP, SF3B6, PRPF40A, CLNS1A, FMR1, SNRPC, SNRPB2, SMU1, USP39, HNRNPF, CSTF2, SRSF1, EIF4A3, RBM7, SNRPE, SNRNP40, SRSF7, LSM6, RBM14, DCPS, PRPF19, RNPS1, RBM47, CSTF3, METTL3, NSRP1, SNU13, RBMX2, RBM25, WDR83, RBM42, TENT2, CDK12, LSM5, SCAF11, CSTF2T, LEO1, AHCYL1, GEMIN8, U2AF1L4, CWC22, RBM39, THOC7, PRPF3, RBBP6, SYMPK, UBL5, ZRANB2, PPIL1, SRSF2, LSM8, C1H9orf78, HNRNPK, PTBP1, BUD31, PAN2, GEMIN2, SDE2, PCF11, PTBP3, RBM15, NCBP2, PABPC1, U2AF2, SSU72, RBM41, SF3A3, LSM2, RBMX, CLP1, SON, CDC40, DDX39B, FRA10AC1, DYRK1A, MFAP1, SNRNP35, SFSWAP, EFTUD2, SF3B4* |

| | | | | | | |
|---|---|---|---|---|---|---|
| GO:0000375 | RNA splicing, via transesterification reactions | 107 | 1.645 | 2.75E-03 | 6.66E-03 | *NHP2, SNRPF, SNRPD1, RBM8A, SNRPA1, SNRPG, SNRPD3, TRA2B, GEMIN6, SNRPA, ZMAT2, STRAP, WTAP, SF3B6, PRPF40A, CLNS1A, FMR1, SNRPC, SNRPB2, SMU1, USP39, SRSF1, EIF4A3, RBM7, SNRPE, SRSF7, LSM6, RBM14, DCPS, PRPF19, METTL3, NSRP1, SNU13, RBMX2, WDR83, RBM42, LSM5, SCAF11, GEMIN8, U2AF1L4, CWC22, RBM39, PRPF3, UBL5, PPIL1, SRSF2, LSM8, C1H9orf78, PTBP1, BUD31, GEMIN2, RBM15, NCBP2, RBM41, SF3A3, LSM2, SON, CDC40, DDX39B, FRA10AC1, DYRK1A, MFAP1, SNRNP35, SFSWAP, EFTUD2, SF3B4, CWC15* |
| GO:0000377 | RNA splicing, via transesterification reactions with bulged adenosine as nucleophile | 107 | 1.645 | 2.75E-03 | 6.66E-03 | *NHP2, SNRPF, SNRPD1, RBM8A, SNRPA1, SNRPG, SNRPD3, TRA2B, GEMIN6, SNRPA, ZMAT2, STRAP, WTAP, SF3B6, PRPF40A, CLNS1A, FMR1, SNRPC, SNRPB2, SMU1, USP39, SRSF1, EIF4A3, RBM7, SNRPE, SRSF7, LSM6, RBM14, DCPS, PRPF19, METTL3, NSRP1, SNU13, RBMX2, WDR83, RBM42, LSM5, SCAF11, GEMIN8, U2AF1L4, CWC22, RBM39, PRPF3, UBL5, PPIL1, SRSF2, LSM8, C1H9orf78, PTBP1, BUD31, GEMIN2, RBM15, NCBP2, RBM41, SF3A3, LSM2, SON, CDC40, DDX39B, FRA10AC1, DYRK1A, MFAP1, SNRNP35, SFSWAP, EFTUD2, SF3B4, CWC15* |
| GO:0000398 | MRNA splicing, via spliceosome | 107 | 1.645 | 2.75E-03 | 6.66E-03 | *NHP2, SNRPF, SNRPD1, RBM8A, SNRPA1, SNRPG, SNRPD3, TRA2B, GEMIN6, SNRPA, ZMAT2, STRAP, WTAP, SF3B6, PRPF40A, CLNS1A, FMR1, SNRPC, SNRPB2, SMU1, USP39, SRSF1, EIF4A3, RBM7, SNRPE, SRSF7, LSM6, RBM14, DCPS, PRPF19, METTL3, NSRP1, SNU13, RBMX2, WDR83, RBM42, LSM5, SCAF11, GEMIN8, U2AF1L4, CWC22, RBM39, PRPF3, UBL5, PPIL1, SRSF2, LSM8, C1H9orf78, PTBP1, BUD31, GEMIN2, RBM15, NCBP2, RBM41, SF3A3, LSM2, SON, CDC40, DDX39B, FRA10AC1, DYRK1A, MFAP1, SNRNP35, SFSWAP, EFTUD2, SF3B4, CWC15* |
| GO:0098771 | Inorganic ion homeostasis | 133 | 1.640 | 1.30E-03 | 3.61E-03 | *CXCL9, CXCL11, CXCL10, IL1A, CCL8, SNX10, CP, CD40, MT-2B, CCL5, SLC39A8, STC2, TFRC, CCR1, FTH1, DDIT3* |
| GO:0045184 | Establishment of protein localization | 452 | 1.639 | 1.36E-05 | 5.95E-05 | *IFNG, NEURL3, ISG15, RSAD2, IL6, IL1A, TNFAIP3, SNX10, RHBDF2, STX11, EGR2, FFAR2, CCL5, CD74, NFKBIA, TFRC, ZFAND2A, STAT3, LRRC32, TIMM8A, ARL5B, PLK3, DDIT3, RAB3D, RHOB, HSPA9, ADRA2A, SEC23B, DERL3, KPNA2, OPTN, FBXO7, TXNIP,* |

| | | | | | | |
|---|---|---|---|---|---|---|
| | | | | | | *TOMM40, SCFD2, TGFB3, SELENOS, IPO4, TLR4, PREB, ARRDC3, LOC100157391, SRPRA, CALR, SEC24A, ARF4, TMEM126A, TLR2, RAB39B, INSIG1, SEC61A1, SEC61B, CHCHD4, TMED5, STXBP1, SYS1, USO1, SEC22B, SSR3, AP3S2, ZPR1, F2RL2, RAB1A, GDAP1, SPCS1, NDFIP2, ARCN1, TMED2, TMED10, DNAJC15, COPE, C7H6orf125, CADPS2, SRP14, C2CD2L, TRAM1, TMED9, TVP23B, GOSR2, XPO6, IFT20, LRRC8A, COPB1, EMC3, VCP, YWHAG, KPNB1, ARF6, COPB2, DESI1, SDAD1, YIF1A, SAR1B, GDI2, NXT1, HYAL2, TIMM13, SRP54, COPZ1, SEC63, XPO5, PHAX, SNX19, MACIR, BCS1L, MTX1, PDCD10, SNF8, SRI, TIMM44, RAB9A, RAB20, AKIRIN2, RAB13, SQSTM1, RAB2A, SNX5, TIMM8B, SRP9, NUTF2, TOMM20, TMED1, CHMP4A, STX17, NUP188, SRP68, ENY2, ARF1, TNPO1, TIMM21, VTI1B, SEC24C, YIPF5, CLTC, NSF, APOD, ARF2, MTX2, ZDHHC9, AP3M1, ELAVL1, PHB2, BBS5, TMEM129, MPC2, VPS26A* |
| GO:0051094 | Positive regulation of developmental process | 269 | 1.639 | 8.12E-05 | 2.98E-04 | *CXCL9, IFNG, CHI3L1, ISG15, CXCL8, IL1A, TNFSF13B, STAT1, CSF3, SOCS1, THBS1, JUNB, EGR2, PIM1, CD40, GDF15, ZFP36, STAT5A, IL10, CEBPD, LTF, TFRC, TNFSF14, GJA1, LIF, CCR1, DUSP10, STAT3, LRRTM2, RHOB, SOX11, BCL6, MAD2L2, CEBPB, TNFSF9, CASP8, LPAR3, YBX3, TGFB3, B4GALT5, MEDAG, C13H21orf91, ITGA5, BMP2, MIEF1* |
| GO:0051049 | Regulation of transport | 313 | 1.629 | 1.07E-04 | 3.77E-04 | *CXCL9, CXCL11, CXCL10, IFNG, CCL2, IL1B, S100A8, RSAD2, IL6, IL1A, THBS1, RHBDF2, IL15, FFAR2, LIPG, NR4A3, CCL5, KCNJ2, PDZK1, P2RY2, C3, NFKBIA, GJA1, LIF, CCR1, PLK3, LRRTM2, BPIFA1, PLA2G1B, SOX11, ADRA2A, GPM6B, DERL3, CEBPB, TGFB3, TLR4, CALR, LYAR, KCNJ14, TLR2, TMEM38B, INSIG1, BMP2, EIPR1, ZPR1, SFTPD, TP53, F2RL2, CRY1, NDFIP2, TMED10, INHBA, ADORA1, NKX3-1, SLC25A5, C7H6orf125, IL16, CADPS2, ATP1B1, NUS1, KCNA2* |
| GO:0010498 | Proteasomal protein catabolic process | 131 | 1.626 | 1.90E-03 | 5.02E-03 | *PSMB9, PSME2, PSME1, PSMA5, RNF122, ZFAND2A, HSP90B1, DDIT3, DNAJB9, PSMB3, DERL3, FBXO7, TNFAIP1, SELENOS, PSMD10, CALR, FBXO4, UBE2D1, KLHDC3, PAQR3, YOD1, PSMD8, SIAH1, RAD23A, USP44, VCP, DESI1, STT3B, UBE2B, PSMC1, PSME3, TAF9,* |

| | | | | | | |
|---|---|---|---|---|---|---|
| | | | | | | *SIAH2, PSMF1, AKIRIN2, PSMD4, RNF5, UBXN4, DDRGK1, OGT, EDEM2, NFE2L2, TMEM129, PSMB7, FBXO9, N4BP1, RAD23B, PSMC5, SOCS5, TOPORS, ARIH2* |
| GO:0046903 | Secretion | 189 | 1.624 | 6.56E-04 | 2.01E-03 | *IFNG, IL1B, S100A8, RSAD2, IL6, IL1A, GNAT1, CCL8, SNX10, RHBDF2, STX11, FFAR2, CCL5, STAT5A, P2RY2, GJA1, LIF, CCR1, NME1, SOX11, ADRA2A, TGFB3, TLR4, HRH2, KIT, TLR2, TMEM38B, BMP2, AQP5, VDR, TP53, F2RL2, KCNK1, CRY1, RAB1A, TMED10, INHBA, ADORA1, NKX3-1, C7H6orf125, CADPS2, KCNA2, C2CD2L, TVP23B, PTGES2, LRRC8A* |
| GO:0045597 | Positive regulation of cell differentiation | 170 | 1.623 | 1.92E-03 | 5.04E-03 | *CXCL9, IFNG, ISG15, STAT1, CSF3, SOCS1, JUNB, EGR2, PIM1, ZFP36, STAT5A, IL10, CEBPD, LTF, TNFSF14, GJA1, LIF, CCR1, DUSP10, STAT3, SOX11, BCL6, CEBPB, TNFSF9, CASP8, LPAR3, TGFB3* |
| GO:0040017 | Positive regulation of locomotion | 143 | 1.623 | 2.34E-03 | 5.94E-03 | *CXCL10, MADCAM1, IFNG, CCL2, CXCL8, CCL4, CCL8, THBS1, CD40, NR4A3, CCL5, BVES, TNFSF14, CCR1, CXCL16, RHOB, BCL6, ADRA2A, CASP8, TLR4, CALR, KIT, TWIST2, BMP2, LOC100516039* |
| GO:0036211 | Protein modification process | 430 | 1.622 | 2.21E-05 | 8.93E-05 | *IFNG, CHI3L1, ISG15, UBA7, USP18, SOCS3, SOCS1, RNF19B, TNFAIP3, IL15, PARP12, EGR2, TRIM21, STAT2, SLC39A8, GALNT15, C3, CISH, LIF, RNF122, PARP11, EGR1, GNL3, TRAF2, GLUL, ST6GALNAC1, PLA2G1B, NEK6, DERL3, TICAM1, FBXO7, TNFAIP1, B4GALT1, PSMD10, B4GALT5, WSB2, USP15, FBXO4, RNF115, MOGS, OSTC, ITGA5, UBE2D1, ST3GAL1, EIPR1, GTPBP4, B3GNT5, POMGNT2, ALG3, RPN1, OTUB2, JAK2, RPN2, CRY1, KLHL25, NDFIP2, USP2, UBE2D3, RXYLT1, KAT2A, FUT2, MPDU1, NCOA7, NUS1, SOCS7, SPRY2, SIAH1, DAD1, MGAT2, PARP3, RNF138, DDX3X, TMEM258, B3GALT6, GALNT1, RNF114, USP44, VCP, NRP1, DESI1, PPP2R2A, MGAT1, A4GALT, STT3B, UBE2B, NPC1, ST8SIA2, DCUN1D5, B3GALNT1, DPAGT1, UCHL5, HINT1, PLAA, SIAH2, PHF23, FBXO33, USP39, PPP2CA, ATM, C1GALT1, PARP8, B4GALT3, UBA3, PPP1CB, SQSTM1, RLIM, DR1, RNF5, OTUB1, OTUD6B, ST6GAL2, BCL10, TRAF7, PIK3R5, DPM1, WBP1L, TPST1, FUT8, SEMA4D, COPS6, GAPDH, CUL5, MAGT1, DDRGK1, OGT,* |

| | | | | | | |
|---|---|---|---|---|---|---|
| | | | | | | *PADI4, DCUN1D3, EDEM2, RNF14, PPIA, PRPF19, NFE2L2, TMEM129, USP5, UFC1, PCNP, MINDY3, FBXO9* |
| GO:0055082 | Intracellular chemical homeostasis | 167 | 1.619 | 1.46E-03 | 4.01E-03 | *CXCL9, CXCL11, CXCL10, IL1A, CCL8, HK3, CP, CD40, MT-2B, CCL5, SLC39A8, STC2, TFRC, CCR1, FTH1, GCK, DDIT3, ADRA2A, GHITM, ORMDL1, SLC31A1, MYC, TMEM38B, SLC41A1, VDR, HK2* |
| GO:0050801 | Monoatomic ion homeostasis | 148 | 1.616 | 1.64E-03 | 4.41E-03 | *CXCL9, CXCL11, CXCL10, IL1A, CCL8, SNX10, CP, CD40, MT-2B, CCL5, SLC39A8, STC2, TFRC, CCR1, FTH1, DDIT3* |
| GO:0055080 | Monoatomic cation homeostasis | 147 | 1.616 | 1.43E-03 | 3.95E-03 | *CXCL9, CXCL11, CXCL10, IL1A, CCL8, SNX10, CP, CD40, MT-2B, CCL5, SLC39A8, STC2, TFRC, CCR1, FTH1, DDIT3* |
| GO:0009896 | Positive regulation of catabolic process | 128 | 1.605 | 4.03E-03 | 9.02E-03 | *SAA3, IFNG, TNFAIP3, ZFP36, SESN2, ZFAND2A, PLK3, ADRA2A, OPTN, TICAM1, FBXO7, PSMD10, ABHD5, STING1, PAQR3, IGF2BP1, SLC25A5, AGO2, RALB, PDE12, RIDA* |
| GO:0015031 | Protein transport | 395 | 1.601 | 9.32E-05 | 3.35E-04 | *IFNG, NEURL3, RSAD2, IL6, IL1A, SNX10, RHBDF2, STX11, EGR2, FFAR2, CCL5, CD74, NFKBIA, TFRC, STAT3, TIMM8A, ARL5B, PLK3, RAB3D, RHOB, HSPA9, ADRA2A, SEC23B, DERL3, KPNA2, OPTN, TXNIP, TOMM40, SCFD2, TGFB3, SELENOS, IPO4, TLR4, PREB, ARRDC3, LOC100157391, SRPRA, CALR, SEC24A, ARF4, TLR2, RAB39B, INSIG1, SEC61A1, SEC61B, CHCHD4, TMED5, STXBP1, SYS1, USO1, SEC22B, AP3S2, ZPR1, F2RL2, RAB1A, NDFIP2, ARCN1, TMED2, TMED10, DNAJC15, COPE, C7H6orf125, CADPS2, C2CD2L, TRAM1, TMED9, TVP23B, GOSR2, XPO6, IFT20, LRRC8A, COPB1, VCP, KPNB1, ARF6, COPB2, DESI1, SDAD1, YIF1A, SAR1B, GDI2, NXT1, HYAL2, TIMM13, SRP54, COPZ1, SEC63, XPO5, PHAX, SNX19, MACIR, MTX1, PDCD10, SNF8, SRI, TIMM44, RAB9A, RAB20, AKIRIN2, RAB13, SQSTM1, RAB2A, SNX5, TIMM8B, NUTF2, TOMM20, TMED1, STX17, NUP188, ENY2, ARF1, TNPO1, TIMM21, VTI1B, SEC24C, YIPF5, CLTC, NSF, APOD, ARF2, MTX2, AP3M1, ELAVL1, PHB2, BBS5, TMEM129, MPC2, VPS26A* |
| GO:0034220 | Monoatomic ion transmembrane transport | 131 | 1.595 | 3.01E-03 | 7.21E-03 | *CXCL9, CXCL11, CXCL10, CALHM6, CNGA4, SLC39A8, KCNJ2, PDZK1, GLRB, KCNMA1, PANX2, BPIFA1, DDIT3, SLC25A28, PLA2G1B, ADRA2A, GHITM, SLC31A1, KCNJ14, TMEM38B, STING1* |

| | | | | | | |
|---|---|---|---|---|---|---|
| GO:0008285 | Negative regulation of cell population proliferation | 156 | 1.583 | 2.62E-03 | 6.56E-03 | *IFNG, CD274, CCL2, IL1A, STAT1, PTGIR, TNFAIP3, THBS1, IFI35, IL10, ATF5, LIF, DUSP10, LRRC32, SOX11, BCL6, IFI30, CEBPB, TGFB3, B4GALT1, MORC3, ARG2, FBXO4, BMP2, GTPBP4, VDR, DLL4, JAK2, KLF9, SMAD1, INHBA, NKX3-1, PPARGC1A* |
| GO:0019725 | Cellular homeostasis | 199 | 1.575 | 2.14E-03 | 5.53E-03 | *CXCL9, CXCL11, CXCL10, IL1A, CCL8, SNX10, HK3, CP, CD40, MT-2B, CCL5, SLC39A8, STC2, TFRC, CCR1, FTH1, GCK, DDIT3, ADRA2A, SELENOS, GHITM, ORMDL1, SLC31A1, FBXO4, MYC, TMEM38B, TXNRD1, SLC41A1, VDR, HK2* |
| GO:0009894 | Regulation of catabolic process | 237 | 1.573 | 9.67E-04 | 2.80E-03 | *SAA3, IFNG, PSME2, TNFAIP3, TIMP1, PSME1, ZFP36, IL10, SESN2, ZFAND2A, PLK3, ARID5A, DRAM2, TRAF2, MAD2L2, TIGAR, ADRA2A, DRAM1, OPTN, TICAM1, FBXO7, PSMD10, MCL1, DDIT4, ABHD5, RAB39B, TIRAP, STING1, PAQR3, RBM8A, PSMD3, DEPP1, IGF2BP1, SLC25A5, AGO2, DDX49, RALB, FBP1, RUBCN, PDE12, TMEM39A, RIDA, TBRG4, NPM1, DESI1, PSMD1, PSME3, TAF9, PHF23, PHAX, FMR1, PPP2CA, SNF8, PTK2B, PHKG2, CDC37, EIF4A3, PSMD2, SQSTM1, CNOT7, RNF5, ORMDL3, TRAF7* |
| GO:0032879 | Regulation of localization | 413 | 1.571 | 8.89E-05 | 3.23E-04 | *CXCL9, CXCL11, CXCL10, IFNG, CCL2, IL1B, S100A8, RSAD2, IL6, IL1A, THBS1, RHBDF2, IL15, FFAR2, LIPG, NR4A3, CCL5, ZFP36, KCNJ2, PDZK1, P2RY2, C3, NFKBIA, GJA1, LIF, CCR1, SLC5A3, SESN2, PLK3, LRRTM2, GNL3, BPIFA1, DDIT3, PLA2G1B, SOX11, ADRA2A, GPM6B, PLIN2, DERL3, CEBPB, TGFB3, GHITM, TLR4, CALR, LYAR, FBXO4, ABHD5, MRAP2, KCNJ14, TLR2, TMEM38B, INSIG1, BMP2, ABHD17B, EIPR1, ZPR1, SFTPD, TP53, F2RL2, HILPDA, CRY1, NDFIP2, TMED10, INHBA, ADORA1, NKX3-1, SLC25A5* |
| GO:0051046 | Regulation of secretion | 116 | 1.565 | 6.25E-03 | 1.32E-02 | *IFNG, IL1B, S100A8, RSAD2, IL6, IL1A, RHBDF2, FFAR2, CCL5, P2RY2, GJA1, LIF, SOX11, ADRA2A, TGFB3, TLR4, TLR2, BMP2, TP53, F2RL2, CRY1, TMED10, INHBA, ADORA1, NKX3-1, C7H6orf125, CADPS2, KCNA2, C2CD2L* |
| GO:2001233 | Regulation of apoptotic signaling pathway | 108 | 1.563 | 8.23E-03 | 1.70E-02 | *S100A8, ATF3, TNFAIP3, THBS1, IL10, FASLG, TRAF2, DDIT3, FBXO7, YBX3, SELENOS, GHITM, PSMD10, MCL1, TLR4, CFLAR, RIPK3, TP53, JAK2, INHBA, NKX3-1, SLC25A5, WNT16, G0S2, SIAH1, ACKR3, DDX3X, NRP1, SLC35F6, HYAL2, SNAI2, PSME3, SIAH2* |

| | | | | | | |
|---|---|---|---|---|---|---|
| GO:0051241 | Negative regulation of multicellular organismal process | 230 | 1.561 | 8.08E-04 | 2.36E-03 | *CXCL10, IFNG, CD274, IRF1, SERPINE1, DHX58, STAT1, SOCS1, TNFAIP3, THBS1, GDF15, ZFP36, IL10, TSKU, LTF, GJA1, CCR1, DNAJB11, DUSP10, LRRC32, KCNMA1, DDIT3, HSPA9, SOX11, BCL6, MAD2L2, RGS2, CEBPB, FBXO7, YBX3, TGFB3, TMEM176B, JAK3, TLR4, ARRDC3, MORC3, CALR, ARG2, CREB3L1, MYC, BMP2, GPR4, TP53, ELF4, CRY1, ANGPT2, CD200, INHBA, ADORA1, ADM, NKX3-1* |
| GO:0045595 | Regulation of cell differentiation | 312 | 1.558 | 5.77E-04 | 1.80E-03 | *CXCL9, CXCL10, IFNG, IRF1, ISG15, IL27, STAT1, CSF3, SOCS1, JUNB, EGR2, PIM1, ZFP36, STAT5A, IL10, CEBPD, NFKBIA, TSKU, LTF, TNFSF14, GJA1, LIF, CCR1, DNAJB11, DUSP10, MAFF, STAT3, DDIT3, TCIM, HSPA9, SOX11, BCL6, MAD2L2, CEBPB, TNFSF9, CASP8, FBXO7, LPAR3, TGFB3, TMEM176B, JAK3, MEDAG, CALR, MYC, INSIG1, TWIST2, BMP2, PRDM1, VDR, DLL4, IKBKB, SMAD1, INHBA, LIG4, ADM, KAT2A, C7H6orf125, KLF10, S1PR2, RASSF2, TFE3, PDCD4, SPRY2, CYP26B1, CLDN5, RUNX1* |
| GO:2000026 | Regulation of multicellular organismal development | 306 | 1.553 | 3.78E-04 | 1.22E-03 | *CXCL10, IFNG, IRF1, CHI3L1, ISG15, CXCL8, IL27, SERPINE1, IL1A, STAT1, CSF3, SOCS1, THBS1, EGR2, PIM1, CD40, ZFP36, STAT5A, IL10, NFKBIA, LTF, GJA1, LIF, CCR1, DNAJB11, DUSP10, MAFF, STAT3, LRRTM2, PHLDA2, VASH2, GLUL, TCIM, RHOB, HSPA9, SOX11, BCL6, CEBPB, TNFSF9, CASP8, FBXO7, LPAR3, TMEM176B, JAK3, B4GALT5, CREB3L1, MYC, ITGA5, BMP2, PRDM1, GPR4, ZPR1, VDR, DLL4, TP53, IKBKB, SMAD1, ANGPT2, INHBA, LIG4, ADM, KAT2A* |
| GO:0030335 | Positive regulation of cell migration | 133 | 1.552 | 3.94E-03 | 8.89E-03 | *CXCL10, MADCAM1, IFNG, CCL2, CCL4, CCL8, THBS1, CD40, NR4A3, CCL5, TNFSF14, CCR1, CXCL16, RHOB, ADRA2A, CASP8, TLR4, CALR, KIT, TWIST2, BMP2, LOC100516039* |
| GO:0061061 | Muscle structure development | 141 | 1.551 | 6.35E-03 | 1.34E-02 | *CXCL9, CXCL10, MYBPH, ATF3, EGR2, MSC, GDF15, BVES, TNFSF14, GJA1, LIF, UNC45B, MAFF, EGR1, DDIT3, SOX11, YBX3* |
| GO:2000147 | Positive regulation of cell motility | 139 | 1.545 | 3.51E-03 | 8.20E-03 | *CXCL10, MADCAM1, IFNG, CCL2, CCL4, CCL8, THBS1, CD40, NR4A3, CCL5, TNFSF14, CCR1, CXCL16, RHOB, BCL6, ADRA2A, CASP8, TLR4, CALR, KIT, TWIST2, BMP2, LOC100516039* |
| GO:0010629 | Negative regulation of gene expression | 279 | 1.543 | 6.23E-04 | 1.93E-03 | *SAA3, IFNG, CXCL8, SERPINE1, DHX58, TNFAIP3, THBS1, ZFP36, IL10, TSKU, LTF, GJA1, CCR1, SESN2, LRRC32, MOV10, ARID5A, EIF4EBP1, TRAF2, DDIT3, SOX11, BCL6, BAZ1A,* |

| | | | | | | |
|---|---|---|---|---|---|---|
| | | | | | | *TGFB3, TLR4, MORC3, CALR, ARG2, CREB3L1, PNPT1, RNASEH2B, TIRAP, BMP2, EIF2S1, PELO, PRDM1, RBM8A, DLL4, ETF1, ELF4, NDFIP2, IGF2BP1, NKX3-1, AGO2, DXO, HAVCR2, EXOSC3, PDCD4, PDE12, RIDA, CLDN5, PARP3, TBRG4, DDX3X* |
| GO:0006508 | Proteolysis | 432 | 1.537 | 3.05E-04 | 1.02E-03 | *IFNG, MMP8, ISG15, MMP12, PSMB9, SERPINE1, USP18, PSME2, PSMB8, MMP1, RNF19B, PSMB10, LOC100522887, CTSL, TNFAIP3, THBS1, CFB, TIMP1, MMP9, PSME1, CASP10, C2, MMP13, IL10, MMP27, LTF, PSMA5, RNF122, ZFAND2A, HSP90B1, ASRGL1, CASP3, PSMA2, DPP3, PSMA3, DDIT3, CNDP2, GZMA, DNAJB9, CASP1, ADRA2A, PSMB3, DERL3, CASP8, PSMA1, FBXO7, TNFAIP1, SELENOS, PSMD10, ADAM17, CFLAR, CALR, USP15, FBXO4, RNF115, TMEM126A, PSMB4, UBE2D1, SPCS3, CFI, KLHDC3, PAQR3, SEC11C, YOD1, OTUB2, PSMA4, ENPEP, PSMD3, CD46, SPCS1, NDFIP2, USP2, UBE2D3, PISD, AEBP1, SPCS2, DNPEP* |
| GO:0006811 | Monoatomic ion transport | 220 | 1.526 | 2.07E-03 | 5.38E-03 | *CXCL9, CXCL11, CXCL10, CCL2, STEAP4, CALHM6, CCL8, SLC39A14, CNGA4, CCL5, SLC39A8, KCNJ2, PDZK1, LTF, TFRC, CCR1, GLRB, FTH1, ATP1B3, KCNMA1, PANX2, BPIFA1, DDIT3, SLC25A28, PLA2G1B, ADRA2A, GHITM, SLC5A6, SLC11A2, SLC31A1, KCNJ14, TMEM38B, STING1, SLC39A7, VDR, KCNK1, KCNK6, NDFIP2, SLC30A7, SLCO4C1, IL16, ATP1B1, KCNA2, STEAP3, SLC10A7, SLC30A4, SLC11A1, SLC38A2, ATOX1, SLC4A4, LRRC8A, SLC41A2* |
| GO:0001701 | In utero embryonic development | 110 | 1.526 | 1.42E-02 | 2.70E-02 | *FOSL1, JUNB, PSPH, IL10, GJA1, LIF, MAFF, CSF2, PHLDA2, RCN1, EIF2S2, CEBPB, YBX3, RNASEH2B, NLE1, BMP2, WDR74, PRDM1, ZPR1, GNA13, LIG4, ADM, KAT2A, RPL7L1, SLC39A1, DAD1, RSPO3, ZFAND5, UBE2B, SF3B6, APBA3, HORMAD1* |
| GO:0043687 | Post-translational protein modification | 268 | 1.518 | 1.16E-03 | 3.25E-03 | *ISG15, UBA7, USP18, SOCS3, SOCS1, RNF19B, TNFAIP3, PARP12, EGR2, TRIM21, CISH, RNF122, PARP11, EGR1, GNL3, TRAF2, TICAM1, FBXO7, TNFAIP1, PSMD10, WSB2, USP15, FBXO4, RNF115, UBE2D1, EIPR1, GTPBP4, OTUB2, JAK2, CRY1, KLHL25, NDFIP2, USP2, UBE2D3, SOCS7, SPRY2, SIAH1, PARP3, RNF138, DDX3X, RNF114, USP44, VCP, DESI1, STT3B, UBE2B, DCUN1D5, UCHL5, HINT1, PLAA, SIAH2, PHF23, FBXO33, USP39, PARP8, UBA3, SQSTM1, RLIM, RNF5, OTUB1, OTUD6B, BCL10, TRAF7, WBP1L, TPST1,* |

| | | | | | | |
|---|---|---|---|---|---|---|
| | | | | | | *COPS6, CUL5, DDRGK1, OGT, PADI4, DCUN1D3, RNF14, PPIA, PRPF19, NFE2L2, TMEM129, USP5, UFC1, PCNP, MINDY3, FBXO9, N4BP1, UBE3A, DCAF11, SOCS5, TOPORS, UCHL3, ARIH2, MSL2* |
| GO:0048646 | Anatomical structure formation involved in morphogenesis | 275 | 1.516 | 1.51E-03 | 4.10E-03 | *CXCL9, CXCL10, CHI3L1, MMP8, CXCL8, SERPINE1, IL1A, TNFSF13B, STAT1, MYBPH, DUSP5, TNFAIP3, THBS1, JUNB, PIM1, CD40, GDF15, ANGPTL4, IL10, TNFSF14, GJA1, CHN2, VASH2, ESM1, GLUL, RHOB, SOX11, BCL6, GARIN1A, B4GALT1, SLC31A1, CREB3L1, ITGA5, NLE1, BMP2, WDR74, GPR4, DLL4, GNA13, SMAD1, ANGPT2, PLEKHO1, INHBA, ADM, KAT2A, NKX3-1* |
| GO:0006812 | Monoatomic cation transport | 196 | 1.516 | 5.51E-03 | 1.20E-02 | *CXCL9, CXCL11, CXCL10, CCL2, STEAP4, CCL8, SLC39A14, CNGA4, CCL5, SLC39A8, KCNJ2, PDZK1, LTF, TFRC, CCR1, FTH1, ATP1B3, KCNMA1, PANX2, BPIFA1, DDIT3, SLC25A28, PLA2G1B, ADRA2A, GHITM, SLC5A6, SLC11A2, SLC31A1, KCNJ14, TMEM38B, SLC39A7, VDR, KCNK1, KCNK6, NDFIP2, SLC30A7, IL16, ATP1B1, KCNA2, STEAP3, SLC10A7, SLC30A4, SLC11A1, SLC38A2, ATOX1, SLC4A4, SLC41A2* |
| GO:0008380 | RNA splicing | 154 | 1.514 | 7.57E-03 | 1.58E-02 | *NHP2, SNRPF, SNRPD1, ZPR1, RBM8A, SNRPA1, SMNDC1, CCNL1, RTCB, SNRPG, SNRNP27, SNRPD3, SLC38A2, TRA2B, GEMIN6, SMN1, SNRPA, ZMAT2, STRAP, MAGOHB, WTAP, SF3B6, PRPF40A, CLNS1A, FMR1, SNRPC, SNRPB2, SMU1, USP39, HNRNPF, SRSF1, EIF4A3, RBM7, SNRPE, SNRNP40, SRSF7, LSM6, RBM14, DCPS, PRPF19, RNPS1, TSEN15, RBM47, MAGOH, METTL3, NSRP1, SNU13, RBMX2, RBM25, WDR83, RBM42, CDK12, LSM5, SCAF11, GEMIN8, U2AF1L4, CWC22, RBM39, MBNL1, PRPF3, UBL5, ZRANB2, PPIL1, SRSF2, LSM8, C1H9orf78, HNRNPK, PTBP1, BUD31, GEMIN2, SDE2, PTBP3, RBM15, NCBP2, ZBTB8OS, U2AF2, RBM41, SF3A3, LSM2, RBMX, CLP1, SON, CDC40, DDX39B, FRA10AC1, DYRK1A, MFAP1, SNRNP35, SFSWAP, EFTUD2, SF3B4* |
| GO:0008284 | Positive regulation of cell population proliferation | 215 | 1.513 | 2.74E-03 | 6.66E-03 | *IL1B, IL6, IL1A, TNFSF13B, STAT1, CSF3, TNFAIP3, THBS1, IL15, PIM1, TIMP1, CD40, NR4A3, CCL5, STAT5A, IL10, LTF, TFRC, GJA1, LIF, NME1, CSF2, ESM1, PLA2G1B, SOX11, BCL6, SLC7A1, TNFSF9, TICAM1, TGFB3, B4GALT1, TLR4, CALR, FAM98A, NOP2,* |

| | | | | | | |
|---|---|---|---|---|---|---|
| | | | | | | *SLC25A33, MYC, RNASEH2B, IGFBP2, TIPIN, BMP2, DLL4, JAK2, HILPDA, LIG4, ADM, NKX3-1, SLC25A5* |
| GO:0098655 | Monoatomic cation transmembrane transport | 121 | 1.510 | 9.25E-03 | 1.90E-02 | *CXCL9, CXCL11, CXCL10, CNGA4, SLC39A8, KCNJ2, PDZK1, KCNMA1, BPIFA1, DDIT3, SLC25A28, PLA2G1B, ADRA2A, GHITM, SLC31A1, KCNJ14, TMEM38B* |
| GO:0010608 | Post-transcriptional regulation of gene expression | 137 | 1.502 | 9.90E-03 | 2.01E-02 | *THBS1, ZFP36, SESN2, ARID5A, EIF4EBP1, TRAF2, RGS2, CALR, FBXO4, TIRAP, EIF2S1, EIF5A, ETF1, PUM3, IGF2BP1, AGO2, C7H6orf125, NAT10, DDX49, EIF1, PDE12, RIDA, TBRG4, DDX3X, BZW1, NPM1, IGF2BP2, ABCF1, PHAX, FMR1, OGFOD1, CDC37, ACO1, EIF4A3, CNOT7, SRP9, SERBP1, SSB, PTCD3, GAPDH, DCPS, OGT, PAIP2B, ELAVL1, FASTKD2, METTL3, DAP, ZFP36L2, CAPRIN1, TENT2, STK35, GUF1, PCID2, PAIP1, UPF3B, NAF1, ROCK1, MTOR, PAIP2, PAN2, IGFBP5, MATR3, USP16, ZFP36L1, FXR1, NCBP2, NCK1, ZC3H14, LSM14A, CLP1, CSDE1, BCDIN3D, SPOUT1, RPS6KB1, NCK2* |
| GO:0030001 | Metal ion transport | 182 | 1.499 | 5.79E-03 | 1.25E-02 | *CXCL9, CXCL11, CXCL10, CCL2, STEAP4, CCL8, SLC39A14, CCL5, SLC39A8, KCNJ2, LTF, TFRC, CCR1, FTH1, ATP1B3, KCNMA1, BPIFA1, DDIT3, SLC25A28, PLA2G1B, ADRA2A, GHITM, SLC5A6, SLC11A2, SLC31A1, KCNJ14, TMEM38B, SLC39A7, VDR, KCNK1, KCNK6, NDFIP2, SLC30A7, IL16, ATP1B1, KCNA2, STEAP3, SLC10A7, SLC30A4, SLC11A1, SLC38A2, ATOX1, SLC4A4, SLC41A2* |
| GO:0032940 | Secretion by cell | 162 | 1.490 | 4.07E-03 | 9.05E-03 | *IFNG, IL1B, RSAD2, IL6, IL1A, GNAT1, CCL8, RHBDF2, STX11, FFAR2, CCL5, GJA1, LIF, CCR1, SOX11, ADRA2A, TGFB3, TLR4, KIT, TLR2, TMEM38B, BMP2, TP53, F2RL2, KCNK1, CRY1, RAB1A, TMED10, INHBA, NKX3-1, C7H6orf125, CADPS2, KCNA2* |
| GO:0040012 | Regulation of locomotion | 233 | 1.490 | 3.15E-03 | 7.47E-03 | *CXCL10, MADCAM1, IFNG, SELE, CCL2, CXCL8, CCL4, CCL8, THBS1, CD40, NR4A3, CCL5, GDF15, BVES, TNFSF14, GJA1, CCR1, DUSP10, CXCL16, PHLDA2, GLUL, RHOB, BCL6, ADRA2A, CASP8, FBXO7, TLR4, ARRDC3, KLRK1, CALR, KIT, RIPK3, TWIST2, BMP2, GTPBP4, LOC100516039* |
| GO:0007155 | Cell adhesion | 326 | 1.476 | 1.76E-03 | 4.71E-03 | *MADCAM1, IFNG, SELE, RND1, CD274, IRF1, IL1B, CXCL8, S100A8, IL1A, TNFSF13B, SOCS1, CLSTN3, THBS1, VCAM1, IL15, NR4A3, SLA-DQB1, SELP, CCL5, SLC39A8, BVES,* |

| | | | | | | |
|---|---|---|---|---|---|---|
| | | | | | | *STAT5A, IL10, SLA-DMB, TFRC, TNFSF14, LIF, TNC, DUSP10, LRRC32, RHOB, OLR1, BCL6, VCAN, MAD2L2, ICAM1, CDH12, SLC7A1, CEBPB, TNFSF9, B4GALT1, JAK3, ICAM4, CALR, ARG2, KIT, ITGA5, AMBN, THBS2, IGFBP2, BMP2* |
| GO:1903530 | Regulation of secretion by cell | 107 | 1.475 | 1.98E-02 | 3.66E-02 | *IFNG, IL1B, RSAD2, IL6, IL1A, RHBDF2, FFAR2, CCL5, GJA1, LIF, SOX11, ADRA2A, TGFB3, TLR4, TLR2, BMP2, TP53, F2RL2, CRY1, TMED10, INHBA, NKX3-1, C7H6orf125, CADPS2, KCNA2, C2CD2L* |
| GO:1902532 | Negative regulation of intracellular signal transduction | 170 | 1.474 | 1.26E-02 | 2.47E-02 | *DHX58, STAT1, ATF3, DUSP5, TNFAIP3, THBS1, IFI35, GDF15, SLC39A8, IL10, NFKBIA, LIF, DUSP10, SESN2, DDIT3, BCL6, OPTN, CASP8, FBXO7, YBX3, TNFAIP1, SELENOS, PSMD10, TLR4, CGAS, CALR, DDIT4, MYC, INSIG1, UBE2D1, PAQR3* |
| GO:0022603 | Regulation of anatomical structure morphogenesis | 188 | 1.468 | 4.60E-03 | 1.01E-02 | *CXCL10, CCL2, CHI3L1, CXCL8, SERPINE1, IL1A, TNFSF13B, STAT1, TNFAIP3, THBS1, CD40, STAT2, BVES, IL10, GJA1, LIF, SLC26A5, VASH2, GLUL, RHOB, LPAR3, KIT, CDC42EP3, CREB3L1, ITGA5, BMP2, MIEF1, CDC42EP2, GPR4, GNA13, CDC42SE1, SMAD1, ANGPT2, PLEKHO1, ADM* |
| GO:0070647 | Protein modification by small protein conjugation or removal | 256 | 1.467 | 3.59E-03 | 8.29E-03 | *ISG15, UBA7, USP18, SOCS3, SOCS1, RNF19B, TNFAIP3, EGR2, TRIM21, CISH, RNF122, EGR1, GNL3, TRAF2, TICAM1, FBXO7, TNFAIP1, PSMD10, WSB2, USP15, FBXO4, RNF115, UBE2D1, EIPR1, GTPBP4, OTUB2, CRY1, KLHL25, NDFIP2, USP2, UBE2D3, SOCS7, SPRY2, SIAH1, RNF138, DDX3X, RNF114, USP44, VCP, DESI1, UBE2B, DCUN1D5, UCHL5, HINT1, PLAA, SIAH2, PHF23, FBXO33, USP39, UBA3, SQSTM1, RLIM, RNF5, OTUB1, OTUD6B, BCL10, TRAF7, WBP1L, COPS6, CUL5, DDRGK1, OGT, DCUN1D3, RNF14, PPIA, PRPF19, NFE2L2, TMEM129, USP5, UFC1, PCNP, MINDY3, FBXO9, N4BP1, UBE3A, DCAF11, SOCS5, TOPORS, UCHL3, ARIH2, MSL2* |
| GO:0141124 | Intracellular signaling cassette | 454 | 1.466 | 9.86E-04 | 2.81E-03 | *SAA3, RND1, S100A12, CHI3L1, RASGEF1A, STAT1, FAS, ATF3, DUSP5, TNFAIP3, TIFA, THBS1, CD40, IFI35, TICAM2, CCL5, ACKR4, GDF15, ZFP36, SLC39A8, NFKBIB, CD74, NFKBIA, TNFSF14, LIF, CCR1, DUSP10, SESN2, FASLG, GADD45G, CSF2, CHN2, CXCR6, TRAF2, DDIT3, RASSF1, PLA2G1B, TCIM, RHOB, TRAF3, DDX21, ACKR2, RHOJ, BCL6, ADRA2A, TNIP1, OPTN, CASP8, TICAM1, MYD88, LPAR3, TNFAIP1, PSMD10, TLR4, TNIP2,* |

| | | | | | | |
|---|---|---|---|---|---|---|
| | | | | | | *ADAM17, CGAS, CFLAR, GPR37, CALR, CDC42EP3, TMEM106A, RIPK3, MYC, TLR2, RAB39B, TMEM38B, TIRAP, BMP2, CDC42EP2, GADD45B, EIF2S1, GPR4, JAK2, GNA13, LAMTOR2, IKBKB, CDC42SE1, SMAD1, SMAD3, INHBA, NKX3-1, WNT16, RIPK1, HAVCR2, TERF2IP, RALB, FBP1, ATP1B1, LTBR, RASSF2, PDCD4* |
| GO:0060284 | Regulation of cell development | 177 | 1.460 | 1.09E-02 | 2.18E-02 | *IFNG, IRF1, ISG15, IL27, STAT1, CSF3, SOCS1, EGR2, ZFP36, STAT5A, IL10, NFKBIA, LTF, LIF, CCR1, DNAJB11, DUSP10, STAT3, TCIM, HSPA9, SOX11, BCL6, CEBPB, TNFSF9, CASP8, FBXO7, LPAR3, TMEM176B, JAK3, MYC, BMP2, PRDM1, DLL4, IKBKB, INHBA, LIG4, KAT2A, KLF10, S1PR2, RASSF2, TFE3, CYP26B1, CLDN5, RUNX1* |
| GO:0009725 | Response to hormone | 167 | 1.460 | 1.18E-02 | 2.33E-02 | *STAT1, SOCS1, JUNB, PIM1, TIMP1, NR4A3, GDF15, ZFP36, STAT2, STAT5A, IL10, SESN2, STAT3, ARID5A, EGR1, PLA2G1B, ADRA2A, TGFB3, JAK3, CALR, DDIT4, SLC2A4, GOT1, SLC25A33, IGFBP2, PTGES3, VDR, JAK2, KLF9, CRY1, NKX3-1, GSTM3* |
| GO:0048585 | Negative regulation of response to stimulus | 401 | 1.454 | 2.56E-03 | 6.46E-03 | *CCL2, ISG15, CXCL8, SERPINE1, DHX58, STAT1, SOCS3, SOCS1, PTGIR, ATF3, DUSP5, BPIFB1, TNFAIP3, THBS1, RGS16, PIM1, IFI35, GDF15, ZFP36, STAT2, SLC39A8, IL10, NFKBIA, TSKU, LTF, CISH, LIF, DUSP10, SESN2, EGR1, DDIT3, PAK1IP1, SERPING1, TCIM, BCL6, DKK1, MAD2L2, ADRA2A, RGS2, OPTN, CASP8, FBXO7, YBX3, TNFAIP1, SELENOS, GHITM, JAK3, PSMD10, MCL1, TLR4, ARRDC3, CGAS, KLRK1, CFLAR, CALR, ARG2, DDIT4, LYAR, CREB3L1, RNF115, MRAP2, MYC, PSMB4, INSIG1, UBE2D1, GPR155, BMP2, PAQR3, DLL4, ELF4, CD46, CRY1, ANGPT2, ADORA1, ADM, NKX3-1, SLC25A5, WNT16, CD59, RASL11B, HAVCR2, NCOA7, FBP1, RASSF2, SOCS7* |
| GO:0030334 | Regulation of cell migration | 209 | 1.443 | 6.04E-03 | 1.29E-02 | *CXCL10, MADCAM1, IFNG, SELE, CCL2, CCL4, CCL8, THBS1, CD40, NR4A3, CCL5, GDF15, TNFSF14, GJA1, CCR1, DUSP10, CXCL16, PHLDA2, GLUL, RHOB, ADRA2A, CASP8, TLR4, KLRK1, CALR, KIT, RIPK3, TWIST2, BMP2, GTPBP4, LOC100516039, DLL4, GNA13* |
| GO:0048871 | Multicellular organismal-level homeostasis | 185 | 1.442 | 1.49E-02 | 2.82E-02 | *IL1B, ISG15, IL1A, TNFSF13B, STAT1, FAS, TNFAIP3, SNX10, AQP3, GDF15, ZFP36, SLC39A8, STAT5A, TSKU, GJA1, STAT3, CASP3, CEBPG, DRAM2, EGR1, SLC1A5, BPIFA1,* |

| | | | | | | |
|---|---|---|---|---|---|---|
| | | | | | | *DDIT3, AQP7, ST6GALNAC1, HSPA9, BCL6, CEBPB, TLR4, ARRDC3, KIT, LYAR, RIPK3, NLE1* |
| GO:0098660 | Inorganic ion transmembrane transport | 132 | 1.428 | 1.73E-02 | 3.23E-02 | *CXCL9, CXCL11, CXCL10, SLC39A8, KCNJ2, GLRB, KCNMA1, BPIFA1, DDIT3, SLC25A28, PLA2G1B, ADRA2A, GHITM, SLC5A6, SLC31A1, KCNJ14, TMEM38B, VDR, KCNK1, KCNK6, SLC30A7* |
| GO:0010648 | Negative regulation of cell communication | 352 | 1.427 | 3.02E-03 | 7.21E-03 | *ISG15, CXCL8, DHX58, STAT1, SOCS3, SOCS1, PTGIR, ATF3, DUSP5, BPIFB1, TNFAIP3, THBS1, RGS16, FFAR2, IFI35, GDF15, STAT2, SLC39A8, IL10, NFKBIA, TSKU, LTF, CISH, GJA1, LIF, DUSP10, SESN2, EGR1, DDIT3, PAK1IP1, TCIM, BCL6, DKK1, MAD2L2, ADRA2A, RGS2, OPTN, CASP8, FBXO7, YBX3, TNFAIP1, SELENOS, GHITM, PSMD10, MCL1, TLR4, ARRDC3, CGAS, CFLAR, CALR, DDIT4, CREB3L1, RNF115, MRAP2, MYC, INSIG1, UBE2D1, GPR155, BMP2* |
| GO:0098662 | Inorganic cation transmembrane transport | 122 | 1.421 | 2.22E-02 | 4.03E-02 | *CXCL9, CXCL11, CXCL10, SLC39A8, KCNJ2, KCNMA1, BPIFA1, DDIT3, SLC25A28, PLA2G1B, ADRA2A, GHITM, SLC31A1, KCNJ14, TMEM38B* |
| GO:0023057 | Negative regulation of signaling | 354 | 1.418 | 3.47E-03 | 8.17E-03 | *ISG15, CXCL8, DHX58, STAT1, SOCS3, SOCS1, PTGIR, ATF3, DUSP5, BPIFB1, TNFAIP3, THBS1, RGS16, FFAR2, IFI35, GDF15, STAT2, SLC39A8, IL10, NFKBIA, TSKU, LTF, CISH, GJA1, LIF, DUSP10, SESN2, EGR1, DDIT3, PAK1IP1, TCIM, BCL6, DKK1, MAD2L2, ADRA2A, RGS2, OPTN, CASP8, FBXO7, YBX3, TNFAIP1, SELENOS, GHITM, PSMD10, MCL1, TLR4, ARRDC3, CGAS, CFLAR, CALR, DDIT4, CREB3L1, RNF115, MRAP2, MYC, INSIG1, UBE2D1, GPR155, BMP2* |
| GO:0046907 | Intracellular transport | 362 | 1.417 | 3.75E-03 | 8.58E-03 | *IFNG, MX1, SNX10, STX11, EGR2, CD74, GOLT1A, NFKBIA, STAT3, TIMM8A, ARL5B, PLK3, RAB3D, RHOB, HSPA9, SEC23B, DERL3, KPNA2, TXNIP, SELENOS, IPO4, PREB, SRPRA, CALR, SEC24A, GOLT1B, RANBP1, INSIG1, SEC61A1, SEC61B, EIPR1, USO1, SEC22B, AP3S2, ZPR1, RAB1A, NDFIP2, NOL6, TMED10, NOP9, COPE, NUS1, SLC30A4, TRAM1, TMED9, XPO6, IFT20, COPB1, VCP, NPM1, KPNB1, ARL1, CHMP5, EIF6, ARF6, COPB2, DESI1, SDAD1, YIF1A, SAR1B, NXT1, NPC1, HYAL2, SRP54, COPZ1, SEC63, XPO5, PHAX,* |

| | | | | | | |
|---|---|---|---|---|---|---|
| | | | | | | *FMR1, HTATIP2, LDLR, PDCD10, SNF8, RAB9A, RAB20, AKIRIN2, RAB13, SQSTM1, SNX5, NUTF2, TOMM20, FYTTD1, TMED1, SSB, CHMP4A, TBC1D20, RINT1, STX17, ENY2, TNPO1, VTI1B, SEC24C, YIPF5, CLTC, APOD, AP3M1, ELAVL1, PHB2, RAE1, TMEM129, VPS26A, M6PR, NHERF1, MAP2K1, MFSD1, CHTOP, ERGIC2, AKAP5, RIC1, STX12, HIKESHI, TNPO3, SEC23A, STX5, IPO13, MED1, THOC3, RAB5A, COPA, AHCYL1, NEMF, TRIM27, SNX9, LMAN2, RANGRF, BLOC1S6, MTCH1, KPNA1, MAP2K2* |
| GO:0009968 | Negative regulation of signal transduction | 333 | 1.408 | 3.53E-03 | 8.20E-03 | *ISG15, CXCL8, DHX58, STAT1, SOCS3, SOCS1, PTGIR, ATF3, DUSP5, BPIFB1, TNFAIP3, THBS1, RGS16, IFI35, GDF15, STAT2, SLC39A8, IL10, NFKBIA, TSKU, LTF, CISH, LIF, DUSP10, SESN2, EGR1, DDIT3, PAK1IP1, TCIM, BCL6, DKK1, MAD2L2, RGS2, OPTN, CASP8, FBXO7, YBX3, TNFAIP1, SELENOS, GHITM, PSMD10, MCL1, TLR4, ARRDC3, CGAS, CFLAR, CALR, DDIT4, CREB3L1, RNF115, MRAP2, MYC, INSIG1, UBE2D1, GPR155, BMP2* |
| GO:0043408 | Regulation of MAPK cascade | 154 | 1.406 | 2.89E-02 | 4.95E-02 | *SAA3, CHI3L1, FAS, ATF3, DUSP5, CD40, GDF15, LIF, CCR1, DUSP10, GADD45G, CSF2, TRAF2, PLA2G1B, ADRA2A, LPAR3, PSMD10, TLR4, ADAM17, GPR37, TMEM106A, MYC, TIRAP, BMP2, GADD45B* |
| GO:0009890 | Negative regulation of biosynthetic process | 481 | 1.400 | 1.88E-03 | 5.01E-03 | *SAA3, IFNG, IRF1, CXCL8, SERPINE1, DHX58, TNFAIP3, THBS1, ZFP36, IL10, CEBPD, ATF5, TSKU, LTF, GJA1, CCR1, SESN2, FASLG, LRRC32, MOV10, RFX5, PLK3, ARID5A, EGR1, EIF4EBP1, TRAF2, DDIT3, SOX11, BCL6, MAD2L2, BAZ1A, CEBPB, YBX3, TXNIP, MFSD12, TGFB3, PSMD10, TLR4, MORC3, ORMDL1, CALR, ARG2, LYAR, CREB3L1, PNPT1, RNASEH2B, INSIG1, TIRAP, BMP2, EIF2S1, PELO, PRDM1, RBM8A, VDR, DLL4, ETF1, TP53, ELF4, NKRF, CRY1, NDFIP2, USP2, UXT, KAT2A, IGF2BP1, NKX3-1, AGO2, DXO, KLF10, NAT10, BASP1, HAVCR2, FBP1, EXOSC3, PDCD4, SUV39H2, PDE12, MTERF3, RIDA, CLDN5, PARP3, TBRG4, DNAJB1, DDX3X* |
| GO:2000145 | Regulation of cell motility | 225 | 1.400 | 1.10E-02 | 2.18E-02 | *CXCL10, MADCAM1, IFNG, SELE, CCL2, CCL4, CCL8, THBS1, CD40, NR4A3, CCL5, GDF15, TNFSF14, GJA1, CCR1, DUSP10, CXCL16, PHLDA2, GLUL, RHOB, BCL6, ADRA2A,* |

| | | | | | | |
|---|---|---|---|---|---|---|
| | | | | | | *CASP8, TLR4, KLRK1, CALR, KIT, RIPK3, TWIST2, BMP2, GTPBP4, LOC100516039, DLL4, GNA13* |
| GO:0010558 | Negative regulation of macromolecule biosynthetic process | 465 | 1.394 | 4.27E-03 | 9.45E-03 | *SAA3, IFNG, IRF1, CXCL8, SERPINE1, DHX58, TNFAIP3, THBS1, ZFP36, IL10, CEBPD, ATF5, TSKU, LTF, GJA1, CCR1, SESN2, FASLG, LRRC32, MOV10, RFX5, PLK3, ARID5A, EGR1, EIF4EBP1, TRAF2, DDIT3, SOX11, BCL6, MAD2L2, BAZ1A, CEBPB, YBX3, TXNIP, TGFB3, PSMD10, TLR4, MORC3, CALR, ARG2, LYAR, CREB3L1, PNPT1, RNASEH2B, TIRAP, BMP2, EIF2S1, PELO, PRDM1, RBM8A, VDR, DLL4, ETF1, TP53, ELF4, NKRF, CRY1, NDFIP2, USP2, UXT, KAT2A, IGF2BP1, NKX3-1, AGO2, DXO, KLF10, NAT10, BASP1, HAVCR2, FBP1, EXOSC3, PDCD4, SUV39H2, PDE12, MTERF3, RIDA, CLDN5, PARP3, TBRG4, DNAJB1, DDX3X* |
| GO:0043933 | Protein-containing complex organization | 432 | 1.391 | 4.68E-03 | 1.02E-02 | *IFNG, ISG15, SAMHD1, MMP1, LPL, TIFA, LIPG, SLA-DQB1, KCNJ2, FGB, CD74, SLA-DMB, MLKL, NDUFAF4, SLC1A5, SOAT1, TRAF2, HIGD1A, FARSA, MSRB1, BAZ1A, TICAM1, KIF18A, SNRPF, MYD88, TNFAIP1, NDUFAF5, PSMD10, HSPA8, TLR4, MRTO4, PREB, SNRPD1, CALR, DDIT4, CDC42EP3, RRS1, SLC31A1, NOP2, SLC25A33, TMEM126A, AVIL, PFDN6, TIRAP, GRWD1, CDC42EP2, EIF5A, PTGES3, ARPC5L, AQP5, ETF1, TP53, PSMG2, KCNK6, EIF3I, NOPCHAP1, PET100, UBLCP1, C7H6orf125, EIF3J, TWF1, UGDH, COX17, SNRPG, KCNA2, TMEM186, ME1, ABT1, RUBCN, TMEM39A, SHQ1, SNRPD3, DDX3X, COX16, LRRC8A, GEMIN6, VCP, DRG1, NPM1, MRPS7, CHMP5, ARF6, CAPZA2, HSD17B10, STRAP, SET, COA6, SAR1B, HSF1, RAP1B, NUFIP1, TAF9, CLNS1A, GRIA3, CAPZA1, PHF23, EIF3A, SNRPC, TMEM177, UQCC4, OGFOD1, USP39, FOXRED1, BCS1L, PPP2CA, PTK2B, SRSF1, PICALM, SIGMAR1, BRD2, PSMG4, OTUD6B, BCL10, SNRPE, CHMP4A, COPS7A, STX17, GTF2A2, ORC4, GTF2B, CHCHD10, TIMM21, OGT, TUBB1, PADI4, CYRIB, SUGT1, NDUFB1, NDUFB4, NSF* |
| GO:0032446 | Protein modification by | 213 | 1.384 | 9.49E-03 | 1.94E-02 | *ISG15, UBA7, SOCS3, SOCS1, RNF19B, TNFAIP3, EGR2, TRIM21, CISH, RNF122, EGR1, GNL3, TRAF2, TICAM1, FBXO7, TNFAIP1, PSMD10, WSB2, FBXO4, RNF115, UBE2D1, EIPR1, GTPBP4, CRY1, KLHL25, NDFIP2, UBE2D3, SOCS7, SPRY2, SIAH1, RNF138,* |

| | | | | | | |
|---|---|---|---|---|---|---|
| | small protein conjugation | | | | | *DDX3X, RNF114, VCP, UBE2B, DCUN1D5, PLAA, SIAH2, PHF23, FBXO33, UBA3, SQSTM1, RLIM, RNF5, BCL10, TRAF7, WBP1L, CUL5, DDRGK1, OGT, DCUN1D3, RNF14, PPIA, PRPF19, NFE2L2, TMEM129, UFC1, PCNP, FBXO9, N4BP1, UBE3A, DCAF11, SOCS5, TOPORS, ARIH2, MSL2* |
| GO:0006753 | Nucleoside phosphate metabolic process | 195 | 1.380 | 2.28E-02 | 4.09E-02 | *IDO1, SAMHD1, GFPT2, HK3, CTPS1, SLC25A22, NAMPT, NME1, GCK, TIGAR, UGP2, CNP, PGK1, DDIT4, GOT1, PRPS1, RNASEH2B, GFPT1, DCTD, HK2, PDE4B, MDH1, RFK, GNPNAT1, ACSL4, ATP5MC1, DCTPP1, LOC100515984, UGDH, FBP1, UAP1, ATP1B1, ME1, NT5C, FITM2, AK3, ENTPD1, NME2, ITPA, RRM2B, PDE4D, ADK, KMO, PDHX, G6PD, HINT1, PDE4A, IMPDH2, ACLY, TKT, PPP2CA, TPI1, NUDT11, DPYS, CTPS2, ATP6V1B2, FUT8, ATP5PB, ATP5F1A, ELOVL1, GAPDH, OGT, MPC2, ADSS2, PDHA1, ATP5MC3, ATP5MF, NT5C3A, PPCDC, MPI, CMPK1, PGAM1, DERA, HPRT1, MFN1, EPHA2, PGM3, PGD, GNPDA2, ATP5PD, UXS1, TGDS, COX11, BLOC1S6, GALT* |
| GO:0000165 | MAPK cascade | 169 | 1.379 | 2.24E-02 | 4.03E-02 | *SAA3, CHI3L1, FAS, ATF3, DUSP5, CD40, GDF15, ZFP36, LIF, CCR1, DUSP10, GADD45G, CSF2, TRAF2, PLA2G1B, ADRA2A, LPAR3, PSMD10, TLR4, ADAM17, GPR37, TMEM106A, MYC, TIRAP, BMP2, GADD45B* |
| GO:0006796 | Phosphate-containing compound metabolic process | 389 | 1.374 | 6.49E-03 | 1.36E-02 | *IFNG, IDO1, CHI3L1, SAMHD1, PPA1, GFPT2, THBS1, IL15, HK3, CCL5, STAT2, C3, CTPS1, SLC25A22, GK, LIF, NAMPT, NME1, STAT3, GCK, PLA2G1B, NEK6, TIGAR, UGP2, PLPPR2, CNP, PGK1, ORMDL1, DDIT4, GOT1, PRPS1, ABHD5, ITGA5, RNASEH2B, TMEM38B, GFPT1, DCTD, PDXK, HK2, PDE4B, MDH1, LPCAT3, JAK2, RFK, GNPNAT1, ACSL4, ATP5MC1, ITPKC, DCTPP1, LOC100515984, PISD, PLSCR3, UGDH, FBP1, UAP1, ATP1B1, NUS1, ME1, NT5C, FITM2, AK3, TAMM41, ENTPD1, NME2, CDS1, CHKA, PIGH, ITPA, RRM2B, AGPAT4, NRP1, PDE4D, ADK, PPP2R2A, SAMD8, KMO, PDHX, G6PD, HINT1, PDE4A, IMPDH2, ITPK1, ACLY, TKT, PPP2CA, TPI1, ATM, NUDT11, DPYS, PPP1CB, CTPS2, PLCH1, ORMDL3, ATP6V1B2, GNB3, BCL10, PIK3R5, DPM1, FUT8, SEMA4D, ATP5PB, ATP5F1A, ELOVL1, BPNT1, INPP1, GAPDH, SGMS1, OGT* |

| | | | | | |
|---|---|---|---|---|---|
| GO:0065003 | Protein-containing complex assembly | 381 | 1.366 | 6.07E-03 | 1.29E-02 | *IFNG, ISG15, SAMHD1, MMP1, TIFA, SLA-DQB1, KCNJ2, FGB, CD74, SLA-DMB, MLKL, NDUFAF4, SLC1A5, SOAT1, TRAF2, HIGD1A, FARSA, MSRB1, BAZ1A, TICAM1, SNRPF, MYD88, TNFAIP1, NDUFAF5, PSMD10, TLR4, MRTO4, PREB, SNRPD1, CALR, CDC42EP3, RRS1, SLC31A1, NOP2, SLC25A33, TMEM126A, AVIL, PFDN6, TIRAP, GRWD1, CDC42EP2, PTGES3, ARPC5L, AQP5, TP53, PSMG2, KCNK6, EIF3I, NOPCHAP1, PET100, UBLCP1, C7H6orf125, EIF3J, TWF1, UGDH, COX17, SNRPG, KCNA2, TMEM186, ME1, ABT1, SHQ1, SNRPD3, DDX3X, COX16, LRRC8A, GEMIN6, DRG1, NPM1, MRPS7, ARF6, CAPZA2, HSD17B10, STRAP, SET, COA6, SAR1B, HSF1, RAP1B, NUFIP1, TAF9, CLNS1A, GRIA3, CAPZA1, EIF3A, SNRPC, TMEM177, UQCC4, USP39, FOXRED1, BCS1L, PPP2CA, PTK2B, SRSF1, PICALM, SIGMAR1, BRD2, PSMG4, OTUD6B, BCL10, SNRPE, CHMP4A, COPS7A, GTF2A2, ORC4, GTF2B, CHCHD10, TIMM21, OGT, TUBB1, PADI4, CYRIB, SUGT1, NDUFB1, NDUFB4* |
| GO:0006793 | Phosphorus metabolic process | 390 | 1.359 | 1.40E-02 | 2.69E-02 | *IFNG, IDO1, CHI3L1, SAMHD1, PPA1, GFPT2, THBS1, IL15, HK3, CCL5, STAT2, C3, CTPS1, SLC25A22, GK, LIF, NAMPT, NME1, STAT3, GCK, PLA2G1B, NEK6, TIGAR, UGP2, PLPPR2, CNP, PGK1, ORMDL1, DDIT4, GOT1, PRPS1, ABHD5, ITGA5, RNASEH2B, TMEM38B, GFPT1, DCTD, PDXK, HK2, PDE4B, MDH1, LPCAT3, JAK2, RFK, GNPNAT1, ACSL4, ATP5MC1, ITPKC, DCTPP1, LOC100515984, PISD, PLSCR3, UGDH, FBP1, UAP1, ATP1B1, NUS1, ME1, NT5C, FITM2, AK3, TAMM41, ENTPD1, NME2, CDS1, CHKA, PIGH, ITPA, RRM2B, AGPAT4, NRP1, PDE4D, ADK, PPP2R2A, SAMD8, KMO, PDHX, G6PD, HINT1, PDE4A, IMPDH2, ITPK1, ACLY, TKT, PPP2CA, TPI1, ATM, NUDT11, DPYS, PPP1CB, CTPS2, PLCH1, ORMDL3, ATP6V1B2, GNB3, BCL10, PIK3R5, DPM1, FUT8, SEMA4D, ATP5PB, ATP5F1A, ELOVL1, BPNT1, INPP1, GAPDH, SGMS1, OGT* |
| GO:0045935 | Positive regulation of nucleobase-containing | 395 | 1.357 | 1.03E-02 | 2.07E-02 | *IRF1, IL1A, IRF7, SAMHD1, STAT1, FOSL1, JUNB, IRF9, IRF8, EGR2, PIM1, CD40, NFIL3, ZFP36, STAT2, STAT5A, IL10, NFKBIA, TFRC, LIF, STAT3, ARID3C, CEBPG, EGR1, GNL3, PLA2G1B, DDX21, SOX11, MAD2L2, TIGAR, BAZ1A, CEBPB, TGFB3, PLSCR1, TLR4, TNIP2, SERTAD1, TBX21, TNFRSF1A, LYAR, FBXO4, MYC, BMP2, WDR75, PTGES3, VDR, TP53,* |

| | | | | | | |
|---|---|---|---|---|---|---|
| | compound metabolic process | | | | | *JAK2, MORF4L2, IKBKB, SMAD1, SMAD3, INHBA, KAT2A, IGF2BP1, NKX3-1, ECD, AGO2, PPARGC1A, NCOA7, TBX18, TFE3, EXOSC3, IER5, CCT4, PDE12, POU2F2, RIDA, PARP3, PTGES2, DDX3X, TRA2B, NME2, NPM1, IER2* |
| GO:0016567 | Protein ubiquitination | 191 | 1.356 | 2.24E-02 | 4.03E-02 | *ISG15, UBA7, SOCS3, SOCS1, RNF19B, TNFAIP3, TRIM21, CISH, RNF122, TRAF2, TICAM1, FBXO7, TNFAIP1, PSMD10, WSB2, FBXO4, RNF115, UBE2D1, EIPR1, GTPBP4, CRY1, KLHL25, NDFIP2, UBE2D3, SOCS7, SPRY2, SIAH1, RNF138, DDX3X, RNF114, VCP, UBE2B, PLAA, SIAH2, PHF23, FBXO33, SQSTM1, RLIM, RNF5, BCL10, TRAF7, WBP1L, CUL5, OGT, RNF14, PPIA, PRPF19, NFE2L2, TMEM129, PCNP, FBXO9, N4BP1, UBE3A, DCAF11, SOCS5, TOPORS, ARIH2, MSL2* |
| GO:0140352 | Export from cell | 184 | 1.348 | 2.85E-02 | 4.91E-02 | *IFNG, IL1B, RSAD2, IL6, IL1A, GNAT1, CCL8, RHBDF2, STX11, FFAR2, CCL5, GJA1, LIF, CCR1, SOX11, ADRA2A, TGFB3, TLR4, KIT, TLR2, TMEM38B, BMP2, TP53, F2RL2, KCNK1, CRY1, RAB1A, TMED10, INHBA, NKX3-1, C7H6orf125, CADPS2, ATP1B1, KCNA2, C2CD2L, SLC30A4, TVP23B, LRRC8A* |
| GO:0009628 | Response to abiotic stimulus | 208 | 1.347 | 2.67E-02 | 4.67E-02 | *IRF1, CHI3L1, IL1A, GNAT1, MMP1, AQP3, TIMP1, CD40, NFKBIA, TNFSF14, GJA1, LIF, AEN, BHLHE40, PLK3, EGR1, TCIM, RHOB, TIGAR, HSPD1, CASP8, YBX3, TGFB3, TLR4, DDIT4, KIT, FBXO4, MYC, NABP1, TIPIN, BMP2, EIF2S1, GPR4, HSP90AA1, AQP5, TP53, TXN, CRY1, USP2, LIG4, DNAJA2, NKX3-1, IER5, EGLN3, SUV39H2, INIP, DNAJB1, DDX3X, IFT20, LRRC8A, VCP, NPM1, DNAJA3, ATP6V0A2, UBE2B, HSF1, KMO, HYAL2, CCND2* |
| GO:0045934 | Negative regulation of nucleobase-containing compound metabolic process | 285 | 1.344 | 1.67E-02 | 3.14E-02 | *SAA3, IFNG, IRF1, ZFP36, CEBPD, ATF5, FASLG, RFX5, PLK3, ARID5A, EGR1, TRAF2, DDIT3, SOX11, BCL6, MAD2L2, TIGAR, CEBPB, YBX3, TXNIP, PSMD10, CGAS, MORC3, CALR, DDIT4, LYAR, TIRAP, BMP2, GTPBP4, VDR, DLL4, TP53, NKRF, CRY1, USP2, UXT, KAT2A, IGF2BP1, NKX3-1, KLF10, NAT10, BASP1, TERF2IP, FBP1, PDCD4, SUV39H2, MTERF3, PARP3, DNAJB1* |
| GO:0051254 | Positive regulation of RNA metabolic process | 348 | 1.343 | 1.29E-02 | 2.53E-02 | *IRF1, IL1A, IRF7, SAMHD1, STAT1, FOSL1, JUNB, IRF9, IRF8, EGR2, PIM1, CD40, NFIL3, ZFP36, STAT2, STAT5A, IL10, NFKBIA, LIF, STAT3, ARID3C, EGR1, GNL3, PLA2G1B, DDX21, SOX11, MAD2L2, CEBPB, TGFB3, PLSCR1, TLR4, TNIP2, SERTAD1, TBX21,* |

| | | | | | | |
|---|---|---|---|---|---|---|
| | | | | | | *TNFRSF1A, LYAR, MYC, BMP2, WDR75, VDR, TP53, JAK2, IKBKB, SMAD1, SMAD3, INHBA, KAT2A, IGF2BP1, NKX3-1, ECD, AGO2, PPARGC1A, NCOA7, TBX18, TFE3, IER5, PDE12, POU2F2, RIDA, PTGES2, DDX3X, TRA2B, NME2, NPM1, IER2* |
| GO:0045893 | Positive regulation of DNA-templated transcription | 317 | 1.338 | 1.74E-02 | 3.23E-02 | *IRF1, IL1A, IRF7, SAMHD1, STAT1, FOSL1, JUNB, IRF9, IRF8, EGR2, PIM1, CD40, NFIL3, STAT2, STAT5A, IL10, NFKBIA, LIF, STAT3, ARID3C, EGR1, GNL3, PLA2G1B, DDX21, SOX11, MAD2L2, CEBPB, TGFB3, PLSCR1, TLR4, TNIP2, SERTAD1, TBX21, TNFRSF1A, LYAR, MYC, BMP2, WDR75, VDR, TP53, JAK2, IKBKB, SMAD1, SMAD3, INHBA, KAT2A, NKX3-1, ECD, PPARGC1A* |
| GO:1902680 | Positive regulation of RNA biosynthetic process | 317 | 1.338 | 1.74E-02 | 3.23E-02 | *IRF1, IL1A, IRF7, SAMHD1, STAT1, FOSL1, JUNB, IRF9, IRF8, EGR2, PIM1, CD40, NFIL3, STAT2, STAT5A, IL10, NFKBIA, LIF, STAT3, ARID3C, EGR1, GNL3, PLA2G1B, DDX21, SOX11, MAD2L2, CEBPB, TGFB3, PLSCR1, TLR4, TNIP2, SERTAD1, TBX21, TNFRSF1A, LYAR, MYC, BMP2, WDR75, VDR, TP53, JAK2, IKBKB, SMAD1, SMAD3, INHBA, KAT2A, NKX3-1, ECD, PPARGC1A* |
| GO:0007399 | Nervous system development | 436 | -1.219 | 2.71E-02 | 4.70E-02 | *BMP7, SRCIN1, LOC100525798, HEYL, KCNE1, CXCR4, EPHB3, SEMA6D, MTHFR, ID2, TYRO3, FUT1, SEMA3E, BMP4, NYAP1, VCL, CDC20, WNT11, BBS10, PKD2, CLDN11, FGF21, WNT2, BBS1, SOX9, TMEM107, DCX, EPOR, AMIGO1, SEMA3B, TPP1, NNAT, HES6, GATA3, CHODL, WNT7A, SLC1A1, SPOCK1, WNT9A, GPRASP3, SEMA6A, BEND6, NRROS, EPHA7, DCT, FOLR3, STMN1, NTF3, MYOC, NTRK3, CRB2, MAP1B, INPP5J, SEMA6C, RTN4R, CNTN1, RASGRF1, ICAM5, GBX1, HOXB8, HEY1, EFNB3, EPHA3, SRPX2, CNR1, WNT4, BDNF, MDGA1, ATCAY, NDP, MYCN, LRRN1, CNTN5, ZNF365, HHIP, CCR2, VXN, RXRG, P2RY12, SLC24A4, VEGFD, SFRP2, CX3CR1* |
| GO:0007167 | Enzyme-linked receptor protein signaling pathway | 232 | -1.296 | 2.72E-02 | 4.70E-02 | *BMP7, EGF, ADAMTSL2, EPHB3, FGF18, TYRO3, SMAD7, IGF2, BMP4, SAMD12, KDR, IGF1, FGF21, SOX9, CD109, GATA3, CAV2, EFNA2, ACVR2B, FRS3, SEMA6A, NRROS, EPHA7, FOLR3, HIP1R, TMEM204, NTF3, EREG, MYOC, NTRK3, CITED1, CRB2, BLK, INPP5K, MZB1, EFNB3, ANGPT1, APLN, EPHA3, AGTR2, GFRA3, BDNF, ANGPTL1, HPGD, WIF1, FGF1, NDP, HHIP, ASPN, CCN3, VEGFD, SFRP2* |

| | | | | | | |
|---|---|---|---|---|---|---|
| GO:0016055 | Wnt signaling pathway | 131 | -1.333 | 2.55E-02 | 4.51E-02 | *EDNRB, WNT2B, WLS, DIXDC1, SFRP1, LGR4, NDRG2, FGF9, SDC1, JUP, LGR5, KREMEN1, EDA, FZD2, APC2, SFRP5, WNT11, PKD2, WNT2, SOX9, CALCOCO1, NRARP, FAM53B, GATA3, WNT10B, TLE4, WNT7A, DKK3, WNT9A, WNT3A, FOLR3, MYOC, CPE, ALPK2, WNT4, WIF1, SFRP4, DKK2, NDP, SFRP2* |
| GO:0030182 | Neuron differentiation | 255 | -1.336 | 1.04E-02 | 2.09E-02 | *ITM2C, ACP4, ARHGEF25, UCHL1, DMD, RTN4RL1, ID4, SYT3, KREMEN1, RND2, BMP7, SRCIN1, LOC100525798, HEYL, EPHB3, SEMA6D, ID2, SEMA3E, BMP4, NYAP1, VCL, WNT11, BBS10, WNT2, BBS1, SOX9, DCX, SEMA3B, CHODL, WNT7A, SPOCK1, WNT9A, GPRASP3, SEMA6A, BEND6, EPHA7, FOLR3, STMN1, NTF3, MYOC, NTRK3, CRB2, MAP1B, INPP5J, SEMA6C, RTN4R, CNTN1, RASGRF1, GBX1, HEY1, EFNB3, EPHA3, CNR1, WNT4, BDNF, ATCAY, NDP, MYCN, CNTN5, ZNF365, VEGFD, SFRP2* |
| GO:0007017 | Microtubule-based process | 245 | -1.338 | 1.31E-02 | 2.54E-02 | *SPDL1, HOOK2, PARD6G, DNAL4, EFHB, CCNB2, KIF19, TEKT1, CFAP20, SPRY1, PIERCE1, BICDL2, TACC3, LCA5, FES, SPC25, MGARP, CCSAP, ATAT1, TUBB2B, RIBC1, EFHC1, SPACA9, KIF20B, TUBE1, CDK1, CFAP206, MID1, SGO1, CFAP141, CCNB1, FOXJ1, UCHL1, CGN, NUF2, ODAD3, WDR62, KIF2C, ARMC3, BIRC5, SPEF2, NCKAP5L, TEKT2, KLC4, CENPH, CFAP58, CFAP91, CFAP276, KIF11, MAP10, CEP250, DYNLT5, CCNF, BICD1, IFT46, DYNLT4, KIFC1, AURKB, BBS1, DCX, KIF22, KIF18B, KIF20A, CCDC68, FAM110A, GEN1, CENPA, IQUB, SLC22A16, WRAP73, SKA1, TUBB4A, KIF24, CFAP97D1, CATSPER4, DNAI2, MEIG1, STMN1, TPPP, DYNC2I2, TPPP3, TUB, MAP1B, RGS14, ODAD1, EPHA3, DEFB1* |
| GO:0007507 | Heart development | 117 | -1.375 | 2.08E-02 | 3.81E-02 | *GJA5, FGF9, BMPR2, FOXJ1, NOX4, CCN1, BMP7, HEYL, HSPB7, NPY1R, SMAD7, BMP4, MESP2, KDR, WNT11, PKD2, WNT2, SOX9, EPOR, GATA3, ACVR2B, COL14A1, CCM2L, FOLR3, GATA5, NTRK3, PDLIM3, CPE, APLN, ALPK2, WIF1, ADRA1A, SFRP2* |
| GO:0007268 | Chemical synaptic transmission | 115 | -1.388 | 2.63E-02 | 4.62E-02 | *WNT7A, SLC1A1, EPHA7, KCNB1, NTF3, SLC6A2, TMOD2, RGS14, RASGRF1, EFNB3, JPH4, P2RX7, BDNF, CHRM2, GRIN3B, CCR2, SYT12, P2RX3, CX3CR1* |
| GO:0098916 | Anterograde trans-synaptic signaling | 115 | -1.388 | 2.63E-02 | 4.62E-02 | *WNT7A, SLC1A1, EPHA7, KCNB1, NTF3, SLC6A2, TMOD2, RGS14, RASGRF1, EFNB3, JPH4, P2RX7, BDNF, CHRM2, GRIN3B, CCR2, SYT12, P2RX3, CX3CR1* |

| | | | | | | |
|---|---|---|---|---|---|---|
| GO:0031344 | Regulation of cell projection organization | 119 | -1.406 | 2.54E-02 | 4.51E-02 | *TACSTD2, SFRP1, PALM, NR2E1, PODXL, BMPR2, NEDD9, ACP4, RTN4RL1, KREMEN1, RND2, BMP7, SRCIN1, EPHB3, ENPP2, F2RL1, CHODL, WNT7A, SPOCK1, WRAP73, GPRASP3, EPHA7, PLD1, NTRK3, INPP5J, RTN4R, CNTN1, EFNB3, EPHA3, P2RX7, BDNF, ZNF365, SFRP2* |
| GO:0048699 | Generation of neurons | 268 | -1.407 | 2.70E-03 | 6.66E-03 | *VEGFA, ITM2C, ACSL6, ACP4, ARHGEF25, UCHL1, DMD, RTN4RL1, ID4, SYT3, KREMEN1, RND2, BMP7, SRCIN1, LOC100525798, HEYL, EPHB3, SEMA6D, ID2, TYRO3, SEMA3E, BMP4, NYAP1, VCL, WNT11, BBS10, WNT2, BBS1, SOX9, DCX, SEMA3B, CHODL, WNT7A, SPOCK1, WNT9A, GPRASP3, SEMA6A, BEND6, EPHA7, DCT, FOLR3, STMN1, NTF3, MYOC, NTRK3, CRB2, MAP1B, INPP5J, SEMA6C, RTN4R, CNTN1, RASGRF1, GBX1, HEY1, EFNB3, EPHA3, CNR1, WNT4, BDNF, MDGA1, ATCAY, NDP, MYCN, CNTN5, ZNF365, HHIP, VEGFD, SFRP2* |
| GO:0099537 | Trans-synaptic signaling | 116 | -1.429 | 1.32E-02 | 2.54E-02 | *WNT7A, SLC1A1, EPHA7, KCNB1, NTF3, SLC6A2, TMOD2, RGS14, RASGRF1, EFNB3, JPH4, CNR1, P2RX7, BDNF, CHRM2, GRIN3B, CCR2, SYT12, P2RX3, CX3CR1* |
| GO:0120035 | Regulation of plasma membrane bounded cell projection organization | 118 | -1.429 | 2.20E-02 | 4.02E-02 | *TACSTD2, SFRP1, PALM, NR2E1, PODXL, BMPR2, NEDD9, ACP4, RTN4RL1, KREMEN1, RND2, BMP7, SRCIN1, EPHB3, ENPP2, F2RL1, CHODL, WNT7A, SPOCK1, WRAP73, GPRASP3, EPHA7, PLD1, NTRK3, INPP5J, RTN4R, CNTN1, EFNB3, EPHA3, P2RX7, BDNF, ZNF365, SFRP2* |
| GO:0099536 | Synaptic signaling | 122 | -1.444 | 1.32E-02 | 2.54E-02 | *WNT7A, SLC1A1, EPHA7, KCNB1, NTF3, RGS10, SLC6A2, TMOD2, RGS14, RASGRF1, EFNB3, JPH4, CNR1, P2RX7, BDNF, CHRM2, GRIN3B, CCR2, SYT12, P2RX3, CX3CR1* |
| GO:0034330 | Cell junction organization | 141 | -1.531 | 3.78E-03 | 8.58E-03 | *SPARC, OCLN, S100A10, SRCIN1, EPHB3, SMAD7, SEMA3E, CLDN18, KDR, VCL, CDC20, WNT11, CLDN11, DCX, F2RL1, C1QC, CLDN6, AMIGO1, CLDN22, WNT7A, SLC1A1, GPRASP3, EPHA7, TRPV4, MYOC, MAP1B, RTN4R, ICAM5, EPHA3, SRPX2, BDNF, MDGA1, LRRN1, CNTN5, ZNF365, CX3CR1* |
| GO:0001501 | Skeletal system development | 117 | -1.534 | 3.78E-03 | 8.58E-03 | *TBX3, PDGFA, HOXA3, WNT2B, HOXA2, SFRP1, HOXA9, FGF9, BMPR2, RFLNA, SMPD3, BMPR1B, CCN1, HOXC4, BMP7, BGN, FGF18, RFLNB, SMAD7, BMP4, KDR, WNT11, BBS1,* |

| | | | | | | |
|---|---|---|---|---|---|---|
| | | | | | | *SOX9, TMEM107, NFATC2, CHAD, WNT7A, ACVR2B, MMP16, OGN, HOXC6, PAPPA2, TRPV4, MYOC, LEPR, OSR2, HOXB8, P2RX7, HOXB7, MYCN, HHIP, CCN3, LRRC17, SFRP2* |
| GO:0120036 | Plasma membrane bounded cell projection organization | 296 | -1.622 | 4.20E-05 | 1.63E-04 | *KIF19, TTC8, CAMK1D, PIK3R1, TEKT1, CFAP20, PIERCE1, FAM161A, BTBD3, ARL6, LCA5, FGD1, FES, SEMA4F, RHOD, NFIB, SPATA6, SPACA9, FSCN1, TPBG, SMO, CFAP206, HOXA2, TACSTD2, UBXN10, FLOT1, SFRP1, PALM, EMP3, NR2E1, PODXL, EHD2, B4GALT6, BMPR2, NEDD9, FOXJ1, ACP4, ARHGEF25, AIF1, UCHL1, ODAD3, CCDC113, RTN4RL1, SYT3, KREMEN1, RND2, SPEF2, BMP7, SRCIN1, TEKT2, EPHB3, SEMA6D, TTC39C, CFAP58, ENPP2, CFAP91, CEP250, SEMA3E, NYAP1, IFT46, VCL, BBS10, PKD2, BBS1, TMEM107, DCX, F2RL1, SEMA3B, IQUB, CHODL, WNT7A, SPOCK1, WRAP73, GPRASP3, SEMA6A, CFAP97D1, EPHA7, DNAI2, FOLR3, MEIG1, CCNO, PLD1, STMN1, NTF3, DYNC2I2, MYOC, NTRK3, TUB, INPP5K, MAP1B, GMNC, INPP5J, SEMA6C, RTN4R, CNTN1, RASGRF1, GBX1, EFNB3, ODAD1, EPHA3, CNR1, P2RX7, BDNF, ATCAY, NDP, CNTN5, ZNF365, P2RY12, SFRP2* |
| GO:0030030 | Cell projection organization | 311 | -1.624 | 1.95E-05 | 8.05E-05 | *KIF19, TTC8, CAMK1D, PIK3R1, TEKT1, CFAP20, PIERCE1, FAM161A, BTBD3, ARL6, LCA5, FGD1, FES, SEMA4F, RHOD, NFIB, TCTN2, SPATA6, SPACA9, FSCN1, TPBG, SMO, PDGFA, CFAP206, HOXA2, TACSTD2, UBXN10, FLOT1, SFRP1, PALM, EMP3, NR2E1, PODXL, EHD2, B4GALT6, BMPR2, NEDD9, FOXJ1, ACP4, ARHGEF25, AIF1, UCHL1, ODAD3, CCDC113, RTN4RL1, SYT3, KREMEN1, RND2, SPEF2, BMP7, SRCIN1, TEKT2, EPHB3, SEMA6D, TTC39C, CFAP58, ENPP2, CFAP91, CEP250, SEMA3E, NYAP1, IFT46, TCTN1, VCL, BBS10, PKD2, BBS1, TMEM107, DCX, F2RL1, SEMA3B, PARVG, IQUB, CHODL, WNT7A, SPOCK1, MKS1, WRAP73, GPRASP3, SEMA6A, CFAP97D1, EPHA7, DNAI2, FOLR3, MEIG1, CCNO, PLD1, STMN1, NTF3, DYNC2I2, MYOC, NTRK3, TUB, INPP5K, MAP1B, GMNC, INPP5J, SEMA6C, RTN4R, CNTN1, RASGRF1, GBX1, EFNB3, ODAD1, EPHA3, CNR1, P2RX7, BDNF, ATCAY, NDP, CNTN5, ZNF365, P2RY12, SFRP2* |
| GO:0031175 | Neuron projection development | 167 | -1.632 | 4.38E-04 | 1.39E-03 | *ACP4, ARHGEF25, UCHL1, RTN4RL1, SYT3, KREMEN1, RND2, BMP7, SRCIN1, EPHB3, SEMA6D, SEMA3E, NYAP1, VCL, BBS1, DCX, SEMA3B, CHODL, WNT7A, SPOCK1,* |

| | | | | | | |
|---|---|---|---|---|---|---|
| | | | | | | *GPRASP3, SEMA6A, EPHA7, FOLR3, STMN1, NTF3, MYOC, NTRK3, MAP1B, INPP5J, SEMA6C, RTN4R, CNTN1, RASGRF1, GBX1, EFNB3, EPHA3, CNR1, BDNF, ATCAY, NDP, CNTN5, ZNF365, SFRP2* |
| GO:0048858 | Cell projection morphogenesis | 113 | -1.695 | 6.77E-04 | 2.04E-03 | *SEMA4F, NFIB, TPBG, SMO, HOXA2, FLOT1, NR2E1, B4GALT6, BMPR2, ARHGEF25, UCHL1, SYT3, RND2, BMP7, SRCIN1, EPHB3, SEMA6D, ENPP2, SEMA3E, NYAP1, VCL, DCX, SEMA3B, CHODL, WNT7A, GPRASP3, SEMA6A, EPHA7, NTF3, MAP1B, SEMA6C, RTN4R, CNTN1, GBX1, EFNB3, EPHA3, BDNF, NDP, CNTN5, ZNF365* |
| GO:0120039 | Plasma membrane bounded cell projection morphogenesis | 113 | -1.695 | 6.77E-04 | 2.04E-03 | *SEMA4F, NFIB, TPBG, SMO, HOXA2, FLOT1, NR2E1, B4GALT6, BMPR2, ARHGEF25, UCHL1, SYT3, RND2, BMP7, SRCIN1, EPHB3, SEMA6D, ENPP2, SEMA3E, NYAP1, VCL, DCX, SEMA3B, CHODL, WNT7A, GPRASP3, SEMA6A, EPHA7, NTF3, MAP1B, SEMA6C, RTN4R, CNTN1, GBX1, EFNB3, EPHA3, BDNF, NDP, CNTN5, ZNF365* |
| GO:0048812 | Neuron projection morphogenesis | 109 | -1.708 | 3.77E-04 | 1.22E-03 | *SEMA4F, NFIB, TPBG, SMO, HOXA2, FLOT1, NR2E1, B4GALT6, BMPR2, ARHGEF25, UCHL1, SYT3, RND2, BMP7, SRCIN1, EPHB3, SEMA6D, SEMA3E, NYAP1, VCL, DCX, SEMA3B, CHODL, WNT7A, GPRASP3, SEMA6A, EPHA7, NTF3, MAP1B, SEMA6C, RTN4R, CNTN1, GBX1, EFNB3, EPHA3, BDNF, NDP, CNTN5, ZNF365* |

Abbreviations: GO:BP: Biological Processes in Gene Ontology; FDR: False discovery rate; MAPK: Mitogen-activated protein kinase; NES: Normalized enrichment score.

81

**Supplementary Table S5. Significantly enriched pathways from the GSEA result of the KEGG database, Low-glucose infection versus Low-glucose control.** Abbreviations: GSEA: Gene set enrichment analysis; KEGG: Kyoto Encyclopedia of Genes and Genomes.

| KEGG ID | Description | Set Size | NES | P-value | FDR | Core enrichment gene |
|---|---|---|---|---|---|---|
| ssc05168 | Herpes simplex virus 1 infection | 140 | 2.719 | 3.60E-24 | 1.24E-21 | *IFNG, OAS1, RIGI, OAS2, TAP1, IRF7, STAT1, SLA-DOA, BST2, TAP2, CCL2, SLA-3, SLA-DRA, IFIH1, EIF2AK2, IRF9, FASLG, SOCS3, FAS, SLA-DQA1, CD74, B2M, SLA-DOB, BIRC3, IL1B, SLA-DQB1, RNASEL, SLA-DMB, TAPBP, STAT2, SLA-DMA, PML, LOC100517129, SLA-7, CCL5, TAPBPL, C3, SLA-DRB1, IL6, STING1, IKBKE, BAK1, IFNAR2, PDIA3, CGAS, SLA-8, CASP3, NFKBIA, TNFRSF1A, CALR, DAXX, ITGA5, TRAF3, MYD88, TNFSF14, TLR2, ZC3HAV1, TLR3, TRAF2, CASP8, TICAM1, IKBKB, JAK2, CYCS, NFKB1* |
| ssc05164 | Influenza A | 127 | 2.664 | 8.93E-21 | 1.54E-18 | *CXCL10, RSAD2, IFNG, MX1, OAS1, RIGI, OAS2, IRF7, STAT1, SLA-DOA, CCL2, CXCL8, SLA-DRA, TNFSF10, IFIH1, EIF2AK2, IRF9, FASLG, SOCS3, FAS, SLA-DQA1, SLA-DOB, IL1B, SLA-DQB1, RNASEL, SLA-DMB, STAT2, SLA-DMA, ADAR, CASP1, PML, CCL5, IL1A, LOC100624226, SLA-DRB1, TRIM25, IL6, IKBKE, NFKBIB, BAK1, TMPRSS4, IFNAR2, CASP3, NFKBIA, TNFRSF1A, TRAF3, MYD88, TLR3, CALCOCO2, CASP8, XPO1, TICAM1, IKBKB, JAK2, CYCS, ICAM1, NFKB1, TLR4, DNAJC3* |
| ssc05169 | Epstein-Barr virus infection | 168 | 2.546 | 1.51E-18 | 1.74E-16 | *CXCL10, ISG15, OAS1, RIGI, OAS2, TAP1, IRF7, STAT1, SLA-DOA, TAP2, SLA-3, SLA-DRA, EIF2AK2, IRF9, FAS, SLA-DQA1, B2M, SLA-DOB, SLA-DQB1, SLA-DMB, TAPBP, STAT2, GADD45G, TNFAIP3, RELB, SLA-DMA, CDKN1A, SLA-7, CD40, TAPBPL, SLA-DRB1, IL6, IKBKE, GADD45A, NFKBIB, BAK1, STAT3, NFKB2, IFNAR2, NFKBIE, PDIA3, SLA-8, JAK3, CASP3, NFKBIA, CALR, TRAF3, MYC, MYD88, TLR2, TRAF2, CASP8, E2F2, IKBKB, CYCS, RIPK1, RUNX3, ICAM1, NFKB1, GADD45B, HES1, ENTPD1, BLNK, CCND3, PSMD3* |
| ssc04668 | TNF signaling pathway | 104 | 2.544 | 2.92E-15 | 2.52E-13 | *CXCL10, SELE, IRF1, AMCF-II, CXCL2, CCL2, RHBDF2, SOCS3, FAS, CX3CL1, FOS, BIRC3, IL1B, BCL3, CASP10, TNFAIP3, JUNB, MAP3K8, VCAM1, CCL5, CASP7, PTGS2, NOD2, CSF2,* |

| | | | | | | |
|---|---|---|---|---|---|---|
| | | | | | | *CSF1, IL6, IL15, LIF, MLKL, PGAM5, CFLAR, CASP3, NFKBIA, TNFRSF1A, CEBPB, ADAM17, TRAF3, MMP9, RIPK3, TNFRSF1B, CREB3L1, TRAF2, CASP8, IKBKB, RIPK1, ICAM1, NFKB1, FRMD8* |
| ssc04620 | Toll-like receptor signaling pathway | 87 | 2.481 | 2.85E-13 | 1.64E-11 | *CXCL9, CXCL11, CXCL10, CCL4, IRF7, STAT1, LBP, CXCL8, IRF9, CCL3L1, FOS, IL1B, STAT2, MAP3K8, CCL5, CD40, SPP1, IL6, IKBKE, TICAM2, IFNAR2, NFKBIA, TRAF3, MYD88, TLR2, TLR3, CD14, CASP8, TICAM1, IKBKB, RIPK1, NFKB1, TLR4* |
| ssc04612 | Antigen processing and presentation | 47 | 2.454 | 1.09E-10 | 3.40E-09 | *IFNG, TAP1, SLA-DOA, TAP2, SLA-3, SLA-DRA, SLA-DQA1, CD74, PSME2, CTSL, B2M, SLA-DOB, SLA-DQB1, SLA-DMB, TAPBP, PSME1, SLA-DMA, IFI30, SLA-7, TAPBPL, SLA-DRB1, HSPA5, RFX5, PDIA3, SLA-8, CALR, HSPA8* |
| ssc04657 | IL-17 signaling pathway | 70 | 2.409 | 6.91E-11 | 2.38E-09 | *CXCL10, IFNG, AMCF-II, CCL11, CXCL2, CCL2, CXCL8, S100A8, S100A9, FOS, IL1B, TNFAIP3, FOSL1, MUC5AC, CSF3, PTGS2, MMP1, CSF2, IL6, IKBKE, HSP90B1, FOSB, TRAF3IP2, CASP3, NFKBIA, CEBPB, TRAF3, MMP9, TRAF2, CASP8, IKBKB, NFKB1, IL17B* |
| ssc04622 | RIG-I-like receptor signaling pathway | 54 | 2.391 | 1.37E-10 | 3.63E-09 | *CXCL10, ISG15, DHX58, RIGI, IRF7, CXCL8, IFIH1, ZNFX1, CASP10, ADAR, TRIM25, STING1, IKBKE, NFKBIB, NFKBIA, TRAF3, LOC100624590, TRAF2, CASP8, IKBKB, RIPK1, NFKB1* |
| ssc04060 | Cytokine-cytokine receptor interaction | 170 | 2.365 | 4.67E-14 | 3.23E-12 | *CXCL9, CXCL11, CXCL10, IFNG, CCL26, CXCL13, CCL19, CCL4, AMCF-II, CCL11, IL27, CXCL2, CCL2, CXCL8, TNFSF10, TNFSF13B, FASLG, CCL3L1, FAS, CX3CL1, IL1B, XCL1, ACKR4, IL4R, CSF3R, IL2RB, LOC110260995, CXCR6, CCL5, IL1A, LOC100624226, CD40, CSF3, CCL8, CCL28, ACKR3, CSF2, IL1RAP, CSF1, IL1R1, IL1R2, IL6, CCR1, IL15, OSMR, CXCL16, LIF, IFNAR2, CSF2RB, GDF15, ACVR1B, CNTFR, IL6R, TNFRSF1A, IL31RA, IL18RAP, TNFRSF11B, TNFSF14, TNFRSF1B, RELT, TNFSF13, IL2RA, LOC397451, IL2RG, IL10RB, IL17B, CXCR2, TNFSF9, IL21R, RELL1, IL10* |
| ssc04621 | NOD-like receptor signaling pathway | 127 | 2.338 | 2.47E-11 | 1.06E-09 | *GBP1, OAS1, GBP2, OAS2, IRF7, STAT1, CXCL2, CCL2, CXCL8, IRF9, LOC100522887, BIRC3, IL1B, RNASEL, STAT2, TNFAIP3, CASP1, CCL5, NAMPT, NOD2, TRPM2, IL6, STING1, IKBKE, NFKBIB, IFNAR2, TXNIP, GSDMD, NFKBIA, TRAF3, MYD88, RIPK3, GP91-PHOX, TRAF2, CASP8, TICAM1, IKBKB, RIPK1, NFKB1, RIPK2, TLR4, TRPV2, GABARAPL1, JUN, TAB3, NLRP3* |

| | | | | | | |
|---|---|---|---|---|---|---|
| ssc05162 | Measles | 108 | 2.337 | 2.11E-11 | 1.04E-09 | *MX1, OAS1, RIGI, OAS2, IRF7, STAT1, IFIH1, EIF2AK2, IRF9, FASLG, FAS, FOS, IL1B, STAT2, TNFAIP3, IL2RB, ADAR, IL1A, SLAMF1, STAT5A, IL6, IKBKE, NFKBIB, BAK1, STAT3, IFNAR2, FCGR2B, JAK3, CASP3, NFKBIA, TRAF3, MYD88, HSPA8, TLR2, CASP8, IKBKB, IL2RA, CYCS, IL2RG, NFKB1, TLR4* |
| ssc05323 | Rheumatoid arthritis | 65 | 2.311 | 6.85E-09 | 1.48E-07 | *IFNG, AMCF-II, SLA-DOA, CXCL2, CCL2, CXCL8, SLA-DRA, TNFSF13B, CCL3L1, SLA-DQA1, FOS, CTSL, SLA-DOB, IL1B, SLA-DQB1, SLA-DMB, CTLA4, SLA-DMA, TCIRG1, CCL5, IL1A, SLA-DRB1, MMP1, CSF2, ATP6V1C2, CSF1, IL6, IL15* |
| ssc04061 | Viral protein interaction with cytokine and cytokine receptor | 60 | 2.285 | 9.11E-09 | 1.75E-07 | *CXCL9, CXCL11, CXCL10, CCL26, CXCL13, CCL19, CCL4, AMCF-II, CCL11, CXCL2, CCL2, CXCL8, TNFSF10, CCL3L1, CX3CL1, XCL1, ACKR4, IL2RB, CCL5, LOC100624226, CCL8, CCL28, ACKR3, CSF1, IL6, CCR1, IL6R, TNFRSF1A, IL18RAP, TNFSF14, TNFRSF1B, IL2RA, IL2RG, IL10RB, CXCR2* |
| ssc04064 | NF-kappa B signaling pathway | 87 | 2.276 | 3.63E-09 | 8.36E-08 | *CCL19, RIGI, CCL4, LBP, CXCL2, CXCL8, TNFSF13B, BIRC3, IL1B, GADD45G, TNFAIP3, RELB, VCAM1, CD40, PTGS2, TRIM25, IL1R1, GADD45A, TICAM2, NFKB2, CFLAR, NFKBIA, TNFRSF1A, BCL2A1, TRAF3, MYD88, TNFSF14, CD14, TRAF2, TICAM1, IKBKB, RIPK1, ICAM1, NFKB1, GADD45B, TLR4, ERC1, BLNK, LTBR, CXCL12, TAB3* |
| ssc05160 | Hepatitis C | 120 | 2.265 | 1.21E-10 | 3.47E-09 | *CXCL10, RSAD2, IFNG, MX1, OAS1, IFIT1, RIGI, OAS2, IRF7, STAT1, EIF2AK2, IRF9, FASLG, SOCS3, FAS, RNASEL, STAT2, CDKN1A, IKBKE, BAK1, STAT3, IFNAR2, CFLAR, CASP3, NFKBIA, CLDN5, TNFRSF1A, TRAF3, MYC, TLR3, TRAF2, CASP8, TICAM1, E2F2, IKBKB, CYCS, RIPK1, NFKB1* |
| ssc04623 | Cytosolic DNA-sensing pathway | 59 | 2.265 | 1.11E-07 | 1.67E-06 | *CXCL10, RIGI, ZBP1, CCL4, IRF7, SAMHD1, IL1B, ADAR, CASP1, CCL5, CASP7, IL6, STING1, IKBKE, NFKBIB, TREX1, MLKL, CGAS, GSDMD, CASP3, NFKBIA, RIPK3, CASP8, IKBKB, RIPK1, NFKB1* |
| ssc05171 | Coronavirus disease - COVID-19 | 181 | 2.229 | 5.04E-11 | 1.93E-09 | *CXCL10, ISG15, MX1, OAS1, RIGI, OAS2, STAT1, CCL2, CXCL8, IFIH1, EIF2AK2, C2, IRF9, FOS, IL1B, C4A, STAT2, ADAR, CASP1, CFB, CSF3, F13A1, C3, MMP1, CSF2, IL6, STING1, IKBKE, C9, NFKBIB, RPL13A, STAT3, IFNAR2, LOC100515049, CGAS, NFKBIA, IL6R, TNFRSF1A, ADAM17, TRAF3, MYD88, SELP, TLR2, TLR3* |

| | | | | | | |
|---|---|---|---|---|---|---|
| ssc05145 | Toxoplasmosis | 94 | 2.206 | 2.53E-08 | 4.16E-07 | *IFNG, STAT1, SLA-DOA, SLA-DRA, SOCS1, SLA-DQA1, SLA-DOB, BIRC3, SLA-DQB1, SLA-DMB, PPIF, LAMB4, SLA-DMA, CD40, SLA-DRB1, NFKBIB, STAT3, CASP3, ITGA6, NFKBIA, TNFRSF1A, MYD88, HSPA8, TLR2, NOS2, ALOX5, CASP8, IKBKB, JAK2, CYCS, IL10RB, NFKB1, TLR4, IL10* |
| ssc04217 | Necroptosis | 109 | 2.198 | 2.15E-08 | 3.71E-07 | *IFNG, ZBP1, STAT1, TNFSF10, EIF2AK2, IRF9, FASLG, FAS, BIRC3, IL1B, STAT2, TNFAIP3, CASP1, IL1A, LOC100624226, STAT5A, GLUL, TICAM2, STAT3, IFNAR2, MLKL, PGAM5, CFLAR, CHMP7, JAK3, TNFRSF1A, TLR3, RIPK3, FTH1, GP91-PHOX, TRAF2, CASP8, TICAM1, JAK2, RIPK1* |
| ssc04630 | JAK-STAT signaling pathway | 110 | 2.195 | 8.96E-09 | 1.75E-07 | *IFNG, STAT1, IL27, SOCS1, IRF9, SOCS3, STAT2, PIM1, IL4R, CSF3R, IL2RB, CDKN1A, CSF3, STAT5A, CSF2, IL6, IL15, OSMR, STAT3, LIF, IFNAR2, CISH, CSF2RB, JAK3, CNTFR, IL6R, IL31RA, MYC, PDGFRA, MCL1, PTPN2, IL2RA, LOC397451, JAK2, PDGFRB, IL2RG, IL10RB, IL21R, IL10, CCND3, SOCS7, CSF2RA* |
| ssc05140 | Leishmaniasis | 59 | 2.186 | 1.25E-06 | 1.56E-05 | *IFNG, STAT1, SLA-DOA, SLA-DRA, SLA-DQA1, FOS, SLA-DOB, IL1B, SLA-DQB1, SLA-DMB, SLA-DMA, IL1A, C3, SLA-DRB1, PTGS2, NFKBIB, FCGR1A, NFKBIA, MYD88, TLR2, NOS2, GP91-PHOX, NCF1, JAK2, NFKB1, TLR4, IL10* |
| ssc04659 | Th17 cell differentiation | 89 | 2.184 | 9.42E-08 | 1.48E-06 | *IFNG, STAT1, SLA-DOA, FOXS1, IL27, SLA-DRA, SLA-DQA1, FOS, SLA-DOB, IL1B, SLA-DQB1, SLA-DMB, SLA-DMA, IL4R, IL2RB, SLA-DRB1, STAT5A, IL1RAP, IL1R1, IL6, NFKBIB, STAT3, NFKBIE, JAK3, NFKBIA, IL6R, TBX21, IKBKB, IL2RA, JAK2, IL2RG, NFKB1, IL21R, RXRA* |
| ssc05321 | Inflammatory bowel disease | 45 | 2.181 | 1.27E-06 | 1.56E-05 | *IFNG, STAT1, SLA-DOA, SLA-DRA, SLA-DQA1, SLA-DOB, IL1B, SLA-DQB1, SLA-DMB, SLA-DMA, IL4R, IL1A, SLA-DRB1, NOD2, IL6, STAT3, IL18RAP, TBX21, TLR2, IL2RG, NFKB1, TLR4, IL21R, IL10, JUN, IL12RB2, TLR5* |
| ssc05152 | Tuberculosis | 137 | 2.174 | 1.47E-08 | 2.66E-07 | *IFNG, LOC110258711, CLEC4E, STAT1, LBP, SLA-DOA, SLA-DRA, SLA-DQA1, CD74, SLA-DOB, IL1B, SLA-DQB1, SLA-DMB, CYP27B1, CASP10, SLA-DMA, TCIRG1, IL1A, C3, SLA-DRB1, NOD2, IL6, RFX5, FCGR2B, CASP3, FCGR1A, TNFRSF1A, CEBPB, MYD88, PLK3, TLR2, CD14, NOS2, CTSS, CASP8, CEBPG, CLEC7A, JAK2, CYCS, MRC1, IL10RB, NFKB1, HSPA9, RIPK2, TLR4, IL10, IRAK2* |

| | | | | | | |
|---|---|---|---|---|---|---|
| ssc04062 | Chemokine signaling pathway | 158 | 2.152 | 1.38E-09 | 3.40E-08 | *CXCL9, CXCL11, CXCL10, CCL26, CXCL13, CCL19, CCL4, AMCF-II, STAT1, CCL11, CXCL2, CCL2, CXCL8, CCL3L1, CX3CL1, XCL1, STAT2, CXCR6, CCL5, CCL8, CCL28, CCR1, NFKBIB, STAT3, CXCL16, JAK3, NFKBIA, LOC100621431, NCF1, IKBKB, HCK, JAK2, NFKB1, FGR, GNG10, ADCY3, CXCR2* |
| ssc03050 | Proteasome | 40 | 2.125 | 4.37E-06 | 4.57E-05 | *IFNG, PSMB9, PSMB8, PSMB10, PSME2, PSME1, PSMA5, PSMA2, PSMA3, PSMA1, PSMB2, PSMD3, PSMB4, PSME4, PSMD14, PSMB3, PSMD11, PSMD12, PSMF1, PSMD7, PSMA4* |
| ssc05332 | Graft-versus-host disease | 23 | 2.114 | 9.56E-06 | 9.70E-05 | *IFNG, SLA-DOA, SLA-3, SLA-DRA, FASLG, FAS, SLA-DQA1, SLA-DOB, IL1B, SLA-DQB1, SLA-DMB, SLA-DMA, SLA-7, IL1A, SLA-DRB1, IL6, PRF1, SLA-8* |
| ssc04940 | Type I diabetes mellitus | 25 | 2.108 | 1.36E-05 | 1.34E-04 | *IFNG, SLA-DOA, SLA-3, SLA-DRA, FASLG, FAS, SLA-DQA1, SLA-DOB, IL1B, SLA-DQB1, SLA-DMB, SLA-DMA, SLA-7, IL1A, SLA-DRB1, PRF1, SLA-8* |
| ssc04658 | Th1 and Th2 cell differentiation | 73 | 2.071 | 3.28E-06 | 3.54E-05 | *IFNG, STAT1, SLA-DOA, SLA-DRA, SLA-DQA1, FOS, SLA-DOB, SLA-DQB1, SLA-DMB, SLA-DMA, IL4R, IL2RB, DLL4, SLA-DRB1, STAT5A, NFKBIB, NFKBIE, JAK3, NFKBIA, TBX21, IKBKB, IL2RA, JAK2, IL2RG, RUNX3, NFKB1* |
| ssc05330 | Allograft rejection | 21 | 2.061 | 2.13E-05 | 1.93E-04 | *IFNG, SLA-DOA, SLA-3, SLA-DRA, FASLG, FAS, SLA-DQA1, SLA-DOB, SLA-DQB1, SLA-DMB, SLA-DMA, SLA-7, CD40, SLA-DRB1, PRF1, SLA-8* |
| ssc05322 | Systemic lupus erythematosus | 48 | 2.049 | 2.10E-05 | 1.93E-04 | *IFNG, SLA-DOA, SLA-DRA, C2, SLA-DQA1, SLA-DOB, TRIM21, SLA-DQB1, SLA-DMB, C4A, SLA-DMA, CD40, C3, SLA-DRB1, C9, FCGR1A* |
| ssc05133 | Pertussis | 63 | 2.042 | 1.48E-05 | 1.42E-04 | *IRF1, AMCF-II, CXCL8, C2, FOS, IL1B, C4A, CASP1, IRF8, SFTPA1, IL1A, CASP7, C3, SERPING1, IL6, C4BPB, TICAM2, CASP3, ITGA5, MYD88, CD14, NOS2, TICAM1, NFKB1, TLR4, IL10, JUN, NLRP3* |
| ssc05320 | Autoimmune thyroid disease | 26 | 2.014 | 4.06E-05 | 3.42E-04 | *SLA-DOA, SLA-3, SLA-DRA, FASLG, FAS, TG, SLA-DQA1, SLA-DOB, SLA-DQB1, SLA-DMB, CTLA4, SLA-DMA, SLA-7, CD40, SLA-DRB1, PRF1, SLA-8* |
| ssc04672 | Intestinal immune network for IgA production | 34 | 1.997 | 1.22E-04 | 8.97E-04 | *MADCAM1, PIGR, SLA-DOA, SLA-DRA, TNFSF13B, SLA-DQA1, SLA-DOB, SLA-DQB1, SLA-DMB, SLA-DMA, CD40, SLA-DRB1, CCL28, IL6, IL15* |

| | | | | | | |
|---|---|---|---|---|---|---|
| ssc05161 | Hepatitis B | 141 | 1.983 | 1.56E-06 | 1.79E-05 | *RIGI, IRF7, STAT1, CXCL8, IFIH1, FASLG, FAS, FOS, STAT2, CASP10, EGR2, CDKN1A, STAT5A, IL6, IKBKE, TICAM2, STAT3, JAK3, CASP3, NFKBIA, TRAF3, MYC, MYD88, TLR2, MMP9, LOC100624590, TLR3, CREB3L1, CASP8, TICAM1, E2F2, IKBKB, JAK2, CYCS, NFKB1, TLR4, PTK2B* |
| ssc05417 | Lipid and atherosclerosis | 171 | 1.973 | 7.03E-07 | 1.01E-05 | *SELE, IRF7, LBP, CXCL2, CCL2, CXCL8, TNFSF10, FASLG, CCL3L1, FAS, FOS, IL1B, CASP1, VCAM1, CCL5, LOC100624226, CD40, CASP7, SOD2, HSPA5, MMP1, OLR1, IL6, IKBKE, TICAM2, HSP90B1, STAT3, VLDLR, CASP3, NFKBIA, TNFRSF1A, TRAF3, MYD88, HSPA8, SELP, TLR2, MMP9, CD14, GP91-PHOX, TRAF2, CASP8, NCF1, TICAM1, IKBKB, JAK2, CYCS, ERN1, ICAM1, NFKB1, TLR4, RXRA, DDIT3* |
| ssc04141 | Protein processing in endoplasmic reticulum | 149 | 1.970 | 1.05E-06 | 1.45E-05 | *HYOU1, HSPA5, PDIA4, BAK1, HSP90B1, SEC24D, DNAJB11, PDIA3, WFS1, HERPUD1, EDEM1, CALR, SEC23B, HSPA8, HSPH1, SEC61A1, LMAN1, CKAP4, TRAF2, SELENOS, DERL3, PREB, MOGS, ERN1, NGLY1, YOD1, SEC24A, DNAJC3, PDIA6, DNAJA1, P4HB, DDIT3, MAN1B1, UGGT1, DNAJC1, RPN2, RPN1, TRAM1, CANX, SSR1, ERO1B, EIF2AK3, STT3A, SEC31B, UBE2D1, SYVN1, OSTC, SSR3, SEC13, SEC24C, RAD23A, VCP, CUL1, HSPA1L, EIF2S1, HSPA4L, UBQLN1, MAGT1, GANAB, EDEM2, NPLOC4, UBE2D3, SEC31A, PLAA, STT3B, MAPK10, XBP1, SAR1B, SEL1L, ERLEC1* |
| ssc05310 | Asthma | 14 | 1.967 | 4.61E-05 | 3.70E-04 | *CCL11, SLA-DOA, SLA-DRA, SLA-DQA1, SLA-DOB, SLA-DQB1, SLA-DMB, SLA-DMA, CD40, SLA-DRB1, FCER1A* |
| ssc05167 | Kaposi sarcoma-associated herpesvirus infection | 156 | 1.956 | 2.65E-06 | 2.95E-05 | *IRF7, STAT1, CXCL2, CXCL8, SLA-3, EIF2AK2, IRF9, FAS, FOS, CLEC2B, STAT2, CDKN1A, SLA-7, ZFP36, C3, PTGS2, CSF2, IL6, CCR1, IKBKE, BAK1, STAT3, IFNAR2, SLA-8, CASP3, NFKBIA, TNFRSF1A, LOC100621431, TRAF3, MYC, TLR3, TRAF2, CASP8, TICAM1, E2F2, IKBKB, HCK, JAK2, CYCS, ICAM1, NFKB1, GNG10* |
| ssc04210 | Apoptosis | 114 | 1.914 | 3.70E-05 | 3.19E-04 | *TNFSF10, FASLG, FAS, FOS, CTSL, BIRC3, PMAIP1, GADD45G, CASP10, LOC100624226, CASP7, PARP3, PRF1, GADD45A, BAK1, CSF2RB, CFLAR, LOC100127131, CASP3, NFKBIA, TNFRSF1A, DAXX, BCL2A1, CTSO, CTSS, TRAF2, CASP8, MCL1, IKBKB, CYCS, ERN1, RIPK1, PTPN13, NFKB1, GADD45B, LMNB2, DDIT3* |

| | | | | | | |
|---|---|---|---|---|---|---|
| ssc04145 | Phagosome | 115 | 1.910 | 2.59E-05 | 2.29E-04 | *TAP1, SLA-DOA, TAP2, SLA-3, SLA-DRA, SLA-DQA1, CTSL, SLA-DOB, THBS1, SLA-DQB1, SLA-DMB, SLA-DMA, TCIRG1, SFTPA1, SLA-7, C3, SLA-DRB1, OLR1, ATP6V1C2, FCGR2B, SLA-8, LOC100127131, FCGR1A, CALR, ITGA5, TFRC, TLR2, CD14, SEC61A1, CTSS, GP91-PHOX, NCF1, CLEC7A, MRC1* |
| ssc05163 | Human cytomegalovirus infection | 190 | 1.904 | 1.44E-06 | 1.72E-05 | *TAP1, CCL4, TAP2, CCL2, CXCL8, SLA-3, FASLG, CCL3L1, FAS, CX3CL1, B2M, IL1B, TAPBP, CDKN1A, SLA-7, CCL5, TAPBPL, PTGS2, IL1R1, IL6, STING1, CCR1, BAK1, STAT3, PDIA3, CGAS, SLA-8, CASP3, NFKBIA, IL6R, TNFRSF1A, CALR, LOC100621431, MYC, CREB3L1, TRAF2, CASP8, PDGFRA, E2F2, IKBKB, CYCS, RIPK1, IL10RB, NFKB1, GNG10, ADCY3, CXCR2* |
| ssc05166 | Human T-cell leukemia virus 1 infection | 194 | 1.900 | 1.16E-06 | 1.54E-05 | *SLA-DOA, SLA-3, SLA-DRA, SLA-DQA1, FOS, B2M, SLA-DOB, SLA-DQB1, SLA-DMB, RELB, SLA-DMA, EGR2, IL2RB, CDKN1A, SLA-7, FOSL1, ZFP36, CD40, SLA-DRB1, STAT5A, EGR1, CSF2, IL1R1, IL1R2, IL6, IL15, NFKB2, MSX2, SLA-8, JAK3, NFKBIA, TNFRSF1A, CALR, MYC, TSPO, CREB3L1, XPO1, E2F2, IKBKB, IL2RA, IL2RG, ICAM1, NFKB1, ETS2, ADCY3, CCND3, LTBR, JUN, KAT2A, CANX, SLC25A5, NRP1, ELK4, E2F3, CREB3L3, TGFB3, MSX1, TBPL1* |
| ssc05142 | Chagas disease | 92 | 1.896 | 5.61E-05 | 4.40E-04 | *IFNG, CCL2, CXCL8, SERPINE1, FASLG, CCL3L1, FAS, FOS, IL1B, CCL5, C3, IL6, LOC100515049, CFLAR, NFKBIA, TNFRSF1A, CALR, MYD88, TLR2, NOS2, CASP8, TICAM1, IKBKB, NFKB1, TLR4, IL10* |
| ssc05134 | Legionellosis | 44 | 1.896 | 8.56E-04 | 5.68E-03 | *CXCL2, CXCL8, IL1B, CASP1, CASP7, C3, IL6, NFKB2, CASP3, NFKBIA, BNIP3, MYD88, HSPA8, TLR2, CD14, CASP8, CYCS, NFKB1, TLR4, HSPD1, TLR5* |
| ssc04936 | Alcoholic liver disease | 109 | 1.871 | 7.89E-05 | 5.92E-04 | *LBP, CXCL2, CXCL8, C2, FASLG, CPT1C, FAS, IL1B, C4A, C3, IL6, IKBKE, CASP3, NFKBIA, ALDH9A1, TNFRSF1A, TRAF3, MYD88, CD14, CASP8, TICAM1, IKBKB, PPARGC1A, RIPK1, NFKB1, TLR4, TAB3, IL17RA, FOXO1, MAP2K3, PPARA, TIRAP, TRA2B, MAP3K14, ALDH3A2, LY96, MLYCD, CPT1B, IRAK1* |
| ssc04380 | Osteoclast differentiation | 115 | 1.861 | 6.25E-05 | 4.79E-04 | *IFNG, STAT1, SOCS1, IRF9, SOCS3, FOS, IL1B, STAT2, RELB, JUNB, OSCAR, FOSL1, IL1A, LOC100622460, CSF1, IL1R1, LOC100522225, FOSB, NFKB2, IFNAR2, FCGR2B, FCGR1A, NFKBIA, TNFRSF1A, FOSL2, TNFRSF11B, TRAF2, NCF1, IKBKB* |

| | | | | | | |
|---|---|---|---|---|---|---|
| ssc05144 | Malaria | 39 | 1.858 | 8.30E-04 | 5.61E-03 | *IFNG, ACKR1, SELE, CCL2, CXCL8, THBS1, IL1B, VCAM1, CD40, CSF3, IL6, MYD88, SELP, TLR2, KLRK1, ICAM1, TLR4, IL10* |
| ssc05416 | Viral myocarditis | 51 | 1.836 | 1.18E-03 | 7.41E-03 | *SLA-DOA, SLA-3, SLA-DRA, SLA-DQA1, SLA-DOB, SLA-DQB1, SLA-DMB, SLA-DMA, SLA-7, CD40, SLA-DRB1, PRF1, SLA-8, CASP3, CASP8, CYCS, ICAM1* |
| ssc04625 | C-type lectin receptor signaling pathway | 90 | 1.827 | 1.96E-04 | 1.41E-03 | *LOC110258711, CLEC4E, IRF1, STAT1, IRF9, IL1B, BCL3, STAT2, RELB, EGR2, CASP1, PTGS2, IL6, IKBKE, NFKB2, NFKBIA, PLK3, CASP8, IKBKB, CLEC7A, NFKB1, IL10, RRAS, JUN, NLRP3* |
| ssc05150 | Staphylococcus aureus infection | 32 | 1.813 | 1.14E-03 | 7.26E-03 | *SLA-DOA, SLA-DRA, C2, SLA-DQA1, SLA-DOB, SLA-DQB1, SLA-DMB, C4A, SLA-DMA, CFB, C3, SLA-DRB1, FCGR2B, FCGR1A, SELP, ICAM1, CFI, IL10* |
| ssc04917 | Prolactin signaling pathway | 59 | 1.746 | 3.82E-03 | 2.03E-02 | *IRF1, STAT1, SOCS1, SOCS3, FOS, STAT5A, STAT3, CISH, GCK, JAK2, NFKB1, SOCS7* |
| ssc05143 | African trypanosomiasis | 28 | 1.743 | 6.17E-03 | 3.09E-02 | *IDO1, IFNG, SELE, FASLG, FAS, IL1B, VCAM1, IL6* |
| ssc00591 | Linoleic acid metabolism | 12 | 1.723 | 8.15E-03 | 3.91E-02 | *PLA2G2D, CYP3A22* |
| ssc05235 | PD-L1 expression and PD-1 checkpoint pathway in cancer | 81 | 1.711 | 1.55E-03 | 9.35E-03 | *BATF2, IFNG, CD274, STAT1, FOS, BATF3, NFKBIB, TICAM2, STAT3, NFKBIE, NFKBIA, MYD88, TLR2, TICAM1, IKBKB, JAK2, NFKB1, TLR4* |
| ssc05203 | Viral carcinogenesis | 149 | 1.702 | 4.44E-04 | 3.06E-03 | *IRF7, SLA-3, EIF2AK2, IRF9, PMAIP1, EGR2, LOC100517129, CDKN1A, SLA-7, C3, STAT5A, BAK1, STAT3, NFKB2, SLA-8, JAK3, CASP3, NFKBIA, TRAF3, LOC100624590, HDAC4, H2BC21, SCIN, CREB3L1, TRAF2, CASP8, NFKB1, CCND3, LTBR, JUN, H2BC15, KAT2A, SND1, CREB3L3, IL6ST, MAPKAPK2, TBPL1* |
| ssc04933 | AGE-RAGE signaling pathway | 93 | 1.701 | 1.90E-03 | 1.11E-02 | *SELE, STAT1, CCL2, CXCL8, SERPINE1, IL1B, PIM1, VCAM1, IL1A, STAT5A, EGR1, IL6, STAT3, CASP3, GP91-PHOX, JAK2, F3, ICAM1, NFKB1, AGTR1, COL4A1, JUN, PLCD4, AGT, FOXO1, TGFB3, COL4A2, MMP2, PLCG1* |

| | | | | | | |
|---|---|---|---|---|---|---|
| | in diabetic complications | | | | | |
| ssc04640 | Hematopoietic cell lineage | 68 | 1.694 | 3.01E-03 | 1.65E-02 | *SLA-DOA, SLA-DRA, SLA-DQA1, SLA-DOB, CD38, IL1B, SLA-DQB1, SLA-DMB, SLA-DMA, IL4R, CSF3R, KIT, IL1A, CSF3, SLA-DRB1, CSF2, CSF1, IL1R1, IL1R2, IL6, ITGA6, FCGR1A, IL6R, ITGA5, TFRC, CD22, CD14* |
| ssc00380 | Tryptophan metabolism | 32 | 1.661 | 7.29E-03 | 3.54E-02 | *IDO1, AOC1, CYP1B1, KYNU, ALDH9A1, KMO, IL4I1* |
| ssc05170 | Human immunodeficiency virus 1 infection | 175 | 1.635 | 1.05E-03 | 6.81E-03 | *TAP1, BST2, TAP2, SLA-3, SAMHD1, FASLG, FAS, FOS, B2M, TAPBP, SLA-7, TAPBPL, STING1, BAK1, PDIA3, CGAS, SLA-8, CASP3, NFKBIA, TNFRSF1A, CALR, LOC100621431, MYD88, TLR2, TNFRSF1B, TRAF2, CASP8, IKBKB, CYCS, RIPK1, NFKB1, GNG10, TLR4, PTK2B* |
| ssc04514 | Cell adhesion molecule (CAM) interaction | 114 | 1.632 | 2.47E-03 | 1.41E-02 | *MADCAM1, SELE, CD274, SLA-DOA, SLA-3, SLA-DRA, PDCD1LG2, SLA-DQA1, SLA-DOB, SLA-DQB1, SLA-DMB, CTLA4, SLA-DMA, VCAM1, SLA-7, CD40, SLA-DRB1, SIGLEC1, PVR, SLITRK2, SLA-8, ITGA6, CLDN5, SELP, CD22, SDC4* |
| ssc05418 | Fluid shear stress and atherosclerosis | 119 | 1.520 | 5.71E-03 | 2.91E-02 | *IFNG, SELE, CCL2, FOS, CTSL, IL1B, LOC100510917, VCAM1, IL1A, IL1R1, IL1R2, HSP90B1, DUSP1, TNFRSF1A, MMP9, SDC4, NCF1, IKBKB, ICAM1, NFKB1, NQO1* |
| ssc05132 | Salmonella infection | 215 | 1.519 | 3.42E-03 | 1.84E-02 | *CXCL8, TNFSF10, FOS, LOC100522887, BIRC3, IL1B, CASP1, LOC100624226, CASP7, IL6, BAK1, HSP90B1, MLKL, MYL7, GSDMD, LOC100127131, CASP3, NFKBIA, TNFRSF1A, RHOB, ACTR3B, MYC, MYD88, RHOG, TLR2, CD14, RIPK3, TRAF2, CASP8, RHOJ, AHNAK2, IKBKB, CYCS, RIPK1, NFKB1, RHOH, RIPK2, TLR4, RRAS, JUN, TAB3, TUBA4A, NLRP3, TLR5* |
| ssc05165 | Human papillomavirus infection | 281 | 1.466 | 2.74E-03 | 1.53E-02 | *ISG15, MX1, IRF1, STAT1, SLA-3, EIF2AK2, IRF9, FASLG, FAS, THBS1, STAT2, LAMB4, FZD9, TCIRG1, CDKN1A, SLA-7, HES4, PTGS2, SPP1, WNT8B, PPP2R5B, ATP6V1C2, IKBKE, COL2A1, BAK1, IFNAR2, ITGA7, TNC, SLA-8, CASP3, ITGA6, TNFRSF1A, ITGA5, TRAF3, FZD4, TLR3, WNT16, APC2, CREB3L1, CASP8, TICAM1, IKBKB, TNN, PDGFRB, NOTCH4, NFKB1, HES1* |
| ssc05200 | Pathways in cancer | 441 | 1.430 | 2.48E-03 | 1.41E-02 | *IFNG, STAT1, CXCL8, FASLG, FAS, FOS, BIRC3, PMAIP1, LOC100510917, STAT2, GADD45G, LAMB4, PIM1, IL4R, CSF3R, IL2RB, FZD9, PML, CDKN1A, F2R, DLL4, KIT, CASP7, ZBTB16, PTGS2, STAT5A, MMP1, EGLN3, WNT8B, IL6, IL15, GADD45A, BAK1, HSP90B1, GLI2, STAT3,* |

| | | | | | | |
|---|---|---|---|---|---|---|
| | | | | | | *NFKB2, IFNAR2, NKX3-1, CSF2RB, JAK3, RALGDS, CASP3, ITGA6, NFKBIA, MAX, IL6R, LOC100621431, TRAF3, MYC, FZD4, MMP9, WNT16, APC2, NOS2, RASSF1, TRAF2, CASP8, PDGFRA, E2F2, IKBKB, IL2RA, JAK2, PDGFRB, CYCS, NOTCH4, TXNRD1, IL2RG, NFKB1, GADD45B, GNG10, HES1, ADCY3, AR, NQO1, RXRA, AGTR1, CCND3, CSF2RA, COL4A1, JUN, AGT, PGF, CXCL12, GNA13, IL12RB2, HMOX1, FOXO1, RASGRP3, E2F3, RUNX1, PPARD, GNB4, TGFB3, IL6ST, FGFR4, IL13RA1, RALB, JAG1, COL4A2, MMP2* |
| ssc04310 | Wnt signaling pathway | 151 | -1.433 | 7.15E-03 | 3.53E-02 | *RSPO1, FZD8, RSPO2, WNT7A, APCDD1, LGR5, PRKCB, LGR4, WNT9A, AXIN2, TLE6, FZD2, SERPINF1, NOTUM, CAMK2G, NFATC1, BAMBI, TCF7, WNT11, WNT3A, PLCB2, TLE4, CTNNBIP1, WNT10B, WIF1, WNT3, PRKCG, WNT5B, RAC3, WNT4, WNT2, PRICKLE1, LOC100624648, CTNND2, NFATC2, SFRP5, SFRP4, DKK2, SFRP2* |
| ssc04020 | Calcium signaling pathway | 186 | -1.456 | 1.82E-03 | 1.08E-02 | *VEGFA, PHKA2, MCOLN1, CXH4orf3, CACNA1G, CALM3, ATP2B2, PLCE1, PLCD3, ADCY4, CAMK4, GNAS, PLCD1, CALM1, PDGFA, ADRA1D, P2RX7, HTR4, ADRA1B, PRKCB, PTGER3, MST1, GRIN2D, CAMK2G, ADCY1, NFATC1, FLT4, PDGFD, PLCB2, TRHR, P2RX5, PTGER1, GRIN3A, CACNA1E, GNA14, ATP2B4, EGF, PRKCG, RET, CASQ1, FGF18, SLC8A3, P2RX6, MCOLN2, NTRK2, RYR3, GRIN2B, CACNA1S, ADRB1, GRIN3B, NFATC2, FGF1, CHRM2, ADRA1A, VEGFD, CALML4, CACNA1B, P2RX3* |
| ssc03010 | Ribosome | 144 | -1.481 | 5.73E-03 | 2.91E-02 | *MRPL9, RPL13, RPS8, LOC100739087, RPL7A, RPL8, RPL7, RPL14, RPL36A, RPL31, RPS4X, RPL24, MRPS26, RPS27A, MRPL11, RPL6, MRPL49, RPL21, RPL18, MRPL22, RPS7, RPS5, RPL10A, RPS23, MRPS17, RPL19, RPLP2, MRPL14, RPS21, MRPS18C, MRPL45, MRPS2, RPS3A, RPS3, FAU, RPL27A, MRPL36, RPS17, RPL37, MRPL13, RPL35A, MRPS24, RPS13, RPS19, RPL39, RPL29, RPS29, MRPL10, RPS15, RPL32, RPS25, RPL10, RPS16, RPLP1, RPL5, RPS20, MRPL12, RPS15A, RPL17, MRPL52, RPS24, RPL38, RPL28, MRPL30, RPL26, RPL34, RPL11, MRPS25, RPL23A, RPL35, MRPS14, RPS28, RPS27, RPS11* |
| ssc04080 | Neuroactive ligand-receptor interaction | 149 | -1.507 | 4.21E-03 | 2.20E-02 | *TRHR, P2RX5, PTGER1, GRIN3A, NPFF, CNR1, F2RL1, LPAR2, P2RX6, P2RY8, LYPD6B, TSHR, S1PR5, GABRR3, LHB, GRIN2B, GRIA1, ADRB1, GRIN3B, GABRA1, CALCB, GRIK2, GABBR2,* |

| | | | | | | |
|---|---|---|---|---|---|---|
| | | | | | | *LYPD6, OPRL1, CHRM2, ADRA1A, VIPR2, PYY, GRIA4, GPR156, EDN3, APLN, QRFPR, ADRA2C, APLNR, AGTR2, P2RX3* |
| ssc04820 | Cytoskeleton in muscle cells | 178 | -1.542 | 1.52E-03 | 9.35E-03 | *DSC2, COL6A3, COMP, SNTB2, DMD, SGCA, FLNC, DIAPH3, VCL, ITGB7, SYNPO2, MYL9, SGCE, ITGB4, FHL2, SPTBN1, CSRP1, MYBPC3, SDC2, COL3A1, FBLN1, DSG2, COL4A4, ANK2, LAMA2, FBN3, ACTN2, MYH14, BGN, LAMA1, DCN, VIM, ITGA3, COL1A2, COL1A1, PDLIM4, COL6A5, COL4A3, ENO3, SDC1, TLN2, SPTBN4, TMOD2, FN1, MYOM3, TNNI1, ITGA4, SDC3, THBS3, COL9A3, FHL1, OBSCN, FBLN2, PDLIM3, TRIM55, COL6A6, TNNI3, MYH2* |
| ssc04360 | Axon guidance | 167 | -1.702 | 4.27E-05 | 3.51E-04 | *CDK5, MYL9, SEMA3A, PRKCZ, SEMA4G, NTNG2, CAMK2G, SEMA6C, SEMA6A, NEO1, EPHA4, EFNA3, PAK3, EFNA5, EFNA2, ROBO1, RHOD, UNC5C, BMPR2, EFNA4, PLXNB3, WNT5B, PLXNA3, BMP7, PARD6G, EFNB3, EPHA7, EPHB2, SHH, SEMA3C, SRGAP3, SEMA3E, RAC3, PAK6, WNT4, SLIT3, SEMA3B, LOC100620540, NFATC2, EPHB1, NTNG1, EPHA3, SEMA3D, NTN3* |
| ssc04974 | Protein digestion and absorption | 70 | -1.931 | 2.61E-04 | 1.84E-03 | *COL6A3, SLC16A10, MME, COL16A1, COL8A1, COL23A1, COL12A1, CTRB2, DPP4, COL14A1, COL3A1, COL4A4, SLC1A1, COL1A2, COL1A1, COL6A5, COL4A3, PRCP, SLC8A3, COL25A1, SLC7A9, COL26A1, COL19A1, COL9A3, COL8A2, COL21A1, FXYD2, SLC7A8, CPA1, COL6A6, COL15A1, KCNJ13* |

Abbreviations: AGE-RAGE: Advanced glycation end products-receptor for AGEs; FDR: False discovery rate; IgA: Immunoglobulin A; IL-17: Interleukin-17; JAK-STAT: Janus kinases-signal transducer and activator of transcription proteins; KEGG: Kyoto Encyclopedia of Genes and Genomes; NES: Normalized enrichment score; NF-kappa B: Nuclear factor-kappa B; NOD: Nucleotide-binding domain; PD-1: Programmed cell death protein 1; PD-L1: Programmed cell death ligand 1; RIG-I: Retinoic acid-inducible gene I; Th1/Th2/Th17: T helper 1/2/17 cells; TNF: Tumor necrosis factor.

**Supplementary Table S6. Significantly enriched pathways from the GSEA result of the GO:BP database, Low-glucose infection versus Low-glucose control.** Abbreviations: GO:BP: Biological Processes in Gene Ontology; GSEA: Gene set enrichment analysis.

| GO:BP ID | Description | Set Size | NES | P-value | FDR | Core enrichment gene |
|---|---|---|---|---|---|---|
| GO:0034097 | Response to cytokine | 214 | 2.639 | 1.00E-10 | 2.37E-09 | *CXCL9, CXCL11, CXCL10, UBD, ISG15, IFNG, MX1, ACKR1, OAS1, CXCL13, SELE, ZBP1, CCL4, IRF7, IRF1, AMCF-II, CHI3L1, STAT1, CCL2, CXCL8, SOCS1, TNFSF13B, GFPT2, MMP8, SNX10, IL1B, SLA-DQB1, MSC, STAT2, PIM1, IL4R, IRF8, TIFA, CCL5, FOSL1, KIT, ZFP36, IL1A, CD40, CCL8, STAT5A, NFIL3, EGR1, IL6, ITIH4, IL15, TICAM2, TIMP1, STAT3, CXCL16, CISH, NKX3-1, JAK3, NFKBIA, IL6R, TNFRSF1A, ADAM17, IL18RAP, TRAF3, TIMP3, RAB20, TLR2, NOS2, YBX3, TRAF2, CASP8, MCL1, IKBKB, JAK2, RIPK1, SPRY2* |
| GO:1901652 | Response to peptide | 214 | 2.639 | 1.00E-10 | 2.37E-09 | *CXCL9, CXCL11, CXCL10, UBD, ISG15, IFNG, MX1, ACKR1, OAS1, CXCL13, SELE, ZBP1, CCL4, IRF7, IRF1, AMCF-II, CHI3L1, STAT1, CCL2, CXCL8, SOCS1, TNFSF13B, GFPT2, MMP8, SNX10, IL1B, SLA-DQB1, MSC, STAT2, PIM1, IL4R, IRF8, TIFA, CCL5, FOSL1, KIT, ZFP36, IL1A, CD40, CCL8, STAT5A, NFIL3, EGR1, IL6, ITIH4, IL15, TICAM2, TIMP1, STAT3, CXCL16, CISH, NKX3-1, JAK3, NFKBIA, IL6R, TNFRSF1A, ADAM17, IL18RAP, TRAF3, TIMP3, RAB20, TLR2, NOS2, YBX3, TRAF2, CASP8, MCL1, IKBKB, JAK2, RIPK1, SPRY2* |
| GO:0019221 | Cytokine-mediated signaling pathway | 129 | 2.591 | 1.00E-10 | 2.37E-09 | *CXCL9, CXCL11, CXCL10, ISG15, IFNG, ACKR1, OAS1, CXCL13, ZBP1, CCL4, IRF7, IRF1, AMCF-II, STAT1, CCL2, CXCL8, SOCS1, TNFSF13B, MMP8, IL1B, STAT2, IL4R, TIFA, CCL5, FOSL1, KIT, IL1A, CD40, CCL8, STAT5A, EGR1, IL6, IL15, TICAM2, STAT3, CISH, JAK3, NFKBIA, IL6R, TNFRSF1A, ADAM17, IL18RAP, TRAF3* |
| GO:0043207 | Response to external biotic stimulus | 365 | 2.582 | 1.00E-10 | 2.37E-09 | *CXCL9, CXCL10, ISG20, GBP1, RSAD2, BATF2, ISG15, IFNG, MX1, MX2, OAS1, IFIT1, GBP2, CXCL13, DHX58, RIGI, OAS2, ZBP1, CCL4, IRF7, IRF1, AMCF-II, STAT1, LBP, BST2, CALHM6, IL27, IFIT2, CCL2, CXCL8, SAMHD1, IFIH1, S100A8, C2, S100A12, BPIFB1, MMP8, CD180, PTGIR, IL1B, SLA-DQB1, C4A, RNF19B, NMI, LTF, IFI35, STAT2, SLC15A3,* |

| | | | | | | |
|---|---|---|---|---|---|---|
| | | | | | | *TNFAIP3, PIM1, TRIM14, IRF8, UNC93B1, TIFA, IFIT5, CCL5, FOSL1, CFB, ZFP36, IL1A, CD40, MR1, C3, CCL8, HAVCR2, SERPING1, NOD2, BPIFA1, SLAMF6, TRIM26, GNLY, STING1, FFAR2, TREM1, C9, NFKBIB, TICAM2, RGS1, OPTN, CXCL16, DUSP10, MLKL, MORC3, CGAS, N4BP3, HP, NFKBIA, CEBPB, IL18RAP, TRAF3, SLC11A1, MYD88, RAB20, MSRB1, TLR2, TLR3, CD14, KLRK1, RIPK3, NOS2, GP91-PHOX, TRAF2, NKG7, PLSCR4, SELENOS, DDX21, CEBPG, PLSCR1, TICAM1, RNF115, CLEC7A, CHGA, JAK2* |
| GO:0051707 | Response to other organism | 365 | 2.582 | 1.00E-10 | 2.37E-09 | *CXCL9, CXCL10, ISG20, GBP1, RSAD2, BATF2, ISG15, IFNG, MX1, MX2, OAS1, IFIT1, GBP2, CXCL13, DHX58, RIGI, OAS2, ZBP1, CCL4, IRF7, IRF1, AMCF-II, STAT1, LBP, BST2, CALHM6, IL27, IFIT2, CCL2, CXCL8, SAMHD1, IFIH1, S100A8, C2, S100A12, BPIFB1, MMP8, CD180, PTGIR, IL1B, SLA-DQB1, C4A, RNF19B, NMI, LTF, IFI35, STAT2, SLC15A3, TNFAIP3, PIM1, TRIM14, IRF8, UNC93B1, TIFA, IFIT5, CCL5, FOSL1, CFB, ZFP36, IL1A, CD40, MR1, C3, CCL8, HAVCR2, SERPING1, NOD2, BPIFA1, SLAMF6, TRIM26, GNLY, STING1, FFAR2, TREM1, C9, NFKBIB, TICAM2, RGS1, OPTN, CXCL16, DUSP10, MLKL, MORC3, CGAS, N4BP3, HP, NFKBIA, CEBPB, IL18RAP, TRAF3, SLC11A1, MYD88, RAB20, MSRB1, TLR2, TLR3, CD14, KLRK1, RIPK3, NOS2, GP91-PHOX, TRAF2, NKG7, PLSCR4, SELENOS, DDX21, CEBPG, PLSCR1, TICAM1, RNF115, CLEC7A, CHGA, JAK2* |
| GO:0098542 | Defense response to other organism | 304 | 2.580 | 1.00E-10 | 2.37E-09 | *CXCL9, CXCL10, ISG20, GBP1, RSAD2, BATF2, ISG15, IFNG, MX1, MX2, OAS1, GBP2, CXCL13, DHX58, RIGI, OAS2, ZBP1, CCL4, IRF7, IRF1, AMCF-II, STAT1, LBP, BST2, CALHM6, IL27, CCL2, CXCL8, SAMHD1, IFIH1, C2, S100A12, BPIFB1, SLA-DQB1, C4A, RNF19B, NMI, LTF, IFI35, STAT2, SLC15A3, TNFAIP3, PIM1, TRIM14, IRF8, UNC93B1, TIFA, CCL5, FOSL1, CFB, CD40, MR1, C3, CCL8, HAVCR2, SERPING1, NOD2, BPIFA1, SLAMF6, TRIM26, GNLY, STING1, FFAR2, TREM1, C9, TICAM2, OPTN, CXCL16, DUSP10, MLKL, MORC3, CGAS, N4BP3, HP, NFKBIA, CEBPB, IL18RAP, TRAF3, MYD88, RAB20, MSRB1, TLR2, TLR3, CD14, KLRK1, RIPK3, NOS2, GP91-PHOX, TRAF2, NKG7, CEBPG, PLSCR1, TICAM1, RNF115, CHGA, JAK2* |

| | | | | | | |
|---|---|---|---|---|---|---|
| GO:0071345 | Cellular response to cytokine stimulus | 199 | 2.572 | 1.00E-10 | 2.37E-09 | *CXCL9, CXCL11, CXCL10, ISG15, IFNG, MX1, ACKR1, OAS1, CXCL13, ZBP1, CCL4, IRF7, IRF1, AMCF-II, CHI3L1, STAT1, CCL2, CXCL8, SOCS1, TNFSF13B, GFPT2, MMP8, SNX10, IL1B, SLA-DQB1, MSC, STAT2, PIM1, IL4R, IRF8, TIFA, CCL5, FOSL1, KIT, ZFP36, IL1A, CD40, CCL8, STAT5A, NFIL3, EGR1, IL6, IL15, TICAM2, STAT3, CISH, NKX3-1, JAK3, NFKBIA, IL6R, TNFRSF1A, ADAM17, IL18RAP, TRAF3, RAB20, TLR2, NOS2, YBX3, TRAF2, IKBKB, JAK2, RIPK1, SPRY2* |
| GO:0009607 | Response to biotic stimulus | 378 | 2.559 | 1.00E-10 | 2.37E-09 | *CXCL9, CXCL10, ISG20, GBP1, RSAD2, BATF2, ISG15, IFNG, MX1, MX2, OAS1, IFIT1, GBP2, CXCL13, DHX58, RIGI, OAS2, ZBP1, CCL4, IRF7, IRF1, AMCF-II, STAT1, LBP, BST2, CALHM6, IL27, IFIT2, CCL2, CXCL8, SAMHD1, IFIH1, S100A8, C2, S100A12, BPIFB1, MMP8, CD180, PTGIR, IL1B, SLA-DQB1, C4A, RNF19B, NMI, LTF, IFI35, STAT2, SLC15A3, TNFAIP3, PIM1, TRIM14, IRF8, UNC93B1, TIFA, IFIT5, CCL5, FOSL1, CFB, ZFP36, IL1A, CD40, MR1, C3, CCL8, HAVCR2, SERPING1, NOD2, BPIFA1, SLAMF6, TRIM26, GNLY, STING1, FFAR2, TREM1, C9, NFKBIB, TICAM2, RGS1, OPTN, CXCL16, DUSP10, MLKL, MORC3, CGAS, TXNIP, N4BP3, HP, NFKBIA, TIGAR, CEBPB, IL18RAP, TRAF3, SLC11A1, MYD88, RAB20, MSRB1, TLR2, TLR3, CD14, KLRK1, RIPK3, NOS2, GP91-PHOX, TRAF2, NKG7, PLSCR4, SELENOS, DDX21, CEBPG, PLSCR1, TICAM1, RNF115, CLEC7A, CHGA, JAK2* |
| GO:0140546 | Defense response to symbiont | 264 | 2.555 | 1.00E-10 | 2.37E-09 | *CXCL9, CXCL10, ISG20, GBP1, RSAD2, ISG15, IFNG, MX1, OAS1, GBP2, CXCL13, DHX58, RIGI, OAS2, ZBP1, CCL4, IRF7, IRF1, AMCF-II, STAT1, LBP, CALHM6, IL27, CCL2, CXCL8, SAMHD1, IFIH1, C2, S100A12, BPIFB1, SLA-DQB1, C4A, RNF19B, NMI, LTF, IFI35, STAT2, SLC15A3, TNFAIP3, PIM1, TRIM14, IRF8, UNC93B1, TIFA, CCL5, FOSL1, CFB, MR1, C3, CCL8, HAVCR2, SERPING1, NOD2, BPIFA1, SLAMF6, TRIM26, STING1, FFAR2, TREM1, C9, TICAM2, CXCL16, DUSP10, MORC3, CGAS, N4BP3, NFKBIA, IL18RAP, TRAF3, MYD88, RAB20, MSRB1, TLR2, TLR3, CD14, KLRK1, NOS2, GP91-PHOX, TRAF2, CEBPG, PLSCR1, TICAM1, RNF115* |

| | | | | | | |
|---|---|---|---|---|---|---|
| GO:0044419 | Biological process involved in interspecies interaction between organisms | 399 | 2.543 | 1.00E-10 | 2.37E-09 | *CXCL9, CXCL10, ISG20, GBP1, RSAD2, BATF2, ISG15, IFNG, MX1, MX2, OAS1, IFIT1, GBP2, CXCL13, DHX58, RIGI, OAS2, ZBP1, CCL4, IRF7, IRF1, AMCF-II, STAT1, LBP, BST2, CALHM6, IL27, IFIT2, CCL2, CXCL8, SAMHD1, IFIH1, S100A8, C2, S100A12, BPIFB1, MMP8, CD180, PTGIR, IL1B, SLA-DQB1, C4A, RNF19B, NMI, LTF, IFI35, STAT2, SLC15A3, TNFAIP3, PIM1, TRIM14, IRF8, UNC93B1, TIFA, IFIT5, CCL5, FOSL1, CFB, ZFP36, IL1A, CD40, MR1, C3, CCL8, HAVCR2, SERPING1, NOD2, BPIFA1, SLAMF6, TRIM26, ST6GALNAC1, GNLY, STING1, FFAR2, TREM1, C9, PRF1, NFKBIB, TICAM2, SIGLEC1, RGS1, OPTN, CXCL16, DUSP10, TMPRSS4, MLKL, MORC3, CGAS, N4BP3, HP, NFKBIA, CEBPB, IL18RAP, TRAF3, SLC11A1, MYD88, RAB20, MSRB1, TLR2, TLR3, CD14, KLRK1, RIPK3, NOS2, GP91-PHOX, TRAF2, NKG7, PLSCR4, SELENOS, DDX21, CEBPG, PLSCR1, TICAM1, RNF115, CLEC7A, CHGA, JAK2* |
| GO:0006955 | Immune response | 468 | 2.532 | 1.00E-10 | 2.37E-09 | *CXCL9, CXCL11, CXCL10, ISG20, GBP1, RSAD2, ISG15, IFNG, MX1, OAS1, CCL26, GBP2, CXCL13, DHX58, CCL19, RIGI, OAS2, ZBP1, CCL4, IRF7, IRF1, AMCF-II, STAT1, LBP, CCL11, SLA-DOA, CALHM6, IL27, CXCL2, CCL2, CXCL8, SLA-3, SAMHD1, SLA-DRA, TNFSF10, IFIH1, TNFSF13B, C2, FASLG, S100A12, CCL3L1, FAS, BPIFB1, SLA-DQA1, CD74, CD180, SLA-DOB, THBS1, IL1B, SLA-DQB1, SLA-DMB, C4A, RNF19B, NMI, LTF, XCL1, CTLA4, IFI35, STAT2, SLC15A3, TNFAIP3, ACKR4, PIM1, SLA-DMA, TRIM14, IRF8, UNC93B1, TIFA, CXCR6, CCL5, FOSL1, CFB, KIT, IL1A, CD40, MR1, CSF3, C3, CCL8, SLA-DRB1, NFIL3, HAVCR2, CCL28, SERPING1, LOC100622460, ACKR3, NOD2, BPIFA1, SLAMF6, CSF2, TRIM26, IL6, PARP3, STING1, CCR1, FFAR2, IL15, TREM1, C9, BMX, SLAMF8, TICAM2, STAT3, CXCL16, DUSP10, LIF, MORC3, CGAS, BCL6, ACKR2, N4BP3, SLA-8, JAK3, MAD2L2, FCGR1A, NFKBIA, IL6R, IL18RAP, TRAF3, MYD88, TFRC, RAB20, MSRB1, TNFSF14, TLR2, TLR3, CD14, ST3GAL1, KLRK1, RIPK3, NOS2, GP91-PHOX, TRAF2, SELENOS, DDX21, CEBPG, PLSCR1, TNFSF13, CD84, TICAM1, RNF115, CHGA, JAK2, CFI, TNFAIP1, TNFSF9, TLR4, TCIM, ATP1B1, LIG4, NCR1, IL10, POLR3D, PTK2B, BLNK, USP44, LRP8, CD200, SFTPD, C7, CD59, CMKLR1, PSMB4* |

| | | | | | | |
|---|---|---|---|---|---|---|
| GO:0006952 | Defense response | 450 | 2.502 | 1.00E-10 | 2.37E-09 | *CXCL9, CXCL11, CXCL10, ISG20, GBP1, SAA3, RSAD2, BATF2, ISG15, IFNG, MX1, ACKR1, MX2, OAS1, GBP2, CXCL13, DHX58, CCL19, RIGI, OAS2, ZBP1, CCL4, IRF7, IRF1, AMCF-II, CHI3L1, STAT1, LBP, CCL11, BST2, CALHM6, IL27, CXCL2, CCL2, CXCL8, SAMHD1, IFIH1, S100A8, C2, S100A12, BPIFB1, CD74, MMP8, THBS1, PTGIR, IL1B, SLA-DQB1, C4A, RNF19B, NMI, LTF, IFI35, STAT2, SLC15A3, TNFAIP3, PIM1, IL4R, TRIM14, CASP1, IRF8, UNC93B1, TIFA, CXCR6, CCL5, FOSL1, CFB, KIT, ZFP36, IL1A, CD40, MR1, APOD, C3, CCL8, STAT5A, HAVCR2, SERPING1, OLR1, NOD2, BPIFA1, SLAMF6, IL1RAP, TRIM26, IL6, GNLY, STING1, CCR1, FFAR2, ITIH4, TREM1, C9, NFKBIB, TICAM2, CD163, OPTN, STAT3, CXCL16, DUSP10, MLKL, MORC3, CGAS, BCL6, N4BP3, HP, FCGR1A, ELF4, NFKBIA, TNFRSF1A, B4GALT1, CEBPB, IL18RAP, TRAF3, MYD88, RAB20, MSRB1, TLR2, TLR3, CD14, KLRK1, RIPK3, NOS2, TNFRSF1B, GP91-PHOX, TRAF2, NKG7, PTN, SELENOS, CEBPG, PLSCR1, TICAM1, RNF115, IL2RA, CHGA, JAK2, SLC39A8, F3, RIPK1, SMAD1, ADORA1, CYP26B1, TLR4, ATP1B1, IL10, POLR3D, USP44, AGTR1, LRP8, SFTPD, C7, CD59, EPHA2, CMKLR1, PSMB4* |
| GO:0045087 | Innate immune response | 248 | 2.455 | 1.00E-10 | 2.37E-09 | *CXCL10, ISG20, GBP1, RSAD2, ISG15, IFNG, MX1, OAS1, GBP2, DHX58, RIGI, OAS2, ZBP1, IRF7, IRF1, STAT1, LBP, CALHM6, IL27, CCL2, SAMHD1, IFIH1, C2, S100A12, BPIFB1, SLA-DQB1, C4A, RNF19B, NMI, LTF, IFI35, STAT2, SLC15A3, TNFAIP3, PIM1, TRIM14, IRF8, UNC93B1, TIFA, CCL5, FOSL1, CFB, MR1, C3, HAVCR2, SERPING1, NOD2, BPIFA1, SLAMF6, TRIM26, STING1, FFAR2, TREM1, C9, TICAM2, CXCL16, DUSP10, MORC3, CGAS, N4BP3, NFKBIA, IL18RAP, TRAF3, MYD88, RAB20, MSRB1, TLR2, TLR3, CD14, KLRK1, NOS2, GP91-PHOX, TRAF2, CEBPG, PLSCR1, TICAM1, RNF115, JAK2* |
| GO:0006954 | Inflammatory response | 217 | 2.265 | 1.00E-10 | 2.37E-09 | *CXCL9, CXCL10, SAA3, IFNG, ACKR1, CXCL13, CCL19, CCL4, AMCF-II, CHI3L1, CCL11, IL27, CXCL2, CCL2, CXCL8, S100A8, S100A12, CD74, MMP8, THBS1, PTGIR, IL1B, C4A, IFI35, TNFAIP3, IL4R, TRIM14, CASP1, CXCR6, CCL5, FOSL1, KIT, ZFP36, IL1A, APOD, C3, CCL8, STAT5A, OLR1, IL1RAP, IL6, CCR1, FFAR2, ITIH4, NFKBIB, TICAM2, CD163, STAT3, DUSP10, BCL6, HP, FCGR1A, ELF4, NFKBIA, TNFRSF1A, B4GALT1, CEBPB, MYD88, TLR2,* |

| | | | | | | |
|---|---|---|---|---|---|---|
| | | | | | | *TLR3, CD14, NOS2, TNFRSF1B, GP91-PHOX, PTN, SELENOS, TICAM1, IL2RA, JAK2, SLC39A8, F3, SMAD1, ADORA1, CYP26B1, TLR4, IL10, AGTR1, EPHA2, CMKLR1, PSMB4* |
| GO:0002831 | Regulation of response to biotic stimulus | 139 | 2.220 | 4.93E-10 | 1.03E-08 | *RSAD2, ISG15, DHX58, ZBP1, IRF7, IRF1, CALHM6, SAMHD1, BPIFB1, CD180, LTF, IFI35, STAT2, SLC15A3, TNFAIP3, PIM1, UNC93B1, TIFA, FOSL1, HAVCR2, SERPING1, SLAMF6, STING1, FFAR2, TICAM2, OPTN, DUSP10, CGAS, NFKBIA, TIGAR, IL18RAP, TRAF3, MYD88, TLR2, KLRK1, PLSCR1, TICAM1, RNF115* |
| GO:0045088 | Regulation of innate immune response | 126 | 2.171 | 1.11E-08 | 1.65E-07 | *RSAD2, ISG15, DHX58, ZBP1, IRF7, IRF1, CALHM6, SAMHD1, BPIFB1, IFI35, STAT2, SLC15A3, TNFAIP3, PIM1, UNC93B1, TIFA, FOSL1, HAVCR2, SERPING1, SLAMF6, STING1, FFAR2, TICAM2, DUSP10, CGAS, NFKBIA, IL18RAP, MYD88, TLR2, KLRK1, PLSCR1, TICAM1, RNF115* |
| GO:1903131 | Mononuclear cell differentiation | 123 | 2.163 | 5.22E-08 | 6.87E-07 | *RSAD2, BATF2, IFNG, IRF1, IL27, SOCS1, TNFSF13B, IL1B, SLA-DQB1, IRF8, DLL4, KIT, IL1A, STAT5A, NFIL3, EGR1, SLAMF6, IL6, PRDM1, STAT3, DUSP10, LIF, TMEM176B, BCL6, FBXO7, JAK3, IL6R, CEBPB, MYC, ST3GAL1, RIPK3, CASP8, CEBPG, GATA2, LRRC8A, CYP26B1, TNFSF9, TLR4, LIG4, IL10, PTK2B, USP44* |
| GO:0033993 | Response to lipid | 156 | 2.156 | 1.03E-08 | 1.60E-07 | *CXCL9, CXCL10, CXCL13, AMCF-II, CXCL8, S100A8, MMP8, CD180, PTGIR, IL1B, LTF, AQP3, TNFAIP3, PIM1, JUNB, IRF8, ZFP36, IL1A, HAVCR2, SPP1, AARD, FFAR2, GPR155, NFKBIB, DUSP10, NKX3-1, GDAP1, NFKBIA, CALR, CEBPB, SLC11A1, AQP1, KLF9, KLRK1, NOS2, PLSCR4, SELENOS, TICAM1, JAK2, PLSCR3, CYP26B1, AR, TLR4, INSIG1, IL10, PTK2B* |
| GO:0001819 | Positive regulation of cytokine production | 134 | 2.144 | 4.66E-08 | 6.40E-07 | *RSAD2, ISG15, IFNG, DHX58, IRF7, IRF1, CHI3L1, STAT1, CD74, MMP8, THBS1, IL1B, ADRA2A, CASP1, IRF8, KIT, IL1A, CD40, STAT5A, HAVCR2, EGR1, SLAMF6, STING1, FFAR2, IL15, TICAM2, CGAS, CEBPB, MYD88, PANX2, TLR2, KLRK1, NOS2, TRAF2, CASP8, DDX21, CEBPG, TICAM1, JAK2, TMEM106A, TLR4, FRMD8, IL10, POLR3D, DDIT3, TMED10* |
| GO:0009617 | Response to bacterium | 128 | 2.105 | 1.33E-07 | 1.36E-06 | *CXCL9, CXCL10, ISG15, CXCL13, DHX58, AMCF-II, LBP, CXCL8, S100A8, MMP8, CD180, PTGIR, IL1B, LTF, TNFAIP3, IRF8, ZFP36, IL1A, HAVCR2, BPIFA1, GNLY, TREM1, NFKBIB,* |

| | | | | | | |
|---|---|---|---|---|---|---|
| | | | | | | *RGS1, OPTN, DUSP10, HP, NFKBIA, CEBPB, SLC11A1, MYD88, TLR2, KLRK1, NOS2, PLSCR4, SELENOS, TICAM1, CHGA, JAK2* |
| GO:0071396 | Cellular response to lipid | 120 | 2.101 | 1.91E-07 | 1.84E-06 | *CXCL9, CXCL10, CXCL13, AMCF-II, CXCL8, MMP8, CD180, IL1B, LTF, TNFAIP3, PIM1, IRF8, ZFP36, IL1A, HAVCR2, SPP1, FFAR2, GPR155, NFKBIB, NKX3-1, NFKBIA, CALR, AQP1, KLF9, KLRK1, NOS2, PLSCR4, SELENOS, TICAM1, JAK2, PLSCR3, CYP26B1, AR, INSIG1, IL10, PTK2B* |
| GO:0001775 | Cell activation | 261 | 2.099 | 1.20E-10 | 2.67E-09 | *CXCL10, RSAD2, BATF2, IFNG, CD274, IRF1, AMCF-II, IL27, CXCL8, SOCS1, TNFSF13B, MMP8, CD180, THBS1, IL1B, SLA-DQB1, ADRA2A, SLA-DMB, CTLA4, IFI35, TNFAIP3, IRF8, CCL5, DLL4, KIT, IL1A, CD40, STAT5A, NFIL3, HAVCR2, LRRC32, EGR1, SLAMF6, IL6, PARP3, PRDM1, IL15, HSH2D, STAT3, DUSP10, CTPS1, BCL6, FBXO7, JAK3, MAD2L2, ELF4, IL6R, CEBPB, IL18RAP, SLC7A1, TFRC, TNFSF14, ST3GAL1, KLRK1, RIPK3, TRAF2, CEBPG, PLSCR1, CD84, TICAM1, LRRC8A, CHGA, JAK2, TMEM106A, CYP26B1, TNFSF9, TLR4, TCIM, LIG4, ADRA2B, ENTPD1, IL10, PTK2B, USP44, IGFBP2, CD200* |
| GO:0002682 | Regulation of immune system process | 390 | 2.095 | 1.00E-10 | 2.37E-09 | *CXCL10, RSAD2, ISG15, IFNG, MADCAM1, DHX58, SELE, ZBP1, CD274, IRF7, IRF1, STAT1, CALHM6, IL27, CCL2, SAMHD1, SOCS1, TNFSF13B, C2, BPIFB1, MMP8, THBS1, IL1B, SLA-DQB1, SLA-DMB, C4A, LTF, CTLA4, AQP3, IFI35, STAT2, SLC15A3, TNFAIP3, PIM1, UNC93B1, TIFA, CCL5, FOSL1, CFB, KIT, ZFP36, IL1A, CD40, APOD, CSF3, C3, STAT5A, HAVCR2, SERPING1, LOC100622460, LRRC32, SLAMF6, PARP3, STING1, PRDM1, CCR1, FFAR2, IL15, C9, BMX, TICAM2, STAT3, DUSP10, LIF, TMEM176B, GDF15, CGAS, BCL6, FBXO7, JAK3, MAD2L2, NFKBIA, CEACAM1, CALR, CEBPB, IL18RAP, MYC, MYD88, SLC7A1, TFRC, TNFSF14, TLR2, KLRK1, RIPK3, NOS2, TRAF2, CASP8, PTN, SELENOS, DDX21, PLSCR1, CD84, TICAM1, RNF115, GATA2, JAK2, HSPA9, CFI, ADORA1, CYP26B1, TNFSF9, TLR4, TCIM, NCR1, IL10, POLR3D, PTK2B, BLNK, IGFBP2, LRP8, CD200, C7, CD59, CMKLR1, PSMB4* |
| GO:0045321 | Leukocyte activation | 226 | 2.082 | 5.31E-10 | 1.05E-08 | *RSAD2, BATF2, IFNG, CD274, IRF1, AMCF-II, IL27, CXCL8, SOCS1, TNFSF13B, MMP8, CD180, THBS1, IL1B, SLA-DQB1, SLA-DMB, CTLA4, IFI35, TNFAIP3, IRF8, CCL5, DLL4,* |

| | | | | | | |
|---|---|---|---|---|---|---|
| | | | | | | *KIT, IL1A, CD40, STAT5A, NFIL3, HAVCR2, LRRC32, EGR1, SLAMF6, IL6, PARP3, PRDM1, IL15, HSH2D, STAT3, DUSP10, CTPS1, BCL6, FBXO7, JAK3, MAD2L2, ELF4, IL6R, CEBPB, IL18RAP, SLC7A1, TFRC, TNFSF14, ST3GAL1, KLRK1, RIPK3, TRAF2, CEBPG, PLSCR1, CD84, TICAM1, LRRC8A, CHGA, JAK2, TMEM106A, CYP26B1, TNFSF9, TLR4, LIG4, IL10, PTK2B, USP44, IGFBP2, CD200* |
| GO:0002521 | Leukocyte differentiation | 147 | 2.082 | 1.44E-07 | 1.42E-06 | *RSAD2, BATF2, IFNG, IRF1, IL27, SOCS1, TNFSF13B, SNX10, IL1B, SLA-DQB1, LTF, JUNB, IRF8, DLL4, KIT, IL1A, STAT5A, NFIL3, EGR1, SLAMF6, IL6, PRDM1, CCR1, STAT3, DUSP10, LIF, TMEM176B, BCL6, FBXO7, JAK3, IL6R, CEBPB, MYC, ST3GAL1, RIPK3, CASP8, CEBPG, GATA2, LRRC8A, CYP26B1, TNFSF9, TLR4, LIG4, IL10, PTK2B, USP44, EPHA2, RRAS* |
| GO:0050776 | Regulation of immune response | 232 | 2.063 | 2.39E-09 | 4.26E-08 | *RSAD2, ISG15, IFNG, DHX58, ZBP1, IRF7, IRF1, CALHM6, IL27, SAMHD1, TNFSF13B, C2, BPIFB1, IL1B, SLA-DQB1, SLA-DMB, C4A, IFI35, STAT2, SLC15A3, TNFAIP3, PIM1, UNC93B1, TIFA, FOSL1, CFB, KIT, CD40, C3, HAVCR2, SERPING1, LOC100622460, SLAMF6, PARP3, STING1, FFAR2, IL15, C9, BMX, TICAM2, DUSP10, CGAS, BCL6, JAK3, MAD2L2, NFKBIA, IL18RAP, MYD88, TFRC, TLR2, KLRK1, RIPK3, NOS2, TRAF2, SELENOS, PLSCR1, TICAM1, RNF115, CFI, TLR4, NCR1, IL10, POLR3D, BLNK, LRP8, CD200, C7, CD59, CMKLR1, PSMB4* |
| GO:0032103 | Positive regulation of response to external stimulus | 166 | 2.049 | 2.50E-07 | 2.34E-06 | *CXCL10, RSAD2, IFNG, DHX58, ZBP1, IRF7, IRF1, CXCL8, S100A8, BPIFB1, CD74, MMP8, CD180, THBS1, IFI35, SLC15A3, TNFAIP3, UNC93B1, TIFA, CCL5, FOSL1, STAT5A, HAVCR2, SLAMF6, STING1, CCR1, FFAR2, TICAM2, OPTN, CGAS, NFKBIA, TNFRSF1A, CALR, CEBPB, IL18RAP, MYD88, TNFSF14, TLR2, KLRK1, PTN, PLSCR1, TICAM1, RNF115* |
| GO:0001816 | Cytokine production | 207 | 2.048 | 9.57E-09 | 1.55E-07 | *RSAD2, ISG15, IFNG, DHX58, IRF7, IRF1, CHI3L1, STAT1, CALHM6, SOCS1, CD74, MMP8, THBS1, IL1B, ADRA2A, LTF, TNFAIP3, CASP1, IRF8, KIT, ZFP36, IL1A, CD40, APOD, STAT5A, HAVCR2, LRRC32, EGR1, SLAMF6, CSF2, IL6, STING1, FFAR2, IL15, TICAM2, MORC3, CGAS, BCL6, ELF4, IL6R, CEBPB, TRAF3, MYD88, PANX2, TLR2, KLRK1, RIPK3,* |

| | | | | | | |
|---|---|---|---|---|---|---|
| | | | | | | *TSKU, NOS2, TRAF2, CASP8, DDX21, CEBPG, CD84, TICAM1, JAK2, TMEM106A, TLR4, FRMD8, IL10, POLR3D, DDIT3, TMED10, EPHA2, CMKLR1* |
| GO:0002250 | Adaptive immune response | 118 | 2.033 | 8.86E-07 | 7.33E-06 | *RSAD2, IFNG, IRF7, SLA-DOA, IL27, SLA-DRA, TNFSF13B, C2, SLA-DQA1, SLA-DOB, IL1B, SLA-DQB1, SLA-DMB, C4A, CTLA4, TNFAIP3, SLA-DMA, CFB, CD40, C3, SLA-DRB1, HAVCR2, SLAMF6, IL6, PARP3, TREM1, C9, BMX, STAT3, DUSP10, BCL6, JAK3, MAD2L2, FCGR1A, IL6R, IL18RAP, TFRC, KLRK1, RIPK3, TRAF2* |
| GO:0032101 | Regulation of response to external stimulus | 258 | 2.032 | 1.96E-09 | 3.66E-08 | *CXCL10, RSAD2, ISG15, IFNG, DHX58, ZBP1, IRF7, IRF1, CALHM6, CCL2, CXCL8, SAMHD1, SERPINE1, S100A8, BPIFB1, CD74, MMP8, CD180, THBS1, LTF, IFI35, STAT2, SLC15A3, TNFAIP3, PIM1, CASP1, UNC93B1, TIFA, CCL5, FOSL1, ZFP36, STAT5A, HAVCR2, SERPING1, SLAMF6, STING1, CCR1, FFAR2, TICAM2, OPTN, DUSP10, CGAS, BCL6, ELF4, NFKBIA, TNFRSF1A, CALR, CEBPB, IL18RAP, TRAF3, MYD88, TNFSF14, TLR2, KLRK1, PTN, SELENOS, PLSCR1, TICAM1, RNF115, JAK2, SLC39A8, ADORA1, TLR4, IL10, POLR3D, PTK2B, LRP8, CMKLR1, PSMB4, ADAM10, CXCL12, NLRP3, FFAR4, TLR5* |
| GO:0031347 | Regulation of defense response | 201 | 2.021 | 4.68E-08 | 6.40E-07 | *RSAD2, ISG15, IFNG, DHX58, ZBP1, IRF7, IRF1, CALHM6, SAMHD1, S100A8, BPIFB1, CD74, MMP8, IFI35, STAT2, SLC15A3, TNFAIP3, PIM1, CASP1, UNC93B1, TIFA, FOSL1, ZFP36, STAT5A, HAVCR2, SERPING1, SLAMF6, STING1, FFAR2, TICAM2, OPTN, DUSP10, CGAS, BCL6, ELF4, NFKBIA, TNFRSF1A, CEBPB, IL18RAP, TRAF3, MYD88, TLR2, KLRK1, SELENOS, PLSCR1, TICAM1, RNF115, JAK2, SLC39A8, RIPK1, ADORA1, TLR4, IL10, POLR3D, LRP8* |
| GO:0001817 | Regulation of cytokine production | 204 | 2.015 | 5.60E-08 | 6.87E-07 | *RSAD2, ISG15, IFNG, DHX58, IRF7, IRF1, CHI3L1, STAT1, CALHM6, SOCS1, CD74, MMP8, THBS1, IL1B, ADRA2A, LTF, TNFAIP3, CASP1, IRF8, KIT, ZFP36, IL1A, CD40, APOD, STAT5A, HAVCR2, LRRC32, EGR1, SLAMF6, STING1, FFAR2, IL15, TICAM2, MORC3, CGAS, BCL6, ELF4, CEBPB, TRAF3, MYD88, PANX2, TLR2, KLRK1, RIPK3, TSKU, NOS2, TRAF2, CASP8, DDX21, CEBPG, CD84, TICAM1, JAK2, TMEM106A, TLR4, FRMD8, IL10, POLR3D, DDIT3, TMED10, EPHA2, CMKLR1* |

| | | | | | | |
|---|---|---|---|---|---|---|
| GO:0002684 | Positive regulation of immune system process | 265 | 2.002 | 7.66E-09 | 1.30E-07 | *CXCL10, RSAD2, IFNG, MADCAM1, DHX58, ZBP1, IRF7, IRF1, SOCS1, TNFSF13B, C2, BPIFB1, MMP8, THBS1, IL1B, SLA-DQB1, SLA-DMB, C4A, AQP3, IFI35, SLC15A3, TNFAIP3, UNC93B1, TIFA, CCL5, FOSL1, CFB, KIT, IL1A, CD40, C3, STAT5A, HAVCR2, SERPING1, SLAMF6, STING1, CCR1, FFAR2, IL15, C9, BMX, TICAM2, DUSP10, LIF, CGAS, BCL6, MAD2L2, NFKBIA, CALR, IL18RAP, MYD88, SLC7A1, TFRC, TNFSF14, TLR2, KLRK1, NOS2, TRAF2, CASP8, PTN, DDX21, PLSCR1, TICAM1, RNF115, JAK2, CFI, TNFSF9, TLR4, IL10, POLR3D, PTK2B, BLNK, IGFBP2, C7, CD59, CMKLR1* |
| GO:0046649 | Lymphocyte activation | 192 | 1.999 | 1.31E-07 | 1.36E-06 | *RSAD2, IFNG, CD274, IRF1, IL27, SOCS1, TNFSF13B, CD180, IL1B, SLA-DQB1, SLA-DMB, CTLA4, TNFAIP3, IRF8, CCL5, DLL4, KIT, IL1A, CD40, STAT5A, NFIL3, HAVCR2, LRRC32, EGR1, SLAMF6, IL6, PARP3, PRDM1, IL15, HSH2D, STAT3, DUSP10, CTPS1, BCL6, FBXO7, JAK3, MAD2L2, ELF4, IL6R, CEBPB, SLC7A1, TFRC, TNFSF14, ST3GAL1, KLRK1, RIPK3, TRAF2, CEBPG, CD84, TICAM1, LRRC8A, JAK2, CYP26B1, TNFSF9, TLR4, LIG4, IL10, PTK2B, USP44, IGFBP2* |
| GO:0002252 | Immune effector process | 159 | 1.990 | 7.36E-07 | 6.24E-06 | *RSAD2, IFNG, DHX58, IRF7, CALHM6, IL27, C2, CD180, IL1B, SLA-DQB1, C4A, RNF19B, IFI35, IRF8, CFB, KIT, CD40, C3, HAVCR2, SERPING1, SLAMF6, IL6, PARP3, FFAR2, TREM1, C9, PRF1, STAT3, DUSP10, BCL6, JAK3, MAD2L2, FCGR1A, IL6R, IL18RAP, TFRC, ST3GAL1, KLRK1, RIPK3, NOS2, TRAF2, DDX21, CEBPG, CD84, TICAM1, CHGA, CFI, TLR4, TCIM, LIG4, IL10, PTK2B, C7, CD59* |
| GO:0002833 | Positive regulation of response to biotic stimulus | 110 | 1.982 | 7.32E-06 | 4.49E-05 | *RSAD2, DHX58, ZBP1, IRF7, IRF1, BPIFB1, CD180, IFI35, SLC15A3, TNFAIP3, UNC93B1, TIFA, FOSL1, HAVCR2, SLAMF6, STING1, FFAR2, TICAM2, CGAS, NFKBIA, IL18RAP, MYD88, TLR2, KLRK1, PLSCR1, TICAM1, RNF115* |
| GO:0045089 | Positive regulation of innate immune response | 105 | 1.977 | 2.31E-05 | 1.26E-04 | *RSAD2, DHX58, ZBP1, IRF7, IRF1, BPIFB1, IFI35, SLC15A3, TNFAIP3, UNC93B1, TIFA, FOSL1, HAVCR2, SLAMF6, STING1, FFAR2, TICAM2, CGAS, NFKBIA, IL18RAP, MYD88, TLR2, KLRK1, PLSCR1, TICAM1, RNF115* |
| GO:0031349 | Positive regulation of defense response | 138 | 1.966 | 5.25E-06 | 3.60E-05 | *RSAD2, IFNG, DHX58, ZBP1, IRF7, IRF1, S100A8, BPIFB1, CD74, MMP8, IFI35, SLC15A3, TNFAIP3, UNC93B1, TIFA, FOSL1, STAT5A, HAVCR2, SLAMF6, STING1, FFAR2, TICAM2,* |

| | | | | | | |
|---|---|---|---|---|---|---|
| | | | | | | *OPTN, CGAS, NFKBIA, TNFRSF1A, CEBPB, IL18RAP, MYD88, TLR2, KLRK1, PLSCR1, TICAM1, RNF115, RIPK1* |
| GO:0006935 | Chemotaxis | 121 | 1.951 | 1.32E-05 | 7.33E-05 | *CXCL9, CXCL11, CXCL10, CXCL13, CCL4, AMCF-II, CCL2, CXCL8, S100A8, THBS1, ACKR4, CXCR6, CCL5, KIT, CCL8, ACKR3, CCR1, FFAR2, TREM1, CXCL16, ACKR2, CALR, TNFSF14, KLRK1, PTN, PDGFRA, CHGA, SEMA4D, PTK2B, AGTR1, EPHA2, CMKLR1, ADAM10, CXCL12* |
| GO:0042330 | Taxis | 121 | 1.951 | 1.32E-05 | 7.33E-05 | *CXCL9, CXCL11, CXCL10, CXCL13, CCL4, AMCF-II, CCL2, CXCL8, S100A8, THBS1, ACKR4, CXCR6, CCL5, KIT, CCL8, ACKR3, CCR1, FFAR2, TREM1, CXCL16, ACKR2, CALR, TNFSF14, KLRK1, PTN, PDGFRA, CHGA, SEMA4D, PTK2B, AGTR1, EPHA2, CMKLR1, ADAM10, CXCL12* |
| GO:0042110 | T cell activation | 133 | 1.942 | 8.72E-06 | 5.09E-05 | *RSAD2, IFNG, CD274, IRF1, IL27, SOCS1, TNFSF13B, IL1B, SLA-DQB1, SLA-DMB, CTLA4, CCL5, DLL4, KIT, IL1A, STAT5A, HAVCR2, LRRC32, EGR1, SLAMF6, IL6, PRDM1, IL15, HSH2D, STAT3, DUSP10, CTPS1, BCL6, JAK3, ELF4, IL6R, CEBPB, SLC7A1, TFRC, TNFSF14, RIPK3, TRAF2, CD84, JAK2, CYP26B1, TNFSF9, LIG4, USP44, IGFBP2* |
| GO:0030097 | Hemopoiesis | 236 | 1.938 | 1.17E-07 | 1.26E-06 | *RSAD2, BATF2, ISG15, IFNG, IRF1, STAT1, IL27, SOCS1, TNFSF13B, SNX10, IL1B, SLA-DQB1, LTF, PIM1, JUNB, IRF8, DLL4, KIT, ZFP36, IL1A, CSF3, STAT5A, NFIL3, EGR1, SLAMF6, CSF2, IL6, PRDM1, CCR1, STAT3, DUSP10, LIF, SLC1A5, CEBPD, TMEM176B, BCL6, FBXO7, JAK3, CASP3, NFKBIA, IL6R, CEBPB, MYC, ST3GAL1, RIPK3, CASP8, PTN, CEBPG, GATA2, LRRC8A, JAK2, HSPA9, GPATCH4, CYP26B1, TNFSF9, TLR4, TCIM, LIG4, IL10, PTK2B, USP44* |
| GO:0050778 | Positive regulation of immune response | 190 | 1.913 | 3.09E-06 | 2.20E-05 | *RSAD2, IFNG, DHX58, ZBP1, IRF7, IRF1, TNFSF13B, C2, BPIFB1, IL1B, SLA-DQB1, SLA-DMB, C4A, IFI35, SLC15A3, TNFAIP3, UNC93B1, TIFA, FOSL1, CFB, CD40, C3, HAVCR2, SERPING1, SLAMF6, STING1, FFAR2, IL15, C9, BMX, TICAM2, CGAS, MAD2L2, NFKBIA, IL18RAP, MYD88, TFRC, TLR2, KLRK1, NOS2, TRAF2, PLSCR1, TICAM1, RNF115, CFI, TLR4, IL10, POLR3D, BLNK, C7, CD59, CMKLR1* |

| | | | | | | |
|---|---|---|---|---|---|---|
| GO:0010628 | Positive regulation of gene expression | 304 | 1.901 | 6.18E-08 | 7.34E-07 | *RSAD2, ISG15, IFNG, DHX58, IRF7, IRF1, CHI3L1, STAT1, CXCL8, CD74, MMP8, THBS1, IL1B, ADRA2A, CASP1, ATF3, IRF8, DLL4, KIT, ZFP36, IL1A, CD40, STAT5A, NFIL3, HAVCR2, LRRC32, EGR1, SLAMF6, STING1, PRDM1, FFAR2, IL15, IGF2BP1, TICAM2, LIF, NKX3-1, CGAS, ARID5A, MAD2L2, CLDN5, CALR, CEBPB, MYD88, TFRC, PANX2, TLR2, WNT16, KLRK1, NOS2, TRAF2, CASP8, DDX21, CEBPG, TICAM1, GATA2, JAK2, PPARGC1A, SPRY2, SMAD1, MYOCD, TMEM106A, CYP26B1, AR, TLR4, FRMD8, FAM98A, EIF5A, IL10, POLR3D, BLNK, DDIT3, TMED10* |
| GO:0002694 | Regulation of leukocyte activation | 134 | 1.890 | 3.79E-05 | 1.99E-04 | *IFNG, CD274, IRF1, IL27, SOCS1, TNFSF13B, MMP8, THBS1, IL1B, SLA-DQB1, SLA-DMB, CTLA4, TNFAIP3, CCL5, IL1A, CD40, STAT5A, HAVCR2, LRRC32, PARP3, PRDM1, IL15, DUSP10, BCL6, FBXO7, JAK3, MAD2L2, CEBPB, SLC7A1, TFRC, TNFSF14, RIPK3, PLSCR1, CD84, TICAM1, JAK2, CYP26B1, TNFSF9, TLR4, IL10, IGFBP2, CD200* |
| GO:0080134 | Regulation of response to stress | 333 | 1.889 | 5.52E-08 | 6.87E-07 | *RSAD2, ISG15, IFNG, DHX58, ZBP1, IRF7, IRF1, CALHM6, SAMHD1, SERPINE1, S100A8, FAS, BPIFB1, CD74, MMP8, THBS1, ADRA2A, IFI35, STAT2, SLC15A3, TNFAIP3, PIM1, CASP1, UNC93B1, TIFA, FOSL1, ZFP36, STAT5A, HAVCR2, SERPING1, ACKR3, SLAMF6, PARP3, STING1, FFAR2, TICAM2, OPTN, DUSP10, CGAS, SESN2, BCL6, MAD2L2, ELF4, NFKBIA, TIGAR, TNFRSF1A, CEBPB, IL18RAP, TRAF3, MYC, MYD88, TLR2, RNASEH2B, KLRK1, YBX3, PTN, SELENOS, CEBPG, PLSCR1, TICAM1, RNF115, FICD, JAK2, SLC39A8, RIPK1, ADORA1, TLR4, INSIG1, IL10, POLR3D, LRP8, PSMD10, PSMB4, CXCL12, CLDN4, NLRP3, FFAR4, TLR5, USP51* |
| GO:0002683 | Negative regulation of immune system process | 116 | 1.878 | 5.23E-05 | 2.62E-04 | *ISG15, DHX58, CD274, IRF1, CCL2, SOCS1, BPIFB1, THBS1, LTF, STAT2, TNFAIP3, PIM1, APOD, HAVCR2, SERPING1, LRRC32, PARP3, DUSP10, TMEM176B, GDF15, CGAS, BCL6, FBXO7, JAK3, CEBPB, MYC, KLRK1, SELENOS, CD84, RNF115, GATA2, HSPA9, ADORA1, IL10, CD200, CD59, PSMB4* |
| GO:0002764 | Immune response-regulating signaling pathway | 121 | 1.874 | 5.96E-05 | 2.95E-04 | *RSAD2, IFNG, DHX58, IRF7, IRF1, BPIFB1, IFI35, SLC15A3, TNFAIP3, UNC93B1, TIFA, FOSL1, KIT, CD40, HAVCR2, LOC100622460, STING1, FFAR2, BMX, TICAM2, CGAS, NFKBIA, MYD88, TLR2, KLRK1, NOS2, PLSCR1, TICAM1, RNF115, TLR4, NCR1, BLNK* |

| | | | | | | |
|---|---|---|---|---|---|---|
| GO:0002253 | Activation of immune response | 145 | 1.837 | 8.47E-05 | 4.02E-04 | *RSAD2, IFNG, DHX58, IRF7, IRF1, C2, BPIFB1, C4A, IFI35, SLC15A3, TNFAIP3, UNC93B1, TIFA, FOSL1, CFB, C3, HAVCR2, SERPING1, STING1, FFAR2, C9, BMX, TICAM2, CGAS, NFKBIA, MYD88, TLR2, KLRK1, NOS2, PLSCR1, TICAM1, RNF115, CFI, TLR4, BLNK, C7, CD59, CMKLR1* |
| GO:0050865 | Regulation of cell activation | 146 | 1.835 | 3.99E-05 | 2.06E-04 | *IFNG, CD274, IRF1, IL27, SOCS1, TNFSF13B, MMP8, THBS1, IL1B, SLA-DQB1, SLA-DMB, CTLA4, TNFAIP3, CCL5, IL1A, CD40, STAT5A, HAVCR2, LRRC32, PARP3, PRDM1, IL15, DUSP10, BCL6, FBXO7, JAK3, MAD2L2, CEBPB, SLC7A1, TFRC, TNFSF14, RIPK3, PLSCR1, CD84, TICAM1, JAK2, CYP26B1, TNFSF9, TLR4, IL10, IGFBP2, CD200* |
| GO:0051249 | Regulation of lymphocyte activation | 120 | 1.833 | 9.92E-05 | 4.53E-04 | *IFNG, CD274, IRF1, IL27, SOCS1, TNFSF13B, IL1B, SLA-DQB1, SLA-DMB, CTLA4, TNFAIP3, CCL5, IL1A, CD40, STAT5A, HAVCR2, LRRC32, PARP3, PRDM1, IL15, DUSP10, BCL6, FBXO7, JAK3, MAD2L2, CEBPB, SLC7A1, TFRC, TNFSF14, RIPK3, TICAM1, JAK2, CYP26B1, TNFSF9, TLR4, IL10, IGFBP2* |
| GO:0042127 | Regulation of cell population proliferation | 374 | 1.832 | 6.73E-08 | 7.73E-07 | *CXCL9, CXCL11, CXCL10, IFNG, CD274, IRF1, STAT1, IL27, CCL2, TNFSF13B, THBS1, PTGIR, IL1B, LTF, CTLA4, IFI35, STAT2, TRNP1, TNFAIP3, PIM1, JUNB, IFI30, CCL5, DLL4, ZFP36, IL1A, CD40, APOD, CSF3, STAT5A, HAVCR2, EGLN3, ACKR3, LRRC32, CSF2, ESM1, ADM, IL6, PRDM1, IL15, TIMP1, GLI2, STAT3, DUSP10, LIF, MORC3, NKX3-1, BCL6, NFKBIA, CLDN5, IL6R, CALR, B4GALT1, CEBPB, MYC, AQP1, SLC7A1, TFRC, KLF9, RNASEH2B, PHLDA2, FUT2, RIPK3, NOS2, PTN, ATF5, TICAM1, GATA2, JAK2, PPARGC1A, F3, SPRY2, SMAD1, PPP1R16B, TNFSF9, AR, TLR4, FAM98A, LIG4, IL10, SLC35F6, PTK2B, IGFBP2, AGTR1, SLC25A33, ADAM10, JUN, PID1, SIX4, IL12RB2, NOP2, HMOX1, SLC25A5, GTPBP4, C4H1orf56, HAPLN1, GRK5* |
| GO:1901700 | Response to oxygen-containing compound | 311 | 1.831 | 5.94E-07 | 5.29E-06 | *CXCL9, CXCL10, CXCL13, AMCF-II, STAT1, LPL, CXCL8, SOCS1, S100A8, MMP8, CD180, THBS1, PTGIR, IL1B, ADRA2A, LTF, AQP3, STAT2, TNFAIP3, PIM1, IRF8, SLC2A4, ZFP36, IL1A, APOD, STAT5A, HAVCR2, SPP1, GPR37, EGR1, AARD, FFAR2, AQP9, GPR155, NFKBIB, STAT3, DUSP10, NKX3-1, GDF15, GDAP1, SESN2, JAK3, NFKBIA, RHOB, CEBPB,* |

| | | | | | | |
|---|---|---|---|---|---|---|
| | | | | | | *SLC11A1, AQP1, KLF9, PLK3, TLR2, KLRK1, RIPK3, NOS2, TRAF2, PLSCR4, SELENOS, TICAM1, JAK2, PLSCR3, CYP26B1, AR, TLR4, INSIG1, IL10, PTK2B, IGFBP2, SLC25A33* |
| GO:0030099 | Myeloid cell differentiation | 128 | 1.826 | 9.53E-05 | 4.41E-04 | *BATF2, ISG15, IFNG, STAT1, SOCS1, SNX10, LTF, JUNB, IRF8, KIT, ZFP36, CSF3, STAT5A, CSF2, CCR1, STAT3, LIF, SLC1A5, BCL6, CASP3, NFKBIA, CEBPB, MYC, CASP8, CEBPG, GATA2, JAK2, HSPA9, TLR4, PTK2B* |
| GO:0040011 | Locomotion | 285 | 1.818 | 2.42E-06 | 1.83E-05 | *CXCL9, CXCL11, CXCL10, IFNG, MADCAM1, CXCL13, SELE, CCL4, AMCF-II, CCL2, CXCL8, S100A8, THBS1, ADRA2A, ACKR4, CXCR6, CCL5, DLL4, KIT, CD40, APOD, CCL8, ACKR3, GPR37, GLUL, CCR1, FFAR2, BVES, TREM1, CXCL16, DUSP10, ARRDC3, GDF15, MYOC, BCL6, ACKR2, FBXO7, CLDN5, RHOB, CALR, TNFSF14, PHLDA2, KLRK1, RIPK3, CASP8, PTN, PDGFRA, GATA2, CHGA, SPRY2, ADORA1, TLR4, SEMA4D, STC1, PTK2B, AGTR1, EPHA2, CMKLR1, ADAM10, RRAS, CXCL12, GNA13, CLDN4, GTPBP4, NRP1, NUS1, MEAK7, XCR1* |
| GO:0002757 | Immune response-activating signaling pathway | 116 | 1.814 | 1.43E-04 | 6.42E-04 | *RSAD2, IFNG, DHX58, IRF7, IRF1, BPIFB1, IFI35, SLC15A3, TNFAIP3, UNC93B1, TIFA, FOSL1, HAVCR2, STING1, FFAR2, BMX, TICAM2, CGAS, NFKBIA, MYD88, TLR2, KLRK1, NOS2, PLSCR1, TICAM1, RNF115* |
| GO:0055085 | Transmembrane transport | 259 | 1.811 | 2.27E-06 | 1.75E-05 | *CXCL9, CXCL11, CXCL10, CALHM6, SLC25A47, AQP7, CNGA4, THBS1, ADRA2A, AQP3, GLRB, ATP1A2, SLC15A3, KCNJ2, SLC25A25, KCNMA1, SLC22A4, SLC2A4, C3, SLC25A22, BPIFA1, STING1, AQP9, GPR155, SLC25A28, SLC1A5, SESN2, SLC7A2, KCNA2, SLC5A6, NALF2, SLC11A1, AQP1, SLC7A1, PANX2, TOMM40, SLC38A5, SEC61A1, ATP13A5, ABCG2, GP91-PHOX, LRRC8A, SLC39A8, SLC31A1, KCNK7, ATP1B1, TMEM38B, KCNJ14, PTK2B, SLC5A1, SLC1A3, DDIT3, SLC25A33, SLC30A7, ATP2A2, PID1, SLC35E4, TRAM1, FFAR4, SLC25A5, SLC47A2, XCR1, SLC9A1, LRRC8B, GJA1, SLC16A13, SLC35B3, SLC4A4, GHITM, ABCB7, SLC33A1, ATP6V1B2, PDZK1, LOC100157391, SCNN1G, KCNJ15* |
| GO:0016477 | Cell migration | 339 | 1.809 | 5.01E-07 | 4.57E-06 | *CXCL9, CXCL11, CXCL10, IFNG, MADCAM1, CXCL13, SELE, CCL4, AMCF-II, RASGEF1A, CCL2, CXCL8, S100A8, S100A12, THBS1, ADRA2A, CDH12, TNFAIP3, ACKR4, CXCR6, CCL5, DLL4, KIT, CD40, APOD, CCL8, ACKR3, GLUL, CCR1, FFAR2, TREM1, CXCL16,* |

| | | | | | | |
|---|---|---|---|---|---|---|
| | | | | | | *DUSP10, GDF15, MYOC, ACKR2, CLDN5, RHOB, CALR, B4GALT1, ITGA5, PLEKHO1, TNFSF14, PHLDA2, KLRK1, RIPK3, SDC4, CASP8, PTN, GATA2, CHGA, SPRY2, ADORA1, TNFAIP1, TLR4, SEMA4D, FSCN1, STC1, PTK2B, AGTR1, LRP8, DDIT3, EPHA2, CMKLR1, ADAM10, RRAS, ENPEP, CXCL12, SIX4, GNA13, CLDN4, GTPBP4, NRP1, NARS1, SEMA4C, NUS1, MEAK7, XCR1* |
| GO:0008283 | Cell population proliferation | 444 | 1.787 | 7.20E-08 | 8.01E-07 | *CXCL9, CXCL11, CXCL10, IFNG, CD274, IRF1, STAT1, IL27, CCL2, GNAT1, TNFSF13B, CD180, THBS1, PTGIR, IL1B, LTF, CTLA4, IFI35, STAT2, TRNP1, TNFAIP3, PIM1, JUNB, IFI30, CCL5, DLL4, ZFP36, IL1A, CD40, APOD, CSF3, STAT5A, HAVCR2, EGLN3, ACKR3, LRRC32, CSF2, ESM1, ADM, IL6, PRDM1, IL15, IGF2BP1, TIMP1, GLI2, STAT3, DUSP10, LIF, CTPS1, MORC3, NKX3-1, SDCBP2, BCL6, ELF4, NFKBIA, CLDN5, IL6R, CALR, B4GALT1, CEBPB, IL18RAP, MYC, AQP1, SLC7A1, TFRC, KLF9, DRAM2, RNASEH2B, PHLDA2, WNT16, FUT2, RIPK3, NOS2, PTN, ATF5, TICAM1, GATA2, JAK2, PPARGC1A, F3, SPRY2, SMAD1, PPP1R16B, TNFSF9, AR, TLR4, FAM98A, LIG4, STC1, IL10, SLC35F6, PTK2B, IGFBP2, AGTR1, DDIT3, SLC25A33, EPHA2, ADAM10, JUN, PID1, ENPEP, SIX4, KAT2A, IL12RB2, NOP2, HMOX1, SLC25A5, GTPBP4, C4H1orf56, HAPLN1, GRK5, TIPIN, MEAK7, TGFB3, ETV6, SQSTM1, NME1, FGFR4, FBXO4, OGFOD1* |
| GO:0042981 | Regulation of apoptotic process | 344 | 1.785 | 6.89E-07 | 5.98E-06 | *CXCL10, IFNG, CD274, STAT1, CCL2, S100A8, FASLG, FAS, LOC100522887, THBS1, BCL2L14, LTF, GADD45G, CASP10, TNFAIP3, PIM1, CASP1, ATF3, CCL5, ZFP36, CD40, STAT5A, ACKR3, NOD2, EGR1, HSH2D, BAK1, GLI2, NKX3-1, ANGPTL4, CFLAR, BCL6, FBXO7, JAK3, CASP3, TIGAR, RHOB, BCL2A1, B4GALT1, ITGA5, CEBPB, TRAF3, MYC, AQP1, PLK3, PHLDA2, ST3GAL1, WNT16, RIPK3, YBX3, TRAF2, CASP8, SELENOS, ATF5, MCL1, GATA2, JAK2, PPARGC1A, F3, SPRY2, UTP11, GADD45B, HSPA9, PLSCR3, RIPK2, AR, TLR4, TCIM, LIG4, IL10, SLC35F6, PTK2B, RNF122, G0S2, LRP8, DDIT3, PSMD10, TNFRSF12A, ATP2A2, HIGD1A, CXCL12, CTH, SIX4, FFAR4, HMOX1, FOXO1, SLC25A5, NRP1, GRK5* |

| | | | | | | |
|---|---|---|---|---|---|---|
| GO:1901701 | Cellular response to oxygen-containing compound | 243 | 1.784 | 7.66E-06 | 4.62E-05 | *CXCL9, CXCL10, CXCL13, AMCF-II, STAT1, CXCL8, SOCS1, MMP8, CD180, IL1B, ADRA2A, LTF, TNFAIP3, PIM1, IRF8, SLC2A4, ZFP36, IL1A, STAT5A, HAVCR2, SPP1, GPR37, FFAR2, AQP9, GPR155, NFKBIB, STAT3, GDF15, SESN2, JAK3, NFKBIA, RHOB, AQP1, KLF9, TLR2, KLRK1, RIPK3, NOS2, TRAF2, PLSCR4, SELENOS, TICAM1, JAK2, PLSCR3, CYP26B1, AR, TLR4, INSIG1, IL10, PTK2B, SLC25A33* |
| GO:0022407 | Regulation of cell-cell adhesion | 103 | 1.771 | 5.56E-04 | 2.20E-03 | *IFNG, SELE, CD274, IRF1, SOCS1, TNFSF13B, IL1B, SLA-DQB1, SLA-DMB, CCL5, IL1A, STAT5A, HAVCR2, LRRC32, IL15, DUSP10, BCL6, JAK3, MAD2L2, CEBPB, SLC7A1, TFRC, TNFSF14* |
| GO:0043067 | Regulation of programmed cell death | 353 | 1.750 | 1.48E-06 | 1.20E-05 | *CXCL10, IFNG, CD274, STAT1, CCL2, S100A8, FASLG, FAS, LOC100522887, THBS1, BCL2L14, LTF, GADD45G, CASP10, TNFAIP3, PIM1, CASP1, ATF3, CCL5, ZFP36, CD40, STAT5A, ACKR3, NOD2, EGR1, HSH2D, BAK1, GLI2, NKX3-1, ANGPTL4, CFLAR, BCL6, FBXO7, JAK3, CASP3, TIGAR, RHOB, BCL2A1, B4GALT1, ITGA5, CEBPB, TRAF3, MYC, AQP1, PLK3, PHLDA2, ST3GAL1, WNT16, RIPK3, FTH1, YBX3, TRAF2, CASP8, SELENOS, ATF5, MCL1, GATA2, JAK2, PPARGC1A, F3, SPRY2, UTP11, GADD45B, HSPA9, PLSCR3, RIPK2, AR, TLR4, TCIM, LIG4, IL10, SLC35F6, PTK2B, RNF122, G0S2, LRP8, DDIT3, PSMD10, TNFRSF12A, ATP2A2, HIGD1A, CXCL12, CTH, SIX4, FFAR4, HMOX1, FOXO1, SLC25A5, NRP1, GRK5, HTATIP2, TCTN3* |
| GO:0051050 | Positive regulation of transport | 149 | 1.734 | 3.35E-04 | 1.37E-03 | *CXCL9, CXCL11, CXCL10, IFNG, CCL2, S100A8, IL1B, ADRA2A, LIPG, CCL5, IL1A, C3, SPP1, IL6, F2RL2, CCR1, FFAR2, IL15, NKX3-1, CADPS2, NFKBIA, CALR, CEBPB, AQP1, PLK3, TLR2* |
| GO:0051240 | Positive regulation of multicellular organismal process | 368 | 1.727 | 2.06E-06 | 1.63E-05 | *RSAD2, ISG15, IFNG, DHX58, IRF7, IRF1, CHI3L1, STAT1, CXCL8, SOCS1, TNFSF13B, CD74, MMP8, THBS1, IL1B, SLA-DQB1, ADRA2A, SLA-DMB, LTF, PIM1, EGR2, LRRTM2, LIPG, CASP1, IRF8, CCL5, KIT, IL1A, CD40, STAT5A, HAVCR2, SPP1, EGR1, SLAMF6, ADM, IL6, STING1, CCR1, FFAR2, IL15, TICAM2, RGS2, SLITRK2, DUSP10, LIF, NKX3-1, CGAS, BCL6, MAD2L2, RHOB, ITGA5, CEBPB, AQP1, MYD88, SLC7A1, TFRC, TNFSF14, PANX2, TLR2, MYMK, HRH2, KLRK1, TSKU, NOS2, YBX3, TRAF2, CASP8, PTN, DDX21, CEBPG,* |

| | | | | | | |
|---|---|---|---|---|---|---|
| | | | | | | *TICAM1, GATA2, JAK2, PPARGC1A, F3, B4GALT5, P2RY2, SMAD1, TMEM106A, TNFSF9, TLR4, FRMD8, SEMA4D, LIG4, IL10, POLR3D, PTK2B, G0S2, IGFBP2, LRP8, DDIT3, TMED10, CMKLR1, RRAS, CXCL12, SIX4, KAT2A, FFAR4, HAPLN1, NRP1, PEAR1, FMR1, TENT5A, RUNX1* |
| GO:0007186 | G protein-coupled receptor signaling pathway | 166 | 1.725 | 3.12E-04 | 1.29E-03 | *CXCL9, CXCL11, CXCL10, ACKR1, CXCL13, CCL4, AMCF-II, CCL2, CXCL8, GNAT1, PTGIR, ADRA2A, GLRB, CCL5, RGS16, C3, CCL8, GPR37, ADM, FFAR2, RGS1, RGS2, PROKR2, ARRDC3* |
| GO:0034220 | Monoatomic ion transmembrane transport | 131 | 1.716 | 5.53E-04 | 2.20E-03 | *CXCL9, CXCL11, CXCL10, CALHM6, CNGA4, ADRA2A, GLRB, ATP1A2, KCNJ2, KCNMA1, BPIFA1, STING1, SLC25A28, KCNA2, NALF2, SLC11A1, PANX2, GP91-PHOX, LRRC8A, SLC39A8, SLC31A1, KCNK7, ATP1B1, TMEM38B, KCNJ14, PTK2B, SLC5A1, DDIT3, SLC30A7, ATP2A2* |
| GO:0009057 | Macromolecule catabolic process | 375 | 1.705 | 3.46E-06 | 2.41E-05 | *ISG20, ISG15, IFNG, PSMB9, CHI3L1, PSMB8, PSMB10, PSME2, ADRA2A, RNF19B, PSME1, TNFAIP3, ZFP36, IGF2BP1, HSP90B1, PSMA5, TIMP1, PSMA2, AGO2, ARID5A, FBXO7, MAD2L2, POP1, PSMA3, CALR, TIMP3, ZFAND2A, PLK3, RNASEH2B, NOS2, TRAF2, CASP8, SELENOS, DERL3, RNF115, DNAJB9, KLHDC3, YOD1, PSMA1, PNPT1, TNFAIP1, OVGP1, IL10, PTK2B, RNF122, USP44, DDIT3, PSMD10, PSMD3, PSMB4, ADAM10, IDE, FOXO1, ETF1, SKIC2, FMR1, TENT5A, UBE2D1, SQSTM1, RLIM, FBXO4, ZFP36L1, PSMB3, LGMN, PSMF1, KLHL15, PGM2, PAQR3, PDE12, PELO, DESI1, TIRAP, LPCAT1, PSMA4, RAD23A, ZPR1, CHMP4A, REXO4, VCP, FOXRED2, NDFIP2, RNF114, CHMP5, EDEM2, PLK2, ZFP36L2, TRAF7, WWP1, CTSB, HYAL2, TENT2, SENP1, UBE2D3, SPATA18, PSMD1, PLAA, STT3B* |
| GO:0030155 | Regulation of cell adhesion | 171 | 1.696 | 4.47E-04 | 1.81E-03 | *IFNG, SELE, CD274, IRF1, CXCL8, RND1, SOCS1, TNFSF13B, THBS1, IL1B, SLA-DQB1, SLA-DMB, CCL5, IL1A, APOD, STAT5A, HAVCR2, LRRC32, IL15, DUSP10, LIF, TNC, MYOC, BCL6, JAK3, MAD2L2, CALR, CEBPB, SLC7A1, TFRC, TNFSF14, FUT2, JAK2, TNFSF9, SEMA4D, IL10, PTK2B, IGFBP2, EPHA2, ADAM10, RRAS, CXCL12, GTPBP4, NRP1* |

| | | | | | | |
|---|---|---|---|---|---|---|
| GO:0048870 | Cell motility | 387 | 1.688 | 6.23E-06 | 4.11E-05 | *CXCL9, CXCL11, CXCL10, IFNG, MADCAM1, CXCL13, SELE, CCL4, AMCF-II, RASGEF1A, CCL2, CXCL8, S100A8, S100A12, THBS1, ADRA2A, CDH12, TNFAIP3, ACKR4, CXCR6, CCL5, DLL4, KIT, CD40, APOD, CCL8, ACKR3, GLUL, CCR1, FFAR2, TREM1, CXCL16, DUSP10, GDF15, MYOC, BCL6, ACKR2, CLDN5, RHOB, CALR, B4GALT1, ITGA5, PLEKHO1, TNFSF14, PHLDA2, KLRK1, RIPK3, SDC4, CASP8, PTN, GATA2, CHGA, SPRY2, ADORA1, TNFAIP1, TLR4, SEMA4D, FSCN1, STC1, PTK2B, AGTR1, LRP8, DDIT3, EPHA2, CMKLR1, DNAAF6, ADAM10, RRAS, ENPEP, CXCL12, SIX4, GNA13, CLDN4, CFAP161, GTPBP4, NRP1, NARS1, SEMA4C, NUS1, MEAK7, XCR1* |
| GO:0006915 | Apoptotic process | 450 | 1.686 | 2.66E-06 | 1.97E-05 | *CXCL10, IFNG, CD274, IRF1, CHI3L1, STAT1, CCL2, S100A8, FASLG, FAS, LOC100522887, THBS1, BCL2L14, LTF, GADD45G, CASP10, TNFAIP3, PIM1, CASP1, ATF3, SHISA5, CCL5, ZFP36, IL1A, LOC100624226, CD40, STAT5A, EGLN3, ACKR3, NOD2, EGR1, HSH2D, BAK1, GLI2, AEN, NKX3-1, ANGPTL4, CFLAR, CSRNP1, BCL6, FBXO7, JAK3, CASP3, TIGAR, TNFRSF1A, RHOB, BCL2A1, B4GALT1, ITGA5, CEBPB, TRAF3, MYC, AQP1, TNFRSF11B, PLK3, PHLDA2, ST3GAL1, WNT16, RIPK3, YBX3, NEK6, TRAF2, CASP8, SELENOS, PLSCR1, ATF5, MCL1, TICAM1, GATA2, JAK2, PPARGC1A, CYCS, F3, SPRY2, UTP11, GADD45B, HSPA9, PLSCR3, RIPK2, AR, TLR4, TCIM, LIG4, IL10, SLC35F6, PTK2B, RNF122, G0S2, LRP8, DDIT3, PSMD10, TNFRSF12A, EPHA2, ATP2A2, HIGD1A, CXCL12, CTH, SIX4, FFAR4, HMOX1, FOXO1, SLC25A5, NRP1, GRK5* |
| GO:1902533 | Positive regulation of intracellular signal transduction | 271 | 1.686 | 4.82E-05 | 2.45E-04 | *SAA3, RSAD2, DHX58, CHI3L1, RASGEF1A, S100A8, FASLG, S100A12, CD74, THBS1, ADRA2A, IFI35, SLC15A3, GADD45G, PIM1, TIFA, CCL5, CD40, HAVCR2, ACKR3, GPR37, CSF2, CCR1, GPR155, TICAM2, LIF, NTRK3, NKX3-1, ARRDC3, GDF15, CFLAR, SESN2, MYOC, CALR, ADAM17, MYD88, TNFSF14, TLR2, WNT16, RIPK3, TRAF2, CASP8, DDX21, MCL1, TICAM1, IKBKB, JAK2, F3, RIPK1, SPRY2, GADD45B, TMEM106A, AR, TLR4, SEMA4D, TCIM, ADRA2B, PTK2B, DDIT3* |
| GO:0070887 | Cellular response to chemical stimulus | 463 | 1.684 | 3.04E-06 | 2.20E-05 | *CXCL9, CXCL11, CXCL10, CXCL13, CCL4, AMCF-II, STAT1, CCL2, CXCL8, SOCS1, S100A8, MMP8, CD180, THBS1, IL1B, ADRA2A, LTF, AQP3, TNFAIP3, ACKR4, PIM1, JUNB, IRF8,* |

| | | | | | | |
|---|---|---|---|---|---|---|
| | | | | | | *CXCR6, CCL5, SLC2A4, KIT, ZFP36, IL1A, CCL8, STAT5A, HAVCR2, SPP1, ACKR3, GPR37, CCR1, FFAR2, AQP9, TREM1, GPR155, MT-2B, NFKBIB, STAT3, CXCL16, NKX3-1, GDF15, CGAS, ARID5A, SESN2, ACKR2, JAK3, NFKBIA, TIGAR, RHOB, CALR, CEBPB, AQP1, ZFAND2A, TNFSF14, KLF9, TLR2, SLC41A1, WNT16, KLRK1, RIPK3, NOS2, YBX3, TRAF2, PLSCR4, PTN, SELENOS, ITPKC, TICAM1, FBP1, CHGA, JAK2, SLC39A8, PPARGC1A, PLSCR3, CYP26B1, AR, TLR4, INSIG1, TMEM38B, IL10, PTK2B, AGTR1, DDIT3, SLC25A33, EPHA2, CMKLR1, ATP2A2, ADAM10, JUN, PID1* |
| GO:0023056 | Positive regulation of signaling | 394 | 1.683 | 8.05E-06 | 4.78E-05 | *SAA3, RSAD2, DHX58, ZBP1, IRF7, CHI3L1, RASGEF1A, CCL2, S100A8, FASLG, S100A12, CD74, MMP8, CD180, THBS1, IL1B, ADRA2A, IFI35, SLC15A3, GADD45G, TNFAIP3, PIM1, LRRTM2, ATF3, TIFA, CCL5, DLL4, KIT, CD40, C3, HAVCR2, SPP1, ACKR3, GPR37, CSF2, ESM1, IL6, F2RL2, CCR1, FFAR2, BVES, GPR155, TICAM2, STAT3, LIF, NTRK3, NKX3-1, ARRDC3, GDF15, CFLAR, SESN2, MYOC, MRAP2, CALR, ADAM17, MYD88, GRIA3, TNFSF14, TLR2, WNT16, RIPK3, TRAF2, CASP8, PTN, DDX21, MCL1, TICAM1, IKBKB, LRRC8A, JAK2, F3, RIPK1, SPRY2, GADD45B, ADORA1, TMEM106A, AR, TLR4, SEMA4D, TCIM, ADRA2B, IL10, PTK2B, G0S2, DDIT3* |
| GO:0010647 | Positive regulation of cell communication | 392 | 1.676 | 5.46E-06 | 3.67E-05 | *SAA3, RSAD2, DHX58, ZBP1, IRF7, CHI3L1, RASGEF1A, CCL2, S100A8, FASLG, S100A12, CD74, MMP8, CD180, THBS1, IL1B, ADRA2A, IFI35, SLC15A3, GADD45G, TNFAIP3, PIM1, LRRTM2, ATF3, TIFA, CCL5, DLL4, KIT, CD40, C3, HAVCR2, SPP1, ACKR3, GPR37, CSF2, ESM1, IL6, F2RL2, CCR1, FFAR2, GPR155, TICAM2, STAT3, LIF, NTRK3, NKX3-1, ARRDC3, GDF15, CFLAR, SESN2, MYOC, MRAP2, CALR, ADAM17, MYD88, GRIA3, TNFSF14, TLR2, WNT16, RIPK3, TRAF2, CASP8, PTN, DDX21, MCL1, TICAM1, IKBKB, LRRC8A, JAK2, F3, RIPK1, SPRY2, GADD45B, ADORA1, TMEM106A, AR, TLR4, SEMA4D, TCIM, ADRA2B, IL10, PTK2B, G0S2, DDIT3* |
| GO:0009967 | Positive regulation of signal transduction | 357 | 1.667 | 2.92E-05 | 1.55E-04 | *SAA3, RSAD2, DHX58, ZBP1, IRF7, CHI3L1, RASGEF1A, S100A8, FASLG, S100A12, CD74, MMP8, CD180, THBS1, ADRA2A, IFI35, SLC15A3, GADD45G, TNFAIP3, PIM1, ATF3, TIFA, CCL5, DLL4, KIT, CD40, C3, HAVCR2, ACKR3, GPR37, CSF2, ESM1, IL6, CCR1, GPR155,* |

| | | | | | | |
|---|---|---|---|---|---|---|
| | | | | | | *TICAM2, STAT3, LIF, NTRK3, NKX3-1, ARRDC3, GDF15, CFLAR, SESN2, MYOC, MRAP2, CALR, ADAM17, MYD88, TNFSF14, TLR2, WNT16, RIPK3, TRAF2, CASP8, DDX21, MCL1, TICAM1, IKBKB, JAK2, F3, RIPK1, SPRY2, GADD45B, TMEM106A, AR, TLR4, SEMA4D, TCIM, ADRA2B, IL10, PTK2B, G0S2, DDIT3* |
| GO:0043065 | Positive regulation of apoptotic process | 130 | 1.662 | 1.27E-03 | 4.52E-03 | *IFNG, CD274, STAT1, CCL2, S100A8, FASLG, THBS1, GADD45G, ATF3, CCL5, CD40, NKX3-1, BCL6, CASP3, TIGAR, RHOB, BCL2A1, B4GALT1, PHLDA2, RIPK3, TRAF2, CASP8, MCL1, JAK2, F3, UTP11, GADD45B, HSPA9, TLR4, IL10, RNF122, G0S2, DDIT3, TNFRSF12A, ATP2A2* |
| GO:0008219 | Cell death | 463 | 1.659 | 7.07E-06 | 4.42E-05 | *CXCL10, IFNG, CD274, IRF1, CHI3L1, STAT1, CCL2, S100A8, FASLG, FAS, LOC100522887, THBS1, BCL2L14, LTF, GADD45G, CASP10, TNFAIP3, PIM1, CASP1, ATF3, SHISA5, CCL5, ZFP36, IL1A, LOC100624226, CD40, STAT5A, EGLN3, ACKR3, NOD2, EGR1, HSH2D, BAK1, GLI2, AEN, MLKL, NKX3-1, ANGPTL4, CFLAR, CSRNP1, BCL6, FBXO7, JAK3, CASP3, TIGAR, TNFRSF1A, RHOB, BCL2A1, B4GALT1, ITGA5, CEBPB, TRAF3, MYC, AQP1, TNFRSF11B, PLK3, PHLDA2, ST3GAL1, WNT16, RIPK3, FTH1, YBX3, NEK6, TRAF2, CASP8, SELENOS, PLSCR1, ATF5, MCL1, TICAM1, GATA2, JAK2, PPARGC1A, CYCS, F3, SPRY2, UTP11, GADD45B, HSPA9, PLSCR3, CYP26B1, RIPK2, AR, TLR4, TCIM, LIG4, IL10, SLC35F6, PTK2B, RNF122, G0S2, LRP8, DDIT3, PSMD10, TNFRSF12A, EPHA2, ATP2A2, HIGD1A, CXCL12, CTH, SIX4, FFAR4, HMOX1, FOXO1, SLC25A5, NRP1, GRK5, HTATIP2* |
| GO:0012501 | Programmed cell death | 463 | 1.659 | 7.07E-06 | 4.42E-05 | *CXCL10, IFNG, CD274, IRF1, CHI3L1, STAT1, CCL2, S100A8, FASLG, FAS, LOC100522887, THBS1, BCL2L14, LTF, GADD45G, CASP10, TNFAIP3, PIM1, CASP1, ATF3, SHISA5, CCL5, ZFP36, IL1A, LOC100624226, CD40, STAT5A, EGLN3, ACKR3, NOD2, EGR1, HSH2D, BAK1, GLI2, AEN, MLKL, NKX3-1, ANGPTL4, CFLAR, CSRNP1, BCL6, FBXO7, JAK3, CASP3, TIGAR, TNFRSF1A, RHOB, BCL2A1, B4GALT1, ITGA5, CEBPB, TRAF3, MYC, AQP1, TNFRSF11B, PLK3, PHLDA2, ST3GAL1, WNT16, RIPK3, FTH1, YBX3, NEK6, TRAF2, CASP8, SELENOS, PLSCR1, ATF5, MCL1, TICAM1, GATA2, JAK2, PPARGC1A, CYCS, F3, SPRY2, UTP11, GADD45B, HSPA9, PLSCR3, CYP26B1, RIPK2, AR, TLR4, TCIM, LIG4, IL10,* |

| | | | | | | |
|---|---|---|---|---|---|---|
| | | | | | | *SLC35F6, PTK2B, RNF122, G0S2, LRP8, DDIT3, PSMD10, TNFRSF12A, EPHA2, ATP2A2, HIGD1A, CXCL12, CTH, SIX4, FFAR4, HMOX1, FOXO1, SLC25A5, NRP1, GRK5, HTATIP2* |
| GO:0051094 | Positive regulation of developmental process | 269 | 1.659 | 7.52E-05 | 3.67E-04 | *CXCL9, ISG15, IFNG, CHI3L1, STAT1, CXCL8, SOCS1, TNFSF13B, THBS1, LTF, PIM1, EGR2, JUNB, LRRTM2, ZFP36, IL1A, CD40, CSF3, STAT5A, ADM, CCR1, BRINP3, GLI2, SLITRK2, STAT3, DUSP10, LIF, CEBPD, GDF15, BCL6, MAD2L2, CLDN5, RHOB, ITGA5, CEBPB, AQP1, TFRC, TNFSF14, YBX3, CASP8, PTN, GATA2, LRRC8A, F3, B4GALT5, C13H21orf91, SMAD1, CYP26B1, TNFSF9, AR, SEMA4D, MIEF1, LIG4, ADRA2B, IL10, PTK2B, AGTR1, LRP8, ZHX3, CMKLR1, RRAS, CXCL12, SCGB3A1, SIX4, FFAR4, HAPLN1, NRP1, FMR1, TENT5A, RUNX1, TGFB3* |
| GO:0031399 | Regulation of protein modification process | 104 | 1.657 | 5.38E-03 | 1.44E-02 | *ISG15, IFNG, CHI3L1, STAT2, TNFAIP3, C3, EGR1, IL15, LIF, ITGA5, GNL3, TICAM1, JAK2, SPRY2, SEMA4D, PSMD10, KAT2A, GTPBP4, NRP1, UBE2D1, SQSTM1, FBXO4* |
| GO:0048878 | Chemical homeostasis | 242 | 1.649 | 2.49E-04 | 1.07E-03 | *CXCL9, CXCL11, CXCL10, LPL, AQP7, HK3, SNX10, ADRA2A, AQP3, ATP1A2, LIPG, CCL5, IL1A, CD40, CP, CCL8, STC2, BPIFA1, IL6, CCR1, FFAR2, MT-2B, HK2, STAT3, ANGPTL4, GCK, SESN2, SOAT1, MYC, AQP1, TFRC, RAB20, SLC41A1, ATP13A5, FTH1, TSKU, LRRC8A, SLC39A8, SLC31A1, ORMDL1, PLSCR3, INSIG1, ATP1B1, TMEM38B, STC1, PTK2B, DDIT3, STIM1, SFTPD, SLC30A7, ATP2A2, ABHD5, FITM2* |
| GO:0072594 | Establishment of protein localization to organelle | 116 | 1.648 | 2.01E-03 | 6.74E-03 | *NEURL3, ISG15, IFNG, TNFAIP3, APOD, STAT3, GDAP1, TXNIP, FBXO7, NFKBIA, ZFAND2A, TOMM40, SEC61A1, IPO4, SRPRA, TIMM8A, DDIT3, TRAM1, SQSTM1, SSR3, KPNB1, IPO13, LOC100157391, SPCS1, KPNA2, ZPR1, CHMP4A* |
| GO:0045597 | Positive regulation of cell differentiation | 170 | 1.646 | 1.12E-03 | 4.01E-03 | *CXCL9, ISG15, IFNG, STAT1, SOCS1, LTF, PIM1, EGR2, JUNB, ZFP36, CSF3, STAT5A, ADM, CCR1, BRINP3, GLI2, STAT3, DUSP10, LIF, CEBPD, BCL6, CLDN5, CEBPB, TNFSF14, CASP8, PTN, GATA2, LRRC8A, SMAD1, CYP26B1, TNFSF9, AR, SEMA4D, LIG4, ADRA2B, IL10, AGTR1, LRP8, ZHX3, CMKLR1, RRAS, FFAR4, HAPLN1, NRP1, FMR1, TENT5A, RUNX1, TGFB3* |

| | | | | | | |
|---|---|---|---|---|---|---|
| GO:0045595 | Regulation of cell differentiation | 312 | 1.639 | 9.50E-05 | 4.41E-04 | *CXCL9, CXCL10, ISG15, IFNG, IRF1, STAT1, IL27, SOCS1, LTF, PIM1, EGR2, JUNB, DLL4, ZFP36, CSF3, MAFF, STAT5A, ADM, PRDM1, CCR1, BRINP3, GLI2, STAT3, DUSP10, LIF, DNAJB11, CEBPD, TMEM176B, BCL6, FBXO7, JAK3, MAD2L2, NFKBIA, CLDN5, CALR, CEBPB, MYC, TNFSF14, TSKU, CASP8, PTN, IKBKB, GATA2, LRRC8A, SPRY2, SMAD1, HSPA9, CYP26B1, TNFSF9, AR, SEMA4D, INSIG1, TCIM, LIG4, ADRA2B, RBPMS2, IL10, PTK2B, AGTR1, LRP8, DDIT3, TMEM182, ZHX3, CMKLR1, RRAS, SIX4, KAT2A, FFAR4, RFLNB, FOXO1, HAPLN1, NRP1, FMR1, TENT5A, RUNX1* |
| GO:0055082 | Intracellular chemical homeostasis | 167 | 1.634 | 8.82E-04 | 3.24E-03 | *CXCL9, CXCL11, CXCL10, HK3, ADRA2A, ATP1A2, CCL5, IL1A, CD40, CP, CCL8, STC2, CCR1, MT-2B, HK2, GCK, MYC, AQP1, TFRC, RAB20, SLC41A1, ATP13A5, FTH1, LRRC8A, SLC39A8, SLC31A1, ORMDL1, ATP1B1, TMEM38B, STC1, PTK2B, DDIT3, STIM1, SLC30A7, ATP2A2, FITM2* |
| GO:0030163 | Protein catabolic process | 266 | 1.632 | 1.48E-04 | 6.51E-04 | *ISG15, IFNG, PSMB9, PSMB8, PSMB10, PSME2, ADRA2A, RNF19B, PSME1, TNFAIP3, HSP90B1, PSMA5, TIMP1, PSMA2, FBXO7, MAD2L2, PSMA3, CALR, TIMP3, ZFAND2A, PLK3, NOS2, TRAF2, CASP8, SELENOS, DERL3, RNF115, DNAJB9, KLHDC3, YOD1, PSMA1, TNFAIP1, IL10, PTK2B, RNF122, USP44, DDIT3, PSMD10, PSMD3, PSMB4, ADAM10, IDE, FOXO1, UBE2D1, SQSTM1, RLIM, FBXO4, PSMB3, LGMN, PSMF1, KLHL15, PAQR3, DESI1, LPCAT1, PSMA4, RAD23A, CHMP4A, VCP, FOXRED2, NDFIP2, RNF114, CHMP5, EDEM2, PLK2, TRAF7, WWP1, CTSB, UBE2D3, SPATA18, PSMD1, PLAA, STT3B* |
| GO:2000026 | Regulation of multicellular organismal development | 306 | 1.631 | 8.46E-05 | 4.02E-04 | *CXCL10, ISG15, IFNG, IRF1, CHI3L1, STAT1, IL27, CXCL8, SERPINE1, SOCS1, THBS1, LTF, PIM1, EGR2, LRRTM2, DLL4, ZFP36, IL1A, CD40, CSF3, MAFF, STAT5A, GLUL, ADM, PRDM1, CCR1, GLI2, SLITRK2, STAT3, DUSP10, LIF, DNAJB11, TMEM176B, BCL6, FBXO7, JAK3, NFKBIA, CLDN5, RHOB, ITGA5, CEBPB, VASH2, MYC, AQP1, PHLDA2, CREB3L1, CASP8, PTN, IKBKB, GATA2, F3, SPRY2, B4GALT5, SMAD1, HSPA9, PPP1R16B, CYP26B1, TNFSF9, SEMA4D, TCIM, LIG4, IL10, PTK2B, LRP8, EPHA2, ADAM10, RRAS, CXCL12, SIX4, KAT2A, RFLNB, HAPLN1, NRP1, HTATIP2, FMR1, TENT5A, RUNX1, DMP1* |

| | | | | | | |
|---|---|---|---|---|---|---|
| GO:0071705 | Nitrogen compound transport | 495 | 1.631 | 6.80E-06 | 4.40E-05 | *NEURL3, RSAD2, IFNG, TAP1, CALHM6, GNAT1, S100A8, RHBDF2, CD74, SNX10, ADRA2A, AQP3, SLC15A3, STX11, EGR2, CCL5, SLC22A4, ZFP36, IL1A, APOD, SLC25A22, IL6, F2RL2, FFAR2, IGF2BP1, RAB3D, OPTN, STAT3, SLC1A5, STXBP1, ARRDC3, CADPS2, TXNIP, SLC7A2, KCNA2, ARL5B, RAB39B, NFKBIA, SELENBP1, RHOB, CALR, SEC23B, AQP1, SLC7A1, TFRC, RAB20, TOMM40, PLK3, TLR2, SLC38A5, SEC61A1, ATP13A5, XPO6, ABCG2, NOS2, IPO4, SELENOS, DERL3, PREB, GPM6B, SRPRA, LRRC8A, CHGA, TIMM8A, SCFD2, HSPA9, IGF2BP2, SEC24A, TLR4, INSIG1, ARF4, SLC18B1, SLC1A3, SLC25A33, TMED10, AP3S2, USO1, SNX19, TRAM1, FFAR4, FOXO1, SLC25A5, STX17, FMR1, RAB11FIP5, NOL6, TMED2, TGFB3, SEC31B, NUP188, SQSTM1, XPO5, SLC38A2, GJA1, SEC22B, TMED5, C2CD2L, KPNB1, IPO13, SLC33A1, DESI1, TP53, TVP23B, PDZK1, LOC100157391, SEC24C, KPNA2, TMED9, ZPR1, COPA* |
| GO:0042592 | Homeostatic process | 401 | 1.622 | 2.47E-05 | 1.33E-04 | *CXCL9, CXCL11, CXCL10, ISG15, STAT1, LPL, TNFSF13B, AQP7, FAS, HK3, SNX10, IL1B, ADRA2A, AQP3, ATP1A2, TNFAIP3, LIPG, CCL5, KIT, ZFP36, IL1A, CD40, CP, CCL8, STC2, STAT5A, SPP1, BPIFA1, EGR1, ST6GALNAC1, IL6, CCR1, FFAR2, MT-2B, HK2, STAT3, SLC1A5, PIWIL4, ARRDC3, GDF15, ANGPTL4, GCK, SESN2, BCL6, CASP3, CLDN5, SOAT1, CEBPB, MYC, AQP1, TFRC, RAB20, DRAM2, SLC41A1, ATP13A5, RIPK3, FTH1, TSKU, SELENOS, CEBPG, GATA2, LRRC8A, JAK2, SLC39A8, PPARGC1A, TXNRD1, SLC31A1, SMAD1, HSPA9, ORMDL1, PLSCR3, TLR4, INSIG1, ATP1B1, TMEM38B, STC1, PTK2B, G0S2, DDIT3, STIM1, SFTPD, SLC30A7, CMKLR1, ATP2A2, ABHD5, IDE, FITM2* |
| GO:0051603 | Proteolysis involved in protein catabolic process | 215 | 1.618 | 7.61E-04 | 2.88E-03 | *ISG15, PSMB9, PSMB8, PSMB10, PSME2, RNF19B, PSME1, TNFAIP3, HSP90B1, PSMA5, PSMA2, FBXO7, PSMA3, CALR, ZFAND2A, CASP8, SELENOS, DERL3, RNF115, DNAJB9, KLHDC3, YOD1, PSMA1, TNFAIP1, PTK2B, RNF122, USP44, DDIT3, PSMD10, PSMD3, PSMB4, IDE, UBE2D1, RLIM, FBXO4, PSMB3, LGMN, PSMF1, KLHL15, PAQR3, DESI1, PSMA4, RAD23A, CHMP4A, VCP, FOXRED2, NDFIP2, RNF114, CHMP5, EDEM2, TRAF7, WWP1, CTSB* |

| | | | | | | |
|---|---|---|---|---|---|---|
| GO:0098771 | Inorganic ion homeostasis | 133 | 1.614 | 2.74E-03 | 8.63E-03 | *CXCL9, CXCL11, CXCL10, SNX10, ATP1A2, CCL5, IL1A, CD40, CP, CCL8, STC2, CCR1, MT-2B, MYC, TFRC, SLC41A1, ATP13A5, FTH1, SLC39A8, SLC31A1, ATP1B1, TMEM38B, STC1, PTK2B, DDIT3, STIM1, SLC30A7, ATP2A2* |
| GO:0098660 | Inorganic ion transmembrane transport | 132 | 1.611 | 2.24E-03 | 7.45E-03 | *CXCL9, CXCL11, CXCL10, ADRA2A, GLRB, ATP1A2, KCNJ2, KCNMA1, BPIFA1, SLC25A28, KCNA2, SLC5A6, NALF2, SLC11A1, AQP1, LRRC8A, SLC39A8, SLC31A1, KCNK7, ATP1B1, TMEM38B, KCNJ14, PTK2B, SLC5A1, DDIT3, SLC30A7, ATP2A2* |
| GO:1902531 | Regulation of intracellular signal transduction | 476 | 1.604 | 9.61E-06 | 5.52E-05 | *SAA3, RSAD2, DHX58, CHI3L1, RASGEF1A, STAT1, S100A8, FASLG, S100A12, FAS, CD74, THBS1, DUSP5, ADRA2A, IFI35, SLC15A3, GADD45G, TNFAIP3, PIM1, ATF3, TIFA, CCL5, CD40, HAVCR2, ACKR3, GPR37, CSF2, CCR1, GPR155, NFKBIB, TICAM2, OPTN, DUSP10, LIF, NTRK3, NKX3-1, ARRDC3, GDF15, CFLAR, CGAS, SESN2, TNIP1, MYOC, BCL6, FBXO7, CHN2, NFKBIA, CALR, ADAM17, TRAF3, MYC, MYD88, TNFSF14, TLR2, RNASEH2B, WNT16, RIPK3, YBX3, TRAF2, CASP8, SELENOS, DDX21, MCL1, TICAM1, IKBKB, FBP1, FICD, JAK2, SLC39A8, F3, RIPK1, SPRY2, GADD45B, PPP1R16B, TMEM106A, CYP26B1, TNFAIP1, CDC42SE1, AR, TLR4, SEMA4D, INSIG1, TCIM, ADRA2B, IL10, PTK2B, USP44, DDIT3, PSMD10, EPHA2, CMKLR1, DUSP4, LTBR, RRAS, CXCL12, CTH, NOP2, NLRP3, FFAR4, HMOX1* |
| GO:0098655 | Monoatomic cation transmembrane transport | 121 | 1.603 | 3.88E-03 | 1.14E-02 | *CXCL9, CXCL11, CXCL10, CNGA4, ADRA2A, ATP1A2, KCNJ2, KCNMA1, BPIFA1, SLC25A28, KCNA2, NALF2, SLC11A1, SLC39A8, SLC31A1, KCNK7, ATP1B1, TMEM38B, KCNJ14, PTK2B, SLC5A1, DDIT3, SLC30A7, ATP2A2* |
| GO:0006811 | Monoatomic ion transport | 220 | 1.600 | 1.40E-03 | 4.83E-03 | *CXCL9, CXCL11, CXCL10, CALHM6, CCL2, SLC39A14, CNGA4, ADRA2A, LTF, GLRB, ATP1A2, KCNJ2, STEAP4, CCL5, KCNMA1, SLC22A4, CCL8, BPIFA1, STING1, CCR1, SLC25A28, KCNA2, SLC5A6, NALF2, SLCO4C1, ATP1B3, SLC11A1, TFRC, PANX2, FTH1, SLC17A7, GP91-PHOX, CD84, LRRC8A, SLC39A8, SLC31A1, SLC11A2, KCNK7, ATP1B1, TMEM38B, KCNJ14, STC1, PTK2B, SLC5A1, DDIT3, STIM1, SLC30A7, SLC39A7, ATP2A2* |

| | | | | | | |
|---|---|---|---|---|---|---|
| GO:0051046 | Regulation of secretion | 116 | 1.599 | 3.61E-03 | 1.07E-02 | *RSAD2, IFNG, S100A8, RHBDF2, IL1B, ADRA2A, CCL5, IL1A, SPP1, IL6, F2RL2, FFAR2, LIF, NKX3-1, CADPS2, KCNA2, AQP1, TLR2, NOS2, CD84, LRRC8A, CHGA, P2RY2, SYN1, ADORA1, TLR4, STC1, TMED10, ATP2A2, FFAR4, FOXO1* |
| GO:0030003 | Intracellular monoatomic cation homeostasis | 133 | 1.597 | 3.36E-03 | 1.03E-02 | *CXCL9, CXCL11, CXCL10, ATP1A2, CCL5, IL1A, CD40, CP, CCL8, STC2, CCR1, MT-2B, MYC, TFRC, RAB20, SLC41A1, ATP13A5, FTH1, SLC39A8, SLC31A1, ATP1B1, TMEM38B, STC1, PTK2B, DDIT3, STIM1, SLC30A7, ATP2A2* |
| GO:0009896 | Positive regulation of catabolic process | 128 | 1.595 | 4.23E-03 | 1.20E-02 | *SAA3, IFNG, ADRA2A, TNFAIP3, ZFP36, STING1, IGF2BP1, OPTN, AGO2, SESN2, FBXO7, ZFAND2A, PLK3, TICAM1, PTK2B, PSMD10, ABHD5, FOXO1, SLC25A5, SQSTM1, ZFP36L1, RALB, PAQR3, PDE12, LPCAT1, ORMDL3, FYCO1, PLK2, ZFP36L2, TRAF7* |
| GO:0051241 | Negative regulation of multicellular organismal process | 230 | 1.593 | 6.12E-04 | 2.39E-03 | *CXCL10, IFNG, DHX58, CD274, IRF1, STAT1, SERPINE1, SOCS1, THBS1, LTF, TNFAIP3, KCNMA1, ZFP36, APOD, HAVCR2, LRRC32, ADM, PARP3, CCR1, RGS2, GLI2, DUSP10, DNAJB11, MORC3, NKX3-1, ARRDC3, TMEM176B, GDF15, BCL6, FBXO7, JAK3, MAD2L2, ELF4, CLDN5, CALR, CEBPB, MYC, TSKU, YBX3, CREB3L1, PTN, CD84, GATA2, SPRY2, HSPA9, ADORA1, TLR4, STC1, IL10, PTK2B, DDIT3, CD200, EPHA2, CMKLR1, ATP2A2* |
| GO:0006873 | Intracellular monoatomic ion homeostasis | 134 | 1.593 | 4.77E-03 | 1.31E-02 | *CXCL9, CXCL11, CXCL10, ATP1A2, CCL5, IL1A, CD40, CP, CCL8, STC2, CCR1, MT-2B, MYC, TFRC, RAB20, SLC41A1, ATP13A5, FTH1, SLC39A8, SLC31A1, ATP1B1, TMEM38B, STC1, PTK2B, DDIT3, STIM1, SLC30A7, ATP2A2* |
| GO:0043068 | Positive regulation of programmed cell death | 134 | 1.587 | 5.39E-03 | 1.44E-02 | *IFNG, CD274, STAT1, CCL2, S100A8, FASLG, THBS1, GADD45G, ATF3, CCL5, CD40, NKX3-1, BCL6, CASP3, TIGAR, RHOB, BCL2A1, B4GALT1, PHLDA2, RIPK3, TRAF2, CASP8, MCL1, JAK2, F3, UTP11, GADD45B, HSPA9, TLR4, IL10, RNF122, G0S2, DDIT3, TNFRSF12A, ATP2A2, FOXO1, HTATIP2, TCTN3* |
| GO:0046903 | Secretion | 189 | 1.586 | 3.52E-03 | 1.05E-02 | *RSAD2, IFNG, GNAT1, S100A8, RHBDF2, SNX10, IL1B, ADRA2A, STX11, CCL5, KIT, IL1A, CCL8, STAT5A, SPP1, IL6, F2RL2, CCR1, FFAR2, LIF, NKX3-1, CADPS2, KCNA2, AQP1, TLR2, HRH2, NOS2, CD84, LRRC8A, CHGA, P2RY2, SYN1, ADORA1, TLR4, TMEM38B, STC1, TMED10, ATP2A2, SNX19, FFAR4, FOXO1* |

| | | | | | | |
|---|---|---|---|---|---|---|
| GO:0030335 | Positive regulation of cell migration | 133 | 1.585 | 3.99E-03 | 1.16E-02 | *CXCL10, IFNG, MADCAM1, CCL4, CCL2, THBS1, ADRA2A, CCL5, KIT, CD40, CCL8, CCR1, CXCL16, MYOC, RHOB, CALR, TNFSF14, CASP8, PTN, GATA2, SPRY2, TLR4, SEMA4D, PTK2B, EPHA2, CMKLR1, ADAM10, RRAS, CXCL12, CLDN4, NRP1, NUS1* |
| GO:0098662 | Inorganic cation transmembrane transport | 122 | 1.585 | 4.06E-03 | 1.18E-02 | *CXCL9, CXCL11, CXCL10, ADRA2A, ATP1A2, KCNJ2, KCNMA1, BPIFA1, SLC25A28, KCNA2, NALF2, SLC11A1, AQP1, SLC39A8, SLC31A1, KCNK7, ATP1B1, TMEM38B, KCNJ14, PTK2B, SLC5A1, DDIT3, SLC30A7, ATP2A2* |
| GO:0019725 | Cellular homeostasis | 199 | 1.584 | 1.40E-03 | 4.83E-03 | *CXCL9, CXCL11, CXCL10, HK3, SNX10, ADRA2A, ATP1A2, CCL5, IL1A, CD40, CP, CCL8, STC2, CCR1, MT-2B, HK2, GCK, MYC, AQP1, TFRC, RAB20, SLC41A1, ATP13A5, FTH1, SELENOS, LRRC8A, SLC39A8, TXNRD1, SLC31A1, ORMDL1, ATP1B1, TMEM38B, STC1, PTK2B, DDIT3, STIM1, SLC30A7, ATP2A2, IDE, FITM2* |
| GO:0051246 | Regulation of protein metabolic process | 316 | 1.575 | 3.13E-04 | 1.29E-03 | *ISG15, IFNG, CHI3L1, SERPINE1, PSME2, THBS1, ADRA2A, PSME1, STAT2, TNFAIP3, LIPG, ZFP36, APOD, C3, EGR1, IL15, IGF2BP1, RGS2, TIMP1, LIF, PIWIL4, AGO2, SESN2, MAD2L2, CASP3, NFKBIA, SOAT1, CALR, ITGA5, TIMP3, ZFAND2A, PLK3, GNL3, NOS2, CASP8, CD84, TICAM1, JAK2, SPRY2, IGF2BP2, EIF5A, SEMA4D, IL10, PTK2B, AGTR1, PSMD10, TMED10, PSMD3, KAT2A, FOXO1, GTPBP4, ETF1, NRP1, FMR1, STK35, UBE2D1, SQSTM1, FGFR4, FBXO4, OGFOD1* |
| GO:0050801 | Monoatomic ion homeostasis | 148 | 1.568 | 4.15E-03 | 1.19E-02 | *CXCL9, CXCL11, CXCL10, SNX10, ATP1A2, CCL5, IL1A, CD40, CP, CCL8, STC2, CCR1, MT-2B, MYC, TFRC, RAB20, SLC41A1, ATP13A5, FTH1, SLC39A8, SLC31A1, ATP1B1, TMEM38B, STC1, PTK2B, DDIT3, STIM1, SLC30A7, ATP2A2* |
| GO:0055080 | Monoatomic cation homeostasis | 147 | 1.567 | 4.26E-03 | 1.20E-02 | *CXCL9, CXCL11, CXCL10, SNX10, ATP1A2, CCL5, IL1A, CD40, CP, CCL8, STC2, CCR1, MT-2B, MYC, TFRC, RAB20, SLC41A1, ATP13A5, FTH1, SLC39A8, SLC31A1, ATP1B1, TMEM38B, STC1, PTK2B, DDIT3, STIM1, SLC30A7, ATP2A2* |
| GO:0098609 | Cell-cell adhesion | 182 | 1.560 | 2.49E-03 | 8.07E-03 | *IFNG, MADCAM1, SELE, CD274, IRF1, SOCS1, TNFSF13B, S100A8, IL1B, SLA-DQB1, SLA-DMB, CDH12, CLSTN3, VCAM1, CCL5, KIT, IL1A, STAT5A, HAVCR2, OLR1, LRRC32, BVES, IL15, SLITRK2, DUSP10, BCL6, JAK3, MAD2L2, ITGA5, CEBPB, SLC7A1, TFRC, TNFSF14, JAK2, SLC39A8, ICAM1, TNFSF9, SEMA4D, ENTPD1, IL10, IGFBP2* |

| | | | | | | |
|---|---|---|---|---|---|---|
| GO:0006812 | Monoatomic cation transport | 196 | 1.557 | 1.94E-03 | 6.57E-03 | *CXCL9, CXCL11, CXCL10, CCL2, SLC39A14, CNGA4, ADRA2A, LTF, ATP1A2, KCNJ2, STEAP4, CCL5, KCNMA1, SLC22A4, CCL8, BPIFA1, CCR1, SLC25A28, KCNA2, SLC5A6, NALF2, ATP1B3, SLC11A1, TFRC, PANX2, FTH1, SLC17A7, CD84, SLC39A8, SLC31A1, SLC11A2, KCNK7, ATP1B1, TMEM38B, KCNJ14, STC1, PTK2B, SLC5A1, DDIT3, STIM1, SLC30A7, SLC39A7, ATP2A2* |
| GO:0030001 | Metal ion transport | 182 | 1.552 | 2.79E-03 | 8.71E-03 | *CXCL9, CXCL11, CXCL10, CCL2, SLC39A14, ADRA2A, LTF, ATP1A2, KCNJ2, STEAP4, CCL5, KCNMA1, SLC22A4, CCL8, BPIFA1, CCR1, SLC25A28, KCNA2, SLC5A6, NALF2, ATP1B3, SLC11A1, TFRC, FTH1, SLC17A7, CD84, SLC39A8, SLC31A1, SLC11A2, KCNK7, ATP1B1, TMEM38B, KCNJ14, STC1, PTK2B, SLC5A1, DDIT3, STIM1, SLC30A7, SLC39A7, ATP2A2* |
| GO:0040017 | Positive regulation of locomotion | 143 | 1.542 | 5.67E-03 | 1.47E-02 | *CXCL10, IFNG, MADCAM1, CCL4, CCL2, CXCL8, THBS1, ADRA2A, CCL5, KIT, CD40, CCL8, CCR1, BVES, CXCL16, MYOC, BCL6, RHOB, CALR, TNFSF14, CASP8, PTN, GATA2, SPRY2, TLR4, SEMA4D, PTK2B, EPHA2, CMKLR1, ADAM10, RRAS, CXCL12, CLDN4, NRP1* |
| GO:0048585 | Negative regulation of response to stimulus | 401 | 1.542 | 1.72E-04 | 7.45E-04 | *ISG15, DHX58, STAT1, CCL2, CXCL8, SERPINE1, SOCS1, SOCS3, BPIFB1, DKK1, THBS1, DUSP5, PTGIR, ADRA2A, LTF, IFI35, STAT2, TNFAIP3, PIM1, ATF3, DLL4, ZFP36, RGS16, APOD, HAVCR2, SERPING1, ACKR3, EGR1, ADM, PARP3, GPR155, RGS1, RGS2, OPTN, GLI2, DUSP10, LIF, CISH, NKX3-1, ARRDC3, GDF15, CFLAR, CGAS, SESN2, MYOC, BCL6, FBXO7, JAK3, MAD2L2, ELF4, NFKBIA, MRAP2, CALR, MYC, WNT16, KLRK1, TSKU, APC2, YBX3, CREB3L1, CASP8, SELENOS, MCL1, RNF115, GATA2, FBP1, SLC39A8, SPRY2, ADORA1, CYP26B1, TNFAIP1, AR, TLR4, INSIG1, TCIM, IL10, SLC35F6, DDIT3, PSMD10, SOCS7, CD59, PSMB4, DUSP4, PID1, CXCL12, NLRP3, FFAR4, HMOX1, SLC25A5, NRP1, PAK1IP1* |
| GO:0010629 | Negative regulation of gene expression | 279 | 1.531 | 9.27E-04 | 3.37E-03 | *SAA3, IFNG, DHX58, CXCL8, SERPINE1, THBS1, LTF, TNFAIP3, DLL4, MOV10, ZFP36, APOD, HAVCR2, LRRC32, PARP3, PRDM1, CCR1, IGF2BP1, PIWIL4, MORC3, NKX3-1, AGO2, ARID5A, SESN2, BCL6, ELF4, CLDN5, CALR, BAZ1A, LOC100624590, RNASEH2B, TSKU, NOS2, CREB3L1, TRAF2, CD84, GATA2, PNPT1, TLR4, IL10, DDIT3, EPHA2,* |

| | | | | | | |
|---|---|---|---|---|---|---|
| | | | | | | *CMKLR1, ADAM10, SETDB2, FFAR4, FOXO1, ETF1, FMR1, TENT5A, TGFB3, RLIM, ZFP36L1, GJA1, SOX11, EIF4EBP1, PDE12, PELO, TIRAP* |
| GO:0060284 | Regulation of cell development | 177 | 1.530 | 3.42E-03 | 1.04E-02 | *ISG15, IFNG, IRF1, STAT1, IL27, SOCS1, LTF, EGR2, DLL4, ZFP36, CSF3, STAT5A, PRDM1, CCR1, STAT3, DUSP10, LIF, DNAJB11, TMEM176B, BCL6, FBXO7, JAK3, NFKBIA, CLDN5, CEBPB, MYC, CASP8, PTN, IKBKB, GATA2, HSPA9, CYP26B1, TNFSF9, SEMA4D, TCIM, LIG4, IL10, PTK2B, LRP8, KAT2A, RFLNB, HAPLN1, NRP1, FMR1, RUNX1* |
| GO:0010648 | Negative regulation of cell communication | 352 | 1.529 | 7.25E-04 | 2.78E-03 | *ISG15, DHX58, STAT1, CXCL8, SOCS1, SOCS3, BPIFB1, DKK1, THBS1, DUSP5, PTGIR, ADRA2A, LTF, IFI35, STAT2, TNFAIP3, ATF3, DLL4, RGS16, APOD, ACKR3, EGR1, ADM, FFAR2, GPR155, RGS1, RGS2, OPTN, GLI2, DUSP10, LIF, CISH, NKX3-1, ARRDC3, GDF15, CFLAR, CGAS, SESN2, MYOC, BCL6, FBXO7, MAD2L2, NFKBIA, MRAP2, CALR, MYC, WNT16, TSKU, APC2, YBX3, CREB3L1, CASP8, SELENOS, MCL1, RNF115, GATA2, FBP1, CHGA, SLC39A8, SPRY2, ADORA1, CYP26B1, TNFAIP1, AR, TLR4, INSIG1, TCIM, IL10, SLC35F6, DDIT3, PSMD10, SOCS7, DUSP4, PID1, CXCL12, NLRP3, FFAR4, HMOX1, FOXO1, SLC25A5, NRP1, PAK1IP1* |
| GO:0051049 | Regulation of transport | 313 | 1.526 | 1.39E-03 | 4.83E-03 | *CXCL9, CXCL11, CXCL10, RSAD2, IFNG, CCL2, S100A8, RHBDF2, THBS1, IL1B, ADRA2A, LRRTM2, LIPG, KCNJ2, CCL5, IL1A, APOD, C3, SPP1, BPIFA1, IL6, F2RL2, CCR1, FFAR2, IL15, SIGLEC1, LIF, NKX3-1, CADPS2, KCNA2, NFKBIA, CALR, CEBPB, AQP1, PLK3, TLR2, NOS2, DERL3, GPM6B, CD84, RUBCN, LRRC8A, CHGA, P2RY2, SYN1, ADORA1, TLR4, INSIG1, ATP1B1, TMEM38B, KCNJ14, STC1, SLC35F6, PTK2B, STIM1, TMED10, SFTPD* |
| GO:1903530 | Regulation of secretion by cell | 107 | 1.525 | 1.48E-02 | 3.42E-02 | *RSAD2, IFNG, RHBDF2, IL1B, ADRA2A, CCL5, IL1A, SPP1, IL6, F2RL2, FFAR2, LIF, NKX3-1, CADPS2, KCNA2, TLR2, NOS2, CD84, LRRC8A, CHGA, SYN1, TLR4, TMED10, ATP2A2, FFAR4, FOXO1* |
| GO:0034655 | Nucleobase-containing compound catabolic process | 148 | 1.524 | 6.70E-03 | 1.73E-02 | *ISG20, SAMHD1, HK3, ZFP36, CNP, IGF2BP1, HK2, AGO2, GCK, ARID5A, DPYS, POP1, TIGAR, RNASEH2B, TRAF2, FBP1, ALDOC, PNPT1, ENTPD1, PGK1, FITM2, PDE4A, ETF1, SKIC2, FMR1, TENT5A, ZFP36L1, PDE12, PELO, ZBTB7A, TIRAP, ZPR1, PDE4B, REXO4, NUDT7, DDIT4, NT5C, DCTPP1, ZFP36L2, TENT2, SENP1* |

| | | | | | | |
|---|---|---|---|---|---|---|
| GO:0008285 | Negative regulation of cell population proliferation | 156 | 1.523 | 7.87E-03 | 2.00E-02 | *IFNG, CD274, STAT1, CCL2, THBS1, PTGIR, IFI35, TNFAIP3, IFI30, DLL4, IL1A, APOD, HAVCR2, ACKR3, LRRC32, DUSP10, LIF, MORC3, NKX3-1, BCL6, B4GALT1, CEBPB, KLF9, PTN, ATF5, GATA2, JAK2, PPARGC1A, SPRY2, SMAD1, AR, IL10, PTK2B* |
| GO:0045596 | Negative regulation of cell differentiation | 143 | 1.522 | 8.26E-03 | 2.09E-02 | *CXCL10, IRF1, STAT1, SOCS1, LTF, ZFP36, STAT5A, GLI2, DNAJB11, TMEM176B, BCL6, FBXO7, JAK3, MAD2L2, NFKBIA, CALR, MYC, TSKU, PTN, GATA2, SPRY2, SMAD1, HSPA9, SEMA4D, INSIG1, RBPMS2, PTK2B, DDIT3, TMEM182, SIX4, RFLNB, FOXO1, NRP1* |
| GO:0030334 | Regulation of cell migration | 209 | 1.520 | 4.34E-03 | 1.22E-02 | *CXCL10, IFNG, MADCAM1, SELE, CCL4, CCL2, THBS1, ADRA2A, CCL5, DLL4, KIT, CD40, APOD, CCL8, GLUL, CCR1, CXCL16, DUSP10, GDF15, MYOC, CLDN5, RHOB, CALR, TNFSF14, PHLDA2, KLRK1, RIPK3, CASP8, PTN, GATA2, SPRY2, ADORA1, TLR4, SEMA4D, STC1, PTK2B, EPHA2, CMKLR1, ADAM10, RRAS, CXCL12, GNA13, CLDN4, GTPBP4, NRP1, NUS1, MEAK7* |
| GO:0061061 | Muscle structure development | 141 | 1.518 | 9.71E-03 | 2.33E-02 | *CXCL9, CXCL10, MYBPH, MSC, UNC45B, EGR2, ATF3, DLL4, MAFF, EGR1, ADM, BVES, LIF, GDF15, PLEKHO1, TNFSF14, MYMK, YBX3, LRRC8A, SMAD1, MYOCD, CYP26B1, RBPMS2, DDIT3, TMEM182, ATP2A2, SCGB3A1, SIX4* |
| GO:0023057 | Negative regulation of signaling | 354 | 1.514 | 8.06E-04 | 2.99E-03 | *ISG15, DHX58, STAT1, CXCL8, SOCS1, SOCS3, BPIFB1, DKK1, THBS1, DUSP5, PTGIR, ADRA2A, LTF, IFI35, STAT2, TNFAIP3, ATF3, DLL4, RGS16, APOD, ACKR3, EGR1, ADM, FFAR2, GPR155, RGS1, RGS2, OPTN, GLI2, DUSP10, LIF, CISH, NKX3-1, ARRDC3, GDF15, CFLAR, CGAS, SESN2, MYOC, BCL6, FBXO7, MAD2L2, NFKBIA, MRAP2, CALR, MYC, WNT16, TSKU, APC2, YBX3, CREB3L1, CASP8, SELENOS, MCL1, RNF115, GATA2, FBP1, CHGA, SLC39A8, SPRY2, ADORA1, CYP26B1, TNFAIP1, AR, TLR4, INSIG1, TCIM, IL10, SLC35F6, DDIT3, PSMD10, SOCS7, DUSP4, PID1, CXCL12, NLRP3, FFAR4, HMOX1, FOXO1, SLC25A5, NRP1, PAK1IP1* |
| GO:0048646 | Anatomical structure formation involved in morphogenesis | 275 | 1.514 | 1.72E-03 | 5.90E-03 | *CXCL9, CXCL10, CHI3L1, STAT1, CXCL8, SERPINE1, TNFSF13B, MYBPH, MMP8, THBS1, DUSP5, TNFAIP3, PIM1, JUNB, DLL4, IL1A, CD40, ACKR3, GLUL, ESM1, ADM, GLI2, NKX3-1, GDF15, ANGPTL4, BCL6, CHN2, CLDN5, RHOB, B4GALT1, ITGA5, VASH2, AQP1, PLEKHO1, TNFSF14, MYMK, GARIN1A, CREB3L1, SAT1, GATA2, F3, SPRY2, SLC31A1,* |

| | | | | | | |
|---|---|---|---|---|---|---|
| | | | | | | *SMAD1, PPP1R16B, AR, IL10, PTK2B, AGTR1, LRP8, TMEM182, EPHA2, RRAS, SCGB3A1, SIX4, GNA13, KAT2A, NRP1, HTATIP2, SEMA4C, DMP1, TBC1D20* |
| GO:0036211 | Protein modification process | 430 | 1.502 | 3.05E-04 | 1.29E-03 | *ISG15, IFNG, UBA7, USP18, CHI3L1, SOCS1, SOCS3, GALNT15, PARP12, TRIM21, RNF19B, STAT2, TNFAIP3, EGR2, C3, EGR1, GLUL, ST6GALNAC1, MGAT4C, PARP3, IL15, PARP11, LIF, CISH, FBXO7, B4GALT1, ITGA5, USP15, POMGNT2, GNL3, ST3GAL1, FUT2, NOS2, NEK6, TRAF2, DERL3, TICAM1, PADI4, RNF115, MOGS, ALG3, JAK2, SLC39A8, WSB2, SPRY2, B4GALT5, ASB5, TNFAIP1, SEMA4D, RNF122, USP44, LOC100625207, PSMD10, SOCS7, B3GAT2, USP2, RPN2, CTH, KAT2A, RPN1, USP51, FOXO1, GTPBP4, NRP1, B3GNT5, NUS1, UBE2D1, SQSTM1, RLIM, FBXO4, OSTC, B4GALT3, DCAF11, ST8SIA4, KLHL15, MGAT1, DESI1, MPDU1, GALNT1, VCP, NDFIP2, WBP1L, RNF114, RNF138, CRY1, MAGT1, EDEM2, TRIM27, OTUB2, FUT8, TRAF7, WWP1, PPP2R2A, ABL1, NPC1, SENP1, KLHL25, ST8SIA2, UBE2D3, ST6GALNAC4, PLAA, CD80, STT3B* |
| GO:0032940 | Secretion by cell | 162 | 1.502 | 9.32E-03 | 2.27E-02 | *RSAD2, IFNG, GNAT1, RHBDF2, IL1B, ADRA2A, STX11, CCL5, KIT, IL1A, CCL8, SPP1, IL6, F2RL2, CCR1, FFAR2, LIF, NKX3-1, CADPS2, KCNA2, TLR2, NOS2, CD84, LRRC8A, CHGA, SYN1, TLR4, TMEM38B, TMED10, ATP2A2, SNX19, FFAR4, FOXO1* |
| GO:0009968 | Negative regulation of signal transduction | 333 | 1.498 | 2.37E-03 | 7.75E-03 | *ISG15, DHX58, STAT1, CXCL8, SOCS1, SOCS3, BPIFB1, DKK1, THBS1, DUSP5, PTGIR, LTF, IFI35, STAT2, TNFAIP3, ATF3, DLL4, RGS16, APOD, ACKR3, EGR1, ADM, GPR155, RGS1, RGS2, OPTN, GLI2, DUSP10, LIF, CISH, NKX3-1, ARRDC3, GDF15, CFLAR, CGAS, SESN2, MYOC, BCL6, FBXO7, MAD2L2, NFKBIA, MRAP2, CALR, MYC, WNT16, TSKU, APC2, YBX3, CREB3L1, CASP8, SELENOS, MCL1, RNF115, GATA2, FBP1, SLC39A8, SPRY2, CYP26B1, TNFAIP1, AR, TLR4, INSIG1, TCIM, IL10, SLC35F6, DDIT3, PSMD10, SOCS7, DUSP4, PID1, CXCL12, NLRP3, HMOX1, SLC25A5, NRP1, PAK1IP1* |
| GO:0051247 | Positive regulation of protein metabolic process | 172 | 1.494 | 6.86E-03 | 1.76E-02 | *IFNG, CHI3L1, THBS1, ADRA2A, TNFAIP3, C3, EGR1, IL15, IGF2BP1, LIF, CASP3, NFKBIA, SOAT1, ITGA5, ZFAND2A, PLK3, GNL3, CASP8, TICAM1, JAK2, EIF5A, SEMA4D, PTK2B, AGTR1, PSMD10, FOXO1, NRP1, FMR1, UBE2D1, FGFR4, FBXO4, PAQR3, LPCAT1,* |

| | | | | | | |
|---|---|---|---|---|---|---|
| | | | | | | *NDFIP2, WBP1L, CRY1, PLK2, TRAF7, NAT10, ABL1, CD80, INHBA, RPUSD3, MTOR, METTL3, HBEGF, PRKAA2, TIPARP, RPS6KB1, MDM2, GUF1, DCUN1D3, CCDC22* |
| GO:0097190 | Apoptotic signaling pathway | 153 | 1.491 | 5.41E-03 | 1.44E-02 | *IFNG, S100A8, FASLG, THBS1, TNFAIP3, ATF3, IL1A, ACKR3, AEN, NKX3-1, CFLAR, FBXO7, BCL2A1, CEBPB, MYC, WNT16, RIPK3, YBX3, TRAF2, CASP8, SELENOS, MCL1, TICAM1, JAK2, PLSCR3, AR, TLR4, IL10, SLC35F6, G0S2, DDIT3, PSMD10, EPHA2, CXCL12, HMOX1, SLC25A5, NRP1* |
| GO:0043412 | Macromolecule modification | 499 | 1.490 | 1.46E-04 | 6.48E-04 | *ISG15, IFNG, UBA7, USP18, CHI3L1, SOCS1, SOCS3, GALNT15, PARP12, TRIM21, RNF19B, STAT2, TNFAIP3, EGR2, C3, EGR1, GLUL, ST6GALNAC1, MGAT4C, PARP3, IL15, PARP11, LIF, CISH, DNAJB11, FBXO7, B4GALT1, ITGA5, USP15, POMGNT2, GNL3, ST3GAL1, FUT2, QTRT2, NOS2, NEK6, TRAF2, DERL3, TICAM1, PADI4, RNF115, MOGS, ALG3, JAK2, SLC39A8, WSB2, SPRY2, B4GALT5, ASB5, TNFAIP1, WDR4, SEMA4D, RNF122, USP44, LOC100625207, PSMD10, SOCS7, B3GAT2, USP2, RPN2, CTH, FBL, KAT2A, RPN1, NOP2, USP51, FOXO1, GTPBP4, NRP1, B3GNT5, NUS1, UBE2D1, SQSTM1, RLIM, FBXO4, QTRT1, PUS3, TRMT6, DKC1, OSTC, B4GALT3, DCAF11, ST8SIA4, KLHL15, MGAT1, DESI1, EMG1, MPDU1, GALNT1, VCP, NDFIP2, APOBEC2, WBP1L, RNF114, RNF138, CRY1, MAGT1, EDEM2, TRIM27, NSUN4, OTUB2, FUT8, TRAF7, WWP1, NAT10, PPP2R2A, BUD23, NSUN2, NSUN3, ABL1, NPC1, SENP1, KLHL25, ST8SIA2, UBE2D3, ST6GALNAC4, PLAA, CD80, STT3B, B3GALT6, A4GALT, RPUSD3, PUS1, NCOA7, RBBP6, ATM, TGS1, USP43, MTOR, METTL3, CBX4* |
| GO:0032879 | Regulation of localization | 413 | 1.479 | 7.79E-04 | 2.92E-03 | *CXCL9, CXCL11, CXCL10, RSAD2, IFNG, CCL2, S100A8, RHBDF2, THBS1, IL1B, ADRA2A, LRRTM2, LIPG, KCNJ2, SLC5A3, CCL5, ZFP36, IL1A, APOD, C3, SPP1, BPIFA1, IL6, F2RL2, PLIN2, CCR1, FFAR2, IL15, SIGLEC1, LIF, NKX3-1, CADPS2, SESN2, KCNA2, NFKBIA, MRAP2, CALR, CEBPB, AQP1, PLK3, TLR2, GNL3, NOS2, DERL3, ABHD17B, GPM6B, CD84, RUBCN, LRRC8A, CHGA, P2RY2, SYN1, ADORA1, AR, TLR4, INSIG1, ATP1B1, TMEM38B, KCNJ14, STC1, SLC35F6, PTK2B, DDIT3, STIM1, TMED10, SFTPD, EPHA2, ATP2A2,* |

| | | | | | | |
|---|---|---|---|---|---|---|
| | | | | | | *ADAM10, ABHD5, PID1, FITM2, SIX4, FFAR4, FOXO1, SLC25A5, FMR1, PPARD, NUS1, TBC1D20, MEAK7, XCR1, TGFB3* |
| GO:2000147 | Positive regulation of cell motility | 139 | 1.479 | 9.53E-03 | 2.31E-02 | *CXCL10, IFNG, MADCAM1, CCL4, CCL2, THBS1, ADRA2A, CCL5, KIT, CD40, CCL8, CCR1, CXCL16, MYOC, BCL6, RHOB, CALR, TNFSF14, CASP8, PTN, GATA2, SPRY2, TLR4, SEMA4D, PTK2B, EPHA2, CMKLR1, ADAM10, RRAS, CXCL12, CLDN4, NRP1, NUS1* |
| GO:0040012 | Regulation of locomotion | 233 | 1.477 | 2.31E-03 | 7.60E-03 | *CXCL10, IFNG, MADCAM1, SELE, CCL4, CCL2, CXCL8, THBS1, ADRA2A, CCL5, DLL4, KIT, CD40, APOD, CCL8, GLUL, CCR1, BVES, CXCL16, DUSP10, ARRDC3, GDF15, MYOC, BCL6, FBXO7, CLDN5, RHOB, CALR, TNFSF14, PHLDA2, KLRK1, RIPK3, CASP8, PTN, GATA2, SPRY2, ADORA1, TLR4, SEMA4D, STC1, PTK2B, EPHA2, CMKLR1, ADAM10, RRAS, CXCL12, GNA13, CLDN4, GTPBP4, NRP1, NUS1, MEAK7* |
| GO:0141124 | Intracellular signaling cassette | 454 | 1.471 | 7.12E-04 | 2.76E-03 | *SAA3, CHI3L1, RASGEF1A, STAT1, RND1, FASLG, S100A12, FAS, CD74, THBS1, DUSP5, ADRA2A, IFI35, GADD45G, TNFAIP3, ACKR4, ATF3, TIFA, CXCR6, CCL5, ZFP36, CD40, HAVCR2, ACKR3, GPR37, CSF2, CCR1, NFKBIB, TICAM2, OPTN, DUSP10, LIF, NTRK3, NKX3-1, GDF15, CFLAR, CGAS, SESN2, TNIP1, MYOC, BCL6, ACKR2, RAB39B, CHN2, NFKBIA, RHOB, CALR, ADAM17, TRAF3, SLC11A1, MYC, AQP1, MYD88, RHOG, TNFSF14, TLR2, WNT16, RIPK3, RASSF1, CDC42EP3, TRAF2, CASP8, DDX21, RHOJ, TICAM1, IKBKB, FBP1, JAK2, SLC39A8, RIPK1, SPRY2, SMAD1, GADD45B, PPP1R16B, TMEM106A, TNFAIP1, CDC42SE1, AR, TLR4, SEMA4D, TCIM, ATP1B1, ADRA2B, TMEM38B, PTK2B, AGTR1, DDIT3, PSMD10, EPHA2, CMKLR1, DUSP4, LTBR, ATP2A2, RRAS, CTH, GNA13, NLRP3, FFAR4, HMOX1, NRP1, PEAR1, SEMA4C, RASGEF1B, XCR1, SQSTM1, CDC42EP1, FGFR4* |
| GO:0051248 | Negative regulation of protein metabolic process | 103 | 1.469 | 2.00E-02 | 4.43E-02 | *ISG15, SERPINE1, THBS1, TNFAIP3, ZFP36, APOD, IGF2BP1, TIMP1, AGO2, SESN2, MAD2L2, CALR, TIMP3, NOS2, CD84, SPRY2* |

| | | | | | | |
|---|---|---|---|---|---|---|
| GO:0022603 | Regulation of anatomical structure morphogenesis | 188 | 1.456 | 1.22E-02 | 2.87E-02 | *CXCL10, CHI3L1, STAT1, CCL2, CXCL8, SERPINE1, TNFSF13B, THBS1, STAT2, TNFAIP3, KIT, IL1A, CD40, GLUL, ADM, BVES, LIF, CLDN5, RHOB, ITGA5, VASH2, AQP1, PLEKHO1, CDC42EP3, CREB3L1, GATA2, SLC26A5, F3, SPRY2, SMAD1, PPP1R16B, CDC42SE1, AR, SEMA4D, MIEF1, IL10, PTK2B, LRP8, EPHA2, RRAS, FITM2, SIX4, GNA13, CLDN4, NRP1, HTATIP2* |
| GO:0033365 | Protein localization to organelle | 237 | 1.454 | 4.53E-03 | 1.25E-02 | *NEURL3, ISG15, IFNG, SNX10, TNFAIP3, APOD, PARP3, OPTN, STAT3, MORC3, GDAP1, TXNIP, SESN2, ARL5B, FBXO7, NFKBIA, CALR, ZFAND2A, TOMM40, GNL3, SEC61A1, IPO4, ABHD17B, SRPRA, TIMM8A, INSIG1, DDIT3, TMED10, ADAM10, SIX4, TRAM1, NRP1, MEAK7, SQSTM1, RRS1, FBXO4, SSR3, KPNB1, IPO13, PAQR3, ZBTB7A, LOC100157391, SPCS1, KPNA2, ZPR1, CHMP4A, VCP, TERF2IP, ORMDL3, HSPA1L* |
| GO:0000165 | MAPK cascade | 169 | 1.451 | 1.61E-02 | 3.68E-02 | *SAA3, CHI3L1, FAS, DUSP5, ADRA2A, GADD45G, ATF3, ZFP36, CD40, HAVCR2, ACKR3, GPR37, CSF2, CCR1, DUSP10, LIF, GDF15, MYOC, ADAM17, SLC11A1, MYC, WNT16, TRAF2, SPRY2, SMAD1, GADD45B, TMEM106A, AR, TLR4, ADRA2B, PTK2B, PSMD10, EPHA2, DUSP4, RRAS, FFAR4, NRP1, SEMA4C* |
| GO:0001525 | Angiogenesis | 131 | 1.449 | 2.20E-02 | 4.71E-02 | *CXCL10, CHI3L1, STAT1, CXCL8, SERPINE1, THBS1, DLL4, IL1A, CD40, ACKR3, GLUL, ESM1, ADM, ANGPTL4, CLDN5, RHOB, B4GALT1, ITGA5, VASH2, AQP1, CREB3L1, SAT1, GATA2, F3, SPRY2, SLC31A1, SMAD1, PPP1R16B, IL10, PTK2B, AGTR1, EPHA2, RRAS, GNA13, NRP1, HTATIP2* |
| GO:1902532 | Negative regulation of intracellular signal transduction | 170 | 1.444 | 1.46E-02 | 3.39E-02 | *DHX58, STAT1, THBS1, DUSP5, IFI35, TNFAIP3, ATF3, ACKR3, OPTN, DUSP10, LIF, GDF15, CGAS, SESN2, MYOC, BCL6, FBXO7, NFKBIA, CALR, MYC, YBX3, CASP8, SELENOS, FBP1, SLC39A8, SPRY2, CYP26B1, TNFAIP1, TLR4, INSIG1, IL10, DDIT3, PSMD10, DUSP4, CXCL12, NLRP3* |
| GO:0008284 | Positive regulation of cell population proliferation | 215 | 1.443 | 1.07E-02 | 2.55E-02 | *STAT1, TNFSF13B, THBS1, IL1B, LTF, TNFAIP3, PIM1, CCL5, DLL4, IL1A, CD40, CSF3, STAT5A, HAVCR2, CSF2, ESM1, ADM, IL6, IL15, TIMP1, GLI2, LIF, NKX3-1, BCL6, CLDN5, IL6R, CALR, B4GALT1, MYC, AQP1, SLC7A1, TFRC, RNASEH2B, PTN, TICAM1, GATA2, JAK2, F3, PPP1R16B, TNFSF9, AR, TLR4, FAM98A, LIG4, IL10, SLC35F6, PTK2B, IGFBP2,* |

| | | | | | | |
|---|---|---|---|---|---|---|
| | | | | | | *AGTR1, SLC25A33, ADAM10, PID1, IL12RB2, NOP2, HMOX1, SLC25A5, HAPLN1, GRK5, TIPIN, TGFB3, NME1, FGFR4* |
| GO:0043408 | Regulation of MAPK cascade | 154 | 1.441 | 2.24E-02 | 4.77E-02 | *SAA3, CHI3L1, FAS, DUSP5, ADRA2A, GADD45G, ATF3, CD40, HAVCR2, ACKR3, GPR37, CSF2, CCR1, DUSP10, LIF, GDF15, MYOC, ADAM17, MYC, WNT16, TRAF2, SPRY2, GADD45B, TMEM106A, AR, TLR4, ADRA2B, PTK2B, PSMD10, EPHA2, DUSP4, RRAS, FFAR4, NRP1, SEMA4C* |
| GO:0009894 | Regulation of catabolic process | 237 | 1.439 | 5.54E-03 | 1.46E-02 | *SAA3, IFNG, PSME2, ADRA2A, PSME1, TNFAIP3, DEPP1, ZFP36, STING1, IGF2BP1, OPTN, TIMP1, AGO2, ARID5A, SESN2, DRAM1, FBXO7, MAD2L2, RAB39B, TIGAR, TIMP3, ZFAND2A, PLK3, DRAM2, NOS2, TRAF2, MCL1, TICAM1, FBP1, RUBCN, IL10, PTK2B, PSMD10, PSMD3, ABHD5, TMEM39A, FOXO1, SLC25A5, FMR1, TENT5A, SQSTM1, ZFP36L1, RALB, PAQR3, PDE12, ZBTB7A, DESI1, TIRAP, LPCAT1* |
| GO:0043687 | Post-translational protein modification | 268 | 1.429 | 9.19E-03 | 2.26E-02 | *ISG15, UBA7, USP18, SOCS1, SOCS3, PARP12, TRIM21, RNF19B, TNFAIP3, EGR2, EGR1, PARP3, PARP11, CISH, FBXO7, USP15, GNL3, TRAF2, TICAM1, PADI4, RNF115, JAK2, WSB2, SPRY2, ASB5, TNFAIP1, RNF122, USP44, LOC100625207, PSMD10, SOCS7, USP2, USP51, GTPBP4, UBE2D1, SQSTM1, RLIM, FBXO4, DCAF11, KLHL15, DESI1, VCP, NDFIP2, WBP1L, RNF114, RNF138, CRY1, TRIM27, OTUB2, TRAF7, WWP1, SENP1, KLHL25, UBE2D3, PLAA, STT3B* |
| GO:0045184 | Establishment of protein localization | 452 | 1.425 | 2.58E-03 | 8.26E-03 | *NEURL3, RSAD2, ISG15, IFNG, RHBDF2, CD74, SNX10, ADRA2A, TNFAIP3, STX11, EGR2, CCL5, IL1A, APOD, LRRC32, IL6, F2RL2, FFAR2, RAB3D, OPTN, STAT3, STXBP1, ARRDC3, GDAP1, CADPS2, TXNIP, ARL5B, FBXO7, RAB39B, NFKBIA, SELENBP1, RHOB, CALR, SEC23B, ZFAND2A, TFRC, RAB20, TOMM40, PLK3, TLR2, SEC61A1, XPO6, NOS2, IPO4, SELENOS, DERL3, PREB, SRPRA, LRRC8A, CHGA, TIMM8A, SCFD2, HSPA9, SEC24A, TLR4, INSIG1, ARF4, DDIT3, TMED10, AP3S2, USO1, SNX19, TRAM1, FOXO1, STX17, RAB11FIP5, TMED2, TGFB3, SEC31B, NUP188, SQSTM1, XPO5, SEC22B, TMED5, SSR3, C2CD2L, KPNB1, IPO13, DESI1, TVP23B, LOC100157391, SEC24C, SPCS1, KPNA2, TMED9, ZPR1, CHMP4A, COPA, VCP, NDFIP2, SNX9, HSPA1L, ARCN1* |

| | | | | | | |
|---|---|---|---|---|---|---|
| GO:0006886 | Intracellular protein transport | 189 | 1.425 | 2.27E-02 | 4.80E-02 | *IFNG, CD74, SNX10, STX11, EGR2, APOD, RAB3D, STAT3, TXNIP, ARL5B, NFKBIA, RHOB, CALR, SEC23B, RAB20, PLK3, SEC61A1, XPO6, IPO4, SELENOS, DERL3, SRPRA, TIMM8A, HSPA9, SEC24A, INSIG1, TMED10, AP3S2, USO1, TRAM1, STX17, SQSTM1, XPO5, KPNB1, IPO13, DESI1, SEC24C, KPNA2, TMED9, ZPR1, COPA, VCP, SNX9, AHCYL1, HYAL2, COPB1, AP4B1, CLTC, SAR1B, COPZ1, SEC63, M6PR, TNPO2, STX12, ARF6, TMED1, VPS18, RIC1, SNX16, SEC61B, RAB1A, VPS26B* |
| GO:0015031 | Protein transport | 395 | 1.412 | 3.48E-03 | 1.05E-02 | *NEURL3, RSAD2, IFNG, RHBDF2, CD74, SNX10, ADRA2A, STX11, EGR2, CCL5, IL1A, APOD, IL6, F2RL2, FFAR2, RAB3D, OPTN, STAT3, STXBP1, ARRDC3, CADPS2, TXNIP, ARL5B, RAB39B, NFKBIA, SELENBP1, RHOB, CALR, SEC23B, TFRC, RAB20, TOMM40, PLK3, TLR2, SEC61A1, XPO6, NOS2, IPO4, SELENOS, DERL3, PREB, SRPRA, LRRC8A, CHGA, TIMM8A, SCFD2, HSPA9, SEC24A, TLR4, INSIG1, ARF4, TMED10, AP3S2, USO1, SNX19, TRAM1, FOXO1, STX17, RAB11FIP5, TMED2, TGFB3, SEC31B, NUP188, SQSTM1, XPO5, SEC22B, TMED5, C2CD2L, KPNB1, IPO13, DESI1, TVP23B, LOC100157391, SEC24C, KPNA2, TMED9, ZPR1, COPA, VCP, NDFIP2, SNX9* |
| GO:0006508 | Proteolysis | 432 | 1.410 | 4.45E-03 | 1.24E-02 | *ISG15, IFNG, USP18, PSMB9, PSMB8, SERPINE1, PSMB10, C2, PSME2, MMP8, CTSL, LOC100522887, THBS1, ADRA2A, RNF19B, LTF, PSME1, CASP10, TNFAIP3, CASP1, MMP12, CFB, MMP1, HSP90B1, CNDP2, PSMA5, TIMP1, DPP3, TMPRSS4, PSMA2, CFLAR, MMP27, FBXO7, CASP3, PSMA3, CALR, ADAM17, TIMP3, USP15, ZFAND2A, MMP9, GZMA, CASP8, ADAMTS9, SELENOS, DERL3, RHBDL2, RNF115, DNAJB9, KLHDC3, PISD, ASRGL1, F3, YOD1, PSMA1, CFI, TNFAIP1, ADAMTS1, IL10, PTK2B, RNF122, USP44, DDIT3, LOC100625207, PSMD10, PSMD3, PSMB4, USP2, ADAM10, IDE, DNPEP, ENPEP, USP51, SPCS3, ADAM15, F10, AEBP1, CLCA1, CNDP1, UBE2D1, THOP1, FGFR4, RLIM, FBXO4, XPNPEP2, PSMB3, MMP2, LGMN, PSMF1, KLHL15, CD46, PAQR3, DESI1, PSMA4, SPCS1, RAD23A, CHMP4A, ACE2, VCP, FOXRED2, NDFIP2, SEC11C, RNF114, CHMP5, CTSC, CPXM1, EDEM2, OTUB2, TRAF7, METAP1, WWP1, CTSB* |

| | | | | | | |
|---|---|---|---|---|---|---|
| GO:2000145 | Regulation of cell motility | 225 | 1.409 | 1.36E-02 | 3.18E-02 | *CXCL10, IFNG, MADCAM1, SELE, CCL4, CCL2, THBS1, ADRA2A, CCL5, DLL4, KIT, CD40, APOD, CCL8, GLUL, CCR1, CXCL16, DUSP10, GDF15, MYOC, BCL6, CLDN5, RHOB, CALR, TNFSF14, PHLDA2, KLRK1, RIPK3, CASP8, PTN, GATA2, SPRY2, ADORA1, TLR4, SEMA4D, STC1, PTK2B, EPHA2, CMKLR1, ADAM10, RRAS, CXCL12, GNA13, CLDN4, GTPBP4, NRP1, NUS1, MEAK7* |
| GO:0048871 | Multicellular organismal-level homeostasis | 185 | 1.406 | 2.33E-02 | 4.89E-02 | *ISG15, STAT1, TNFSF13B, AQP7, FAS, SNX10, IL1B, AQP3, TNFAIP3, KIT, ZFP36, IL1A, STAT5A, SPP1, BPIFA1, EGR1, ST6GALNAC1, STAT3, SLC1A5, PIWIL4, ARRDC3, GDF15, BCL6, CASP3, CLDN5, CEBPB, AQP1, DRAM2, RIPK3, TSKU, CEBPG, GATA2, JAK2, SLC39A8, PPARGC1A, HSPA9, TLR4, PTK2B, G0S2, DDIT3, SFTPD, CMKLR1, FFAR4, HMOX1, SLC25A5* |
| GO:0007155 | Cell adhesion | 326 | 1.399 | 9.09E-03 | 2.25E-02 | *IFNG, MADCAM1, SELE, CD274, IRF1, CXCL8, RND1, SOCS1, TNFSF13B, S100A8, THBS1, IL1B, SLA-DQB1, SLA-DMB, CDH12, AMBN, CLSTN3, VCAM1, CCL5, KIT, IL1A, APOD, STAT5A, HAVCR2, SPP1, OLR1, LRRC32, BVES, IL15, SIGLEC1, LYVE1, SLITRK2, DUSP10, LIF, TNC, MYOC, BCL6, JAK3, MAD2L2, CLDN5, RHOB, CALR, B4GALT1, ITGA5, CEBPB, SLC7A1, TFRC, SELP, TNFSF14, SNED1, FUT2, JAM3, JAK2, AMIGO2, LSAMP, SLC39A8, ICAM1, TNFSF9, SEMA4D, ATP1B1, ENTPD1, IL10, PTK2B, IGFBP2, EPHA2, ADAM10, RRAS, CXCL12, CLDN4, GTPBP4, HAPLN1, NRP1, PEAR1, THBS2, DMP1, ICAM4, VCAN, TFE3, FBLIM1* |
| GO:0006914 | Autophagy | 155 | 1.397 | 2.16E-02 | 4.67E-02 | *IFNG, S100A8, FUNDC1, DEPP1, STING1, OPTN, SESN2, DRAM1, FBXO7, RAB39B, TIGAR, TMEM74, PLK3, DRAM2, CALCOCO2, ATG2A, MCL1, TICAM1, RUBCN, IL10, GABARAPL1, ATP2A2, TMEM39A, FOXO1, SLC25A5, STX17, SQSTM1, RALB, CHMP4A, VCP, ORMDL3, EIF2S1, FYCO1, CHMP5, TRIM27, PLK2, SPATA18, PIM2, PLAA* |
| GO:0061919 | Process utilizing autophagic mechanism | 155 | 1.397 | 2.16E-02 | 4.67E-02 | *IFNG, S100A8, FUNDC1, DEPP1, STING1, OPTN, SESN2, DRAM1, FBXO7, RAB39B, TIGAR, TMEM74, PLK3, DRAM2, CALCOCO2, ATG2A, MCL1, TICAM1, RUBCN, IL10, GABARAPL1, ATP2A2, TMEM39A, FOXO1, SLC25A5, STX17, SQSTM1, RALB, CHMP4A, VCP, ORMDL3, EIF2S1, FYCO1, CHMP5, TRIM27, PLK2, SPATA18, PIM2, PLAA* |

| | | | | | | |
|---|---|---|---|---|---|---|
| GO:0051093 | Negative regulation of developmental process | 197 | 1.385 | 1.93E-02 | 4.32E-02 | *CXCL10, IRF1, STAT1, SERPINE1, SOCS1, THBS1, LTF, ZFP36, STAT5A, PARP3, CCR1, GLI2, DUSP10, DNAJB11, NKX3-1, TMEM176B, GDF15, BCL6, FBXO7, JAK3, MAD2L2, NFKBIA, CLDN5, CALR, MYC, TSKU, YBX3, CREB3L1, PTN, GATA2, SPRY2, SMAD1, HSPA9, SEMA4D, INSIG1, RBPMS2, PTK2B, DDIT3, TMEM182, EPHA2, SIX4, RFLNB, FOXO1, NRP1* |
| GO:0043069 | Negative regulation of programmed cell death | 213 | 1.382 | 2.00E-02 | 4.43E-02 | *CCL2, FAS, THBS1, LTF, TNFAIP3, PIM1, CCL5, STAT5A, ACKR3, HSH2D, GLI2, ANGPTL4, CFLAR, BCL6, FBXO7, TIGAR, BCL2A1, ITGA5, CEBPB, MYC, AQP1, PLK3, ST3GAL1, WNT16, FTH1, YBX3, CASP8, SELENOS, ATF5, MCL1, GATA2, JAK2, PPARGC1A, SPRY2, AR, TCIM, LIG4, IL10, SLC35F6, PTK2B, PSMD10, HIGD1A, CXCL12, CTH, SIX4, FFAR4, HMOX1, SLC25A5, NRP1, GRK5* |
| GO:0016567 | Protein ubiquitination | 191 | 1.378 | 2.31E-02 | 4.88E-02 | *ISG15, UBA7, SOCS1, SOCS3, TRIM21, RNF19B, TNFAIP3, CISH, FBXO7, TRAF2, TICAM1, RNF115, WSB2, SPRY2, ASB5, TNFAIP1, RNF122, PSMD10, SOCS7, GTPBP4, UBE2D1, SQSTM1, RLIM, FBXO4, DCAF11, KLHL15, VCP, NDFIP2, WBP1L, RNF114, RNF138, CRY1, TRIM27, TRAF7, WWP1, KLHL25, UBE2D3, PLAA* |
| GO:0010558 | Negative regulation of macromolecule biosynthetic process | 465 | 1.377 | 4.91E-03 | 1.33E-02 | *SAA3, IFNG, DHX58, IRF1, CXCL8, SERPINE1, FASLG, THBS1, LTF, TNFAIP3, DLL4, MOV10, ZFP36, APOD, HAVCR2, LRRC32, EGR1, PARP3, PRDM1, CCR1, RFX5, IGF2BP1, GLI2, PIWIL4, MORC3, CEBPD, NKX3-1, AGO2, ARID5A, TXNIP, SESN2, BCL6, MAD2L2, ELF4, CLDN5, CALR, CEBPB, BAZ1A, PLK3, LOC100624590, RNASEH2B, TSKU, NOS2, YBX3, CREB3L1, TRAF2, ATF5, CD84, GATA2, FBP1, HES1, ZMYND15, PNPT1, AR, TLR4, SEMA4D, IL10, DDIT3, PSMD10, ZHX3, EPHA2, CMKLR1, USP2, CIPC, ADAM10, SETDB2, SIX4, KAT2A, KDM8, FFAR4, FOXO1, ETF1, NKRF, FMR1, TENT5A, PPARD, TGFB3, SQSTM1, RLIM, ZFP36L1, GJA1, LYAR, SOX11, EIF4EBP1, PDE12, MTERF3, PELO, TP53, TIRAP* |
| GO:0009890 | Negative regulation of biosynthetic process | 481 | 1.376 | 3.31E-03 | 1.03E-02 | *SAA3, IFNG, DHX58, IRF1, CXCL8, SERPINE1, FASLG, THBS1, LTF, TNFAIP3, DLL4, MOV10, ZFP36, APOD, HAVCR2, LRRC32, EGR1, MFSD12, PARP3, PRDM1, CCR1, RFX5, IGF2BP1, GLI2, PIWIL4, MORC3, CEBPD, NKX3-1, AGO2, ARID5A, TXNIP, SESN2, BCL6,* |

| | | | | | | |
|---|---|---|---|---|---|---|
| | | | | | | *MAD2L2, ELF4, CLDN5, CALR, CEBPB, BAZ1A, PLK3, LOC100624590, RNASEH2B, TSKU, NOS2, YBX3, CREB3L1, TRAF2, ATF5, CD84, GATA2, FBP1, ORMDL1, HES1, ZMYND15, PNPT1, AR, TLR4, SEMA4D, INSIG1, IL10, DDIT3, PSMD10, ZHX3, EPHA2, CMKLR1, USP2, CIPC, ADAM10, PID1, SETDB2, SIX4, KAT2A, KDM8, FFAR4, FOXO1, ETF1, NKRF, FMR1, TENT5A, PPARD* |
| GO:0070647 | Protein modification by small protein conjugation or removal | 256 | 1.361 | 1.83E-02 | 4.12E-02 | *ISG15, UBA7, USP18, SOCS1, SOCS3, TRIM21, RNF19B, TNFAIP3, EGR2, EGR1, CISH, FBXO7, USP15, GNL3, TRAF2, TICAM1, RNF115, WSB2, SPRY2, ASB5, TNFAIP1, RNF122, USP44, LOC100625207, PSMD10, SOCS7, USP2, USP51, GTPBP4, UBE2D1, SQSTM1, RLIM, FBXO4, DCAF11, KLHL15, DESI1, VCP, NDFIP2, WBP1L, RNF114, RNF138, CRY1, TRIM27, OTUB2, TRAF7, WWP1, SENP1, KLHL25, UBE2D3* |
| GO:0006796 | Phosphate-containing compound metabolic process | 389 | 1.294 | 2.09E-02 | 4.58E-02 | *IDO1, IFNG, CHI3L1, SAMHD1, GFPT2, PPA1, HK3, THBS1, STAT2, CCL5, C3, SLC25A22, NAMPT, CNP, IL15, HK2, STAT3, LIF, CTPS1, GCK, DPYS, PLPPR2, TIGAR, UGP2, ITGA5, KMO, GK, RNASEH2B, NEK6, CHKA, ITPKC, FBP1, ALDOC, JAK2, PISD, CDS1, AASS, PDXK, ORMDL1, PLSCR3, ACSL4, AR, SEMA4D, ATP1B1, ENTPD1, TMEM38B, FMO1, PRPS1, PGK1, EPHA2, AMPD3, ABHD5, PID1, FITM2, PDE4A, FOXO1, NRP1, DCTD, GFPT1, NUS1, NME1, ACLY, AGPAT4, SAMD8, UAP1, ITPK1, ZBTB7A, ATP6V1B2, LPCAT1, AK4, PDE4B, TKT, NUDT7, ORMDL3, DDIT4, NT5C, PLCH1, RFK, DCTPP1, NT5C3A, LPCAT3, PDHX, MDH1, FUT8, GNPNAT1, ME1, PPP2R2A, UGDH, GPAM, AK3, NAPEPLD, RPE, ABL1, PLA2G4B, MEF2C, CD80, PGD, G6PD, GOT1, PPCDC, ADSS2, ATM, PIK3CD, MTOR, UXS1, GPAT4, AGPAT3, MTMR1, KCTD20* |
| GO:0007167 | Enzyme-linked receptor protein signaling pathway | 232 | -1.290 | 2.06E-02 | 4.53E-02 | *SAMD12, IGF2, BAMBI, CDKN2B, BLVRB, EFNA2, ANGPTL1, INPP5K, BMPR2, PHF14, EFNA4, WIF1, EGF, GATA3, CD109, TMEM100, FGF18, BMP7, EFNB3, TMEM204, EPHA7, BDNF, CAV2, CITED1, FOLR3, BLK, FGF1, GFRA3, EPHA3, ANGPT1, HPGD, HHIP, VEGFD, NDP, APLN, IGF1, ASPN, SFRP2, AGTR2, CCN3, CRB2* |
| GO:0031175 | Neuron projection development | 167 | -1.301 | 1.60E-02 | 3.66E-02 | *BORCS7, RAB11A, PLPPR4, GBX1, BBS1, SNAPIN, FLOT1, NTN4, GFAP, KREMEN1, BTBD3, NTF4, VCL, WNT7A, RTN4RL1, NFIB, HOXA2, EXT1, ACP4, NDNF, B4GALT6,* |

| | | | | | | |
|---|---|---|---|---|---|---|
| | | | | | | *SEMA6C, SEMA6A, DCX, GPRASP3, NR2E1, NYAP1, BMPR2, EFNA4, CNTN5, CNR1, CNTN1, SPOCK1, BMP7, EFNB3, EPHA7, STMN1, MAP1B, BDNF, SEMA3E, FOLR3, UCHL1, SEMA3B, ZNF365, EPHA3, ATCAY, NDP, SFRP2* |
| GO:0071363 | Cellular response to growth factor stimulus | 142 | -1.317 | 1.76E-02 | 3.98E-02 | *GATA3, CD109, TMEM100, FGF18, BMP7, TMEM204, BDNF, CAV2, CITED1, FOLR3, WNT2, FGF1, HPGD, HHIP, VEGFD, NDP, APLN, ASPN, SFRP2, AGTR2, CRB2* |
| GO:0007017 | Microtubule-based process | 245 | -1.396 | 2.72E-03 | 8.63E-03 | *CFAP298, SPACA9, TUBB4B, TACC3, BICDL2, SGO1, CCDC61, ABRAXAS1, ODAD3, DTNBP1, GEN1, CFAP68, PPP1R35, DNAI2, CEP250, NCKAP5L, RAB11A, NUF2, DYNC1I2, KIF11, SPC25, DCLK1, BBS1, PIERCE1, MID1IP1, CCNB1, SNAPIN, TBCB, CCNF, CDK1, TUBB4A, CFAP20, CCNB2, HAUS1, CATSPER4, HOOK2, DYNC2LI1, MEIG1, MID1, KIF20A, CGN, ATAT1, FAM110A, MAD2L1, KIFC1, IFT46, WRAP73, TUBB2B, SPDL1, TUB, TSSK4, AURKB, BICD1, CENPA, DCX, KLC4, CENPH, CFAP141, HSPB1, DYNC2I2, KIF22, RGS14, CFAP97D1, BIRC5, SKA1, PARD6G, CFAP276, TPPP3, STMN1, MAP1B, UCHL1, ODAD1, TPPP, EPHA3, DEFB1* |
| GO:0016055 | Wnt signaling pathway | 131 | -1.404 | 1.17E-02 | 2.77E-02 | *PORCN, HNF1A, FRZB, DKK3, SOX9, ARL6, RSPO2, KREMEN1, PFDN5, WNT7A, MDFIC, EXT1, TAX1BP3, LGR5, LGR4, WNT9A, FZD2, WLS, CALCOCO1, BAMBI, WNT11, WNT3A, TLE4, WNT10B, WIF1, FAM53B, GATA3, CPE, SDC1, FOLR3, WNT4, WNT2, ALPK2, SFRP5, SFRP4, DKK2, NDP, SFRP2* |
| GO:0048858 | Cell projection morphogenesis | 113 | -1.495 | 8.75E-03 | 2.18E-02 | *BTBD3, NTF4, VCL, WNT7A, NFIB, HOXA2, EXT1, B4GALT6, SEMA6C, SEMA6A, DCX, GPRASP3, NR2E1, NYAP1, BMPR2, EFNA4, CNTN5, CNTN1, BMP7, EFNB3, EPHA7, MAP1B, BDNF, SEMA3E, UCHL1, SEMA3B, ZNF365, EPHA3, NDP* |
| GO:0120039 | Plasma membrane bounded cell projection morphogenesis | 113 | -1.495 | 8.75E-03 | 2.18E-02 | *BTBD3, NTF4, VCL, WNT7A, NFIB, HOXA2, EXT1, B4GALT6, SEMA6C, SEMA6A, DCX, GPRASP3, NR2E1, NYAP1, BMPR2, EFNA4, CNTN5, CNTN1, BMP7, EFNB3, EPHA7, MAP1B, BDNF, SEMA3E, UCHL1, SEMA3B, ZNF365, EPHA3, NDP* |

| | | | | | | |
|---|---|---|---|---|---|---|
| GO:0048812 | Neuron projection morphogenesis | 109 | -1.536 | 5.56E-03 | 1.46E-02 | *BTBD3, NTF4, VCL, WNT7A, NFIB, HOXA2, EXT1, B4GALT6, SEMA6C, SEMA6A, DCX, GPRASP3, NR2E1, NYAP1, BMPR2, EFNA4, CNTN5, CNTN1, BMP7, EFNB3, EPHA7, MAP1B, BDNF, SEMA3E, UCHL1, SEMA3B, ZNF365, EPHA3, NDP* |

Abbreviations: GO:BP: Biological Processes in Gene Ontology; FDR: False discovery rate; MAPK: Mitogen-activated protein kinase; NES: Normalized enrichment score.

**Supplementary Table S7. Significantly enriched pathways from the GSEA result of the KEGG database, High-glucose infection versus Low-glucose infection.** Abbreviations: GSEA: Gene set enrichment analysis; KEGG: Kyoto Encyclopedia of Genes and Genomes.

| KEGG ID | Description | Set Size | NES | P-value | FDR | Core enrichment gene |
|---|---|---|---|---|---|---|
| ssc04657 | IL-17 signaling pathway | 70 | 2.601 | 3.39E-11 | 1.95E-09 | *IL6, AMCF-II, IL1B, CXCL2, MMP13, CXCL8, CCL2, CSF3, FOSL1, S100A8, MMP1, CXCL10, TNFAIP3, NFKBIA, FOSB, FOS, HSP90AA1, S100A9, IFNG, PTGS2, CASP3, MMP9, HSP90B1, CHUK, LCN2, MAPK6, MAPK4, TRAF2, NFKB1, CEBPB* |
| ssc04141 | Protein processing in endoplasmic reticulum | 149 | 2.599 | 4.26E-16 | 1.47E-13 | *HSPA6, HSPH1, DNAJA1, DNAJB11, HSP90AA1, HSPA8, UGGT2, CKAP4, DDIT3, DNAJC1, DNAJA2, CRYAB, DNAJB1, HSP90B1, SEC23B, NGLY1, HSPBP1, SEC24A, EIF2S1, HSPA5, HYOU1, MOGS, UBE2D3, UBE2D1, SEC61B, PREB, RNF5, UBQLN1, ATF6, STT3B, TRAF2, LMAN1, DAD1, SEC23A, SSR1, RAD23A, OSTC, SEC13, SEC62, YOD1, PDIA4, HSPA4L, SSR3, DNAJC3, RRBP1, UBXN4, SELENOS, SEL1L, SEC24D, EDEM3, ERO1B, STT3A, UBE2D2, SAR1B, RPN1, RAD23B, CANX, RPN2, SEC63, EIF2AK3, MAPK8, DNAJC10, PDIA6, CALR, SAR1A, VCP, SSR4, UBXN8, DERL3, UBE2J1, UGGT1, BAK1, MAPK9, PPP1R15A, TRAM1, BAG1, PDIA3, HERPUD1, ERLEC1, DERL2, BCAP31, GANAB, HSP90AB1, BAG2, SEC61A2, XBP1, UBQLN4, TMEM258* |
| ssc03008 | Ribosome biogenesis in eukaryotes | 69 | 2.488 | 4.12E-10 | 1.58E-08 | *NHP2, GAR1, SPATA5, FBL, EMG1, GNL3, RAN, WDR43, NOP58, NOB1, WDR36, UTP6, RBM28, BMS1, MPHOSPH10, WDR75, WDR3, NXT2, AK6, DKC1, EIF6, NOP56, CSNK2A1, UTP18, NXT1, UTP4, IMP3, RCL1, UTP15, HEATR1, GTPBP4, NOL6, GNL2, FCF1, RIOK1, RPP30, SNU13, NAT10, NMD3, XRN1, TBL3, SBDS, RPP38, NOP10, GNL3L, RPP40, POP1, CSNK2B, RRP7A, POP7* |
| ssc03010 | Ribosome | 144 | 2.409 | 2.79E-12 | 3.21E-10 | *MRPL55, MRPL40, MRPS17, RSL24D1, MRPL35, MRPL52, RPS27L, MRPL2, MRPL51, MRPL13, MRPL22, MRPL34, MRPS35, MRPL20, MRPL42, MRPS24, MRPS18C, MRPS11, MRPL19, MRPS27, MRPL17, MRPS15, MRPL15, MRPL32, RPL27, MRPL47, MRPL44, MRPS23, MRPS31,* |

| | | | | | | |
|---|---|---|---|---|---|---|
| | | | | | | *MRPL30, MRPS30, MRPS14, MRPS33, MRPL46, MRPL28, MRPS25, MRPS5, MRPL38, RPS21, MRPL24, RPL26L1, MRPL48, MRPL36, MRPL50, MRPS7, MRPL49, MRPS36, MRPL18, MRPS6, RPL35A, RPS23, RPL23A, MRPS18A, RPS15A, MRPS10, MRPL11, MRPS22, DAP3, MRPL43, RPL38, RPL23, MRPL33, MRPS26, MRPL3, RPLP1, RPS27, RPS25, MRPL27, MRPL37, RPL21, MRPL39, RPS18, RPS24, RPL36A, RPL14, RPLP2, MRPL1, RPL10A, RPS2, RPS16, RPS27A, RPL34* |
| ssc00970 | Aminoacyl-tRNA biosynthesis | 43 | 2.402 | 1.51E-07 | 2.98E-06 | *SARS1, AARS1, MARS1, TARS1, GARS1, CARS1, WARS1, YARS1, IARS1, NARS1, EPRS1, FARSA, YARS2, LARS1, RARS1, FARSB, VARS1, QRSL1, DARS1* |
| ssc00190 | Oxidative phosphorylation | 113 | 2.400 | 1.15E-10 | 4.94E-09 | *ATP12A, CYCS, NDUFB2, PPA1, NDUFB1, NDUFC2, ATP5MC1, ND4L, NDUFB3, NDUFA6, COX6B, NDUFB9, COX17, COX5B, ATP6V1C1, NDUFS4, COX11, LOC100522678, ATP5PD, ATP5PF, ATP5MF, UQCRC1, COX7A2, NDUFB5, ATP5F1B, ATP6V1F, NDUFA5, NDUFA11, ATP5F1C, UQCR11, COX7B, ATP5MC3, ATP5ME, COX5A, NDUFB6, NDUFB10, NDUFA9, NDUFA2, ATP5F1A, NDUFV2, ATP5PO, UQCR10, NDUFB11, NDUFB7, UQCRQ, NDUFS5, COX7C, NDUFS1, ATP6V0A2, NDUFA4, ATP5F1D, NDUFB4, NDUFS8, UQCRC2, NDUFA8, ATP5PB, LOC100524873, NDUFS2, SDHD, CYC1, NDUFB8, ATP6V1H* |
| ssc04668 | TNF signaling pathway | 104 | 2.399 | 5.96E-10 | 2.06E-08 | *IL6, AMCF-II, IL1B, CXCL2, SELE, CCL2, BIRC3, LIF, CXCL10, TNFAIP3, FAS, NFKBIA, SOCS3, BCL3, CREB5, FOS, JUNB, MAP3K8, PTGS2, IRF1, NOD2, CASP3, MMP9, LOC100511937, CASP7, CHUK, VCAM1, ICAM1, MLKL, ADAM17, TRAF2, PIK3CA, NFKB1, IL15, CEBPB* |
| ssc04932 | Non-alcoholic fatty liver disease | 135 | 2.373 | 1.63E-11 | 1.12E-09 | *IL6, IL1A, IL1B, CXCL8, CYCS, FAS, SOCS3, FOS, NDUFB2, NDUFB1, DDIT3, NDUFC2, NDUFB3, CASP3, LOC100511937, CASP7, NDUFA6, EIF2S1, COX6B, NDUFB9, COX5B, NDUFS4, LOC100522678, UQCRC1, COX7A2, TRAF2, NDUFB5, PIK3CA, NFKB1, NDUFA5, NDUFA11, UQCR11, COX7B, COX5A, NDUFB6, NDUFB10, NDUFA9, NDUFA2, NDUFV2, UQCR10, NDUFB11, NDUFB7, UQCRQ, NDUFS5, PRKAA2, COX7C, NDUFS1, NDUFA4, EIF2AK3, MAPK8, PIK3R2, NDUFB4, NDUFS8, CASP8, UQCRC2, NDUFA8, MLXIPL, PKLR, CDC42, PRKAB2, LOC100524873, MAPK9, NDUFS2, SDHD, CYC1, NDUFB8, PRKAA1* |

| | | | | | | |
|---|---|---|---|---|---|---|
| ssc03040 | Spliceosome | 123 | 2.296 | 1.39E-09 | 4.37E-08 | *HSPA6, SNRPF, SNRPD1, HSPA8, RBM8A, SNRPG, SNRNP27, HNRNPC, SNRPB2, NCBP1, SNRPE, THOC3, DHX15, SF3B6, SMNDC1, SRSF3, U2AF1, SNRPD3, MAGOH, MAGOHB, HNRNPA3, TXNL4A, CWC15, SNRPA1, SNRPC, LSM8, CDC40, ZMAT2, LSM6, SF3B3, TRA2B, LSM5, LSM7, PRPF40A, HNRNPK, SRSF1, EIF4A3, PHF5A, USP39, PPIL1, HNRNPM, NCBP2, ALYREF, SF3A3, SRSF10, BUD31, THOC2, SNRPA, RBMX2, SNU13, THOC6, LOC100738822, SRSF7, DDX23, LSM2, LOC102162880, LSM3, PRPF19, SNRPD2, CTNNBL1, THOC7, SNIP1, DDX5, CRNKL1, SF3B4, DHX38, SNRNP40, U2AF2, WBP11* |
| ssc01230 | Biosynthesis of amino acids | 61 | 2.290 | 1.56E-07 | 2.98E-06 | *PSAT1, PSPH, PHGDH, ASNS, SHMT2, ARG1, GPT2, CBS, PYCR1, ARG2, PAH, GOT1, BCAT1, SDS, PGK1, IDH3A* |
| ssc05012 | Parkinson disease | 220 | 2.272 | 3.86E-13 | 6.66E-11 | *LOC100127131, CYCS, SLC39A14, NDUFB2, NDUFB1, DDIT3, NDUFC2, PSMB3, ATP5MC1, ND4L, PSMD14, UCHL1, NDUFB3, SLC39A6, CASP3, PSMD12, SLC11A2, KIF5B, LRRK2, NDUFA6, EIF2S1, COX6B, NDUFB9, SOD1, HSPA5, PSMA3, PSMA5, COX5B, CALM2, NDUFS4, SLC39A8, TUBAL3, LOC100522678, TUBB1, ATP5PD, ATP5PF, PSMB6, ATF6, UQCRC1, COX7A2, NDUFB5, ATP5F1B, PSMD6, NDUFA5, TUBB, NDUFA11, PSMC5, ATP5F1C, SLC18A1, GNAI1, UQCR11, PSMC6, COX7B, ATP5MC3, COX5A, NDUFB6, PSMA4, NDUFB10, PARK7, PSMA2, PSMD8, PSMA1, NDUFA9, NDUFA2, PSMC4, ATP5F1A, NDUFV2, ATP5PO, UQCR10, SLC39A7, NDUFB11, NDUFB7, UBE2L3, UQCRQ, PSMD2, NDUFS5, PSMC2, PSMB1, RYR3, COX7C, PSMC1, NDUFS1, PSMD13, VDAC3, PSMB4, NDUFA4, ATP5F1D, EIF2AK3, MAPK8, VDAC2, PSMD1, NDUFB4, NDUFS8, PSMD4, UBB, UQCRC2, NDUFA8, VDAC1, TUBB4B, MCU, UBE2J1, SLC25A5, PSMD7, SLC39A1, SLC39A10, ATP5PB, PSMB7, LOC100524873, MAPK9, GNAI3, NDUFS2, SDHD, CYC1, NDUFB8, CALM1, TP53* |
| ssc03013 | Nucleocytoplasmic transport | 95 | 2.269 | 5.85E-08 | 1.44E-06 | *KPNA2, TMEM33, RBM8A, XPOT, RAN, KPNA4, IPO7, NCBP1, THOC3, NXT2, PHAX, NUP54, NXT1, MAGOH, SUMO2, SAP18, MAGOHB, SEC13, NUP35, TNPO1, PYM1, KPNB1, AHCTF1, KPNA1, SUMO1, IPO11, EIF4A3, NUP50, CSE1L, NCBP2, ALYREF, TPR, XPO5, THOC2, NUP42, AAAS, NUP43, NUP155, THOC6, IPO4, NMD3, NUP37, IPO5, RANGAP1, NUP205,* |

| | | | | | | |
|---|---|---|---|---|---|---|
| | | | | | | *KPNA6, IPO8, SENP2, THOC7, SNUPN, NDC1, POM121C, NUP153, UPF2, NUP188, TNPO3, NUP160, UBE2I, RNPS1, UPF3B, LOC100511376* |
| ssc03050 | Proteasome | 40 | 2.263 | 1.50E-06 | 2.73E-05 | *IFNG, PSMB3, PSMD14, PSMD12, PSMA3, PSMA5, PSMB6, PSME3, PSMD6, PSMC5, PSMC6, PSMA4, PSMA2, PSMD8, PSMA1, PSMC4, PSMD2, PSMC2, PSMB1, PSMC1, PSMD13, PSMB4, PSMD1, PSMD4, PSME2, PSMD7, PSMB7, PSMB2, PSMD11, PSMD3* |
| ssc05144 | Malaria | 39 | 2.189 | 3.30E-06 | 5.42E-05 | *IL6, IL1B, SELE, CXCL8, CCL2, CSF3, SELP, IL10, ACKR1, THBS1, IFNG, GYPA, TLR4, TGFB3, CD40, VCAM1, ICAM1* |
| ssc05134 | Legionellosis | 44 | 2.131 | 2.27E-05 | 2.90E-04 | *IL6, IL1B, CXCL2, HSPA6, CXCL8, CYCS, NFKBIA, NFKB2, HSPA8, HSPD1, CASP3, TLR4, CASP7, SEC22B, ARF2, RAB1A, HBS1L* |
| ssc05020 | Prion disease | 226 | 2.090 | 9.13E-11 | 4.50E-09 | *IL6, IL1A, IL1B, HSPA6, LOC100127131, CYCS, CREB5, NDUFB2, HSPA8, NDUFB1, DDIT3, NDUFC2, PSMB3, ATP5MC1, ND4L, STIP1, PSMD14, NDUFB3, CASP3, EGR1, PSMD12, LOC100511937, KIF5B, NDUFA6, EIF2S1, COX6B, NDUFB9, SOD1, HSPA5, PSMA3, PPP3CA, PSMA5, COX5B, NDUFS4, TUBAL3, LOC100522678, CSNK2A1, TUBB1, ATP5PD, ATP5PF, PSMB6, UQCRC1, COX7A2, NDUFB5, PIK3CA, ATP5F1B, PSMD6, NDUFA5, TUBB, NDUFA11, PSMC5, ATP5F1C, UQCR11, PSMC6, COX7B, ATP5MC3, COX5A, NDUFB6, PSMA4, NDUFB10, PSMA2, PSMD8, PSMA1, PRNP, NDUFA9, NDUFA2, PSMC4, ATP5F1A, NDUFV2, ATP5PO, UQCR10, GRIN3A, NDUFB11, NDUFB7, UQCRQ, PSMD2, NDUFS5, FYN, PSMC2, PSMB1, RYR3, COX7C, CREB3, CREB3L1, PSMC1, NDUFS1, PSMD13, VDAC3, PSMB4, NDUFA4, ATP5F1D, EIF2AK3, MAPK8, VDAC2, PSMD1, PPP3CB, PIK3R2, NDUFB4, NDUFS8, PSMD4, UQCRC2, NDUFA8, VDAC1, TUBB4B, MCU, SLC25A5, PSMD7, ATP5PB, PSMB7, LOC100524873, MAPK9, NDUFS2, PPP3R1, SDHD, CYC1, NDUFB8* |
| ssc04064 | NF-kappa B signaling pathway | 87 | 2.053 | 5.30E-06 | 7.94E-05 | *CCL19, IL1B, CXCL2, CXCL8, BIRC3, TNFAIP3, NFKBIA, PLAU, NFKB2, RELB, PTGS2, LBP, IL1R1, TNFSF13B, TLR4, TICAM2, TNFSF14, CHUK, CD40, VCAM1, ICAM1, GADD45A, CSNK2A1, GADD45B, TRAF2, ATM, NFKB1, IRAK4, EDARADD* |
| ssc03060 | Protein export | 27 | 2.048 | 2.83E-04 | 3.15E-03 | *HSPA5, SEC61B, SRP14, SRPRB, IMMP1L, SRP54, SEC62, SRPRA, SPCS2, TMEM208, SEC11C, SEC63, GET3, GET1, SRP9, SEC11A, SPCS3, SEC61A2, SRP19, SRP72, SPCS1* |

| | | | | | | |
|---|---|---|---|---|---|---|
| ssc04060 | Cytokine-cytokine receptor interaction | 170 | 2.043 | 1.24E-08 | 3.28E-07 | *IL6, AMCF-II, CCL19, IL1A, IL1B, CXCL2, CXCL8, INHBA, GDF15, CCL2, CSF3, CCL3L1, LIF, CXCL10, IL1RAP, IL10, PPBP, INHA, CXCL13, BMP2, IL1R2, FAS, CNTFR, IFNG, RELL1, OSMR, TNFRSF11B, TNFSF13, TNFRSF12A, ACKR4, CCL8, NGFR, CXCL11, IL1R1, TNFSF13B, IL6ST, TGFB3, TNFSF14, IFNE, TNFRSF18, IFNAR2, IL33, CD40, IFNAR1, BMPR2, IFNGR1, LOC397451, IL27RA, IL4R* |
| ssc03015 | mRNA surveillance pathway | 80 | 2.041 | 9.98E-06 | 1.38E-04 | *PPP2R2C, PAPOLA, RBM8A, NCBP1, NXT2, CSTF2, HBS1L, NXT1, MAGOH, ETF1, FIP1L1, PPP1CC, SAP18, MAGOHB, PPP2CA, PELO, PYM1, CLP1, CSTF2T, EIF4A3, RNGTT, PPP1CB, GSPT1, NCBP2, ALYREF, PABPC4, CPSF2, PPP2CB, PPP2R5E, PPP2R2A, PPP2R1B, SMG5, WDR82, WDR33, CSTF1, SYMPK, PPP2R5B, PPP2R5D, PAPOLG, TARDBP, CSTF3, UPF2, CPSF6, RNPS1, MSI2, UPF3B* |
| ssc05010 | Alzheimer disease | 319 | 2.006 | 1.19E-11 | 1.02E-09 | *IL6, IL1A, IL1B, LOC100127131, CYCS, FAS, SLC39A14, NDUFB2, NDUFB1, DDIT3, PTGS2, NDUFC2, PSMB3, ATP5MC1, ND4L, PSMD14, PPID, NDUFB3, SLC39A6, CASP3, PSMD12, SLC11A2, LOC100511937, KIF5B, CASP7, CHUK, FZD9, BACE2, NDUFA6, EIF2S1, COX6B, NDUFB9, PSMA3, PPP3CA, WNT9B, PSMA5, COX5B, ADAM17, CALM2, NDUFS4, SLC39A8, TUBAL3, LOC100522678, CSNK2A1, TUBB1, FZD10, ATP5PD, ATP5PF, PSMB6, ATF6, UQCRC1, COX7A2, TRAF2, NDUFB5, PIK3CA, WNT7B, ATP5F1B, NFKB1, PSMD6, NDUFA5, WNT16, TUBB, NDUFA11, PSMC5, ATP5F1C, UQCR11, PSMC6, FZD3, COX7B, ATP5MC3, COX5A, NDUFB6, PSMA4, NDUFB10, PSMA2, PSMD8, PSMA1, NDUFA9, NDUFA2, PSMC4, KRAS, ATP5F1A, NDUFV2, ATP5PO, UQCR10, SLC39A7, NDUFB11, RB1CC1, NDUFB7, UQCRQ, PSMD2, NDUFS5, PSMC2, PSMB1, RYR3, COX7C, PSMC1, NDUFS1, ATG101, PSMD13, VDAC3, PSMB4, NDUFA4, ATP5F1D, MAP2K1, EIF2AK3, MAPK8, VDAC2, PSMD1, PPP3CB, PIK3R2, NDUFB4, NDUFS8, PSMD4, CASP8, UQCRC2, CSNK1A1, NDUFA8, VDAC1, TUBB4B, MCU, GNAQ, IDE, SLC25A5, PSMD7, SLC39A1, SLC39A10, ATP5PB, PSMB7, LOC100524873, MAPK9, NDUFS2, PPP3R1, SDHD, HSD17B10, CYC1, APOE, NDUFB8, CALM1, NRBF2, APP* |

| | | | | | | |
|---|---|---|---|---|---|---|
| ssc00270 | Cysteine and methionine metabolism | 42 | 2.004 | 9.03E-05 | 1.04E-03 | *PSAT1, PHGDH, CBS, GOT1, BCAT1, MDH1, SDS, AMD1, APIP, MDH2, ENOPH1, GCLM, LDHA, MAT1A, MTAP, SMS, MAT2B* |
| ssc05323 | Rheumatoid arthritis | 65 | 1.999 | 4.15E-05 | 5.12E-04 | *IL6, AMCF-II, IL1A, IL1B, CXCL2, CXCL8, CCL2, CTSL, CCL3L1, MMP1, FOS, IFNG, TNFSF13, TNFSF13B, TLR4, TGFB3, ICAM1, ATP6V1C1* |
| ssc05417 | Lipid and atherosclerosis | 171 | 1.964 | 1.03E-07 | 2.23E-06 | *IL6, IL1B, CXCL2, SELE, HSPA6, CXCL8, CCL2, SOD2, SELP, CCL3L1, MMP1, CYCS, FAS, NFKBIA, HSPA4, FOS, HSP90AA1, HSPA8, HSPD1, DDIT3, LBP, CASP3, MMP9, TLR4, LOC100511937, TICAM2, HSP90B1, CASP7, CHUK, STAT3, EIF2S1, CD40, HSPA5, PPP3CA, VCAM1, ICAM1, CALM2, VLDLR, TANK, ATF6, TRAF2, ROCK2, PIK3CA, RAP1B, NFKB1, IRAK4* |
| ssc00220 | Arginine biosynthesis | 15 | 1.919 | 2.02E-03 | 1.51E-02 | *ARG1, GPT2, ARG2, GOT1, GLUD1* |
| ssc04061 | Viral protein interaction with cytokine and cytokine receptor | 60 | 1.906 | 7.59E-05 | 9.03E-04 | *IL6, AMCF-II, CCL19, CXCL2, CXCL8, CCL2, CCL3L1, CXCL10, IL10, PPBP, CXCL13, ACKR4, CCL8, CXCL11, IL6ST, TNFSF14* |
| ssc03020 | RNA polymerase | 30 | 1.901 | 7.78E-04 | 7.25E-03 | *POLR1G, POLR3D, POLR1F, POLR2M, POLR3E, POLR2H, CRCP, POLR2G, POLR1C, POLR2J, POLR1H, POLR3H, POLR2K, POLR1B, POLR1A, POLR2I, POLR2F, POLR2D, POLR2E, POLR2B* |
| ssc04215 | Apoptosis - multiple species | 25 | 1.868 | 2.18E-03 | 1.57E-02 | *BIRC3, PMAIP1, CYCS, NGFR, CASP3, CASP7, SEPTIN4, BBC3, XIAP, MAPK8, CASP8, BAK1, MAPK9, BIRC5, DIABLO, BECN1, TNFRSF1A* |
| ssc03022 | Basal transcription factors | 36 | 1.846 | 1.36E-03 | 1.17E-02 | *TAF4B, TAF13, TAF12, TAF15, GTF2H3, TAF9, MNAT1, TAF2, TAF5, GTF2H5, TBP, GTF2A2, GTF2A1, TBPL1, GTF2F2, TAF3, CDK7, GTF2E2, GTF2F1, GTF2B, CCNH, TAF11, TAF9B* |
| ssc05014 | Amyotrophic lateral sclerosis | 296 | 1.842 | 7.96E-09 | 2.29E-07 | *NEFH, LOC100127131, GRIA1, CYCS, NDUFB2, NDUFB1, DDIT3, NDUFC2, PSMB3, ATP5MC1, RAB39B, ND4L, PSMD14, NDUFB3, CASP3, PSMD12, KIF5B, HNRNPA2B1, NCBP1, NDUFA6, EIF2S1, COX6B, NDUFB9, SOD1, HSPA5, PSMA3, PPP3CA, NRG1, NXT2, PSMA5,* |

| | | | | | | |
|---|---|---|---|---|---|---|
| | | | | | | *COX5B, RAB1A, GPX7, NDUFS4, GPX8, NUP54, SRSF3, TUBAL3, LOC100522678, TUBB1, TANK, ATP5PD, ATP5PF, PSMB6, UBQLN1, ATF6, NXT1, UQCRC1, DCTN6, COX7A2, TRAF2, NDUFB5, TOMM40, ATP5F1B, PSMD6, SEC13, HNRNPA3, NDUFA5, TUBB, NDUFA11, PSMC5, ATP5F1C, UQCR11, NUP35, PSMC6, COX7B, ATP5MC3, COX5A, NDUFB6, PSMA4, NDUFB10, PSMA2, PSMD8, PSMA1, PFN2, NDUFA9, NDUFA2, PSMC4, ATP5F1A, RAB5A, NDUFV2, ATP5PO, UQCR10, NDUFB11, RB1CC1, NDUFB7, UQCRQ, PSMD2, NDUFS5, NUP50, PSMC2, PSMB1, C9orf72, COX7C, PSMC1, NDUFS1, ATG101, PSMD13, PSMB4, ATXN2L, ALYREF, TPR, NDUFA4, CHMP2B, ATP5F1D, EIF2AK3, PSMD1, PPP3CB, NDUFB4, NDUFS8, PSMD4, MATR3, NUP43, UQCRC2, NDUFA8, VDAC1, VCP, NUP155, TUBB4B, MCU, ACTR1A, PSMD7, ATP5PB, SRSF7, PSMB7, LOC100524873, PFN1, NDUFS2, NUP37, PPP3R1, SDHD, CYC1, NDUFB8, TP53, NUP205, NRBF2, MAP2K3, PSMB2, ALS2, DCTN5, DCTN3, NDUFV1, PSMD11, MAPK14, BECN1, XBP1, PSMD3, UBQLN4, NDC1, TUBA1B, POM121C, NDUFA7, NUP153, TUBB2B, TNFRSF1A, TARDBP, NDUFAB1* |
| ssc05016 | Huntington disease | 258 | 1.796 | 8.33E-08 | 1.92E-06 | *SOD2, LOC100127131, GRIA1, TAF4B, CYCS, CREB5, NDUFB2, NDUFB1, NDUFC2, PSMB3, ATP5MC1, ND4L, PSMD14, NDUFB3, CASP3, PSMD12, KIF5B, NDUFA6, COX6B, NDUFB9, SOD1, PSMA3, HDAC2, PSMA5, COX5B, GPX7, NDUFS4, GPX8, TUBAL3, LOC100522678, TUBB1, ATP5PD, ATP5PF, PSMB6, UQCRC1, DCTN6, COX7A2, TRAF2, NDUFB5, ATP5F1B, PSMD6, NDUFA5, TUBB, NDUFA11, PSMC5, ATP5F1C, UQCR11, POLR2H, PSMC6, COX7B, ATP5MC3, COX5A, NDUFB6, PSMA4, NDUFB10, PSMA2, BBC3, PSMD8, PSMA1, NDUFA9, TBP, NDUFA2, PSMC4, ATP5F1A, CLTC, NDUFV2, ATP5PO, UQCR10, NDUFB11, RB1CC1, NDUFB7, UQCRQ, PSMD2, NDUFS5, POLR2G, PSMC2, PSMB1, COX7C, POLR2J, POLR2K, CREB3, CREB3L1, PSMC1, TGM2, NDUFS1, ATG101, PPARGC1A, PSMD13, VDAC3, PSMB4, TBPL1, NDUFA4, ATP5F1D, MAPK8, VDAC2, PSMD1, NDUFB4, NDUFS8, PSMD4, CASP8, UQCRC2, NDUFA8, VDAC1, TUBB4B, ACTR1A, GNAQ, SLC25A5, PSMD7, SP1, ATP5PB, POLR2I, PSMB7, LOC100524873, POLR2F, MAPK9, NDUFS2, POLR2D, CLTA, SDHD, CYC1,* |

| | | | | | | |
|---|---|---|---|---|---|---|
| | | | | | | *NDUFB8, TFAM, TP53, NRBF2, POLR2E, PSMB2, DCTN5, DCTN3, NDUFV1, PSMD11, BECN1, POLR2B, PSMD3, CREB1, SLC1A3, TUBA1B, NDUFA7* |
| ssc00260 | Glycine, serine and threonine metabolism | 30 | 1.776 | 3.30E-03 | 2.19E-02 | *PSAT1, PSPH, PHGDH, SHMT2, CBS, GLDC, SDS* |
| ssc05208 | Chemical carcinogenesis - reactive oxygen species | 193 | 1.773 | 7.38E-06 | 1.06E-04 | *SOD2, CYP1B1, NFKBIA, FOS, NDUFB2, AKR1C1, NDUFB1, HIF1A, NDUFC2, ATP5MC1, ND4L, NDUFB3, LOC100511937, CHUK, NDUFA6, COX6B, NDUFB9, SOD1, GSTM3, COX5B, NDUFS4, PTPN11, LOC100522678, EGF, ATP5PD, ATP5PF, PRKD3, UQCRC1, COX7A2, GSTO1, NDUFB5, PIK3CA, ATP5F1B, NFKB1, NDUFA5, NDUFA11, ATP5F1C, UQCR11, GSTO2, COX7B, ATP5MC3, COX5A, NDUFB6, NDUFB10, NDUFA9, NDUFA2, KRAS, ATP5F1A, NDUFV2, ATP5PO, AHR, UQCR10, NDUFB11, NDUFB7, UQCRQ, NDUFS5, COX7C, NDUFS1, VDAC3, NDUFA4, ATP5F1D, MAP2K1, MAPK8, VDAC2, PIK3R2, NDUFB4, NDUFS8, UQCRC2, NDUFA8, VDAC1, SLC25A5, ATP5PB, LOC100524873, MAPK9, NDUFS2, SDHD, CYC1, NDUFB8* |
| ssc05162 | Measles | 108 | 1.734 | 3.60E-04 | 3.76E-03 | *IL6, IL1A, IL1B, HSPA6, SLAMF1, TNFAIP3, CYCS, CD209, FAS, NFKBIA, FOS, HSPA8, FCGR2B, CASP3, TLR4, LOC100511937, CHUK, STAT3, IFNAR2, EIF2S1, IFNAR1, NFKBIB, CSNK2A1, PIK3CA, NFKB1, IRAK4* |
| ssc00983 | Drug metabolism - other enzymes | 44 | 1.708 | 3.61E-03 | 2.22E-02 | *TPMT, NME1, CES1, UPP1, LOC100515741, GSTM3, NME2, CMPK1, GSTO1, UMPS, GSTO2, IMPDH2, GMPS, RRM2B, UCK2* |
| ssc01524 | Platinum drug resistance | 58 | 1.687 | 1.44E-03 | 1.18E-02 | *BIRC3, PMAIP1, CYCS, CDKN1A, FAS, CASP3, LOC100511937, SLC31A1, GSTM3, GSTO1, PIK3CA, ATM, GSTO2, BBC3, MSH2, XPA, TOP2B, MSH3, XIAP, PIK3R2, CASP8, BAK1, BIRC5, TP53* |
| ssc04115 | p53 signaling pathway | 65 | 1.665 | 2.61E-03 | 1.80E-02 | *SERPINE1, IGF1, PMAIP1, CYCS, CDKN1A, FAS, SESN2, THBS1, CASP3, STEAP3, SIAH1, ZMAT3, TP53I3, GADD45A, GADD45B, CCNG1, ATM, GORAB, BBC3* |
| ssc05142 | Chagas disease | 92 | 1.636 | 1.95E-03 | 1.50E-02 | *IL6, IL1B, CXCL8, CCL2, SERPINE1, CCL3L1, IL10, PPP2R2C, FAS, NFKBIA, FOS, IFNG, TLR4, LOC100511937, TGFB3, CHUK, IFNGR1, PIK3CA, NFKB1, IRAK4, IFNGR2, PPP2CA, GNAI1* |

| | | | | | | |
|---|---|---|---|---|---|---|
| ssc05133 | Pertussis | 63 | 1.616 | 3.37E-03 | 2.19E-02 | *IL6, AMCF-II, IL1A, IL1B, CXCL8, IL10, FOS, IRF1, CASP3, TLR4, TICAM2, CASP7* |
| ssc04068 | FoxO signaling pathway | 116 | 1.610 | 6.67E-04 | 6.39E-03 | *IL6, SOD2, PCK2, IGF1, IL10, CDKN1A, HOMER1, PLK3, SGK1, S1PR1, LOC100511937, TGFB3, CHUK, STAT3, PRMT1, LOC100521447, GADD45A, GADD45B, EGF, SMAD3, PIK3CA, ATM, NLK* |
| ssc05146 | Amoebiasis | 84 | 1.609 | 2.51E-03 | 1.77E-02 | *IL6, IL1B, CXCL2, ARG1, CXCL8, IL10, ARG2, IL1R2, IFNG, PRDX1, CASP3, IL1R1, TLR4, LOC100511937, TGFB3* |
| ssc05418 | Fluid shear stress and atherosclerosis | 119 | 1.608 | 1.69E-03 | 1.33E-02 | *IL1A, IL1B, SELE, CCL2, CTSL, IL1R2, FOS, HSP90AA1, IFNG, MMP9, IL1R1, LOC100511937, HSP90B1, CHUK, BMPR2, VCAM1, ICAM1, GSTM3, CALM2, GSTO1, SUMO2, PIK3CA, NFKB1* |
| ssc04210 | Apoptosis | 114 | 1.605 | 1.32E-03 | 1.17E-02 | *BIRC3, CTSL, LOC100127131, PMAIP1, CYCS, FAS, NFKBIA, FOS, DDIT3, LMNB2, CASP3, LOC100511937, CASP7, CHUK, EIF2S1, GADD45A, TUBAL3, GADD45B, SEPTIN4, TRAF2, PIK3CA, ATM, NFKB1, LMNB1, MCL1, BBC3, LOC100739080, KRAS, AIFM1, XIAP, MAP2K1, EIF2AK3, MAPK8, PIK3R2, LMNA, CASP8, BAK1, CFLAR, MAPK9, BIRC5, TP53, DIABLO* |
| ssc04625 | C-type lectin receptor signaling pathway | 90 | 1.577 | 3.57E-03 | 2.22E-02 | *IL6, IL1B, CLEC4E, IL10, CD209, NFKBIA, BCL3, EGR2, NFKB2, PLK3, RELB, PTGS2, IRF1, LOC100511937, CHUK, PPP3CA, CALM2, PTPN11* |
| ssc03018 | RNA degradation | 68 | 1.572 | 5.63E-03 | 3.08E-02 | *BTG3, HSPA9, HSPD1, MPHOSPH6, DCPS, EXOSC3, SKIC8, CNOT7, LSM8, DHX36, EXOSC5, C1D, EXOSC8, EXOSC4, LSM6, LSM5, LSM7, PNPT1, ENO1, PABPC4, MTREX, EXOSC2, LSM2, XRN1, LSM3, EXOSC1, ENO3, DIS3L, SKIC3, EXOSC9, TENT4A, PATL1, ZCCHC7, CNOT9* |
| ssc03420 | Nucleotide excision repair | 53 | 1.558 | 7.39E-03 | 3.76E-02 | *ERCC6, POLD3, POLR2M, GTF2H3, RAD23A, MNAT1, POLR2H, GTF2H5, RFC4, XPA, POLR2G, RAD23B, POLE3, POLR2J, POLR2K, ERCC4, POLR2I, POLR2F, POLR2D, POLR2E, CDK7, RPA3, POLR2B, RFC2, POLE4* |
| ssc05167 | Kaposi sarcoma-associated herpesvirus infection | 156 | 1.551 | 5.33E-04 | 5.41E-03 | *IL6, CXCL2, CXCL8, ANGPT2, CYCS, CDKN1A, FAS, NFKBIA, FGF2, FOS, CLEC2B, GNG4, HIF1A, PTGS2, RCAN1, CASP3, IL6ST, LOC100511937, ZFP36, CHUK, STAT3, IFNAR2, IFNAR1, PPP3CA, ICAM1, CALM2, IFNGR1, TRAF2, PIK3CA, SLA-7, NFKB1, GNG10* |

| | | | | | | |
|---|---|---|---|---|---|---|
| ssc04714 | Thermogenesis | 205 | 1.547 | 2.97E-04 | 3.20E-03 | *NDUFAF4, CREB5, NDUFB2, NDUFB1, NDUFAF5, ACTL6A, NDUFC2, ATP5MC1, ND4L, NDUFB3, COX16, NDUFA6, COX6B, NDUFB9, COX17, COX5B, NDUFS4, COX11, ACSL1, LOC100522678, COA6, ATP5PD, ACSL4, ATP5PF, ATP5MF, UQCRC1, COX7A2, NDUFB5, ATP5F1B, NDUFA5, NDUFA11, ATP5F1C, UQCR11, RPS6KA6, COX7B, ATP5MC3, ATP5ME, COX5A, NDUFB6, NDUFB10, NDUFA9, NDUFA2, KRAS, ATP5F1A, NDUFV2, ATP5PO, PRKG2, UQCR10, NDUFB11, NDUFB7, UQCRQ, NDUFS5, PRKAA2, COX7C, SMARCC1, CREB3, CREB3L1, NDUFAF7, NDUFS1, PPARGC1A, DPF3, SMARCA4, NDUFAF6, NDUFA4, SMARCE1, RPS6KA3, ATP5F1D, ACSL3, NDUFB4, NDUFS8, UQCRC2, SLC25A20, NDUFA8, ACSL6* |
| ssc04630 | JAK-STAT signaling pathway | 110 | 1.529 | 4.71E-03 | 2.66E-02 | *IL6, CSF3, LIF, IL10, CDKN1A, SOCS3, CNTFR, IFNG, OSMR, CISH, IL6ST, LOC100511937, IFNE, STAT3, IFNAR2, IFNAR1, PTPN2, IFNGR1, PTPN11, LOC397451, IL27RA, PIM1, EGF, IL4R, PIK3CA, IFNGR2, SOCS7, IL15, IL10RB, MCL1, SOCS4, IL21R* |
| ssc04066 | HIF-1 signaling pathway | 96 | 1.515 | 6.52E-03 | 3.52E-02 | *IL6, SERPINE1, ANGPT2, EIF4EBP1, IGF1, TIMP1, CDKN1A, TFRC, IFNG, EIF4E, HIF1A, PFKFB3, TLR4, LOC100511937, STAT3, IFNGR1, PGK1, EGF, PIK3CA, NFKB1, IFNGR2, HK2, PDHA1, EDN1, LDHA, MKNK2, VHL, ENO1, ELOC, MAP2K1, PIK3R2, FLT1* |
| ssc05164 | Influenza A | 127 | 1.514 | 3.42E-03 | 2.19E-02 | *IL6, IL1A, IL1B, CXCL8, CCL2, CXCL10, KPNA2, CYCS, FAS, NFKBIA, SOCS3, IFNG, CASP3, TLR4, LOC100511937, DNAJB1, CHUK, IFNAR2, EIF2S1, IL33, IFNAR1, ICAM1, NXT2, NFKBIB, IFNGR1, NXT1, PIK3CA, NFKB1, IRAK4, IFNGR2* |
| ssc05235 | PD-L1 expression and PD-1 checkpoint pathway in cancer | 81 | 1.512 | 7.11E-03 | 3.72E-02 | *BATF3, CD274, NFKBIA, FOS, IFNG, HIF1A, NFKBIE, RASGRP1, TLR4, LOC100511937, TICAM2, CHUK, STAT3, PPP3CA, NFKBIB, IFNGR1, PTPN11, CSNK2A1, EGF, PIK3CA, NFKB1, IFNGR2* |
| ssc05022 | Pathways of neurodegeneration - multiple diseases | 393 | 1.480 | 1.58E-05 | 2.09E-04 | *IL6, IL1A, IL1B, NEFH, LOC100127131, GRIA1, CYCS, FAS, NDUFB2, NDUFB1, DDIT3, PTGS2, NDUFC2, PSMB3, ATP5MC1, RAB39B, ND4L, PSMD14, PPID, UCHL1, NDUFB3, CASP3, PSMD12, KIF5B, CASP7, LRRK2, FZD9, NDUFA6, EIF2S1, COX6B, NDUFB9, SOD1, HSPA5, PSMA3, PPP3CA, WNT9B, PSMA5, COX5B, CALM2, RAB1A, GPX7, NDUFS4, GPX8, TUBAL3,* |

| | | | | | | |
|---|---|---|---|---|---|---|
| | | | | | | *LOC100522678, CSNK2A1, TUBB1, TANK, FZD10, ATP5PD, ATP5PF, PSMB6, ATF6, UQCRC1, DCTN6, COX7A2, TRAF2, NDUFB5, TOMM40, WNT7B, ATP5F1B, NFKB1, PSMD6, NDUFA5, WNT16, TUBB, NDUFA11, PSMC5, ATP5F1C, UQCR11, PSMC6, FZD3, COX7B, ATP5MC3, COX5A, NDUFB6, PSMA4, NDUFB10, PARK7, PSMA2, PSMD8, PSMA1, PRNP, NDUFA9, NDUFA2, PSMC4, KRAS, ATP5F1A, RAB5A, NDUFV2, ATP5PO, UQCR10, NDUFB11, RB1CC1, NDUFB7, UBE2L3, UQCRQ, PSMD2, NDUFS5, PSMC2, PSMB1, RYR3, C9orf72, COX7C, PSMC1, NDUFS1, ATG101, PSMD13, VDAC3, PSMB4, ATXN2L, NDUFA4, CHMP2B, ATP5F1D, MAP2K1, EIF2AK3, MAPK8, VDAC2, PSMD1, PPP3CB, NDUFB4, NDUFS8, PSMD4, UBB, CASP8, UQCRC2, CSNK1A1, NDUFA8, VDAC1, TRPC3, VCP, TUBB4B, MCU, ACTR1A, GNAQ, UBE2J1, SLC25A5, PSMD7, ATP5PB, BAK1, PSMB7, LOC100524873, MAPK9, NDUFS2, PPP3R1, SDHD, HSD17B10, CYC1, NDUFB8, CALM1, NRBF2, APP* |
| ssc01200 | Carbon metabolism | 97 | 1.470 | 9.50E-03 | 4.62E-02 | *PSAT1, PSPH, PHGDH, SHMT2, GPT2, GOT1, GLDC, MDH1, SDS, PGK1, GLUD1, IDH3A, ME1, SUCLA2, HK2, ME2, MDH2, PDHA1, ACAT2, TPI1, PRPS1, PGAM1, PCCB* |
| ssc04933 | AGE-RAGE signaling pathway in diabetic complications | 93 | 1.448 | 8.36E-03 | 4.12E-02 | *IL6, IL1A, IL1B, SELE, CXCL8, CCL2, SERPINE1, CASP3, EGR1, LOC100511937, TGFB3, STAT3, VCAM1, ICAM1, PIM1, SMAD3, PIK3CA, NFKB1* |
| ssc04217 | Necroptosis | 109 | 1.446 | 1.03E-02 | 4.94E-02 | *IL1A, IL1B, BIRC3, TNFAIP3, FAS, H2AZ1, HSP90AA1, IFNG, PPID, TLR4, TICAM2, STAT3, IFNAR2, IL33, IFNAR1, MLKL, IFNGR1, GLUD1, TRAF2, FTH1, IFNGR2, PPIA, VPS4A, AIFM1, H2AX, VDAC3, PGAM5, XIAP, CHMP2B, MAPK8, VDAC2, CASP8, VDAC1, SLC25A5, CFLAR, MAPK9, RIPK3, STAT1, CHMP7, CHMP6, HSP90AB1, JAK2, CHMP4B, TICAM1* |
| ssc04218 | Cellular senescence | 139 | 1.441 | 4.07E-03 | 2.46E-02 | *IL6, IL1A, CXCL8, SERPINE1, EIF4EBP1, CDKN1A, PPID, LOC100511937, TGFB3, PPP3CA, CALM2, GADD45A, GADD45B, E2F5, PPP1CC, SMAD3, PIK3CA, SLA-7, ATM, NFKB1, HIPK3, KRAS, LIN9, PPP1CB, RBL1, CDK4, RRAS2, VDAC3, CCND2, MAP2K1, NBN, VDAC2, PPP3CB, PIK3R2, TGFBR1, VDAC1, RAD1, MCU, SLC25A5, HIPK1, E2F3, MRAS, RBBP4, PPP3R1, MYC, LIN54, E2F4, CALM1, TP53, CDK2, MAP2K3* |

| | | | | | | |
|---|---|---|---|---|---|---|
| ssc05161 | Hepatitis B | 141 | 1.375 | 8.30E-03 | 4.12E-02 | *IL6, CXCL8, CYCS, CDKN1A, FAS, NFKBIA, CREB5, FOS, EGR2, DDX3X, CASP3, MMP9, TLR4, LOC100511937, TICAM2, TGFB3, CHUK, STAT3, IFNAR1, SMAD3, PIK3CA, NFKB1, IRAK4, KRAS, IRAK1, MAP3K7, SMAD4, CREB3, CREB3L1, VDAC3, TIRAP, MAP2K1, MAPK8, PIK3R2, TGFBR1, IKBKE, CASP8, E2F3, MAPK9, STAT1, BIRC5, MYC, TP53, CDK2, YWHAZ, MAP2K3, JAK2, YWHAQ, TICAM1, MAPK14, PRKCA* |
| ssc05132 | Salmonella infection | 215 | 1.332 | 7.41E-03 | 3.76E-02 | *IL6, IL1B, CXCL8, BIRC3, LOC100127131, CYCS, NFKBIA, FOS, HSP90AA1, RHOJ, CASP3, RIPK2, ARPC5L, TLR4, KPNA4, HSP90B1, KIF5B, CASP7, CHUK, RALA, PAK3, LOC100522887, ARF2, RHOB, MLKL, TUBAL3, DYNLRB1, TUBB1, DCTN6, TRAF2, ROCK2, PIK3CA, NFKB1, IRAK4, DYNC1LI2, TUBB, NCKAP1, PFN2, RAB5A, IRAK1, MAP3K7, ACTR3, ARF6, DYNC1LI1, IL18, KPNA1, ACBD3, CSE1L, EXOC5, TIRAP, MAP2K1, MAPK8, DYNC1I2, CASP8, ARPC3, TUBB4B, ACTR1A, CDC42, ACTR2, EXOC4, PIK3C2A, CD14, BAK1, MAPK9, PFN1, SNX18, RIPK3, RHOA, MYC* |
| ssc04082 | Neuroactive ligand signaling | 116 | -1.491 | 6.93E-03 | 3.68E-02 | *PLCB4, ADRA1B, ADCY7, SLC6A13, GRIK5, P2RY11, POMC, LOC100621431, P2RX3, P2RX1, HTR4, ADORA2A, ADORA3, P2RY14, GRM8, ADRA1D, SLC1A7, GABRG1, P2RY13, PRKCG, GNG8, GRIN3B, GABBR1, ADRA2B, ADCY5, GRM2, OPRL1, PRKCB, P2RY6, GABRR3, GRIK4, PLCB2, ADRA2C, P2RY12, SLC6A4, SLC6A2, P2RX6, GABRD, ADCY8* |
| ssc04142 | Lysosome | 219 | -1.544 | 8.21E-04 | 7.45E-03 | *SLC36A1, ABCA5, VPS11, LAPTM4B, GBA1, FIG4, OSBPL1A, GNPTG, MEAK7, IDUA, ROGDI, MAN2B1, AP3M2, ACP2, MCOLN1, AP5S1, SLC29A3, VPS39, NPC2, FUCA2, SLC29A1, LIPA, PPT1, SGSH, AP2A2, ATP6V1C2, CTSO, TELO2, ACP5, ZFYVE26, AP1G2, TPCN1, DNASE2, OSBPL5, CLCN7, CTSS, SLC11A1, CLN6, ATP6V0A4, ATP6V1G2, CTSB, NAGPA, ASAH1, ATP6V0A1, GGA2, NAGA, TMEM175, OSBPL7, CTSZ, TPP1, MITF, MCOLN2, PLEKHM1, LAPTM5, CTSD, CD68, ABCB9, NAPSA, CTSW, TPCN2, ABCA2, ARSA, GALNS, TLR9, CTSF, LOC100520753, TLR7, MANBA, GM2A, TLR8* |
| ssc04148 | Efferocytosis | 138 | -1.568 | 4.32E-03 | 2.55E-02 | *MAPK12, SLC66A1, CPT1B, MERTK, NFATC1, AXL, STAB2, CAMK2A, PLA2G6, CAMK2G, ADGRB1, TYRO3, CASP1, STAB1, ANO3, LOC100523747, ANO9, C1QA, LOC102161654, GAS6,* |

| | | | | | | |
|---|---|---|---|---|---|---|
| | | | | | | *LOC100620540, C1QB, C1QC, P2RY6, ATP2A3, P2RY12, LOC100511343, TIMD4, CX3CR1, ALOX5* |
| ssc05414 | Dilated cardiomyopathy | 81 | -1.622 | 4.41E-03 | 2.55E-02 | *ITGB5, CACNA1C, ADCY3, TGFB2, ADRB1, DTNA, ITGA3, ADCY7, ITGA9, CACNB4, TPM1, CACNB2, TNNI3, CACNA1F, ITGA8, CACNA2D2, ITGB4, AGT, LAMA1, ADCY5, CACNA1D, CACNG6, CACNG5, ACTC1, ATP2A3, SGCA, SLC8A2, LOC100736765, ITGB7, ADCY8* |
| ssc04081 | Hormone signaling | 128 | -1.629 | 1.58E-03 | 1.27E-02 | *INSR, PLCB4, LEPR, RAPGEF4, HEPH, ADRA1B, ADCY7, GUCY2C, NPR2, AMH, PRLR, ADRB3, POMC, LOC100621431, GPER1, GNA15, AGTR1, AMHR2, IRS2, SLC16A10, LHB, RAMP2, THRA, RAPGEF3, ADRA1D, PIK3CD, VIPR2, SLC16A2, TSHR, PTH1R, AGT, GNG8, NPR1, GIPR, ADRA2B, ADCY5, OPRL1, SLC40A1, CRHR2, GCGR, THPO, PLCB2, ADRA2C, ESR1, VIPR1, ADCY8* |
| ssc04911 | Insulin secretion | 62 | -1.639 | 5.23E-03 | 2.91E-02 | *CACNA1F, ITPR3, STX1A, CREB3L3, ATP1A2, CAMK2A, CAMK2G, VAMP2, PRKCG, TRPM4, ADCY5, PCLO, CACNA1D, KCNMB4, PRKCB, PLCB2, ADCY8, ATP1B2* |
| ssc04925 | Aldosterone synthesis and secretion | 79 | -1.707 | 2.08E-03 | 1.53E-02 | *CACNA1C, LIPE, ADCY3, KCNK3, PLCB4, ADCY7, DAGLA, CACNA1H, POMC, AGTR1, CACNA1F, ITPR3, CREB3L3, ATP1A2, CALML4, CAMK2A, CAMK2G, AGT, PRKCG, NPR1, CYP11A1, ADCY5, ATP2B3, CACNA1D, PRKCB, PDE2A, PLCB2, ADCY8, ATP1B2, CAMK1G* |
| ssc00430 | Taurine and hypotaurine metabolism | 14 | -1.736 | 4.43E-03 | 2.55E-02 | *FMO3, CSAD, FMO4, GGT7, FMO2, GGT1, FMO5, FMO1, LOC100523745, GADL1* |
| ssc04973 | Carbohydrate digestion and absorption | 32 | -1.788 | 2.72E-03 | 1.84E-02 | *PIK3CD, ATP1A2, TAS1R3, SLC2A5, CACNA1D, GNAT3, PRKCB, PLCB2, ATP1B2, MGAM* |
| ssc04740 | Olfactory transduction | 38 | -1.819 | 1.39E-03 | 1.17E-02 | *CALML4, CAMK2A, CAMK2G, CNGA2, PDE1B, PDE2A, SLC8A2, SLC24A4, LOC100736710, LOC100738112* |
| ssc04020 | Calcium signaling pathway | 186 | -1.831 | 4.16E-06 | 6.53E-05 | *ORAI2, PDGFC, FLT4, CACNA1A, PDE1A, CACNA1C, P2RX5, FGFR2, ADCY3, GRIN2D, MCOLN1, EGFR, ADRB1, ITPKB, CHRM2, PTK2B, PLCB4, FGF1, ADRA1B, ADCY7, CYSLTR1, CACNA1H, TGFA, ADRB3, PLCG1, TPCN1, P2RX3, PDGFA, GNA15, AGTR1, CD38, ERBB3,* |

| | | | | | | |
|---|---|---|---|---|---|---|
| | | | | | | *FGFR3, TBXA2R, P2RX1, HTR4, ERBB2, PLCG2, FGFR4, PLCD3, ITPKA, CACNA1F, ADORA2A, ITPR3, ADRA1D, HRC, GDNF, MCOLN2, NFATC1, CALML4, CAMK2A, PHKA2, CAMK2G, TNNC2, CASQ1, MST1, MST1R, CACNA1E, FGF21, PRKCG, CYSLTR2, TPCN2, PHKG1, GRIN3B, ATP2B3, CACNA1D, PLCD1, MYLK4, PRKCB, CACNA1B, PDE1B, P2RX7, ATP2A3, LTB4R2, SLC8A2, PLCB2, P2RX6, NTRK3, ADCY8, CAMK1G, VEGFD* |
| ssc02010 | ABC transporters | 40 | -1.870 | 5.83E-04 | 5.74E-03 | *ABCG5, ABCC11, ABCC10, ABCG2, ABCA7, ABCB9, ABCC6, ABCA2, ABCC5, ABCA4, ABCA14, ABCA15, ABCA6, ABCA9, ABCA8* |
| ssc04080 | Neuroactive ligand-receptor interaction | 149 | -1.904 | 1.58E-06 | 2.73E-05 | *PRLR, TSPO, F2RL1, ADRB3, P2RY11, POMC, KNG1, P2RX3, AGTR1, TBXA2R, LHB, P2RX1, HTR4, GALR1, P2RY10, LYPD6, THRA, ADORA2A, ADORA3, P2RY14, GRM8, ADRA1D, P2RY8, GABRG1, S1PR4, RXFP4, NPFF, VIPR2, TSHR, PTH1R, GRID2, NMUR1, P2RY13, AGT, PAQR6, GPR156, F2, CYSLTR2, GIPR, GRIN3B, GABBR1, ADRA2B, CNR2, GRM2, OPRL1, APLNR, P2RY6, GABRR3, P2RX7, CRHR2, GCGR, GRID1, LTB4R2, GRIK4, ADRA2C, CNR1, VIPR1, P2RX6, GABRD, EDN3, GLP2R, CHRNA1* |

Abbreviations: ABC: ATP-binding cassette; AGE-RAGE: Advanced glycation end products-Receptor for AGEs; FDR: False discovery rate; FoxO: Forkhead box O; HIF-1: Hypoxia-inducible factor 1; IL-17: Interleukin-17; JAK-STAT: Janus kinases-signal transducer and activator of transcription proteins; KEGG: Kyoto Encyclopedia of Genes and Genomes; NES: Normalized enrichment score; NF-kappa B: Nuclear factor-kappa B; PD-1: Programmed cell death protein 1; PD-L1: Programmed cell death ligand 1; TNF: Tumor necrosis factor.

**Supplementary Table S8. Significantly enriched pathways from the GSEA result of the GO:BP database, High-glucose infection versus Low-glucose infection.** Abbreviations: GO:BP: Biological Processes in Gene Ontology; GSEA: Gene set enrichment analysis.

| GO:BP ID | Description | Set Size | NES | P-value | FDR | Core enrichment gene |
|---|---|---|---|---|---|---|
| GO:0022613 | Ribonucleoprotein complex biogenesis | 172 | 2.957 | 1.00E-10 | 7.12E-09 | *NPM3, SNRPF, SNRPD1, NHP2, GAR1, FBL, NIP7, EMG1, EIF3J, DDX3X, RRS1, WDR12, LYAR, ESF1, RRP9, SNRPG, NOPCHAP1, DIMT1, NOB1, NOP9, WDR55, WDR36, RSL24D1, UTP6, NLE1, PTGES3, REXO4, RRP36, BMS1, MRTO4, SNRPE, WDR75, EXOSC3, CLNS1A, AK6, GRWD1, WDR74, GEMIN6, EIF6, PPAN, NOP56, EIF3A, WDR46, STRAP, DDX52, EIF3I, NOP2, UTP18, KRR1, SNRPD3, DENR, RRP15, TAF9, ABT1, DDX49, TMA16, IMP3, RCL1, UTP11, SNRPC, PNO1, NUFIP1, RPL27, EBNA1BP2, FASTKD2, UTP23, LOC100738684, NPM1, UTP15, EXOSC5, EIF3M, C1D, CLP1, EXOSC4, LSM6, RPL7L1, BUD23, SRSF1, GTPBP4, EIF4A3, MRM1, PWP1, GTPBP10, SHQ1, USP39, RPS21, GEMIN2, NAF1, SF3A3, MTREX, TRMT112, NOL6, MRPS7, FCF1, RIOK1, DDX31, EXOSC2, SNU13, NAT10* |
| GO:0042254 | Ribosome biogenesis | 129 | 2.842 | 1.00E-10 | 7.12E-09 | *NPM3, NHP2, GAR1, FBL, NIP7, EMG1, DDX3X, RRS1, WDR12, LYAR, ESF1, RRP9, DIMT1, NOB1, NOP9, WDR55, WDR36, RSL24D1, UTP6, NLE1, REXO4, RRP36, BMS1, MRTO4, WDR75, EXOSC3, AK6, GRWD1, WDR74, EIF6, PPAN, NOP56, WDR46, DDX52, NOP2, UTP18, KRR1, RRP15, ABT1, DDX49, TMA16, IMP3, RCL1, UTP11, PNO1, RPL27, EBNA1BP2, FASTKD2, UTP23, NPM1, UTP15, EXOSC5, C1D, EXOSC4, LSM6, RPL7L1, BUD23, GTPBP4, EIF4A3, MRM1, PWP1, GTPBP10, RPS21, NAF1, MTREX, TRMT112, NOL6, MRPS7, FCF1, RIOK1, DDX31, EXOSC2, SNU13, NAT10, RPL35A, RPS23, SBDS, SDAD1, NOP10, TSR2, MTERF3, RPP40, USP16, SDE2, MYG1, KRI1, POP7, NOC4L, WDR18, LOC100621115* |

| | | | | | | |
|---|---|---|---|---|---|---|
| GO:0006396 | RNA processing | 357 | 2.531 | 1.00E-10 | 7.12E-09 | *WT1, NPM3, SNRPF, SNRPD1, NHP2, GAR1, WDR4, SLC38A2, FBL, EMG1, PAPOLA, RTCA, RRS1, WDR12, LYAR, RBM8A, ESF1, RRP9, SNRPG, SNRNP27, QTRT1, DCPS, ELP4, DIMT1, NOB1, NOP9, TYW1, SNRPB2, WDR55, WDR36, UTP6, SMN1, REXO4, RRP36, PUS3, SNRPE, WDR75, LEO1, EXOSC3, CLNS1A, AK6, SF3B6, WDR74, SMNDC1, GEMIN6, DKC1, PPAN, RTCB, AGO2, CSTF2, ISG20L2, WDR46, STRAP, DDX52, NOP2, UTP18, KRR1, SNRPD3, SSB, MAGOH, RRP15, QTRT2, METTL8, ABT1, MAGOHB, DDX49, TXNL4A, IMP3, RAMAC, RCL1, UTP11, CWC15, RBM7, SNRPA1, SNRPC, TP53RK, ZPR1, LSM8, TYW5, RPRD1B, INTS8, RPL27, CDC40, MFAP1, FASTKD2, UTP23, ZMAT2, UTP15, EXOSC5, THUMPD1, ELP1, C1D, TRMT6, CLP1, EXOSC4, MRPL44, LSM6, CSTF2T, TRA2B, RPL7L1, LSM5, NSUN2, PRPF40A, BUD23, HNRNPK, SRSF1, GTPBP4, RBM42, EIF4A3, MRM1, SMU1, PWP1, PNPT1, USP39, PPIL1, RPS21, WTAP, TSEN15, SCNM1, GEMIN2, NCBP2, ELP5, NAF1, SF3A3, RBM15, TSNAX, BUD31, THOC2, SNRPA, MTREX, TRMT112, NOL6, UBL5, FCF1, RBMX2, RPP30, EXOSC2, SNU13, PTBP3, NAT10, METTL3, RBM41, CWC22, FTSJ1, LOC100738822, SRSF7, TPRKB, TSN, RPL35A, LSM2, NSRP1, HNRNPF, HSD17B10, SBDS, RPP38, LSM3, CSTF1, TRMT5, SYMPK, PRPF19, EXOSC1, NOP10, SNRPD2, TRMT11, THOC7, ELP2, TSR2, RPP40, QKI, INTS10, SCAF11, POP1, PAPOLG, SDE2, MYG1, SF3B4, TRMT2A, CSTF3, KRI1, RPP21, SNRNP40, POP7, U2AF2, FRA10AC1, WBP11, CPSF6, WDR18, PRKRA, RNPS1, PUS1, C1H9orf78, ADAT2, SNRNP35, NOL9, RBMX, RPP25L, EXOSC9, MBNL1, RPS25, SLX9* |
| GO:0000375 | RNA splicing, via transesterification reactions | 107 | 2.309 | 1.90E-09 | 5.64E-08 | *SNRPF, SNRPD1, NHP2, RBM8A, SNRPG, DCPS, SNRPB2, SNRPE, CLNS1A, SF3B6, GEMIN6, STRAP, SNRPD3, TXNL4A, CWC15, RBM7, SNRPA1, SNRPC, LSM8, CDC40, MFAP1, ZMAT2, LSM6, TRA2B, LSM5, PRPF40A, SRSF1, RBM42, EIF4A3, SMU1, USP39, PPIL1, WTAP, GEMIN2, NCBP2, SF3A3, RBM15, BUD31, SNRPA, UBL5, RBMX2, SNU13, METTL3, RBM41, CWC22, SRSF7, LSM2, NSRP1, LSM3, PRPF19* |
| GO:0000377 | RNA splicing, via transesterification | 107 | 2.309 | 1.90E-09 | 5.64E-08 | *SNRPF, SNRPD1, NHP2, RBM8A, SNRPG, DCPS, SNRPB2, SNRPE, CLNS1A, SF3B6, GEMIN6, STRAP, SNRPD3, TXNL4A, CWC15, RBM7, SNRPA1, SNRPC, LSM8, CDC40,* |

| | | | | | | |
|---|---|---|---|---|---|---|
| | reactions with bulged adenosine as nucleophile | | | | | *MFAP1, ZMAT2, LSM6, TRA2B, LSM5, PRPF40A, SRSF1, RBM42, EIF4A3, SMU1, USP39, PPIL1, WTAP, GEMIN2, NCBP2, SF3A3, RBM15, BUD31, SNRPA, UBL5, RBMX2, SNU13, METTL3, RBM41, CWC22, SRSF7, LSM2, NSRP1, LSM3, PRPF19* |
| GO:0000398 | MRNA splicing, via spliceosome | 107 | 2.309 | 1.90E-09 | 5.64E-08 | *SNRPF, SNRPD1, NHP2, RBM8A, SNRPG, DCPS, SNRPB2, SNRPE, CLNS1A, SF3B6, GEMIN6, STRAP, SNRPD3, TXNL4A, CWC15, RBM7, SNRPA1, SNRPC, LSM8, CDC40, MFAP1, ZMAT2, LSM6, TRA2B, LSM5, PRPF40A, SRSF1, RBM42, EIF4A3, SMU1, USP39, PPIL1, WTAP, GEMIN2, NCBP2, SF3A3, RBM15, BUD31, SNRPA, UBL5, RBMX2, SNU13, METTL3, RBM41, CWC22, SRSF7, LSM2, NSRP1, LSM3, PRPF19* |
| GO:0006412 | Translation | 229 | 2.279 | 1.00E-10 | 7.12E-09 | *WT1, GARS1, EIF4EBP1, WARS1, YARS1, EIF2S2, NARS1, SESN2, THBS1, EIF3J, DDX3X, AIMP2, C7H6orf125, ZFP36, EIF1, MRPS17, RSL24D1, FARSA, EIF2S1, MRPL35, BZW1, NIBAN1, MRPL52, RPS27L, PUM3, MRPL2, AGO2, MRPL13, RIDA, EIF3A, EIF3I, GFM2, DENR, MRPL20, RARS1, SSB, ETF1, EIF4A1, MRPS18C, SKIC8, ATM, MRPS11, DRG1, IGFBP5, MRPL17, PAIP1, NEMF, DDRGK1, VARS1, MRPL15, QRSL1, MRPL32, RPL27, MRPL47, FASTKD2, LOC100738684, NPM1, DARS1, EIF3M, MRPL44, PAIP2B, CAPRIN1, ABCF1, EIF4A3, MRPS5, EIF2S3, RPS21, MRPL24, NCBP2, RPL26L1, SERBP1, SRP9, MRPL36, LTN1, GFM1, MRPS7, CALR, MTRF1, GATC, HARS1, NAT10, METTL3, FXR1, FTSJ1, MRPL18, MRPL58, RPL35A, PPP1R15A, RGS2, RPS23, CHCHD1, APP, ELAVL1, EIF5A, RNF14, RPL23A, MRPS18A, MRPL11, QKI, LSM14A, IGF2BP1, USP16, MRPL43, RPL38, RPL23, MRPL33, PTCD3, IMPACT, MRPL3, RPLP1, GAPDH, RPS27, NCK1, UPF3B, DHX29, MRPL27, RPL21, FXR2, RPS18, RPS24, RPS6KB1, RPL14, RPLP2, MRPL1, RPL10A, NARS2, DDX6, CSDE1* |
| GO:0008380 | RNA splicing | 154 | 2.210 | 8.46E-10 | 4.94E-08 | *WT1, SNRPF, SNRPD1, NHP2, SLC38A2, RBM8A, SNRPG, SNRNP27, DCPS, SNRPB2, SMN1, SNRPE, CLNS1A, SF3B6, SMNDC1, GEMIN6, RTCB, STRAP, SNRPD3, MAGOH, MAGOHB, TXNL4A, CWC15, RBM7, SNRPA1, SNRPC, ZPR1, LSM8, CDC40, MFAP1, ZMAT2, CLP1, LSM6, TRA2B, LSM5, PRPF40A, HNRNPK, SRSF1, RBM42, EIF4A3, SMU1, USP39, PPIL1, WTAP, TSEN15, SCNM1, GEMIN2, NCBP2, SF3A3, RBM15, BUD31, SNRPA, UBL5, RBMX2,* |

| | | | | | | |
|---|---|---|---|---|---|---|
| | | | | | | *SNU13, PTBP3, METTL3, RBM41, CWC22, LOC100738822, SRSF7, LSM2, NSRP1, HNRNPF, LSM3, PRPF19, SNRPD2* |
| GO:0016071 | MRNA metabolic process | 244 | 2.130 | 1.00E-10 | 7.12E-09 | *SNRPF, SNRPD1, NHP2, CEBPG, DNAJB11, PAPOLA, RBM8A, SNRPG, SNRNP27, DCPS, SNRPB2, ZFP36, SMN1, PUS3, SNRPE, LEO1, EXOSC3, CLNS1A, SF3B6, SMNDC1, GEMIN6, AGO2, RIDA, CSTF2, STRAP, SNRPD3, SSB, ETF1, TRAF2, TAF9, SKIC8, ATM, ZBTB1, TXNL4A, MED1, CNOT7, SMG8, PDE12, PAIP1, CWC15, RBM7, SNRPA1, SNRPC, LSM8, RPRD1B, CDC40, HIPK3, MFAP1, FASTKD2, PELO, ZMAT2, NPM1, PNRC2, CLP1, EXOSC4, LSM6, CSTF2T, TRA2B, LSM5, PRPF40A, ROCK1, HNRNPK, CAPRIN1, SRSF1, RBM42, EIF4A3, SMU1, PNPT1, USP39, PPIL1, WTAP, SCNM1, GEMIN2, NCBP2, SF3A3, TIRAP, RBM15, PABPC4, BUD31, THOC2, SNRPA, UBL5, DXO, RBMX2, SNU13, PTBP3, METTL3, RBM41, CWC22, VDR, SRSF7, LSM2, NSRP1, HNRNPF, LSM3, APP, CSTF1, ELAVL1, SYMPK, PRPF19, ARID5A, SNRPD2, THOC7, QKI, IGF2BP1, SCAF11, PAPOLG, SDE2, MYG1, SF3B4, CSTF3, SNRNP40, U2AF2, FRA10AC1, CPSF6, RNPS1, PUS1, C1H9orf78, UPF3B, SNRNP35, RBMX* |
| GO:0006397 | MRNA processing | 169 | 2.105 | 3.21E-09 | 8.79E-08 | *SNRPF, SNRPD1, NHP2, PAPOLA, RBM8A, SNRPG, SNRNP27, DCPS, SNRPB2, SMN1, SNRPE, LEO1, CLNS1A, SF3B6, SMNDC1, GEMIN6, CSTF2, STRAP, SNRPD3, TXNL4A, CWC15, RBM7, SNRPA1, SNRPC, LSM8, RPRD1B, CDC40, MFAP1, ZMAT2, CLP1, LSM6, CSTF2T, TRA2B, LSM5, PRPF40A, HNRNPK, SRSF1, RBM42, EIF4A3, SMU1, USP39, PPIL1, WTAP, SCNM1, GEMIN2, NCBP2, SF3A3, RBM15, BUD31, THOC2, SNRPA, UBL5, RBMX2, SNU13, PTBP3, METTL3, RBM41, CWC22, SRSF7, LSM2, NSRP1, HNRNPF, LSM3, CSTF1, SYMPK, PRPF19, SNRPD2, THOC7, QKI, SCAF11, PAPOLG, SDE2, MYG1, SF3B4, CSTF3, SNRNP40, U2AF2, FRA10AC1, CPSF6, RNPS1, C1H9orf78, SNRNP35, RBMX, IK, SRSF2, TBRG4, CDK12, PSIP1, RBM14, NUDT21, XRN2, SRSF6, ZRSR2, RBM17* |
| GO:0019221 | Cytokine-mediated signaling pathway | 129 | 2.073 | 6.77E-08 | 1.42E-06 | *IL6, AMCF-II, IL1A, IL1B, CXCL8, CCL2, FOSL1, DUOX1, TIFA, MMP8, CXCL10, DUOX2, IL10, PPBP, CXCL13, ACKR1, NFKBIA, IFNG, CCL8, IRF1, TXNDC17, CXCL11, EGR1,* |

| | | | | | | |
|---|---|---|---|---|---|---|
| | | | | | | *CISH, TNFSF13B, TICAM2, IFNE, STAT3, CD40, IFNAR1, ADAM17, IFNGR1, IL4R, TRAF2, CDC37, IL15, MED1, CNOT7* |
| GO:0034097 | Response to cytokine | 214 | 2.036 | 6.46E-09 | 1.53E-07 | *IL6, AMCF-II, IL1A, IL1B, SELE, CXCL8, GFPT2, CCL2, UBD, FOSL1, DUOX1, TIFA, MMP8, CXCL10, DUOX2, MSC, IL10, PPBP, TIMP1, CXCL13, CACYBP, ACKR1, NFKBIA, NFIL3, FABP4, IFNG, STIP1, CCL8, IRF1, TXNDC17, CXCL11, EGR1, CISH, TNFSF13B, TLR4, TICAM2, IFNE, ZFP36, STAT3, CD40, IFNAR1, ADAM17, IFNGR1, PIM1, IL4R, PDCD10, SLC12A2, TRAF2, TAF9, CDC37, IL15, MED1, CNOT7, MCL1, PDE12, SOCS4, MRPL15, CCL4, RNF138, SOCS5, NFYB, IRAK1, IRF8, MAP3K7, MYNN* |
| GO:1901652 | Response to peptide | 214 | 2.036 | 6.46E-09 | 1.53E-07 | *IL6, AMCF-II, IL1A, IL1B, SELE, CXCL8, GFPT2, CCL2, UBD, FOSL1, DUOX1, TIFA, MMP8, CXCL10, DUOX2, MSC, IL10, PPBP, TIMP1, CXCL13, CACYBP, ACKR1, NFKBIA, NFIL3, FABP4, IFNG, STIP1, CCL8, IRF1, TXNDC17, CXCL11, EGR1, CISH, TNFSF13B, TLR4, TICAM2, IFNE, ZFP36, STAT3, CD40, IFNAR1, ADAM17, IFNGR1, PIM1, IL4R, PDCD10, SLC12A2, TRAF2, TAF9, CDC37, IL15, MED1, CNOT7, MCL1, PDE12, SOCS4, MRPL15, CCL4, RNF138, SOCS5, NFYB, IRAK1, IRF8, MAP3K7, MYNN* |
| GO:0071345 | Cellular response to cytokine stimulus | 199 | 1.983 | 3.24E-08 | 7.20E-07 | *IL6, AMCF-II, IL1A, IL1B, CXCL8, GFPT2, CCL2, FOSL1, DUOX1, TIFA, MMP8, CXCL10, DUOX2, MSC, IL10, PPBP, CXCL13, CACYBP, ACKR1, NFKBIA, NFIL3, FABP4, IFNG, STIP1, CCL8, IRF1, TXNDC17, CXCL11, EGR1, CISH, TNFSF13B, TLR4, TICAM2, IFNE, ZFP36, STAT3, CD40, IFNAR1, ADAM17, IFNGR1, PIM1, IL4R, PDCD10, SLC12A2, TRAF2, CDC37, IL15, MED1, CNOT7, PDE12, SOCS4, MRPL15, CCL4, RNF138, SOCS5, NFYB, IRAK1, IRF8, MAP3K7, MYNN* |
| GO:0001701 | In utero embryonic development | 110 | 1.920 | 7.15E-06 | 8.78E-05 | *PSPH, GJA1, FOSL1, LIF, EIF2S2, IL10, BMP2, RSPO3, JUNB, PHLDA2, RCN1, ACTL6A, NLE1, GATA1, SF3B6, WDR74, BMI1, GNA13, PRDM1, DAD1, KAT2A, LIG4, MED1, CEBPB, MED21, PLPP4, ZPR1, TMEM100, SMAD4, RPL7L1, MIB1, MAFF, XRCC2, MATR3, ZFAND5, SLC25A20, CSF2, SLC39A1, BMPR1A, UBE2B, APBA3, SBDS, MDFI* |

| | | | | | | |
|---|---|---|---|---|---|---|
| GO:0015980 | Energy derivation by oxidation of organic compounds | 113 | 1.882 | 2.42E-05 | 2.09E-04 | *IGF1, CYCS, MTFR1, PHLDA2, COQ10B, NDUFC2, ND4L, MDH1, PGM2, C7H6orf125, PTGES3, NDUFB9, COX5B, IDH3A, COA6, ATP5MF, COX7A2, PPP1CC, NDUFA5, UQCR11, SLC25A33, COX7B, ATP5ME, COX5A, NDUFB6, PDHA1, PARK7, COQ10A, NDUFV2, ATP5PO, UQCR10, UQCRQ, PPP1CB, COX7C, UGP2, NDUFS1, PPARGC1A, CS, NDUFS8, IDE, TIGAR, ATP5PB, LOC100524873, SDHAF2, CHCHD2, RHOA, IDH3B, SDHD, CYC1, NDUFB8, NDUFV1, PHKG2, ETFB* |
| GO:0043065 | Positive regulation of apoptotic process | 130 | 1.863 | 9.92E-06 | 1.10E-04 | *WT1, INHBA, CCL2, AGTR2, ATF3, S100A8, RNF122, IL10, CD274, BMP2, PLA2G1B, THBS1, PHLDA2, HSPA9, IFNG, DDIT3, TNFRSF12A, DDX3X, CASP3, TLR4, SIAH1, RASSF2, CD40, RHOB, GADD45B, TRAF2, MCL1, UTP11, PARK7, BBC3, COPS5, PRNP* |
| GO:0097190 | Apoptotic signaling pathway | 153 | 1.845 | 9.37E-06 | 1.08E-04 | *IL1A, INHBA, AGTR2, ATF3, IGF1, S100A8, TNFAIP3, IL10, THBS1, IFNG, DDIT3, PSMD10, DDX3X, DDIT4, GHITM, NR4A2, TLR4, HINT1, SIAH1, AEN, GATA1, SNAI2, RPS27L, BMI1, PDCD10, PSME3, TRAF2, ATM, WNT16, CEBPB, MCL1, PARK7, BBC3, SELENOS, PPIA, SMAD4, FEM1B, IL18, SIAH2, GSKIP, NRP1, MIF, BCLAF1, VDAC2, FAM162A, NHERF1, UBB, CASP8, SLC25A5, SP1, HYAL2, CFLAR, MAPK9, SRPX, RIPK3, MYC, TP53, BCAP31, CTSC, JAK2, ING2, TICAM1, BECN1, NKX3-1, BCL10* |
| GO:2001233 | Regulation of apoptotic signaling pathway | 108 | 1.831 | 3.14E-05 | 2.43E-04 | *INHBA, AGTR2, ATF3, IGF1, S100A8, TNFAIP3, IL10, THBS1, DDIT3, PSMD10, DDX3X, GHITM, NR4A2, TLR4, SIAH1, GATA1, SNAI2, BMI1, PSME3, TRAF2, WNT16, MCL1, PARK7, BBC3, SELENOS, PPIA, FEM1B, SIAH2, NRP1, MIF, BCLAF1, NHERF1, UBB, SLC25A5, SP1, HYAL2, CFLAR, MAPK9, SRPX, RIPK3, TP53, BCAP31, CTSC, JAK2, ING2, BECN1, NKX3-1, BCL10* |
| GO:0043068 | Positive regulation of programmed cell death | 134 | 1.805 | 1.74E-05 | 1.72E-04 | *WT1, INHBA, CCL2, AGTR2, ATF3, S100A8, RNF122, IL10, CD274, BMP2, PLA2G1B, THBS1, PHLDA2, HSPA9, IFNG, DDIT3, TNFRSF12A, DDX3X, CASP3, TLR4, SIAH1, RASSF2, CD40, RHOB, GADD45B, TRAF2, MCL1, UTP11, PARK7, BBC3, COPS5, PRNP* |
| GO:0031399 | Regulation of protein | 104 | 1.763 | 8.06E-05 | 5.74E-04 | *LIF, TNFAIP3, PLA2G1B, IFNG, PSMD10, DDX3X, GNL3, CRY1, EGR1, NDFIP2, NCOA7, GATA1, UBE2D1, PTPN11, FBXO4, DCUN1D5, HSPB1, PRKAR2A, RNF180, KAT2A, IL15, PARK7, PRNP, PPIA, DR1, GTPBP4, PHF23, FYN, NRP1, PRKAA2, ARHGEF2, FBXO33,* |

| | | | | | | |
|---|---|---|---|---|---|---|
| | modification process | | | | | *SPRED1, UBB, ITGA5, MBIP, MAPK9, UBE2B, RAMP1, JAK2, BAG2, TICAM1, UBE3A, DCUN1D4, BCL10, ZZZ3, HBEGF, TAOK1* |
| GO:0008219 | Cell death | 463 | 1.747 | 1.11E-09 | 4.94E-08 | *IL1A, WT1, GSDMA, INHBA, CCL2, AGTR2, ATF3, ATF5, IGF1, S100A8, CXCL10, TNFAIP3, RNF122, IL10, CYCS, CD274, ARG2, HIGD1A, BMP2, PLA2G1B, FAS, THBS1, HSP90AA1, PHLDA2, HSPA9, IFNG, PLK3, HTR2B, TNFAIP8L3, SGK1, TCIM, DDIT3, TNFRSF11B, TNFRSF12A, PSMD10, DDX3X, DDIT4, CSRNP1, GHITM, NDNF, IRF1, NOD2, AIMP2, CASP3, EGR1, NR4A2, RIPK2, TLR4, CRYAB, HINT1, ZFP36, SIAH1, MAD2L1, RASSF2, AEN, CD40, GATA1, SOD1, BMPR2, SNAI2, LOC100522887, RHOB, RPS27L, MLKL, IL27RA, PIM1, BMI1, GADD45B, PDCD10, NME2, LGALS1, TNIP2, PSME3, HSPB1, BNIP1, TRAF2, TAF9, FTH1, DAD1, ATM, WNT16, LIG4, MED1, CEBPB, APIP, ST3GAL1, MCL1, PDCL3, EBAG9, UTP11, FZD3, ZPR1, PARK7, BBC3, COPS5, SELENOS, PRNP, HIPK3, FASTKD2, PPIA, KRAS, NPM1, ETV6, MAP3K7, ARF6, SMAD4, API5, FEM1B, IL18, PRDX3, BNIPL, NPPC, BCL2L12, KPNA1, ROCK1, SIAH2, USP53, AIFM1, GSKIP, NRP1, PRKAA2, MIF, ANGPTL4, PPARGC1A, ATG5, CCND2, XIAP, BCLAF1, MAPK8, VDAC2, FAM162A, NHERF1, TGFBR1, XRCC2, SGMS1, UBB, CASP8, VDAC1, TCHP, ITGA5, TIGAR, RAD21, SLC25A5, SP1, VDR, PEA15, ZFAND6, HYAL2, BAK1, GPX4, NTN1, CFLAR, MAPK9, SRPX, PPP1R15A, GLO1, UBE2B, RIPK3, STAT1, BIRC5, RHOA, MYC, PDCD4* |
| GO:0012501 | Programmed cell death | 463 | 1.747 | 1.11E-09 | 4.94E-08 | *IL1A, WT1, GSDMA, INHBA, CCL2, AGTR2, ATF3, ATF5, IGF1, S100A8, CXCL10, TNFAIP3, RNF122, IL10, CYCS, CD274, ARG2, HIGD1A, BMP2, PLA2G1B, FAS, THBS1, HSP90AA1, PHLDA2, HSPA9, IFNG, PLK3, HTR2B, TNFAIP8L3, SGK1, TCIM, DDIT3, TNFRSF11B, TNFRSF12A, PSMD10, DDX3X, DDIT4, CSRNP1, GHITM, NDNF, IRF1, NOD2, AIMP2, CASP3, EGR1, NR4A2, RIPK2, TLR4, CRYAB, HINT1, ZFP36, SIAH1, MAD2L1, RASSF2, AEN, CD40, GATA1, SOD1, BMPR2, SNAI2, LOC100522887, RHOB, RPS27L, MLKL, IL27RA, PIM1, BMI1, GADD45B, PDCD10, NME2, LGALS1, TNIP2, PSME3, HSPB1, BNIP1, TRAF2, TAF9, FTH1, DAD1, ATM, WNT16, LIG4, MED1, CEBPB, APIP, ST3GAL1, MCL1, PDCL3, EBAG9, UTP11, FZD3, ZPR1, PARK7, BBC3, COPS5, SELENOS, PRNP, HIPK3, FASTKD2,* |

| | | | | | | |
|---|---|---|---|---|---|---|
| | | | | | | *PPIA, KRAS, NPM1, ETV6, MAP3K7, ARF6, SMAD4, API5, FEM1B, IL18, PRDX3, BNIPL, NPPC, BCL2L12, KPNA1, ROCK1, SIAH2, USP53, AIFM1, GSKIP, NRP1, PRKAA2, MIF, ANGPTL4, PPARGC1A, ATG5, CCND2, XIAP, BCLAF1, MAPK8, VDAC2, FAM162A, NHERF1, TGFBR1, XRCC2, SGMS1, UBB, CASP8, VDAC1, TCHP, ITGA5, TIGAR, RAD21, SLC25A5, SP1, VDR, PEA15, ZFAND6, HYAL2, BAK1, GPX4, NTN1, CFLAR, MAPK9, SRPX, PPP1R15A, GLO1, UBE2B, RIPK3, STAT1, BIRC5, RHOA, MYC, PDCD4* |
| GO:0006954 | Inflammatory response | 217 | 1.736 | 4.73E-06 | 6.02E-05 | *IL6, AMCF-II, CCL19, ORM1, IL1A, IL1B, CXCL2, DUOXA2, GSDMA, CXCL8, CCL2, AGTR2, FOSL1, MMP8, PTGER3, S100A8, HP, CXCL10, IL1RAP, TNFAIP3, SAA3, IL10, PPBP, CXCL13, BMP2, S100A12, ACKR1, NFKBIA, THBS1, FABP4, IFNG, CCL8, FFAR2, DUSP10, PTGIR, TLR4, TICAM2, ZFP36, STAT3, KCNK6, IL33* |
| GO:0051248 | Negative regulation of protein metabolic process | 103 | 1.735 | 3.57E-04 | 1.90E-03 | *WT1, SERPINE1, EIF4EBP1, TNFAIP3, IL10, TIMP1, SESN2, THBS1, DDX3X, CRY1, PFDN6, ZFP36, MAD2L1, NCOA7, EIF2S1, AGO2, RIDA, PRKAR2A, TAF9, AZIN1, IGFBP5, PDCL3, DDRGK1, PARK7, STYX, PPIA, PAIP2B, ROCK1, CAPRIN1, GTPBP4, FYN, EIF4A3, PRKAA2, MIF, PFDN5, SERBP1, SRP9* |
| GO:0051603 | Proteolysis involved in protein catabolic process | 215 | 1.732 | 4.52E-06 | 5.96E-05 | *TP53INP2, TNFAIP3, RNF122, DNAJB9, DDIT3, RNF19B, PSMB3, PSMD10, UCHL1, HSP90B1, KLF1, NDFIP2, TNFAIP1, ZFAND2A, SIAH1, PSMA3, PSMA5, UBE2D3, TMEM126A, UBE2D1, PAQR3, UCHL3, USP44, RNF11, FBXO4, RNF5, PSMB6, PSME3, STT3B, TAF9, RNF180, SUMO2, RAD23A, KLHL2, PSMC5, UCHL5, YOD1, PDCL3, CUL5, NEMF, PSMA4, DDRGK1, UBXN4, PARK7, PSMA2, PSMD8, PSMA1, SOCS4, SELENOS, RNF115, STYX, SOCS5, FEM1B, MIB1, SIAH2, RAD23B, PSMC2, PSMB1, AKIRIN2, PSMC1, TMEM168, PSMB4, ELOC, CHMP2B, ANAPC10, LTN1, PSMD4, UBB, CASP8, CALR, VCP, IDE, DERL3, RNFT2, PSME2, PSMB7, MAPK9, UBE2B, WAC, SNF8, SPOP, WWP1, KLHDC3, ERLEC1, BCAP31, RNF14, ATE1, ARK2N, BAG2, RLIM, UBE3A, PSMD3, UBQLN4, PPP1R11, AREL1, UBE2G2, TOPORS, L3MBTL3, PJA2, ANAPC13, N4BP1, FBXO9, SPOPL* |

| | | | | | | |
|---|---|---|---|---|---|---|
| GO:0006511 | Ubiquitin-dependent protein catabolic process | 170 | 1.710 | 2.17E-05 | 1.98E-04 | *TP53INP2, TNFAIP3, RNF122, DDIT3, RNF19B, PSMB3, PSMD10, UCHL1, KLF1, NDFIP2, TNFAIP1, ZFAND2A, SIAH1, PSMA3, PSMA5, UBE2D3, UBE2D1, PAQR3, UCHL3, USP44, RNF11, FBXO4, RNF5, TAF9, RNF180, SUMO2, RAD23A, KLHL2, PSMC5, UCHL5, PDCL3, CUL5, NEMF, PSMA4, DDRGK1, PARK7, PSMA2, PSMD8, PSMA1, SOCS4, RNF115, STYX, SOCS5, FEM1B, MIB1, SIAH2, RAD23B, PSMC2, AKIRIN2, PSMC1, TMEM168, ELOC, CHMP2B, ANAPC10, LTN1, PSMD4, VCP, PSMB7, MAPK9, UBE2B, WAC, SNF8, SPOP, WWP1, KLHDC3, BCAP31, RNF14, ATE1, ARK2N, RLIM, UBE3A, PSMD3, UBQLN4, PPP1R11, AREL1, TOPORS, L3MBTL3, PJA2, ANAPC13, N4BP1, FBXO9, SPOPL* |
| GO:0019941 | Modification-dependent protein catabolic process | 173 | 1.710 | 1.46E-05 | 1.48E-04 | *TP53INP2, TNFAIP3, RNF122, DDIT3, RNF19B, PSMB3, PSMD10, UCHL1, KLF1, NDFIP2, TNFAIP1, ZFAND2A, SIAH1, PSMA3, PSMA5, UBE2D3, UBE2D1, PAQR3, UCHL3, USP44, RNF11, FBXO4, RNF5, TAF9, RNF180, SUMO2, RAD23A, KLHL2, PSMC5, UCHL5, PDCL3, CUL5, NEMF, PSMA4, DDRGK1, PARK7, PSMA2, PSMD8, PSMA1, SOCS4, RNF115, STYX, SOCS5, FEM1B, MIB1, SIAH2, RAD23B, PSMC2, AKIRIN2, PSMC1, TMEM168, ELOC, CHMP2B, ANAPC10, LTN1, PSMD4, UBB, VCP, PSMB7, MAPK9, UBE2B, WAC, SNF8, SPOP, WWP1, KLHDC3, BCAP31, RNF14, ATE1, ARK2N, RLIM, UBE3A, PSMD3, UBQLN4, PPP1R11, AREL1, TOPORS, L3MBTL3, PJA2, ANAPC13, N4BP1, FBXO9, SPOPL* |
| GO:0006915 | Apoptotic process | 450 | 1.707 | 1.47E-09 | 5.64E-08 | *IL1A, WT1, INHBA, CCL2, AGTR2, ATF3, ATF5, IGF1, S100A8, CXCL10, TNFAIP3, RNF122, IL10, CYCS, CD274, ARG2, HIGD1A, BMP2, PLA2G1B, FAS, THBS1, HSP90AA1, PHLDA2, HSPA9, IFNG, PLK3, HTR2B, TNFAIP8L3, SGK1, TCIM, DDIT3, TNFRSF11B, TNFRSF12A, PSMD10, DDX3X, DDIT4, CSRNP1, GHITM, NDNF, IRF1, NOD2, AIMP2, CASP3, EGR1, NR4A2, RIPK2, TLR4, CRYAB, HINT1, ZFP36, SIAH1, MAD2L1, RASSF2, AEN, CD40, GATA1, SOD1, BMPR2, SNAI2, LOC100522887, RHOB, RPS27L, IL27RA, PIM1, BMI1, GADD45B, PDCD10, NME2, LGALS1, TNIP2, PSME3, HSPB1, BNIP1, TRAF2, TAF9, DAD1, ATM, WNT16, LIG4, MED1, CEBPB, APIP, ST3GAL1, MCL1, PDCL3, EBAG9, UTP11, FZD3, ZPR1, PARK7, BBC3, COPS5, SELENOS, PRNP, HIPK3, FASTKD2, PPIA, KRAS, NPM1, ETV6, MAP3K7, ARF6, SMAD4, API5, FEM1B, IL18, PRDX3, BNIPL, NPPC, BCL2L12, KPNA1,* |

| | | | | | | |
|---|---|---|---|---|---|---|
| | | | | | | *ROCK1, SIAH2, USP53, AIFM1, GSKIP, NRP1, PRKAA2, MIF, ANGPTL4, PPARGC1A, ATG5, CCND2, XIAP, BCLAF1, MAPK8, VDAC2, FAM162A, NHERF1, TGFBR1, XRCC2, SGMS1, UBB, CASP8, VDAC1, TCHP, ITGA5, TIGAR, RAD21, SLC25A5, SP1, VDR, PEA15, ZFAND6, HYAL2, BAK1, NTN1, CFLAR, MAPK9, SRPX, PPP1R15A, GLO1, UBE2B, RIPK3, STAT1, BIRC5, RHOA, MYC, PDCD4* |
| GO:0051604 | Protein maturation | 146 | 1.699 | 1.48E-04 | 9.65E-04 | *HSPA6, DUOXA2, SERPINE1, MMP16, THBS1, DNAJB11, PLAU, HSP90AA1, HSPA9, CCT6A, HSPD1, PPID, DNAJB5, PFDN6, DNAJA2, CRYAB, DNAJB1, HSP90B1, CCT4, PTGES3, CCT2, GRPEL1, DHH, DNAJC2, HSPB1, IMMP1L, CDC37, DPM1, HSPA14, CCT3, PDCL3, PIGP, METAP2, PPIC, SPCS2, TOR1A, YIPF5, SDF2, PPIA, PIGK, NPPC, VBP1, PPIL3, DNAJB4, METAP1, FKBP2, PPIL1, SEC11C, PPWD1, PFDN5, PHEX, SPCS3, CASP8, CALR* |
| GO:0043632 | Modification-dependent macromolecule catabolic process | 174 | 1.689 | 1.99E-05 | 1.86E-04 | *TP53INP2, TNFAIP3, RNF122, DDIT3, RNF19B, PSMB3, PSMD10, UCHL1, KLF1, NDFIP2, TNFAIP1, ZFAND2A, SIAH1, PSMA3, PSMA5, UBE2D3, UBE2D1, PAQR3, UCHL3, USP44, RNF11, FBXO4, RNF5, TAF9, RNF180, SUMO2, RAD23A, KLHL2, PSMC5, UCHL5, PDCL3, CUL5, NEMF, PSMA4, DDRGK1, PARK7, PSMA2, PSMD8, PSMA1, SOCS4, RNF115, STYX, SOCS5, FEM1B, MIB1, SIAH2, RAD23B, PSMC2, AKIRIN2, PSMC1, TMEM168, ELOC, CHMP2B, ANAPC10, LTN1, PSMD4, UBB, VCP, PSMB7, MAPK9, UBE2B, WAC, SNF8, SPOP, WWP1, KLHDC3, BCAP31, RNF14, ATE1, ARK2N, RLIM, UBE3A, PSMD3, UBQLN4, PPP1R11, AREL1, TOPORS, L3MBTL3, PJA2, ANAPC13, N4BP1, FBXO9, SPOPL* |
| GO:0006091 | Generation of precursor metabolites and energy | 155 | 1.669 | 1.77E-04 | 1.09E-03 | *TYRP1, IGF1, CYCS, MTFR1, PHLDA2, COQ10B, NDUFC2, ND4L, DDIT4, UCHL1, NDUFB3, MDH1, PGM2, C7H6orf125, PTGES3, NDUFB9, COX5B, PGK1, IDH3A, COA6, ATP5MF, COX7A2, SLC4A1, PPP1CC, HK2, NDUFA5, PPP2CA, UQCR11, SLC25A33, COX7B, ATP5ME, COX5A, NDUFB6, PDHA1, PARK7, TPI1, COQ10A, NDUFV2, ATP5PO, UQCR10, PGAM1, UQCRQ, PPP1CB, COX7C, UGP2, NDUFS1, PPARGC1A, CS, NDUFS8, IDE, TIGAR, ATP5PB, LOC100524873, SDHAF2, CHCHD2, RHOA, IDH3B, SDHD, CYC1, NDUFB8, APP, FDX1, NDUFV1, PHKG2, ETFB, ENO3* |

| | | | | | | |
|---|---|---|---|---|---|---|
| GO:0042981 | Regulation of apoptotic process | 344 | 1.667 | 7.87E-07 | 1.22E-05 | *WT1, INHBA, CCL2, AGTR2, ATF3, ATF5, IGF1, S100A8, CXCL10, TNFAIP3, RNF122, IL10, CD274, ARG2, HIGD1A, BMP2, PLA2G1B, FAS, THBS1, HSP90AA1, PHLDA2, HSPA9, IFNG, PLK3, HTR2B, TNFAIP8L3, TCIM, DDIT3, TNFRSF12A, PSMD10, DDX3X, GHITM, NDNF, NOD2, CASP3, EGR1, NR4A2, RIPK2, TLR4, CRYAB, ZFP36, SIAH1, MAD2L1, RASSF2, CD40, GATA1, SOD1, SNAI2, LOC100522887, RHOB, IL27RA, PIM1, BMI1, GADD45B, PDCD10, NME2, TNIP2, PSME3, HSPB1, TRAF2, TAF9, DAD1, WNT16, LIG4, MED1, CEBPB, ST3GAL1, MCL1, UTP11, FZD3, ZPR1, PARK7, BBC3, COPS5, SELENOS, PRNP, HIPK3, PPIA, KRAS, NPM1, API5, FEM1B, PRDX3, NPPC, BCL2L12, KPNA1, ROCK1, SIAH2, NRP1, PRKAA2, MIF, ANGPTL4, PPARGC1A, CCND2, BCLAF1, MAPK8, FAM162A, NHERF1, XRCC2, UBB, CASP8, ITGA5, TIGAR, SLC25A5, SP1, VDR, PEA15, ZFAND6, HYAL2, BAK1, CFLAR, MAPK9, SRPX, GLO1, UBE2B, RIPK3, STAT1, BIRC5, RHOA, MYC, PDCD4* |
| GO:0007005 | Mitochondrion organization | 134 | 1.659 | 2.23E-04 | 1.34E-03 | *IGF1, NDUFAF4, HIGD1A, TIMM8A, MTFR1, SESN2, NDUFB1, NDUFAF5, NDUFC2, PSMD10, GHITM, MICOS13, C7H6orf125, COX16, COX17, PET100, AGK, TMEM126A, MTX2, NDUFB5, APOOL, NDUFA5, NDUFA11, MTX1, APOO, SLC25A33, MCL1, PARK7, BBC3, DDHD1, MIEF1, NDUFB7, PRDX3, NDUFS5, ARMCX3, PPARGC1A, BCS1L, TIMM21, ROMO1, VDAC2, FAM162A, NDUFB4, NDUFS8* |
| GO:0051246 | Regulation of protein metabolic process | 316 | 1.659 | 1.41E-06 | 1.93E-05 | *WT1, INHBA, SERPINE1, EIF4EBP1, LIF, TNFAIP3, IL10, TIMP1, ADRA2A, PLA2G1B, SESN2, NFKBIA, THBS1, LIPG, IFNG, PLK3, PSMD10, DDX3X, GNL3, CRY1, SOAT1, PFDN6, CASP3, EGR1, C7H6orf125, NDFIP2, ZFAND2A, ZFP36, EIF1, MAD2L1, NCOA7, EIF2S1, BZW1, GATA1, HDAC2, NIBAN1, PPP1R15B, PUM3, UBE2D1, PAQR3, PTPN11, AGO2, RIDA, FBXO4, DCUN1D5, PSME3, SSB, HSPB1, ETF1, PRKAR2A, TAF9, RNF180, SUMO2, KAT2A, AZIN1, IL15, IGFBP5, PDCL3, PAIP1, DDRGK1, PARK7, SOCS4, PRNP, STYX, FASTKD2, PPIA, NPM1, SOCS5, DR1, PAIP2B, PSMD2, ROCK1, CAPRIN1, ABCF1, GTPBP4, PHF23, FYN, EIF4A3, NRP1, PRKAA2, ARHGEF2, MIF, NCBP2, TMEM168, PFDN5, NRDC, SERBP1, TMED10, SRP9, MAPK8, PSMD1, FBXO33, SPRED1, UBB, CASP8,* |

| | | | | | | |
|---|---|---|---|---|---|---|
| | | | | | | *CALR, ITGA5, RNFT2, PSME2, NAT10, METTL3, FXR1, SP1, MBIP, MAPK9, PPP1R15A, UBE2B, WAC, RGS2, SNF8, SPOP, RAMP1, APOE, PLGRKT, APP, ELAVL1, RIC1, BCAP31, EIF5A, PIN1, JAK2, BAG2, TICAM1, UBE3A, PSMD3, DCUN1D4, PFDN1, QKI, UBQLN4, LSM14A, BCL10, IGF2BP1, USP16* |
| GO:0043933 | Protein-containing complex organization | 432 | 1.654 | 1.64E-07 | 3.25E-06 | *FGB, TIFA, SAMHD1, AQP5, KIF18A, NDUFAF4, SNRPF, MMP1, HIGD1A, SNRPD1, LIPG, IFNG, HSPA8, NDUFB1, PRND, NDUFAF5, NDUFC2, EIF3J, PSMD10, DDX3X, TWF1, RRS1, DDIT4, BAZ1A, MID1IP1, SOAT1, SNRPG, PFDN6, NOPCHAP1, HMBOX1, UGDH, ARPC5L, TLR4, C7H6orf125, COX16, TNFAIP1, KCNJ2, SLC1A5, KCNK6, FARSA, PTGES3, SLC31A1, COX17, MRTO4, SNRPE, PSMG2, VIL1, PET100, CCT2, MLKL, TMEM126A, CLNS1A, GRWD1, GEMIN6, PTPN11, SET, CDC42EP3, CAND1, PREB, TUBB1, COA6, EIF3A, STRAP, ME1, EIF3I, NOP2, SNRPD3, CAPZA2, DENR, ETF1, PICALM, TRAF2, TAF9, NDUFB5, ABT1, RAP1B, ZBTB1, NDUFA5, NDUFA11, DRG1, PPP2CA, SLC25A33, TMEM39A, NEMF, SNRPC, PARK7, BBC3, NUFIP1, G3BP2, PRNP, FASTKD2, ABITRAM, LOC100738684, NPM1, PYM1, EIF3M, CLP1, GTF2A2, ARF6, SAR1B, NDUFB7, NDUFS5, SRSF1, PLTP, PHF23, POLE3, CAPZA1, SHQ1, USP39, NSF, ARHGEF2, OTUD6B, MIF, CENPP, GEMIN2, SUGT1, BCS1L, AVIL, NAF1, SF3A3, CHMP2B, TIMM21, TIRAP, MAPK8, DSTN, ASF1A, NDUFB4, CDC42EP2, NDUFS8, PSMG4, MRPS7, CALR, ARPC3, MTRF1, SAR1A, VCP, HJURP, CDC123, COPS8, PDCL, CENPC, PARD6B, MAPK9, MRPL58, UBLCP1, SHPRH, LSM2, SNAP23, RHOA, HSD17B10, APOE, NDUFB8, TP53, SLAIN2, ELAVL1, TMEM186, KCNA2, EIF5A, SUPT16H, SMYD3, TERF1, HIRA, TICAM1, BECN1, SNUPN, RASA1, UBQLN4, SPAST, BCL10, SRP19, USP16, CFL1, SCAF11* |
| GO:0007423 | Sensory organ development | 105 | 1.650 | 9.94E-04 | 4.60E-03 | *WT1, CRB2, INHBA, GJA1, SOX11, DCLK1, AQP5, BMP2, C5H12orf57, PROX1, IFT20, RCN1, ACTL6A, DRAM2, STAT3, CEBPD, GNAT1, NECTIN1, SOD1, BMPR2, TSKU, PTPN11* |
| GO:0043067 | Regulation of programmed cell death | 353 | 1.648 | 7.10E-07 | 1.15E-05 | *WT1, INHBA, CCL2, AGTR2, ATF3, ATF5, IGF1, S100A8, CXCL10, TNFAIP3, RNF122, IL10, CD274, ARG2, HIGD1A, BMP2, PLA2G1B, FAS, THBS1, HSP90AA1, PHLDA2, HSPA9, IFNG, PLK3, HTR2B, TNFAIP8L3, TCIM, DDIT3, TNFRSF12A, PSMD10, DDX3X, GHITM, NDNF,* |

| | | | | | | |
|---|---|---|---|---|---|---|
| | | | | | | *NOD2, CASP3, EGR1, NR4A2, RIPK2, TLR4, CRYAB, ZFP36, SIAH1, MAD2L1, RASSF2, CD40, GATA1, SOD1, SNAI2, LOC100522887, RHOB, IL27RA, PIM1, BMI1, GADD45B, PDCD10, NME2, TNIP2, PSME3, HSPB1, TRAF2, TAF9, FTH1, DAD1, WNT16, LIG4, MED1, CEBPB, ST3GAL1, MCL1, UTP11, FZD3, ZPR1, PARK7, BBC3, COPS5, SELENOS, PRNP, HIPK3, PPIA, KRAS, NPM1, API5, FEM1B, PRDX3, NPPC, BCL2L12, KPNA1, ROCK1, SIAH2, NRP1, PRKAA2, MIF, ANGPTL4, PPARGC1A, CCND2, XIAP, BCLAF1, MAPK8, FAM162A, NHERF1, XRCC2, UBB, CASP8, ITGA5, TIGAR, SLC25A5, SP1, VDR, PEA15, ZFAND6, HYAL2, BAK1, GPX4, CFLAR, MAPK9, SRPX, GLO1, UBE2B, RIPK3, STAT1, BIRC5, RHOA, MYC, PDCD4* |
| GO:0009057 | Macromolecule catabolic process | 375 | 1.640 | 1.17E-06 | 1.66E-05 | *TP53INP2, TNFAIP3, RNF122, IL10, TIMP1, ADRA2A, DNAJB9, IFNG, PLK3, DDIT3, RNF19B, PSMB3, PSMD10, RBM8A, UCHL1, DCPS, PGM2, HSP90B1, KLF1, NDFIP2, TNFAIP1, ZFAND2A, ZFP36, SIAH1, MAD2L1, REXO4, PSMA3, EXOSC3, PSMA5, UBE2D3, TMEM126A, UBE2D1, PAQR3, PHAX, UCHL3, AGO2, RIDA, USP44, RNF11, FBXO4, RNF5, PSMB6, PSME3, SSB, ETF1, STT3B, TRAF2, TAF9, RNF180, SUMO2, SKIC8, ATM, RAD23A, KLHL2, DDX49, AZIN1, PSMC5, UCHL5, CNOT7, SMG8, YOD1, PDCL3, PDE12, CUL5, PAIP1, NEMF, ZPR1, PSMA4, DDRGK1, UBXN4, PARK7, PSMA2, PSMD8, PSMA1, SOCS4, SELENOS, RNF115, STYX, FASTKD2, PELO, NPM1, SOCS5, PNRC2, EXOSC4, FEM1B, LSM5, MIB1, PSMD2, ROCK1, CAPRIN1, SIAH2, FYN, EIF4A3, RAD23B, PSMC2, PSMB1, PNPT1, AKIRIN2, PSMC1, NCBP2, TMEM168, PSMB4, NRDC, NAF1, ELOC, CHMP2B, TIRAP, ANAPC10, PABPC4, PSMD1, LTN1, MTREX, PSMD4, UBB, CASP8, CALR, DXO, VCP, IDE, DERL3, RNFT2, PSME2, HYAL2, PSMB7, MAPK9, LSM2, UBE2B, WAC, SNF8, SPOP, WWP1, KLHDC3, ERLEC1, ELAVL1, RIC1, BCAP31, ARID5A, RNF14, PIN1, ATE1, ARK2N, BAG2, PHKG2, RLIM, BECN1, UBE3A, PSMD3, QKI, UBQLN4, IGF2BP1* |
| GO:0065003 | Protein-containing complex assembly | 381 | 1.635 | 6.93E-07 | 1.15E-05 | *FGB, TIFA, SAMHD1, AQP5, NDUFAF4, SNRPF, MMP1, HIGD1A, SNRPD1, IFNG, NDUFB1, PRND, NDUFAF5, NDUFC2, EIF3J, PSMD10, DDX3X, TWF1, RRS1, BAZ1A, MID1IP1, SOAT1, SNRPG, PFDN6, NOPCHAP1, HMBOX1, UGDH, ARPC5L, TLR4,* |

| | | | | | | |
|---|---|---|---|---|---|---|
| | | | | | | *C7H6orf125, COX16, TNFAIP1, KCNJ2, SLC1A5, KCNK6, FARSA, PTGES3, SLC31A1, COX17, MRTO4, SNRPE, PSMG2, VIL1, PET100, CCT2, MLKL, TMEM126A, CLNS1A, GRWD1, GEMIN6, PTPN11, SET, CDC42EP3, CAND1, PREB, TUBB1, COA6, EIF3A, STRAP, ME1, EIF3I, NOP2, SNRPD3, CAPZA2, DENR, PICALM, TRAF2, TAF9, NDUFB5, ABT1, RAP1B, ZBTB1, NDUFA5, NDUFA11, DRG1, PPP2CA, SLC25A33, NEMF, SNRPC, PARK7, BBC3, NUFIP1, G3BP2, PRNP, FASTKD2, ABITRAM, LOC100738684, NPM1, EIF3M, CLP1, GTF2A2, ARF6, SAR1B, NDUFB7, NDUFS5, SRSF1, PLTP, POLE3, CAPZA1, SHQ1, USP39, OTUD6B, MIF, CENPP, GEMIN2, SUGT1, BCS1L, AVIL, NAF1, SF3A3, TIMM21, TIRAP, MAPK8, ASF1A, NDUFB4, CDC42EP2, NDUFS8, PSMG4, MRPS7, CALR, ARPC3, SAR1A, HJURP, CDC123, COPS8, PDCL, CENPC, PARD6B, MAPK9, UBLCP1, SHPRH, LSM2, SNAP23, RHOA, HSD17B10, APOE, NDUFB8, TP53, SLAIN2, ELAVL1, TMEM186, KCNA2* |
| GO:0010498 | Proteasomal protein catabolic process | 131 | 1.616 | 4.70E-04 | 2.43E-03 | *RNF122, DNAJB9, DDIT3, PSMB3, PSMD10, HSP90B1, TNFAIP1, ZFAND2A, SIAH1, PSMA5, UBE2D1, PAQR3, USP44, FBXO4, RNF5, PSME3, STT3B, TAF9, RNF180, SUMO2, RAD23A, KLHL2, PSMC5, YOD1, NEMF, DDRGK1, UBXN4, PARK7, PSMD8, SOCS4, SELENOS, STYX, SOCS5, FEM1B, SIAH2, RAD23B, AKIRIN2, PSMC1, ANAPC10, LTN1, PSMD4, UBB, CALR, VCP, DERL3, RNFT2, PSME2, PSMB7, MAPK9, UBE2B, WAC, SPOP, WWP1, KLHDC3, ERLEC1, BCAP31, ATE1, BAG2* |
| GO:0010608 | Post-transcriptional regulation of gene expression | 137 | 1.611 | 2.89E-04 | 1.62E-03 | *WT1, EIF4EBP1, SESN2, THBS1, DDX3X, DCPS, C7H6orf125, ZFP36, EIF1, EIF2S1, BZW1, NIBAN1, PUM3, PHAX, AGO2, RIDA, FBXO4, SSB, ETF1, TRAF2, DDX49, CDC37, IGFBP5, CNOT7, PDE12, PAIP1, FASTKD2, NPM1, CLP1, PAIP2B, ROCK1, CAPRIN1, ABCF1, EIF4A3, NCBP2, SERBP1, NAF1, TIRAP, SRP9, PABPC4, MATR3, CALR, NAT10, METTL3, FXR1, PPP1R15A, RGS2, APP, ELAVL1, EIF5A, ARID5A, QKI, LSM14A, IGF2BP1, USP16* |
| GO:0009617 | Response to bacterium | 128 | 1.596 | 1.41E-03 | 6.21E-03 | *AMCF-II, IL1A, IL1B, GSDMA, CXCL8, MMP8, S100A8, HP, CXCL10, TNFAIP3, IL10, PPBP, ARG2, CXCL13, BMP2, NFKBIA, FABP4, LBP, DUSP10, PTGIR, TLR4, IFNE, ZFP36, IFNAR1, RAB1A, NFKBIB, PTPN11, IL27RA, RNF5, TNIP2, BPIFA1, CEBPB, RAB14,* |

| | | | | | | |
|---|---|---|---|---|---|---|
| | | | | | | *SELENOS, LEAP2, IRF8, MAP3K7, LSM5, IL18, PRDX3, THRSP, AKIRIN2, CDK4, MIF, TIRAP, ADAM9, MAPK8, ROMO1* |
| GO:0045786 | Negative regulation of cell cycle | 111 | 1.588 | 2.03E-03 | 8.71E-03 | *INHBA, CCL2, ATF5, LIF, BMP2, BTG3, PROX1, PLK3, TIPIN, CRY1, MAD2L1, NLE1, RHOB, RPS27L, PTPN11, TIPRL, USP44, FBXO4, SPDL1, FBXO5, ATM, KAT2A, FZD3, NAA10, WEE1, NPM1, FEM1B, NUBP1, NPPC, CDK2AP2* |
| GO:0008283 | Cell population proliferation | 444 | 1.582 | 5.93E-07 | 1.06E-05 | *IL6, IL1A, IL1B, NR4A3, WT1, APLN, INHBA, CCL2, CSF3, GJA1, SOX11, ATF5, NME1, IGF1, LIF, FAM83B, CXCL10, EIF2S2, TNFAIP3, IL10, TIMP1, CD274, ARG2, INHA, BMP2, PLA2G1B, MMP16, TFRC, NFKBIA, SLC7A1, THBS1, JUNB, FABP4, PHLDA2, PROX1, IER5, IFNG, HTR2B, IFT20, CTPS1, DDIT3, UCHL1, IRF1, TIPIN, CXCL11, DUSP10, DRAM2, ENPEP, FAM98A, PTGIR, TNFSF13B, TLR4, TGFB3, CKS2, ESM1, ZFP36, FAM98B, STAT3, GNAT1, PTGES3, CD40, GATA1, COX17, BMPR2, HDAC2, SNAI2, HILPDA, PIM1, FUT2, BMI1, EGF, FBXO4, PDCD10, CLEC11A, NOP2, CCN3, PRDM1, STRN, RAP1B, KAT2A, WNT16, IL15, LIG4, MED1, IGFBP5, CEBPB, SDR16C5, CNOT7, SLC25A33, PDCL3, FZD3, ZPR1, RPRD1B, DDRGK1, MORF4L1, HAPLN1, CALCRL, KRAS, NPM1, ETV6, IGFBP2* |
| GO:0045184 | Establishment of protein localization | 452 | 1.572 | 9.29E-07 | 1.38E-05 | *IL6, IL1A, DUOXA2, NEURL3, KPNA2, TNFAIP3, ADRA2A, ARL4D, TIMM8A, TFRC, NFKBIA, EGR2, HSPA9, ARL5B, IFNG, PLK3, LOC100157391, IFT20, TMED5, EMC9, DDIT3, RAB39B, KCNJ11, FFAR2, TLR4, C7H6orf125, TGFB3, SEC23B, NDFIP2, ZFAND2A, EMC4, STAT3, ENY2, SEC22B, SEC24A, TIMM8B, ARCN1, HIKESHI, ARF4, ARF2, AGK, RHOB, NXT2, TMEM126A, PHAX, RAB1A, COPB2, SEC61B, CHCHD4, SYS1, PTPN11, MMGT1, COPB1, DNAJC15, SRP14, PREB, YWHAG, PDCD10, BBS5, RAB13, HSPB1, NXT1, MTX2, CCN3, IMMP1L, TIMM10B, SRP54, TOMM40, SEC23A, TIMM13, SEC62, MED1, STX11, MTX1, INSIG1, EMC3, RAB14, SRPRA, VTI1B, SSR3, ZPR1, USO1, PARK7, BBC3, AKAP5, YIPF5, SELENOS, CALCRL, TNPO1, CLTC, RAB5A, KPNB1, STX7, AP1S2, ATG3, ARF6, DNAJC19, KTN1, COPE, SAR1B, KPNA1, BET1, FYN, AKIRIN2, TXNIP, NSF, STAM2, SEC63, VPS26A, GET3, TOMM20, PRKAR1A, SNX19, ZDHHC9, BCS1L, AP3M1, CHMP2B, TIMM21, GET1, TMED10, ADAM9, SRP9, BLOC1S6, ROMO1, SNX5, XPO5, RAB2A, CALR,* |

| | | | | | | |
|---|---|---|---|---|---|---|
| | | | | | | *TMED2, LIN7C, SAR1A, VCP, TOMM22, NUTF2, DERL3, YIF1A, TVP23B, STX2, RAB10, ARL3, ZFAND6, HYAL2, C2CD2L, IPO4, RAB34, TMED9, TRAM1, PAFAH1B1, GDI2, BIRC5, SNF8, RAMP1, YIF1B, RAB3D, RAB27A, BLZF1, ELAVL1, VPS29, RIC1, BCAP31, SDAD1, IPO8, SMYD3, RAB9A, AP3S2, TERF1, TRIM23, SFT2D1, MAPK14, SNUPN, SEC61A2* |
| GO:0071705 | Nitrogen compound transport | 495 | 1.568 | 2.44E-07 | 4.58E-06 | *IL6, IL1A, DUOXA2, GJA1, SLC38A5, S100A8, NEURL3, KPNA2, ADRA2A, ARL4D, TIMM8A, TFRC, NFKBIA, SLC7A1, AQP3, EGR2, HSPA9, ARL5B, IFNG, PLK3, SLC38A2, LOC100157391, EIF4E, SLC18B1, IFT20, TMED5, RAB39B, RBM8A, KCNJ11, SLC28A2, FFAR2, GPM6B, TLR4, C7H6orf125, TGFB3, SEC23B, NDFIP2, ZFP36, STAT3, SLC1A5, ENY2, SEC22B, GNAT1, SEC24A, TIMM8B, ARCN1, THOC3, HIKESHI, ARF4, ARF2, AGK, RHOB, NXT2, PDZK1, PHAX, RAB1A, COPB2, SEC61B, CHCHD4, SYS1, PTPN11, COPB1, DNAJC15, PREB, PDCD10, BBS5, RAB13, SLC12A2, SSB, HSPB1, NXT1, MAGOH, MTX2, CCN3, TIMM10B, SRP54, TOMM40, SEC23A, KCNK1, TIMM13, MAGOHB, SEC62, AZIN1, MED1, STX11, MTX1, SLC15A3, INSIG1, SLC25A33, RAB14, SLC33A1, SRPRA, VTI1B, ZPR1, USO1, PARK7, SHOC2, AKAP5, YIPF5, SELENOS, CALCRL, TNPO1, CLTC, RAB5A, KPNB1, STX7, AP1S2, ATG3, ARF6, DNAJC19, KTN1, COPE, RHCE, SAR1B, KPNA1, BET1, EIF4A3, AKIRIN2, TXNIP, NSF, STAM2, TAP1, SEC63, VPS26A, TOMM20, PRKAR1A, SNX19, NCBP2, AP3M1, CHMP2B, TIMM21, TMED10, ADAM9, BLOC1S6, ROMO1, SNX5, XPO5, RAB2A, NHERF1, THOC2, NOL6, CALR, SLC25A20, TMED2, LIN7C, SAR1A, VCP, TOMM22, NUTF2, DERL3, YIF1A, TVP23B, SLC25A5, STX2, RAB10, ARL3, MRPL18, HYAL2, C2CD2L, IPO4, RAB34, TMED9, TRAM1, PAFAH1B1, GDI2, SNAP23, BIRC5, SNF8, RAMP1, YIF1B, SLC22A4, RAB3D, RAB27A, BLZF1, TP53, ELAVL1, VPS29, RIC1, BCAP31, SDAD1, KCNA2, IPO8* |
| GO:0008285 | Negative regulation of cell population proliferation | 156 | 1.568 | 6.32E-04 | 3.13E-03 | *IL1A, WT1, APLN, INHBA, CCL2, SOX11, ATF5, LIF, TNFAIP3, IL10, CD274, ARG2, BMP2, THBS1, PROX1, IFNG, DUSP10, PTGIR, TGFB3, GATA1, BMPR2, SNAI2, FBXO4, PDCD10, CCN3, STRN, MED1, IGFBP5, CEBPB, CNOT7* |

| | | | | | | |
|---|---|---|---|---|---|---|
| GO:0051050 | Positive regulation of transport | 149 | 1.564 | 5.78E-04 | 2.90E-03 | *IL6, IL1A, IL1B, NR4A3, APLN, CCL2, SOX11, IGF1, S100A8, CXCL10, ADRA2A, BMP2, PLA2G1B, NFKBIA, LIPG, IFNG, PLK3, LYAR, ATP1B1, EIPR1, FFAR2, CXCL11, TLR4, TGFB3* |
| GO:0008284 | Positive regulation of cell population proliferation | 215 | 1.561 | 1.14E-04 | 7.96E-04 | *IL6, IL1A, IL1B, NR4A3, APLN, CSF3, GJA1, SOX11, NME1, IGF1, LIF, TNFAIP3, IL10, TIMP1, BMP2, PLA2G1B, TFRC, SLC7A1, THBS1, FABP4, PROX1, HTR2B, TIPIN, FAM98A, TNFSF13B, TLR4, TGFB3, ESM1, FAM98B, CD40, GATA1, COX17, HDAC2, HILPDA, PIM1, BMI1, EGF, PDCD10, CLEC11A, NOP2, IL15, LIG4, MED1, IGFBP5, CNOT7, SLC25A33, PDCL3, FZD3, RPRD1B, DDRGK1, HAPLN1, CALCRL, KRAS, NPM1, IGFBP2* |
| GO:0033993 | Response to lipid | 156 | 1.560 | 7.74E-04 | 3.73E-03 | *AMCF-II, IL1A, IL1B, CXCL8, MMP8, S100A8, CXCL10, TNFAIP3, IL10, PPBP, CXCL13, NFKBIA, GOT1, AQP3, JUNB, DDIT4, CRY1, FFAR2, DUSP10, CFTR, PTGIR, TLR4, TFPI, TGFB3, ZFP36, PTGES3, IFNAR1, SNAI2, NFKBIB, PTPN11, AARD, PIM1* |
| GO:0046907 | Intracellular transport | 362 | 1.560 | 1.39E-05 | 1.45E-04 | *GOLT1A, KPNA2, ARL4D, TIMM8A, NFKBIA, EGR2, HSPA9, ARL5B, IFNG, PLK3, IFT20, GOLT1B, SLC30A4, UCHL1, EIPR1, RANBP1, NOP9, SEC23B, NDFIP2, STAT3, ENY2, SEC22B, SEC24A, THOC3, HIKESHI, AGK, RHOB, NXT2, PHAX, RAB1A, COPB2, SEC61B, PTPN11, EIF6, COPB1, PREB, PDCD10, RAB13, SSB, HSPB1, NXT1, SRP54, SEC23A, LMNB1, ERGIC2, MED1, STX11, INSIG1, RAB14, SRPRA, NEMF, VTI1B, ZPR1, USO1, PARK7, DPY30, AKAP5, YIPF5, NPC1, SELENOS, NUS1, TNPO1, NPM1, ARL1, CLTC, RAB5A, KPNB1, STX7, AP1S2, ARF6, COPE, SAR1B, KPNA1, ARMCX3, AKIRIN2, TXNIP, SEC63, VPS26A, TOMM20, RHOBTB3, NCBP2, AP3M1, CHMP2B, MAP2K1, TMED10, BLOC1S6, ROMO1, SNX5, XPO5, NHERF1, THOC2, NOL6, UBB, LDLR, CALR, SAR1A, VCP, TOMM22, NUTF2, DERL3, YIF1A, STX2, ASB15, HYAL2, IPO4, TMED9, TRAM1, PAFAH1B1, SNF8, RAMP1, IFT74, YIF1B, RAB3D, APP, ELAVL1, VPS29, RIC1, BCAP31, SDAD1, IPO8, RAB9A, AP3S2, TRIM23, TMEM41B, FYTTD1, MAPK14, BECN1, SNUPN* |
| GO:0042127 | Regulation of cell population proliferation | 374 | 1.559 | 1.15E-05 | 1.24E-04 | *IL6, IL1A, IL1B, NR4A3, WT1, APLN, INHBA, CCL2, CSF3, GJA1, SOX11, ATF5, NME1, IGF1, LIF, CXCL10, TNFAIP3, IL10, TIMP1, CD274, ARG2, INHA, BMP2, PLA2G1B, TFRC, NFKBIA, SLC7A1, THBS1, JUNB, FABP4, PHLDA2, PROX1, IER5, IFNG, HTR2B, IRF1,* |

| | | | | | | |
|---|---|---|---|---|---|---|
| | | | | | | *TIPIN, CXCL11, DUSP10, FAM98A, PTGIR, TNFSF13B, TLR4, TGFB3, ESM1, ZFP36, FAM98B, STAT3, CD40, GATA1, COX17, BMPR2, HDAC2, SNAI2, HILPDA, PIM1, FUT2, BMI1, EGF, FBXO4, PDCD10, CLEC11A, NOP2, CCN3, PRDM1, STRN, IL15, LIG4, MED1, IGFBP5, CEBPB, CNOT7, SLC25A33, PDCL3, FZD3, RPRD1B, DDRGK1, HAPLN1, CALCRL* |
| GO:0015031 | Protein transport | 395 | 1.550 | 8.02E-06 | 9.51E-05 | *IL6, IL1A, DUOXA2, NEURL3, KPNA2, ADRA2A, ARL4D, TIMM8A, TFRC, NFKBIA, EGR2, HSPA9, ARL5B, IFNG, PLK3, LOC100157391, IFT20, TMED5, RAB39B, KCNJ11, FFAR2, TLR4, C7H6orf125, TGFB3, SEC23B, NDFIP2, STAT3, ENY2, SEC22B, SEC24A, TIMM8B, ARCN1, HIKESHI, ARF4, ARF2, AGK, RHOB, NXT2, PHAX, RAB1A, COPB2, SEC61B, CHCHD4, SYS1, PTPN11, COPB1, DNAJC15, PREB, PDCD10, BBS5, RAB13, HSPB1, NXT1, MTX2, CCN3, TIMM10B, SRP54, TOMM40, SEC23A, TIMM13, SEC62, MED1, STX11, MTX1, INSIG1, RAB14, SRPRA, VTI1B, ZPR1, USO1, PARK7, AKAP5, YIPF5, SELENOS, CALCRL, TNPO1, CLTC, RAB5A, KPNB1, STX7, AP1S2, ATG3, ARF6, DNAJC19, KTN1, COPE, SAR1B, KPNA1, BET1, AKIRIN2, TXNIP, NSF, STAM2, SEC63, VPS26A, TOMM20, PRKAR1A, SNX19, AP3M1, CHMP2B, TIMM21, TMED10, ADAM9, BLOC1S6, ROMO1, SNX5, XPO5, RAB2A, CALR, TMED2, LIN7C, SAR1A, VCP, TOMM22, NUTF2, DERL3, YIF1A, TVP23B, STX2, RAB10, ARL3, HYAL2, C2CD2L, IPO4, RAB34, TMED9, TRAM1, PAFAH1B1, GDI2, BIRC5, SNF8, RAMP1, YIF1B, RAB3D, RAB27A, BLZF1, ELAVL1, VPS29, RIC1, BCAP31, SDAD1, IPO8* |
| GO:0009790 | Embryo development | 240 | 1.549 | 1.18E-04 | 8.08E-04 | *PSPH, WT1, CXCL8, APLN, CRB2, INHBA, GJA1, SOX11, FOSL1, MMP8, LIF, EIF2S2, IL10, BMP2, MMP16, RSPO3, JUNB, DUSP5, B4GALT5, PHLDA2, PROX1, HTR2B, RCN1, ACTL6A, UGDH, NLE1, NECTIN1, GATA1, SOD1, BMPR2, LEO1, SF3B6, WDR74, TBX18, BMI1, ACSL4, GNA13, PRDM1, DAD1, KAT2A, LIG4, MED1, CEBPB, INSIG1, RAB14, MED21, PLPP4, FZD3, ZPR1, LUZP1* |
| GO:0006886 | Intracellular protein transport | 189 | 1.536 | 1.07E-03 | 4.83E-03 | *KPNA2, ARL4D, TIMM8A, NFKBIA, EGR2, HSPA9, ARL5B, IFNG, PLK3, SEC23B, STAT3, SEC24A, HIKESHI, AGK, RHOB, RAB1A, COPB2, SEC61B, PTPN11, COPB1, PDCD10, HSPB1, NXT1, SRP54, SEC23A, MED1, STX11, INSIG1, RAB14, SRPRA, VTI1B, ZPR1, USO1,* |

| | | | | | | |
|---|---|---|---|---|---|---|
| | | | | | | *PARK7, AKAP5, SELENOS, TNPO1, CLTC, RAB5A, KPNB1, STX7, AP1S2, ARF6, SAR1B, KPNA1, AKIRIN2, TXNIP, SEC63, VPS26A, TOMM20, AP3M1, TMED10, ROMO1, SNX5, XPO5, CALR, SAR1A, VCP, TOMM22, DERL3, STX2, HYAL2, IPO4, TMED9, TRAM1, SNF8, RAMP1, RAB3D, ELAVL1, RIC1, BCAP31, IPO8, AP3S2, TRIM23, MAPK14, SNUPN* |
| GO:0009628 | Response to abiotic stimulus | 208 | 1.528 | 4.10E-04 | 2.15E-03 | *IL1A, GJA1, AQP5, SLC8A3, LIF, MMP1, TIMP1, BMP2, NFKBIA, AQP3, HSP90AA1, IER5, PLK3, HSPD1, EFHD1, IFT20, TCIM, DDX3X, DDIT4, NDNF, IRF1, TIPIN, CRY1, EGR1, NR4A2, DNAJA2, TLR4, CRYAB, DNAJB1, TGFB3, SUV39H2, TNFSF14, GNAT1, AEN, EIF2S1, CD40, SOD1, HIKESHI, RHOB, GPR4* |
| GO:0044283 | Small molecule biosynthetic process | 171 | 1.527 | 1.07E-03 | 4.83E-03 | *PSPH, PHGDH, ASNS, INHBA, PTGIS, ATF3, PCK2, ADK, BMP2, PLA2G1B, SESN2, GOT1, LIPG, FADS1, PROX1, IFNG, FADS2, RFK, PTGS2, MID1IP1, CRY1, EGR1, UGDH, CYP27B1, PGM2, SLC35B4, SDS, RBP1, PTGES3, ASNSD1, SNAI2, PGK1, CYP51A1, EBP, KAT2A, LTA4H, PPP4R3B, ALDH18A1, APIP, INSIG1, MTHFD1, PARK7, TPI1, NUS1, PGAM1, TPK1, CYP39A1, ACACA, MTAP, SPTLC1, COQ2, PRKAA2, PTGES2, MIF, PPARGC1A, ACER2, IMPA1* |
| GO:0030163 | Protein catabolic process | 266 | 1.512 | 1.75E-04 | 1.09E-03 | *TP53INP2, TNFAIP3, RNF122, IL10, TIMP1, ADRA2A, DNAJB9, IFNG, PLK3, DDIT3, RNF19B, PSMB3, PSMD10, UCHL1, HSP90B1, KLF1, NDFIP2, TNFAIP1, ZFAND2A, SIAH1, MAD2L1, PSMA3, PSMA5, UBE2D3, TMEM126A, UBE2D1, PAQR3, UCHL3, USP44, RNF11, FBXO4, RNF5, PSMB6, PSME3, STT3B, TRAF2, TAF9, RNF180, SUMO2, RAD23A, KLHL2, AZIN1, PSMC5, UCHL5, YOD1, PDCL3, CUL5, NEMF, PSMA4, DDRGK1, UBXN4, PARK7, PSMA2, PSMD8, PSMA1, SOCS4, SELENOS, RNF115, STYX, SOCS5, FEM1B, MIB1, PSMD2, SIAH2, FYN, RAD23B, PSMC2, PSMB1, AKIRIN2, PSMC1, TMEM168, PSMB4, NRDC, ELOC, CHMP2B, ANAPC10, PSMD1, LTN1, PSMD4, UBB, CASP8, CALR, VCP, IDE, DERL3, RNFT2, PSME2, PSMB7, MAPK9, UBE2B, WAC, SNF8, SPOP, WWP1, KLHDC3, ERLEC1, RIC1, BCAP31, RNF14, PIN1, ATE1, ARK2N* |
| GO:0051046 | Regulation of secretion | 116 | 1.512 | 3.13E-03 | 1.21E-02 | *IL6, IL1A, IL1B, INHBA, GJA1, SOX11, PTGER3, S100A8, LIF, ADRA2A, BMP2, IFNG, KCNJ11, CRY1, FFAR2, TLR4, C7H6orf125, TGFB3* |

| | | | | | | |
|---|---|---|---|---|---|---|
| GO:0045935 | Positive regulation of nucleobase-containing compound metabolic process | 395 | 1.510 | 2.65E-05 | 2.14E-04 | *IL1A, WT1, APLN, INHBA, SOX11, FOSL1, SAMHD1, LIF, TP53INP2, IL10, BMP2, PLA2G1B, CEBPG, TFRC, NFKBIA, NFIL3, EGR2, JUNB, PROX1, IER5, MORF4L2, DDX21, ACTL6A, DDX3X, BAZ1A, LYAR, GNL3, UCHL1, IRF1, HMBOX1, EGR1, DCPS, TLR4, DIMT1, ECD, TGFB3, KLF1, ZFP36, STAT3, NCOA7, CCT4, PTGES3, CD40, GATA1, BMPR2, HDAC2, ILF2, SERTAD1, WDR75, CCT2, LEO1, EXOSC3, AGO2, TBX18, PIM1, RIDA, FBXO4, NME2, RHOQ, TNIP2, TAF9, IER2, SUMO2, SMAD3, KAT2A, MED1, CEBPB, CNOT7, MED4, MED21, PDE12, PAIP1, RPRD1B, DDRGK1, MORF4L1, PARK7* |
| GO:0007049 | Cell cycle | 406 | 1.501 | 2.53E-05 | 2.09E-04 | *IL1A, INHBA, CCL2, SERPINE1, GJA1, ATF5, LIF, KIF18A, IL10, INHA, CDKN1A, BMP2, BTG3, JUNB, PROX1, PLK3, MORF4L2, TCIM, DDIT3, ACTL6A, DDX3X, RRS1, WDR12, IRF1, GEM, TIPIN, CRY1, CHORDC1, CKS2, NAA50, MAD2L1, STAT3, NLE1, PARD6G, RHOB, RPS27L, CALM2, GADD45A, PTPN11, TIPRL, GADD45B, USP44, TUBB1, FBXO4, CCNG1, PSME3, SPDL1, DCTN6, PRKAR2A, FBXO5, ATM, KAT2A, MNAT1, WNT16, TUBB, DRG1, PPP2CA, MED1, PRCC, HAUS1, PPM1G, MCMBP, FZD3, ZPR1, RPRD1B, MORF4L1, NAA10, AKAP5, CDC25C, WEE1, RBBP8, NPM1, NCAPG, KPNB1, RMI1, DR1, ARF6, TRNP1, FEM1B, NUBP1, REEP4, NPPC, HAUS6, RCC1, CDK2AP2, SEPTIN11, TBRG1, CDK4, SKA2, STAG2, PRKAR1A, SUGT1, SMC6, CCND2, CHMP2B, ROMO1, ENSA, NHERF1, ING3, XRCC2, UBB, CALR, VCP, RAD1, PSME2, HJURP, NAT10, CDC42, RAD21, SLC25A5, THAP1, ARL3, REEP3, MBIP, PPP2R2A, UBE2B, WAC, PAFAH1B1, BIRC5, RHOA, MYC, SBDS, CALM1, APP, NEK6, CDK2, CDK7, PIN1, CETN4, DCTN3, TERF1, ING2, RHNO1, MAPK14, SEPTIN8, BECN1, EML4, NKX3-1, SPAST, LSM14A, USP16, CFL1, ZZZ3, MITD1, TUBA1B, BUB3, RB1, MZT1, SGO2, TAOK1, UBXN2B* |
| GO:0051254 | Positive regulation of RNA metabolic process | 348 | 1.481 | 7.85E-05 | 5.70E-04 | *IL1A, WT1, APLN, INHBA, SOX11, FOSL1, SAMHD1, LIF, TP53INP2, IL10, BMP2, PLA2G1B, NFKBIA, NFIL3, EGR2, JUNB, PROX1, IER5, DDX21, ACTL6A, DDX3X, LYAR, GNL3, IRF1, HMBOX1, EGR1, DCPS, TLR4, DIMT1, ECD, TGFB3, KLF1, ZFP36, STAT3, NCOA7, CD40, GATA1, BMPR2, HDAC2, ILF2, SERTAD1, WDR75, LEO1, AGO2, TBX18, PIM1, RIDA,* |

| | | | | | | |
|---|---|---|---|---|---|---|
| | | | | | | *NME2, RHOQ, TNIP2, TAF9, IER2, SUMO2, SMAD3, KAT2A, MED1, CEBPB, CNOT7, MED4, MED21, PDE12, PAIP1, RPRD1B, DDRGK1, PARK7* |
| GO:0034655 | Nucleobase-containing compound catabolic process | 148 | 1.478 | 5.58E-03 | 1.87E-02 | *SAMHD1, LOC100515984, PDE4D, DDIT4, RBM8A, UCHL1, DCPS, HINT1, ZFP36, REXO4, PDE4B, EXOSC3, PHAX, PGK1, AGO2, RIDA, SSB, ETF1, SLC4A1, TRAF2, SKIC8, DCTPP1, ATM, DDX49, HK2, PPP2CA, CNOT7, SMG8, PDE12, PAIP1, ZPR1, NUDT11, TPI1, FASTKD2, PELO, NPM1, PNRC2, EXOSC4, PGAM1, LSM5, ROCK1, CAPRIN1, EIF4A3, PNPT1, FITM2, NCBP2, NAF1, TIRAP, PABPC4, MTREX, DXO* |
| GO:1903530 | Regulation of secretion by cell | 107 | 1.475 | 4.79E-03 | 1.69E-02 | *IL6, IL1A, IL1B, INHBA, GJA1, SOX11, LIF, ADRA2A, BMP2, IFNG, KCNJ11, CRY1, FFAR2, TLR4, C7H6orf125, TGFB3* |
| GO:0010817 | Regulation of hormone levels | 108 | 1.472 | 5.81E-03 | 1.93E-02 | *IL6, IL1B, DUOXA2, INHBA, GJA1, SOX11, DUOX1, LIF, DUOX2, ADRA2A, BMP2, IFNG, KCNJ11, CRY1, FFAR2, EGR1, C7H6orf125, RBP1, RAB1A, PTPN11, TUBB1, GCNT4, CCN3* |
| GO:0010629 | Negative regulation of gene expression | 279 | 1.472 | 2.44E-04 | 1.45E-03 | *WT1, CXCL8, APLN, SERPINE1, GJA1, SOX11, EIF4EBP1, TNFAIP3, SAA3, IL10, ARG2, BMP2, SESN2, H2AZ1, THBS1, IFNG, DDIT3, DDX3X, BAZ1A, RBM8A, DCPS, TLR4, TGFB3, NDFIP2, CBX1, ZFP36, HAT1, EIF2S1, REL, HDAC2, EXOSC3, TSKU, CNPY2, PTPN11, AGO2, IL27RA, BMI1, RIDA, PDCD10, SSB, ETF1, TRAF2, PRDM1, SKIC8, ATM, LMNB1, IGFBP5, SDR16C5, CNOT7, SMG8, PDE12, PAIP1, DDRGK1, PARK7, C5AR2, FASTKD2, PELO, NPM1, SOCS5, PNRC2, CLP1, EXOSC4, PAIP2B, LSM5, CBX5, ROCK1, CAPRIN1, TASOR, RBM42, FYN, EIF4A3, PNPT1, PRKAA2, MIF, CREB3L1, PRKAR1A, NCBP2, SERBP1, TIRAP, SRP9, PABPC4, TSNAX* |
| GO:0072594 | Establishment of protein localization to organelle | 116 | 1.472 | 5.17E-03 | 1.75E-02 | *NEURL3, KPNA2, TNFAIP3, TIMM8A, NFKBIA, IFNG, LOC100157391, DDIT3, ZFAND2A, STAT3, HIKESHI, AGK, TMEM126A, CHCHD4, SRP14, IMMP1L, SRP54, TOMM40, MED1, SRPRA, SSR3, ZPR1, BBC3, TNPO1, KPNB1, KPNA1, AKIRIN2, TXNIP, SEC63, BCS1L, TIMM21, SRP9, ROMO1, ZFAND6, HYAL2, IPO4, TRAM1, SNF8, ELAVL1, BCAP31, TERF1, MAPK14, SNUPN, SRP19* |
| GO:0043412 | Macromolecule modification | 499 | 1.466 | 1.86E-05 | 1.79E-04 | *LIF, TNFAIP3, RNF122, B3GNT5, PLA2G1B, NHP2, GAR1, C1GALT1, SOCS3, WDR4, ST6GALNAC1, EGR2, DNAJB11, B4GALT5, IFNG, RXYLT1, FBL, EMG1, TPST1, RNF19B,* |

| | | | | | | |
|---|---|---|---|---|---|---|
| | | | | | | *PSMD10, DDX3X, PUS7L, GNL3, UCHL1, EIPR1, CRY1, KLHL25, QTRT1, EGR1, ELP4, CISH, DIMT1, OTUB2, HINT1, ST6GAL2, NDFIP2, TNFAIP1, SIAH1, NCOA7, PUS3, GATA1, EPHA4, MOGS, EXOSC3, UBE2D3, UBE2D1, UCHL3, PTPN11, SLC39A8, DKC1, B3GALNT1, PCMT1, FUT2, USP44, RNF11, FBXO4, DCUN1D5, RNF5, NOP2, HSPB1, PCNP, STT3B, PRKAR2A, TRAF2, QTRT2, RNF180, SUMO2, FBXO5, DAD1, ATM, OSTC, SOCS7, KAT2A, IL15, PPP2CA, UCHL5, ST8SIA2, COPS6, DPM1, RAMAC, ST3GAL1, CUL5, TYW5, DDRGK1, PARK7, SOCS4, NPC1, NUS1, RNF115, PRNP, RNF138, PPIA, GALNT1, SOCS5, THUMPD1, HECTD2, ELP1, TRMT6, DR1, EXOSC4, MGAT2, ST3GAL3, B3GALT6, FEM1B, MIB1, NSUN2, SUMO1, BUD23, DCAF13, SIAH2, RPN1, USP53, GTPBP4, PHF23, FYN, SOCS1, UBA6, MRM1, NRP1, PRKAA2, PPP1CB, USP5, USP39, ARHGEF2, OTUD6B, RPN2, WTAP, ELP5, NAF1, FBXO33, USP15, LTN1, ALG14, TRMT112, SPRED1, UBB, UBL5, VCP, ITGA5, DERL3, NAT10, METTL3, COPS8, ASB15, FTSJ1, WSB2, MBIP, ALG3, MAPK9, PPP2R2A, UBA3, UBE2B, BIRC5, SPOP, RAMP1, HSD17B10, WWP1, MINDY3, SAE1, NEK6, TRMT5, PRPF19, RNF14, NOP10, UFL1, JAK2, BAG2, TRIM23, RLIM, ELP2, TICAM1, UBE3A, OTUB1, COPS4, TNKS2, DCUN1D4* |
| GO:0051247 | Positive regulation of protein metabolic process | 172 | 1.464 | 2.62E-03 | 1.05E-02 | *INHBA, LIF, TNFAIP3, ADRA2A, PLA2G1B, NFKBIA, THBS1, IFNG, PLK3, PSMD10, DDX3X, GNL3, CRY1, SOAT1, CASP3, EGR1, C7H6orf125, NDFIP2, ZFAND2A, GATA1, HDAC2, NIBAN1, UBE2D1, PAQR3, PTPN11, FBXO4, DCUN1D5, SSB, RNF180, SUMO2, IL15, PAIP1, DDRGK1, SOCS4, PRNP, FASTKD2, PPIA, NPM1, SOCS5, ABCF1, PHF23, EIF4A3, NRP1, PRKAA2, ARHGEF2, NRDC, MAPK8, FBXO33, UBB, CASP8, ITGA5, RNFT2, NAT10, METTL3, SP1, MAPK9, SNF8, RAMP1, PLGRKT, APP, ELAVL1, BCAP31, EIF5A, JAK2, BAG2, TICAM1, UBE3A, DCUN1D4, BCL10, IGF2BP1, USP16, HBEGF, TAOK1* |
| GO:0070887 | Cellular response to chemical stimulus | 463 | 1.463 | 2.32E-05 | 2.07E-04 | *AMCF-II, IL1A, IL1B, NR4A3, WT1, CXCL8, GDF15, CCL2, GJA1, CHRNA3, MT-2B, POSTN, MMP8, AQP5, SLC8A3, S100A8, CXCL10, RDH12, TNFAIP3, IL10, PPBP, ADRA2A, CXCL13, PLA2G1B, SESN2, NFKBIA, GOT1, AQP3, THBS1, JUNB, FABP4, SLC38A2, HTR2B, CPNE8, EFHD1, DDIT3, ACKR4, DDX3X, DDIT4, CCL8, UCHL1, NDNF, CRY1, FFAR2, CXCL11,* |

| | | | | | | |
|---|---|---|---|---|---|---|
| | | | | | | *NR4A2, CFTR, TLR4, TMEM38B, TFPI, ACKR2, LOC100511937, SUV39H2, LAMTOR2, TNFSF14, ZFAND2A, ZFP36, HBE1, STAT3, NCOA7, PTGES3, EIF2S1, SOD1, SNAI2, GSTM3, RHOB, NFKBIB, PTPN11, SLC39A8, PIM1, PDCD10, RAB13, SLC12A2, RHOQ, TNIP2, CCN3, SLC41A1, GSTO1, TRAF2, RAP1B, WNT16, MED1, IGFBP5, CEBPB, INSIG1, SLC25A33, PDE12, DDRGK1, PARK7, BBC3, SELENOS, CCL4, CALCRL, PRNP, C5AR2, IL16, PPIA* |
| GO:0016477 | Cell migration | 339 | 1.461 | 2.61E-04 | 1.52E-03 | *AMCF-II, MADCAM1, RASGEF1A, NR4A3, SELE, CXCL8, CRB2, GDF15, CCL2, GJA1, THY1, ANGPT2, DCLK1, S100A8, CXCL10, IGFBP6, TNFAIP3, PPBP, ADRA2A, CXCL13, BMP2, PLA2G1B, S100A12, NARS1, THBS1, PHLDA2, PROX1, IFNG, TWIST2, HTR2B, DDIT3, ACKR4, CCL8, NDNF, FFAR2, CXCL11, DUSP10, ENPEP, NR4A2, TLR4, ACKR2, LOC100511937, LAMTOR2, TNFSF14, TNFAIP1, CD40, BMPR2, PTP4A1, SNAI2, RHOB, RAB1A, PTPN11, IL27RA, PDCD10, LRP12, RAB13, SLC12A2, SPDL1, CCN3, GNA13, LIMA1* |
| GO:0045893 | Positive regulation of DNA-templated transcription | 317 | 1.460 | 2.96E-04 | 1.62E-03 | *IL1A, WT1, APLN, INHBA, SOX11, FOSL1, SAMHD1, LIF, TP53INP2, IL10, BMP2, PLA2G1B, NFKBIA, NFIL3, EGR2, JUNB, PROX1, IER5, DDX21, ACTL6A, DDX3X, LYAR, GNL3, IRF1, HMBOX1, EGR1, TLR4, ECD, TGFB3, KLF1, STAT3, NCOA7, CD40, GATA1, BMPR2, HDAC2, ILF2, SERTAD1, WDR75, LEO1, TBX18, PIM1, NME2, RHOQ, TNIP2, TAF9, IER2, SUMO2, SMAD3, KAT2A, MED1, CEBPB, CNOT7, MED4, MED21, RPRD1B, DDRGK1, PARK7, MDFIC* |
| GO:1902680 | Positive regulation of RNA biosynthetic process | 317 | 1.460 | 2.96E-04 | 1.62E-03 | *IL1A, WT1, APLN, INHBA, SOX11, FOSL1, SAMHD1, LIF, TP53INP2, IL10, BMP2, PLA2G1B, NFKBIA, NFIL3, EGR2, JUNB, PROX1, IER5, DDX21, ACTL6A, DDX3X, LYAR, GNL3, IRF1, HMBOX1, EGR1, TLR4, ECD, TGFB3, KLF1, STAT3, NCOA7, CD40, GATA1, BMPR2, HDAC2, ILF2, SERTAD1, WDR75, LEO1, TBX18, PIM1, NME2, RHOQ, TNIP2, TAF9, IER2, SUMO2, SMAD3, KAT2A, MED1, CEBPB, CNOT7, MED4, MED21, RPRD1B, DDRGK1, PARK7, MDFIC* |
| GO:0009117 | Nucleotide metabolic process | 143 | 1.452 | 4.85E-03 | 1.69E-02 | *SAMHD1, NME1, LOC100515984, ADK, GOT1, RFK, CTPS1, PDE4D, ATP5MC1, DDIT4, UCHL1, ATP1B1, MDH1, NAMPT, HINT1, TP53I3, PDE4B, AK3, PGK1, COX11, ATP5PD,* |

| | | | | | | |
|---|---|---|---|---|---|---|
| | | | | | | *NME2, PPAT, ME1, ATP5MF, RHOQ, CMPK1, SLC4A1, DCTPP1, HK2, DCTD, PPP2CA, MTHFD1, ATP5MC3, ATP5ME, PARK7, NUDT11, TPI1, PRPS1, CTPS2, IMPDH2, ATP5F1A, ATP5PO, PGAM1, NPPC, RRM2B* |
| GO:0033554 | Cellular response to stress | 500 | 1.452 | 3.05E-05 | 2.42E-04 | *IL1A, CXCL8, GDF15, ATF3, SAMHD1, AQP5, SLC8A3, IGFBP6, XRCC4, TNFAIP3, ARG2, CEBPG, FAS, SESN2, DNAJB9, AQP3, HSP90AA1, IER5, PLK3, SLC38A2, EFHD1, MORF4L2, APEX1, TCIM, DDIT3, ACTL6A, KLF10, PSMD10, DDX3X, DDIT4, NDNF, TIPIN, CRY1, DUSP10, NR4A2, CFTR, TLR4, DNAJB1, SUV39H2, HSP90B1, CBX1, NCOA7, AEN, REXO4, EIF2S1, SOD1, HIKESHI, BMPR2, NIBAN1, SNAI2, PPP1R15B, GSTM3, RHOB, RPS27L, ALB, GADD45A, PTPN11, TIPRL, YWHAG, FBXO4, PDCD10, DCUN1D5, RNF5, GTF2H3, STT3B, NEIL3, DCTPP1, ATM, ZBTB1, RAD23A, MNAT1, WNT16, TERF2IP, PPP4R3B, LIG4, UCHL5, CEBPB, INSIG1, PLIN2, MCL1, YOD1, GTF2H5, DDRGK1, UBXN4, MORF4L1, PARK7, BBC3, SELENOS, RFC4, RNF138, RBBP8, MSH2, PPIA, NPM1, SOCS5, RMI1, MAP3K7, PRIMPOL, FEM1B, SAR1B, PRDX3, CBX5, XPA, SUMO1, RAD23B, PRKAA2, AKIRIN2, HAPSTR1, MIF, PPARGC1A, ATG5, SMC6, BCLAF1, MAP2K1, EIF2AK3, MAPK8, ROMO1, FAM162A, ASF1A, ING3, MTREX, XRCC2, SSRP1, SMDT1, SPRED1, CALR, SAR1A, VCP, RAD1, DERL3, RNFT2, METTL3, TIGAR, RAD21, STX2, FICD, BMPR1A, FAN1, POLR2I, MAPK9, SRPX, CHCHD2, MORC2, UBE2B, WAC, RIPK3, RHOA, PRDX5, MYC* |
| GO:1901137 | Carbohydrate derivative biosynthetic process | 198 | 1.451 | 2.66E-03 | 1.05E-02 | *GFPT2, NME1, ADK, B3GNT5, C1GALT1, ST6GALNAC1, B4GALT5, RXYLT1, RFK, CTPS1, ATP5MC1, GFPT1, GNPNAT1, SOAT1, UGDH, PGM2, ST6GAL2, GATA1, BMPR2, MOGS, AK3, MPI, COX11, SLC39A8, B3GALNT1, FUT2, ATP5PD, NME2, PPAT, ATP5MF, STT3B, DAD1, OSTC, ST8SIA2, DPM1, PGM3, ST3GAL1, PIGP, ATP5MC3, ATP5ME, TYW5, TPI1, NPC1, NUS1, PRPS1, CTPS2, GALNT1, IMPDH2, PIGK, ATP5F1A, ATP5PO, MGAT2, ST3GAL3, B3GALT6, NPPC, MTAP, RRM2B, RPN1* |
| GO:0051726 | Regulation of cell cycle | 280 | 1.450 | 5.46E-04 | 2.78E-03 | *IL1A, INHBA, CCL2, ATF5, LIF, IL10, INHA, CDKN1A, BMP2, BTG3, JUNB, PROX1, PLK3, MORF4L2, TCIM, DDIT3, ACTL6A, DDX3X, WDR12, IRF1, TIPIN, CRY1, MAD2L1, STAT3, NLE1, RHOB, RPS27L, CALM2, GADD45A, PTPN11, TIPRL, GADD45B, USP44, FBXO4,* |

| | | | | | | |
|---|---|---|---|---|---|---|
| | | | | | | *PSME3, SPDL1, PRKAR2A, FBXO5, ATM, KAT2A, MNAT1, DRG1, PPP2CA, MED1, PRCC, PPM1G, FZD3, ZPR1, RPRD1B, MORF4L1, NAA10, AKAP5, CDC25C, WEE1, NPM1, DR1, ARF6, TRNP1, FEM1B, NUBP1, NPPC, RCC1, CDK2AP2, TBRG1, CDK4, PRKAR1A, SMC6, CCND2, CHMP2B, NHERF1, ING3, CALR, RAD1, PSME2, NAT10, CDC42, RAD21, THAP1, MBIP, PPP2R2A, UBE2B, WAC, BIRC5, RHOA, CALM1, APP, NEK6, CDK7* |
| GO:0030155 | Regulation of cell adhesion | 171 | 1.446 | 3.74E-03 | 1.36E-02 | *IL1A, IL1B, RND1, NR4A3, SELE, CXCL8, ANGPT2, LIF, IL10, CD274, ARG2, BMP2, TFRC, SLC7A1, THBS1, PLAU, IFNG, TNC, NDNF, IRF1, DUSP10, TNFSF13B, TNFSF14, SNAI2, GPR4, PTPN11, FUT2, BMI1, YWHAG* |
| GO:0006753 | Nucleoside phosphate metabolic process | 195 | 1.444 | 2.38E-03 | 9.72E-03 | *GFPT2, SAMHD1, NME1, LOC100515984, ADK, GOT1, RFK, CTPS1, PDE4D, ATP5MC1, DDIT4, GFPT1, UCHL1, ATP1B1, MDH1, GNPNAT1, NAMPT, UGDH, HINT1, TP53I3, PDE4B, AK3, PGK1, MPI, COX11, ATP5PD, NME2, ACSL4, PPAT, ME1, ATP5MF, RHOQ, CMPK1, SLC4A1, DCTPP1, HK2, DCTD, PPP2CA, PGM3, MTHFD1, ATP5MC3, ATP5ME, PDHA1, PARK7, NUDT11, TPI1, PRPS1, PANK3, CTPS2, IMPDH2, ATP5F1A, ATP5PO, PGAM1, ACACA, NPPC, PDHX, RRM2B* |
| GO:0006952 | Defense response | 450 | 1.443 | 2.48E-05 | 2.09E-04 | *IL6, AMCF-II, CCL19, ORM1, IL1A, IL1B, CXCL2, DUOXA2, GSDMA, CXCL8, CCL2, AGTR2, FOSL1, DUOX1, TIFA, MMP8, PTGER3, SAMHD1, S100A8, NDUFAF4, HP, CXCL10, IL1RAP, DUOX2, TNFAIP3, SAA3, IL10, PPBP, ARG2, CXCL13, BMP2, S100A12, CEBPG, FCN2, ACKR1, NFKBIA, THBS1, HSP90AA1, FABP4, IFNG, RNF19B, LBP, DDX3X, DDIT4, LYAR, CCL8, IRF1, NOD2, ATP1B1, FFAR2, CXCL11, DUSP10, PTGIR, CFB, TLR4, BPIFB1, TICAM2, IFNE, ZFP36, STAT3, KCNK6, IL33, CD40, IFNAR1, POLR3D, MLKL, GPR4, TMEM126A, RAB1A, HEXIM1, NFKBIB, IFNGR1, SLC39A8, IL27RA, PIM1, USP44, IL4R* |
| GO:0022407 | Regulation of cell-cell adhesion | 103 | 1.442 | 9.41E-03 | 2.75E-02 | *IL1A, IL1B, NR4A3, SELE, IL10, CD274, ARG2, BMP2, TFRC, SLC7A1, IFNG, IRF1, DUSP10, TNFSF13B, TNFSF14* |
| GO:0043161 | Proteasome-mediated ubiquitin- | 100 | 1.441 | 1.26E-02 | 3.31E-02 | *RNF122, DDIT3, PSMB3, PSMD10, TNFAIP1, ZFAND2A, SIAH1, PSMA5, UBE2D1, PAQR3, USP44, FBXO4, TAF9, RNF180, SUMO2, RAD23A, KLHL2, PSMC5, NEMF, DDRGK1,* |

| | | | | | | |
|---|---|---|---|---|---|---|
| | dependent protein catabolic process | | | | | *PARK7, PSMD8, SOCS4, STYX, SOCS5, FEM1B, SIAH2, RAD23B, AKIRIN2, PSMC1, ANAPC10, LTN1, PSMD4, VCP, PSMB7, MAPK9, UBE2B, WAC, SPOP, WWP1, KLHDC3* |
| GO:0040008 | Regulation of growth | 121 | 1.440 | 9.06E-03 | 2.71E-02 | *WT1, INHBA, GDF15, GJA1, MT-2B, SESN2, LPAR3, PROX1, PSMD10, DDX3X, DUSP10, LAMTOR2, SOD1, BMPR2, SERTAD1, PIM1, DCUN1D5, CCN3, DDX49, IGFBP5, SLC25A33, ZPR1, BBC3, IGFBP2, EXOSC4, SMAD4, NUBP1, BNIPL, NPPC, NRP1, ZNF639* |
| GO:1901700 | Response to oxygen-containing compound | 311 | 1.437 | 3.45E-04 | 1.86E-03 | *AMCF-II, IL1A, IL1B, NR4A3, CXCL8, GDF15, GJA1, CHRNA3, POSTN, DUOX1, MMP8, S100A8, CXCL10, DUOX2, RDH12, TNFAIP3, IL10, PPBP, ADRA2A, CXCL13, PLA2G1B, SESN2, NFKBIA, GOT1, AQP3, THBS1, PLK3, SLC38A2, HTR2B, DDIT4, KCNJ11, CRY1, FFAR2, DUSP10, EGR1, NR4A2, CFTR, PTGIR, TLR4, LOC100511937, TGFB3, ZFP36, STAT3, SOD1, IFNAR1, SNAI2, PPP1R15B, RHOB, NFKBIB, PTPN11, AARD, PIM1, PDCD10, RHOQ, TNIP2* |
| GO:0009890 | Negative regulation of biosynthetic process | 481 | 1.436 | 6.62E-05 | 5.02E-04 | *WT1, CXCL8, APLN, SERPINE1, GJA1, SOX11, ATF5, EIF4EBP1, TNFAIP3, SAA3, IL10, ARG2, BMP2, SESN2, H2AZ1, THBS1, FABP4, PROX1, IFNG, PLK3, DDIT3, KLF10, PSMD10, DDX3X, BAZ1A, LYAR, RBM8A, IRF1, CRY1, HMBOX1, EGR1, DCPS, NR4A2, TLR4, CRYAB, BASP1, DNAJB1, TGFB3, SUV39H2, NDFIP2, CBX1, ORMDL1, NKRF, ZFP36, HAT1, ID1, CEBPD, EIF2S1, REL, IL33, GATA1, HDAC2, SNAI2, EXOSC3, HEXIM1, TSKU, CNPY2, PTPN11, AGO2, IL27RA, BMI1, RIDA, PDCD10, STRAP, ZNF24, UXT, SSB, ETF1, ID3, TRAF2, PRDM1, SKIC8, ATM, ZBTB1, LMNB1, KAT2A, PSMC5, MED1, IGFBP5, CEBPB, SDR16C5, CNOT7, INSIG1, ZBTB33, SMG8, PDE12, PAIP1, ZC3H8, FRK, DDRGK1, PARK7, MDFIC, ZBTB2, C5AR2, NAB2, FASTKD2, PELO, NPM1, SOCS5, PNRC2, CLP1, DR1, EXOSC4, PAIP2B, SMAD4, LSM5, CBX5, ROCK1, CAPRIN1, TASOR, RBM42, FYN, EIF4A3, PNPT1, PRKAA2, POLE3, TXNIP, ZNF639, MIF, CREB3L1, PRKAR1A, NCBP2, NACC1, PFDN5, ZHX1, BTAF1, SERBP1, BCLAF1, TIRAP* |
| GO:0090407 | Organophosphate biosynthetic process | 177 | 1.433 | 2.33E-03 | 9.66E-03 | *GFPT2, NME1, ADK, HTR2B, RFK, CTPS1, ATP5MC1, GFPT1, GNPNAT1, NAMPT, UGDH, ABHD5, TMEM38B, ORMDL1, AK3, MPI, INPP1, COX11, LPGAT1, ATP5PD, NME2, PPAT, ATP5MF, CMPK1, PTPMT1, TAMM41, LPCAT3, DPM1, PGM3, MTHFD1, PIGP, ATP5MC3,* |

| | | | | | | |
|---|---|---|---|---|---|---|
| | | | | | | *ATP5ME, PDHA1, TPI1, NUS1, PRPS1, BPNT1, PANK3, CTPS2, IMPDH2, PIGK, ATP5F1A, ATP5PO, TPK1, ACACA, NPPC, CRLS1, PDHX, RRM2B* |
| GO:1901987 | Regulation of cell cycle phase transition | 105 | 1.432 | 1.46E-02 | 3.68E-02 | *INHBA, CCL2, ATF5, PROX1, PLK3, TCIM, DDX3X, TIPIN, CRY1, MAD2L1, RPS27L, PTPN11, TIPRL, USP44, FBXO4, PSME3, SPDL1, ATM, MNAT1, PPP2CA, CDC25C, WEE1, NPM1* |
| GO:0010558 | Negative regulation of macromolecule biosynthetic process | 465 | 1.432 | 6.97E-05 | 5.17E-04 | *WT1, CXCL8, APLN, SERPINE1, GJA1, SOX11, ATF5, EIF4EBP1, TNFAIP3, SAA3, IL10, ARG2, BMP2, SESN2, H2AZ1, THBS1, FABP4, IFNG, PLK3, DDIT3, KLF10, PSMD10, DDX3X, BAZ1A, LYAR, RBM8A, IRF1, CRY1, HMBOX1, EGR1, DCPS, NR4A2, TLR4, CRYAB, BASP1, DNAJB1, TGFB3, SUV39H2, NDFIP2, CBX1, NKRF, ZFP36, HAT1, ID1, CEBPD, EIF2S1, REL, IL33, GATA1, HDAC2, EXOSC3, HEXIM1, TSKU, CNPY2, PTPN11, AGO2, IL27RA, BMI1, RIDA, PDCD10, STRAP, ZNF24, UXT, SSB, ETF1, ID3, TRAF2, PRDM1, SKIC8, ATM, ZBTB1, LMNB1, KAT2A, PSMC5, MED1, IGFBP5, CEBPB, SDR16C5, CNOT7, ZBTB33, SMG8, PDE12, PAIP1, ZC3H8, FRK, DDRGK1, PARK7, MDFIC, ZBTB2, C5AR2, NAB2, FASTKD2, PELO, NPM1, SOCS5, PNRC2, CLP1, DR1, EXOSC4, PAIP2B, SMAD4, LSM5, CBX5, ROCK1, CAPRIN1, TASOR, RBM42, FYN, EIF4A3, PNPT1, PRKAA2, POLE3, TXNIP, ZNF639, MIF, CREB3L1, PRKAR1A, NCBP2, NACC1, PFDN5, ZHX1, BTAF1, SERBP1, BCLAF1, TIRAP, SRP9, PABPC4* |
| GO:0009967 | Positive regulation of signal transduction | 357 | 1.431 | 1.76E-04 | 1.09E-03 | *IL6, RASGEF1A, CRB2, INHBA, GDF15, AGTR2, SOX11, ATF3, TIFA, MMP8, IGF1, S100A8, LIF, FAM83B, IGFBP6, TNFAIP3, SAA3, IL10, ADRA2A, BMP2, PLA2G1B, S100A12, RSPO3, SESN2, LPAR3, THBS1, PROX1, HTR2B, DDX21, TCIM, DDIT3, DDX3X, TLR4, TICAM2, LAMTOR2, TNFSF14, ESM1, SIAH1, MAD2L1, RASSF2, STAT3, NLE1, KCNK6, REL, CD40, SOD1, BMPR2, GPR4, ADAM17, HEXIM1, PTPN11, DHH, PIM1, GADD45B, EGF, PDCD10, FZD10, TNIP2, CCN3, TRAF2, RAP1B, WNT16, TERF2IP, PPP2CA, MED1, IGFBP5, SLC15A3, MCL1, PARK7, SHOC2, BBC3, AVPI1, PRNP, PPIA, KRAS* |
| GO:0044057 | Regulation of system process | 105 | 1.431 | 1.47E-02 | 3.70E-02 | *IL6, IL1B, INHBA, GJA1, PTGER3, SLC8A3, ADRA2A, C5H12orf57, KCNE4, KCNE3, EGR2, ATP1B1, CRY1, TMEM38B, KCNJ2, SOD1, BMPR2, CALM2, PTPN11, TBX18* |

| | | | | | | |
|---|---|---|---|---|---|---|
| GO:0023056 | Positive regulation of signaling | 394 | 1.425 | 1.49E-04 | 9.65E-04 | *IL6, IL1B, RASGEF1A, CRB2, INHBA, GDF15, CCL2, AGTR2, SOX11, ATF3, TIFA, MMP8, IGF1, SLC8A3, S100A8, LIF, FAM83B, IGFBP6, TNFAIP3, SAA3, IL10, ADRA2A, BMP2, PLA2G1B, S100A12, RSPO3, SESN2, LPAR3, THBS1, PROX1, HTR2B, DDX21, TCIM, DDIT3, DDX3X, FFAR2, TLR4, TICAM2, LAMTOR2, TNFSF14, ESM1, SIAH1, MAD2L1, RASSF2, STAT3, BVES, NLE1, KCNK6, REL, CD40, SOD1, BMPR2, GPR4, ADAM17, HEXIM1, PTPN11, DHH, PIM1, GADD45B, EGF, PDCD10, FZD10, TNIP2, CCN3, TRAF2, RAP1B, WNT16, TERF2IP, PPP2CA, MED1, IGFBP5, SLC15A3, MCL1, PARK7, SHOC2, BBC3, AKAP5, AVPI1, PRNP, PPIA, KRAS* |
| GO:0033365 | Protein localization to organelle | 237 | 1.418 | 2.40E-03 | 9.72E-03 | *NEURL3, KPNA2, TNFAIP3, TIMM8A, SESN2, NFKBIA, ARL5B, IFNG, LOC100157391, IFT20, EMC9, DDIT3, DDX3X, RRS1, GNL3, ZFAND2A, EMC4, STAT3, HIKESHI, AGK, TMEM126A, PAQR3, CHCHD4, MMGT1, SRP14, FBXO4, SSB, SPDL1, IMMP1L, SRP54, TOMM40, LMNB1, TERF2IP, MED1, INSIG1, EMC3, SRPRA, SSR3, ZPR1, DDRGK1, PARK7, BBC3, TNPO1, ARL1, KPNB1, ARF6, EDN1, KPNA1, TASOR, FYN, NRP1, PRKAA2, TBRG1, AKIRIN2, TXNIP, ABHD17B, SEC63, GET3, RAB33B, BCS1L, TIMM21, GET1, TMED10, SRP9, ROMO1, CALR, VCP, RAD21, ARL3, ZFAND6, HYAL2, IPO4, TRAM1, GDI2, SNF8, RAP1A, ELAVL1, BCAP31, PIN1, TERF1, MAPK14, SNUPN, TNKS2, SRP19, SNX10, BUB3, ANKRD13C* |
| GO:0002521 | Leukocyte differentiation | 147 | 1.418 | 7.60E-03 | 2.35E-02 | *IL6, IL1A, IL1B, LIF, IL10, BMP2, CEBPG, NFIL3, JUNB, IFNG, KLF10, IRF1, DUSP10, EGR1, TNFSF13B, TLR4, RASSF2, STAT3, GATA1, BMI1, USP44, SRP54, LRRC17, PRDM1, ATM, ZBTB1, KAT2A, LIG4, MED1, CEBPB, ST3GAL1* |
| GO:0010647 | Positive regulation of cell communication | 392 | 1.413 | 1.44E-04 | 9.65E-04 | *IL6, IL1B, RASGEF1A, CRB2, INHBA, GDF15, CCL2, AGTR2, SOX11, ATF3, TIFA, MMP8, IGF1, SLC8A3, S100A8, LIF, FAM83B, IGFBP6, TNFAIP3, SAA3, IL10, ADRA2A, BMP2, PLA2G1B, S100A12, RSPO3, SESN2, LPAR3, THBS1, PROX1, HTR2B, DDX21, TCIM, DDIT3, DDX3X, FFAR2, TLR4, TICAM2, LAMTOR2, TNFSF14, ESM1, SIAH1, MAD2L1, RASSF2, STAT3, NLE1, KCNK6, REL, CD40, SOD1, BMPR2, GPR4, ADAM17, HEXIM1, PTPN11, DHH, PIM1, GADD45B, EGF, PDCD10, FZD10, TNIP2, CCN3, TRAF2, RAP1B, WNT16, TERF2IP,* |

| | | | | | | |
|---|---|---|---|---|---|---|
| | | | | | | *PPP2CA, MED1, IGFBP5, SLC15A3, MCL1, PARK7, SHOC2, BBC3, AKAP5, AVPI1, PRNP, PPIA, KRAS* |
| GO:0009894 | Regulation of catabolic process | 237 | 1.404 | 3.54E-03 | 1.33E-02 | *TNFAIP3, SAA3, IL10, TIMP1, ADRA2A, SESN2, IFNG, PLK3, PSMD10, RAB39B, DDIT4, RBM8A, UCHL1, DRAM2, DCPS, ABHD5, ZFAND2A, ZFP36, MAD2L1, PAQR3, PHAX, AGO2, RIDA, RNF5, PSME3, SLC4A1, TRAF2, TAF9, RNF180, SUMO2, DDX49, CDC37, AZIN1, PPP2CA, CNOT7, MCL1, PDCL3, TMEM39A, PDE12, PAIP1, DDRGK1, PARK7, SOCS4, STYX, FASTKD2, NPM1, SOCS5, PSMD2, ROCK1, CAPRIN1, SPTLC1, PHF23, FYN, EIF4A3, PRKAA2, TMEM168, DRAM1, NRDC, NAF1, TIRAP, PABPC4, PSMD1, UBB, ITGB1, STK38L, RNFT2, PSME2, TIGAR, SLC25A5, CISD2, MAPK9, GNAI3, WAC, SNF8, APP, ELAVL1, RIC1, BCAP31, ARID5A, PIN1, BAG2, TRIM23, PHKG2, TICAM1, BECN1, UBE3A, PSMD3, QKI, UBQLN4, IGF2BP1* |
| GO:0030097 | Hemopoiesis | 236 | 1.402 | 1.61E-03 | 6.97E-03 | *IL6, IL1A, IL1B, INHBA, CSF3, SLC8A3, LIF, IL10, BMP2, CEBPG, NFKBIA, NFIL3, JUNB, HSPA9, IFNG, TCIM, KLF10, LYAR, IRF1, DUSP10, CASP3, EGR1, TNFSF13B, TLR4, KLF1, ZFP36, RASSF2, STAT3, CEBPD, SLC1A5, GATA1, LEO1, PTPN11, PIM1, BMI1, USP44, GPATCH4, TUBB1, SRP54, SLC4A1, LRRC17, PRDM1, SKIC8, ATM, ZBTB1, EBP, KAT2A, LIG4, MED1, CEBPB, PGM3, ST3GAL1* |
| GO:0055086 | Nucleobase-containing small molecule metabolic process | 205 | 1.389 | 3.38E-03 | 1.28E-02 | *GFPT2, SAMHD1, NME1, LOC100515984, ADK, GOT1, RFK, CTPS1, PDE4D, ATP5MC1, DDIT4, GFPT1, UCHL1, ATP1B1, MDH1, GNPNAT1, NAMPT, UGDH, PGM2, HINT1, TP53I3, PDE4B, AK3, PGK1, MPI, COX11, ATP5PD, NME2, ACSL4, PPAT, ME1, ATP5MF, RHOQ, CMPK1, SLC4A1, DCTPP1, HK2, DCTD, PPP2CA, PGM3, MTHFD1, ATP5MC3, ATP5ME, PDHA1, PARK7, NUDT11, TPI1, PRPS1, PANK3, CTPS2, IMPDH2, ATP5F1A, ATP5PO, PGAM1, ACACA, NPPC, MTAP, PDHX, RRM2B* |
| GO:0050673 | Epithelial cell proliferation | 117 | 1.389 | 1.07E-02 | 3.04E-02 | *IL6, APLN, CCL2, SOX11, NME1, TNFAIP3, IL10, THBS1, PROX1, HTR2B, DUSP10, ZFP36, BMPR2, SNAI2, FUT2, PDCD10, CCN3, PRDM1, WNT16, MED1, IGFBP5, CEBPB, SDR16C5, PDCL3* |

| | | | | | | |
|---|---|---|---|---|---|---|
| GO:0071396 | Cellular response to lipid | 120 | 1.388 | 1.45E-02 | 3.68E-02 | *AMCF-II, IL1A, IL1B, CXCL8, MMP8, CXCL10, TNFAIP3, IL10, PPBP, CXCL13, NFKBIA, DDIT4, CRY1, FFAR2, CFTR, TFPI, ZFP36, PTGES3, SNAI2, NFKBIB, PTPN11, PIM1, TNIP2* |
| GO:1901701 | Cellular response to oxygen-containing compound | 243 | 1.387 | 2.25E-03 | 9.42E-03 | *AMCF-II, IL1A, IL1B, NR4A3, CXCL8, GDF15, GJA1, CHRNA3, POSTN, MMP8, CXCL10, RDH12, TNFAIP3, IL10, PPBP, ADRA2A, CXCL13, PLA2G1B, SESN2, NFKBIA, GOT1, SLC38A2, HTR2B, DDIT4, FFAR2, NR4A2, CFTR, TLR4, LOC100511937, ZFP36, STAT3, SOD1, SNAI2, RHOB, NFKBIB, PTPN11, PIM1, PDCD10, RHOQ, TNIP2, TRAF2, RAP1B, MED1, IGFBP5, INSIG1, SLC25A33, PARK7, SELENOS, CALCRL, PRNP, IRF8, TRA2B, PRDX3, NPPC, ROCK1, FYN, SOCS1, PRKAA2, CDK4, ARHGEF2, SLC2A8, MIF, PRKAR1A, GRB10, ADAM9, MAPK8, ROMO1, GNA14* |
| GO:0098609 | Cell-cell adhesion | 182 | 1.386 | 4.26E-03 | 1.53E-02 | *MADCAM1, IL1A, IL1B, NR4A3, CLSTN3, SELE, CRB2, S100A8, PRTG, IL10, CD274, ARG2, BMP2, TFRC, ICAM4, SLC7A1, IFNG, IRF1, PCDH11X, DUSP10, TMEM47, TNFSF13B, TNFSF14, BVES, NECTIN1, GATA1, VCAM1, ICAM1, SLC39A8, BMI1, YWHAG, TUBB1* |
| GO:0042592 | Homeostatic process | 401 | 1.383 | 9.11E-04 | 4.27E-03 | *IL6, IL1A, IL1B, CRB2, INHBA, GDF15, GJA1, MT-2B, STC2, SLC8A3, CXCL10, TP53INP2, TNFAIP3, ADRA2A, STEAP2, CEBPG, TFRC, FAS, SESN2, GOT1, AQP3, ST6GALNAC1, KCNE3, LIPG, FABP4, HSPA9, HTR2B, PRND, DDIT3, DDX3X, LYAR, GHITM, CCL8, ATP1B1, CRY1, SLC28A2, FFAR2, SOAT1, CXCL11, DRAM2, CASP3, EGR1, CFTR, TNFSF13B, ABHD5, TLR4, TMEM38B, KLF1, ORMDL1, ZFP36, RASSF2, STAT3, SLC1A5, NLE1, SLC31A1, CD40, GATA1, SOD1, CALM2, CLNS1A, TSKU, FXYD2, PTPN11, COX11, SLC39A8, SLC30A5, FBXO4, GCNT4, STRAP, SLC12A2, TXNRD1, CCN3, SLC41A1, BPIFA1, LIMA1, SLC4A1, MINPP1, FTH1, ATP6V1F, HK2, PPP2CA, MED1, IGFBP5, CEBPB, INSIG1* |
| GO:0061061 | Muscle structure development | 141 | 1.376 | 1.21E-02 | 3.21E-02 | *WT1, GDF15, GJA1, SOX11, ATF3, LIF, CXCL10, MSC, BMP2, EGR2, PROX1, IFT20, DDIT3, UCHL1, RCAN1, EGR1, C7H6orf125, TNFSF14, BVES, BMPR2* |
| GO:0051276 | Chromosome organization | 133 | 1.360 | 1.20E-02 | 3.21E-02 | *KIF18A, GAR1, ACTL6A, RRS1, GNL3, GEM, HMBOX1, NAA50, MAD2L1, CCT4, PTGES3, CCT2, USP44, FBXO4, SPDL1, ATM, DRG1, TERF2IP, LIG4, MCMBP, NAA10, NCAPG, KPNB1, NPPC, RCC1, SKA2, TOP2B, STAG2, CENPP, SUGT1, SMC6, NAF1, CHMP2B,* |

| | | | | | | |
|---|---|---|---|---|---|---|
| | | | | | | *HJURP, NAT10, RAD21, CENPC, UBE2B, BIRC5, RHOA, MYC, NEK6, BANF1, NOP10, TERF1, BECN1, EML4, TNKS2, LSM14A, SMC3, MZT1* |
| GO:0051649 | Establishment of localization in cell | 478 | 1.358 | 2.87E-04 | 1.62E-03 | *GOLT1A, GJA1, KIF18A, KPNA2, ADRA2A, STEAP2, ARL4D, TIMM8A, NFKBIA, AQP3, EGR2, HSPA9, ARL5B, IFNG, PLK3, SLC18B1, IFT20, GOLT1B, SLC30A4, RRS1, UCHL1, GEM, EIPR1, RANBP1, GPM6B, TMEM38B, NOP9, SEC23B, NDFIP2, MAD2L1, STAT3, ENY2, SEC22B, SEC24A, SLC31A1, THOC3, HIKESHI, AGK, RHOB, NXT2, TMEM126A, PHAX, CALM2, RAB1A, COPB2, SEC61B, PTPN11, SLC39A8, EIF6, COPB1, PREB, PDCD10, RAB13, SSB, SPDL1, HSPB1, NXT1, SRP54, SEC23A, RAP1B, ATP6V1F, LMNB1, ERGIC2, MED1, STX11, INSIG1, RAB14, SRPRA, NEMF, VTI1B, ZPR1, USO1, PARK7, DPY30, AKAP5, YIPF5, NPC1, SELENOS, NUS1, TNPO1, NPM1, ARL1, CLTC, RAB5A, KPNB1, STX7, AP1S2, ARF6, COPE, SAR1B, KPNA1, ROCK1, ARMCX3, AKIRIN2, TXNIP, SKA2, SEC63, VPS26A, TOMM20, SNX19, RHOBTB3, NCBP2, BCS1L, AP3M1, CHMP2B, MAP2K1, TMED10, BLOC1S6, ROMO1, SNX5, XPO5, NHERF1, THOC2, NOL6, UBB, LDLR, CALR, ARPC3, LIN7C, SAR1A, VCP, TOMM22, STARD3NL, NUTF2, DERL3, YIF1A, CDC42, STX2, ASB15, HYAL2, C2CD2L, IPO4, TMED9, UBE2B, TRAM1, PAFAH1B1, SNAP23, BIRC5, SNF8, RAMP1, RAP1A, IFT74, YIF1B, ATP6V1H, RAB3D, RAB27A, CALM1, APP, ELAVL1, VPS29, RIC1, BCAP31, SDAD1, IPO8, RAB9A, AP3S2, TERF1, TRIM23, TMEM41B, FYTTD1, MAPK14, BECN1, SNUPN, EML4, SPAST, CFL1, SNX10, BUB3, STARD4, AFTPH, UBXN2B, STX6, NUP153, DPCD* |
| GO:0048871 | Multicellular organismal-level homeostasis | 185 | 1.357 | 6.46E-03 | 2.05E-02 | *IL1A, IL1B, CRB2, INHBA, GDF15, GJA1, TP53INP2, TNFAIP3, CEBPG, FAS, AQP3, ST6GALNAC1, FABP4, HSPA9, DDIT3, LYAR, SLC28A2, DRAM2, CASP3, EGR1, CFTR, TNFSF13B, TLR4, KLF1, ZFP36, RASSF2, STAT3, SLC1A5, NLE1, GATA1, SOD1, TSKU, PTPN11, SLC39A8, GCNT4, STRAP, CCN3, BPIFA1, SLC4A1* |
| GO:0043066 | Negative regulation of apoptotic process | 206 | 1.350 | 9.33E-03 | 2.74E-02 | *WT1, CCL2, ATF5, IGF1, TNFAIP3, IL10, ARG2, HIGD1A, FAS, THBS1, PLK3, HTR2B, TCIM, PSMD10, DDX3X, GHITM, NDNF, NR4A2, CRYAB, MAD2L1, GATA1, SOD1, SNAI2, IL27RA, PIM1, BMI1, PDCD10, TNIP2, PSME3, HSPB1, TAF9, DAD1, WNT16, LIG4, MED1, CEBPB,* |

| | | | | | | |
|---|---|---|---|---|---|---|
| | | | | | | *ST3GAL1, MCL1, FZD3, ZPR1, PARK7, SELENOS, HIPK3, PPIA, KRAS, NPM1, API5, PRDX3, NPPC, ROCK1, SIAH2, NRP1, PRKAA2, MIF, ANGPTL4, PPARGC1A* |
| GO:0022402 | Cell cycle process | 327 | 1.349 | 2.14E-03 | 9.08E-03 | *IL1A, INHBA, CCL2, SERPINE1, GJA1, ATF5, LIF, KIF18A, PROX1, PLK3, TCIM, DDX3X, RRS1, GEM, TIPIN, CRY1, CHORDC1, CKS2, NAA50, MAD2L1, PARD6G, RHOB, RPS27L, CALM2, PTPN11, TIPRL, USP44, TUBB1, FBXO4, CCNG1, PSME3, SPDL1, DCTN6, PRKAR2A, FBXO5, ATM, KAT2A, MNAT1, WNT16, TUBB, DRG1, PPP2CA, MED1, HAUS1, MCMBP, ZPR1, RPRD1B, NAA10, AKAP5, CDC25C, WEE1, NPM1, NCAPG, KPNB1, RMI1, ARF6, FEM1B, NUBP1, REEP4, NPPC, HAUS6, RCC1, CDK2AP2, SEPTIN11, CDK4, SKA2, STAG2, PRKAR1A, SUGT1, SMC6, CCND2, CHMP2B, ROMO1, ENSA, XRCC2, UBB, CALR, VCP, RAD1, PSME2, HJURP, NAT10, CDC42, RAD21, SLC25A5, ARL3, REEP3, PPP2R2A, UBE2B, WAC, PAFAH1B1, BIRC5, RHOA, MYC, SBDS, CALM1, APP, NEK6, CDK7, PIN1, CETN4, DCTN3, TERF1, ING2, RHNO1, MAPK14, SEPTIN8, BECN1, EML4, SPAST, LSM14A, USP16, CFL1, MITD1, BUB3, MZT1, SGO2, TAOK1, UBXN2B* |
| GO:0051301 | Cell division | 186 | 1.346 | 9.16E-03 | 2.72E-02 | *IL1A, IL1B, PPBP, PLK3, HTR2B, GNL3, TGFB3, CKS2, MAD2L1, PARD6G, RHOB, CALM2, BMI1, USP44, CCNG1, SPDL1, SEPTIN4, PPP1CC, FBXO5, KAT2A, CDC37, GNAI1, HAUS1, CDC25C, WEE1, RBBP8, NCAPG, DR1, ARF6, VPS4A, CDK2AP2, SEPTIN11, PPP1CB, SEPTIN7, TXNIP, CDK4, ARHGEF2, SKA2, STAG2, CCND2, CHMP2B, ANAPC10, ENSA, BOD1, PGF, MAPRE1, CDC42, CDC123, RAD21, ARL3, OIP5, PARD6B, REEP3, MBIP, GNAI3, PAFAH1B1, BIRC5, RHOA, MYC, CALM1, NEK6, CDK2, PIN1, DCTN3, ING2, SEPTIN8, BECN1, NKX3-1, SPAST, USP16, SMC3, CFL1, ZZZ3, MITD1, SGO2, SDE2* |
| GO:0045934 | Negative regulation of nucleobase-containing compound metabolic process | 285 | 1.341 | 3.70E-03 | 1.36E-02 | *WT1, SOX11, ATF5, SAA3, BMP2, FABP4, IFNG, PLK3, DDIT3, KLF10, PSMD10, DDIT4, LYAR, IRF1, CRY1, HMBOX1, EGR1, NR4A2, CRYAB, BASP1, DNAJB1, SUV39H2, CBX1, NKRF, ZFP36, ID1, CEBPD, IL33, GATA1, HDAC2, PHAX, HEXIM1, BMI1, STRAP, ZNF24, UXT, ID3, SLC4A1, TRAF2, ZBTB1, KAT2A, PSMC5, TERF2IP, PPP2CA, MED1, CEBPB, SDR16C5, ZBTB33, PAIP1, ZC3H8, FRK, MDFIC, ZBTB2, NAB2, DR1, SMAD4, CBX5, TASOR, GTPBP4, RBM42, POLE3, TXNIP, ZNF639, NACC1, PFDN5, ZHX1, BTAF1, NAF1,* |

| | | | | | | |
|---|---|---|---|---|---|---|
| | | | | | | *BCLAF1, TIRAP, CALR, NAT10, TIGAR, VDR, SRSF7, SUPT4H1, MBIP, DLL4, MORC2, MTA2, PHF14, BIRC5, PDCD4, WWP1, TP53, MAGEH1, ELAVL1, ARID5A, SHC1, SCML2, TERF1, EAPP, RLIM, OTUB1, MTERF3, TNKS2, NKX3-1, QKI, UBQLN4, IGF2BP1, PARPBP* |
| GO:1901698 | Response to nitrogen compound | 193 | 1.341 | 1.39E-02 | 3.56E-02 | *NR4A3, GDF15, GJA1, CHRNA3, DUOX1, DUOX2, TNFAIP3, PLA2G1B, SESN2, NFKBIA, DNAJB9, GOT1, HTR2B, DDX21, KCNJ11, CRY1, EGR1, NR4A2, CFTR, TLR4, TMEM38B, LOC100511937, HSP90B1, IFNE, STAT3, GSTM3, PTPN11, RNF5, RHOQ, STT3B, TRAF2, RAP1B, IGFBP5, SLC25A33, YOD1, PDE12, UBXN4, SELENOS, PRNP, SOCS5, IGFBP2, NPPC, ROCK1, FYN, SOCS1, HCFC2, ARHGEF2, SLC2A8, PRKAR1A, GRB10, BLOC1S6, GNA14, PANX1, NHERF1, CALR, TRPC3, VCP, GNAQ, DERL3, ITGB1, RNFT2, KCNE1* |
| GO:0006163 | Purine nucleotide metabolic process | 117 | 1.335 | 1.95E-02 | 4.66E-02 | *SAMHD1, NME1, LOC100515984, ADK, GOT1, PDE4D, ATP5MC1, DDIT4, UCHL1, ATP1B1, MDH1, NAMPT, HINT1, TP53I3, PDE4B, AK3, PGK1, COX11, ATP5PD, NME2, PPAT, ME1, ATP5MF, RHOQ, SLC4A1, HK2, PPP2CA, MTHFD1, ATP5MC3, ATP5ME, NUDT11, TPI1, IMPDH2, ATP5F1A, ATP5PO, PGAM1, NPPC* |
| GO:0045944 | Positive regulation of transcription by RNA polymerase II | 218 | 1.334 | 8.58E-03 | 2.59E-02 | *IL1A, INHBA, SOX11, LIF, IL10, BMP2, PLA2G1B, NFKBIA, EGR2, JUNB, PROX1, IER5, DDX3X, IRF1, TLR4, ECD, STAT3, NCOA7, CD40, GATA1, BMPR2, HDAC2, SERTAD1, LEO1, NME2, RHOQ, TNIP2, TAF9, IER2, SUMO2, SMAD3, KAT2A, MED1, CNOT7, MED4, MED21, RPRD1B, DDRGK1, PARK7* |
| GO:0048870 | Cell motility | 387 | 1.332 | 1.30E-03 | 5.79E-03 | *AMCF-II, MADCAM1, RASGEF1A, NR4A3, SELE, DUOXA2, CXCL8, CRB2, GDF15, CCL2, GJA1, THY1, ANGPT2, DCLK1, S100A8, CXCL10, IGFBP6, TNFAIP3, PPBP, ADRA2A, CXCL13, BMP2, PLA2G1B, S100A12, NARS1, CCNYL1, THBS1, PHLDA2, PROX1, IFNG, TWIST2, HTR2B, DDIT3, ACKR4, CCL8, NDNF, FFAR2, CXCL11, DUSP10, ENPEP, NR4A2, TLR4, ACKR2, LOC100511937, LAMTOR2, TNFSF14, TNFAIP1, CD40, BMPR2, PTP4A1, SNAI2, RHOB, RAB1A, PTPN11, IL27RA, PDCD10, LRP12, RAB13, SLC12A2, SPDL1, CCN3, GNA13, LIMA1* |

| | | | | | | |
|---|---|---|---|---|---|---|
| GO:0043069 | Negative regulation of programmed cell death | 213 | 1.331 | 1.20E-02 | 3.21E-02 | *WT1, CCL2, ATF5, IGF1, TNFAIP3, IL10, ARG2, HIGD1A, FAS, THBS1, PLK3, HTR2B, TCIM, PSMD10, DDX3X, GHITM, NDNF, NR4A2, CRYAB, MAD2L1, GATA1, SOD1, SNAI2, IL27RA, PIM1, BMI1, PDCD10, TNIP2, PSME3, HSPB1, TAF9, FTH1, DAD1, WNT16, LIG4, MED1, CEBPB, ST3GAL1, MCL1, FZD3, ZPR1, PARK7, SELENOS, HIPK3, PPIA, KRAS, NPM1, API5, PRDX3, NPPC, ROCK1, SIAH2, NRP1, PRKAA2, MIF, ANGPTL4, PPARGC1A, CCND2, MAPK8, XRCC2, CASP8, ITGA5, TIGAR, SLC25A5, ZFAND6, GPX4, CFLAR, GLO1, UBE2B, BIRC5, RHOA, MYC* |
| GO:0044770 | Cell cycle phase transition | 129 | 1.329 | 1.94E-02 | 4.66E-02 | *INHBA, CCL2, ATF5, PROX1, PLK3, TCIM, DDX3X, TIPIN, CRY1, CKS2, MAD2L1, RPS27L, CALM2, PTPN11, TIPRL, USP44, FBXO4, CCNG1, PSME3, SPDL1, ATM, MNAT1, PPP2CA, CDC25C, WEE1, NPM1, FEM1B, RCC1, CDK2AP2, CDK4, CCND2, ENSA, RAD1, PSME2, RAD21, WAC, BIRC5, MYC, CALM1, APP, NEK6* |
| GO:0051253 | Negative regulation of RNA metabolic process | 252 | 1.322 | 4.48E-03 | 1.60E-02 | *WT1, SOX11, ATF5, SAA3, BMP2, FABP4, IFNG, PLK3, DDIT3, KLF10, PSMD10, LYAR, IRF1, CRY1, HMBOX1, EGR1, NR4A2, CRYAB, BASP1, DNAJB1, SUV39H2, CBX1, NKRF, ZFP36, ID1, CEBPD, IL33, GATA1, HDAC2, PHAX, HEXIM1, BMI1, STRAP, ZNF24, UXT, ID3, TRAF2, ZBTB1, KAT2A, PSMC5, MED1, CEBPB, SDR16C5, ZBTB33, PAIP1, ZC3H8, FRK, MDFIC, ZBTB2, NAB2, DR1, SMAD4, CBX5, TASOR, RBM42, POLE3, TXNIP, ZNF639, NACC1, PFDN5, ZHX1, BTAF1, NAF1, BCLAF1, TIRAP* |
| GO:1902533 | Positive regulation of intracellular signal transduction | 271 | 1.320 | 4.88E-03 | 1.69E-02 | *RASGEF1A, INHBA, GDF15, SOX11, TIFA, IGF1, S100A8, LIF, IGFBP6, SAA3, ADRA2A, BMP2, PLA2G1B, S100A12, SESN2, LPAR3, THBS1, PROX1, HTR2B, DDX21, TCIM, DDIT3, DDX3X, TLR4, TICAM2, LAMTOR2, TNFSF14, SIAH1, MAD2L1, RASSF2, KCNK6, REL, CD40, SOD1, BMPR2, GPR4, ADAM17, HEXIM1, PTPN11, PIM1, GADD45B, EGF, PDCD10, FZD10, TNIP2, TRAF2, RAP1B, WNT16, TERF2IP, PPP2CA, MED1, IGFBP5, SLC15A3, MCL1, PARK7, SHOC2, BBC3, AVPI1, PRNP, PPIA, KRAS* |
| GO:0009792 | Embryo development | 168 | 1.317 | 1.51E-02 | 3.76E-02 | *PSPH, CRB2, GJA1, SOX11, FOSL1, LIF, EIF2S2, IL10, BMP2, MMP16, RSPO3, JUNB, PHLDA2, RCN1, ACTL6A, NLE1, GATA1, BMPR2, SF3B6, WDR74, TBX18, BMI1, ACSL4, GNA13, PRDM1, DAD1, KAT2A, LIG4, MED1, CEBPB, MED21, PLPP4, FZD3, ZPR1, LUZP1* |

| | | | | | | |
|---|---|---|---|---|---|---|
| | ending in birth or egg hatching | | | | | |
| GO:0040011 | Locomotion | 285 | 1.315 | 6.27E-03 | 2.05E-02 | *AMCF-II, MADCAM1, NR4A3, SELE, DUOXA2, CXCL8, GDF15, CCL2, GJA1, THY1, ANGPT2, S100A8, CXCL10, PPBP, ADRA2A, CXCL13, BMP2, PLA2G1B, THBS1, PHLDA2, PROX1, IFNG, TWIST2, ACKR4, CCL8, RCAN1, FFAR2, CXCL11, DUSP10, TLR4, ACKR2, TNFSF14, BVES, CD40, BMPR2, PTP4A1, SNAI2, RHOB* |
| GO:1902679 | Negative regulation of RNA biosynthetic process | 230 | 1.311 | 1.11E-02 | 3.11E-02 | *WT1, SOX11, ATF5, SAA3, BMP2, FABP4, IFNG, PLK3, DDIT3, KLF10, PSMD10, LYAR, IRF1, CRY1, HMBOX1, EGR1, NR4A2, CRYAB, BASP1, DNAJB1, SUV39H2, CBX1, NKRF, ZFP36, ID1, CEBPD, IL33, GATA1, HDAC2, HEXIM1, BMI1, STRAP, ZNF24, UXT, ID3, ZBTB1, KAT2A, PSMC5, MED1, CEBPB, SDR16C5, ZBTB33, ZC3H8, FRK, MDFIC, ZBTB2, NAB2, DR1, SMAD4, CBX5, TASOR, POLE3, TXNIP, ZNF639, NACC1, PFDN5, ZHX1, BTAF1, BCLAF1* |
| GO:0048878 | Chemical homeostasis | 242 | 1.311 | 6.67E-03 | 2.10E-02 | *IL6, IL1A, MT-2B, STC2, SLC8A3, CXCL10, ADRA2A, STEAP2, TFRC, SESN2, GOT1, AQP3, KCNE3, LIPG, FABP4, HTR2B, PRND, DDIT3, DDX3X, GHITM, CCL8, ATP1B1, CRY1, FFAR2, SOAT1, CXCL11, CFTR, ABHD5, TMEM38B, ORMDL1, STAT3, SLC31A1, CD40, SOD1, CALM2, TSKU, FXYD2, PTPN11, COX11, SLC39A8, SLC30A5, SLC12A2, SLC41A1, BPIFA1, LIMA1, SLC4A1, MINPP1, FTH1, HK2, IGFBP5, INSIG1, ABCB7, PARK7, NPC1, NUS1, RMI1, EDN1, TRA2B, SMAD4, RHCE, NUBP1, SAR1B, NPPC, PDZD8, ATP1A1, PRKAA2, ANGPTL4, FITM2, SLC4A4* |
| GO:0051049 | Regulation of transport | 313 | 1.309 | 3.33E-03 | 1.27E-02 | *IL6, IL1A, IL1B, NR4A3, APLN, INHBA, CCL2, NKAIN1, GJA1, SOX11, PTGER3, IGF1, S100A8, LIF, CXCL10, ADRA2A, BMP2, PLA2G1B, NFKBIA, KCNE3, THBS1, LIPG, IFNG, PLK3, LYAR, ATP1B1, KCNJ11, EIPR1, CRY1, FFAR2, CXCL11, KCNJ14, GPM6B, TLR4, TMEM38B, C7H6orf125, TGFB3, NDFIP2, KCNJ2* |
| GO:0045892 | Negative regulation of DNA-templated transcription | 228 | 1.303 | 1.26E-02 | 3.31E-02 | *WT1, SOX11, ATF5, SAA3, BMP2, FABP4, IFNG, PLK3, DDIT3, KLF10, PSMD10, LYAR, IRF1, CRY1, HMBOX1, EGR1, NR4A2, CRYAB, BASP1, DNAJB1, SUV39H2, CBX1, NKRF, ZFP36, ID1, CEBPD, IL33, GATA1, HDAC2, HEXIM1, BMI1, STRAP, ZNF24, UXT, ID3,* |

| | | | | | | |
|---|---|---|---|---|---|---|
| | | | | | | *ZBTB1, KAT2A, PSMC5, MED1, CEBPB, SDR16C5, ZBTB33, ZC3H8, FRK, MDFIC, ZBTB2, NAB2, DR1, SMAD4, CBX5, POLE3, TXNIP, ZNF639, NACC1, PFDN5, ZHX1, BTAF1, BCLAF1* |
| GO:0043009 | Chordate embryonic development | 167 | 1.297 | 1.84E-02 | 4.49E-02 | *PSPH, CRB2, GJA1, SOX11, FOSL1, LIF, EIF2S2, IL10, BMP2, MMP16, RSPO3, JUNB, PHLDA2, RCN1, ACTL6A, NLE1, GATA1, BMPR2, SF3B6, WDR74, TBX18, BMI1, GNA13, PRDM1, DAD1, KAT2A, LIG4, MED1, CEBPB, MED21, PLPP4, FZD3, ZPR1, LUZP1* |
| GO:0072359 | Circulatory system development | 245 | 1.297 | 1.12E-02 | 3.11E-02 | *IL1A, WT1, CXCL8, APLN, CRB2, SERPINE1, GJA1, SOX11, FOSL1, ANGPT2, LIF, CXCL10, IL10, BMP2, CACYBP, RSPO3, THBS1, JUNB, PROX1, HTR2B, IFT20, DDIT3, NDNF, LOC100511937, ESM1, BVES, SLC31A1, CD40, VASH2, BMPR2, RHOB, GPR4, HEXIM1, PTPN11, TBX18, PIM1, PDCD10, CCN3, GNA13, PRDM1, SERPINF1, MNAT1, WNT16, MED1, PDCL3, LUZP1, FKBPL, CALCRL, TMEM100, NDUFV2, SMAD4, IL18, MIB1, NPPC, ROCK1, GJC1, FYN, CDH11, NRP1, NAXE, CREB3L1, ANGPTL4, PRKAR1A* |
| GO:0036211 | Protein modification process | 430 | 1.296 | 2.93E-03 | 1.15E-02 | *LIF, TNFAIP3, RNF122, B3GNT5, PLA2G1B, C1GALT1, SOCS3, ST6GALNAC1, EGR2, B4GALT5, IFNG, RXYLT1, TPST1, RNF19B, PSMD10, DDX3X, GNL3, UCHL1, EIPR1, CRY1, KLHL25, EGR1, CISH, OTUB2, HINT1, ST6GAL2, NDFIP2, TNFAIP1, SIAH1, NCOA7, GATA1, EPHA4, MOGS, UBE2D3, UBE2D1, UCHL3, PTPN11, SLC39A8, B3GALNT1, PCMT1, FUT2, USP44, RNF11, FBXO4, DCUN1D5, RNF5, HSPB1, PCNP, STT3B, PRKAR2A, TRAF2, RNF180, SUMO2, FBXO5, DAD1, ATM, OSTC, SOCS7, KAT2A, IL15, PPP2CA, UCHL5, ST8SIA2, COPS6, DPM1, ST3GAL1, CUL5, DDRGK1, PARK7, SOCS4, NPC1, NUS1, RNF115, PRNP, RNF138, PPIA, GALNT1, SOCS5, HECTD2, DR1, MGAT2, ST3GAL3, B3GALT6, FEM1B, MIB1, SUMO1, DCAF13, SIAH2, RPN1, USP53, GTPBP4, PHF23, FYN, SOCS1, UBA6, NRP1, PRKAA2, PPP1CB, USP5, USP39, ARHGEF2, OTUD6B, RPN2* |
| GO:0019752 | Carboxylic acid metabolic process | 248 | 1.294 | 1.14E-02 | 3.11E-02 | *PSPH, PHGDH, ASNS, ARG1, GPT2, GDF15, PTGIS, VNN3, TYRP1, VNN1, ALDH1L2, ARG2, PLA2G1B, SESN2, GOT1, LIPG, FADS1, PROX1, FADS2, CTPS1, PTGS2, DDIT4, UCHL1, MID1IP1, MDH1, UGDH, ABHD5, SDS, RBP1, PTGES3, ASNSD1, PGK1, GLUD1, SLC39A8, ACSL4, ME1, GSTO1, SLC4A1, HK2, AZIN1, LTA4H, PPP2CA, ALDH18A1, APIP, ST3GAL1,* |

| | | | | | | |
|---|---|---|---|---|---|---|
| | | | | | | *INSIGI1, MTHFD1, PDHA1, PARK7, TPI1, NPC1, CTPS2, PTGR2, PGAM1, PCCB, LDHA, ACACA, MTAP, PDHX* |
| GO:0032879 | Regulation of localization | 413 | 1.290 | 3.66E-03 | 1.36E-02 | *IL6, IL1A, IL1B, NR4A3, APLN, INHBA, CCL2, NKAIN1, GJA1, SOX11, PTGER3, IGF1, S100A8, LIF, SLC5A3, CXCL10, TP53INP2, ADRA2A, BMP2, PLA2G1B, SESN2, NFKBIA, KCNE3, THBS1, LIPG, IFNG, PLK3, HTR2B, DDIT3, LYAR, GNL3, GHITM, ATP1B1, KCNJ11, EIPR1, CRY1, FFAR2, CXCL11, KCNJ14, GPM6B, ABHD5, TLR4, TMEM38B, C7H6orf125, TGFB3, NDFIP2, ZFP36, KCNJ2, CCT4, PARD6G, CCT2, PDZK1, CALM2, FXYD2, PTPN11, HILPDA, YWHAG, FBXO4, PDCD10, RHOQ, CCN3, BPIFA1, RAP1B, AZIN1, IL15, CEBPB, INSIG1, RAB14, PLIN2, PLPP4, ZPR1, DDRGK1, PARK7, BBC3, AKAP5, YIPF5, NUS1* |
| GO:0009607 | Response to biotic stimulus | 378 | 1.277 | 8.23E-03 | 2.53E-02 | *AMCF-II, IL1A, IL1B, GSDMA, CXCL8, CCL2, FOSL1, TIFA, MMP8, SAMHD1, S100A8, NDUFAF4, HP, CXCL10, TNFAIP3, IL10, PPBP, ARG2, CXCL13, BMP2, S100A12, CEBPG, FCN2, NFKBIA, HSP90AA1, FABP4, IFNG, DDX21, DDIT3, RNF19B, LBP, DDX3X, DDIT4, LYAR, CCL8, IRF1, NOD2, ATP1B1, FFAR2, DUSP10, PTGIR, CFB, TLR4, BPIFB1, TICAM2, IFNE, ZFP36, KCNK6, CD40, IFNAR1, POLR3D, PPP1R15B, MLKL, TMEM126A, RAB1A, HEXIM1, NFKBIB, IFNGR1, PTPN11, IL27RA, PIM1, USP44, YWHAG, RNF5, TNIP2, BPIFA1, TRAF2, S100A6, IRAK4, CDC37, PPP2CA, MED1, CEBPB, CNOT7, SLC15A3, RAB14, PDE12, FRK, SELENOS, CCL4, RNF115, LEAP2, TRIM10, IRAK1, IRF8, MAP3K7, EXOSC4, BATF2, CD46* |
| GO:0043207 | Response to external biotic stimulus | 365 | 1.273 | 6.46E-03 | 2.05E-02 | *AMCF-II, IL1A, IL1B, GSDMA, CXCL8, CCL2, FOSL1, TIFA, MMP8, SAMHD1, S100A8, NDUFAF4, HP, CXCL10, TNFAIP3, IL10, PPBP, ARG2, CXCL13, BMP2, S100A12, CEBPG, FCN2, NFKBIA, HSP90AA1, FABP4, IFNG, DDX21, RNF19B, LBP, DDX3X, DDIT4, LYAR, CCL8, IRF1, NOD2, ATP1B1, FFAR2, DUSP10, PTGIR, CFB, TLR4, BPIFB1, TICAM2, IFNE, ZFP36, KCNK6, CD40, IFNAR1, POLR3D, MLKL, TMEM126A, RAB1A, HEXIM1, NFKBIB, IFNGR1, PTPN11, IL27RA, PIM1, USP44, RNF5, TNIP2, BPIFA1, TRAF2, S100A6, IRAK4, CDC37, PPP2CA, MED1, CEBPB, CNOT7, SLC15A3, RAB14, PDE12, FRK, SELENOS, CCL4,* |

| | | | | | | |
|---|---|---|---|---|---|---|
| | | | | | | *RNF115, LEAP2, TRIM10, IRAK1, IRF8, MAP3K7, EXOSC4, BATF2, CD46, LSM5, IL18, PRDX3* |
| GO:0051707 | Response to other organism | 365 | 1.273 | 6.46E-03 | 2.05E-02 | *AMCF-II, IL1A, IL1B, GSDMA, CXCL8, CCL2, FOSL1, TIFA, MMP8, SAMHD1, S100A8, NDUFAF4, HP, CXCL10, TNFAIP3, IL10, PPBP, ARG2, CXCL13, BMP2, S100A12, CEBPG, FCN2, NFKBIA, HSP90AA1, FABP4, IFNG, DDX21, RNF19B, LBP, DDX3X, DDIT4, LYAR, CCL8, IRF1, NOD2, ATP1B1, FFAR2, DUSP10, PTGIR, CFB, TLR4, BPIFB1, TICAM2, IFNE, ZFP36, KCNK6, CD40, IFNAR1, POLR3D, MLKL, TMEM126A, RAB1A, HEXIM1, NFKBIB, IFNGR1, PTPN11, IL27RA, PIM1, USP44, RNF5, TNIP2, BPIFA1, TRAF2, S100A6, IRAK4, CDC37, PPP2CA, MED1, CEBPB, CNOT7, SLC15A3, RAB14, PDE12, FRK, SELENOS, CCL4, RNF115, LEAP2, TRIM10, IRAK1, IRF8, MAP3K7, EXOSC4, BATF2, CD46, LSM5, IL18, PRDX3* |
| GO:0045321 | Leukocyte activation | 226 | 1.268 | 1.57E-02 | 3.89E-02 | *IL6, AMCF-II, IL1A, IL1B, CXCL8, GJA1, SOX11, MMP8, TNFAIP3, IL10, CD274, ARG2, CEBPG, TFRC, SLC7A1, NFIL3, THBS1, IFNG, CTPS1, IRF1, DUSP10, EGR1, TNFSF13B, TLR4, TNFSF14, IFNE, STAT3, IL33, CD40, EXOSC3, IL27RA, BMI1, USP44, YWHAG, TRAF2, PRDM1, ATM, ZBTB1, KAT2A, IL15, LIG4, CEBPB, ST3GAL1* |
| GO:1901135 | Carbohydrate derivative metabolic process | 305 | 1.266 | 8.55E-03 | 2.59E-02 | *GFPT2, SAMHD1, NME1, GK, LOC100515984, ITIH2, ADK, B3GNT5, BMP2, C1GALT1, ST6GALNAC1, B4GALT5, IFNG, RXYLT1, RFK, CTPS1, PDE4D, ATP5MC1, DDIT4, GFPT1, UCHL1, ATP1B1, GNPNAT1, SOAT1, UGDH, PGM2, HINT1, ST6GAL2, GATA1, BMPR2, PDE4B, MOGS, AK3, PGK1, MPI, COX11, SLC39A8, B3GALNT1, FUT2, ATP5PD, NME2, PPAT, ATP5MF, RHOQ, STT3B, SLC4A1, DAD1, DCTPP1, OSTC, HK2, PPP2CA, ST8SIA2, DPM1, PGM3, ST3GAL1, PIGP, ATP5MC3, ATP5ME, TYW5, NUDT11, TPI1, NPC1, NUS1, PRPS1, CTPS2, GALNT1, IMPDH2, PIGK, ATP5F1A, ATP5PO, MGAT2, PGAM1, ST3GAL3, B3GALT6, NPPC, MTAP, RRM2B, RPN1* |
| GO:0010628 | Positive regulation of gene expression | 304 | 1.262 | 9.72E-03 | 2.79E-02 | *IL1A, IL1B, CXCL8, INHBA, GJA1, SOX11, ATF3, MMP8, LIF, IL10, ADRA2A, BMP2, PLA2G1B, CEBPG, TFRC, NFIL3, THBS1, HSP90AA1, IFNG, SLC38A2, HTR2B, DDX21, DDIT3, DDX3X, IRF1, FFAR2, EGR1, FAM98A, TLR4, LOC100511937, TICAM2, C7H6orf125,* |

| | | | | | | |
|---|---|---|---|---|---|---|
| | | | | | | *ZFP36, FAM98B, CHUK, IL33, CD40, SOD1, BMPR2, POLR3D, NIBAN1, EXOSC3, RAB1A, PTPN11, HILPDA, IL27RA, PDCD10, SSB, TRAF2, PRDM1, KAT2A, WNT16, IL15, MED1, CEBPB, PDCL3, PAIP1, ZPR1, DDRGK1, PARK7, DPY30, IL16, FASTKD2, KRAS, NPM1, IRF8, MAP3K7* |
| GO:0006796 | Phosphate-containing compound metabolic process | 389 | 1.257 | 5.99E-03 | 1.97E-02 | *GFPT2, SAMHD1, NME1, LIF, GK, LOC100515984, ADK, PLA2G1B, GOT1, THBS1, IFNG, HTR2B, PPA1, RFK, CTPS1, PDE4D, ATP5MC1, DDIT4, GFPT1, UCHL1, ATP1B1, MDH1, GNPNAT1, NAMPT, UGDH, ABHD5, TMEM38B, HINT1, ORMDL1, TP53I3, STAT3, GATA1, PDE4B, EPHA4, AGK, AK3, PGK1, PTPN11, MPI, INPP1, COX11, LPGAT1, ATP5PD, NME2, ACSL4, PPAT, ME1, ATP5MF, RHOQ, PLPPR2, CMPK1, PTPMT1, HSPB1, PRKAR2A, SLC4A1, TAMM41, DCTPP1, ATM, LPCAT3, HK2, IL15, DCTD, PPP2CA, DPM1, PGM3, MTHFD1, PIGP, PLPP4, ATP5MC3, ATP5ME, PDHA1, PARK7, NUDT11, TPI1, NUS1, PRPS1, PRNP, BPNT1, PANK3, CTPS2, PPIA, IMPDH2, PIGK, ATP5F1A, ATP5PO, PGAM1, TPK1, ACACA, NPPC, CRLS1, ROCK1, PDHX, RRM2B, FYN, NRP1, PRKAA2, PPP1CB, NAXE, UGP2, ARHGEF2, MIF, FITM2* |
| GO:0001775 | Cell activation | 261 | 1.257 | 1.12E-02 | 3.11E-02 | *IL6, AMCF-II, IL1A, IL1B, FGB, CXCL8, GJA1, SOX11, MMP8, CXCL10, TNFAIP3, IL10, CD274, ARG2, ADRA2A, CEBPG, TFRC, SLC7A1, NFIL3, THBS1, IFNG, CTPS1, TCIM, IRF1, DUSP10, EGR1, TNFSF13B, TLR4, TNFSF14, IFNE, STAT3, IL33, CD40, GATA1, EXOSC3, IL27RA, BMI1, USP44, YWHAG, TUBB1, GNA13, TRAF2, PRDM1, ATM, ZBTB1, KAT2A, IL15, LIG4, CEBPB, ST3GAL1* |
| GO:0043436 | Oxoacid metabolic process | 250 | 1.256 | 1.39E-02 | 3.56E-02 | *PSPH, PHGDH, ASNS, ARG1, GPT2, GDF15, PTGIS, VNN3, TYRP1, VNN1, ALDH1L2, ARG2, PLA2G1B, SESN2, GOT1, LIPG, FADS1, PROX1, FADS2, CTPS1, PTGS2, DDIT4, UCHL1, MID1IP1, MDH1, UGDH, ABHD5, SDS, RBP1, PTGES3, ASNSD1, PGK1, GLUD1, SLC39A8, ACSL4, ME1, GSTO1, SLC4A1, HK2, AZIN1, LTA4H, PPP2CA, ALDH18A1, APIP, ST3GAL1, INSIG1, MTHFD1, PDHA1, PARK7, TPI1, NPC1, CTPS2, PTGR2, PGAM1, PCCB, LDHA, ACACA, MTAP, PDHX* |

| | | | | | | |
|---|---|---|---|---|---|---|
| GO:0006793 | Phosphorus metabolic process | 390 | 1.251 | 6.91E-03 | 2.16E-02 | *GFPT2, SAMHD1, NME1, LIF, GK, LOC100515984, ADK, PLA2G1B, GOT1, THBS1, IFNG, HTR2B, PPA1, RFK, CTPS1, PDE4D, ATP5MC1, DDIT4, GFPT1, UCHL1, ATP1B1, MDH1, GNPNAT1, NAMPT, UGDH, ABHD5, TMEM38B, HINT1, ORMDL1, TP53I3, STAT3, GATA1, PDE4B, EPHA4, AGK, AK3, PGK1, PTPN11, MPI, INPP1, COX11, LPGAT1, ATP5PD, NME2, ACSL4, PPAT, ME1, ATP5MF, RHOQ, PLPPR2, CMPK1, PTPMT1, HSPB1, PRKAR2A, SLC4A1, TAMM41, DCTPP1, ATM, LPCAT3, HK2, IL15, DCTD, PPP2CA, DPM1, PGM3, MTHFD1, PIGP, PLPP4, ATP5MC3, ATP5ME, PDHA1, PARK7, NUDT11, TPI1, NUS1, PRPS1, PRNP, BPNT1, PANK3, CTPS2, PPIA, IMPDH2, PIGK, ATP5F1A, ATP5PO, PGAM1, TPK1, ACACA, NPPC, CRLS1, ROCK1, PDHX, RRM2B, FYN, NRP1, PRKAA2, PPP1CB, NAXE, UGP2, ARHGEF2, MIF, FITM2* |
| GO:1902531 | Regulation of intracellular signal transduction | 476 | 1.251 | 5.15E-03 | 1.75E-02 | *RASGEF1A, INHBA, GDF15, SOX11, ATF3, TIFA, IGF1, S100A8, LIF, IGFBP6, TNFAIP3, SAA3, IL10, ADRA2A, BMP2, PLA2G1B, S100A12, FAS, SESN2, NFKBIA, LPAR3, THBS1, DUSP5, PROX1, HTR2B, DDX21, TCIM, DDIT3, PSMD10, DDX3X, RRS1, DDIT4, RCAN1, SAMD5, CRY1, DUSP10, TLR4, TICAM2, LAMTOR2, TNFSF14, TNFAIP1, SIAH1, MAD2L1, RASSF2, KCNK6, REL, CD40, SOD1, BMPR2, SNAI2, GPR4, UBE2D1, PAQR3, ADAM17, CALM2, HEXIM1, NFKBIB, PTPN11, SLC39A8, PIM1, GADD45B, USP44, EGF, YWHAG, FBXO4, PDCD10, FZD10, NOP2, TNIP2, CHN2, CCN3, TRAF2, ATM, RAP1B, CRK, WNT16, TERF2IP, PPP2CA, MED1, IGFBP5, SLC15A3, INSIG1, MCL1, DDRGK1, PARK7, SHOC2, BBC3, AKAP5, NPC1, SELENOS, MDFIC, AVPI1, PRNP, STYX, PPIA, KRAS* |
| GO:0006508 | Proteolysis | 432 | 1.236 | 1.14E-02 | 3.11E-02 | *MMP12, MMP13, SERPINE1, CTSL, MMP8, MMP1, TP53INP2, TNFAIP3, RNF122, IL10, TIMP1, ADRA2A, MMP16, DNAJB9, THBS1, PLAU, IFNG, ASRGL1, DDIT3, RNF19B, PSMB3, PSMD10, DESI2, MMP27, UCHL1, CASP3, ENPEP, PCOLCE, MMP9, CFB, OTUB2, HSP90B1, KLF1, NDFIP2, TNFAIP1, ZFAND2A, SIAH1, BACE2, ADAMTS1, ADAMTS9, PSMA3, HDAC2, LOC100522887, PSMA5, UBE2D3, TMEM126A, UBE2D1, PAQR3, ADAM17, UCHL3, DHH, ADAMTS20, USP44, RNF11, FBXO4, RNF5, PSMB6, PSME3, IMMP1L, STT3B, TAF9, RNF180, SUMO2, RAD23A, KLHL2, LTA4H, PSMC5, UCHL5, YOD1,* |

| | | | | | | |
|---|---|---|---|---|---|---|
| | | | | | | *PDCL3, METAP2, CUL5, NEMF, SPCS2, PSMA4, PREPL, DDRGK1, UBXN4, PARK7, PSMA2, PSMD8, PSMA1, SOCS4, YIPF5, SELENOS, RNF115, LOC100739080, STYX, SOCS5, PIGK, CD46, OMA1, FEM1B, MIB1, DPP3, SIAH2, THOP1, RAD23B, PSMC2, PSMB1, METAP1, USP5, AKIRIN2, USP39, SEC11C, ARHGEF2, OTUD6B, PSMC1, TMEM168, PSMB4, NRDC, CPA1, ELOC, CHMP2B, ADAM9, ANAPC10, USP15, LTN1, PHEX, PSMD4, SPCS3, UBB, CASP8, CALR, UQCRC2, VCP, FOLH1B, IDE, DERL3, RNFT2, PSME2, RHBDL2* |
| GO:0007155 | Cell adhesion | 326 | 1.230 | 9.58E-03 | 2.77E-02 | *MADCAM1, IL1A, IL1B, RND1, NR4A3, CLSTN3, SELE, CXCL8, CRB2, THY1, POSTN, SELP, ANGPT2, S100A8, LIF, PRTG, VCAN, IL10, CD274, ARG2, BMP2, TFRC, ICAM4, SLC7A1, THBS1, PLAU, IFNG, TNC, NDNF, IRF1, PCDH11X, ATP1B1, DUSP10, TMEM47, TNFSF13B, TNFSF14, BVES, NECTIN1, GATA1, EPHA4, SNAI2, VCAM1, ICAM1, RHOB, GPR4, PTPN11, SLC39A8, FUT2, BMI1, YWHAG, TUBB1* |
| GO:0002682 | Regulation of immune system process | 390 | 1.227 | 1.19E-02 | 3.21E-02 | *MADCAM1, ORM1, IL1A, IL1B, SELE, INHBA, GDF15, CCL2, CSF3, SOX11, FOSL1, TIFA, MMP8, SAMHD1, LIF, CXCL10, TNFAIP3, IL10, CD274, ARG2, PLA2G1B, TFRC, FCN2, NFKBIA, SLC7A1, AQP3, THBS1, HSP90AA1, HSPA9, IFNG, DDX21, TCIM, KLF10, DDX3X, LYAR, IRF1, ULBP1, FFAR2, DUSP10, TNFSF13B, CFB, TLR4, BPIFB1, TICAM2, TGFB3, TNFSF14, ZFP36, RASSF2, STAT3, KCNK6, LOC100622460, IL33, CD40, GATA1, POLR3D, LEO1, EXOSC3, TMEM126A, HEXIM1, IL27RA, PIM1, BMI1, YWHAG, TNIP2, CCN3, TRAF2, LRRC17, PRDM1, SKIC8, ZBTB1, KAT2A, CDC37, IL15, PPP2CA, MED1, CEBPB, CNOT7, SLC15A3, ZBTB33* |
| GO:0006082 | Organic acid metabolic process | 252 | 1.225 | 1.81E-02 | 4.45E-02 | *PSPH, PHGDH, ASNS, ARG1, GPT2, GDF15, PTGIS, VNN3, TYRP1, VNN1, ALDH1L2, ARG2, PLA2G1B, SESN2, GOT1, LIPG, FADS1, PROX1, FADS2, CTPS1, PTGS2, DDIT4, UCHL1, MID1IP1, MDH1, UGDH, ABHD5, SDS, RBP1, PTGES3, ASNSD1, PGK1, GLUD1, SLC39A8, ACSL4, ME1, GSTO1, SLC4A1, HK2, AZIN1, LTA4H, PPP2CA, ALDH18A1, APIP, ST3GAL1, INSIG1, MTHFD1, PDHA1, PARK7, TPI1, NPC1* |
| GO:0044419 | Biological process involved in | 399 | 1.212 | 1.98E-02 | 4.69E-02 | *AMCF-II, IL1A, IL1B, GSDMA, CXCL8, CCL2, FOSL1, TIFA, MMP8, SAMHD1, S100A8, NDUFAF4, HP, CXCL10, TNFAIP3, IL10, PPBP, ARG2, CXCL13, BMP2, S100A12, CEBPG,* |

| | | | | | | |
|---|---|---|---|---|---|---|
| | interspecies interaction between organisms | | | | | *FCN2, NFKBIA, ST6GALNAC1, HSP90AA1, FABP4, IFNG, DDX21, RNF19B, LBP, DDX3X, DDIT4, LYAR, CCL8, IRF1, NOD2, ATP1B1, FFAR2, DUSP10, PTGIR, CFB, TLR4, BPIFB1, TICAM2, IFNE, ZFP36, KCNK6, NECTIN1, CD40, IFNAR1, POLR3D, MLKL, TMEM126A, RAB1A, HEXIM1, NFKBIB, IFNGR1, PTPN11, IL27RA, PIM1, USP44, RNF5, TNIP2, BPIFA1, TRAF2, S100A6, IRAK4, CDC37, PPP2CA, MED1, CEBPB, CNOT7, SLC15A3, RAB14, PDE12, PAIP1, FRK, SELENOS, CCL4, RNF115, LEAP2, TRIM10, RAB5A, IRAK1, IRF8, MAP3K7, EXOSC4, BATF2, CD46, LSM5, IL18, PRDX3, FYN, NRP1, HCFC2, THRSP, LOC100516039, AKIRIN2, ZNF639, CDK4, ARHGEF2, MIF* |
| GO:0003008 | System process | 300 | 1.185 | 1.87E-02 | 4.52E-02 | *IL6, IL1B, INHBA, CCL2, AGTR2, GJA1, PTGER3, AQP5, SLC8A3, GLRB, RDH12, ARG2, ADRA2A, SLC26A5, C5H12orf57, MYH2, AQP3, KCNE4, KCNE3, EGR2, HTR2B, F5, IFT20, DDIT3, UCHL1, RCAN1, ATP1B1, KCNJ11, CRY1, DRAM2, ENPEP, HTR7, TLR4, TMEM38B, NOB1, KCNJ2, GNAT1, KCNK6, GRK7, SOD1, BMPR2, SNAI2, GPR4, CALM2* |
| GO:0030031 | Cell projection assembly | 139 | -1.438 | 1.28E-02 | 3.31E-02 | *TEKT1, TEKT2, RAC2, F2RL1, PIERCE1, PDGFA, SPACA9, VSTM5, GMNC, CEP250, TMEM107, CCNO, LIMK2, SPEF2, IQUB, CFAP58, UBXN10, AIF1, CFAP97D1, BBS10, PLD1, TACSTD2, WRAP73, PARVG, CCDC113, CFAP91, INPP5K, FSCN1, BBS1, MEIG1, DNAI2, DYNC2I2, ODAD1, P2RX7, P2RY12* |
| GO:0120036 | Plasma membrane bounded cell projection organization | 296 | -1.491 | 8.50E-04 | 4.03E-03 | *FAM161A, ACP4, ARHGEF25, TEKT1, TEKT2, SEMA6A, GPR37, RAC2, F2RL1, EPHB3, WNT7A, FES, TUB, PIERCE1, SPACA9, SRCIN1, PRKN, VSTM5, RTN4RL1, ENPP2, PLPPR4, SLITRK2, SEMA3G, GMNC, CEP250, NEDD9, TMEM107, CCNO, EFNB3, FOLR3, BDNF, IFT46, CHODL, KIF19, LIMK2, SPEF2, IQUB, CFAP58, UBXN10, CAMK2A, CNTN5, SYT3, CNTN1, AIF1, GPRASP3, CFAP97D1, SEMA6D, BBS10, PLD1, TACSTD2, WRAP73, CCDC113, CFAP91, INPP5K, FSCN1, BBS1, MEIG1, DNAI2, NTF3, SFRP2, DYNC2I2, ZNF365, SEMA6C, GLDN, RASGRF1, RTN4R, ODAD1, P2RX7, RND2, ATCAY, P2RY12, CNR1, GBX1, MYOC, NTRK3, INPP5J, APOD* |
| GO:0030030 | Cell projection organization | 311 | -1.515 | 7.76E-04 | 3.73E-03 | *FAM161A, ACP4, ARHGEF25, TEKT1, TEKT2, SEMA6A, GPR37, RAC2, F2RL1, EPHB3, WNT7A, FES, TUB, PIERCE1, PDGFA, SPACA9, SRCIN1, PRKN, VSTM5, RTN4RL1, ENPP2,* |

*PLPPR4, SLITRK2, SEMA3G, GMNC, CEP250, NEDD9, TMEM107, CCNO, EFNB3, FOLR3, BDNF, IFT46, CHODL, KIF19, LIMK2, TCTN1, SPEF2, IQUB, CFAP58, UBXN10, CAMK2A, CNTN5, SYT3, CNTN1, AIF1, GPRASP3, CFAP97D1, SEMA6D, BBS10, PLD1, TACSTD2, WRAP73, PARVG, CCDC113, CFAP91, INPP5K, FSCN1, BBS1, MEIG1, DNAI2, NTF3, SFRP2, DYNC2I2, ZNF365, SEMA6C, GLDN, RASGRF1, RTN4R, MKS1, ODAD1, P2RX7, RND2, ATCAY, P2RY12, CNR1, GBX1, MYOC, NTRK3, INPP5J, APOD*

**Abbreviations:** GO:BP: Biological Processes in Gene Ontology; FDR: False discovery rate; NES: Normalized enrichment score.

93
